# Targeting a Pleckstrin Homology Domain with a Lysine-Reactive Covalent Binder

**DOI:** 10.64898/2025.12.17.694903

**Authors:** Rebekah M. West, Radu Costin Bizga Nicolescu, Paul Brear, James Wagstaff, Beata K. Blaszczyk, Tomas Deingruber, Matthew G. Sanders, F. Javier Pérez-Areales, David R. Spring, Marko Hyvönen

## Abstract

Bruton’s Tyrosine Kinase (BTK) is a validated target for haematological malignancies, with numerous FDA approved inhibitors on the market. Current therapies target the highly conserved ATP binding site and hence limit the therapeutic index given the site’s highly conserved nature across the kinome. We explore a novel approach for BTK inhibition, by targeting the PH domain-mediated membrane recruitment and activation of BTK. We have identified a fragment which covalently labels a lysine in the inositol phosphate (PIP3) binding site. Fragment growth and an extensive structure-binding relationship study uncovered 27 crystal structures and a best-in-class analog, **24**. Evaluation of pKa values of the targeted lysine in BTK and other PH domains suggests this as a more general approach to PH domain inhibition.

---

Inhibition of signaling proteins, such as protein kinases, is typically achieved through targeting of their active sites, such as ATP-binding sites in kinases. As these sites tend to be under high evolutionary pressure to preserve binding to natural co-factor ATP, development of specific inhibitors can be challenging. An alternative approach is to identify other functionally important sites in the protein, which would offer better selectivity. Such sites might include protein-protein interaction sites or binding sites for other co-factors. In the case of BCR:ABL inhibition, the myristoyl binding site in the kinase lobe has been successfully used to develop a treatment for chronic myeloid leukaemia (CML).^1^ We have recently shown how targeting of the TPX2 binding site on Aurora A kinase can result in highly specific inhibition through a novel mechanism.^2^

Many eukaryotic signaling proteins are composed of multiple independent domains that modulate the activity and localization of the proteins. Among these, Pleckstrin Homology (PH) domains are one of the most common, with an estimated 250 of them in the human proteome.^3^ PH domains are typically involved in membrane association, through binding to phosphoinositides on the inner leaflet of the plasma membrane. In the case of Akt, the N-terminal PH domain can interact with the kinase domain, rendering it inactive, but can release this activation upon lipid binding. The importance of this mechanism has been highlighted by identification of molecules that inhibit Akt by locking the PH and kinase domains in their inactive state.^4^

The Tec family of tyrosine kinases is another example of PH domain-containing signaling proteins where membrane association is critically important for their activity. As members of the larger Src superfamily, they share a core structure of SH2-SH3-kinase domains, but contain an additional PH domain and a Zn^2+^ binding Tec-homology domain (also called as BTK-motif) in their N-termini (Figure 1A).^5^ However, unlike other Src-like kinases, which are usually myristoylated at their N-termini and hence intrinsically membrane associated, the membrane association of Tec kinases is mediated by the PH domain through interactions with phosphatidylinositol (3,4,5)-trisphosphate (PIP3), the product of PI3-kinase activity on the plasma membrane.^6^

**Figure 1.**
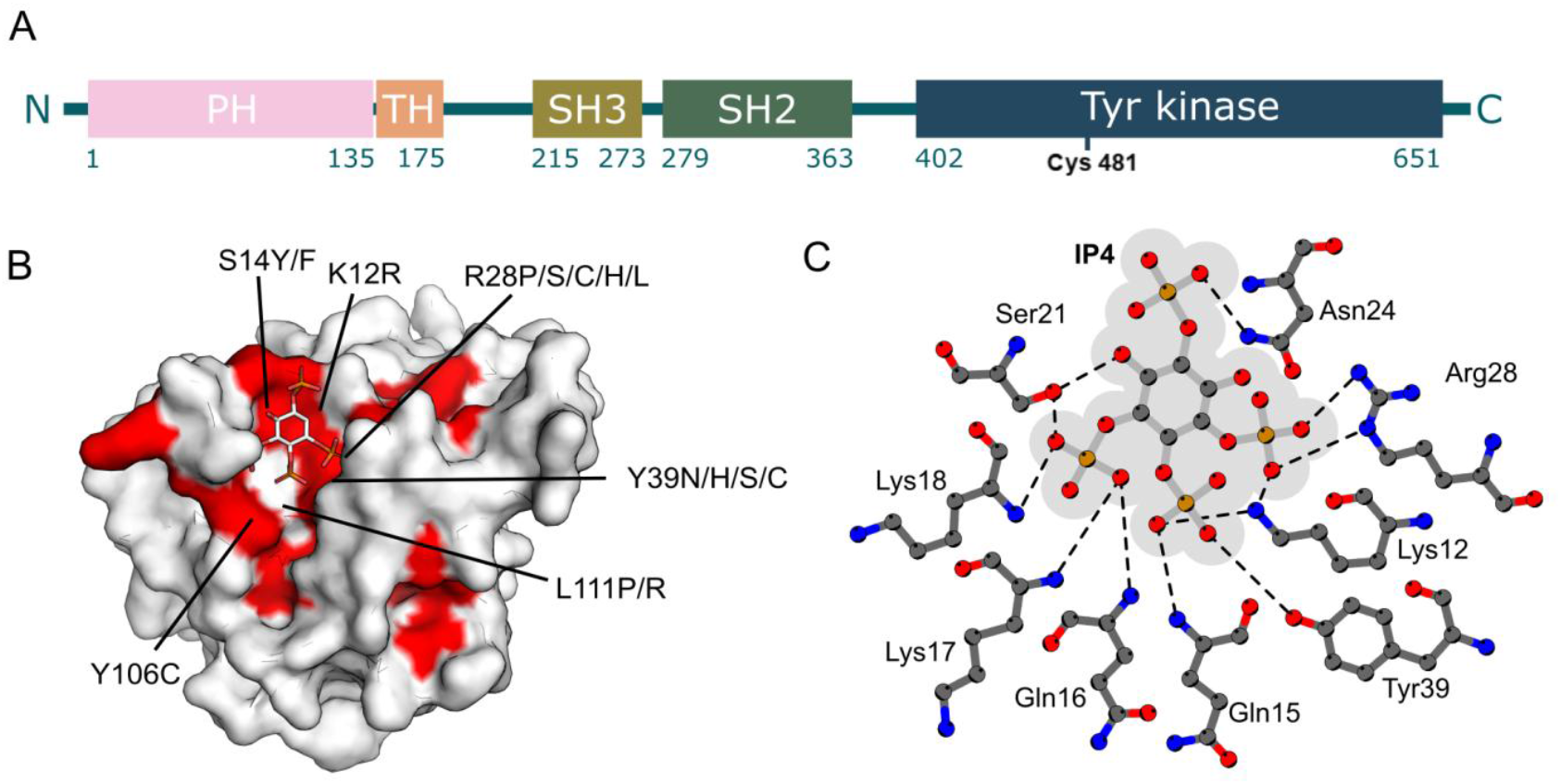
BTK as a target. **A.**From N- to C-terminus, the 659 amino acid BTK are divided into five domains - Pleckstrin homology (PH), Tec homology (TH), Src homology 3 (SH3), Src homology 2 (SH2) and tyrosine kinase. Lysine 12 and cysteine 481, targets of covalent inhibitors from this work and of ibrutinib, respectively, are highlighted. **B**. Structure of BTK PH domain in complex with IP4 (PDB: 1B55) with ligand show as sticks and domain with molecular surface. Residues around the IP4 binding site are found to cause XLA are coloured red (as listed in LOVD database https://databases.lovd.nl/shared/genes/BTK). **C**. Details of the IP4 binding site in BTK PH domain, showing the electrostcic and hydrogen bonding interactions between the protein and the domain. Figure modified from Ligplot output.

Bruton’s Tyrosine Kinase (BTK) is one of the best studied Tec family kinases.^7^ BTK plays a key role in B-cell maturation and its inactivation by mutations results in X-linked agammaglobulinemia (XLA), a condition in which (typically) male individuals lack B-cells and antibody-mediated humoral immune protection.^5,7^

BTK is also studied for its role in cancer. Primarily expressed in haematopoietic cells, aberrant BTK interferes with healthy BCR signaling leading to uncontrolled B-cell survival and proliferation in chronic lymphocytic leukaemia (CLL), mantle cell lymphoma (MCL) and other blood cancers.^8^ Interestingly, isoforms of BTK (p65BTK) are also thought to drive tumour progression in solid tumours (such as colon, breast and prostate), through promoting resistance to chemotherapy, contributing to tumour-stroma communication and manipulating macrophage activity and immune suppression.^8^

Activation of BTK is dependent on PH domain binding to PIP3. Membrane recruitment mediated dimerization is thought to release BTK from its inhibited form, resulting in trans-autophosphorylation and activation of BTK.^9^ Given that a large number of mutations are found in the inositol phosphate binding site and abolishing lipid binding can result in XLA, inhibition of membrane association by small molecules would be a valid approach to BTK inhibition (Figure 1B). For example, R28, one of the key residues involved in PIP3 binding, is mutated in a number of XLA cases^10^ and mutation of the same residue to cysteine (R28C) in mice results in X-linked immunodeficiency (XID).^11^ In addition to validating the PH domain as a target for BTK inhibition, interaction with these conserved ligand-binding residues would potentially make it difficult for BTK to acquire resistance as their mutation would also disrupt binding to natural ligands.

Targeting PH domains is not expected to be straightforward as the binding site for phosphorylated phosphoinositides is highly charged and their interactions with the PH domain are largely ionic (Figure 1C). Also, the binding occurs in a relatively shallow pocket on the surface of the domain, offering limited three-dimensionality. In the early 2000s, the purine analog, tricirbine, was repurposed upon the discovery that its mono-phosphate active form prevented the membrane recruitment of Akt. As might be expected for inhibitors binding a highly charged pocket, tricirbine had limited bioavailability and, disappointingly, was discontinued from clinical trials due to toxicity and poor efficacy.^12^ Peptides which are better able to mimic PIP3 interaction motifs have been developed against Akt PH domain E17K mutants, but they suffer from poor stability, permeability, and half-life in comparison to small molecules.^13^

At the time of writing, no direct PH domain inhibitors have advanced to late-stage clinical trials, despite the recent development of preclinical candidates against PDK1^14^ and BRAG2^15^. With the PH domain being found in large numbers of human proteins whilst being relatively poorly conserved, its inhibition has been recognised as an interesting alternative to more conventional targets sites in BTK.^16^

With this background in mind, we decided to use BTK as a model for PH domain-targeted inhibitor development. We used fragment-based ligand discovery, aiming to generate from the outset binders that have drug-like properties and are suitable for further development. A reactive lysine residue was discovered in the middle of the PIP3 binding site and used as the anchor point for the development of inhibitors that block PIP3 binding to the PH domain.

## RESULTS AND DISCUSSION

### Fragment screening against PH domain

The fragment screening campaign against the BTK PH domain consisted of a thermal shift assay, followed by validation with X-ray crystallography and biophysical techniques (ITC).

An in-house library of 720 fragments was screened against both the wild-type (WT) and mutant (R28C) BTK PH domain constructs by differential scanning fluorimetry (DSF). DSF assays have varied success in fragment-based screens; however, soluble PIP3 analog inositol-1,3,4,5-tetrakisphosphate (IP4) resulted in +20 °C stabilization of the domain, suggesting that productive binding to this site would be detected by thermal shift. As R28 is one of the key residues for IP4 binding and frequently mutated in XLA patients (Figure 1B), we used a PH domain with an R28C mutation as a control to identify hits that bound to the targeted IP4 binding site. A fragment that stabilized the WT PH domain but not the R28C mutant was likely to be binding at the IP4 site. The original screen identified 7 hits that stabilized the WT PH domain by more than 5 °C. These were further validated by DSF and X-ray crystallography. Testing the fragments at different concentrations by DSF identified two compounds (**i/1** and **v**) that showed significant stabilization of the WT domain while fully inactive against the R28C mutant. Compound **vii** also showed modest stabilization of the WT PH domain compared to the mutant (Figure 2A).

**Figure 2.**
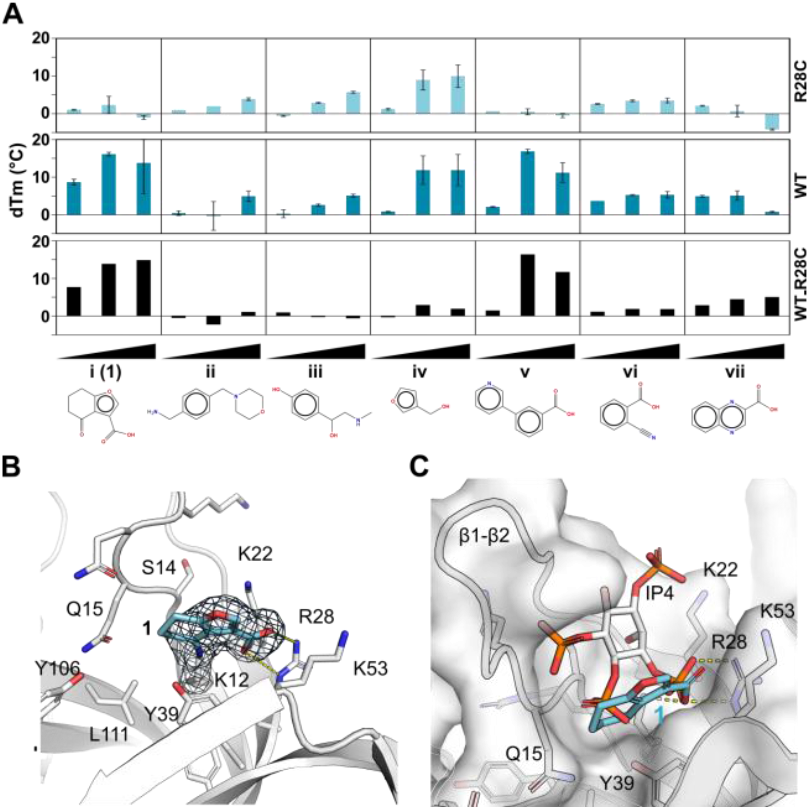
Fragment hit validation. **A.**DSF analysis of hit fragments against WT and R28C mutant of BTK PH domain at three different concentrations. **B**. Crystal structure of **1** covalently bound to K12 in the BTK PH domain (PDB:6TUH). **C**. Overlay of IP4 (PDB:1B55) and hit **1**.

While the WT PH domain had not been crystallized before in unliganded form, well diffracting, soakable crystals were obtained through serial seeding using crystals of R28C mutant (PDB: 1BTK).^5^ The four fragment hits were soaked to these crystals but data from **v, vi** and **vii** showed only partial density for the fragment, most likely of the carboxylic acid moiety, close to R28 side chain.

In contrast, soaking the crystals with fragment **i/1** showed good density for the entire fragment (Figure 2B). Unexpectedly, **1** forms a covalent bond to terminal amino group of K12 (PDB:6TUH). The carboxylic acid of **1** interacts with R28 as expected, confirming its failure to stabilise R28C mutant. The binding site is overlapping with IP4 binding site, engaging with key residues for IP4 interactions (Figure 2C). Compound **1** is sandwiched between a loop connecting β-strands 1 and 2 (β1-β2 loop) and residues Y39 and K53 from strands 3 and 4. The 1-2 loop undergoes significant reorganization upon binding to IP4 and is seen also in a more closed conformation with Q15 side chain hydrogen binding to Y106, compared to unliganded BTK PH domain. Our crystal system contained four domains in the asymmetric unit, which showed variable levels of modification by **1**, with one of the domains containing a second modified lysine (K19) next to K12. In subsequent elaborations only K12 was ever modified.

### Biophysical validation of fragments

To further investigate the contribution of covalent bond formation to inhibition of IP4 binding to the BTK PH domain, we characterized this hit using isothermal titration calorimetry (ITC). While direct binding of **1** to the BTK PH domain could not be detected in ITC (data not shown), we could measure binding of IP4 to the PH domain with K_d_ of ca. 36 nM (Figure 3A). When the same titration was performed in the presence of 1 mM of **1**, IP4 binding was completely inhibited, confirming interaction with the target site (Figure 3B). The same ITC competition study was performed with **1z** where the electrophilic cyclohexanone section of **1** was removed (Figure 3C). As expected, **1z** did not inhibit the binding of IP4 to BTK PH and confirmed the importance of the covalent interaction observed in the crystal structure.

**Figure 3.**
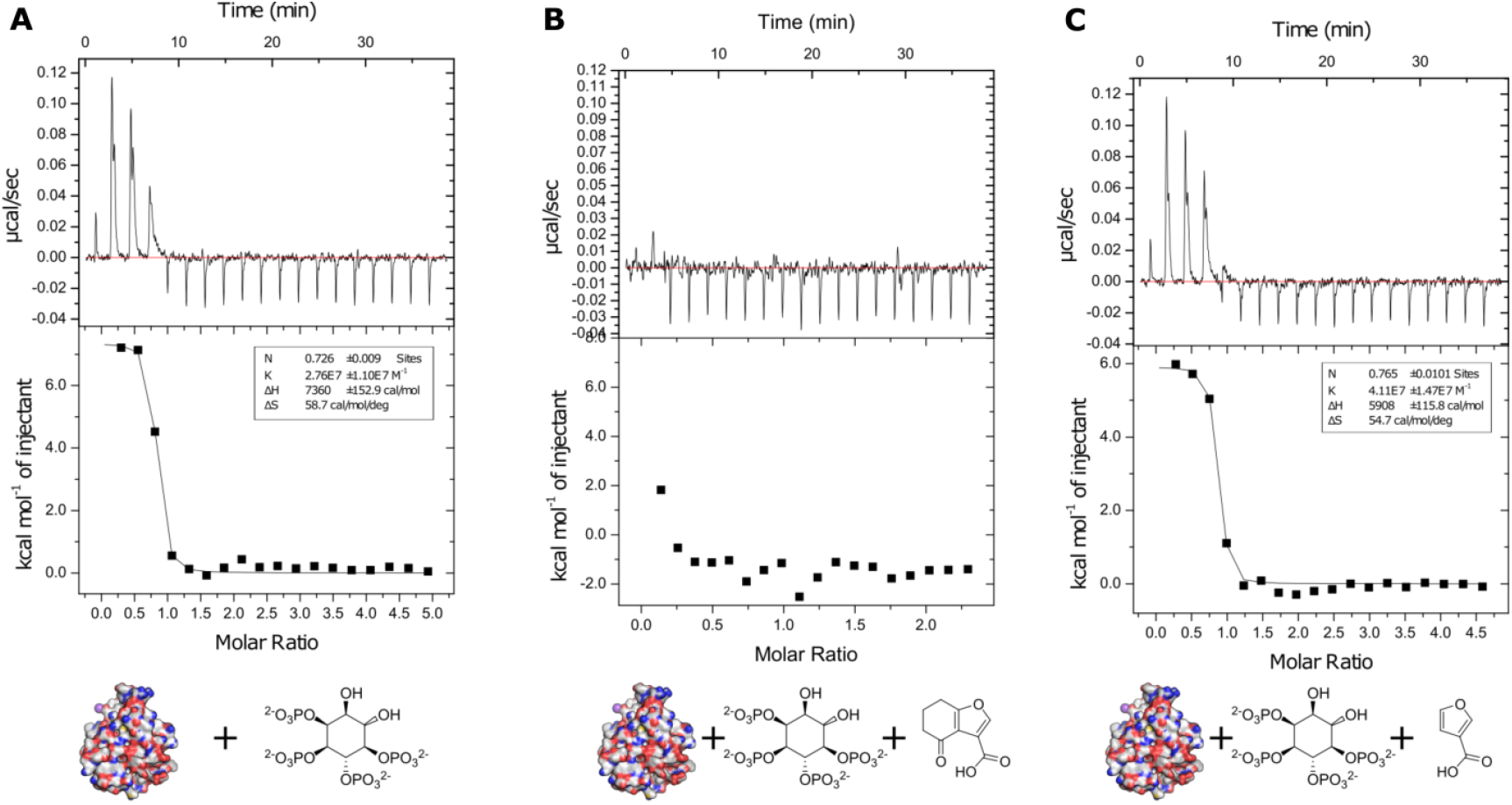
Isothermal titration calorimetry experiments. **A.**BTK PH domain titrated with IP4. **B**. Pre-treatment with 1 mM furan fragment **1. C**. Pre-treatment with 1 mM furan fragment without ketone **1z**.

Mixed solvent molecular dynamics (MxMD) were performed to assess the druggability of the BTK PH domain. Given its function as a membrane anchor, the PH domain is a positively charged domain with shallow, charged pockets. Probing the surface of BTK PH domain with three co-solvents (isopropanol, pyrimidine and acetonitrile) 5 % in water uncovered a pocket immediately adjacent to the three position of the cyclohexanone ring of the analogues (with a promising MxMD score of 31000). The pocket has an area of 286 Å2 and a volume of 94 Å3, with strong pyrimidine and isopropanol clustering. Importantly, this highlighted the potential of growing with *ortho* substituted phenyl rings, which productively fill the uncovered pocket with pi-stacking potential and hydrophobicity (Figure S108). Virtual screening and R-group enumeration using this phenyl scaffold also indicated that *ortho* substitutions may be advantageous (Figure S109).

### Development of fragment hit 1 to give parent 2

The crystal structure of **1** showed significant space at the end of the six-membered ring, towards Y106 (Figure 4A, S2 A). Given the flexibility of β1-β2 loop, as seen in the unliganded and IP4 bound structures before, we anticipated a more substantial pocket could open up with a suitable ligand. To explore this, a synthetic route was developed to grow the fragment from the six-membered ring. The most obvious derivative of **1** was to add a phenyl ring to it, yielding **2**. Soaking this into BTK PH domain crystals yielded a high-resolution co-crystal structure of a singly covalently modified target protein. Like fragment **1, 2** shows the crystal ligand covalently bound to K12 (Figure 4B, S2 B) and making a salt bridge with R28, but with the β1-β2 loop moving away from the binding site, making space for the phenyl ring. The phenyl ring is lying on top of L111, sandwiched between β1β2 loop and Y106 sidechain.

**Figure 4.**
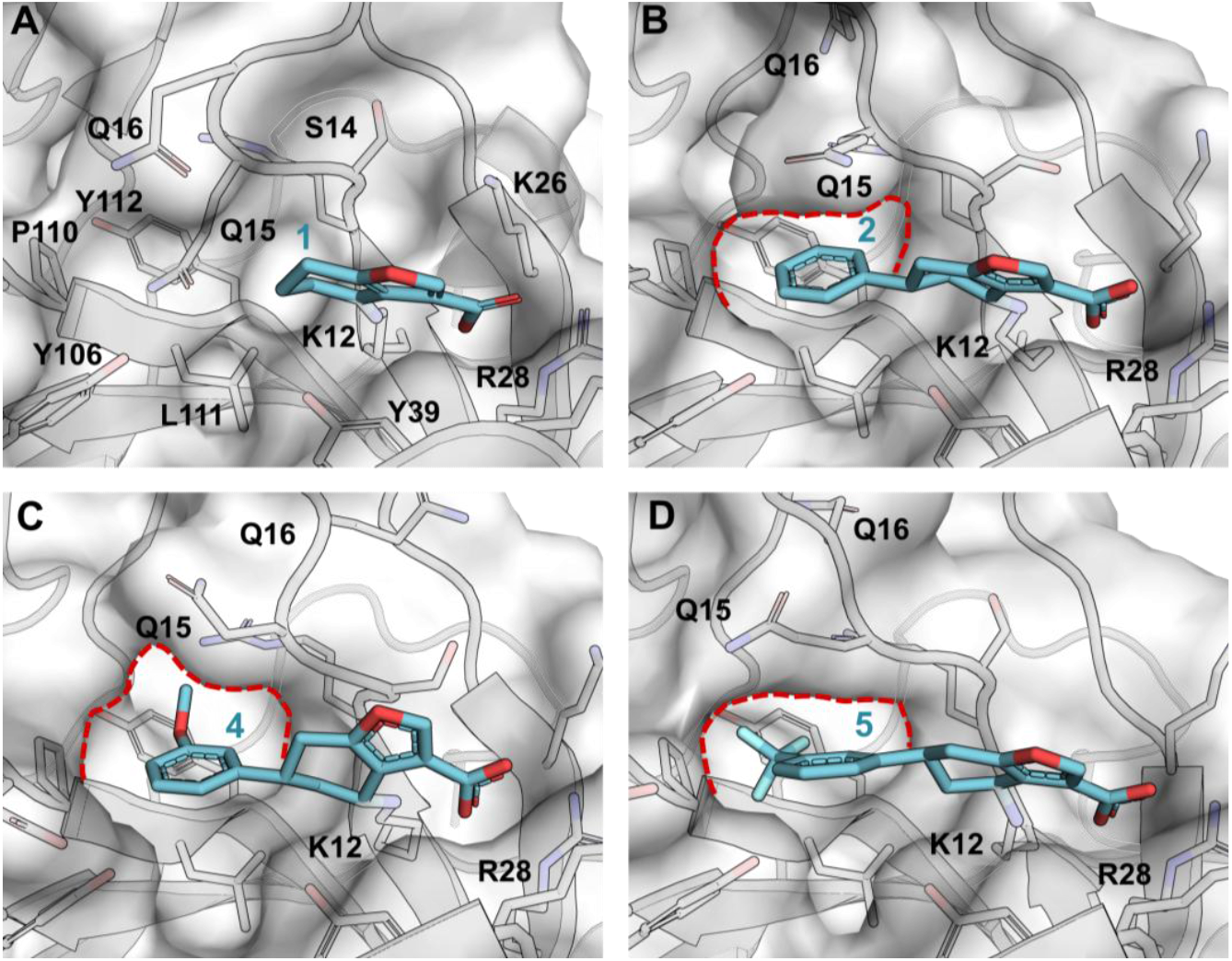
First iteration analogs. **A.**Complex of **1** with BTK PH domain with semi-transparent surface of the domain and key binding site residue shown as sticks and labeled. **B**. Complex of **2** with BTK PH domain in the same view as in A showing expanded pocket to accommodate the phenyl ring. **C**. Complex of **4** with BTK PH domain in the same view as in A. **D**. Complex of **4** with BTK PH domain in the same view as in A. The pocket that opens up with β1-β2 loop moving away is indicated with red dotted line in panels B, C, and D.

Importantly, the phenyl group of **2** presents various potential growing vectors as illustrated by analogs **4** and **5** (Figure 4C,D). Firstly, a large hydrophobic pocket gated by Y39 could potentially hydrogen bond with substituents in the *ortho* position. Varying ring substitution patterns would also perturb electron distribution and could influence C–H···π interactions between the side chain of Q15 and the aromatic system. The potential substitution of the phenyl ring and scaffold hopping to investigate key interactions provided scope for our subsequent structure-binding relationship study and led to the synthesis of several iterations of analogs (Figure 5).

**Figure 5.**
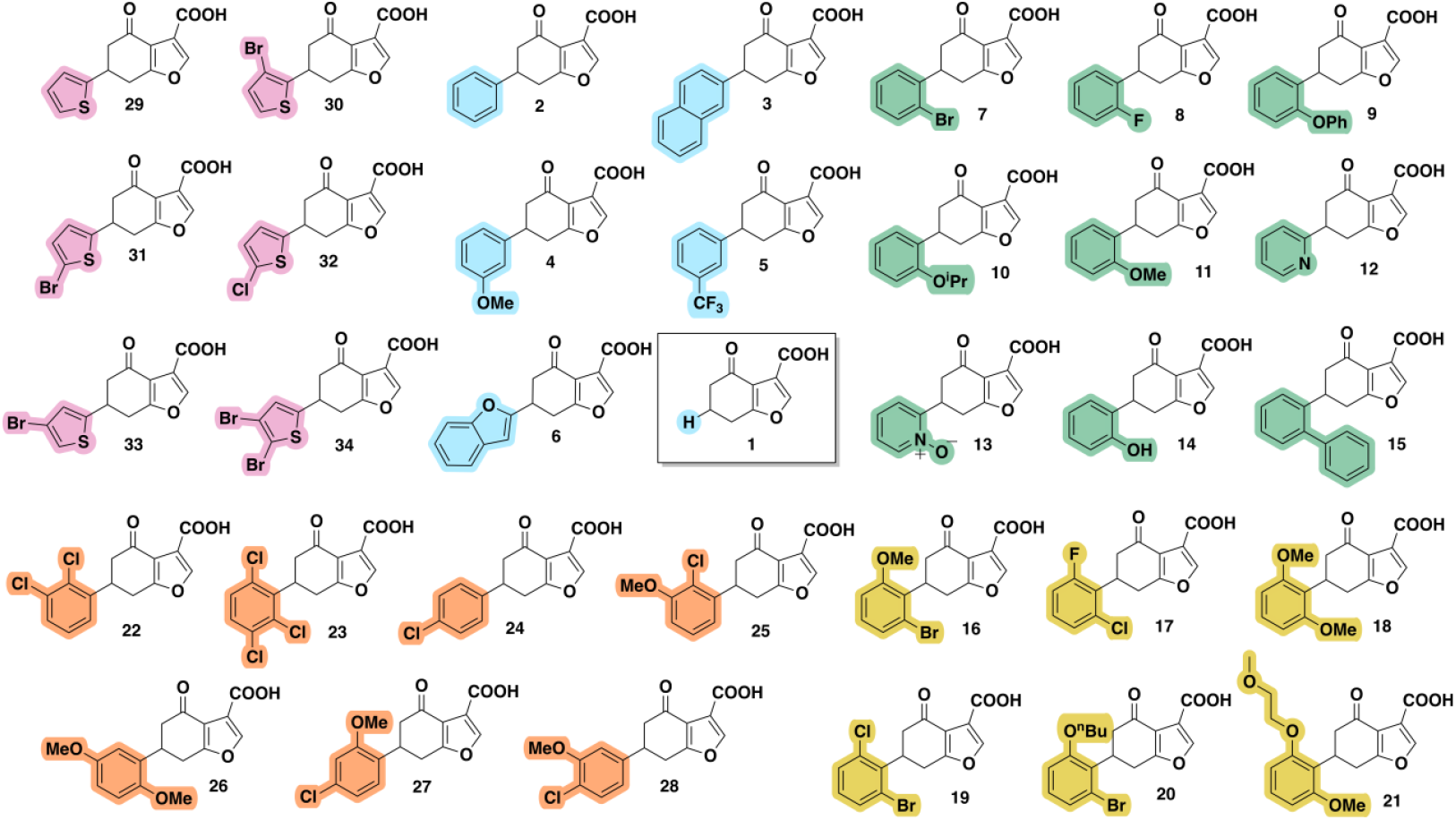
All analogs. The fragment hit **1**, surrounded by first-iteration analogs (compound **2**-**6**, pale blue) exploring fused rings and *meta* substitution, second-iteration analogs (compound **7**-**15**, green) exploring *ortho* substitution, third-iteration analogs (compound **16**-**21**, yellow) exploring 2,6 *ortho* disubstitution, fourth-iteration analogs (compound **22**-**28**, orange) exploring other substitution patterns on a benzene scaffold and fifth-iteration analogs (compound **29**-**34**, pink) exploring substitution on a thiophene scaffold.

### Chemistry

The furan fragment hit **1** could be synthesized in one step from readily available diketone starting material **1c** (**Scheme 1**). Potassium hydroxide generates the enolate of the diketone to attack ethyl bromopyruvate forming a furan ester which is hydrolysed *in situ*.

### Scheme 1. Synthesis of the furan fragment hit 1.^a^

**Scheme 1.**
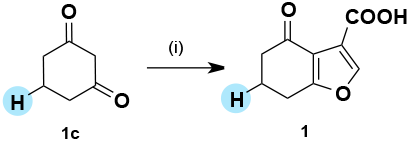
^a^Reagents and conditions (i) ethyl bromopyruvate (2 equiv), KOH (3 equiv), EtOH, rt; 50 wt % NaOH (4.5 equiv).

The synthetic route was extended to access the phenyl parent compound **2**, for which the diketone **2c** had to be synthesized from α,β-unsaturated compound **2b** in three stages (**Scheme 2a**). The diethyl malonate Michael addition product cyclises under the reaction conditions to give the 1,3-cyclohexanedione moiety with an additional ester group after the first stage. The additional ester is removed in the final two stages by hydrolysis and decarboxylation to form diketone **2c** (**Scheme 2b**). As before, the furan **2** is formed with ethyl bromopyruvate, however solubility issues were addressed by using a stronger base (NaOEt) (**Scheme 2c**).

### Scheme 2. a. Synthesis of the parent phenyl 2. b. Reaction pathway towards diketone formation. c. Furan formation from diketone

**Scheme 2.**
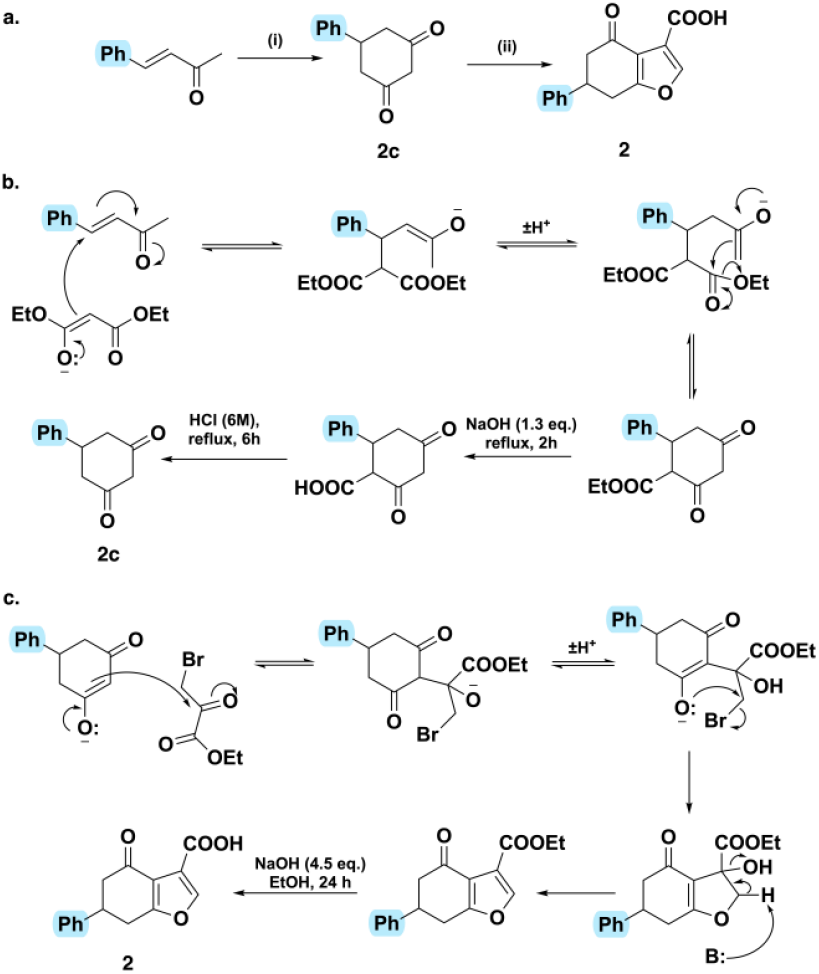
^b^Reagents and conditions (i) diethyl malonate (1.1 equiv), 7 % NaOEt in EtOH (1.1 equiv), rt; 50 wt % NaOH (1.3 equiv), reflux; 1 M – 6 M HCl, reflux; (ii) ethyl bromopyruvate (1.3 equiv), 7 % NaOEt in EtOH (1.1 equiv), rt/reflux; 50 wt % NaOH (4.5 equiv), rt.

One main synthetic route was then developed to obtain subsequent analogs **3** – **12, 16** – **34** (**Scheme 3**). The α,β-unsaturated compounds (**b**) were synthesized from aldehydes (**a**) by Wittig reactions.

In order to interrogate the functionalities on the molecule, matched-pair compounds were synthesized and tested. Diketones **16c, 24c** and **33c** were subjected to furan formation conditions without subsequent hydrolysis to yield esters **16d, 24d** and **33d** respectively (**Scheme 3**). For analog **17**, the electrophilic ketone was reduced to an alcohol to form **17e**. The aldehyde precursors of the O-alkylated analogs **11a, 20a, 21a** were synthesized by adding an alkylation step before the Wittig reaction (**Scheme 4a**). Similarly, the aldehyde for the methoxy analog **16a** could not be obtained commercially so was synthesized by methylating the respective hydroxy form **16f** (**Scheme 4b**). Analogs **13, 14** and **15** were synthesized via oxidation of **12** (**Scheme 4c**), demethylation of **11** (**Scheme 4d**) and cross-coupling with **7**, respectively (**Scheme 4e**). Finally, negative control **37**, based on the structure of alpha tetralone **35** but with a carboxylic acid to mimic that in the fragment hit, was synthesized (**Scheme 5**).

### Scheme 3. General synthetic route to analogs 3-12, 16-34.^c^

**Scheme 3.**
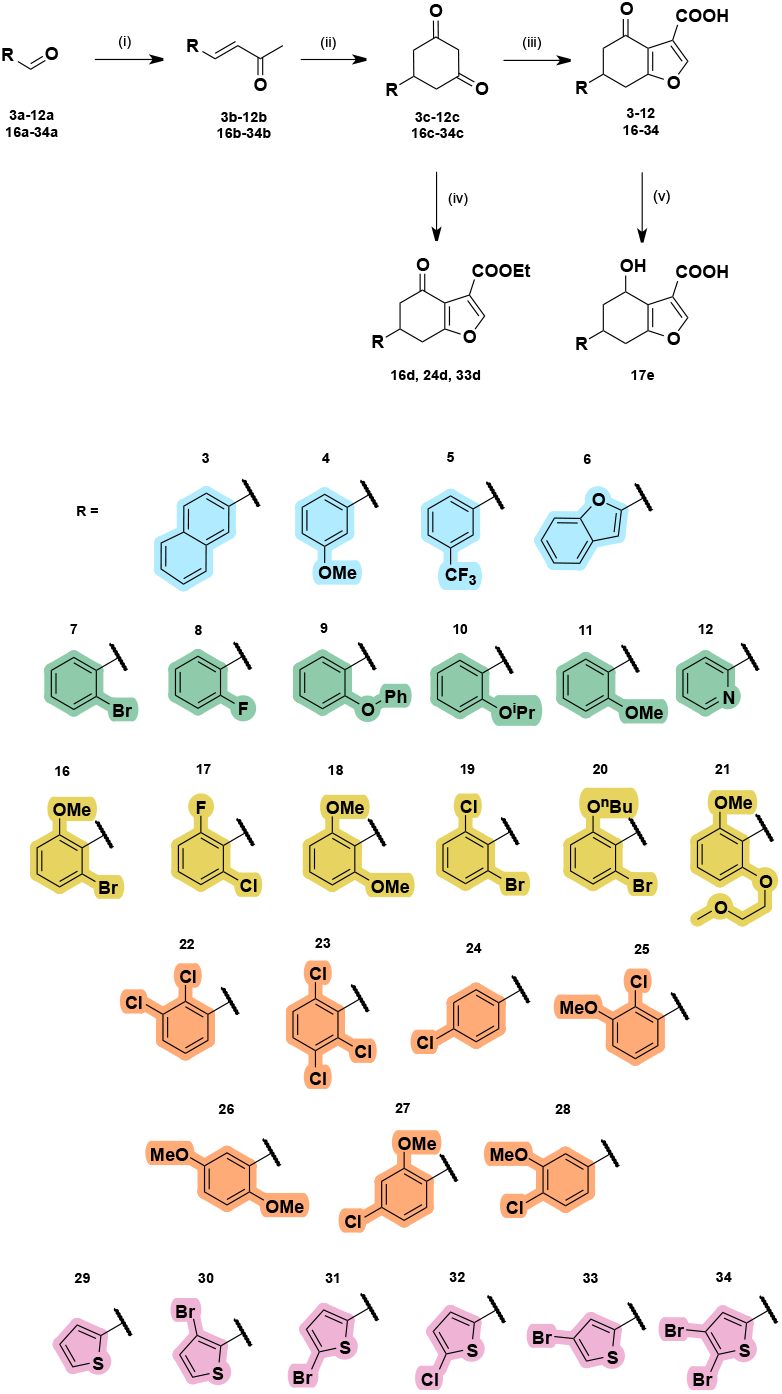
^c^Reagents and conditions (i) 1-(triphenyl-λ5-phosphaneylidene)propan-2-one (1 equiv), THF, reflux; (ii) diethyl malonate (1.1 equiv), 7 % NaOEt in EtOH (1.1 equiv), rt; 50 wt % NaOH (1.3 equiv), reflux; 1 M – 6 M HCl, reflux; (iii) ethyl bromopyruvate (1.3 equiv), 7 % NaOEt in EtOH (1.1 equiv), rt/reflux; 50 wt % NaOH (4.5 equiv), rt; (iv) ethyl bromopyruvate, 7 % NaOEt in EtOH (1.1 equiv), rt/reflux; (v) NaBH_4_ (1.5 equiv), EtOH, 0 °C – rt.

### Scheme 4. Additional synthetic steps towards 10a, 20a, 21a, 16a and 13-15. ^d^

**Scheme 4.**
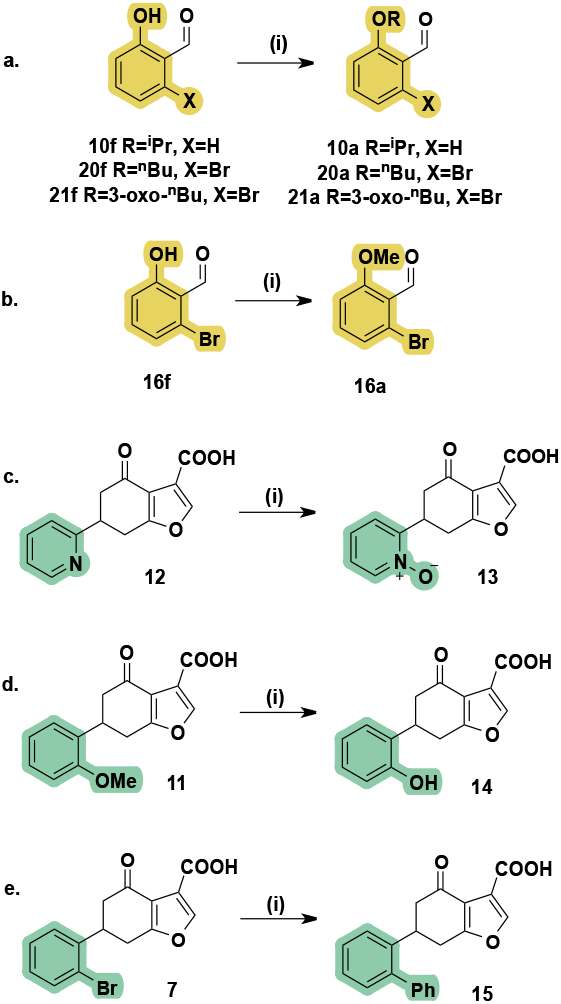
^d^Reagents and conditions **(a)** (i) RBr (1.5 equiv), DMF, K_2_CO_3_ (2 equiv), 0 °C – rt. **(b)** (i) K_2_CO_3_ (3.8 equiv), MeI (17 equiv), acetone, rt. **(c)** (i) mCPBA (1.5 equiv), DCM, rt. **(d)** (i) BBr_3_ (2.3 equiv), DCM, −78 °C – rt. **(e)** (i) Phenyl boronic acid (1.2 equiv), Pd(OAc)_2_ (0.075 equiv), s-phos (0.1 equiv), K_3_PO_4_ (2 equiv), 20:3 toluene/H_2_O, µwave, 60 °C.

### Scheme 5. Synthetic route to negative control 37.^e^

**Scheme 5.**
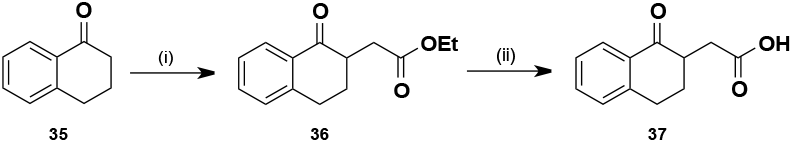
^e^Reagents and conditions (i) LDA (1.1 equiv), ethyl 2-bromoacetate (1.1 equiv), THF, hexane, −60 °C – rt; (ii) 3M NaOH (3 equiv), 2:1 THF/MeOH, rt.

### Structure–Binding Relationship Study

Crystal structures were obtained for 24 analogs (Supporting Information S1 X-ray Crystallography). All show similar binding modes as illustrated with the phenyl parent, **2** (Figure 4B, S2 B). As well as the covalent label at K12, the salt bridge of the carboxylic acid with R28 is preserved in all cases; no crystal structures could be obtained for the ethyl ester precursors **16d, 27d** and **33d**.

The first-iteration analogs (**3-6**) were designed to explore accessibility to a pocket next to the *meta* position of the phenyl ring. Crystal structures were obtained for **4** and **5**, which show the methoxy and trifluoromethyl group expand the pocket opened up by the phenyl group of **2**, with Q15 in β1-β2 loop making room for the ligands. The pocket is lined by P110, L111 and Y112, with the aliphatic part of R13 side chain at the far end (Figure 4, S2 C,D). The fused rings in **3** and **6** appear to be too large or having non-ideal geometry for this malleable pocket and we were unable to get crystal structures with these compounds.

The second-iteration analogs (**7-15**) explored the *ortho* position and yielded promising results with well defined binding pose evident in crystal structures for **7 – 14** (Figure S1, S2 E-L). No crystal structures were obtained for the biphenyl analog **15**, suggesting that the rigid system was not accommodated. However, with an oxygen spacer, the phenoxy substituted **9** projects the phenyl ring further into the pocket lined by R13 and Y106, underneath Q15 (Figure 6A). This pocket is also occupied by the *ortho* halogens in **7** (Figure 6B) and **8**. Notably, the bromine atom in **7** creates a strong water-mediated halogen bond with the backbone carbonyl of R13, indicating a strong preference for bulky, soft halogens in the ortho position. Pyridine-containing analog **12** and its oxidised counterpart, **13**, project their heteroatoms towards the same protein pocket and β1-β2 loop, as does the hydroxyl group of **14**.

**Figure 6.**
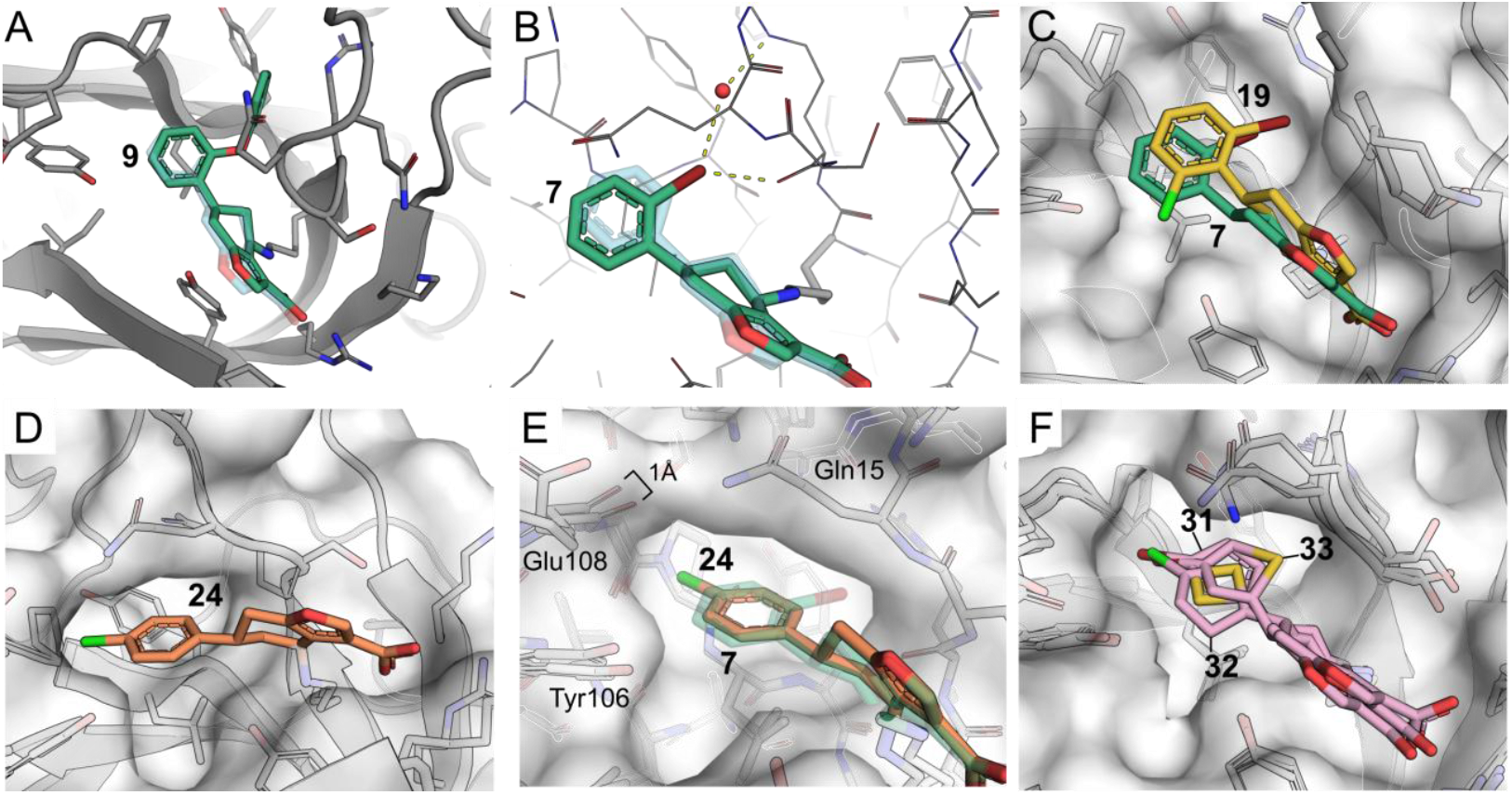
Elaborated inhibitors. **A.**Second-iteration analog **9** (green) overlayed with transparent **2**. An oxygen spacer allows the large phenyl group to point into the hydrophobic pocket under Q15. **B**. Structure of **7** demonstrating *ortho* halogens also occupying this pocket and the possibility of halogen bonds with water or carbonyl of S14. **C**. A third-iteration analog **19** with the larger halogen (Br) points deeper than monosubstituted **7** into the hydrophobic pocket, possibly pushed by the chloro substituent on the other side of the ring. **D**. *Para*-chloro analogue **24** makes a weak halogen bond with backbone carbonyl of E108. **E**. Comparison of **7** and **24** showing how **24** pushes E108 by 1 Å to make room for the *para*-chloro group, with Q15 from the β1-β2 loop folding over **24. F**. Halogenated tiophenes **31, 32** and **33** all form similar halogen bonds with backbone of E108, with thiophene ring flipping depending on the position of the halogen.

The alkoxy substituted analogs behave differently. The isopropoxy group of **10** and methoxy of **11** project in the oppostite direction, towards Y39 (Figure S2 I-J), without making additional contacts with the protein. Given the propensity of *ortho* substituents to point towards either inside (**7** Br, **8** F, **9** OPh, **12** N, **13**, N^+^-O^-^, **14** OH) or outside (**10** OiPr, **11** OMe,) of the protein, a third series of analogs with 2,6-disubstitutions (**16 – 21**) was constructed.

The crystal structures of **16, 17** and **19** demonstrate that larger halogen atoms (bromo or chloro) go into the hydrophobic pocket adjacent to the *ortho* position, while the smaller substituents (methoxy and, respectively, fluoro and chloro) point towards Y39 (Figure 6C). The second substituent causes the phenyl ring to move towards the β1-β2 loop, pushing the loop further back. The halogen interaction made by **7** and **8** with a water is subsequently lost. The symmetric analog **18** with two methoxy groups binds in a similar manner to the halogenated analogs. The methoxy group on the opposite side of the β1-β2 loop is inserted into a small pocket between the terminal hydroxyl groups of Y39 and Y106, lined by I56 and L111 side chains (Figure S2 M-P).

The fourth-iteration analogs explored various chloro and methoxy substitution patterns on the phenyl ring. With adjacent *ortho* and *meta* positions occupied in **22** and **25**, the substituents were forced inside the hydrophobic pocket towards L111. The non-adjacent *ortho* and *meta*-methoxy substituents of **26** oriented in the same way as the monosubstituted analogs **11** and **4**, with the methoxy in the *ortho* position pointing into the small pocket between Y39 and Y106, while the methoxy in the *meta* points towards the hydrophobic pocket by the β1-β2 loop (Figure S2 Q-U).

The crystal structure of **24** reveals that the system tolerates also a chloro substituent in the *para* position, pushing residue E108 by ca. 1 Å, to make space for the halogen, with the phenyl ring shifting further towards Y106 compared to ortho-substitued compounds (Figure 6D,E). With the *para-*substitution β1-β2 loop is also folding over the compound, creating a partially closed cavity for the inhibitor. Retaining the interaction the chlorine makes, a methoxy group was added to decorate the *ortho* position in **27** and it points, expectedly, towards Y106 and shifts the phenyl ring slightly more to that same direction. Interestingly, the *para*-chloro halogen bond allows a *meta*-methoxy, which was previously observed to point towards β1-β2 loop in analogs **11** and **26**, to also point towards Y39 in **28**.

Scaffold hopping at the phenyl ring led to a series of thiophene analogs, beginning with **29**. The introduction of a 3-bromo substituent on the thiophene in **30** allows an interaction with S14, similar to that with the *ortho* halogenated phenyls in **17** and **19**. 4-bromo substitution in **33** flips the ring suggesting the bulky Br atom is not accommodated in the *meta* position without flipping the heterocycle (Figure 6F). In the 5-bromo and 5-chloro thiophene analogs, **31** and **32**, the halogen interacts with a pocket formed by the main chain carbonyls of E108, G109 and P110 and the side chain of Y106. Like **33, 32** is ring flipped relative to the unsubstituted thiophene recapitulating the preference for a *para*-chloro substituted phenyl ring. We were unable to obtain a well defined structure of double-brominated thiophene **34** bound to the target, even though there appears to be room for a second substituent.

Overall, while the binding is dominated by ionic interactions with R28 and covalent linkage to K12, the binding site with its highly flexible β1-β2 loop offers significant opportunities for further elaboration. The β1-β2 loop seems be able to mold itself to many different substitutions, with Q15 either engaging with the ligand directly or moving away from the binding site, as exemplified most dramatically by bromo-substitutions in the *ortho* position of the phenyl ring in (e.g. in **19**), opening a new pocket where large substitutions can be accommodated (Figure 6C).

### Mass Spectrometry

All analogs were tested at a concentration of 150 μM with 2.5 μM of WT BTK PH domain and, with the exeption of the fifth iteration compounds, using R28C mutant as a negative control. The complexes were incubated at 37 °C for five minutes. With compound **2**, these conditions gave rise to mostly a singly labeled covalent ligand-protein complex (Figure 7), while a small proportion of the protein showed a second labeling. All analogs labeled the BTK WT protein and compounds **1**-**4** failed to covalently label the R28C mutant. This suggests that the analogs specifically label K12 and occupy the native inositol phosphate binding site, relying on the key ionic interaction between R28 and the carboxylic acid in the binders, which is disrupted in the R28C mutant.

**Figure 7.**
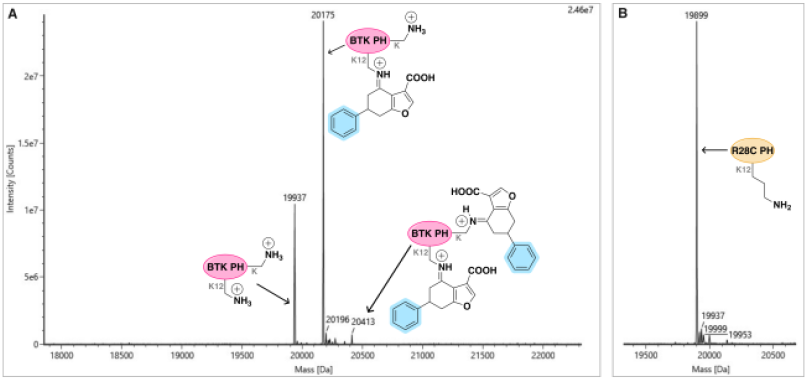
Protein mass spectrometry. Compound **2** (150 μM) labels the WT BTK PH domain (2.5 μM) after five minutes incubation (shaking at 37 °C).

Intact protein mass spectrometry has been previously used for approximating the extent of protein labeling by a covalent inhibitor.^17^ The relative intensities of peaks of mass corresponding to singly, doubly and unlabeled WT protein were recorded after five minutes incubation with the first-to fourth-iteration analogs (Figure 8A, Figures S42-S70).

**Figure 8.**
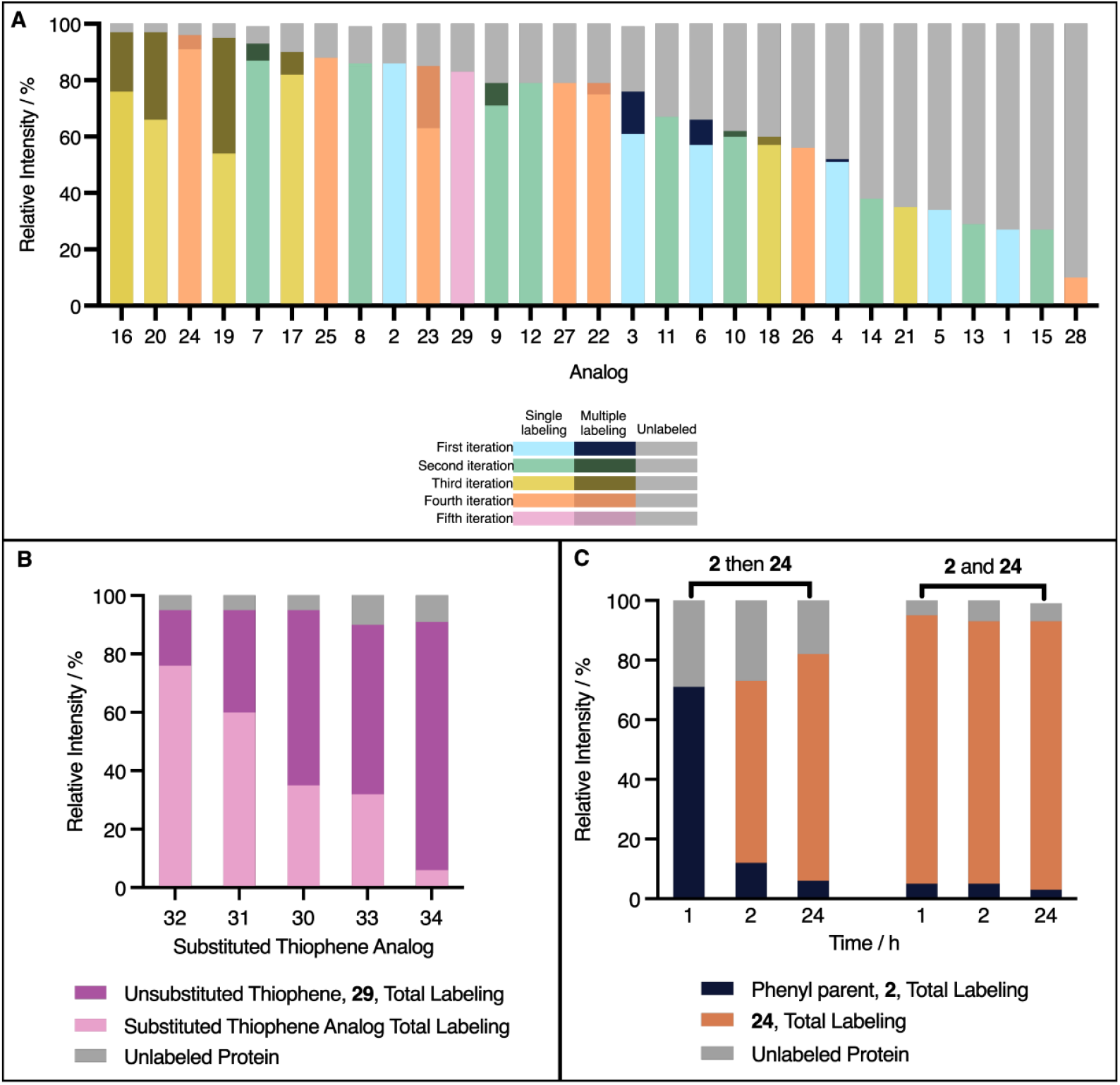
Mass spectrometry studies. **A.**Mass spectrometric evaluation of WT protein labeling after 5 min. First-to fourth-iteration analogs **1** – **28** and thiophene **29** (150 μM) were incubated with WT BTK (2.5 μM) for 5 min, with shaking, at 37 °C. The intensities of peaks were compared within each sample to show the proportion of unlabeled and singly, doubly and triply labeled proteins. **B**. Thiophene analog competition experiments. Each fifth-iteration thiophene analog (150 μM) was incubated along with the unsubstituted thiophene **29** (150 μM) and WT BTK PH domain (2.5 μM) and shaken at 37 °C for 1 h. The intensity of single labeling peaks for the unsubstituted and substituted thiophenes relative to the total protein intensity are shown in dark and pale purple, respectively. **C**. A graph to summarise the results of the three treatments in reversibility studies. The protein (2.5 μM) was incubated at 37 °C for 1 h with either analog **2** (150 μM), analog **24** (150 μM) or both (150 μM each). A mass spectrum was recorded before addition of analog **24** (150 μM), addition of analog **2** (150 μM) or no addition, respectively, and further incubation for 1 h. A second mass spectrum was recorded for each treatment. The samples were incubated at 37 °C with no further additions for another 22 h. In each case, the same equilibrium is reached indicating the reversibility of the labeling.

The total labeling relative intensity of parent analog **2** is almost three times thaf of the initial hit **1** (86% compared to 27%), demonstrating the benefit of introducing the phenyl ring. Fused ring analogs **3** and **6** also show efficient labeling relative to **1**. The other first-iteration analogs display less desirable behavior with *meta* substituents [*meta*-methoxy (**4**) and trifluoromethyl (**5**)] giving relatively low peak intensities for labeled species (52% and 34%, respectively).

The *ortho* analogs of the second-iteration display a range of outcomes. Halogens, bromo **7** and fluoro **8**, exhibit high total labeling relative intensities of 94% and 87% respectively with minimal non-specific binding. However, the alkoxy substituents yield moderate responses, 62% with the isopropoxyl analog **10** and 67% with the methoxy analog **11**. The oxygen spacer in the phenoxy group of **9** improves the relative intensity of total K12 labeling (79%) compared to the larger, more rigid biphenyl system in **15** (27%). Total binding of the pyridine analog **12** (78%) is reduced by oxidation to **13** (29%) and the hydroxy analog **14**, structurally similar to N-oxide **13**, also behaves poorly (38%).

The third and fourth generations achieve relatively high intensities for total labeling, except for disubstituted analogs **18, 21** and **26** (59%, 35% and 56% respectively) which all contain *ortho*-methoxy and alkoxy substituent. Interestingly, *ortho*-bromo methoxy **16** and *ortho*-bromo n-butoxy **20** show the highest relative intensity of total labeling (both 97%) albeit inducing high intensity non-specific labeling (21% and 31% respectively). Disubstituted *ortho*-chloro containing analogs **17, 19, 22**, and **25** performed well (90%, 95%, 78% and 88%) as do disubstituted *para* chloro containing analogs **27** and **28** (79% and 83%) and trichloro analog **23** (84%). Finally, singly substituted *para*-chloro analog **24** shows the highest relative intensity of total labeling (96%) with minimal non-specific binding events (4%).

Scaffold hopping at the phenyl ring to give thiophene analog **29** proved successful and the relative intensities of labeled protein peaks are comparable to the phenyl series in mass spectrometry. The subsequent thiophene analogs (**30**-**34**) were incubated in succession with the unsubstituted thiophene **29** to create a competition study (Figure 8B, Table S2). Based on labeling efficiency, the analogs show clear ranking **32** > **31** > **30** > **33** > **34**, with **31** and **32** out-performing the unsubstituted thiophene **29**.

The reversibility of the reaction was studied by using two ligands in succession. The total labeling relative intensity of the phenyl parent analog, **2**, reached approximately 75% after 1 hour incubation at 37 °C. Subsequent incubation with high performing *para*-chloro analog, **24**, caused the system to re-equilibrate. Irrespective of order of addition, the same distribution of labeling was achieved after 24 h incubation at 37 °C suggesting an equilibrium is reached, indicative of reversible imine formation (Figure 8C. figures S71-79).

The time and pH dependence of protein labeling was further explored to investigate the nature of the covalent bond between the ligand and the protein (Figure S80-S90). Basic pH environments favor the reaction, which was found to reach > 95% completion in less than 20 minutes for pH values of 9 and 7.8. While the reaction is slower at physiological pH 7.4, the protein labeling reaches > 90% completion in less than an hour, highlighting the potential of our compounds for in vivo labeling of K12. As expected for lysine labeling, the reaction becomes significantly slower at pH = 6, where a significant proportion of K12 is protonated, and reaches 50% conversion after 1 h.

Mass spectrometry was also used to determine the effect of scaffold hopping and chirality on binding. All ester (**16d, 24d, 33d**), reduced (**17e**) and alpha tetralone based (**37**) analogs showed no binding to the target (Figures S93-S95). The covalent binding of the parent phenyl, **2**, was further confirmed using MALDI (Figure S100).

To evaluate the overall reactivity of lysines in BTK PH domain, we used a non-specific electrophilic lysine label 4-methylbenzenesulfonyl fluoride **38** (150 μM), as a probe. We observed no covalent adduct when the reaction was incubated at 37 °C for five minutes (Figure S96). With increased concentration, up to 1 mM, and overnight incubation at room temperature, **38** gave basrely detectable labeling (Figure S97) This is in stark contrast to the nearly 100% labeling by **24** under the same conditions (Figure S98).

### Differential Scanning Fluorimetry

The analogs were screened against the WT BTK PH domain (Supporting Information S6.2, Figure S101A) and the R28C mutant (Supporting Information S6.2, Figure S101B). The upper limit of the ligand concentration was around 1.5 mM in the assay, constrained by solubility of the compounds. All analogs induce different thermal shift profiles in the WT versus R28C, indicating the specificity of binding in the IP4 binding site, consistent with the requirement for the salt bridge between R28 and the inhibitor carboxylate for labeling of K12.

The first-iteration analogs (**3** and **5**) induce small shifts in protein melting point (T_m_) compared to parent **2** (ΔT_m_ at 750 μM ligand = 7 °C), indicating weaker binding to the protein (Table S5). Higher T_m_ shifts (ΔT_m_ at 750 μM ligand = 7.5–10.5 °C) are seen for all second-iteration analogs (**7** – **15**) except the phenoxy and biphenyl analogs, **9** and **15** (ΔT_m_ at 750 μM ligand = 1 °C and 3 °C respectively). Notably, **16, 17** and **19** in the third iteration produce even larger shifts (ΔT_m_ at 750 μM ∼ 14 °C), with selectivity confirmed by the R28C mutant. The fourth-iteration analogs (**22** – **28**) induce significant stabilization of the protein (all ΔT_m_ at 750 μM > 8 °C) with the exception of **22** and **28**, which induce small shifts (ΔT_m_ at 750 μM = 3-5 °C) performing poorly as in mass spectrometry.

K_d_ values were approximated from the DSF data to be used only as a metric for comparison with mass spectrometry. Encouragingly, approximate log(K_d_) correlates well with total labeling relative intensities from mass spectrometry (Pearson, r = −0.56, R squared = 0.32, P value [two tailed] = 0.003, Figure S102), indicating higher K_d_ values for compounds with low total labeling relative intensities.

To further evaluate the importance of the covalent interaction between the compounds and the target, we synthesized **17e**, an analog of **17** in which the ketone has been reduced to an alcohol (Figure 9). This compound failed to induce a potent positive response in the WT BTK, instead inducing a negative shift of – 5 °C. This suggests **17e** engages the with the PH domain in an alternative binding mode, confirming the ketone moiety is essential for covalent binding at K12 through an imine forming carbonyl group.

**Figure 9.**
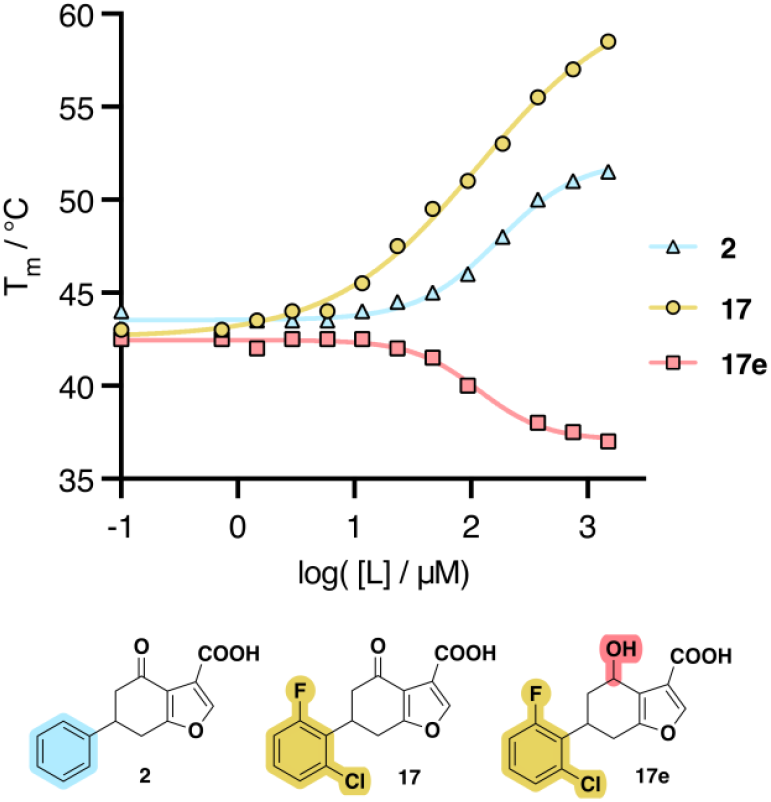
Differential scanning fluorimetric evaluation of binders. A dose-response curve of selected inhibitors, measured as change in melting temperature, T_m_, measured using 5 μM BTK PH domain.

From the crystal structures, the stereochemistry at the chiral carbon in the binders cannot be ascertained, thus the enantiomers of **19** were separated to investigate binding. As expected, the enantiomers (**19p1** and **19p2**) were shown not to interconvert in HEPES buffer (Figures S103-S107). **19p1**, showed a higher total labeling after five minutes than the other **19p2** in protein mass spectrometry (Figure S99), which was in line with the electron density in X-ray crystallography.

## DISCUSSION

We herein report the discovery of a novel mode of targeting BTK through its PH domain. A reversible covalent binding mode was uncovered, with crystallographic analysis proving an iminium bond formation through K12 with the ketone electrophile.

X-ray crystal structures of BTK PH domain with various analogs demonstrate how the binding mode of the furan fragment is consistently determined by the imine formation with K12 and the salt bridge interaction with R28 – both of which are key residues interacting with PH domain ligand IP4.

The phenyl ring of the analogs is versatile with respect to the orientation of its *ortho* substituents relative to the protein backbone. Soft halogens (bromine and chlorine) in **17** and **19** point towards the protein backbone, with bromine displaying a water-mediated halogen bond with the carbonyl of Arg13. The large phenoxy substituent in **9** unexpectedly points in the same direction, albeit with notable β1-β2 loop rearrangement (Figure 10). On the other hand, alkoxy substituents (methoxy, isopropoxy) and hydroxyl substituents orient their donor atoms towards the hydroxyl of Tyr106, indicating a propensity towards H bonding.

**Figure 10.**
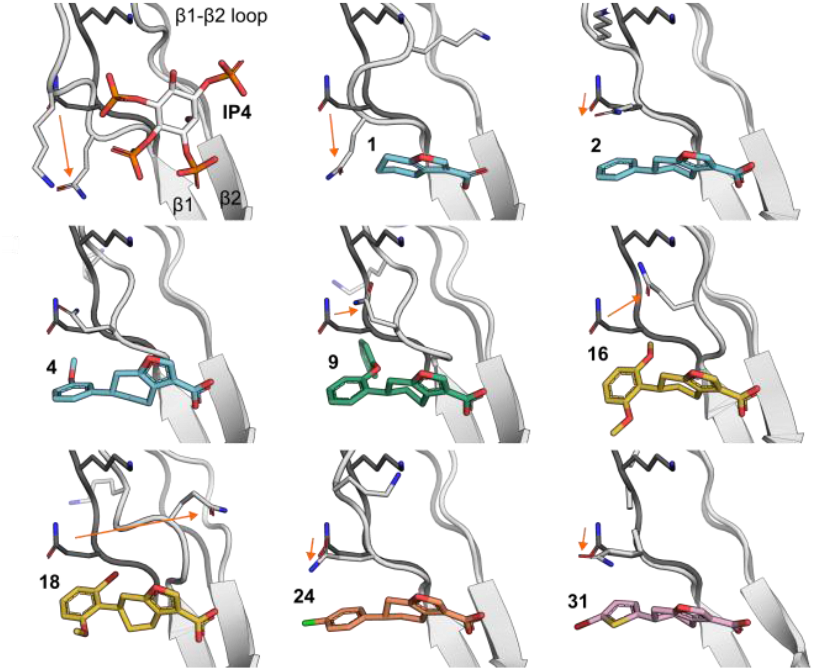
Movement of β1-β2 loop. Representative compounds in complex with BTK PH domain are shown with β1-β2 loop and Q15 residue. In each panel the β1-β2 loop with **1** is shown with darker grey and movement of Q15 in each complex relative to that is indicated with a red arrow.

The trends at the *meta* position are less defined with respect to the in-or-out positioning relative to the protein backbone. Notably, the methoxy and trifluoromethyl groups in **4** and **5** point inwards and induce a pocket via β1-β2 loop rearrangement, whereas in **28** the *meta*-methoxy group points towards the solvent.

The *para* position emerges as a privileged one for halogens, with the chlorine in **28** forming a halogen bond with the the backbone carbonyl of Gly109. Thiophene analogs in the fifth generation proved to be potent bioisosteres of the phenyl ring and are well tolerated, while introducing a slightly different angularity around substituents.

One of the key features in our exploration of structure-activity relationship around our compounds is the movement of β1-β2 loop. It has been seen to be highly mobile and adopt significantly different conformations in unliganded and IP4-bound structures.^18^ Depending on the compound presented here, the β1-β2 loop can either engage with the compound or move away, exposing a wider pocket underneath it where substituents can go, with Q15 as one of the most mobile residues (Figure 10).

This work demonstrates the feasibility to design small molecule inhibitors for PH domains in general and acts as a springboard for the development of more potent and selective BTK PH domain inhibitors. We expect this also to act as a proof-of-concept for inhibition of the PH domain more broadly.

In general, clear trends emerged from MS and DSF, indicating that certain bulky groups or ring systems are not tolerated on the phenyl ring, while halogens and alkoxy substituents are. Straight carbon to nitrogen swaps are also tolerated in the ring. The binding specificity of the analogs was confirmed by the point mutant R28C which ablates the key arginine-carboxylate salt bridge. Mass spectrometry studies converge with DSF studies and point to *ortho* and *para* halogen substituted phenyl rings being the most beneficial for binding. The covalent labeling was proven to be highly dependent on the pH of the reaction buffer, with higher pHs inducing faster kinetics. The reversible covalent nature of the binding was proven by the reversibility study, in which **24** fully labeled the protein regardless of the order of addition.

This novel class of BTK PH domain binders represents an important starting point for further development. With binding specificity validated by crystallography, MS and DSF, further medicinal chemistry modifications have the potential to yield potent and selective BTK inhibitors, complementary to the current ATP binding site targeting ones.

Two obvious questions arise from this as well. Firstly, why is Lys12 modified? Is it particularly reactive? Following on from that, could this be more generalized method for targeting PH domains?

To answer the first question, we analyzed 12 BTK PH domain structures (PDBs 1B55, 1BWN, 2Z0B, 4Y94) using two computational pKa calculation methods, DeepKa^19^ and pyPKa^20^. Both methods predicted significantly lowered pKa compared to the other 14 lysines in this domain. pKa of K12 was predicted to be 7.05 ± 1.29 and 8.82 ± 0.80, with the two methods, respectively, while other lysines had average pKas of 10.50 ± 0.97 and 10.77 ± 0.90 (Figure S110, Table S10). This suggests K12 is particularly reactive in BTK PH domain. To answer the second question, we analyzed also other PH domains (Akt, Grb1, DAPP1 and pRex1) for which structures are available and which bind inositol phosphates in the same site using the DeepKa server. In all cases, the lysines that are equivalent to K12 in BTK PH domain were unique in having their pKa values 2.2-3.2 units lower than other lysines in those structures. This suggest the covalent approach we describe here as a more universal approach to PH domain inhibition.

## CONCLUSIONS

In conclusion, we developed a novel class of lysine targeting covalent binders against the PH domain of BTK, a previously unreported strategy. Five families of analogs were generated in a structure-binding relationship study incrementally guided by emerging X-ray crystallographic data. Following on from the unsubstituted phenyl of parent compound **2**, *ortho* and *meta* substituents were explored, as well as fused rings. *Meta* substituents and fused rings turned out to be detrimental for binding, while *ortho* substituents proved to be beneficial (particularly halogens and alkoxy groups).

The binding selectivity was validated by protein mass spectrometry and differential scanning fluorimetry, whereby all analogs induce expected responses with the WT BTK, but fail to induce a response with the loss of function mutant R28C. The DSF results suggest an increase in binding affinity of slightly more than two orders of magnitude from hit compound **1** to lead compound **24**.

## METHODS

### Expression and purification

WT and R28C mutant of the BTK PH-Btk motif construct (residues 1-170, Uni-prot:Q06187) was expressed and purified as described previously^5^, with the exception that final ion exchange chromatography could be left out without compromising the quality of the preparations. To facilitate crystallization, a variant form of the protein was produced in which exposed C145 in the TH domain was mutated to a serine or alanine, creating proteins Btk_S and Btk_A, respectively.

### X-ray crystallography

Btk_wt/Btk_S/Btk_A were crystalized at 20-25 mg/mL in 100 mM NaCl, 20 mM CHES-NaOH, pH 9.5. The reservoir used was 0.1 M TRIS 8.5 pH, 32.5% w/v PEG 3350, 200 mM MgCl_2_ in a 1:0.9 ratio with a total volume of 0.4 μL by the site-in drop method using the mosquito robotics system (SPT Labtech). In the absence of suitable crystals, previously described crystals of R28C mutant were used as the start for serial seeding process.^5^ Seeds were generated from past crystal screens and diluted in 0.1 M TRIS 8.5 pH, 32.5% w/v PEG 3350, 200mM MgCl_2_. Seeds were added to the drop at 0.01 μL using the mosquito robotics system (SPT Labtech) The fragments were soaked as singletons at 2-100 mM into these crystals for 15–20 h in 0.1 M TRIS 8.5 pH, 32.5% w/v PEG 3350, 200mM MgCl_2_ after which the crystals were cryo-cooled in liquid nitrogen for data collection. X-ray diffraction data was collected at Diamond synchrotron radiation sources and then processed using the pipedream package by Global Phasing Ltd; structures were solved using Phaser^21^ from the CCP4 package.^22^ Models were iteratively refined and rebuilt by using Refmac^23^, Buster^24^ and Coot^25^ programs. Ligand coordinates and restraints were generated from their SMILES strings using the AceDRG^26^ software from the CCP4 package.

### Isothermal Titration Calorimetry

All ITC experiments were performed at 25 °C using a MicroCal iTC200 instrument (GE Healthcare). WT BTK PH domain (20 mg/mL, 50mM HEPES pH 8.0, 100mM NaCl) was diluted in HEPES buffer (50mM HEPES pH 8.0, 100mM NaCl, 5% DMSO, **1**/**1z** 1mM/0mM) and concentrated to 5 μM of WT BTK PH domain. IP4 in 10 mM stock solutions was diluted into the buffer to 100 μM (50mM HEPES pH 8.0, 100mM NaCl, 5% DMSO, 1/1z 5mM/0mM), ensuring that the DMSO concentrations were carefully matched. In a typical experiment WT BTK PH domain (5 μM) was loaded into the sample cell and 100 μM of the ligand was titrated in eighteen 2 μL injections of 2 s duration at 150 s intervals, stirring at 750 rpm. Heats of dilution were determined in identical experiments, but without protein in the cell. The data fitting was performed with a single-site binding model using the Origin software package.

### Protein MS experiments

2.5 μM of BTK PH domain was incubated with 150 μM ligand(s), 20 mM HEPES buffer, 100 mM NaCl and 0.5 mM TCEP, pH 8.0 in 25 µL total volume. The DMSO concentration was kept constant throughout all experiments, at 1.5 %. The samples were then incubated at 37 °C either for five minutes (individual ligand experiments) or one hour (competition ligand experiments). All samples were injected directly into the liquid chromatography mass spectrometer and analyzed. Protein MS measurements were performed on a Waters LCT Premier Time of Flight mass spectrometer, with errors within ± 5 ppm. All mass spectra are shown in supplementary data, Figures S42-S70.

### Defining Relative Intensity Terms from MS

The un-, singly, doubly and triply labeled protein peaks can be described as P, P+L, P+2L and P+3L with intensities of *I*_*P*_, *I*_*P*+*L*_, *I*_*P*+2*L*_ and *I*_*P*+3*L*_ respectively.

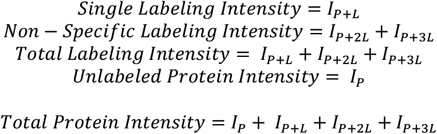

Under the assumption that the primary label is happening in all labeled cases, the relative intensities can be calculated (by dividing by the total protein intensity) as follows:

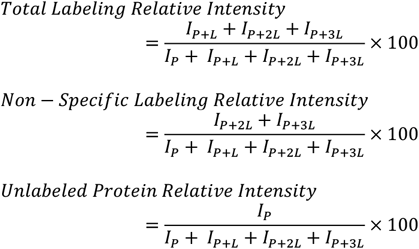

### MALDI analysis

ZipTip® Pipette tips (Merck Millipore, C18 resin, bed volume 0.6 μL, tip volume 10 μL) were washed by aspirating and dispensing MeCN (2 x 10 µL) and 98 % H_2_O, 2 % MeCN, 0.1 % TFA (2 x 10 µL). The protein (2.5 μM) was bound by aspirating samples (10 µL) from mass spectrometry 3 - 7 times. The tip was washed with 98 % H_2_O, 2 % MeCN, 0.1 % TFA (2 x 10 µL). The sample was eluted by aspirating and dispensing 3 uL of matrix solution, 2-cyano-4-OH-cinnamic acid in 30 % MeCN: H_2_O (0.1 % TFA). This was pipetted (1.5 - 2 µL) on a stainless steel target plate, which was dried under vacuum and analysed on a Bruker UltrafleXtreme MALDI-TOF.

### Differential Scanning Fluorimetry

Protein (5 μM) was mixed with corresponding ligand (range of concentrations), in the presence of SYPRO orange (8 x), in pH 8 HEPES (20 mM) buffer. The total concentration of DMSO was 5% in all the samples. The total volume of each sample was 25 μL. The samples were incubated in a RT-qPCR machine fitted with a fluorescent plate reader. Fluorescence was constantly measured while the temperature increased following a gradient defined as follows: 15 minutes at 25 °C, then a constant increase of 1.15 degrees/minute between at 25 °C and 100 °C. DSF measurements were performed on a Bio-Rad CXF96 RT-qPCR machine, with fluorescent readings taken every 15 seconds.

### Determination of approximate K_d_ values from DSF

The T_m_ values for each concentration of ligand were converted to ΔT_m_ values, by subtracting T_m_ with no ligand present. Non-linear regression for “One site – Specific Binding” (performed in Prism10; GraphPad Software Inc.) fit the data to the simplest saturation model, Y = B_max_*X/(K_d_ + X).

### Statistics

For statistical comparisons of total labeling relative intensity from MS and approximate log(K_d_) values from DSF, normal distribution of data was confirmed using Shapiro−Wilk’s tests and analyzed using Pearson correlation tests; P value (two-tailed) < 0.05 is significant. Analysis was performed in Prism10; GraphPad Software Inc.^27^

### Mixed solvent molecular dynamics (MxMD)

Simulations were conducted in Maestro (Schrödinger, release 2025-2) using Desmond with the OPLS4 force field to probe ligandable hotspots on the prepared protein. The holo structure was protein-prepared, then stripped of ligands, waters and nonessential cofactors; essential ions/waters were retained as needed. For each selected probe (default set: acetonitrile, isopropanol, pyrimidine), independent systems were built by placing a ∼7 Å shell of the cosolvent around the solute and solvating in water to achieve ∼5 % cosolvent (v/v). Each system was minimized and equilibrated by the built-in MxMD workflow, followed by production MD on Linux GPU nodes; frames were saved for analysis. Probe occupancy maps were computed on a 3 Å grid, clustered into probe “spots,” and merged across probes to define “hotspots.” Hotspots were ranked by the MxMD score (aggregate probe occupancy) and characterized by surface area, volume and probe composition; predicted hotspots were compared to the co-crystallized ligand.

### Virtual screening and R-group enumeration

Covalent docking, virtual screening and R-group enumeration were performed in Maestro (Schrödinger, release 2025-3) using Glide and Prime. The receptor (BTK PH domain, PDB 7I9I) was protein-prepared (hydrogens added, termini capped, missing loops built with Prime), and ligands were standardized and converted to 3D with LigPrep at physiological pH. A covalent docking model was defined with Lys12 as the reactive residue and a predefined chemistry for imine condensation; the cocrystal-derived template was first docked in Pose Prediction (thorough) mode to validate covalent bond formation and pose fidelity, with outputs ranked by Prime energies and Glide cdock metrics. For library design, a template scaffold was enumerated at the *ortho, meta* and *para* positions of the phenyl ring, using a curated R-group library to generate analogs. This was followed by LigPrep and covalent docking in Virtual Screening (fast) mode under the validated receptor grid and reaction settings. Docked complexes were analyzed in Pose Viewer and cdock affinity/docking score distributions were compared across *ortho, meta* and *para* substitution patterns to assess substituent positioning, pocket compatibility and covalent pose quality.

### Synthetic Chemistry

All experiments were performed in oven-dried glassware and under an atmosphere of nitrogen, unless stated otherwise. Commercial starting materials were used without further purification. Dry solvents were distilled from mixtures containing CaH_2_ or LiAlH_4_ as drying agents. Yields refer to spectroscopically and chromatographically pure compounds unless otherwise specified. Analytical thin layer chromatography (TLC) was carried out on glass Merck Kieselgel 60 F254 plates. The plates were visualised under direct UV irradiation (254 nm). R_*f*_ values are quoted to the nearest 0.1. Preparative thin layer chromatography was performed on commercially available Analtech plates. Flash column chromatography was undertaken on silica gel 60 (230-400 mesh) under a positive air pressure. The eluent systems are reported as % (v/v) of the solvent components.

Reversed-phase column chromatography was carried out using a Combiflash Rf200 automated chromatography system with Redisep® reverse-phase C18-silica flash columns (20-40 μm). Preparative high-performance liquid chromatography (HPLC) was performed on an Agilent 1260 infinity machine. The samples were eluted using a Supelcosil ABZ+PLUS column (250 mm × 21.2 mm, 5 µm). The used linear gradient (for 20 min and a flow rate of 20 mL/min) was: solvent A - 0.1% (v/v) TFA in water, solvent B - 0.05% (v/v) TFA in MeCN. The diodes used the wavelength of 220 nm and 254 nm in order to detect absorbance. HPLC traces of all key compounds are shown in supplementary data, Figures S4-S41.

^1^H NMR spectra were recorded under an internal deuterium lock at rt on Bruker Advance III HD (400 MHz, 500 MHz, 700 MHz; Smart probe). Assignments are supported by ^1^H-^1^H COSY, ^1^H-^13^C HSQC and ^1^H-^13^C HMBC spectra. Chemical shifts (**δ**) are given in ppm quoted to the nearest 0.01 ppm (**δ**_**H**_). The residual solvent peaks are 7.26 for CDCl_3_, 5.32 for CD_2_Cl_2_, 3.31 for CD_3_OD and 2.51 for (CD_3_)_2_SO. Coupling constants for mutually coupling protons are reported in Hertz, rounded to the nearest 0.1 Hz. Data are reported as: chemical shift, multiplicity (br, broad; s, singlet; d, doublet; t, triplet; q, quartet; m, multiplet; or a combination of them), coupling constants, number of nuclei. Spectra were processed using TopSpin v.4.0.6(Bruker) or MestReNova v 14.3.3-33362 (Mestrelab Research S.L.). ^13^C NMR spectra were recorded using an internal deuterium lock at rt on Bruker Avance III HD (101 MHz) with broadband proton decoupling. Chemical shifts (**δ**_**C**_) are quoted to the nearest 0.1 ppm and the solvent reference peaks (in ppm) are 77.2 (CDCl_3_), 53.5 (CD_2_Cl_2_), 49.1 (CD_3_OD), 33.0 (CD_3_)_2_SO). ^19^F NMR spectra were recorded using an internal deuterium lock at rt on Bruker Avance III HD (101 MHz) with broad-band proton decoupling. Chemical shifts (**δ**_**F**_) are quoted to the nearest 0.1 ppm. For fluorine containing compounds, data are reported as: chemical shift, multiplicity, coupling constant. Spectra were processed using TopSpin v.4.0.6(Bruker). All key spectra are shown in the supplementary data file.

High resolution mass spectrometry (HRMS) was performed on a Waters LCT Premier Time of Flight mass spectrometer, with errors within ± 5 ppm.

Analysis of chiral species was performed analytically with the Agilent 1260 Infinity II supercritical fluid chromatography (SFC) system: Chiralpak IC-3 (0.3 cm × 10 cm). Mobile phase: sc-CO_2_/2-propanol = 70/30, flow rate = 1.2 mL/min, wavelength = 254 nm, column temperature = 40 °C. For preparative separation, the chiral species were dissolved in 1:2 MeOH/DMSO and purified on Sepiatec using the following SFC conditions: Column: Chiralpak AD, 21 x 250 mm, 5 micron, Mobile phase: 40% MeOH / 60% scCO_2_, Flow rate: 60 ml/min, BPR: 120 bar, Column temperature: 40 °C, UV max 265 nm.

General synthetic procedure for aldehydes **10a, 20a, 21a**. To a stirred suspension of K_2_CO_3_ (2 eq.) in DMF was added corresponding aldehyde (1 eq., 0.4 M) and the solution was left stirring at room temperature for 1 h. Alkyl halide (1.5 eq.) was added dropwise, on ice, at a rate of 0.1 mL/min and the solution stirred at room temperature for 24 h. Subsequently, the reaction was refluxed for 8 h until full conversion of the starting material. The solution was filtered and subsequently solvent removed *in vacuo*, before purification by silica gel chromatography (hexane / EtOAc and mixtures) to yield the corresponding aldehydes **a**.

General synthetic procedure for enones **3b** – **12b, 16b** – **34b**.

A solution of 1-(triphenyl-λ5-phosphaneylidene)propan-2-one (1 eq.) and corresponding aldehydes **a** (1 eq., 0.4 M) in THF was refluxed for 48 h. The solvent was subsequently removed *in vacuo* before purification by silica gel chromatography (hexane / EtOAc mixtures) to yield the corresponding α,β unsaturated ketones **b**.

General synthetic procedure for diketones **2c** – **12c, 16c** – **34c**.

To a stirred solution of 7% EtONa in EtOH (1.1 eq.) was added a solution of diethyl malonate (1.1 eq., 1.2 M in EtOH) dropwise, on ice, at a rate of 0.4 mL / min. The solution was subsequently left stirring at room temperature for 30 minutes. A solution of corresponding α,β unsaturated ketone **b** in EtOH (0.8 M) was added dropwise on ice, at a rate of 0.4 mL/min. The solution was stirred at rt for 24 h, and subsequently a solution of 50% wt NaOH (1.3 eq.) was added. The resulting mixture was refluxed for 3 h and the solvent subsequently removed *in vacuo*. 6 M HCl (150 ml) was added and the mixture subsequently refluxed for 6 h. The solution was extracted with EtOAc, dried over anhydrous Na_2_SO_4_ and the solvent removed *in vacuo*. The mixtures were purified by reversed-phase column chromatography (2% HCOOH in H_2_O / MeCN 9:1 – 1:1) to yield the corresponding 1,3 diketones **c** as white powders.

General synthetic procedure for carboxy furans **2** – **12, 16** – **34**.

To a stirred solution of 7% NaOEt (1.1 eq.) was added a solution of corresponding 1,3 diketone **c** (1 eq., 0.4 M in EtOH) dropwise, on ice, at a rate of 0.4 mL/min. The mixture was subsequently stirred at rt for 30 min before ethyl bromopyruvate^†^ (1.3 eq.) was added dropwise, on ice, at a rate of 0.4 mL/min. The mixture was subsequently stirred at rt for 24 h before 50% NaOH (4.5 eq.) was added. The reaction was subsequently stirred at rt for 24 h before the solvent was removed *in vacuo* and the resulting mixture dissolved in water. The mixture was acidified to pH 1 using HCl 3 M. The solution was extracted with EtOAc, dried over anhydrous Na_2_SO_4_ and the solvent removed *in vacuo*. The mixtures were purified by reversed-phase column chromatography (H_2_O / MeCN 9:1 – 1:1) to yield the corresponding carboxy-furans as white powders.

General synthetic procedure for esters **16d, 24d, 33d**.

To a stirred solution of 7% NaOEt (1.1 eq.) was added a solution of corresponding 1,3 diketone **c** (1 eq., 0.4 M in EtOH) dropwise, on ice, at a rate of 0.4 mL/min. The mixture was subsequently stirred at rt for 30 min before ethyl bromopyruvate^†^ (1.3 eq.) was added dropwise, on ice, at a rate of 0.4 mL/min. The mixture was subsequently stirred at rt for 24 h before adding water and acidifying to pH 3 using 3 M HCl. The solution was extracted with EtOAc, dried over. anhydrous Na_2_SO_4_ and the solvent removed *in vacuo*. The mixtures were purified by reversed-phase column chromatography (H_2_O / MeCN 9:1 – 1:1) to yield the esters as white powders.

4-oxo-4,5,6,7-tetrahydrobenzofuran-3-carboxylic acid (**1**) (3.3 mg, 18 µmol, 2%) ^**1**^**H NMR** (400 MHz, CDCl_3_) δ 13.22 (s, 1H), 8.08 (s, 1H), 2.99 (t, *J* = 6.3 Hz, 2H), 2.69 (dd, *J* = 7.1, 5.9 Hz, 2H), 2.29 (p, *J* = 6.4 Hz, 2H). ^**13**^**C NMR** (101 MHz, CDCl_3_) δ 199.42, 170.67, 161.50, 150.40, 118.20, 117.47, 36.69, 23.40, 22.55. **HRMS** (ESI+): m/z [M+H]^+^ calculated for C_9_H_9_O_4_ 181.0501, found: 181.0495, error: −3.3 ppm. **HPLC purity** 100%.

5-phenylcyclohexane-1,3-dione (**2c**) (221 mg, 1.17 mmol, 12 %). ^**1**^**H NMR** (500 MHz, CD_3_OD) δ 7.33 – 7.23 (m, 5H), 3.37 (m, 1H), 2.68 (dd, *J* = 17.1, 11.7 Hz, 2H), 2.55 (dd, *J* = 17.1, 4.7 Hz, 2H). 5 % open form impurity [δ 1.99 (s, methyl ketone)]. ^**13**^**C NMR** (125 MHz, CD_3_OD) δ 143.0, 128.3, 126.6, 126.5, 102.7, 39.5, 39.4. 3-carboxy-6-phenyl-4,5,6,7-tetrahydrobenzofuran-4-one (**2**) (3.4 mg, 13 µmol, 2 %). ^**1**^**H NMR** (500 MHz, CDCl_3_) δ 13.14 (s, 1H), 8.15 (s, 1H), 7.41 (t, *J* = 7.8 Hz, 2H), 7.36 – 7.31 (m, 3H), 3.70 (m, *J* = 5.0 Hz, 1H), 3.32 (dd, *J* = 17.6, 5.0 Hz, 1H), 3.20 (dd, *J* = 17.6, 11.0 Hz, 1H), 2.99 (s, 1H), 2.98 (d, *J* = 3.6 Hz, 1H). ^**13**^**C NMR** (125 MHz, CDCl_3_) δ 197.5, 169.6, 161.2, 150.7, 140.8, 129.2, 127.9, 126.7, 118.1, 117.3, 43.8, 41.3, 31.1. **HRMS** (ESI-): m/z [M-H]^-^ calculated for C_15_H_11_O_4-_ 255.0663, found: 255.0642, error: 2.1 ppm. **HPLC purity** 96 %.

(*E*)-4-(naphthalen-2-yl)but-3-en-2-one (**3b**) (610 mg; 3.09 mmol; 33 %), white solid. ^**1**^**H NMR** (500 MHz, CDCl_3_) δ 7.98 (s, 1H), 7.91 – 7.83 (m, 3H), 7.73 – 7.67 (m, 1H), 7.70 (d, *J* = 16.3 Hz, 1H), 7.58 – 7.52 (m, 2H), 6.86 (d, *J* = 16.3 Hz, 1H), 2.45 (s, 3H). ^**13**^**C NMR** (126 MHz, CDCl_3_) δ 198.3, 143.5, 134.3, 133.3, 131.9, 130.3, 128.8, 128.5, 127.8, 127.4, 127.3, 126.8, 123.5, 27.6. 5-(naphthalen-2-yl)cyclohexane-1,3-dione (**3c**) (158 mg; 0.66 mmol; 26 %). ^**1**^**H NMR** (500 MHz, d_6_-DMSO) δ 7.93 – 7.79 (m, 4H), 7.56 (dd, *J* = 8.5, 1.8 Hz, 1H), 7.54 – 7.43 (m, 2H), 5.34 (s, 1H), 3.50 (tt, *J* = 11.7, 4.4 Hz, 1H), 2.92 (dd, *J* = 7.2, 1.4 Hz, 2H), 2.59 (dd, *J* = 15.7, 8.7 Hz, 2H). ^**13**^**C NMR** (126 MHz, d_6_-DMSO) δ 141.6, 133.5, 132.4, 128.4, 128.0, 127.9, 126.6, 126.2, 126.1, 125.4, 104.0, 30.7. 6-(naphthalen-2-yl)-4-oxo-4,5,6,7-tetrahydrobenzofuran-3-carboxylic acid (**3**) (12 mg; 0.04 mmol; 6 %). ^**1**^**H NMR** (500 MHz, CDCl_3_) δ 13.15 (s, 1H), 8.17 (s, 1H), 7.92 (d, *J* = 8.5 Hz, 1H), 7.90 – 7.83 (m, 2H), 7.74 (s, 1H), 7.58 – 7.51 (m, 2H), 7.44 (dd, *J* = 8.5, 1.8 Hz, 1H), 3.92 – 3.82 (m, 1H), 3.41 (dd, *J* = 17.6, 5.2 Hz, 1H), 3.31 (dd, *J* = 17.6, 10.9 Hz, 1H), 3.16 – 3.05 (m, 2H). ^**13**^**C NMR** (126 MHz, CDCl_3_) δ 197.8, 169.6, 161.2, 150.71, 138.1, 133.4, 132.8, 129.1, 127.74, 127.72, 126.7, 126.4, 125.4, 124.5, 118.1, 117.3, 43.8, 41.4, 31.0. **HRMS** (ESI+): m/*z* [M+H]^+^ calculated for C_19_H_15_O_4_ 307.0965; found 307.0970. error: 1.6 ppm. **HPLC purity** 98 %.

(*E*)-4-(3-methoxyphenyl)but-3-en-2-one (**4b**) (2.45 g; 13.91 mmol; 93 %). ^**1**^**H NMR** (500 MHz, CDCl_3_) δ 7.50 (d, *J* = 16.3 Hz, 1H), 7.34 (t, *J* = 7.9 Hz, 1H), 7.16 (d, *J* = 7.6 Hz, 1H), 7.10 – 7.07 (m, 1H), 6.97 (ddd, *J* = 8.3, 2.6, 0.8 Hz, 1H), 6.72 (d, *J* = 16.3 Hz, 1H), 3.86 (s, 3H), 2.41 (s, 3H). ^**13**^**C NMR** (126 MHz, CDCl_3_) δ 198.4, 159.9, 143.3, 135.8, 129.9, 127.4, 121.0, 116.4, 113.0, 55.3, 27.5. 5-(3-methoxyphenyl)cyclohexane-1,3-dione (**4c**) (140 mg; 0.64 mmol; 6 %). ^**1**^**H NMR** (500 MHz, CD_3_CN) δ 7.28 (t, *J* = 7.9 Hz, 1H), 6.94 – 6.80 (m, 3H), 5.39 (s), 3.82 (s, 3H), 3.37 (tt, *J* = 11.8, 4.7 Hz, 1H), 2.63 (dd, *J* = 16.8, 11.9 Hz, 2H), 2.51 (dd, *J* = 17.1, 4.6 Hz, 2H). ^**13**^**C NMR** (126 MHz, CD_3_CN) δ 159.9, 145.3, 129.7, 119.1, 112.8, 111.9, 103.9, 54.8, 39.3.

6-(3-methoxyphenyl)-4-oxo-4,5,6,7-tetrahydrobenzofuran-3-carboxylic acid (**4**) (2.4 mg; 8.17 μmol; 2 %) ^**1**^**H NMR** (500 MHz, CDCl_3_) δ 13.1 (s, 1H), 8.15 (s, 1H), 7.34 (t, *J* = 7.9 Hz, 1H), 6.9 – 6.8 (m, 3H), 3.83 (s, 3H), 3.7 – 3.6 (m, 1H), 3.31 (dd, *J* = 17.6, 5.1 Hz, 1H), 3.19 (dd, *J* = 17.6, 11.1 Hz, 1H), 3.00 – 2.90 (m, 2H). ^**13**^**C NMR** (126 MHz, CDCl_3_) δ 197.8, 169.6, 161.2, 160.1, 150.7, 142.4, 130.3, 118.8, 118.1, 117.2, 113.1, 112.5, 55.3, 43.8, 41.3, 31.0. **HPLC purity** 98 %.

(*E*)-4-(3-(trifluoromethyl)phenyl)but-3-en-2-one (**5b**) (3.48 g; 16.2 mmol; 85 %) ^**1**^**H NMR** (500 MHz, CDCl_3_) δ 7.81 (d, *J* = 0.4 Hz, 1H), 7.74 (d, *J* = 7.8 Hz, 1H), 7.67 (d, *J* = 7.8 Hz, 1H), 7.55 (t, *J* = 7.8 Hz, 1H), 7.54 (d, *J* = 16.4 Hz, 1H), 6.79 (d, *J* = 16.3 Hz, 1H), 2.42 (s, 3H). ^**13**^**C NMR** (126 MHz, CDCl_3_) δ 197.8 (s), 141.3 (s), 135.3 (s), 131.5 (q, *J* = 32.6 Hz), 131.2 (q, *J* = 1.3 Hz), 129.5 (s), 128.5 (s), 126.8 (q, *J* = 3.7 Hz), 124.8 (q, *J* = 3.8 Hz), 124.5 (q, *J* = 273.6 Hz), 27.8 (s). ^**19**^**F NMR** (470 MHz, CDCl_3_) δ −63.9.

5-(3-(trifluoromethyl)phenyl)cyclohexane-1,3-dione (**5c**) (117 mg; 0.456 mmol; 4 %) ^**1**^**H NMR** (500 MHz, CD_3_OD) δ 7.70 – 7.60 (m, 2H), 7.61 – 7.45 (m, 2H), 3.60 – 3.50 (m, 1H), 2.74 (dd, *J* = 17.5, 11.7 Hz, 2H), 2.74 (dd, *J* = 17.5, 4.7 Hz, 2H) Open ring impurities are present (20%). ^**13**^**C NMR** (126 MHz, CD_3_OD) δ 144.6 (s), 130.5 (d, J = 33 Hz), 130.5 (s), 129.2 (s), 124.2 (q, *J* = 274 Hz), 123.4 (q, *J* = 4.1 Hz), 39.2 (s), 38.0 (s). ^**19**^**F NMR** (470 MHz, CDCl_3_) δ −65.0 (-C**F**_**3**_). **HRMS** (ESI+): m/*z* [M+H]^+^ calculated for C_13_H_12_F_3_O_2_ 257.0784; found 257.0782. error: – 0.7 ppm. 4-oxo-6-(3-(trifluoromethyl)phenyl)-4,5,6,7-tetrahydrobenzofuran-3-carboxylic acid (**5**) (1.7 mg; 5.25 μmol; 1 %). ^**1**^**H NMR** (500 MHz, CDCl_3_) δ 13.0 (s, 1H), 8.17 (s, 1H), 7.64 (d, *J* = 7.7 Hz, 1H), 7.61 – 7.47 (m, 3H), 3.77 – 3.71 (m, 1H), 3.36 (dd, *J* = 17.5, 5.1 Hz, 1H), 3.23 (dd, *J* = 17.5, 11.2 Hz, 1H), 3.05 – 2.93 (m, 2H). ^**13**^**C NMR** (126 MHz, CDCl_3_) δ 197.0 (s), 169.0 (s), 161.0 (s), 150.9 (s), 141.7 (s), 131.5 (q, *J* = 32.7 Hz), 130.1 (s), 129.8 (s), 124.9 (q, *J* = 3.5 Hz) 123.5 (q, *J* = 3.5 Hz), 124.0 (q, *J* = 274 Hz), 118.0 (s), 117.3 (s), 43.5 (s), 41.0 (s), 30.9 (s). ^**19**^**F NMR** (470 MHz, CDCl_3_) δ −63.7. **HRMS** (ESI+): m/*z* [M+H]^+^ calculated for C_15_H_12_F_3_O_4_ 325.0682; found 325.0686. error: 1.2 ppm. **HPLC purity** 96 %.

(E)-4-(benzofuran-2-yl)but-3-en-2-one (**6b**) (3.10 g, 16.7 mmol, 81 %) ^**1**^**H NMR** (400 MHz, CDCl_3_) δ 7.57 (d, J = 7.5 Hz, 1H), 7.47 (dd, J = 7.5, 1.0 Hz, 1H), 7.38 (d, J = 15.7 Hz, 1H), 7.36 (td, J = 7.5, 1.0 Hz, 1H), 7.24 (td, J = 7.5, 1.0 Hz, 1H), 6.97 (s, 1H), 6.86 (d, J = 15.7 Hz, 1H), 2.37 (s, 3H). ^**13**^**C NMR** (101 MHz, CDCl_3_) δ 197.5, 155.6, 152.4, 129.5, 128.4, 126.7, 126.7, 123.4, 121.8, 112.0, 111.4, 28.3.

5-(benzofuran-2-yl)cyclohexane-1,3-dione (**6c**) (338 mg, 1.87 mmol, 50 %) ^**1**^**H NMR** (400 MHz, CD_3_OD) δ 7.50 (d, J = 7.5 Hz, 1H), 7.41 (d, J = 8.0 Hz, 1H), 7.22 (td, J = 8.0, 1.4 Hz, 1H), 7.17 (td, J = 7.5, 1.2, 1H), 6.55 (t, J = 0.80 Hz, 1H), 5.42 (s, weak), 3.64 (m, J = 5.0 Hz, 1H), 2.80 (dd, J = 17.0, 5.2 Hz, 2H), 2.73 (dd, J = 17.0, 3.9 Hz, 2H). ^**13**^**C NMR** (101 MHz, CD_3_OD) δ 160.5, 156.1, 129.8, 124.9, 123.8, 121.8, 111.7, 102.9, 38.6, 34.5. Open form impurity 30 %. HRMS (ESI+): m/z [M+H]+ calculated for C_14_H_13_O_3_^+^: 229.0859, found: 229.0865, error: 0.4 ppm

4’-oxo-4’,5’,6’,7’-tetrahydro-[2,6’-bibenzofuran]-3’-carboxylic acid (**6**) (24 mg, 81 µmol, 8.7 %). ^**1**^**H NMR** (500 MHz, CD_3_OD) δ 8.11 (s, 1H), 7.52 (dd, *J* = 7.7, 0.8 Hz, 1H), 7.43 (dd, *J* = 8.0, 0.8 Hz, 1H), 7.25 (td, *J* = 8.0, 1.5 Hz, 1H), 7.19 (td, *J* = 7.7, 0.8 Hz, 1H), 6.65 (t, *J* = 1.0 Hz, 1H), 3.97 (m, 1H), 3.48 (dd, *J* = 17.3, 5.3 Hz, 1H), 3.35 (dd, *J* = 17.6, 9.3 Hz, 1H), 3.07 (s, 1H), 3.05 (d, *J* = 4.8 Hz, 1H). ^**13**^**C NMR** (125 MHz, CD_3_OD) δ 128.3, 123.9, 122.5, 120.6, 110.5, 102.2, 40.9, 34.4, 27.4. **HRMS** (ESI+): m/z [M+H]^+^ calculated mass for C_17_H_13_O_5_^+^: 297.0757, found: 297.07423, error: 1.4 ppm. **HPLC purity** 97 %.

(*E*)-4-(2-bromophenyl)but-3-en-2-one (**7b**) (6.8 g; 30.2 mmol; 78 %). ^**1**^**H NMR** (500 MHz, CDCl_3_) δ 7.91 (d, *J* = 16.3 Hz, 1H), 7.69 – 7.61 (m, 2H), 7.40 – 7.33 (m, 1H), 7.26 (dd, *J* = 7.8 Hz, 1.7 Hz, 1H), 6.64 (d, *J* = 16.3 Hz, 1H), 2.45 (s, 3H). ^**13**^**C NMR** (126 MHz, CDCl_3_) δ 198.3, 141.9, 134.4, 133.4, 131.4, 129.8, 127.8, 127.7, 125.6, 27.2. 5-(2-bromophenyl)cyclohexane-1,3-dione (**7c**) (387 mg; 1.448 mmol; 5 %). ^**1**^**H NMR** (500 MHz, CD_3_OD) δ 7.63 (dd, *J* = 8.0, 1.2 Hz, 1H), 7.46 (dd, *J* = 7.8, 1.6 Hz, 1H), 7.39 (td, *J* = 7.7, 1.2 Hz, 1H), 7.19 (td, *J* = 7.8, 1.7 Hz, 1H), 3.84 (tt, *J* = 11.5, 4.6 Hz, 1H), 2.69 (dd, *J* = 17.1, 11.5 Hz, 2H), 2.60 (dd, *J* = 17.1, 4.7 Hz, 2H). ^**13**^**C NMR** (126 MHz, CD_3_OD) δ 141.4, 133.0, 128.4, 127.8, 127.3, 123.7, 38.5, 37.9. **HRMS** (ESI+): m/*z* [M+H]^+^ calculated for C_12_H_12_^79^BrO_2_ 267.0015; found 267.0018. error: 1.0 ppm.

6-(2-bromophenyl)-4-oxo-4,5,6,7-tetrahydrobenzofuran-3-carboxylic acid (**7**) (143 mg; 0.426 mmol; 30 %) ^**1**^**H NMR** (500 MHz, CDCl_3_) δ 13.10 (s, 1H), 8.16 (s, 1H), 7.66 (dd, *J* = 8.0, 1.2 Hz, 1H), 7.39 (td, *J* = 7.6, 1.2 Hz, 1H), 7.34 (dd, *J* = 7.8, 1.7 Hz, 1H), 7.22 (ddd, *J* = 8.0, 7.4, 1.7 Hz, 1H), 4.22 – 4.13 (m, 1H), 3.39 (dd, *J* = 17.6, 5.1 Hz, 1H), 3.15 (dd, *J* = 17.6, 10.8 Hz, 1H), 3.02 – 2.97 (m, 2H). ^**13**^**C NMR** (126 MHz, CDCl_3_) δ 197.5, 169.4, 161.1, 150.8, 139.5, 133.8, 129.4, 128.3, 127.1, 124.2, 118.1, 117.2, 42.4, 40.0, 29.6. **HRMS** (ESI+): m/*z* [M+H]^+^ calculated for C_15_H_12_^79^BrO_4_ 334.9913; found 334.9926. error: – 3.8 ppm. **HPLC purity** 97 %.

(*E*)-4-(2-fluorophenyl)but-3-en-2-one (**8b**) (2.34 g; 14.26 mmol; 74 %). ^**1**^**H NMR** (500 MHz, CDCl_3_) δ 7.69 (d, *J* = 16.5 Hz, 1H), 7.59 (td, *J* = 7.6, 1.7 Hz, 1H), 7.39 (dddd, *J* = 8.3, 7.2, 5.3, 1.7 Hz, 1H), 7.19 (td, *J* = 7.6, 0.9 Hz, 1H), 7.13 (ddd, *J* = 10.6, 8.3, 1.1 Hz, 1H), 6.80 (d, *J* = 16.5 Hz, 1H), 2.42 (s, 3H). ^**13**^**C NMR** (126 MHz, CDCl_3_) δ 198.4 (s), 161.4 (d, *J* = 254 Hz), 135.7 (d, *J* = 3.5 Hz), 131.9 (d, *J* = 8.8 Hz), 129.2 (d, *J* = 5.4 Hz), 128.7 (d, *J* = 2.8 Hz), 124.5 (d, *J* = 3.7 Hz), 122.5 (d, *J* = 11.6 Hz), 116.2 (d, *J* = 21.8 Hz), 27.5 (s). ^**19**^**F NMR** (376 MHz, CDCl_3_) δ −115.8.

5-(2-fluorophenyl)cyclohexane-1,3-dione (**8c**) (350 mg; 1.7 mmol; 12 %). ^**1**^**H NMR** (500 MHz, CD_3_OD) δ 7.39 (td, *J* = 7.7, 1.5 Hz, 1H), 7.30 (tdd, *J* = 7.3, 5.3, 1.7 Hz, 1H), 7.18 (td, *J* = 7.6, 0.8 Hz, 1H), 7.15 – 7.08 (m, 1H), 3.68 (tt, *J* = 11.7, 4.6 Hz, 1H), 2.74 (dd, *J* = 17.1, 11.7 Hz, 2H), 2.57 (dd, *J* = 17.2, 4.6 Hz, 2H). ^**13**^**C NMR** (126 MHz, CD_3_OD) δ 160.9 (d, *J* = 247 Hz), 129.5 (d, *J* = 13.6 Hz), 128.4 (d, *J* = 8.6 Hz), 127.8 (d, *J* = 4.5 Hz), 124.3 (d, *J* = 3.5 Hz), 115.2 (d, *J* = 23 Hz), 37.7 (s), 33.1 (d, *J* = 2.5 Hz). ^**19**^**F NMR** (376 MHz, CDCl_3_) δ −120.2. **HRMS** (ESI+): m/*z* [M+H]^+^ calculated for C_12_H_12_FO_2_ 207.0816; found 207.0817. error: 0.7 ppm.

6-(2-fluorophenyl)-4-oxo-4,5,6,7-tetrahydrobenzofuran-3-carboxylic acid (**8**) (250 mg; 0.912 mmol; 53 %). ^**1**^**H NMR** (500 MHz, CDCl_3_) δ 13.12 (s, 1H), 8.15 (s, 1H), 7.39 – 7.31 (m, 1H), 7.31 – 7.25 (m, 1H), 7.20 (td, *J* = 7.6, 1.1 Hz, 1H), 7.14 (ddd, *J* = 10.8, 8.2, 1.1 Hz, 1H), 3.99 – 3.88 (m, 1H), 3.32 (dd, *J* = 16.8, 5.2 Hz, 1H), 3.27 (dd, *J* = 16.8, 9.3 Hz, 1H), 3.08 (dd, *J* = 17.2, 12.3 Hz, 1H), 2.96 (dd, *J* = 17.2, 4.2 Hz, 1H). ^**13**^**C NMR** (126 MHz, CDCl_3_) δ 197.8 (s), 169.6 (s), 161.2 (s), 160.8 (d, *J* = 246 Hz), 150.7 (s), 129.6 (d, *J* = 8.6 Hz), 128.0 (d, *J* = 4.3 Hz), 127.6 (d, *J* = 13.4 Hz), 124.8 (d, *J* = 3.6 Hz), 118.1 (s), 117.2 (s), 116.3 (d, *J* = 22.1 Hz), 42.1 (d, *J* = 1.8 Hz), 35.6 (d, *J* = 1.9 Hz), 29.4 (d, *J* = 2.2 Hz). ^**19**^**F NMR** (376 MHz, CDCl_3_) δ −116.4. **HRMS** (ESI+): m/*z* [M+H]^+^ calculated for C_15_H_12_FO_4_ 275.0714; found 275.0719. error: 1.8 ppm. **HPLC purity** >99 %.

(*E*)-4-(2-phenoxyphenyl)but-3-en-2-one (**9b**) (3.56 g; 14.95 mmol; 78 %). ^**1**^**H NMR** (500 MHz, CDCl_3_) δ 7.90 (d, *J* = 16.5 Hz, 1H), 7.69 (dd, *J* = 7.8, 1.6 Hz, 1H), 7.37 – 7.32 (m, 3H), 7.20 – 7.12 (m, 2H), 7.06 – 7.01 (m, 2H), 6.91 (dd, *J* = 8.3, 1.0 Hz, 1H), 6.80 (d, *J* = 16.5 Hz, 1H), 2.37 (s, 3H). ^**13**^**C NMR** (126 MHz, CDCl_3_) δ 198.9, 156.8, 156.1, 137.9, 131.7, 130.0, 128.5, 128.2, 125.9, 123.8, 123.7, 118.97, 118.95, 27.1.

5-(2-phenoxyphenyl)cyclohexane-1,3-dione (**9c**) (1.41 g; 4.04 mmol; 27 %). ^**1**^**H NMR** (500 MHz, CD_3_OD) δ 7.44 (dd, *J* = 7.7, 1.6 Hz, 1H), 7.39 – 7.32 (m, 2H), 7.28 – 7.22 (m, 1H), 7.17 (td, *J* = 7.5, 1.2 Hz, 1H), 7.14 – 7.07 (m, 1H), 6.98 – 6.92 (m, 2H), 6.88 (dd, *J* = 8.1, 1.2 Hz, 1H), 3.78 – 3.65 (m, 1H), 2.76 (dd, *J* = 17.2, 11.7 Hz, 2H), 2.55 (dd, *J* = 17.3, 4.5 Hz, 2H). ^**13**^**C NMR** (126 MHz, CD_3_OD) δ 157.6, 154.4, 133.7, 129.6, 128.0, 127.6, 123.9, 122.8, 119.2, 117.6, 33.6. Two quaternary carbon signals are missing or overlapping. **HRMS** (ESI+): m/*z* [M+H]^+^ calculated for C_18_H_17_O_3_ 281.1172; found 281.1170. error: – 0.7 ppm.

4-oxo-6-(2-phenoxyphenyl)-4,5,6,7-tetrahydrobenzofuran-3-carboxylic acid (**9**) (250 mg; 0.717 mmol; 17 %). ^**1**^**H NMR** (400 MHz, CDCl_3_) δ 13.19 (s, 1H), 8.12 (s, 1H), 7.43 – 7.25 (m, 4H), 7.16 (t, *J* = 7.4 Hz, 2H), 7.00 (d, *J* = 7.8 Hz, 2H), 6.95 – 6.87 (m, 1H), 4.25 – 3.88 (m, 1H), 3.42 – 3.23 (m, 2H), 3.16 (dd, *J* = 17.2, 12.3 Hz, 1H), 2.97 (dd, *J* = 17.2, 4.1 Hz, 1H). ^**13**^**C NMR** (101 MHz, CDCl_3_) δ 198.3, 170.1, 161.3, 156.6, 155.0, 150.5, 131.3, 130.1, 129.1, 127.8, 123.9, 123.8, 119.0, 118.7, 118.0, 117.1, 42.3, 36.3, 29.5. **HRMS** (ESI+): m/*z* [M+H]^+^ calculated for C_21_H_17_O_5_ 349.1071; found 349.1085. error: 4.3 ppm. **HPLC purity** 91 %.

2-isopropoxybenzaldehyde (**10a**) (7.87 g; 48 mmol; 80 %). ^**1**^**H NMR** (500 MHz, CDCl_3_) δ 10.51 (d, *J* = 0.8 Hz, 1H), 7.84 (dd, *J* = 7.9, 1.9 Hz, 1H), 7.53 (ddd, *J* = 8.5, 7.3, 1.9 Hz, 1H), 7.03 – 6.99 (m, 2H), 4.70 (septet, *J* = Hz, 1H), 1.42 (d, *J* = 6.1 Hz, 6H). ^**13**^**C NMR** (126 MHz, CDCl_3_) δ 190.2, 160.6, 135.7, 128.3, 125.7, 120.4, 114.0, 71.1, 22.0.

(E)-4-(2-isopropoxyphenyl)but-3-en-2-one (**10b**) (6.2 g; 30.06 mmol; 78 %). ^**1**^**H NMR** (500 MHz, CDCl_3_) δ 7.92 (d, *J* = 16.5 Hz, 1H), 7.57 (dd, *J* = 7.7, 1.7 Hz, 1H), 7.35 (ddd, *J* = 8.4, 7.4, 1.7 Hz, 1H), 6.96 (t, *J* = 7.6 Hz, 1H), 6.95 (d, *J* = 8.4 Hz, 1H), 6.77 (d, *J* = 16.5 Hz, 1H), 4.64 (septet, *J* = 6.1 Hz, 1H), 2.40 (s, 3H), 1.42 (d, *J* = 6.1 Hz, 6H). ^**13**^**C NMR** (126 MHz, CDCl_3_) δ 199.1, 156.8, 139.1, 131.6, 128.4, 127.5, 124.3, 120.6, 113.8, 71.0, 27.1, 22.1.

5-(2-isopropoxyphenyl)cyclohexane-1,3-dione (**10c**) (1.18 g; 4.81 mmol; 16 %). ^**1**^**H NMR** (500 MHz, CD_3_OD) δ 7.25 – 7.19 (m, 2H), 6.98 (d, *J* = 8.1 Hz, 1H), 6.91 (td, *J* = 7.5, 1.0 Hz, 1H), 4.68 (septet. *J* = 5.9 Hz, 1H), 3.70 (tt, *J* = 11.7, 4.5 Hz, 1H), 2.73 (dd, *J* = 17.2, 11.7 Hz, 2H), 2.53 (dd, *J* = 17.3, 4.5 Hz, 2H), 1.36 (d, *J* = 6.0 Hz, 6H). ^**13**^**C NMR** (126 MHz, CD_3_OD) δ 155.2, 131.2, 127.6, 126.9, 120.1, 112.7, 69.5, 47.6, 33.9, 21.0. **HRMS** (ESI+): m/*z* [M+H]^+^ calculated for C_15_H_19_O_3_ 247.1329; found 247.1324. error: – 1.8 ppm. 6-(2-isopropoxyphenyl)-4-oxo-4,5,6,7-tetrahydrobenzofuran-3carboxylic acid (**10**) (300 mg; 0.954 mmol; 20 %). ^**1**^**H NMR** (500 MHz, CDCl_3_) δ 13.30 (s, 1H), 8.13 (s, 1H), 7.32 – 7.27 (m, 1H), 7.19 (dd, *J* = 7.8, 1.6 Hz, 1H), 6.95 (m, 2H), 4.75 – 4.57 (m, 1H), 4.01 – 3.88 (m, 1H), 3.32 (dd, *J* = 17.6, 10.5 Hz, 1H), 3.24 (dd, *J* = 17.6, 5.5 Hz, 1H), 3.14 (dd, *J* = 17.2, 12.3 Hz, 1H), 2.90 (dd, *J* = 17.2, 4.0 Hz, 1H), 1.39 (dd, *J* = 6.0, 2.6 Hz, 6H). ^**13**^**C NMR** (126 MHz, CDCl_3_) δ 199.1, 170.6, 161.4, 155.3, 150.4, 129.2, 128.7, 127.6, 120.5, 118.1, 117.0, 112.7, 69.8, 42.1, 36.6, 29.0, 22.2. **HRMS** (ESI+): m/*z* [M+H]^+^calculated for C_18_H_19_O_5_ 315.1227; found 315.1234. error: 2.2 ppm. **HPLC purity** 97 %.

(*E*)-4-(2-methoxyphenyl)but-3-en-2-one (**11b**) (2.19 g; 12.44 mmol; 65 %). ^**1**^**H NMR** (500 MHz, CDCl_3_) δ 7.91 (d, *J* = 16.5 Hz, 1H), 7.57 (dd, *J* = 7.7, 1.6 Hz, 1H), 7.42 – 7.36 (m, 1H), 7.00 (t, *J* = 7.5 Hz, 1H), 6.95 (d, *J* = 8.3 Hz, 1H), 6.78 (d, *J* = 16.5 Hz, 1H), 3.92 (s, 3H), 2.41 (s, 3H). ^**13**^**C NMR** (126 MHz, CDCl_3_) δ 199.1, 158.2, 138.7, 131.8, 128.3, 127.8, 123.3, 120.8, 111.1, 55.5, 27.1. 5-(2-methoxyphenyl)cyclohexane-1,3-dione (**11c**) (164 mg; 0.751 mmol; 7 %). ^**1**^**H NMR** (500 MHz, CD_3_OD) δ 7.29 – 7.21 (m, 2H), 6.99 (dd, *J* = 8.1, 0.6 Hz, 1H), 6.94 (td, *J* = 7.5, 1.0 Hz, 1H), 3.87 (s, 3H), 3.74 – 3.66 (m, 1H), 2.72 (dd, *J* = 17.2, 11.7 Hz, 2H), 2.53 (dd, *J* = 17.3, 4.5 Hz, 2H). ^**13**^**C NMR** (126 MHz, CD_3_OD) δ 157.2, 130.4, 127.7, 126.6, 120.4, 110.4, 54.4, 37.7, 33.7. 6-(2-methoxyphenyl)-4-oxo-4,5,6,7-tetrahydrobenzofuran-3-carboxylic acid (**11**) (108 mg; 0.375 mmol; 50 %). ^**1**^**H NMR** (500 MHz, CDCl_3_) δ 13.3 (s, 1H), 8.13 (s, 1H), 7.36 – 7.30 (m, 1H), 7.20 (dd, *J* = 7.6, 1.6 Hz, 1H), 7.00 (td, *J* = 7.5, 1.0 Hz, 1H), 6.96 (d, *J* = 8.3 Hz, 1H), 4.02 – 3.92 (m, 1H), 3.30 (dd, *J* = 17.6, 10.3 Hz, 1H), 3.24 (dd, *J* = 17.6, 5.7 Hz, 1H), 3.12 (dd, *J* = 17.2, 12.4 Hz, 1H), 2.90 (dd, *J* = 17.2, 3.8 Hz, 1H). ^**13**^**C NMR** (126 MHz, CDCl_3_) δ 198.9, 170.6, 161.4, 157.0, 150.4, 128.9, 128.7, 127.3, 120.9, 118.0, 117.0, 111.0, 55.3, 42.0, 36.3, 29.0. **HRMS** (ESI+): m/*z* [M+H]^+^ calculated for C_16_H_15_O_5_ 287.0914; found 287.0912. error: – 0.7 ppm. **HPLC purity** 97 %.

(*E*)-4-(pyridin-2-yl)but-3-en-2-one (**12b**) (2.4 g; 16.32 mmol; 85 %). ^**1**^**H NMR** (500 MHz, CDCl_3_) δ 8.67 (dd, *J* = 4.7, 0.7 Hz, 1H), 7.74 (td, *J* = 7.7, 1.8 Hz, 1H), 7.54 (d, *J* = 16.0 Hz, 1H), 7.50 (d, *J* = 7.8 Hz, 1H), 7.32 – 7.27 (m, 1H), 7.16 (d, *J* = 16.0 Hz, 1H), 2.42 (s, 3H). ^**13**^**C NMR** (126 MHz, CDCl_3_) δ 198.5, 153.1, 150.2, 141.9, 136.8, 130.2, 124.3, 124.2, 28.1.

5-(pyridin-2-yl)cyclohexane-1,3-dione (**12c**) (87 mg; 0.46 mmol; 3 %). ^**1**^**H NMR** (500 MHz, CD_3_OD) δ 8.54 (ddd, *J* = 4.9, 1.6, 0.8 Hz, 1H), 7.82 (td, *J* = 7.7, 1.8 Hz, 1H), 7.42 (d, *J* = 7.9 Hz, 1H), 7.32 (ddd, *J* = 7.6, 4.9, 1.1 Hz, 1H), 3.57 (ddd, *J* = 11.5, 8.1, 4.7 Hz, 1H), 2.83 (dd, *J* = 17.3, 11.6 Hz, 2H), 2.62 (dd, *J* = 17.4, 4.7 Hz, 2H). ^**13**^**C NMR** (126 MHz, CD_3_OD) δ 161.4, 148.7, 137.5, 122.3, 122.0, 41.1, 37.6. **HRMS** (ESI+): m/*z* [M+H]^+^ calculated for C_11_H_12_NO_2_ 190.0863; found 190.0861. error: – 0.8 ppm.

4-oxo-6-(pyridin-2-yl)-4,5,6,7-tetrahydrobenzofuran-3-carboxylic acid (**12**) (30 mg; 0.116 mmol; 25 %). ^**1**^**H NMR** (500 MHz, CDCl_3_) δ 13.22 (s, 1H), 8.74 – 8.39 (m, 1H), 8.12 (s, 1H), 7.72 (td, *J* = 7.7, 1.8 Hz, 1H), 7.27 – 7.23 (m, 2H), 3.91 – 3.80 (m, 1H), 3.50 (dd, *J* = 17.7, 9.7 Hz, 1H), 3.32 (dd, *J* = 17.7, 5.3 Hz, 1H), 3.20 (dd, *J* = 17.3, 11.1 Hz, 1H), 2.96 (dd, *J* = 17.3, 4.2 Hz, 1H). ^**13**^**C NMR** (126 MHz, CDCl_3_) δ 197.9, 169.4, 161.3, 159.4, 150.6, 149.6, 137.1, 122.7, 121.9, 118.0, 117.1, 42.7, 42.3, 29.3. **HRMS** (ESI+): m/*z* [M+H]^+^calculated for C_14_H_12_NO_4_ 258.0761; found 258.0764. error: 1.8 ppm. **HPLC purity** >99 %.

2-(3-carboxy-4-oxo-4,5,6,7-tetrahydrobenzofuran-6-yl)pyridine 1-oxide (**13**)(0.89 mg; 3.25 μmol; 12 %). To a solution of **12** (6.8 mg; 0.0264 mmol) in CH_2_Cl_2_ (0.5 mL) was added mCPBA (6.8 mg; 0.04 mmol) and the mixture stirred at rt for 5h. The solvent was subsequently removed under a stream of nitrogen. The resulting mixture was purified by reversed-phase preparative HPLC (0.1 % TFA H_2_O / MeCN 2:3 – 1:9) to yield **13** as a white powder. ^**1**^**H NMR** (500 MHz, CDCl_3_) δ 8.51 (dd, *J* = 6.5, 0.9 Hz, 1H), 8.15 (s, 1H), 7.60 (td, *J* = 7.8, 1.2 Hz, 1H), 7.49 – 7.44 (m, 1H), 7.39 (dd, *J* = 7.9, 1.9 Hz, 1H), 4.57 – 4.33 (m, 1H), 3.52 (dd, *J* = 17.7, 5.6 Hz, 1H), 3.48 (dd, *J* = 17.7, 9.0 Hz, 1H), 3.27 (dd, *J* = 17.1, 10.5 Hz, 1H), 3.08 (dd, *J* = 17.1, 4.3 Hz, 1H). ^**13**^**C NMR** (126 MHz, CDCl_3_) δ 196.7, 168.7, 161.0, 151.0, 140.9, 130.4, 125.5, 124.8, 118.0, 117.1, 39.0, 36.1, 26.1. **HRMS** (ESI+): m/*z* [M+H]^+^ calculated for C_14_H_12_NO_5_ 274.0710; found 274.0720. error: 3.5 ppm. **HPLC purity** >99 %.

6-(2-hydroxyphenyl)-4-oxo-4,5,6,7-tetrahydrobenzofuran-3-carboxylic acid (**14**) (2.63 mg; 9.66 μmol; 12 %). To a solution of **11** (24.8 mg; 0.0866 mmol) in CH_2_Cl_2_ at – 78 °C was added a 1 M solution of BBr_3_ in CH_2_Cl_2_ (2.28 eq., 0.197 mmol; 50 mg; 0.2 mL) dropwise and the reaction left to warm to rt and stirred for 24 h. The solvent was removed under a stream of nitrogen and the resulting mixture was purified by reversed-phase preparative HPLC (0.1 % TFA H_2_O / MeCN 2:3 – 1:9) to yield **14** as a white powder. ^**1**^**H NMR** (500 MHz, CDCl_3_) δ 8.15 (s, 1H), 7.25 – 7.16 (m, 2H), 6.98 (td, *J* = 7.5, 1.1 Hz, 1H), 6.83 (dd, *J* = 8.0, 1.0 Hz, 1H), 4.03 – 3.88 (m, 1H), 3.38 (dd, *J* = 17.6, 10.7 Hz, 1H), 3.29 (dd, *J* = 17.6, 5.3 Hz, 1H), 3.20 (dd, *J* = 17.2, 12.4 Hz, 1H), 2.94 (dd, *J* = 17.2, 4.0 Hz, 1H). ^**13**^**C NMR** (126 MHz, CDCl_3_) δ 198.9, 170.6, 161.7, 153.3, 150.5, 128.8, 127.9, 127.0, 121.3, 117.9, 117.0, 115.9, 41.8, 36.7, 28.8. **HRMS** (ESI+): m/*z* [M+H]^+^ calculated for C_15_H_13_O_5_ 273.0757; found 273.0750. error: – 2.8 ppm. **HPLC purity** >99 %.

6-([1,1’-biphenyl]-2-yl)-4-oxo-4,5,6,7-tetrahydrobenzofuran-3-carboxylic acid (**15**) (6.9 mg; 0.021 mmol; 35 %). **7** (20 mg; 0.06 mmol), phenyl boronic acid (1.2 eq.; 8.8 mg; 0.072 mmol), Pd(OAc)_2_ (1 mg; 4.45 μmol), s-phos (2.5 mg; 6.0 μmol) and K_3_PO_4_ (26 mg; 0.12 mmol) were added in toluene (120 μl) and H_2_O (18 μL) and heated in the microwave at 60 °C for 18h. The resulting mixture was filtered through celite, washed with Et_2_O (3 x 20 ml) and the solution concentrated *in vacuo*. The resulting mixture was purified by reversed-phase preparative HPLC (0.1 % TFA H_2_O / MeCN 2:3 – 1:9) to yield **15** as a white powder. ^**1**^**H NMR** (500 MHz, CDCl_3_) δ 8.09 (s, 1H), 7.51 – 7.35 (m, 6H), 7.32 – 7.25 (m, 3H), 3.88 – 3.75 (m, 1H), 3.14 (dd, *J* = 17.8, 8.3 Hz, 1H), 3.06 (dd, *J* = 17.8, 5.4 Hz, 1H), 2.98 (dd, *J* = 17.3, 12.9 Hz, 1H), 2.83 (dd, *J* = 17.3, 3.7 Hz, 1H). ^**13**^**C NMR** (126 MHz, CDCl_3_) δ 197.7, 169.5, 161.3, 150.6, 142.0, 140.5, 138.2, 131.0, 128.9, 128.6, 128.16, 127.6, 127.4, 125.7, 117.9, 117.0, 43.8, 37.0, 31.1. **HRMS** (ESI+): m/*z* [M+H]^+^ calculated for C_21_H_17_O_4_ 333.1121; found 333.1131. error: 3.0 ppm. **HPLC purity** 97 %.

2-bromo-6-methoxybenzaldehyde (**16a**) (1.21 g, 5.6 mmol, 75 %). A mixture of 6-bromor-2-hydroxybenzaldehyde (1.5 g, 7.5 mmol, 1 equiv.), potassium carbonate (3.95 g, 28.6 mmol, 3.8 equiv.), methyl iodide (7.9 ml, 126 mmol, 17 equiv.) and 40 ml acetone was stirred for 96h, diluted with H_2_O (80 ml) and extracted with ether. The organic extract was dried with anhydrous MgSO_4_, evaporated and purified by silica gel chromatography (0 - 60 % EtOAc/pet. ether) to yield crystalline 2-bromo-6-methoxybenzaldehyde. *R*_f_ = 0.6 (1:1 pet. Ether/EtOAc). ^**1**^**H NMR** (500 MHz, CDCl_3_) δ 10.4 (s, 1H), 7.32 (t, *J* = 8.2 Hz, 1H), 7.24 (dd, *J* = 8.0, 1.0 Hz, 1H), 6.95 (dd, *J* = 8.4, 1.0 Hz, 1H), 3.91 (s, 3H). ^**13**^**C NMR** (126 MHz, CDCl_3_) δ 190.6, 162.1, 134.9, 126.6, 125.1, 123.6, 111.2, 56.4. (*E*)-4-(2-bromo-6-methoxyphenyl)but-3-en-2-one (**16b**) (814 mg; 3.189 mmol; 68 %). ^**1**^**H NMR** (500 MHz, CDCl_3_) δ 7.82 (d, *J* = 16.5 Hz, 1H), 7.28 (dd, *J* = 7.8, 0.9 Hz, 1H), 7.18 (t, *J* = 8.2 Hz, 1H), 7.08 (d, *J* = 16.5 Hz, 1H), 6.92 (d, *J* = 8.4 Hz, 1H), 3.91 (s, 3H), 2.43 (s, 3H). ^**13**^**C NMR** (126 MHz, CDCl_3_) δ 199.7, 159.7, 139.0, 133.2, 131.0, 127.1, 125.7, 123.2, 110.3, 55.9, 27.0. **HRMS** (ESI+): m/*z* [M+H]^+^ calculated for C_11_H_11_^79^BrO_2_ 255.0015; found 255.0017. error: 0.5 ppm.

5-(2-bromo-6-methoxyphenyl)cyclohexane-1,3-dione (**16c**) (100 mg; 0.336 mmol; 11 %). ^**1**^**H NMR** (500 MHz, d_6_-DMSO) δ 7.21 (dd, *J* = 8.1, 1.1 Hz, 1H), 7.18 (t, *J* = 8.2 Hz, 1H), 7.07 (dd, *J* = 8.3, 0.8 Hz, 1H), 3.90 (m, 1H), 3.84 (s, 3H), 3.09 (t, *J* = 14.6 Hz, 2H), 2.40 (dd, *J* = 17.4, 4.7 Hz, 2H). ^**13**^**C NMR** (126 MHz, d_6_-DMSO): δ 159.2, 129.4, 128.6, 125.1, 111.8, 103.3, 55.9. **HRMS** (ESI+): m/*z* [M+H]^+^ calculated for C_13_H_14_^79^BrO_3_ 297.0121; found 297.0125. error: 1.4 ppm. 6-(2-bromo-6-methoxyphenyl)-4-oxo-4,5,6,7-tetrahydrobenzofuran-3-carboxylic acid (**16**) (35 mg; 0.095 mmol; 28 %). ^**1**^**H NMR** (500 MHz, CDCl_3_) δ 13.41 (s, 1H), 8.15 (s, 1H), 7.27 (dd, *J* = 8.1, 1.0 Hz, 1H), 7.18 (t, *J* = 8.2 Hz, 1H), 6.93 (dd, *J* = 8.3, 0.8 Hz, 1H), 4.37 (m, 1H), 3.90 (s, 3H), 3.79 – 3.51 (m, 2H), 3.03 (dd, *J* = 17.6, 5.4 Hz, 1H), 2.71 (dd, *J* = 17.5, 4.2 Hz, 1H). ^**13**^**C NMR** (126 MHz, CDCl_3_) δ 199.2, 170.7, 161.6, 159.0, 150.4, 129.7, 127.5, 125.9, 125.5, 118.1, 116.9, 110.8, 55.6, 39.5 x 2, 26.4. **HRMS** (ESI+): m/*z* [M+H]^+^ calculated for C_16_H_14_^79^BrO_5_ 365.0019; found 365.0030. error: 3.1 ppm. **HPLC purity** >99 %.

ethyl 6-(2-bromo-6-methoxyphenyl)-4-oxo-4,5,6,7-tetrahydro-benzofuran-3-carboxylate (**16d**) (44 mg, 0.11 mmol, 33 %). ^**1**^**H NMR** (500 MHz, CDCl_3_) δ 7.90 (s, 1H), 7.21 (dd, *J* = 8.1, 1.1 Hz, 1H), 7.10 (t, *J* = 8.2 Hz, 1H), 6.87 (dd, *J* = 8.3, 1.1 Hz, 1H), 4.36 (q, *J* = 7.1 Hz, 2H), 4.25 (m, 1H), 3.84 (s, 3H), 3.63 (t, *J* = 14.8 Hz, 1H), 3.42 (t, *J* = 14.8 Hz, 1H), 2.91 (dd, *J* = 17.1 Hz, 5.3 Hz, 1H), 2.54 (dd, *J* = 16.6 Hz, 4.1 Hz, 1H), 1.38 (t, *J* = 7.1 Hz, 3H). ^**13**^**C NMR** (126 MHz, CDCl_3_) δ 192.0, 168.6, 162.2, 159.3, 148.0, 129.3, 128.7, 125.9, 125.7, 118.4, 117.2, 110.9, 61.1, 55.7, 41.7, 39.7, 26.6, 14.4. **HRMS** (ESI+): m/*z* [M+H]^+^ neutral calculated C_18_H_17_^79^BrO_5_ 392.02594; neutral found 392.0254. error: −1.4 ppm. **HPLC purity** 95 %. (*E*)-4-(2-chloro-6-fluorophenyl)but-3-en-2-one (**17b**) (3.25 g; 16.4 mmol; 85 %). ^**1**^**H NMR** (500 MHz, CDCl_3_) δ 7.75 (d, *J* = 16.6 Hz, 1H), 7.32 – 7.22 (m, 2H), 7.13 – 7.03 (m, 1H), 6.99 (d, *J* = 16.6 Hz, 1H), 2.43 (s, 3H). ^**13**^**C NMR** (126 MHz, CDCl_3_) δ 198.5 (s), 162.0 (d, *J* = 255 Hz), 136.2 (d, *J* = 5.1 Hz), 133.4 (d, *J* = 13.5 Hz), 133.3 (d, *J* = 2.05 Hz), 130.9 (d, *J* = 10.4 Hz), 126.01 (d, *J* = 3.5 Hz), 121.72 (d, *J* = 13.9 Hz), 115.0 (d, *J* = 23.5 Hz), 27.9 (s). ^**19**^**F NMR** (470 MHz, CDCl_3_) δ −107.6.

5-(2-chloro-6-fluorophenyl)cyclohexane-1,3-dione (**17c**) (685 mg; 2.846 mmol; 17 %). ^**1**^**H NMR** (500 MHz, CD_3_OD) δ 7.36 – 7.26 (m, 2H), 7.18 – 7.08 (m, 1H), 4.05 (ttd, *J* = 13.0, 4.5, 1.1 Hz, 1H), 3.02 (dd, *J* = 17.0, 13.1 Hz, 2H), 2.44 (dd, *J* = 17.4, 4.6 Hz, 2H). ^**13**^**C NMR** (126 MHz, CD_3_OD) δ 162.5 (d, *J* = 247 Hz), 134.2 5 (d, *J* = 7.6 Hz), 129.2 5 (d, *J* = 10.4 Hz), 127.1 (d, *J* = 15.1 Hz), 125.8 (d, *J* = 3.2 Hz), 115.1 (d, *J* = 23.8 Hz), 35.5 (s), 33.9 (s). ^**19**^**F NMR** (470 MHz, CD_3_OD) δ −110.8. **HRMS** (ESI+): m/*z* [M+H]^+^ calculated for C_12_H_11_F^35^ClO_2_ 241.0426; found 241.0429. error: 1.3 ppm.

6-(2-chloro-6-fluorophenyl)-4-oxo-4,5,6,7-tetrahydrobenzofuran-3-carboxylic acid (**17**) (102 mg; 0.33 mmol; 12 %). ^**1**^**H NMR** (400 MHz, CDCl_3_) δ 13.17 (s, 1H), 8.16 (s, 1H), 7.41 – 7.18 (m, 2H), 7.15 – 7.04 (m, 1H), 4.32 (ddd, *J* = 16.6, 12.3, 4.2 Hz, 1H), 3.56 (dd, *J* = 17.5, 12.0 Hz, 1H), 3.38 (dd, *J* = 17.2, 13.7 Hz, 1H), 3.17 (dd, *J* = 17.5, 5.2 Hz, 1H), 2.84 (dd, *J* = 17.4, 4.1 Hz, 1H). ^**13**^**C NMR** (101 MHz, CDCl_3_) δ 197.7 (s), 169.64 (s), 161.2 x 2 (s), 150.6 (s), 134.6 (d, *J* = 7.1 Hz), 129.7 (d, *J* = 10.4 Hz), 126.32 (d, *J* = 3.1 Hz), 118.1 (s), 117.1 (s), 115.55 (d, *J* = 25 Hz), 40.3 (d, *J* = 4.4 Hz), 34.9 (s), 27.4 (d, *J* = 4.5 Hz). ^**19**^**F NMR** (376 MHz, CDCl_3_) δ −108.0. **HRMS** (ESI+): m/*z* [M+H]^+^ calculated for C_15_H_11_^35^ClFO_4_ 309.0324; found 309.0326. error: 0.5 ppm. **HPLC purity** 99 %.

6-(2-chloro-6-fluorophenyl)-4-hydroxy-4,5,6,7-tetrahydrobenzo-furan-3-carboxylic acid (**17e**) NaBH_4_ (1.47 eq.) was added portion-wise to a stirred solution of 6-(2-chloro-6-fluorophenyl)-4-oxo-4,5,6,7-tetrahydrobenzofuran-3-carboxylic acid **17** (20 mg, 1 eq.) in EtOH (1 mL) at 0 °C. The reaction was warmed to rt and stirred for 3 h before cooling back to 0 °C. The cooled mixture was quenched with HCl (1 M, ≈ 5 drops), extracted with EtOAc (3 x 2 mL), dried over Na_2_SO_4_ and the organic layer concentrated in vacuo. The crude mixture was purified by reversed-phase column chromatography (H_2_O / MeCN 9:1 – 1:1) to yield **17e** as a white powder (4.4 mg, 14 µmol, 40 %). ^**1**^**H NMR** (500 MHz, CDCl_3_) δ 8.02 (s, 1H), 7.24-7.18 (m, 2H), 7.01 (m, 1H), 5.09 (m, 1H), 3.77 (m, 1H), 3.14 (m, 1H), 2.78 (m, 1H), 2.41 (m, 1H), 2.33 (m, 1H). ^**13**^**C NMR** (126 MHz, CDCl_3_) δ 167.4, 163.3, 161.3, 153.0, 149.0, 134.6, 128.6, 125.9, 120.6, 117.2, 115.3, 64.5, 35.2, 34.2, 27.1. **HPLC purity** 96 %.

(*E*)-4-(2,6-dimethoxyphenyl)but-3-en-2-one (**18b**) (2.88 g; 13.96 mmol; 66 %). ^**1**^**H NMR** (500 MHz, CDCl_3_) δ 8.00 (d, *J* = 16.6 Hz, 1H), 7.30 (t, *J* = 8.0 Hz, 1H), 7.19 (d, *J* = 16.6 Hz, 1H), 6.59 (d, *J* = 8.4 Hz, 2H), 3.91 (s, 6H), 2.40 (s, 3H). ^**13**^**C NMR** (126 MHz, CDCl_3_) δ 200.6, 160.1, 134.8, 131.5, 130.4, 112.2, 103.7, 55.8, 27.0. 5-(2,6-dimethoxyphenyl)cyclohexane-1,3-dione (**18c**) (538 mg; 2.16 mmol; 16 %). ^**1**^**H NMR** (500 MHz, CD_3_OD) δ 7.21 (t, *J* = 8.4 Hz, 1H), 6.66 (d, *J* = 8.4 Hz, 2H), 4.06 (tt, *J* = 13.0, 4.6 Hz, 1H), 3.84 (s, 6H), 3.22 (dd, *J* = 17.4, 13.0 Hz, 2H), 2.18 (dd, *J* = 17.5, 4.6 Hz, 2H). ^**13**^**C NMR** (126 MHz, CD_3_OD) δ 158.6, 128.0, 117.3, 104.1, 54.7, 47.6, 29.3. **HRMS** (ESI+): m/*z* [M+H]^+^ calculated for C_14_H_17_O_4_ 249.1121; found 249.1120. error: – 0.4 ppm.

6-(2,6-dimethoxyphenyl)-4-oxo-4,5,6,7-tetrahydrobenzofuran-3-carboxylic acid (**18**) (61 mg; 0.192 mmol; 10 %). ^**1**^**H NMR** (500 MHz, CDCl_3_) δ 13.52 (s, 1H), 8.11 (s, 1H), 7.31 – 7.24 (m, 1H), 6.61 (m, 2H), 4.43 – 4.30 (m, 1H), 3.85 (s, 6H), 3.74 – 3.64 (dd, *J* = 17.6, 11.7 Hz, 1H), 3.60 – 3.52 (dd, *J* = 17.6, 13.1 Hz, 1H), 2.94 (dd, *J* = 17.6, 5.2 Hz, 1H), 2.63 (dd, *J* = 17.5, 4.1 Hz, 1H). ^**13**^**C NMR** (126 MHz, CDCl_3_) δ 200.3, 171.7, 161.7, 150.1, 128.8, 118.1, 116.8, 116.3, 104.2, 55.6, 40.4, 30.8, 27.1. **HRMS** (ESI+): m/*z* [M+H]^+^ calculated for C_17_H_17_O_6_ 317.1020; found 317.1026. error: 1.9 ppm. **HPLC purity** 95 %.

(E)-4-(2-bromo-6-chlorophenyl)but-3-en-2-one (**19b**) (1.38 g, 5.3 mmol, 83 %). ^**1**^**H NMR** (500 MHz, CDCl_3_) δ 7.58 (dd, *J* = 8.0, 1.0 Hz, 1H), 7.56 (d, *J* = 16.6 Hz, 1H), 7.43 (dd, *J* = 8.0, 1.0 Hz, 1H), 7.15 (t, *J* = 8.0 Hz, 1H), 6.74 (d, *J* = 16.6 Hz, 1H), 2.45 (s, 3H). ^**13**^**C NMR** (126 MHz, CDCl_3_): δ 198.0, 139.1, 135.1, 124.4, 133.9, 131.9, 130.3, 129.5, 124.6, 27.6. **HRMS** (ESI+): m/z [M+H]^+^ calculated for C_10_H_8_^79^Br^35^ClO: 258.95199, found: 258.9528, mass error: 3.1 ppm. 5-(2-bromo-6-chlorophenyl)cyclohexane-1,3-dione (**19c**) (497 mg, 16.5 mmol, 31 %). ^**1**^**H NMR** (500 MHz, CD_3_OD) δ 7.45-7.66 (d, 2H), 7.19 (t, *J* = 8.0 Hz, 1H), 4.41 (m, 1H), 3.45 (dd, *J* = 17.0, 13.5 Hz, 2H), 2.34 (dd, *J* = 17.0, 4.5 Hz, 2H). ^**13**^**C NMR** (126 MHz, CD_3_OD) δ 132.4, 131.2, 129.5, 39.6. **HRMS** (ESI+): m/z [M+H]^+^ calculated for C_12_H_10_^79^Br^35^ClO_2_: 300.96255, found: 300.9630, mass error: 1.4 ppm.

6-(2-bromo-6-chlorophenyl)-4-oxo-4,5,6,7-tetrahydrobenzofuran-3-carboxylic acid (**19**) (81 mg, 220 µmol, 14 %). ^**1**^**H NMR** (500 MHz, CDCl_3_) δ 13.2 (s, 1H), 8.15 (s, 1H), 7.61 (dd, *J* = 8.0, 1.3 Hz, 1H), 7.40 (dd, *J* = 8.1, 1.3 Hz, 1H), 7.15 (t, *J* = 8.1 Hz, 1H), 4.63 (m, 1H), 4.12 - 3.76 (m, 2H), 3.07 (dd, *J* = 17.8, 5.7 Hz, 1H), 2.74 (dd, *J* = 17.8, 4.6 Hz, 1H). ^**13**^**C NMR** (126 MHz, CDCl_3_) δ 197.9, 169.5, 161.2, 150.8, 135.8, 134.7, 132.7, 131.4, 129.9, 126.6, 118.2, 117.1, 40.3, 38.6, 25.7. **HRMS** (ESI+): m/z [M+H]^+^ calculated for C_15_H_10_^79^Br^35^ClO_4_: 368.95238, found: 368.9532, mass error: 2.3 ppm. **HPLC purity** 98 %.

2-bromo-6-butoxybenzaldehyde (**20a**) (1.5 g; 5.81 mmol; 78 %). ^**1**^**H NMR** (500 MHz, CDCl_3_) δ 10.46 (s, 1H), 7.31 (t, *J* = 8.0 Hz, 1H), 7.24 (dd, *J* = 8.0, 0.5 Hz, 1H), 6.96 (dd, *J* = 8.3, 0.7 Hz, 1H), 4.08 (t, *J* = 6.4 Hz, 2H), 1.89 – 1.79 (m, 2H), 1.58 – 1.48 (m, 2H), 1.00 (t, *J* = 7.4 Hz, 3H). ^**13**^**C NMR** (126 MHz, CDCl_3_) δ 190.1, 161.9, 134.6, 126.4, 123.74, 123.66, 111.8, 68.9, 31.0, 19.2, 13.8. **HRMS** (ESI+): m/*z* [M+H]^+^ calculated for C_11_H_14_^79^BrO_2_ 257.0172; found 257.0168. error: – 1.5 ppm.

(*E*)-4-(2-bromo-6-butoxyphenyl)but-3-en-2-one (**20b**) (1.14 g; 3.854 mmol; 66 %). ^**1**^**H NMR** (500 MHz, CDCl_3_) δ 7.83 (d, *J* = 16.4 Hz, 1H), 7.28 (dd, *J* = 8.0, 1.0 Hz), 7.15 (t, *J* = 8.2 Hz), 7.12 (d, *J* = 16.4 Hz), 6.90 (d, *J* = 8.3 Hz, 1H), 4.07 (t, *J* = 6.5 Hz, 2H), 2.41 (s, 3H), 1.86 (tt, *J* = 12.9, 6.5 Hz, 2H), 1.59 – 1.49 (m, 2H), 1.01 (t, *J* = 7.4 Hz, 3H). ^**13**^**C NMR** (126 MHz, CDCl_3_) δ 199.5, 159.3, 139.1, 132.9, 130.9, 127.2, 125.4, 123.1, 111.1, 68.7, 31.0, 27.2, 19.3, 13.7.

5-(2-bromo-6-butoxyphenyl)cyclohexane-1,3-dione (**20c**) (618 mg; 1.82 mmol; 47 %). ^**1**^**H NMR** (500 MHz, CD_3_OD) δ 7.21 (dd, *J* = 8.0, 1.0 Hz, 1H), 7.14 (dd, *J* = 9.5, 6.8 Hz, 1H), 7.03 (d, *J* = 8.2 Hz, 1H), 4.09 (t, *J* = 6.4 Hz, 2H), 3.35 – 3.30 (m, 1H) 3.28 (dd, *J* = 17.4, 4.8 Hz, 2H), 1.86 – 1.78 (m, 2H),), 1.60 – 1.50 (m, 2H), 1.53 (dd, *J* = 15.1, 7.5 Hz, 2H), 1.00 (t, *J* = 7.4 Hz, 3H). ^**13**^**C NMR** (126 MHz, CD_3_OD) δ 158.7, 129.0, 128.4, 125.1, 111.4, 31.0, 19.2, 12.7. Some carbon signals are missing. **HRMS** (ESI+): m/*z* [M+H]^+^ calculated for C_16_H_19_^79^BrO_3_ 339.0590; found 339.0592. error: 0.6 ppm.

6-(2-bromo-6-butoxyphenyl)-4-oxo-4,5,6,7-tetrahydrobenzofuran-3-carboxylic acid (**20**) (26 mg; 63.63 μmol; 4 %). ^**1**^**H NMR** (500 MHz, CDCl_3_) δ 13.35 (s, 1H), 8.15 (s, 1H), 7.25 (dd, *J* = 8.1, 1.0 Hz, 1H), 7.15 (t, *J* = 8.2 Hz, 1H), 6.92 (d, *J* = 7.8 Hz, 1H), 4.37 (m, 1H), 4.07 (t, *J* = 6.7 Hz, 2H), 3.81 – 3.55 (m, 2H), 3.02 (dd, *J* = 17.6, 5.3 Hz, 1H), 2.71 (dd, *J* = 17.5, 4.1 Hz, 1H), 1.86 – 1.76 (m, 2H), 1.51 – 1.39 (m, 2H), 0.97 (t, *J* = 7.4 Hz, 3H). ^**13**^**C NMR** (126 MHz, CDCl_3_) δ 199.2, 170.7, 161.4, 150.4, 129.6, 127.2, 125.6, 118.1, 116.8, 111.3, 68.3, 40.0, 39.7, 31.1, 26.4, 19.3, 13.7. **HRMS** (ESI+): m/*z* [M+H]^+^ calculated for C_19_H_19_^79^BrO_5_ 407.0489; found 407.0488. error: – 0.1 ppm. **HPLC purity** 96 %.

2-methoxy-6-(2-methoxyethoxy)benzaldehyde (**21a**) (1.36 g; 6.48 mmol; 87 %). ^**1**^**H NMR** (500 MHz, CDCl_3_) δ 10.51 (s, 1H), 7.34 (d, *J* = 3.3 Hz, 1H), 7.14 (dd, *J* = 9.0, 3.3 Hz, 1H), 6.98 (d, *J* = 9.1 Hz, 1H), 4.25 – 4.19 (m, 2H), 3.82 (s, 3H), 3.81 – 3.78 (m, 2H), 3.47 (s, 3H). ^**13**^**C NMR** (126 MHz, CDCl_3_) δ 189.6, 156.0, 153.9, 125.5, 123.5, 115.0, 110.1, 70.9, 69.1, 59.3, 55.7. **HRMS** (ESI+): m/*z* [M+Na]^+^ calculated for C_11_H_14_NaO_4_ 233.0784; found 233.0788. error: 1.7 ppm.

(*E*)-4-(2-methoxy-6-(2-methoxyethoxy)phenyl)but-3-en-2-one (**21b**) (1.11 g; 5.27 mmol; 81 %). ^**1**^**H NMR** (500 MHz, CDCl_3_) δ 7.92 (d, *J* = 16.5 Hz, 1H), 7.08 (d, *J* = 2.5 Hz, 1H), 6.95 – 6.89 (m, 2H), 6.75 (d, *J* = 16.5 Hz, 1H), 4.18 – 4.13 (m, 2H), 3.81 (s, 3H), 3.80 – 3.77 (m, 2H), 3.48 (s, 3H), 2.40 (s, 3H). ^**13**^**C NMR** (126 MHz, CDCl_3_) δ 199.0, 154.0, 152.0, 138.6, 128.0, 125.0, 117.6, 114.9, 112.3, 71.1, 69.3, 59.2, 55.7, 27.0. **HRMS** (ESI+): m/*z* [M+Na]^+^ calculated for C_11_H_14_NaO_4_ 233.0784; found 233.0788. error: 1.7 ppm. 5-(2-methoxy-6-(2-methoxyethoxy)phenyl)cyclohexane-1,3-dione (**21c**) (221 mg; 0.756 mmol; 14 %). ^**1**^**H NMR** (500 MHz, CD_3_OD) δ 6.91 (d, *J* = 8.9 Hz, 1H), 6.82 (d, *J* = 3.0 Hz, 1H), 6.77 (dd, *J* = 8.8, 3.0 Hz, 1H), 4.12 – 4.07 (m, 2H), 3.75 (s, 3H), 3.74 – 3.72 (m, 2H), 3.72 – 3.65 (m, 1H), 3.41 (s, 3H), 2.75 (dd, *J* = 17.1, 11.8 Hz, 2H), 2.53 (dd, *J* = 17.2, 4.5 Hz, 2H). ^**13**^**C NMR** (126 MHz, CD_3_OD) δ 154.1, 150.5, 132.2, 113.6, 113.2, 111.6, 71.0, 68.1, 57.8, 54.7, 37.5, 34.1. **HRMS** (ESI+): m/*z* [M+H]^+^ calculated for C_16_H_21_O_5_: 293.1384; found 293.1388. error: 1.4 ppm.

6-(2-methoxy-6-(2-methoxyethoxy)phenyl)-4-oxo-4,5,6,7-tetrahydrobenzofuran-3-carboxylic acid (**21**) (60 mg; 0.16 mmol; 22 %). ^**1**^**H NMR** (500 MHz, CDCl_3_) δ 13.29 (s, 1H), 8.18 (s, 1H), 6.89 (d, *J* = 8.9 Hz, 1H), 6.81 (dd, *J* = 8.9, 3.1 Hz, 1H), 6.75 (d, *J* = 3.1 Hz, 1H), 4.14 (t, *J* = 4.7 Hz, 2H), 3.98 – 3.90 (m, 1H), 3.79 (s, 3H), 3.75 – 3. 71 (m, 2H), 3.40 (s, 3H), 3.32 (dd, *J* = 17.5, 10.3 Hz, 1H), 3.26 (dd, *J* = 17.5, 5.7 Hz, 1H), 3.14 (dd, *J* = 17.5, 12.6 Hz, 1H), 2.91 (dd, *J* = 17.5, 4.0 Hz, 1H). ^**13**^**C NMR** (126 MHz, CDCl_3_) δ 198.9, 170.7, 161.5, 153.9, 150.5, 150.4, 130.4, 118.1, 116.8, 114.4, 113.4, 112.3, 71.0, 68.2, 59.1, 55.7, 40.0, 41.9, 36.7, 28.9. **HRMS** (ESI+): m/*z* [M+H]^+^ calculated for C_19_H20O_7_ 361.1282; found 361.1284. error: 0.7 ppm. **HPLC purity** 94 %.

(*E*)-4-(2,3-dichlorophenyl)but-3-en-2-one (**22b**) (3.44 g; 16 mmol; 83 %). ^**1**^**H NMR** (500 MHz, CDCl_3_) δ 7.94 (d, *J* = 16.3 Hz, 1H), 7.55 (dd, *J* = 7.9, 1.5 Hz, 1H), 7.52 (dd, *J* = 7.9, 1.5 Hz, 1H), 7.26 (t, *J* = 7.9 Hz, 1H), 6.66 (d, *J* = 16.3 Hz, 1H), 2.45 (s, 3H). ^**13**^**C NMR** (126 MHz, CDCl_3_) δ 198.1, 139.1, 135.1, 134.1, 133.1, 131.7, 130.7, 127.6, 125.8, 27.4.

5-(2,3-dichlorophenyl)cyclohexane-1,3-dione (**22c**) (194 mg; 0.754 mmol; 5 %). ^**1**^**H NMR** (500 MHz, d_6_-DMSO) δ 7.56 (dd, *J* = 8.0, 1.5 Hz, 1H), 7.50 (dd, *J* = 7.9, 1.4 Hz, 1H), 7.39 (t, *J* = 7.9 Hz, 1H), 5.32 (s, 1H), 3.80 – 3.70 (m, 1H), 2.70 – 2.54 (m, 2H), 2.44 (dd, *J* = 16.5, 3.7 Hz, 2H). Open ring impurities present (20%). ^**13**^**C NMR** (126 MHz, d_6_-DMSO) δ 143.3, 132.5, 131.1, 129.4, 128.9, 127.0, 104.9, 36.9. **HRMS** (ESI+): m/*z* [M+H]^+^ calculated for C_12_H_10_^35^Cl_2_O_2_ 257.0131; found 257.0137. error: 2.7 ppm.

6-(2,3-dichlorophenyl)-4-oxo-4,5,6,7-tetrahydrobenzofuran-3-carboxylic acid (**22**) (80 mg; 0.246 mmol; 32 %). ^**1**^**H NMR** (500 MHz, CDCl_3_) δ 13.05 (s, 1H), 8.16 (s, 1H), 7.49 (dd, *J* = 8.1, 2.9 Hz, 1H), 7.29 (t, 1H), 7.25 (dd, *J* = 7.8, 1.6 Hz, 1H), 4.33 – 4.13 (m, 1H), 3.39 (dd, *J* = 17.5, 5.1 Hz, 1H), 3.23 – 3.06 (dd, *J*_1_ = 17.5 Hz, *J*_2_ = 6.9 Hz, 1H), 3.00 (d, *J* = 8.4 Hz, 2H). ^**13**^**C NMR** (126 MHz, CDCl_3_) δ 197.2, 169.1, 161.1, 150.8, 140.2, 134.3, 132.0, 129.9, 127.9, 125.1, 118.1, 117.2, 42.1, 38.3, 29.4. **HRMS** (ESI+): m/*z* [M+H]^+^ calculated for C_15_H_11_^35^Cl_2_O_4_ 325.0029; found 325.0038. error: 2.9 ppm. **HPLC purity** >99 %.

(E)-4-(2,3,6-trichlorophenyl)but-3-en-2-one (**23b**) (3.00 g, 12.0 mmol, 91 %). ^**1**^**H NMR** (500 MHz, CDCl_3_) δ 7.56 (d, *J* = 16.6 Hz, 1H), 7.41 (d, *J* = 8.7 Hz, 1H,), 7.34 (d, *J* = 8.7 Hz, 1H), 6.75 (d, *J* = 16.6 Hz, 1H), 2.45 (s, 3H). ^**13**^**C NMR** (126 MHz, CDCl_3_) δ 197.8, 136.8, 135.6, 134.2, 133.2, 132.7, 132.6, 130.3, 129.0, 27.8. 5-(2,3,6-trichlorophenyl)cyclohexane-1,3-dione (**23c**) (419 mg, 1.4 mmol, 12 %). ^**1**^**H NMR** (500 MHz, CD_3_OD) δ 7.41-7.53 (d, 2H), 4.49 (m, 1H), 3.45 (dd, *J* = 17.2, 13.5 Hz, 2H), 2.35 (dd, *J* = 17.2, 4.3 Hz, 2H).^**13**^**C NMR** (126 MHz, CD_3_OD) δ 137.9, 130.7, 129.7, 36.8, 33.4. **HRMS** (ESI+): m/z [M+H]^+^ calculated for C_12_H_9_^35^Cl_3_O_2_: **290.97409**, found: **290.9743**, mass error: 0.6 ppm.

4-oxo-6-(2,3,6-trichlorophenyl)-4,5,6,7-tetrahydrobenzofuran-3-carboxylic acid (**23**) (91 mg, 250 µmol, 13 %). ^**1**^**H NMR** (500 MHz, CDCl_3_) δ 13.2 (s, 1H), 8.15 (s, 1H), 7.44 (d, *J* = 8.7 Hz, 1H), 7.31 (d, *J* = 8.1 Hz, 1H), 4.74 (m, 1H), 3.95 (dd, *J* = 17.8, 12.2 Hz, 1H), 3.80 (dd, *J* = 17.8, 13.7 Hz, 1H), 3.06 (dd, *J* = 17.8, 6.0 Hz, 1H), 2.72 (dd, *J* = 17.6, 4.6 Hz, 1H). ^**13**^**C NMR** (126 MHz, CDCl_3_) δ 197.6, 169.2, 161.1, 150.9, 136.4, 134.4, 133.3, 133.1, 130.6, 130.4, 118.2, 117.1, 38.4, 37.6, 25.5. **HRMS** (ESI+): m/z [M+H]^+^ calculated for C_15_H_9_^35^Cl_3_O_4_: 358.96392, found: 358.9568, mass error: 0.5 ppm. **HPLC purity** 99 %.

(E)-4-(4-chlorophenyl)but-3-en-2-one (**24b**) (3.4 g, 19 mmol, 88 %). ^**1**^**H NMR** (400 MHz, CDCl_3_) δ 7.50 – 7.41 (m, 3H), 7.37 (d, *J* = 8.6 Hz, 2H), 6.68 (d, *J* = 16.3 Hz, 1H), 2.40 (s, 3H). ^**13**^**C NMR** (101 MHz, CDCl_3_) δ 198.2, 142.0, 136.6, 133.1, 129.4, 129.4, 127.6, 27.8.

5-(4-chlorophenyl)cyclohexane-1,3-dione (**24c**) (662 mg, 2.9 mmol, 36 %) ^**1**^**H NMR** (500 MHz, d_6_-DMSO) δ 7.37 (s, 4H), 5.28 (s, 1H), 3.42 – 3.28 (accounting for overlap with D_2_O peak, m, 1H), 2.57 (dd, *J* = 16.3, 11.9 Hz, 2H), 2.38 (dd, *J* = 16.3, 2.5 Hz, 2H). ^**13**^**C NMR** (126 MHz, d_6_-DMSO) δ 142.6, 131.1, 128.9, 128.4, 103.6, 38.1. **HRMS** (ESI+): m/z [M+H]^+^ calculated for C_12_H_11_^35^ClO_2_: 222.04476, found: 222.0450, mass error: 0.9 ppm.

6-(4-chlorophenyl)-4-oxo-4,5,6,7-tetrahydrobenzofuran-3-carboxylic acid (**24**) (80 mg, 0.27 mmol, 38 %). ^**1**^**H NMR** (500 MHz, CDCl_3_) δ 13.04 (s, 1H), 8.12 (s, 1H), 7.39 – 7.35 (m, 2H), 7.25 – 7.21 (m, 2H), 3.66 (m, 1H), 3.33 – 3.24 (dd, *J* = 17.6, 5.0 Hz, 1H), 3.14 (dd, *J* = 17.6, 11.0 Hz, 1H), 2.99 – 2.86 (m, 2H). ^**13**^**C NMR** (126 MHz, CDCl_3_) δ 197.6, 169.5, 161.2, 150.9, 139.4, 133.9, 129.5, 128.2, 118.1, 117.4, 43.8, 40.8, 31.1. **HRMS** (ESI+): m/*z* [M+H]^+^ neutral calculated for C_15_H_11_^35^ClO_4_ 290.03459; neutral found 290.0341. error: −1.7 ppm. **HPLC purity** >99 %.

ethyl 6-(4-chlorophenyl)-4-oxo-4,5,6,7-tetrahydrobenzofuran-3-carboxylate (**24d**) (78 mg, 0.24 mmol, 42 %). ^**1**^**H NMR** (700 MHz, CDCl_3_) δ 7.92 (s, 1H), 7.33 (d, *J* = 8.4 Hz, 2H), 7.21 (d, *J* = 8.4 Hz, 2H), 4.35 (q, *J* = 7.1 Hz, 2H), 3.55 (m, 1H), 3.20 (dd, *J* = 17.0, 5.2 Hz, 1H), 3.04 (dd, *J* = 17.0, 11.0 Hz, 1H), 2.84 – 2.74 (m, 2H), 1.37 (t, *J* = 7.1 Hz, 3H). ^**13**^**C NMR** (176 MHz, CDCl_3_) δ 190.3 (**C**=O), 167.3, 161.9, 148.5, 140.5, 133.3, 129.2, 128.2, 118.9, 117.7, 61.2, 45.9, 40.2, 31.4, 14.3. **HRMS** (ESI+): m/*z* [M+H]^+^ neutral calculated for C_17_H_15_^35^ClO_4_ 318.06589; found 318.0643. error: −1.6 ppm. **HPLC purity** 99 %.

(E)-4-(2-chloro-3-methoxyphenyl)but-3-en-2-one (**25b**) (3.08 g, 14.6 mmol, 83 %). ^**1**^**H NMR** (500 MHz, CDCl_3_) δ 7.99 (d, *J* = 16.2 Hz, 1H), 7.27 (m, 2H), 6.99 (m, 1H), 6.67 (d, *J* = 16.2 Hz, 1H), 3.95 (s, 3H), 2.44 (s, 3H). ^**13**^**C NMR** (126 MHz, CDCl_3_) δ 198.5, 155.6, 139.6, 134.1, 130.1, 127.4, 123.6, 119.3, 113.1, 56.4, 27.2. **HRMS** (ESI+): m/z [M+H]^+^ calculated for C_11_H_11_^35^ClO_2_: 211.05204, found: 211.0519, mass error: −0.4 ppm.

5-(2-chloro-3-methoxyphenyl)cyclohexane-1,3-dione (**25c**) (450 mg, 1.8 mmol, 14 %). ^**1**^**H NMR** (500 MHz, CD_3_OD) δ 7.30 (t, *J* = 8.0 Hz, 1H), 7.04 (dd, *J* = 8.0, 1.1 Hz, 1H), 7.01 (dd, *J* = 8.0, 1.1 Hz, 1H), 3.90 (m, 1H), 3.90 (s, 3H), 2.68 (dd, *J* = 17.1, 11.4 Hz, 2H), 2.58 (dd, *J* = 17.1, 4.6 Hz, 2H). ^**13**^**C NMR** (126 MHz, CD_3_OD) δ 155.5, 141.2, 127.4, 121.5, 118.6, 110.5, 55.3, 37.8, 35.9. **HRMS** (ESI+): m/z [M+H]^+^ calculated for C_13_H_13_^35^ClO_3_: 253.06260, found: 253.0626, mass error: −0.1 ppm.

6-(2-chloro-3-methoxyphenyl)-4-oxo-4,5,6,7-tetrahydrobenzofuran-3-carboxylic acid (**25**) (233 mg, 73 µmol, 46 %). ^**1**^**H NMR** (500 MHz, CDCl_3_) δ 13.12 (s, 1H), 8.14 (s, 1H), 7.28 (t, *J* = 8.0 Hz, 1H), 6.94 (2 x d, 2H), 4.24 (m, 1H), 3.95 (s, 3H), 3.36 (dd, *J* = 17.6, 5.1 Hz, 1H), 3.16 (dd, *J* = 17.6, 10.7 Hz, 1H), 2.99 (m, 2H). ^**13**^**C NMR** (126 MHz, CDCl_3_) δ 197.8, 169.6, 161.2, 155.7, 150.7, 139.5, 127.9, 122.0, 118.6, 118.0, 117.2, 111.2, 56.4, 42.2, 37.6, 29.4. **HRMS** (ESI+): m/z [M+H]^+^ calculated for C_16_H_13_^35^ClO_5_: 321.05243, found: 321.0529, mass error: 1.4 ppm. **HPLC purity** 98 %.

(E)-4-(2,5-dimethoxyphenyl)but-3-en-2-one (**26b**) (1.00 g, 4.8 mmol, 81 %). ^**1**^**H NMR** (500 MHz, CDCl_3_) δ 7.88 (d, *J* = 16.6 Hz, 1H), 7.09 (d, *J* = 3.0 Hz, 1H), 6.95 (dd, *J* = 9.0, 3.0 Hz, 1H), 6.88 (d, *J* = 9.0 Hz, 1H), 6.73 (d, *J* = 16.6 Hz, 1H), 3.88 (s, 3H), 3.81 (s, 3H), 2.41 (s, 3H). ^**13**^**C NMR** (126 MHz, CDCl_3_) δ 199.1, 153.6, 152.8, 138.5, 127.9, 123.9, 117.6, 112.6, 112.4, 56.1, 55.8, 27.1.

5-(2,5-dimethoxyphenyl)cyclohexane-1,3-dione (**26c**) (510 mg, 2.1 mmol, 56 %). ^**1**^**H NMR** (500 MHz, CD_3_OD) δ 6.92 (d, *J* = 8.8 Hz, 1H), 6.82 (m, 1H), 6.80 (m, 1H), 3.82 (s, 3H), 3.76 (s, 3H), 3.67 (m, 1H), 2.71 (dd, *J* = 17.1, 11.6 Hz, 2H), 2.51 (dd, *J* = 17.1, 4.5 Hz, 2H). ^**13**^**C NMR** (126 MHz, CD_3_OD) δ 153.8, 151.3, 131.6, 113.5, 111.4, 111.4, 54.9, 54.7, 33.8. **HRMS** (ESI+): m/z [M+H]^+^ calculated for C_14_H16O_4_: 249.11214, found: 249.1120, mass error: −0.4 ppm.

6-(2,5-dimethoxyphenyl)-4-oxo-4,5,6,7-tetrahydrobenzofuran-3-carboxylic acid (**26**) (12 mg, 37 µmol, 2 %). ^**1**^**H NMR** (500 MHz, d_6_-DMSO) δ 8.62 (s, 1H), 6.96 (d, *J* = 8.9 Hz, 1H), 6.90 (d, *J* = 3.0 Hz, 1H), 6.83 (dd, *J* = 8.9, 3.0 Hz, 1H), 3.88 (m, 1H), 3.76 (s, 1H), 3.70 (s, 1H), 3.26 (dd, *J* = 17.1, 10.8 Hz, 1H), 3.15 (dd, *J* = 17.1, 5.0 Hz, 1H), 3.05 (dd, *J* = 16.6, 12.4 Hz, 1H), 2.62 (dd, *J* = 16.6, 3.8 Hz, 1H). ^**13**^**C NMR** (126 MHz, d_6_-DMSO) δ 196.4, 170.4, 162.0, 153.6, 151.1, 150.5, 131.1, 117.6, 117.2, 114.4, 112.6, 112.4, 56.4, 55.8, 43.1, 34.9, 28.9. **HRMS** (ESI+): m/z [M+H]^+^ calculated for C_17_H_16_O_6_: 317.10197, found: 317.1024, mass error: 1.3 ppm. **HPLC purity** 94 %.

(E)-4-(4-chloro-2-methoxyphenyl)but-3-en-2-one (**27b**) (1.04 g, 4.9 mmol, 74 %). ^**1**^**H NMR** (500 MHz, CDCl_3_) δ 7.81 (d, *J* = 16.5 Hz, 1H), 7.49 (d, *J* = 8.3 Hz, 1H), 6.99 (dd, *J* = 8.3, 2.0 Hz, 1H), 6.93 (d, *J* = 2.0 Hz, 1H), 6.75 (d, *J* = 16.5 Hz, 1H), 3.92 (s, 3H), 2.40 (s, 3H). ^**13**^**C NMR** (126 MHz, CDCl_3_) δ 198.8, 158.7, 137.5, 137.3, 129.2, 127.9, 122.1, 121.1, 112.0, 55.9, 27.3.

5-(4-chloro-2-methoxyphenyl)cyclohexane-1,3-dione (**27c**) (255 mg, 1.0 mmol, 21 %). ^**1**^**H NMR** (500 MHz, d_6_-DMSO) δ 7.25 (d, *J* =8.2 Hz, 1H), 7.05 (d, *J* = 2.1 Hz, 1H), 6.98 (dd, *J* = 8.2, 2.1 Hz, 1H), 5.28 (s, 1H), 3.82 (s, 3H), 2.55 (m, 2H), 2.35 (m, 2H). ^**13**^**C NMR** (126 MHz, d_6_-DMSO) δ 158.0, 132.5, 130.4, 128.7, 120.7, 111.9, 103.8, 56.4, 32.8. **HRMS** (ESI+): m/z [M+H]^+^ calculated for C_13_H_13_^35^ClO_3_: 253.06260, found: 253.0630, mass error: 1.7 ppm. 6-(4-chloro-2-methoxyphenyl)-4-oxo-4,5,6,7-tetrahydrobenzofuran-3-carboxylic acid (**27**) (17 mg, 50 µmol, 6 %). ^**1**^**H NMR** (500 MHz, CDCl_3_) δ 13.20 (s, 1H), 8.13 (s, 1H), 7.12 (d, *J* = 8.2 Hz, 1H), 6.98 (dd, *J* = 8.2, 2.0 Hz, 1H), 6.94 (d, *J* = 2.0 Hz, 1H), 3.93 (m, 1H), 3.89 (s, 3H), 3.24 (m, 2H), 3.07 (dd, *J* = 17.1, 12.6 Hz, 1H), 2.88 (dd, *J* = 17.1, 4.0 Hz, 1H). ^**13**^**C NMR** (126 MHz, CDCl_3_) δ 198.5, 170.2, 161.3, 157.6, 150.5, 134.4, 128.2, 127.3, 121.0, 118.0, 117.1, 55.7, 41.9, 35.9, 28.9. **HRMS** (ESI+): m/z [M+H]^+^ calculated for C_16_H_13_^35^ClO_5_: 321.05243, found: 321.0527, mass error: 0.8 ppm. **HPLC purity** 93 %.

(E)-4-(4-chloro-3-methoxyphenyl)but-3-en-2-one (**28b**) (2.5 g, 12 mmol, 51 %). ^**1**^**H NMR** (500 MHz, CDCl_3_) δ 7.44 (d, *J* = 16.2 Hz, 1H), 7.37 (d, *J* = 8.0 Hz, 1H), 7.11 – 7.04 (m, 2H), 6.68 (d, *J* = 16.2 Hz, 1H), 3.93 (s, 3H), 2.38 (s, 3H). ^**13**^**C NMR** (126 MHz, CDCl_3_) δ 198.47, 155.70, 142.69, 134.69, 131.03, 127.94, 125.39, 121.92, 111.23, 56.53, 27.98. **HRMS (ESI):** m/z [M+H]^+^ calculated for C_11_H_11_^35^ClO_2_: 210.04476, found: 210.0447, mass error: −0.4 ppm.

5-(4-chloro-3-methoxyphenyl)cyclohexane-1,3-dione (**28c**) (202 mg, 0.80 mmol, 17 %). ^**1**^**H NMR** (500 MHz, CD_3_OD) δ 7.30 (d, *J* = 8.1 Hz, 1H), 7.03 (d, *J* = 2.0 Hz, 1H), 6.88 (dd, *J* = 8.2, 2.0 Hz, 1H), 3.88 (s, 3H), 3.38 (s, 1H), 2.74 – 2.64 (m, 2H), 2.60 – 2.52 (m, 2H). ^**13**^**C NMR** (126 MHz, CD_3_OD) δ 156.56, 144.93, 131.14, 121.87, 120.60, 112.38, 56.60, 40.78. **HRMS (ESI+):** m/z [M+H]^+^ calculated for C_13_H_14_O_3_^35^Cl 253.0631, found: 253.0631, mass error: 0.0 ppm.

6-(4-chloro-3-methoxyphenyl)-4-oxo-4,5,6,7-tetrahydrobenzofuran-3-carboxylic acid (**28**) (72 mg, 0.22 mmol, 36 %). ^**1**^**H NMR** (500 MHz, d_6_-DMSO): δ 8.45 (s, 1H), 7.38 (d, *J* = 8.1 Hz, 1H), 7.22 (d, *J* = 1.9 Hz, 1H), 6.97 (dd, *J* = 8.2, 2.0 Hz, 1H), 3.86 (s, 3H), 3.67 (m, 1H), 3.28 – 3.22 (m, 2H), 3.06 (dd, *J* = 16.5, 12.7 Hz, 1H), 2.69 (dd, *J* = 16.5, 3.9 Hz, 1H). ^**13**^**C NMR** (126 MHz, d_6_-DMSO): δ 195.27, 169.73, 161.66, 154.57, 150.16, 143.07, 129.83, 119.90, 119.62, 117.26, 117.00, 112.01, 56.14, 44.05, 30.08. **HRMS (ESI):** m/z [M+H]^+^ calculated for C_16_H_13_^35^ClO_5_: 320.04515, found: 320.0447, mass error: −1.3 ppm. **HPLC purity** >99 %.

5-(thiophen-2-yl)cyclohexane-1,3-dione (**29c**) (276 mg, 14.2 mmol, 21 %). ^**1**^**H NMR** (500 MHz, CD_3_OD) δ 7.27 (dd, *J* = 4.0, 2.5 Hz, 1H), 6.98 – 6.97 (m, 2H), 5.45 (s, 2H), 3.71 (m, 1H), 2.76 (dd, *J* = 16.8, 4.8 Hz, 2H), 2.67 (dd, *J* = 17.1, 10.4 Hz, 2H). ^**13**^**C NMR** (126 MHz, CD_3_OD) δ 146.6, 126.4, 123.3, 123.1, 40.1, 39.3.

4-oxo-6-(thiophen-2-yl)-4,5,6,7-tetrahydrobenzofuran-3-carboxylic acid (**29**) (42 mg, 161 mmol, 16 %). ^**1**^**H NMR** (500 MHz, CDCl_3_) δ 13.06 (s, 1H), 8.14 (s, 1H), 7.27 (dd, *J* = 5.0, 1.2 Hz, 1H), 7.02 (dd, *J* = 5.0, 3.5 Hz, 1H), 6.96 (dd, *J* = 3.5, 1.2 Hz, 1H), 4.01 (m, 1H), 3.47 (dd, *J* = 17.6, 5.2 Hz, 1H), 3.24 (dd, *J* = 17.6, 9.9 Hz, 1H), 3.13 (dd, *J* = 17.2, 4.1 Hz, 1H), 2.99 (dd, *J* = 17.2, 11.0 Hz, 1H). ^**13**^**C NMR** (126 MHz, CDCl_3_) δ 197.1, 168.9, 161.1, 150.8, 144.4, 127.2, 124.4, 124.3, 118.0, 117.4, 44.6, 36.5, 31.9. **HRMS** (ESI+): m/z [M+H]^+^ calculated for C_13_H10O_4_S: 261.0215, found: 261.0221, mass error: 2.5 ppm. **HPLC purity:** 96 %.

(E)-4-(3-bromothiophen-2-yl)but-3-en-2-one (**30b**) (1.50 g, 6.5 mmol, 83 %). ^**1**^**H NMR** (500 MHz, CDCl_3_) δ 7.70 (d, *J* = 16.3 Hz, 1H), 7.39 (d, *J* = 5.3 Hz, 1H), 7.08 (d, *J* = 5.3 Hz, 1H), 6.58 (d, *J* = 16.3 Hz, 1H), 2.40 (s, 3H). ^**13**^**C NMR** (126 MHz, CDCl_3_) δ 197.6, 134.4, 133.8, 131.6, 128.2, 127.5, 116.7, 27.6. **HRMS** (ESI+): m/z [M+H]^+^ calculated for C_8_H_7_BrOS: 230.94738, found: 230.9476, mass error: 1.0 ppm.

5-(3-bromothiophen-2-yl)cyclohexane-1,3-dione (**30c**) (135 mg, 0.5 mmol, 9 %). ^**1**^**H NMR** (500 MHz, CD_3_OD) δ 7.40 (d, *J* = 5.3 Hz, 1H), 7.01 (dd, *J* = 5.3, 1.0 Hz, 1H), 3.85 (m, 1H), 2.72 (dd, *J* = 16.9, 5.0 Hz, 2H), 2.62 (dd, *J* = 16.9, 10.7 Hz, 2H). ^**13**^**C NMR** (126 MHz, CD_3_OD) δ 140.3, 129.8, 123.9, 108.2, 38.6, 34.5. **HRMS** (ESI+): m/z [M+H]^+^ calculated for C_10_H_9_^79^BrO_2_S: 272.95794, found: 272.9579, mass error: 0.0 ppm.

6-(3-bromothiophen-2-yl)-4-oxo-4,5,6,7-tetrahydrobenzofuran-3-carboxylic acid (**30**) (34 mg, 99 µmol, 34 %). ^**1**^**H NMR** (500 MHz, CDCl_3_) δ 13.03 (s, 1H), 8.17 (s, 1H), 7.28 (d, *J* = 5.2 Hz, 1H), 7.03 (d, *J* = 5.2 Hz, 1H), 4.14 (m, 1H), 3.45 (dd, *J* = 17.6, 5.1 Hz, 1H), 3.19 (dd, *J* = 17.6, 10.4 Hz, 1H), 3.09 (dd, *J* = 17.2, 4.2 Hz, 1H), 2.94 (dd, *J* = 17.2, 11.6 Hz, 1H). ^**13**^**C NMR** (126 MHz, CDCl_3_) δ 195.6, 168.6, 161.0, 150.9, 138.1, 130.6, 124.3, 118.1, 117.2, 109.8, 43.3, 36.0, 30.5. **HRMS (ESI):** m/z [M+H]^+^ calculated for C_13_H_9_^79^BrO_4_S: 339.94049, found: 339.9403, mass error: −0.6 ppm. **HPLC purity** >99 %.

(E)-4-(5-bromothiophen-2-yl)but-3-en-2-one (**31b**) (3.25 g, 14.0 mmol, 67 %). ^**1**^**H NMR** (500 MHz, CDCl_3_) δ 7.52 (d, *J* = 15.9 Hz, 1H), 7.05 (s, 2H), 6.44 (d, *J* = 15.9 Hz, 1H), 2.35 (s, 3H). ^**13**^**C NMR** (126 MHz, CDCl_3_) δ 197.4, 141.3, 134.7, 131.9, 131.3, 125.9, 116.5, 27.9. **HRMS** (ESI+): m/z [M+H]^+^ calculated for C_8_H_7_^79^BrOS: 230.94748, found: 230.9475, mass error: 0.4 ppm.

5-(5-bromothiophen-2-yl)cyclohexane-1,3-dione (**31c**) (264 mg, 1.0 mmol, 14 %). ^**1**^**H NMR** (500 MHz, CD_3_OD) δ 6.96 (d, *J* = 3.8 Hz, 1H), 6.78 (dd, *J* = 3.8, 1.0 Hz, 1H), 3.66 (m, 1H), 2.74 (dd, *J* = 17.1, 4.8 Hz, 2H), 2.63 (dd, *J* = 17.1, 10.3 Hz, 2H). ^**13**^**C NMR** (126 MHz, CD_3_OD) δ 148.7, 129.5, 124.2, 109.3, 35.0. **HRMS** (ESI+): m/z [M+H]^+^ calculated for C_10_H_9_^79^BrO_2_S: 272.95794, found: 272.9581, mass error: 0.6 ppm.

6-(5-bromothiophen-2-yl)-4-oxo-4,5,6,7-tetrahydrobenzofuran-3-carboxylic acid (**31**) (12 mg, 34 µmol, 4 %). ^**1**^**H NMR** (500 MHz, d_6_-DMSO) δ 8.44 (s, 1H), 7.10 (d, *J* =3.8 Hz, 1H), 6.89 (d, *J* =3.8, 1.0 Hz, 1H), 3.95 (m, 1H), 3.19 (dd, *J* = 17.2, 9.6 Hz, 1H), 2.93 (dd, *J* = 16.5, 10.3 Hz, 1H), 2.87 (dd, *J* = 16.5, 4.7 Hz, 1H). ^**13**^**C NMR** (126 MHz, d_6_-DMSO): δ 194.3, 169.0, 162.1, 150.5, 148.2, 130.6, 125.9, 117.7, 117.7, 109.6, 44.9, 35.9, 30.9. **HRMS** (ESI+): m/z [M+H]^+^ calculated for C_13_H_9_^79^BrO_4_S: 340.94777, found: 340.9481, mass error: 1.0 ppm. **HPLC purity** 90 %.

(E)-4-(5-chlorothiophen-2-yl)but-3-en-2-one (**32b**) (0.93 g, 5.0 mmol, 83 %). ^**1**^**H NMR** (500 MHz, CDCl_3_) δ 7.49 (d, *J* = 16.0 Hz, 1H), 7.08 (d, *J* = 3.9 Hz, 1H), 6.90 (d, *J* = 3.9 Hz, 1H), 6.41 (d, *J* = 16.0 Hz, 1H), 2.34 (s, 3H). ^**13**^**C NMR** (126 MHz, CDCl_3_) δ 197.3, 138.5, 135.0, 133.7, 131.2, 127.6, 125.6, 27.9.

5-(5-chlorothiophen-2-yl)cyclohexane-1,3-dione (**32c**) (439 mg, 1.9 mmol, 42 %). ^**1**^**H NMR** (500 MHz, CD_3_OD) δ 6.83 (d, *J* = 3.8 Hz, 1H), 6.79 (dd, *J* = 3.8, 1.0 Hz, 1H), 3.63 (m, 1H), 2.73 (dd, *J* = 17.2, 4.8 Hz, 2H), 2.62 (dd, *J* = 17.2, 10.2 Hz, 2H). ^**13**^**C NMR** (126 MHz, CD_3_OD) δ 145.9, 127.2, 125.7, 123.1, 39.6, 35.0. **HRMS** (ESI+): m/z [M+H]^+^ calculated for C_10_H_9_^35^ClO_2_S: 229.00846, found: 229.0083, mass error: −0.6 ppm.

6-(5-chlorothiophen-2-yl)-4-oxo-4,5,6,7-tetrahydrobenzofuran-3-carboxylic acid (**32**) (66 mg, 220 µmol, 13 %). ^**1**^**H NMR** (500 MHz, d_6_-DMSO) δ 8.44 (s, 1H), 7.00 (d, *J* = 3.8 Hz, 1H), 6.92 (d, *J* = 3.8, 1.0 Hz, 1H), 3.93 (m, 1H), 3.16 (dd, *J* = 17.2, 9.6 Hz, 1H), 2.93 (dd, *J* = 16.5, 10.4 Hz, 1H), 2.87 (dd, *J* = 16.5, 4.6 Hz, 1H). ^**13**^**C NMR** (126 MHz, d_6_-DMSO) δ 194.3, 169.0, 162.1, 150.5, 145.5, 127.1, 126.5, 124.9, 117.7, 117.7, 44.9, 35.9, 30.9. **HRMS** (ESI+): m/z [M+H]^+^ calculated for C_13_H_9_^35^ClO_4_S: 296.99829, found: 296.9982, mass error: −0.3 ppm. **HPLC purity** 95 %.

(E)-4-(4-bromothiophen-2-yl)but-3-en-2-one (**33b**) (2.37 g, 10.2 mmol, 89 %). ^**1**^**H NMR** (500 MHz, CDCl_3_) δ 7.54 (d, *J* = 16.0 Hz, 1H,), 7.31 (br s, 1H), 7.22 (br s, 1H), 6.55 (d, *J* = 16.0 Hz, 1H), 2.36 (s, 3H). ^**13**^**C NMR** (126 MHz, CDCl_3_) δ 197.3, 140.5, 134.1, 132.9, 126.6, 125.6, 111.1, 27.9.

5-(4-bromothiophen-2-yl)cyclohexane-1,3-dione (**33c**) (1.40 g, 5.2 mmol, 61 %). ^**1**^**H NMR** (500 MHz, CD_3_OD) δ 7.27 (s, 1H), 6.94 (s, 1H), 3.70 (m, 1H), 2.75 (dd, *J* = 17.0, 4.8 Hz, 2H), 2.63 (dd, *J* = 17.0, 10.3 Hz, 2H). ^**13**^**C NMR** (126 MHz, CD_3_OD) δ 148.4, 126.3, 120.8, 108.8, 39.5, 34.6. **HRMS** (ESI+): m/z [M+H]^+^ calculated for C_10_H_9_^79^BrO_2_S: 272.95794, found: 272.9580, mass error: 0.0 ppm. 6-(4-bromothiophen-2-yl)-4-oxo-4,5,6,7-tetrahydrobenzofuran-3-carboxylic acid (**33**) (26 mg, 76 µmol, 3 %). ^**1**^**H NMR** (500 MHz, d_6_-DMSO) δ 13.01 (s, 1H), 8.46 (s, 1H), 7.56 (d, *J* = 1.5 Hz, 1H), 7.10 (dd, *J* = 1.5, 1.0 Hz, 1H), 3.98 (m, 1H), 3.43 (dd, *J* = 17.2, 5.4 Hz, 1H & D_2_O peak), 3.22 (dd, *J* = 17.2, 9.8 Hz, 1H), 2.96 (dd, *J* = 16.5, 10.8 Hz, 1H), 2.86 (dd, *J* = 16.5, 4.4 Hz, 1H). ^**13**^**C NMR** (126 MHz, d_6_-DMSO) δ 194.2, 169.1, 162.1, 150.5, 148.1, 127.3, 122.4, 117.6, 117.7, 108.8, 44.8, 35.6, 30.8. **HRMS** (ESI+): m/z [M+H]^+^ calculated for C_13_H_9_^79^BrO_4_S: 340.94777, found: 340.9482, mass error: 1.3 ppm. **HPLC purity** >99 %.

ethyl 6-(4-bromothiophen-2-yl)-4-oxo-4,5,6,7-tetrahydrobenzo-furan-3-carboxylate (**33d**) (140 mg, 0.38 mmol, 15 %). ^**1**^**H NMR** (500 MHz, d_6_-DMSO) δ 7.95 (s, 1H), 7.14 (d, *J* = 1.3 Hz, 1H), 6.86 (d, *J* = 1.3, 0.9 Hz, 1H), 4.37 (q, *J* = 7.2 Hz, 2H), 3.83 (m, 1H), 3.37 (dd, *J* = 17.0, 5.0 Hz, 1H), 3.11 (dd, *J* = 17.0, 10.0 Hz, 1H), 2.96 (dd, *J* = 16.1, 4.1 Hz, 1H), 2.81 (dd, *J* = 16.1, 11.6 Hz, 1H), 1.39 (t, *J* = 7.2 Hz, 3H). ^**13**^**C NMR** (126 MHz, d_6_-DMSO) δ 198.1, 166.3, 161.6, 148.5, 146.9, 126.8, 121.2, 118.9, 117.6, 109.6, 61.1, 46.2, 35.9, 31.8, 14.2. **HRMS (ESI):** m/z [M+H]^+^ calculated for C_15_H_13_^79^BrO_4_S: 390.96101, found: 390.9613, mass error: 0.7 ppm. **HPLC purity** 95 %.

(E)-4-(4,5-dibromothiophen-2-yl)but-3-en-2-one (**34b**) (3.08 g, 9.9 mmol, 89 %). ^**1**^**H NMR** (500 MHz, CDCl_3_) δ 7.45 (dd, *J* = 16.1, 0.5 Hz, 1H), 7.10 (s, 1H), 6.46 (d,, *J* = 16.1 Hz, 1H), 2.35 (s, 3H). ^**13**^**C NMR** (126 MHz, CDCl_3_) δ 197.0, 140.5, 133.5, 133.1, 126.6, 115.3, 114.7, 28.1. **HRMS** (ESI+): m/z [M+H]^+^ calculated for C_8_H_6_^79^Br_2_OS: 308.85789, found: 308.8580, mass error: 0.3 ppm.

5-(4,5-dibromothiophen-2-yl)cyclohexane-1,3-dione (**34c**) (508 mg, 1.4 mmol, 11 %). ^**1**^**H NMR** (500 MHz, d_6_-DMSO): δ 6.98 (s, 1H), 5.28 (s, 1H), 2.48-2.63 (m, 4H). ^**13**^**C NMR** (126 MHz, d_6_-DMSO): δ 149.8, 127.4, 113.3, 108.5, 104.2, 34.8. **HRMS** (ESI+): m/z [M+H]^+^ calculated for C_10_H_8_^79^Br_2_O_2_S: 350.86846, found: 250.8680, mass error: −1.2 ppm.

6-(4,5-dibromothiophen-2-yl)-4-oxo-4,5,6,7-tetrahydrobenzofuran-3-carboxylic acid (**34**) (26 mg, 62 µmol, 5 %). ^**1**^**H NMR** (500 MHz, d_6_-DMSO) δ 12.97 (s, 1H), 8.46 (s, 1H), 7.12 (s, 1H), 3.95 (m, 1H), 3.41 (dd, *J* = 17.1, 5.1 Hz, 1H), 3.21 (dd, *J* = 17.6, 10.2 Hz, 1H), 2.94 (dd, *J* = 16.4, 10.7 Hz, 1H), 2.87 (dd, *J* = 16.4, 4.4 Hz, 1H). ^**13**^**C NMR** (126 MHz, d_6_-DMSO) δ 193.8, 168.8, 162.1, 150.5, 148.4, 127.9, 117.7, 117.7, 113.5, 109.1, 44.5, 36.0, 30.5. **HRMS** (ESI+): m/z [M+H]^+^ calculated for C_13_H_8_^79^Br_2_O_4_S: 418.85829, found: 418.8577, mass error: −1.3 ppm. **HPLC purity** 90 %.

ethyl 2-(1-oxo-1,2,3,4-tetrahydronaphthalen-2-yl)acetate (**36**). A solution of 2.5 M nBuLi (7.3 mL, 18.3 mmol, 1.1 eq.) in hexane was slowly added to a cold (−78 °C) solution of diisopropylamine (2.569 mL, 18.3 mmol, 1.1 eq.) in THF (8.3 mL). After 0.5 h, alpha tetralone **35** (2.22mL, 16.7 mmol, 1 eq.) in THF (0.83 mL) was added dropwise, followed by addition of dry ethyl bromopyruvate (2.03 mL, 18.3 mmol, 1.1 eq.). The reaction was allowed to warm from −78 °C to rt and stirred for 16 h. The reaction was quenched by dilution with H_2_O, extracted with EtOAc, dried over anhydrous MgSO_4_ and purified by reversed-phase column chromatography (H_2_O / MeCN 9:1 – 9:11) to yield ester **36** as a white powder (555 mg, 2.39 mmol, 14.3 %). ^**1**^**H NMR** (500 MHz, CDCl_3_) δ 8.05 (dd, *J* = 7.9, 1.3 Hz, 1H), 7.49 (t, *J* = 7.7 Hz, 1H), 7.33 (t, *J* = 7.7 Hz, 1H), 7.27 (d, *J* = 7.6 Hz, 1H), 4.21 (m, 2H), 3.15 (m, 1H), 3.10 (m, 1H), 3.07 (m, 1H), 3.02 (m, 1H), 2.44 (m, 1H), 2.27 (m, 1H), 1.99 (m, 1H), 1.31 (t, *J* = 7.1 Hz, 3H). ^**13**^**C NMR** (126 MHz, CDCl_3_) δ 198.4, 172.6, 144.0, 133.4, 132.2, 128.8, 127.5, 126.7, 60.6, 44.8, 35.2, 29.3, 14.2.

2-(1-oxo-1,2,3,4-tetrahydronaphthalen-2-yl)acetic acid (**37**). To a solution of the ester **36** (555 mg, 2.39 mmol, 1 eq.) in a 2:1 mix of THF/MeOH (8 mL) was added 3M NaOH (2.39 mL, 7.17 mmol, 3 eq.) and the reaction was stirred for 2 days at rt. The crude mixture was subjected to a stream of N_2_ to remove the solvent and dissolved in water (26 mL). An orangey/pink solution resulted and was acidified to pH 2 (6M HCl) before extraction with EtOAc, drying over anhydrous Na_2_SO_4_ and purification by HPLC to yield the product of the hydrolysis as a white powder (175 mg, 0.86 mmol, 36 %). ^**1**^**H NMR** (500 MHz, CDCl_3_) δ 8.05 (dd, *J* = 7.6, 1.3 Hz, 1H), 7.50 (t, *J* = 7.7 Hz, 1H), 7.33 (t, *J* = 7.7 Hz, 1H), 7.27 (d, *J* = 7.6 Hz, 1H), 3.16 (m, 1H), 3.10 (m, 1H), 3.07 (m, 1H), 3.02 (m, 1H), 2.52 (m, 1H), 2.30 (m, 1H), 2.02 (m, 1H). ^**13**^**C NMR** (126 MHz, CDCl_3_) δ 198.5, 178.3, 144.1, 133.6, 132.0, 128.8, 127.6, 126.8, 44.7, 35.0, 29.3. **HPLC purity** >99 %.

## ASSOCIATED CONTENT

### Supporting Information

The Supporting Information is available free of charge on the ACS Publications website.

^1^H NMR, ^13^C NMR and HPLC spectra for the compounds, as well as mass spectrometry and MALDI-TOF spectra, raw data from differential scanning fluorimetry, statistical analysis and supercritical fluid chromatography spectra of enantiomers. (PDF) Molecular formula strings (CSV)

### Accession Codes

PDB code for BTK PH domain with bound **1** is 6TUH.

PDB code for BTK PH domain with bound **2** is 7I9I

PDB code for BTK PH domain with bound **4** is 7I96

PDB code for BTK PH domain with bound **5** is 9T23

PDB code for BTK PH domain with bound **7** is 7I90

PDB code for BTK PH domain with bound **8** is 7I97

PDB code for BTK PH domain with bound **9** is 7I92

PDB code for BTK PH domain with bound **10** is 7I98

PDB code for BTK PH domain with bound **11** is 9T3M

PDB code for BTK PH domain with bound **12** is 973G

PDB code for BTK PH domain with bound **13** is 7I91

PDB code for BTK PH domain with bound **14** is 7I9D

PDB code for BTK PH domain with bound **16** is 7I9F

PDB code for BTK PH domain with bound **17** is 9T1V

PDB code for BTK PH domain with bound **18** is 7I9E

PDB code for BTK PH domain with bound **19** is 9T0T

PDB code for BTK PH domain with bound **22** is 7I99

PDB code for BTK PH domain with bound **24** is 7I9B

PDB code for BTK PH domain with bound **25** is 7I95

PDB code for BTK PH domain with bound **26** is 7I9H

PDB code for BTK PH domain with bound **27** is 7I93

PDB code for BTK PH domain with bound **28** is 7I9G

PDB code for BTK PH domain with bound **29** is 9T21

PDB code for BTK PH domain with bound **30** is 7I9C

PDB code for BTK PH domain with bound **31** is 9RM0

PDB code for BTK PH domain with bound **32** is 7I9B

PDB code for BTK PH domain with bound **33** is 9RN5

## AUTHOR INFORMATION

### Author Contributions

M. Hyvönen conceptualized the project. The first draft was written by R. C. Bizga Nicolescu and R. M. West, with P. Brear contributing the X-ray crystallography section. The manuscript was written through contributions of all authors. All authors contributed to proof reading and have given approval to the final version of the manuscript.

### Notes

The authors declare no competing financial interest.

## ACKNOWLEDGMENT

We would like to thank Peter Gierth, Andrew Mason and Duncan Howe for their support in NMR analysis, Dijana Matak-Vinkovic, Roberto Canales and Asha Boodhun for maintaining the instruments and methods for the protein mass spectrometry, Kristina Kostadinova (Spring Group) for the MALDI spectra, Linwei Zeng (Spring Group) for the analytical chiral LCMS of compound **19** and Arqum Anwar (Babraham Institute) for conducting biological studies. We thank X-ray crystallographic and Biophysics facilities at the Department of Biochemistry for access to instrumentation and support. We are grateful for access to beamlines I04, I03 and I04-1 at Diamond Light Source (Proposals mx14043, mx25402, mx33658 and mx40158). The project was supported by MRC Confidence in Concept award. R.

M. West acknowledges support from the UK Engineering and Physical Sciences Research Council (EPSRC) Centre of Doctoral Training in Automated Chemical Synthesis Enabled by Digital Molecular Technologies (SynTech) [EP/S024220/1]. R. C. Bizga Nicolescu acknowledges support from Trinity College Cambridge. T. Deingruber acknowledges a studentship from Astra-Zeneca. F.J.Pérez-Areales acknowledges Fundación Ramón Areces (reference BEVP31A6160) and Marie Skłodowska-Curie Individual Fellowships (MSCA-IF-2020, grant number 101025271). The Spring group research was supported by grants from UKRI (EP/P020291/1). For the purpose of Open Access, the author has applied a CC BY public copyright licence to any Author Accepted Manuscript (AAM) version arising.

## ABBREVIATIONS

ATP: Adenosine triphosphate
BTK: Bruton’s Tyrosine Kinase
DCM: dichloromethane
DMF: dimethyl formamide
DMSO: di-methyl sulfoxide
DSF: Differential Scanning Fluorimery
ERBB4: erythroblastic leukemia viral oncogene 4
FCC: flash column chromatography
FDA: Food and Drug Administration
HEPES: 4-(2-hydroxyethyl)-1-piperazineethanesulfonic acid
HPLC: high-performance liquid chromatography
HRMS: high resolution mass spectrometry
IC_50_: half maximal inhibitory concentration
IP4: Inositol 1,3,4,5-tetrakisphosphate
ITC: iso-thermal titration calorimetry
K_d_: dissociation constant
LCMS: liquid chromatography-mass spectrometry
LDA: Lithium diisopropylamide
MALDI-TOF: Matrix-assisted laser desorption/ionization Time of Flight
MOE: Molecular Operating Environment
NMR: nuclear magnetic resonance
PDB: Protein Data Bank
PE: petroleum ether
PH: Pleckstrin Homology
PIP3: Phosphatidylinositol (3,4,5)-trisphosphate
PLC: Phopholipase C
rt: room temperature
SH2: Src Homology 2
SH3: Src Homology 3
SFC: supercritical fluid chromatography
s-phos: 2-Dicyclohexylphosphino-2′,6′-dimethoxybiphenyl
TCEP: tris(2-carboxyethyl)phosphine
THF: tetrahydrofuran
TLC: thin layer chromatography
WT: wildtype

## Footnotes

† Ethyl bromopyruvate was purchased from Sigma Aldrich. The same chemical from other sources didn’t give a successful reaction.

