## Supplemental data for "Targeting a Pleckstrin Homology Domain with a Lysine-Reactive Covalent Binder"

### Supporting Information

### S1 X-Ray Crystallography

**Table S1.** Crystal structure data collection, data reduction and refinement statistics.

| Protein | BtkS | BtkS | BtkS |
|---|---|---|---|
| PDB code | 7I9I | 7I96 | 9T23 |
| Ligand | **2** | **4** | **5** |
| **Data Collection:** | | | |
| Beamline | DIAMOND BEAMLINE i04-1 | DIAMOND BEAMLINE i04-1 | DIAMOND BEAMLINE i03 |
| Wavelength (Å) | 0.9763 | 0.9179 | 0.9763 |
| Resolution range (Å) | 56.08 - 1.83 (1.86- 1.83) | 78.67 - 2.52 (2.56 - 2.52) | 57.12 - 1.84 (1.88 - 1.84) |
| Space group | $P\,1\,2_1\,1$ | $P\,1\,2_1\,1$ | $P\,1\,2_1\,1$ |
| Cell (a b c) (Å) | 67.95 66.14 79.84 | 69.10 67.57 79.97 | 68.89 66.27 79.57 |
| Cell (α β γ) (°) | 90.00 100.24 90.00 | 90.00 100.38 90.00 | 90.00 101.88 90.00 |
| Total reflections | 1895375 (89115) | 190655 (9769) | 415394 (17983) |
| Unique reflections | 61602 (3078) | 24550 (1217) | 60236 (2737) |
| Multiplicity | 30.8 (29.0) | 7.8 (8.0) | 6.9 (6.6) |
| Completeness (%) | 100.0 (100.0) | 99.4 (98.5) | 99.5 (91.9) |
| Mean I/sigma(I) | 8.7 (0.2) | 8.7 (0.5) | 7.1 (0.3) |
| R-merge | 0.195 (8.65) | 0.199 (3.23) | 0.145 (3.44) |
| R-pim | 0.035 (1.62) | 0.077 (1.22) | 0.060 (1.44) |
| CC-half | 0.999 (0.45) | 0.991 (0.17) | 0.994 (0.33) |
| **Refinement:** | | | |
| Refinement resolution range (Å) | 26.53 - 1.83 (1.94 - 1.83) | 78.67 - 2.52 (2.54 - 2.52) | 33.71 - 1.84 (1.87 - 1.84) |
| No. reflections | 50423 (1009) | 24507 (491) | 58330 (1167) |
| No. reflections (Rfree) | 2541 (64) | 1246 (30) | 1943 (91) |
| R-factor | 0.242 (0.373) | 0.250 (0.403) | 0.265 (0.420) |
| Rfree | 0.292 (0.423) | 0.308 (0.425) | 0.319 (0.476) |
| Number of total atoms | 5562 | 5472 | 5749 |
| atoms for macromolecules | 5307 | 5352 | 5552 |
| atoms for ligands | 79 | 40 | 26 |
| atoms for waters | 176 | 80 | 171 |
| Average B-factor ($Å^2$) | 57.4 | 73.8 | 53.7 |
| RMS(bonds) (Å) | 0.009 | 0.007 | 0.009 |
| RMS(bond angles) (°) | 1.04 | 1 | 1.02 |
| RMS(dihedral angles) (°) | 3.49 | 3.2 | 3.51 |

| Protein | BtkA | BtkS | BtkS |
|---|---|---|---|
| PDB code | 7I90 | 7I97 | 7I92 |
| Ligand | **7** | **8** | **9** |
| **Data Collection:** | | | |
| Beamline | DIAMOND BEAMLINE i04-1 | DIAMOND BEAMLINE i03 | DIAMOND BEAMLINE i03 |
| Wavelength (Å) | 0.9762 | 0.9762 | 0.9763 |
| Resolution range (Å) | 56.64 - 1.99 (2.03 - 1.99) | 47.11 - 1.95 (1.98 - 1.95) | 54.88 - 2.63 (2.68 - 2.63) |
| Space group | P 1 $2_1$ 1 | P 1 $2_1$ 1 | P 1 $2_1$ 1 |
| Cell (a b c) (Å) | 68.96 67.06 79.92 | 68.94 67.21 79.72 | 66.31 65.85 78.91 |
| Cell (α β γ) (°) | 90.00 100.36 90.00 | 90.00 100.35 90.00 | 90.00 99.93 90.00 |
| Total reflections | 338150 (16487) | 95286 (4578) | 139076 (6593) |
| Unique reflections | 49146 (2382) | 38801 (1939) | 19995 (924) |
| Multiplicity | 6.9 (6.9) | 2.5 (2.4) | 7.0 (7.1) |
| Completeness (%) | 99.9 (97.4) | 75.7 (76.4) | 99.8 (94.4) |
| Mean I/sigma(I) | 3.5 (0.1) | 5.8 (0.3) | 4.3 (0.4) |
| R-merge | 0.184 (5.77) | 0.095 (2.44) | 0.420 (3.75) |
| R-pim | 0.076 (2.35) | 0.068 (1.80) | 0.171 (1.50) |
| CC-half | 0.990 (0.33) | 0.990 (0.12) | 0.978 (0.35) |
| **Refinement:** | | | |
| Refinement resolution range (Å) | 37.02 - 1.99 (2.12 - 1.99) | 56.57 - 1.69 (1.71 - 1.69) | 54.88 - 2.63 (2.65 - 2.63) |
| No. reflections | 36799 (736) | 77676 (1554) | 19946 (416) |
| No. reflections (Rfree) | 1800 (36) | 3802 (75) | 967 (25) |
| R-factor | 0.240 (0.538) | 0.249 (0.466) | 0.273 (0.450) |
| Rfree | 0.314 (0.508) | 0.291 (0.505) | 0.363 (0.472) |
| Number of total atoms | 5613 | 5603 | 5548 |
| atoms for macromolecules | 5272 | 5225 | 5352 |
| atoms for ligands | 83 | 83 | 100 |
| atoms for waters | 258 | 295 | 96 |
| Average B-factor ($Å^2$) | 71.5 | 44.5 | 70.1 |
| RMS(bonds) (Å) | 0.009 | 0.011 | 0.007 |
| RMS(bond angles) (°) | 1.2 | 1.25 | 0.98 |
| RMS(dihedral angles) (°) | 3.06 | 3.73 | 3.14 |

| Protein | BtkS | BtkS | BtkS |
|---|---|---|---|
| PDB code | 7I98 | 9T3M | 9T3G |
| Ligand | **10** | **11** | **12** |
| **Data Collection:** | | | |
| Beamline | DIAMOND BEAMLINE i03 | DIAMOND BEAMLINE i03 | DIAMOND BEAMLINE i04 |
| Wavelength (Å) | 0.9763 | 0.9763 | 0.9762 |
| Resolution range (Å) | 57.05 - 2.07 (2.11 - 2.07) | 67.60 - 1.80 (1.83 - 1.80) | 47.11 - 1.95 (1.98 - 1.95) |
| Space group | $P\ 1\ 2_1\ 1$ | $P\ 1\ 2_1\ 1$ | $P\ 1\ 2_1\ 1$ |
| Cell (a b c) (Å) | 68.98 66.56 79.46 | 68.59 67.07 79.93 | 68.39 66.23 79.56 |
| Cell (α β γ) (°) | 90.00 101.69 90.00 | 90.00 99.77 90.00 | 90.00 101.46 90.00 |
| Total reflections | 215394 (10032) | 464122 (23522) | 95286 (4578) |
| Unique reflections | 43163 (2112) | 66443 (3312) | 38801 (1939) |
| Multiplicity | 5.0 (4.8) | 7.0 (7.1) | 2.5 (2.4) |
| Completeness (%) | 100.0 (99.5) | 100.0 (99.4) | 75.7 (76.4) |
| Mean I/sigma(I) | 10.9 (0.5) | 9.0 (0.2) | 5.8 (0.3) |
| R-merge | 0.122 (2.47) | 0.130 (4.41) | 0.095 (2.44) |
| R-pim | 0.061 (1.28) | 0.053 (1.76) | 0.068 (1.80) |
| CC-half | 0.993 (0.24) | 0.998 (0.27) | 0.990 (0.12) |
| **Refinement:** | | | |
| Refinement resolution range (Å) | 30.12 - 2.07 (2.08 - 2.07) | 67.60 - 1.80 (1.82 - 1.80) | 47.11 - 1.95 (1.97 - 1.95) |
| No. reflections | 42853 (858) | 65010 (1301) | 36201 (725) |
| No. reflections (Rfree) | 2135 (45) | 3277 (154) | 1724 (81) |
| R-factor | 0.249 (0.387) | 0.245 (0.570) | 0.261 (0.477) |
| Rfree | 0.303 (0.365) | 0.298 (0.566) | 0.334 (0.591) |
| Number of total atoms | 5592 | 5471 | 5265 |
| atoms for macromolecules | 5296 | 5113 | 5090 |
| atoms for ligands | 88 | 47 | 77 |
| atoms for waters | 208 | 311 | 98 |
| Average B-factor ($Å^2$) | 58.9 | 47.1 | 57.1 |
| RMS(bonds) (Å) | 0.008 | 0.009 | 0.009 |
| RMS(bond angles) (°) | 1.02 | 1.05 | 1.07 |
| RMS(dihedral angles) (°) | 3.48 | 3.73 | 3.44 |

| Protein | BtkS | BtkS | Btk |
|---|---|---|---|
| PDB code | 7I91 | 7I9D | 7I9F |
| Ligand | **13** | **14** | **16** |
| **Data Collection:** | | | |
| Beamline | DIAMOND BEAMLINE i03 | DIAMOND BEAMLINE i03 | DIAMOND BEAMLINE i03 |
| Wavelength (Å) | 0.9763 | 0.9763 | 0.9763 |
| Resolution range (Å) | 57.05 - 1.93 (1.96 - 1.93) | 67.23 - 1.80 (1.83 - 1.80) | 56.14 - 2.04 (2.07 - 2.04) |
| Space group | P 1 $2_1$ 1 | P 1 $2_1$ 1 | P 1 $2_1$ 1 |
| Cell (a b c) (Å) | 68.83 66.42 79.59 | 68.52 66.60 79.63 | 68.58 67.63 79.53 |
| Cell (α β γ) (°) | 90.00 101.76 90.00 | 90.00 101.09 90.00 | 90.00 99.83 90.00 |
| Total reflections | 2560839 (126031) | 452977 (22089) | 320897 (14914) |
| Unique reflections | 52937 (2572) | 65293 (3144) | 45711 (2213) |
| Multiplicity | 48.4 (49.0) | 6.9 (7.0) | 7.0 (6.7) |
| Completeness (%) | 99.9 (96.9) | 99.9 (98.2) | 99.7 (97.8) |
| Mean I/sigma(I) | 5.3 (0.2) | 15.8 (0.3) | 8.2 (0.4) |
| R-merge | 0.421 (9.99) | 0.066 (3.27) | 0.172 (6.63) |
| R-pim | 0.061 (1.57) | 0.027 (1.32) | 0.070 (2.77) |
| CC-half | 0.997 (0.36) | 0.999 (0.34) | 0.997 (0.34) |
| **Refinement:** | | | |
| Refinement resolution range (Å) | 21.94 - 1.93 (1.97 - 1.93) | 67.23 - 1.80 (1.81 - 1.80) | 56.14 - 2.04 (2.06 - 2.04) |
| No. reflections | 49641 (993) | 64777 (1296) | 43795 (876) |
| No. reflections (Rfree) | 2502 (45) | 3271 (65) | 2165 (41) |
| R-factor | 0.255 (0.572) | 0.245 (0.442) | 0.263 (0.450) |
| Rfree | 0.322 (0.542) | 0.297 (0.449) | 0.323 (0.420) |
| Number of total atoms | 5546 | 5544 | 5591 |
| atoms for macromolecules | 5289 | 5274 | 5352 |
| atoms for ligands | 57 | 65 | 84 |
| atoms for waters | 200 | 205 | 155 |
| Average B-factor ($Å^2$) | 62.2 | 56.1 | 73.3 |
| RMS(bonds) (Å) | 0.009 | 0.009 | 0.008 |
| RMS(bond angles) (°) | 1.03 | 1.01 | 1.04 |
| RMS(dihedral angles) (°) | 3.57 | 3.76 | 3.46 |

| Protein | BtkS | BtkWT | BtkS |
|---|---|---|---|
| PDB code | 9T1V | 7I9E | 9T0T |
| Ligand | **17** | **18** | **19** |
| **Data Collection:** | | | |
| Beamline | DIAMOND BEAMLINE i03 | DIAMOND BEAMLINE i03 | DIAMOND BEAMLINE i03 |
| Wavelength (Å) | 0.9763 | 0.9763 | 0.9184 |
| Resolution range (Å) | 47.41 - 1.95 (1.98 -1.95) | 56.41 - 1.91 (1.94 - 1.91) | 50.93 - 1.73 (1.76 - 1.73) |
| Space group | P 1 $2_1$ 1 | P 1 $2_1$ 1 | P 1 $2_1$ 1 |
| Cell (a b c) (Å) | 66.82 66.28 79.26 | 68.75 67.32 79.56 | 68.99 66.66 79.86 |
| Cell (α β γ) (°) | 90.00 99.95 90.00 | 90.00 100.28 90.00 | 90.00 98.68 90.00 |
| Total reflections | 95286 (4769) | 388145 (19500) | 523123 (26377) |
| Unique reflections | 38802 (1939) | 55583 (2738) | 74878 (3698) |
| Multiplicity | 2.5 (2.4) | 7.0 (7.1) | 7.0 (7.1) |
| Completeness (%) | 75.7 (76.0) | 99.9 (98.5) | 100.0 (99.7) |
| Mean I/sigma(I) | 5.8 (0.3) | 7.6 (0.2) | 13.2 (0.2) |
| R-merge | 0.095 (2.43) | 0.147 (5.29) | 0.080 (4.95) |
| R-pim | 0.068 (1.81) | 0.060 (2.12) | 0.033 (1.99) |
| CC-half | 0.99 (0.12) | 0.998 (0.27) | 0.998 (0.32) |
| **Refinement:** | | | |
| Refinement resolution range (Å) | 55.24 - 2.41 (2.43 - 2.41) | 56.41 - 1.91 (1.96 - 1.91) | 24.66 - 1.73 (1.76 - 1.73) |
| No. reflections | 26468 (530) | 51914 (1039) | 71430 (1429) |
| No. reflections (Rfree) | 1351 (63) | 2602 (50) | 1970 (92) |
| R-factor | 0.263 (0.433) | 0.266 (0.530) | 0.248 (0.513) |
| Rfree | 0.329 (0.454) | 0.318 (0.540) | 0.301 (0.560) |
| Number of total atoms | 5535 | 5579 | 5981 |
| atoms for macromolecules | 5316 | 5344 | 5561 |
| atoms for ligands | 80 | 66 | 128 |
| atoms for waters | 139 | 169 | 292 |
| Average B-factor (Å$^2$) | 61.2 | 63.3 | 55.3 |
| RMS(bonds) (Å) | 0.007 | 0.009 | 0.009 |
| RMS(bond angles) (°) | 0.98 | 1.01 | 1.03 |
| RMS(dihedral angles) (°) | 2.99 | 3.54 | 3.58 |

| Protein | Btk | Btk | BtkA |
|---|---|---|---|
| PDB code | 7I99 | 7I9B | 7I95 |
| Ligand | **22** | **24** | **25** |
| **Data Collection:** | | | |
| Beamline | DIAMOND BEAMLINE i03 | DIAMOND BEAMLINE i03 | DIAMOND BEAMLINE i03 |
| Wavelength (Å) | 0.9763 | 0.9763 | 0.9763 |
| Resolution range (Å) | 56.37 - 1.97 (2.00 - 1.97) | 56.24 - 2.11 (2.15 - 2.11) | 56.32 - 1.99 (2.02 - 1.99) |
| Space group | $P\ 1\ 2_1\ 1$ | $P\ 1\ 2_1\ 1$ | $P\ 1\ 2_1\ 1$ |
| Cell (a b c) (Å) | 68.51 67.51 79.46 | 68.34 66.38 79.70 | 67.96 66.11 79.63 |
| Cell (α β γ) (°) | 90.00 100.55 90.00 | 90.00 100.25 90.00 | 90.00 100.97 90.00 |
| Total reflections | 352128 (18044) | 284078 (14307) | 332943 (16493) |
| Unique reflections | 50397 (2504) | 40587 (1998) | 47744 (2319) |
| Multiplicity | 7.0 (7.2) | 7.0 (7.2) | 7.0 (7.1) |
| Completeness (%) | 99.6 (98.3) | 99.9 (99.3) | 100.0 (98.8) |
| Mean I/sigma(I) | 8.7 (0.3) | 10.3 (0.4) | 6.7 (0.3) |
| R-merge | 0.130 (6.34) | 0.123 (2.71) | 0.152 (3.65) |
| R-pim | 0.053 (2.53) | 0.050 (1.09) | 0.062 (1.47) |
| CC-half | 0.998 (0.36) | 0.998 (0.31) | 0.997 (0.42) |
| **Refinement:** | | | |
| Refinement resolution range (Å) | 56.37 - 1.97 (2.00 - 1.97) | 30.56 - 2.11 (2.12 - 2.11) | 56.32 - 1.99 (2.02 - 1.99) |
| No. reflections | 48289 (966) | 40484 (810) | 46051 (922) |
| No. reflections (Rfree) | 2339 (39) | 2029 (37) | 2004 (40) |
| R-factor | 0.263 (0.452) | 0.262 (0.437) | 0.254 (0.510) |
| Rfree | 0.316 (0.520) | 0.333 (0.573) | 0.316 (0.486) |
| Number of total atoms | 5632 | 5768 | 5851 |
| atoms for macromolecules | 5373 | 5552 | 5552 |
| atoms for ligands | 80 | 76 | 92 |
| atoms for waters | 179 | 140 | 207 |
| Average B-factor ($Å^2$) | 66.3 | 69.1 | 61.3 |
| RMS(bonds) (Å) | 0.008 | 0.008 | 0.008 |
| RMS(bond angles) (°) | 1.09 | 1.01 | 1 |
| RMS(dihedral angles) (°) | 3.58 | 3.3 | 3.41 |

| Protein | Btk | BtkS | Btk |
|---|---|---|---|
| PDB code | 7I9H | 7I93 | 9RL9 |
| Ligand | **26** | **27** | **28** |
| **Data Collection:** | | | |
| Beamline | DIAMOND BEAMLINE i04 | DIAMOND BEAMLINE i04 | DIAMOND BEAMLINE i03 |
| Wavelength (Å) | 0.9537 | 0.9537 | 0.9763 |
| Resolution range (Å) | 56.38 - 2.59 (2.63 - 2.59) | 56.58 - 1.91 (1.94 - 1.91) | 56.24 - 2.11 (2.15 - 2.11) |
| Space group | $P\ 1\ 2_1\ 1$ | $P\ 1\ 2_1\ 1$ | $P\ 1\ 2_1\ 1$ |
| Cell (a b c) (Å) | 68.26 66.72 79.62 | 68.48 67.15 79.82 | 68.34 66.38 79.70 |
| Cell (α β γ) (°) | 90.00 100.76 90.00 | 90.00 100.86 90.00 | 90.00 100.25 90.00 |
| Total reflections | 153878 (7760) | 384433 (18903) | 284078 (14307) |
| Unique reflections | 22122 (1107) | 54217 (2626) | 40587 (1998) |
| Multiplicity | 7.0 (7.0) | 7.1 (7.2) | 7.0 (7.2) |
| Completeness (%) | 100.0 (99.1) | 98.0 (96.3) | 99.9 (99.3) |
| Mean I/sigma(I) | 7.8 (0.4) | 9.6 (0.2) | 10.3 (0.4) |
| R-merge | 0.189 (2.73) | 0.115 (4.55) | 0.123 (2.71) |
| R-pim | 0.077 (1.11) | 0.046 (1.83) | 0.050 (1.09) |
| CC-half | 0.997 (0.28) | 0.998 (0.38) | 0.998 (0.31) |
| **Refinement:** | | | |
| Refinement resolution range (Å) | 56.38 - 2.59 (2.61 - 2.59) | 27.29 - 1.91 (1.93 - 1.91) | 30.56 - 2.11 (2.12 - 2.11) |
| No. reflections | 22087 (442) | 52008 (1041) | 40484 (810) |
| No. reflections (Rfree) | 1175 (18) | 2603 (61) | 2029 (37) |
| R-factor | 0.251 (0.398) | 0.258 (0.589) | 0.262 (0.437) |
| Rfree | 0.343 (0.364) | 0.317 (0.614) | 0.333 (0.573) |
| Number of total atoms | 5459 | 5516 | 5768 |
| atoms for macromolecules | 5282 | 5186 | 5552 |
| atoms for ligands | 24 | 88 | 76 |
| atoms for waters | 153 | 242 | 140 |
| Average B-factor ($Å^2$) | 79.9 | 61.2 | 69.1 |
| RMS(bonds) (Å) | 0.007 | 0.008 | 0.008 |
| RMS(bond angles) (°) | 0.89 | 1.01 | 1.01 |
| RMS(dihedral angles) (°) | 2.68 | 3.42 | 3.3 |

| Protein | Btk | BtkS | BtkA |
|---|---|---|---|
| PDB code | 7I9G | 9T21 | 7I9C |
| Ligand | **28** | **29** | **30** |
| **Data Collection:** | | | |
| Beamline | DIAMOND BEAMLINE i03 | DIAMOND BEAMLINE i04 | DIAMOND BEAMLINE i03 |
| Wavelength (Å) | 0.9763 | 0.9537 | 0.9188 |
| Resolution range (Å) | 57.74 - 1.70 (1.73 - 1.70) | 47.99 - 1.87 (1.90 - 1.87) | 56.40 - 1.92 (1.950- 1.92) |
| Space group | P 1 $2_1$ 1 | P 1 $2_1$ 1 | P 1 $2_1$ 1 |
| Cell (a b c) (Å) | 69.63 66.43 79.46 | 69.29 67.57 79.81 | 68.15 66.36 79.62 |
| Cell (α β γ) (°) | 90.00 102.63 90.00 | 90.00 100.28 90.00 | 90.00 100.95 90.00 |
| Total reflections | 542465 (26474) | 417902 (20600) | 362968 (16508) |
| Unique reflections | 77629 (3701) | 60135 (2930) | 53350 (2635) |
| Multiplicity | 7.0 (7.2) | 6.9 (7.0) | 6.8 (6.3) |
| Completeness (%) | 99.4 (96.0) | 100.0 (98.7) | 99.8 (99.2) |
| Mean I/sigma(I) | 14.2 (0.4) | 8.7 (0.2) | 6.3 (0.2) |
| R-merge | 0.077 (3.21) | 0.138 (14.58) | 0.176 (3.19) |
| R-pim | 0.031 (1.28) | 0.056 (5.93) | 0.074 (1.40) |
| CC-half | 0.999 (0.26) | 0.997 (0.37) | 0.980 (0.25) |
| **Refinement:** | | | |
| Refinement resolution range (Å) | 57.74 - 1.70 (1.71 - 1.70) | 37.00 - 1.87 (1.91 - 1.87) | 22.30 - 1.94 (1.95 - 1.94) |
| No. reflections | 77532 (1551) | 56561 (1132) | 47517 (951) |
| No. reflections (Rfree) | 1997 (54) | 2787 (132) | 1722 (32) |
| R-factor | 0.259 (0.439) | 0.248 (0.496) | 0.304 (0.660) |
| Rfree | 0.285 (0.476) | 0.307 (0.536) | 0.344 (0.711) |
| Number of total atoms | 5836 | 6145 | 5500 |
| atoms for macromolecules | 5552 | 5561 | 5356 |
| atoms for ligands | 21 | 74 | 77 |
| atoms for waters | 263 | 510 | 67 |
| Average B-factor (Å$^2$) | 46.5 | 53.8 | 52.4 |
| RMS(bonds) (Å) | 0.009 | 0.01 | 0.008 |
| RMS(bond angles) (°) | 1.05 | 1.09 | 0.99 |
| RMS(dihedral angles) (°) | 3.91 | 3.41 | 3.2 |

| Protein | BtkA | Btk | BtkA |
|---|---|---|---|
| PDB code | 9RMO | 7I9b | 9RN5 |
| Ligand | **31** | **32** | **33** |
| **Data Collection:** | | | |
| Beamline | DIAMOND BEAMLINE i04-1 | DIAMOND BEAMLINE i04 | DIAMOND BEAMLINE i04 |
| Wavelength (Å) | 0.9762 | 0.9537 | 0.9537 |
| Resolution range (Å) | 56.53 - 1.80 (1.83 - 1.80) | 56.29 - 2.77 (2.82 - 2.77) | 51.00 - 1.83 (1.86 - 1.83) |
| Space group | $P\ 1\ 2_1\ 1$ | $P\ 1\ 2_1\ 1$ | $P\ 1\ 2_1\ 1$ |
| Cell (a b c) (Å) | 68.92 66.88 79.94 | 68.00 66.03 79.69 | 68.98 67.15 79.78 |
| Cell (α β γ) (°) | 90.00 100.11 90.00 | 90.00 100.81 90.00 | 90.00 100.58 90.00 |
| Total reflections | 464304 (23663) | 124445 (6027) | 439862 (22282) |
| Unique reflections | 66462 (3318) | 17891 (879) | 63456 (3134) |
| Multiplicity | 7.0 (7.1) | 7.0 (6.9) | 6.9 (7.1) |
| Completeness (%) | 100.0 (99.8) | 100.0 (99.2) | 99.8 (98.1) |
| Mean I/sigma(I) | 7.8 (0.2) | 6.5 (0.5) | 10.3 (0.3) |
| R-merge | 0.148 (4.59) | 0.263 (2.50) | 0.105 (3.41) |
| R-pim | 0.060 (1.84) | 0.107 (1.03) | 0.043 (1.37) |
| CC-half | 0.997 (0.32) | 0.990 (0.28) | 0.998 (0.57) |
| **Refinement:** | | | |
| Refinement resolution range (Å) | 56.53 - 1.80 (1.83 - 1.80) | 56.29 - 2.77 (2.79 - 2.77) | 51.01 - 1.83 (1.84 - 1.83) |
| No. reflections | 64254 (1286) | 17852 (380) | 62580 (1252) |
| No. reflections (Rfree) | 3187 (74) | 951 (24) | 3105 (58) |
| R-factor | 0.251 (0.508) | 0.243 (0.378) | 0.269 (0.544) |
| Rfree | 0.293 (0.517) | 0.331 (0.371) | 0.322 (0.526) |
| Number of total atoms | 5610 | 5420 | 5441 |
| atoms for macromolecules | 5239 | 5309 | 5188 |
| atoms for ligands | 79 | 75 | 78 |
| atoms for waters | 292 | 36 | 175 |
| Average B-factor ($Å^2$) | 47.5 | 77.8 | 54.1 |
| RMS(bonds) (Å) | 0.009 | 0.007 | 0.009 |
| RMS(bond angles) (°) | 1 | 0.91 | 1.04 |
| RMS(dihedral angles) (°) | 3.66 | 2.72 | 3.56 |

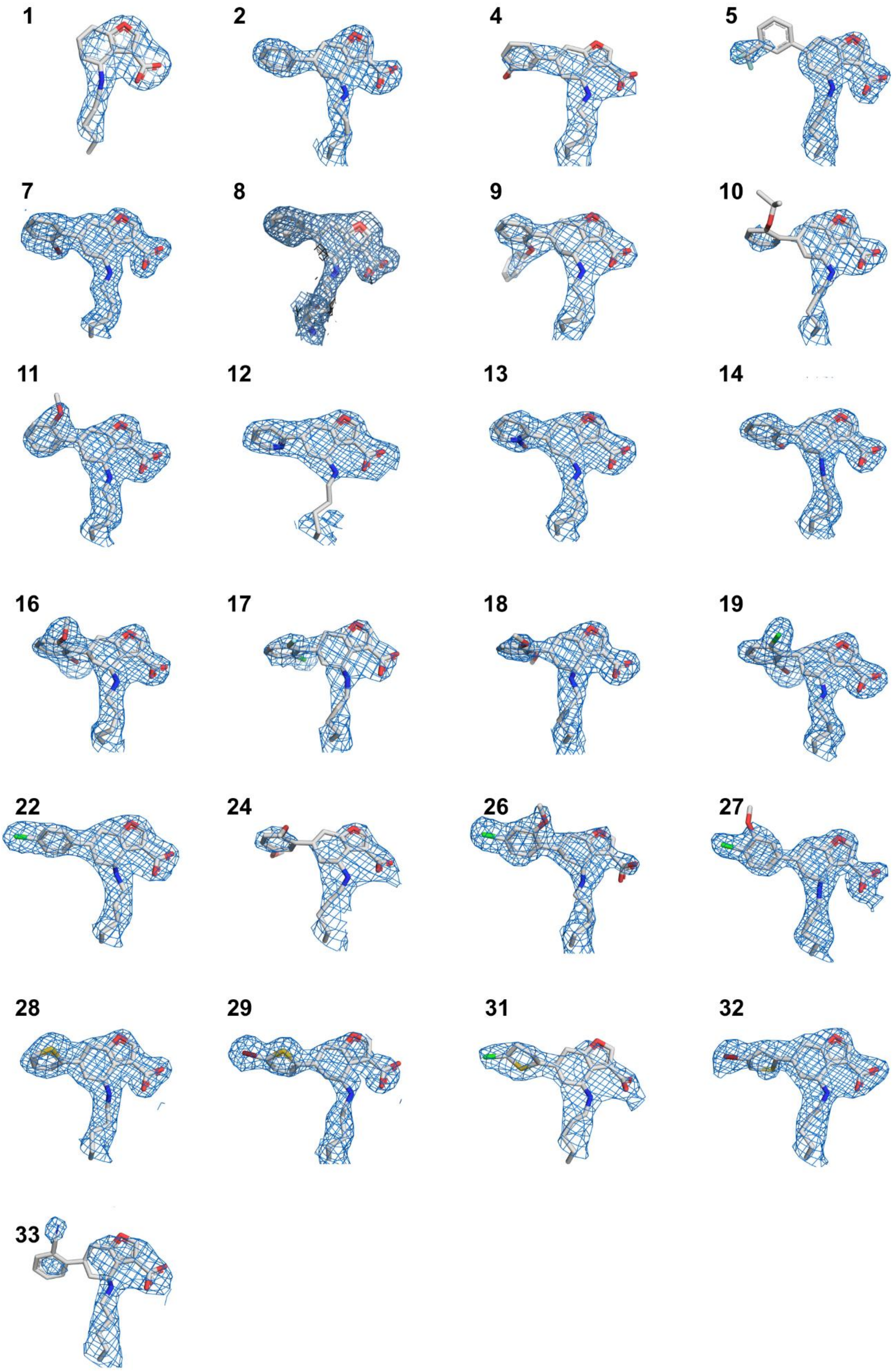

Figure S1. The 2mFo-DFc electron density maps of BTK PH domain binder complexes contoured at 1.0σ before ligand fitting for the crystal structures described.

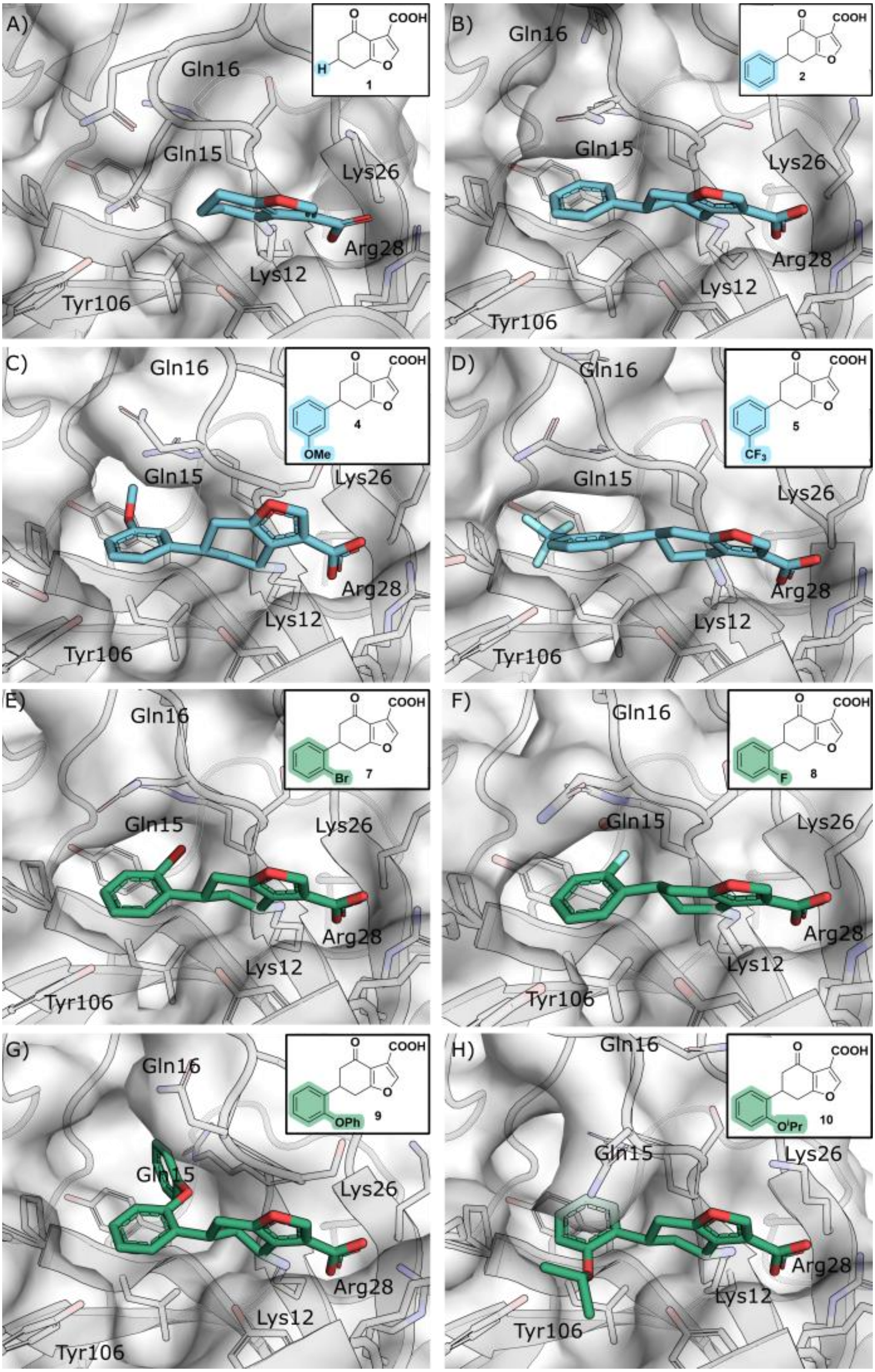

A)
O
COOH
H
O
1
Gln16
Gln15
Lys26
Arg28
Lys12
Tyr106
B)
O
COOH
O
2
Gln16
Gln15
Lys26
Arg28
Lys12
Tyr106
C)
O
COOH
O
4
OMe
Gln16
Gln15
Lys26
Arg28
Lys12
Tyr106
D)
O
COOH
O
5
CF3
Gln16
Gln15
Lys26
Arg28
Lys12
Tyr106
E)
O
COOH
O
Br
7
Gln16
Gln15
Lys26
Arg28
Lys12
Tyr106
F)
O
COOH
O
F
8
Gln16
Gln15
Lys26
Arg28
Lys12
Tyr106
G)
O
COOH
O
OPh
9
Gln16
Gln15
Lys26
Arg28
Lys12
Tyr106
H)
O
COOH
O
OiPr
10
Gln16
Gln15
Lys26
Arg28
Lys12
Tyr106

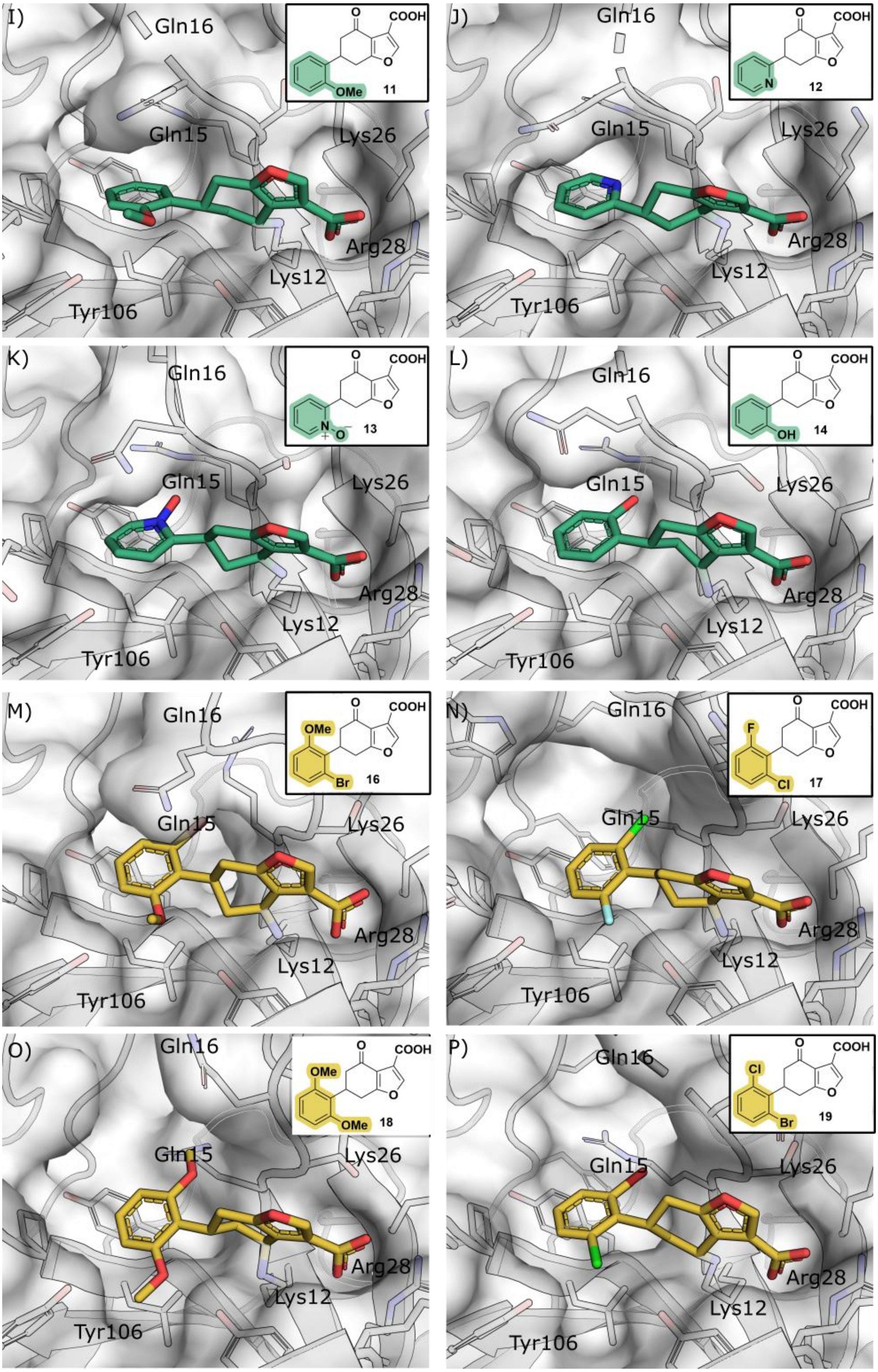
I)
Gln16
Gln15
Lys26
Arg28
Lys12
Tyr106
COOH
O
OMe
11
J)
Gln16
Gln15
Lys26
Arg28
Lys12
Tyr106
COOH
N
12
K)
Gln16
Gln15
Lys26
Arg28
Lys12
Tyr106
COOH
13
L)
Gln16
Gln15
Lys26
Arg28
Lys12
Tyr106
COOH
OH
14
M)
Gln16
Gln15
Lys26
Arg28
Lys12
Tyr106
COOH
OMe
Br
16
N)
Gln16
Gln15
Lys26
Arg28
Lys12
Tyr106
COOH
F
Cl
17
O)
Gln16
Gln15
Lys26
Arg28
Lys12
Tyr106
COOH
OMe
OMe
18
P)
Gln16
Gln15
Lys26
Arg28
Lys12
Tyr106
COOH
Cl
Br
19

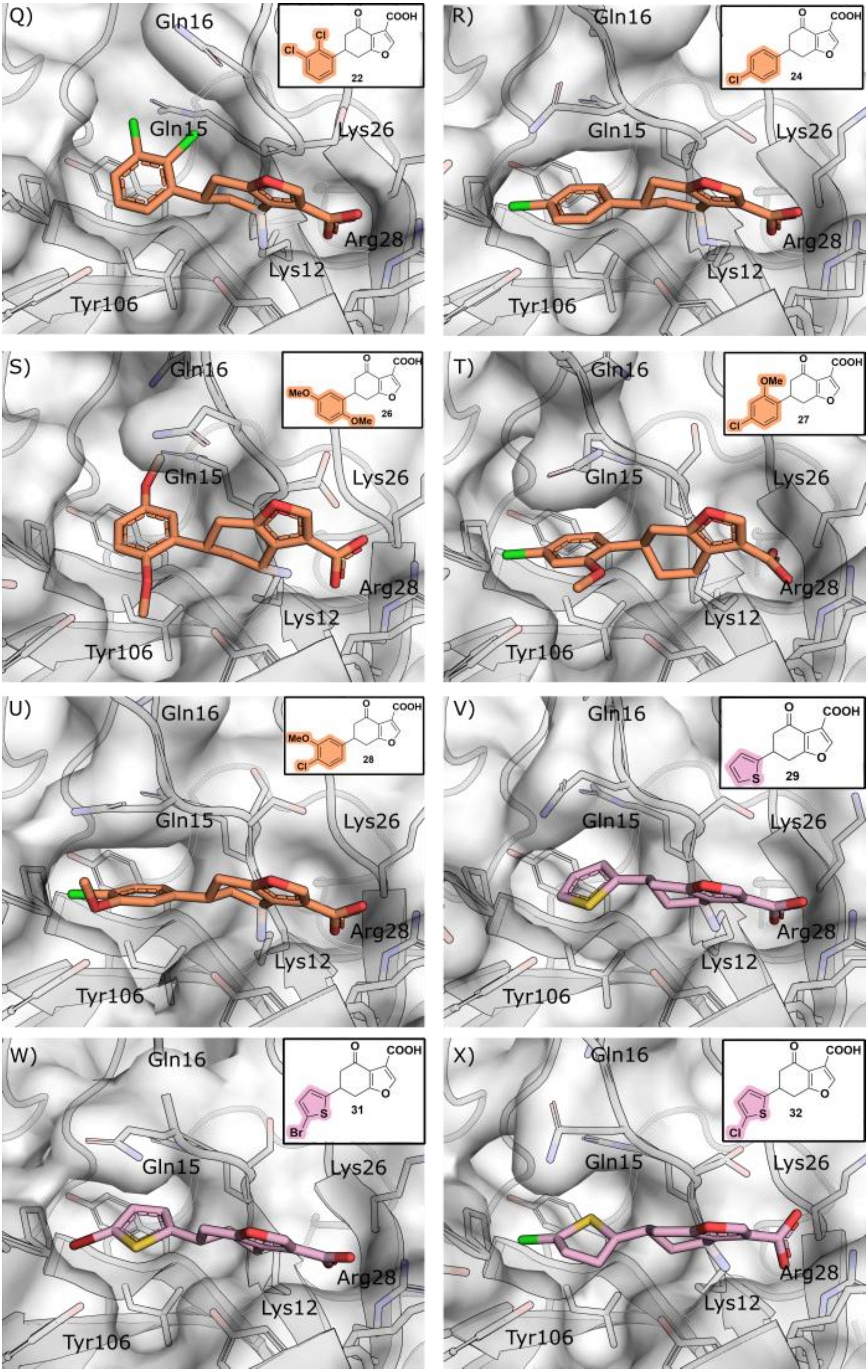

Q)
Gln16
Gln15
Lys26
Arg28
Lys12
Tyr106
O
COOH
Cl
Cl
O
22
R)
Gln16
Gln15
Lys26
Arg28
Lys12
Tyr106
O
COOH
Cl
O
24
S)
Gln16
Gln15
Lys26
Arg28
Lys12
Tyr106
O
COOH
MeO
OMe
O
26
T)
Gln16
Gln15
Lys26
Arg28
Lys12
Tyr106
O
COOH
OMe
Cl
O
27
U)
Gln16
Gln15
Lys26
Arg28
Lys12
Tyr106
O
COOH
MeO
Cl
O
28
V)
Gln16
Gln15
Lys26
Arg28
Lys12
Tyr106
O
COOH
S
O
29
W)
Gln16
Gln15
Lys26
Arg28
Lys12
Tyr106
O
COOH
S
Br
O
31
X)
Gln16
Gln15
Lys26
Arg28
Lys12
Tyr106
O
COOH
S
Cl
O
32

--

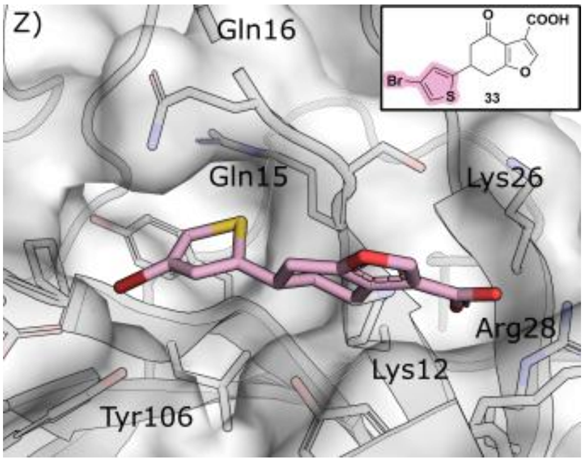

Figure S2. The crystal structures of all the compounds in this study bound to the BTK PH domain. Key residues are highlighted.

### S2 Compounds NMR Spectra

4-oxo-4,5,6,7-tetrahydrobenzofuran-3-carboxylic acid (**1**) $^1$H NMR (400 MHz, $CDCl_3$)

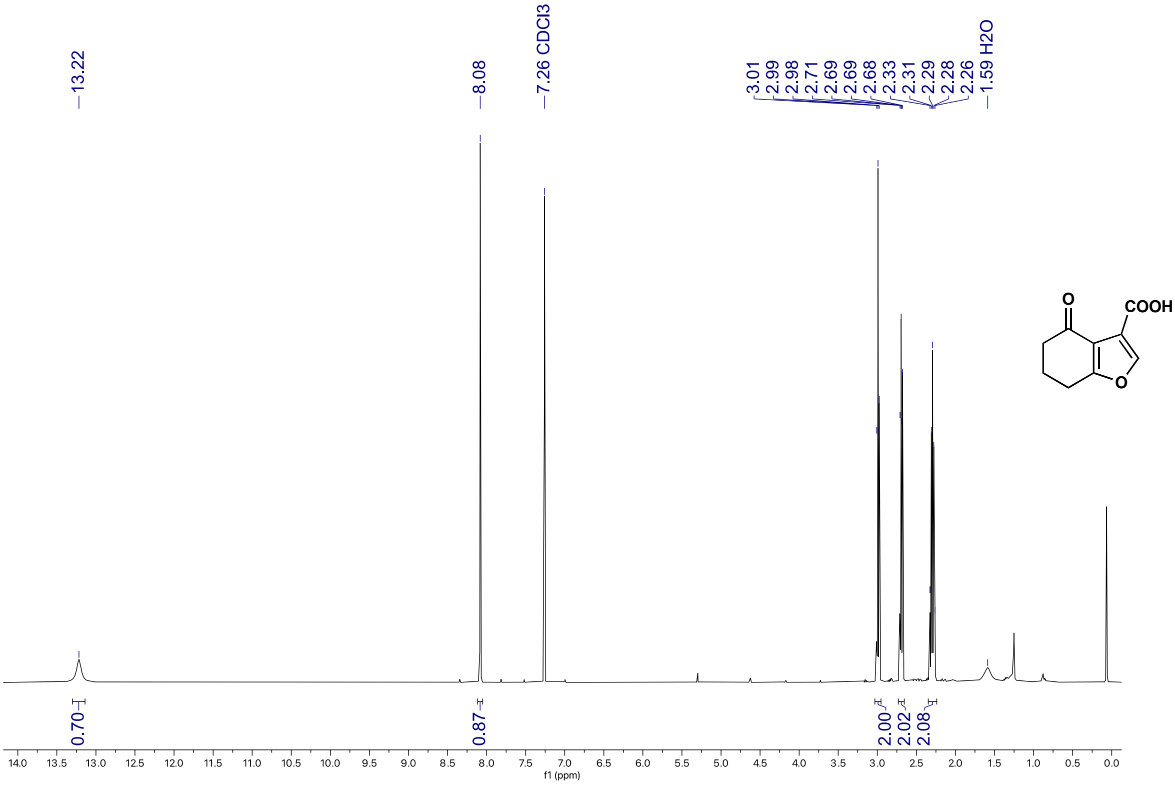

4-oxo-4,5,6,7-tetrahydrobenzofuran-3-carboxylic acid (**1**) $^{13}C$ NMR (101 MHz, $CDCl_3$)

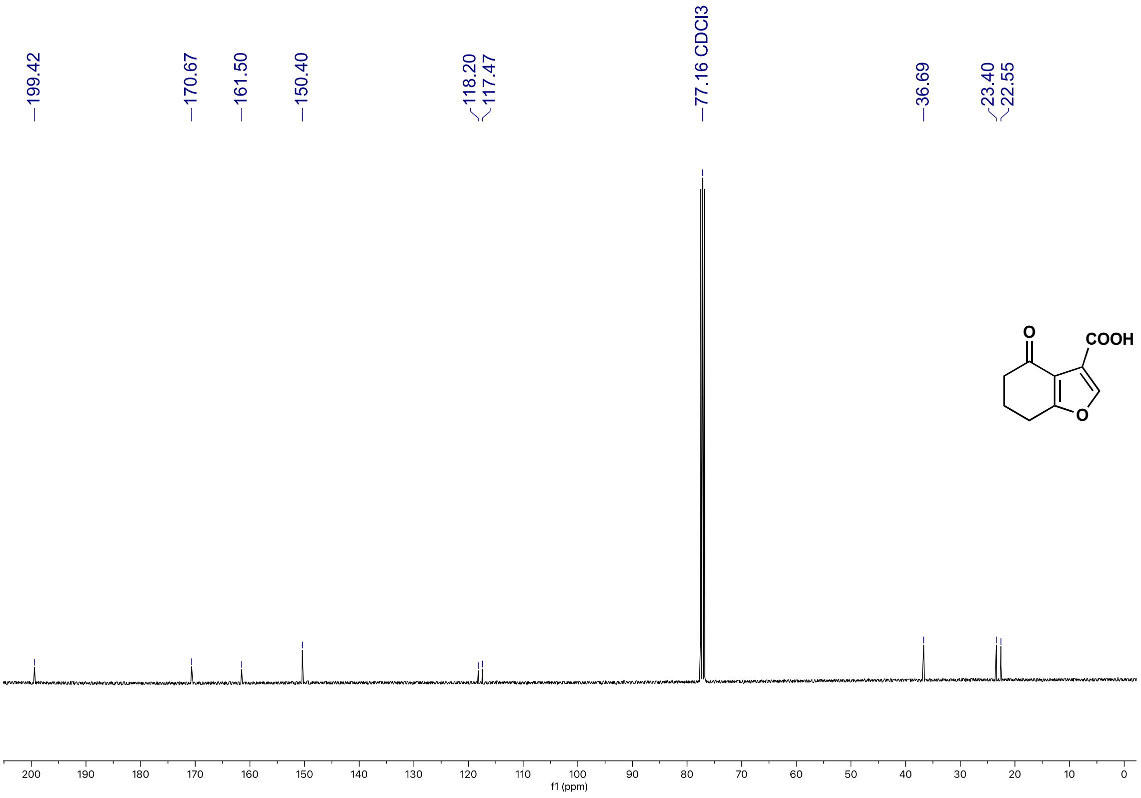

5-phenylcyclohexane-1,3-dione (**2c**) $^{1}H$ NMR (500 MHz, $CD_3OD$)

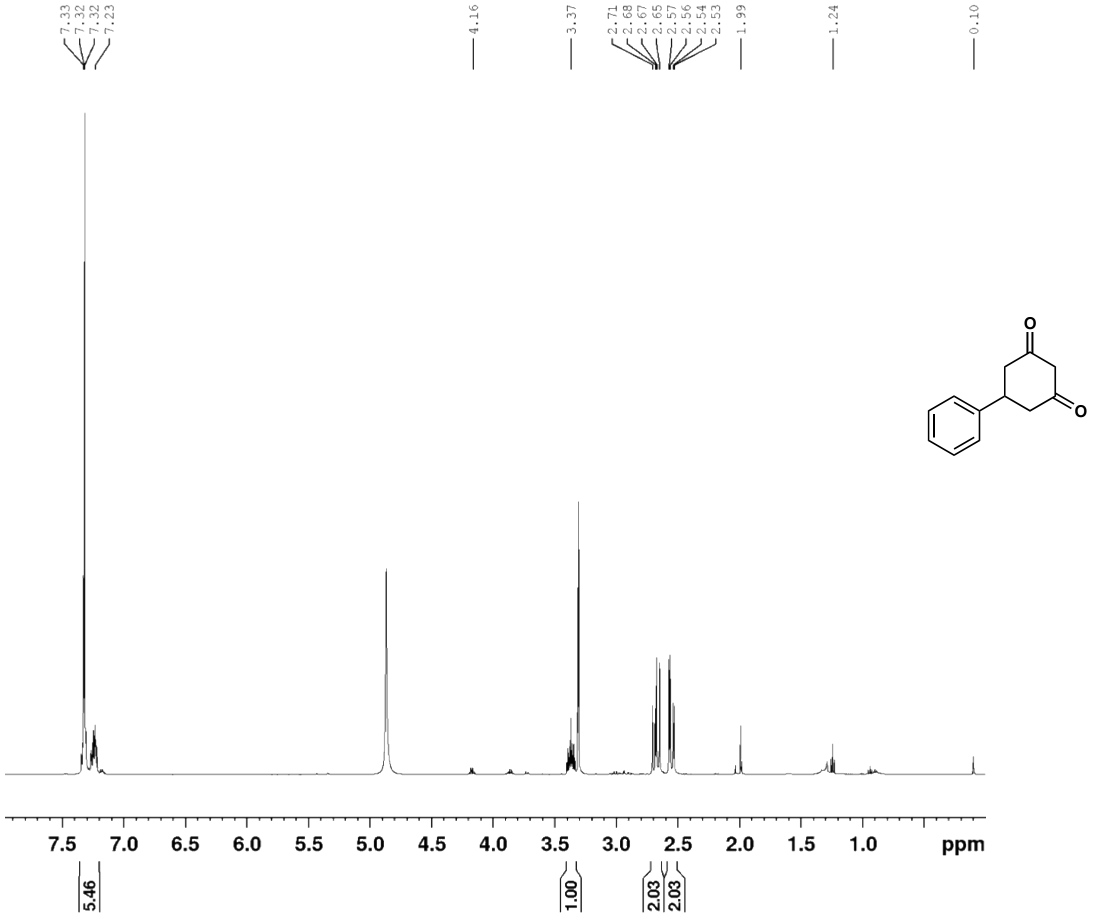

5-phenylcyclohexane-1,3-dione (**2c**) $^{13}C$ NMR (125 MHz, $CD_3OD$)

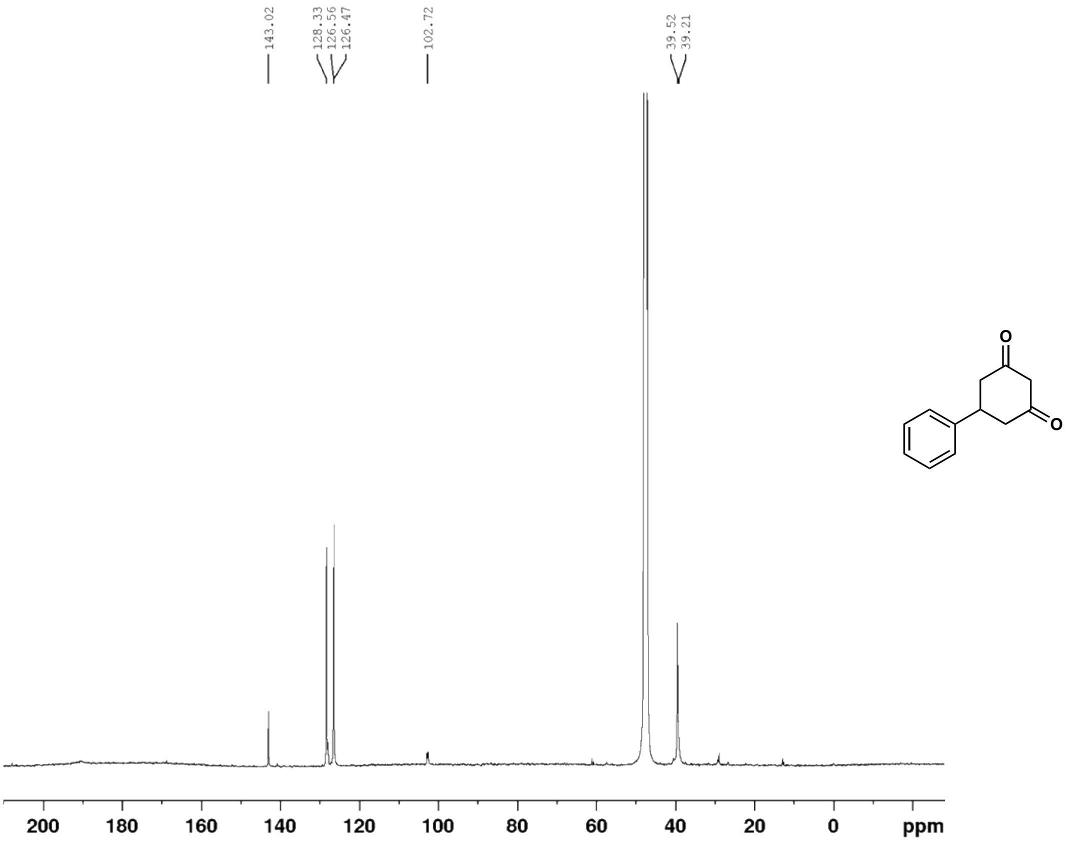

3-carboxy-6-phenyl-4,5,6,7-tetrahydrobenzofuran-4-one (**2**) $^{1}$H NMR (500 MHz, $CDCl_3$)

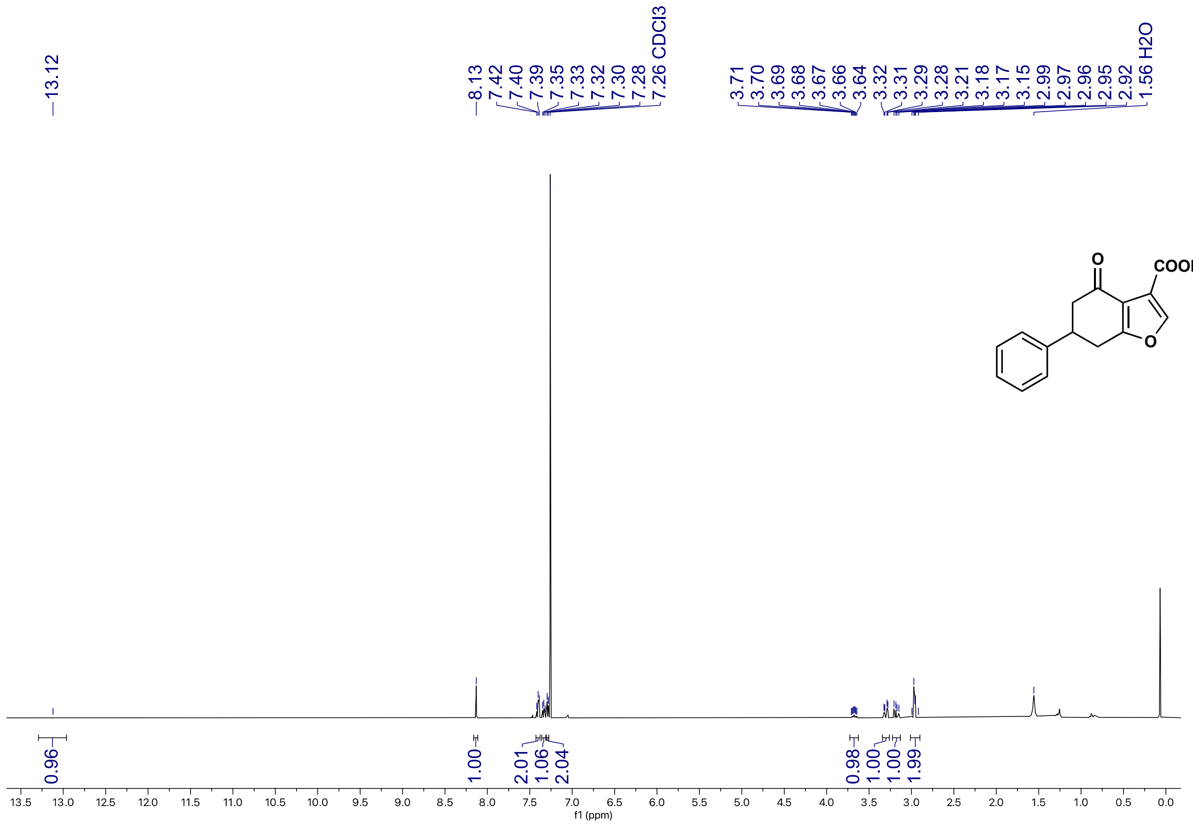

3-carboxy-6-phenyl-4,5,6,7-tetrahydrobenzofuran-4-one (**2**) $^{13}$C NMR (126 MHz, $CDCl_3$)

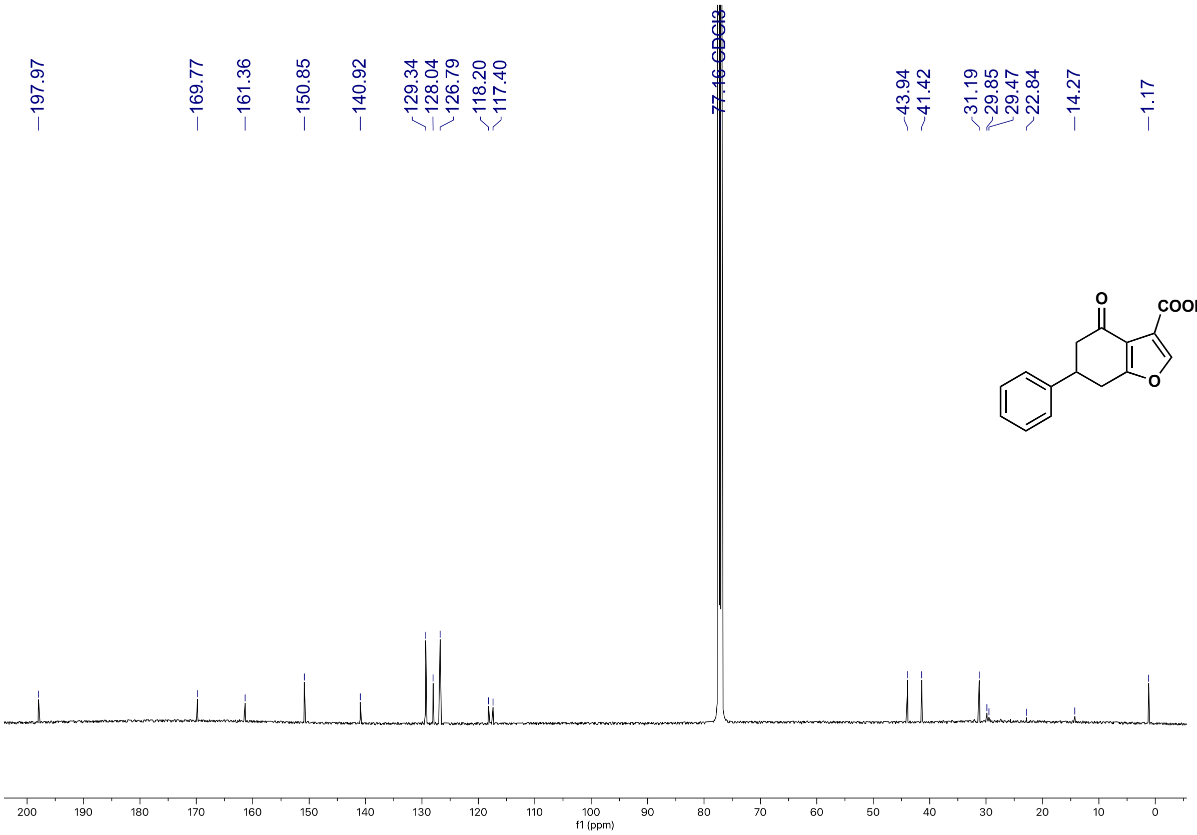

(*E*)-4-(naphthalen-2-yl)but-3-en-2-one (**3b**) $^{1}$H NMR (500 MHz, $CDCl_3$)

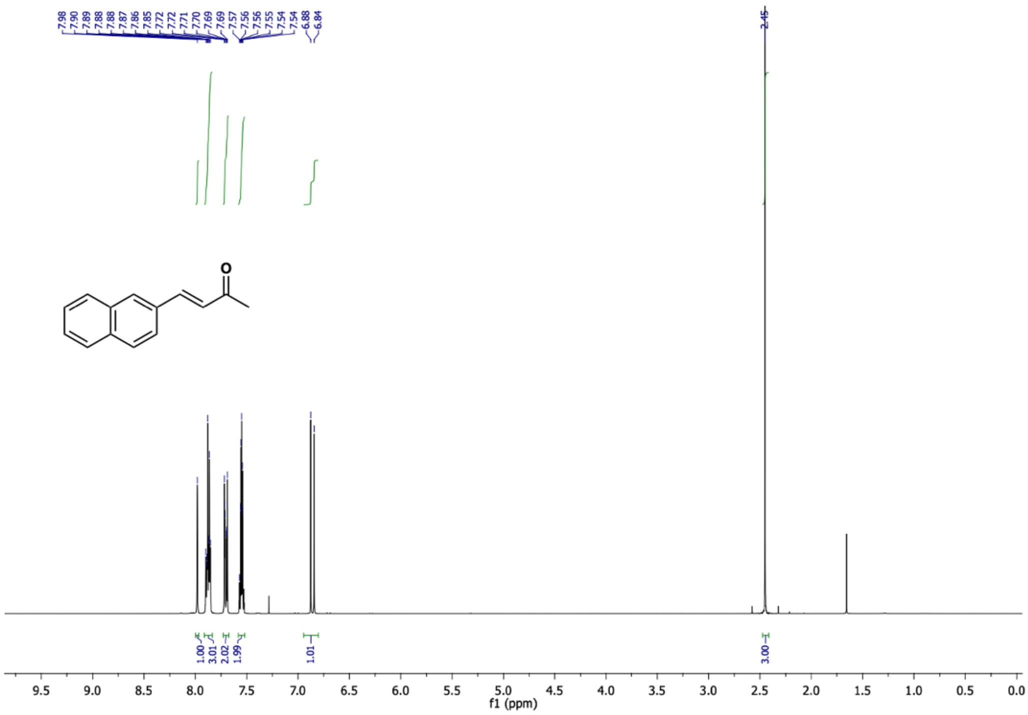

(*E*)-4-(naphthalen-2-yl)but-3-en-2-one (**3b**) $^{13}$C NMR (126 MHz, $CDCl_3$)

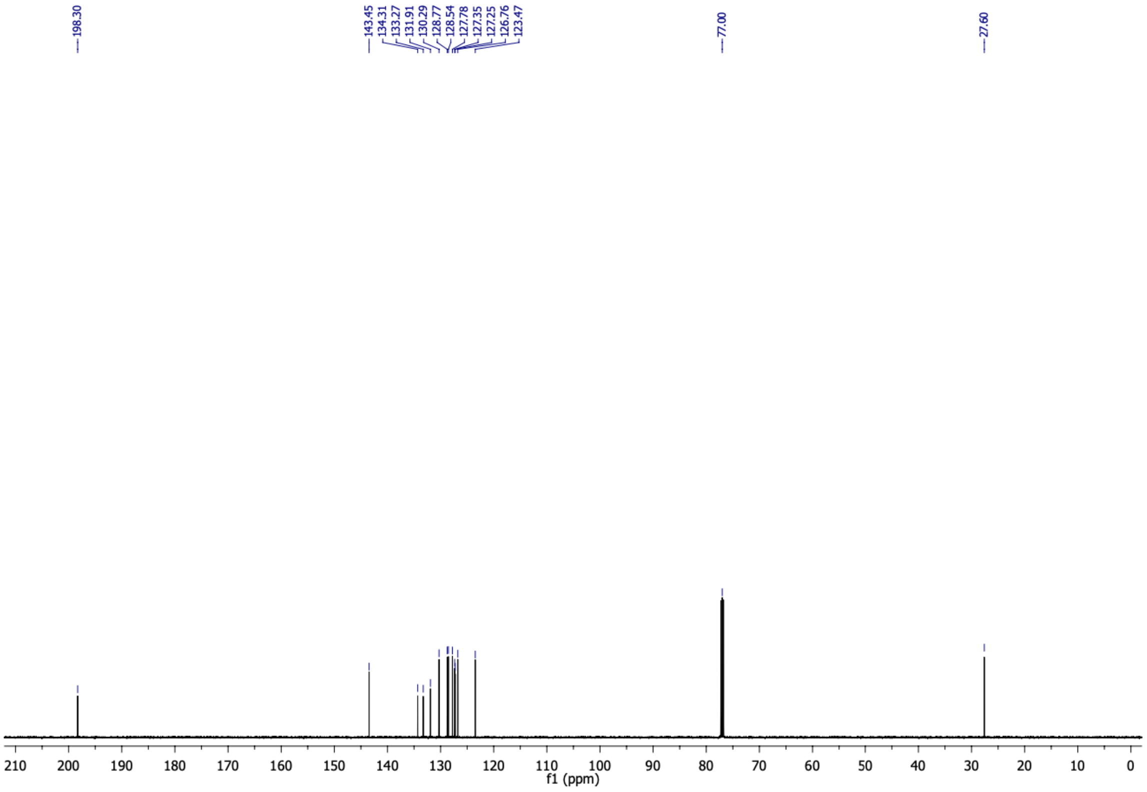

5-(naphthalen-2-yl)cyclohexane-1,3-dione (**3c**) $^{1}H$ NMR (500 MHz, $d_6$-DMSO)

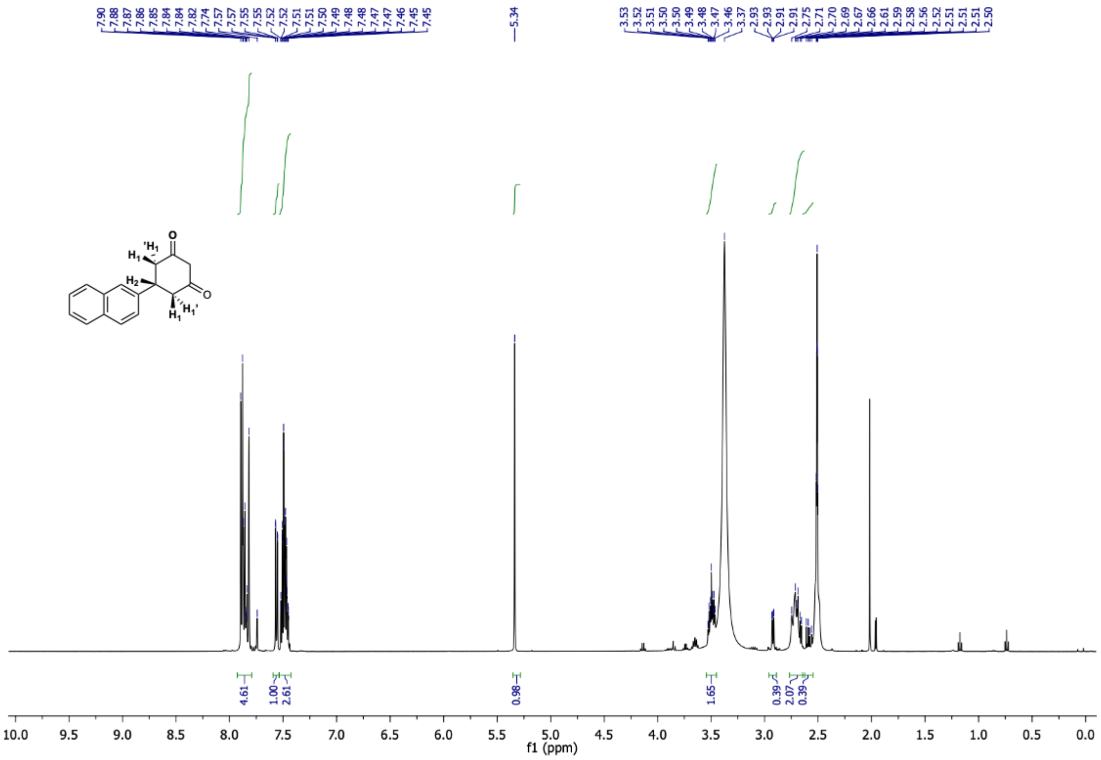

5-(naphthalen-2-yl)cyclohexane-1,3-dione (**3c**) $^{13}C$ NMR (126 MHz, $d_6$-DMSO)

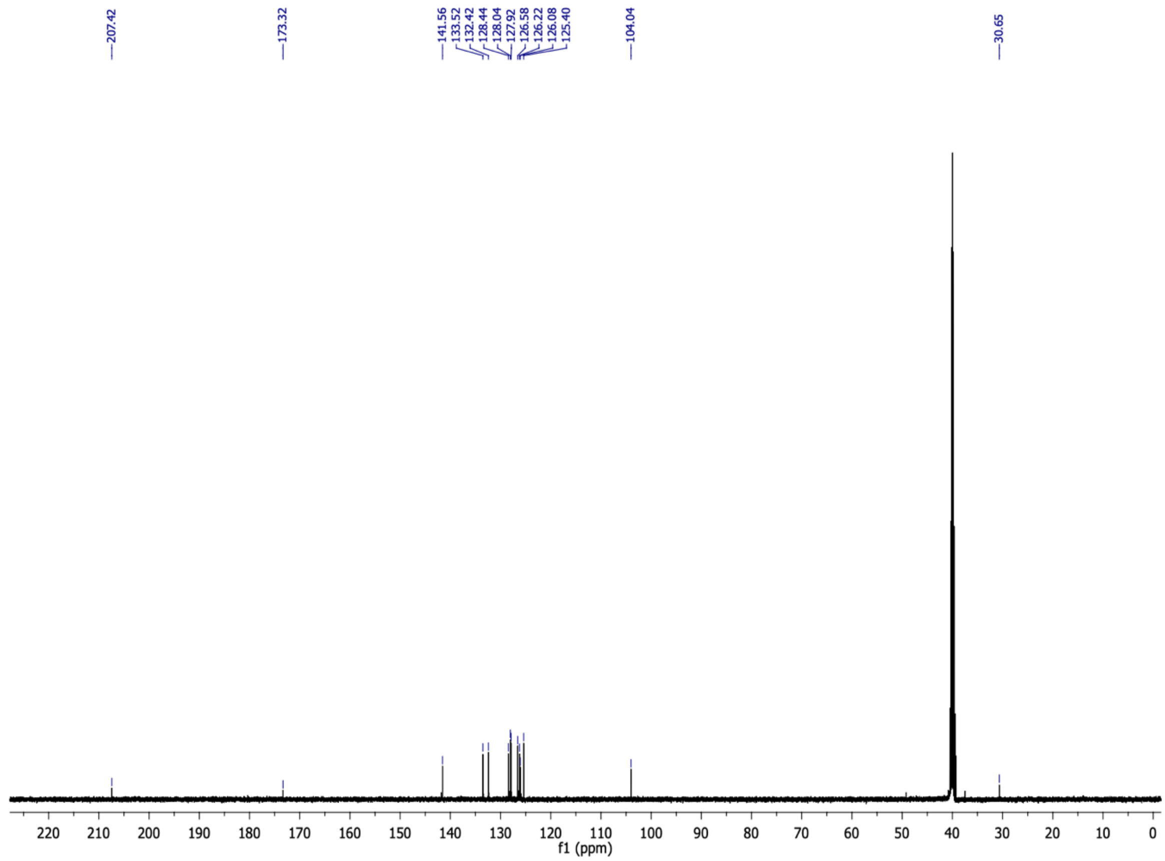

6-(naphthalen-2-yl)-4-oxo-4,5,6,7-tetrahydrobenzofuran-3-carboxylic acid (**3**) $^{1}H$ NMR (500 MHz, $CDCl_3$)

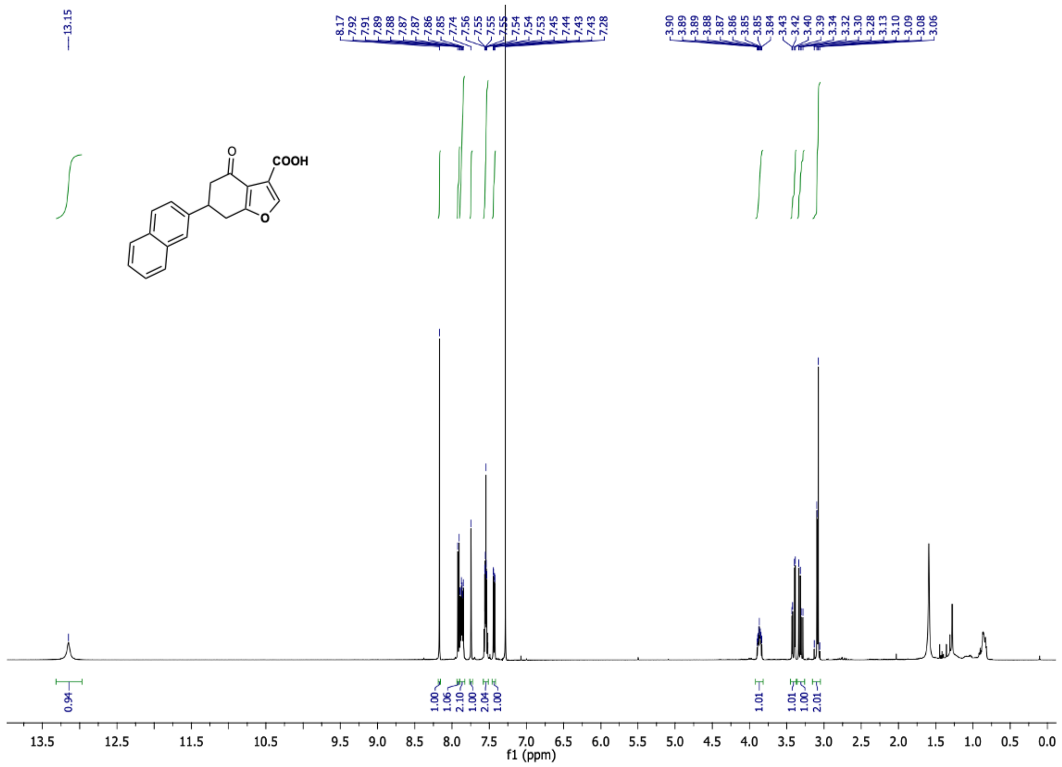

6-(naphthalen-2-yl)-4-oxo-4,5,6,7-tetrahydrobenzofuran-3-carboxylic acid (**3**) $^{13}C$ NMR (126 MHz, $CDCl_3$)

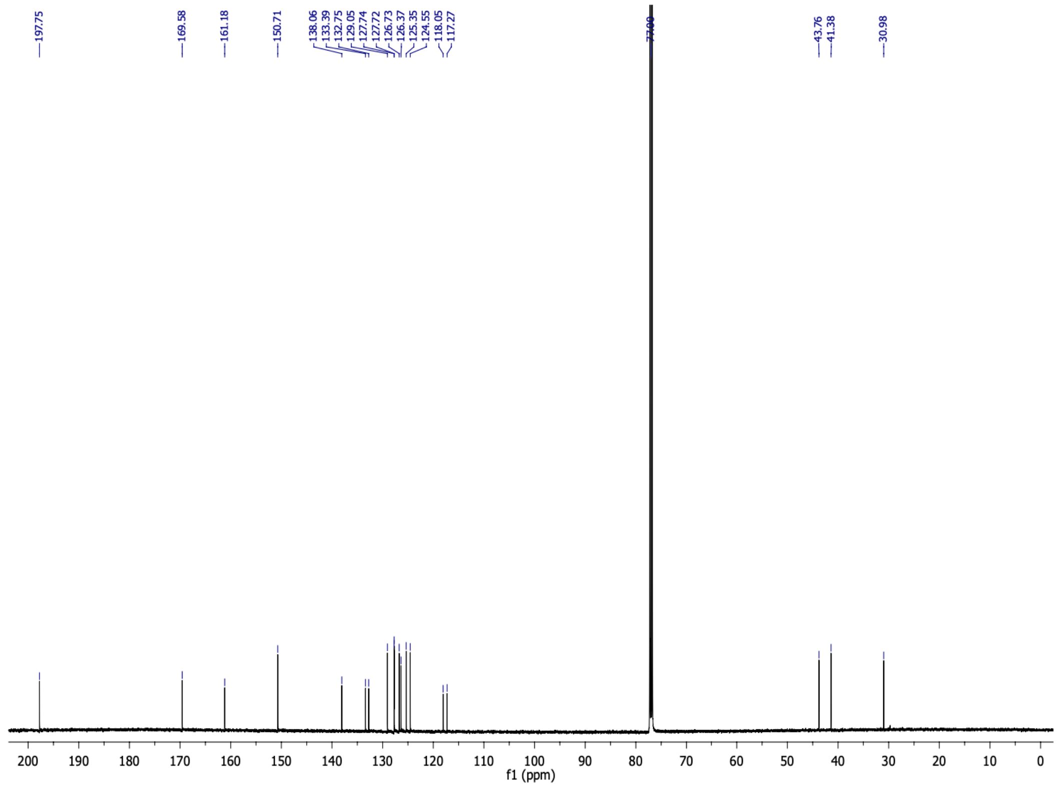

(*E*)-4-(3-methoxyphenyl)but-3-en-2-one (**4b**) $^{1}$H NMR (500 MHz, $CDCl_3$)

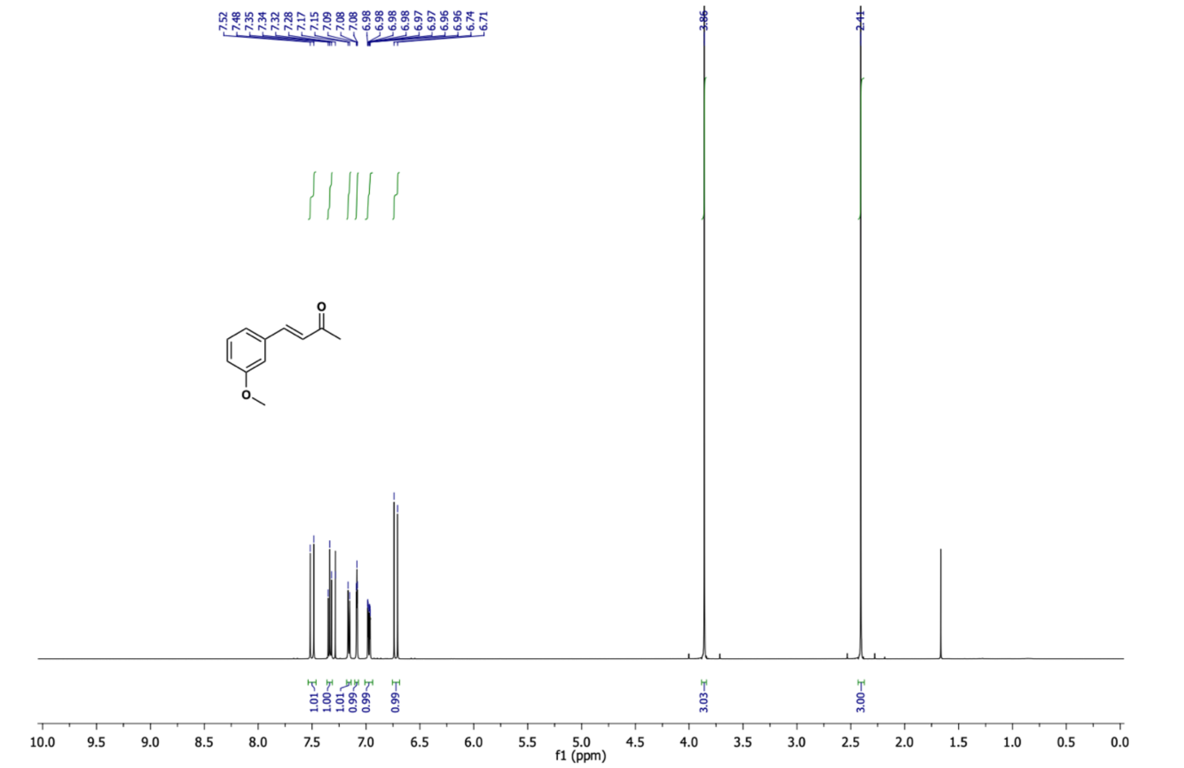

(*E*)-4-(3-methoxyphenyl)but-3-en-2-one (**4b**) $^{13}$C NMR (126 MHz, $CDCl_3$)

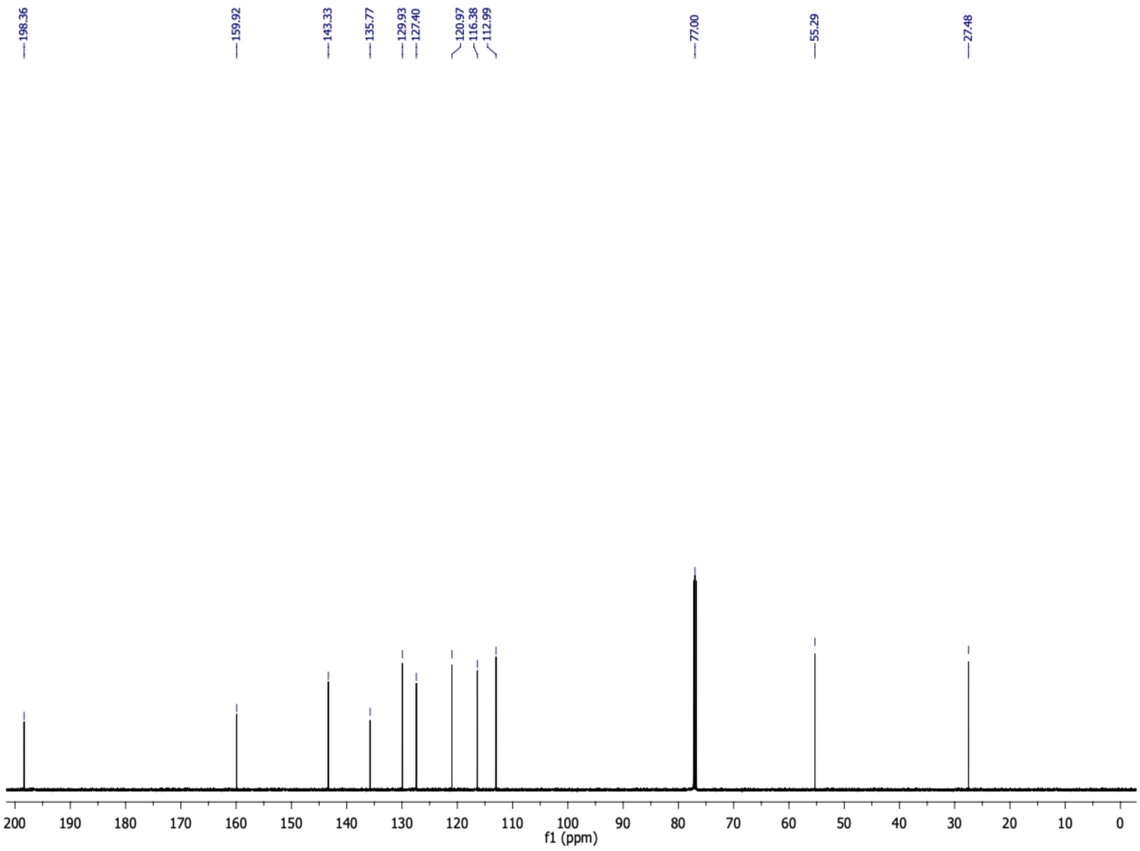

5-(3-methoxyphenyl)cyclohexane-1,3-dione (**4c**) $^{1}H$ NMR (500 MHz, $CD_3CN$)

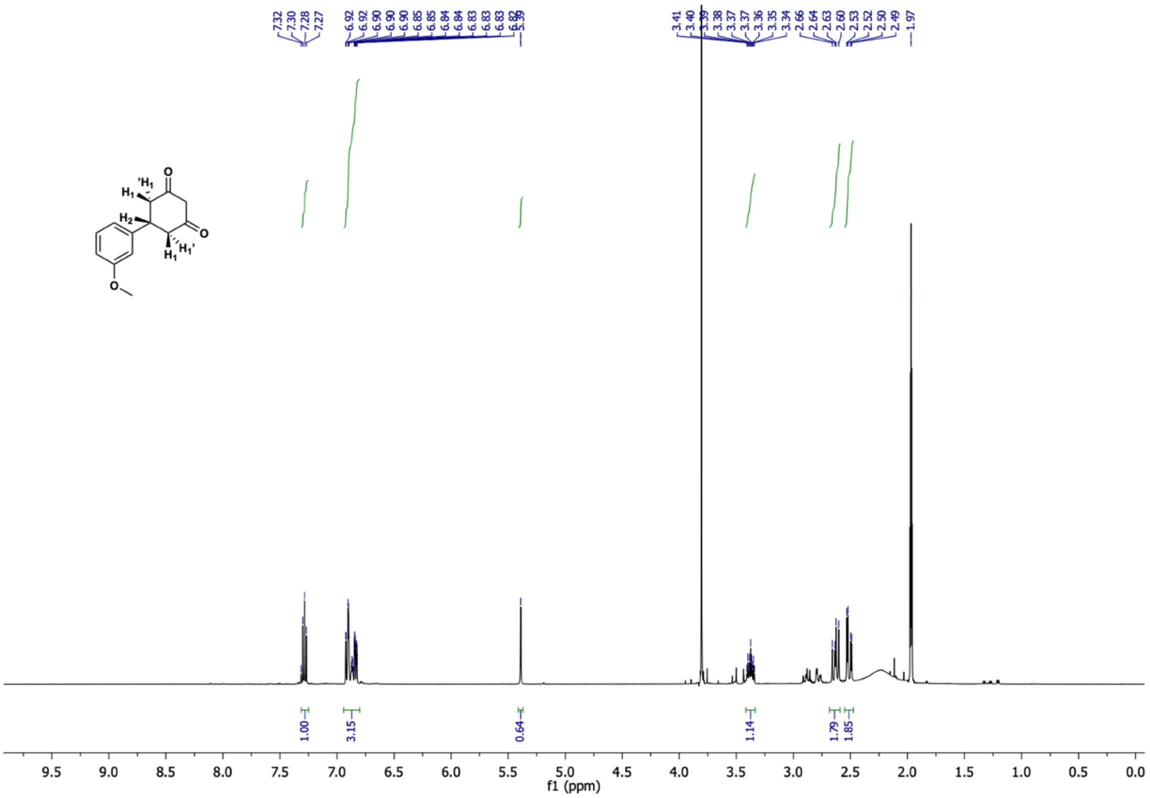

5-(3-methoxyphenyl)cyclohexane-1,3-dione (**4c**) $^{13}C$ NMR (126 MHz, $CD_3CN$)

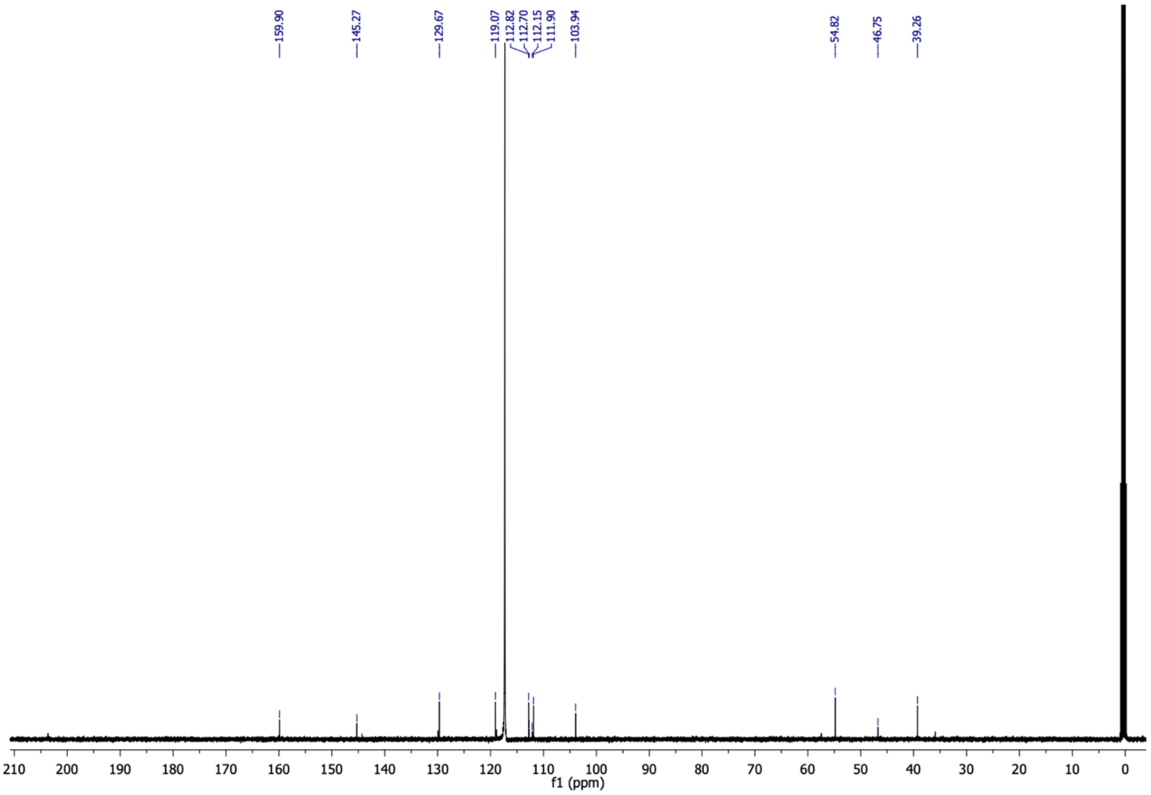

6-(3-methoxyphenyl)-4-oxo-4,5,6,7-tetrahydrobenzofuran-3-carboxylic acid (**4**) $^{1}$H NMR (500 MHz, $CDCl_3$)

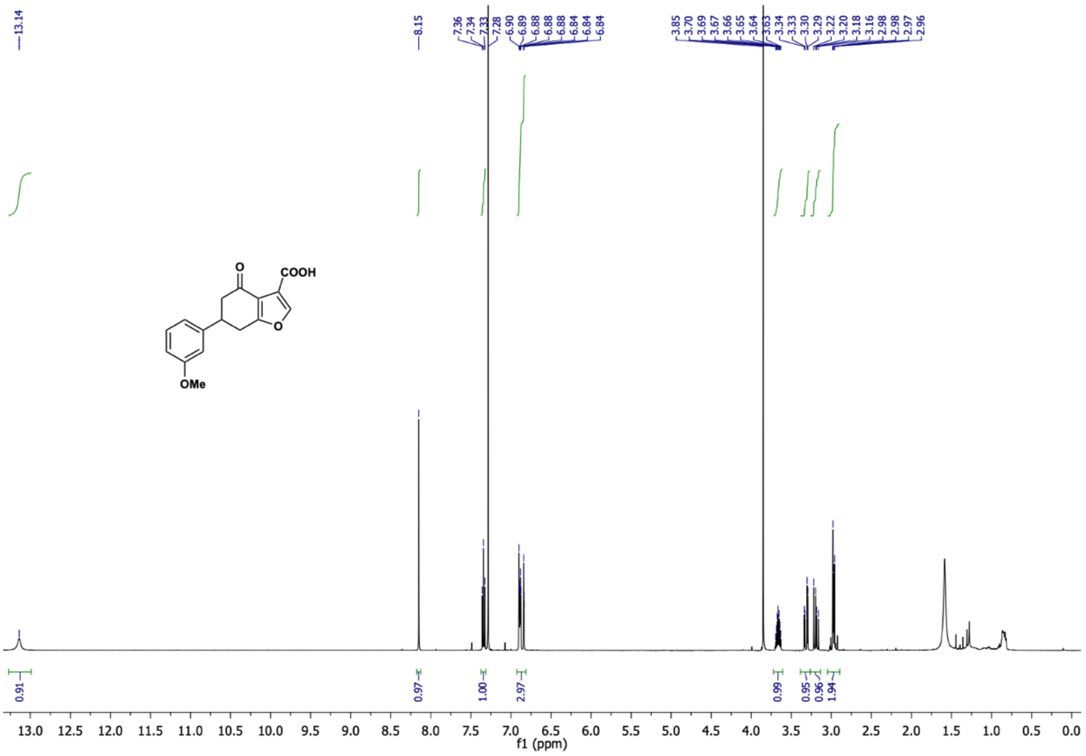

6-(3-methoxyphenyl)-4-oxo-4,5,6,7-tetrahydrobenzofuran-3-carboxylic acid (**4**) $^{13}$C NMR (126 MHz, $CDCl_3$)

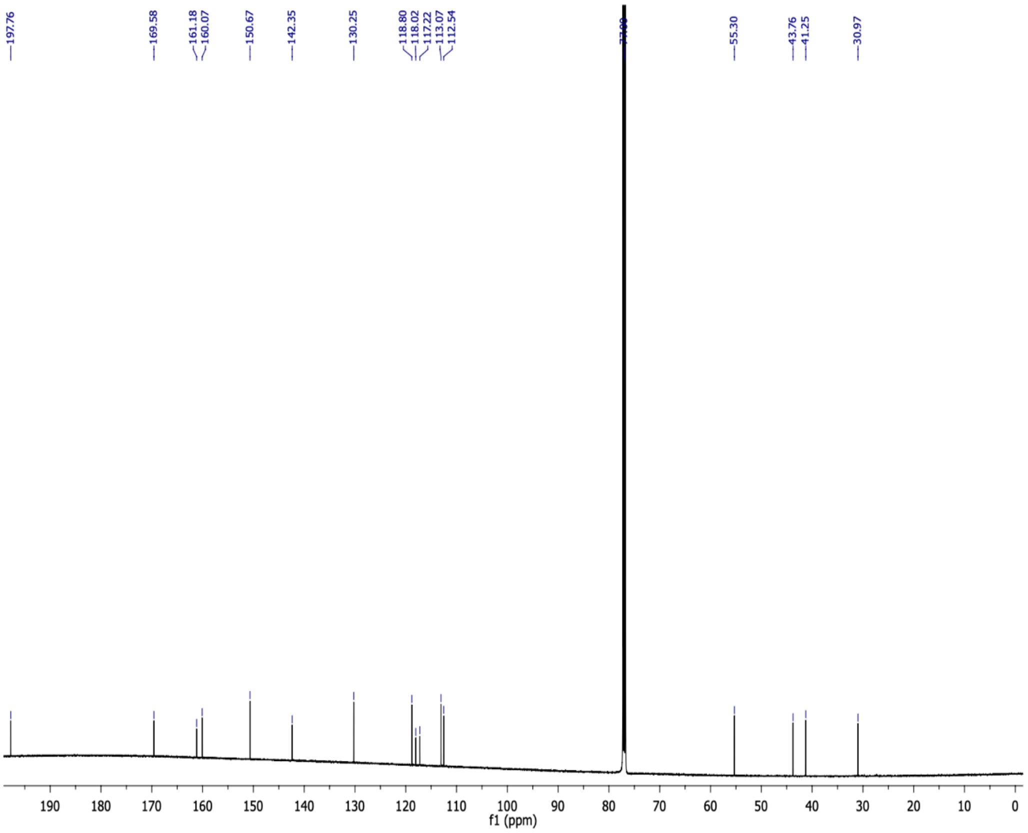

(*E*)-4-(3-(trifluoromethyl)phenyl)but-3-en-2-one (**5b**) $^{1}$H NMR (500 MHz, $CDCl_3$)

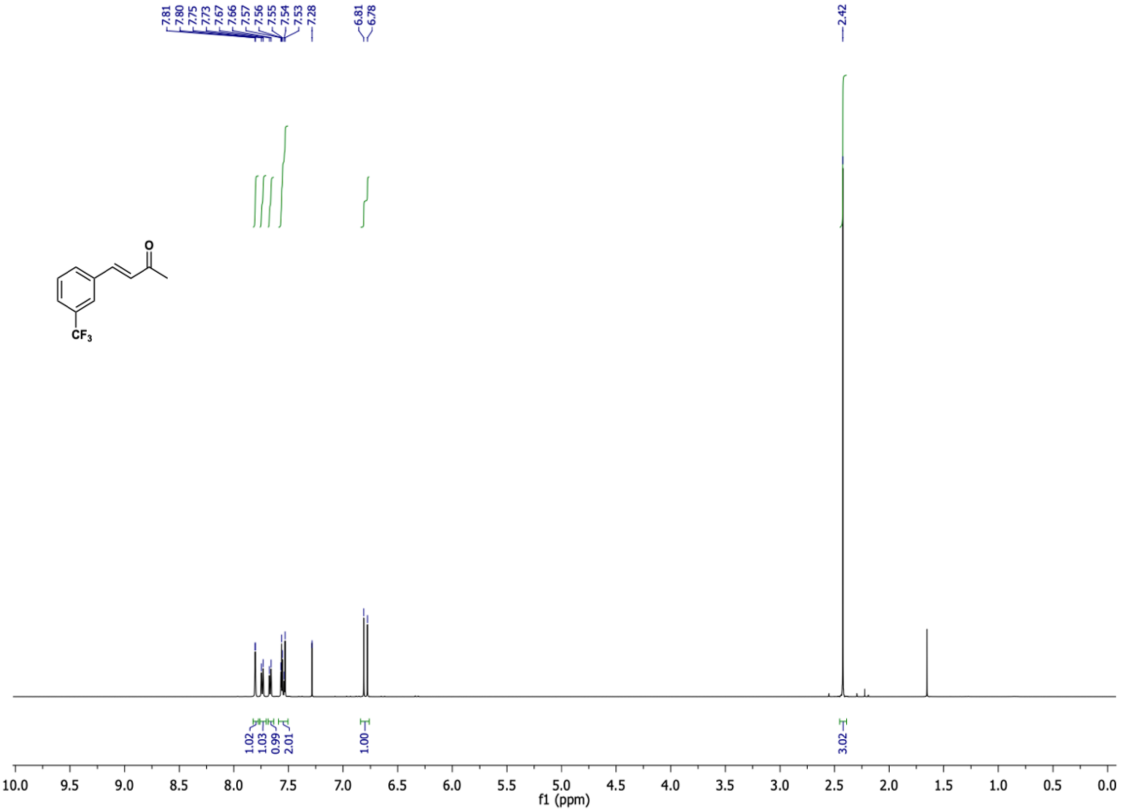

(*E*)-4-(3-(trifluoromethyl)phenyl)but-3-en-2-one (**5b**) $^{13}$C NMR (126 MHz, $CDCl_3$)

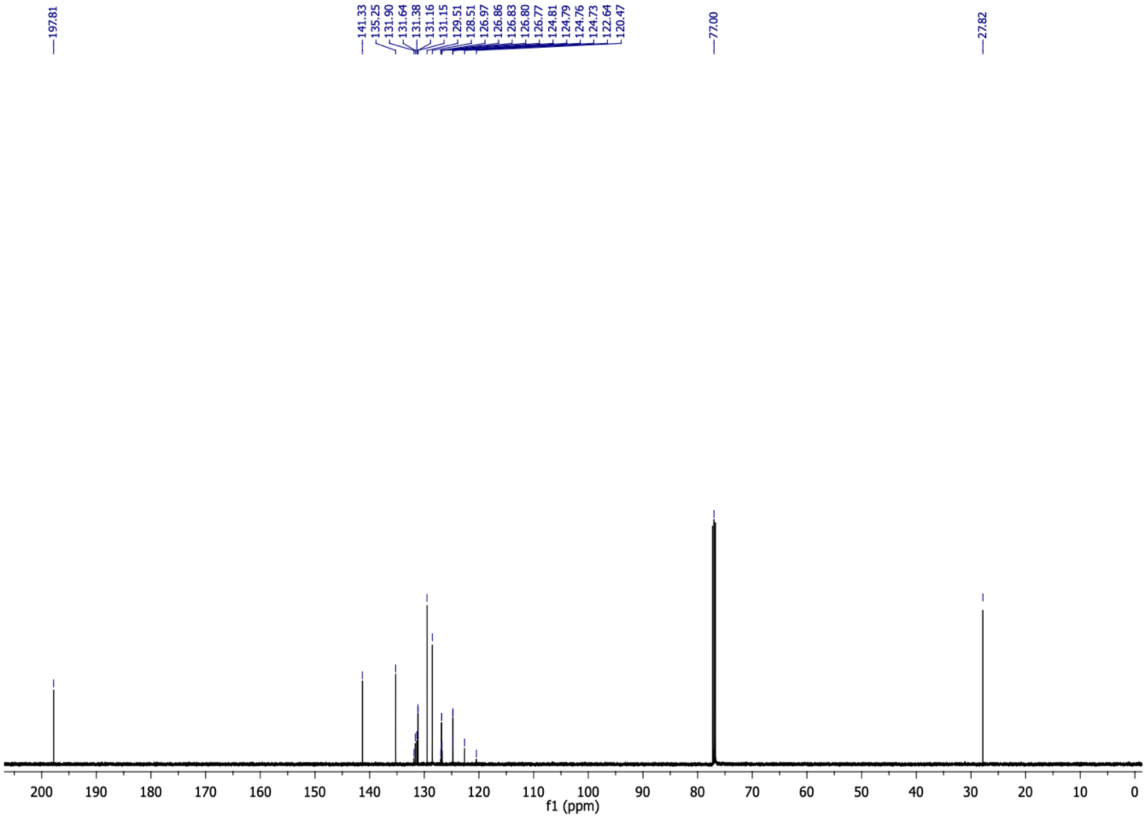

(*E*)-4-(3-(trifluoromethyl)phenyl)but-3-en-2-one (**5b**) **$^{19}$F NMR** (470 MHz, $CDCl_3$)

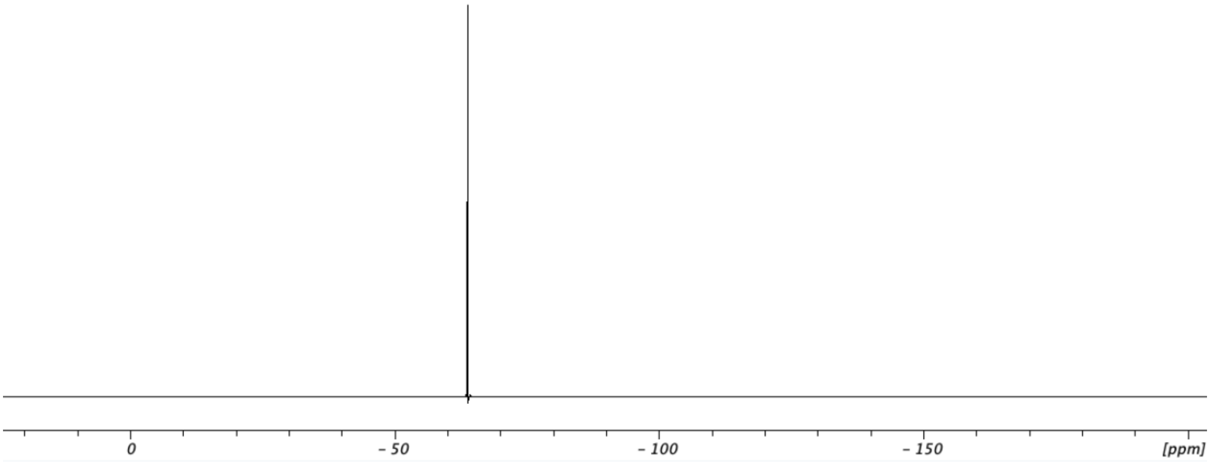

5-(3-(trifluoromethyl)phenyl)cyclohexane-1,3-dione (**5c**) $^{1}$H NMR (500 MHz, $CD_3OD$)

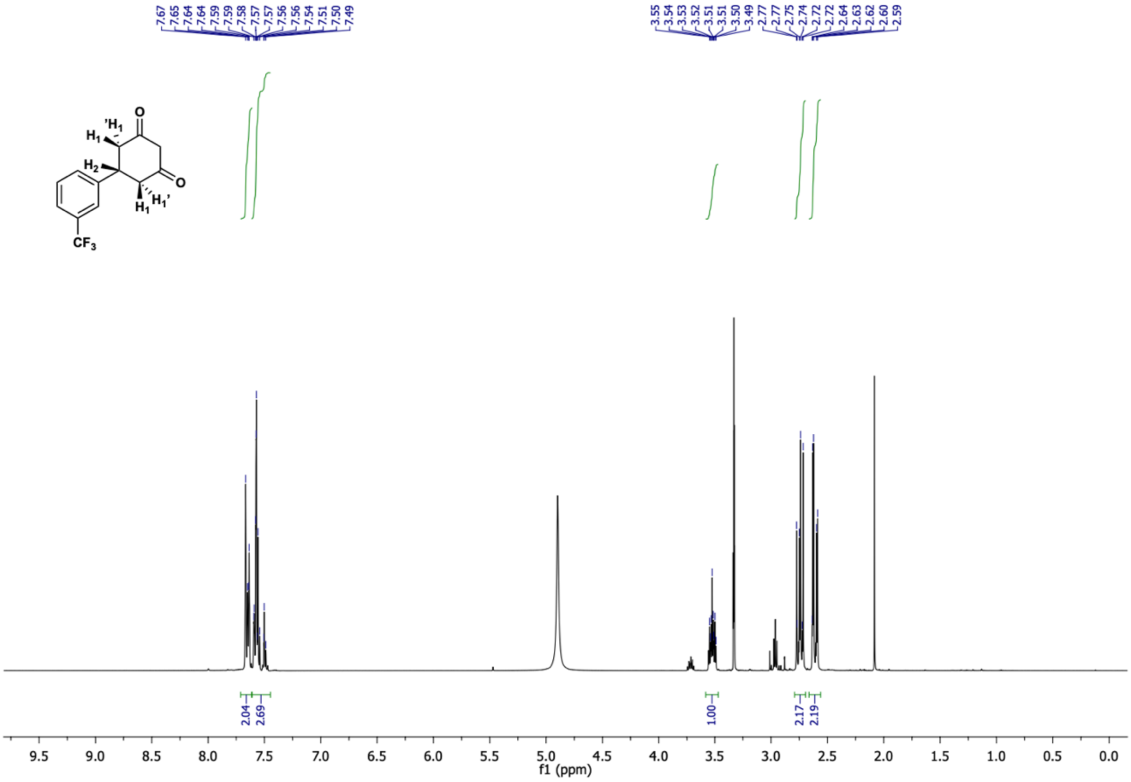

5-(3-(trifluoromethyl)phenyl)cyclohexane-1,3-dione (**5c**) $^{13}$C NMR (126 MHz, $CD_3OD$)

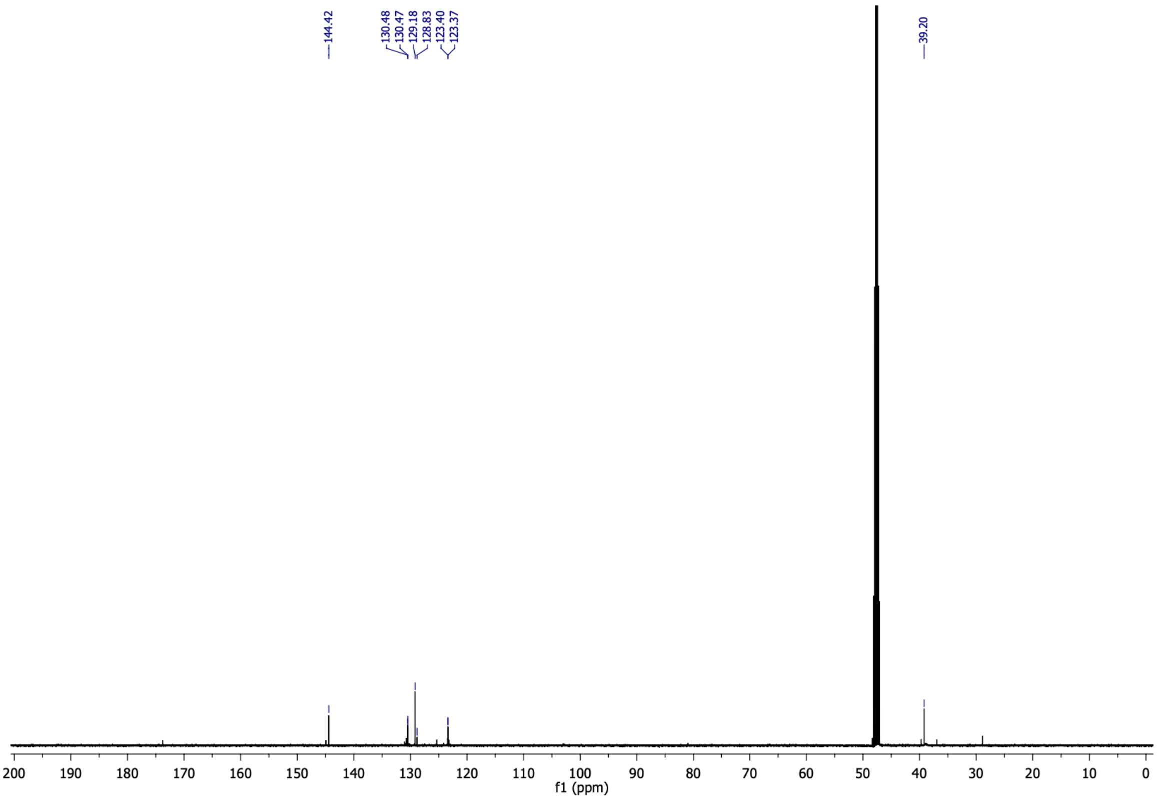

5-(3-(trifluoromethyl)phenyl)cyclohexane-1,3-dione (**5c**) $^{19}$F NMR (470 MHz, $CDCl_3$)

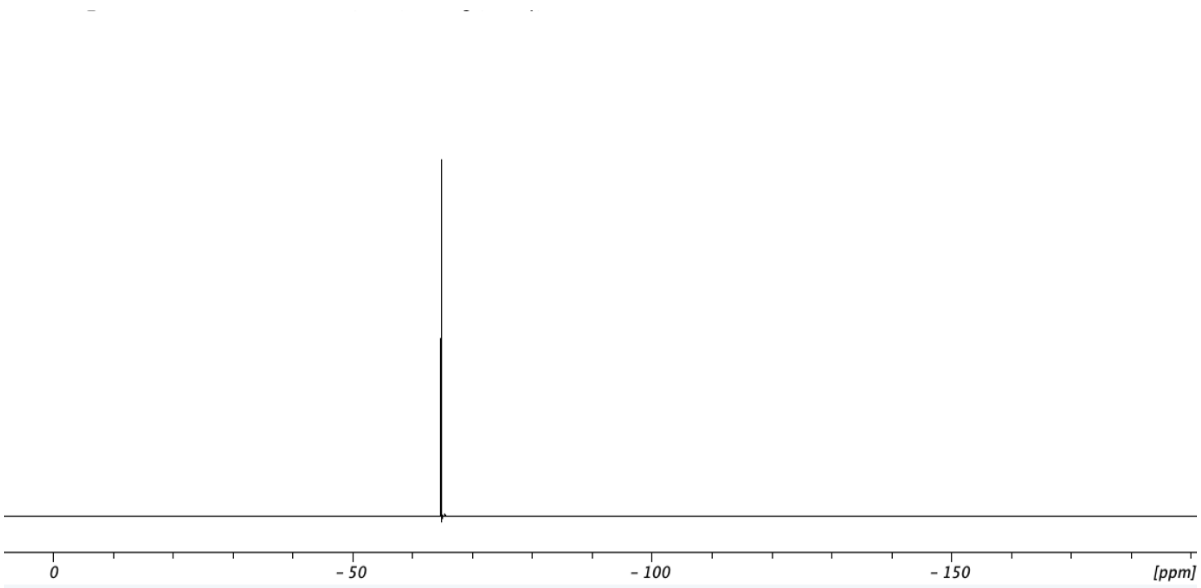

4-oxo-6-(3-(trifluoromethyl)phenyl)-4,5,6,7-tetrahydrobenzofuran-3-carboxylic acid (**5**) $^{1}H$ NMR (500 MHz, $CDCl_3$)

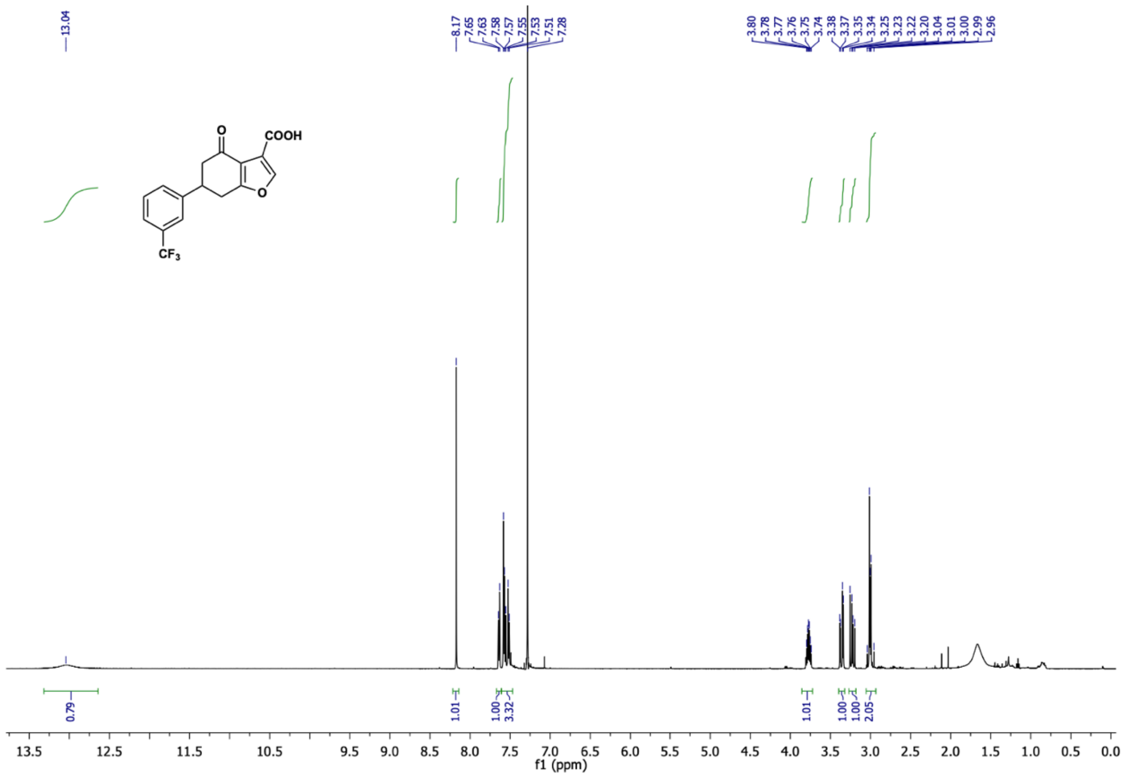

4-oxo-6-(3-(trifluoromethyl)phenyl)-4,5,6,7-tetrahydrobenzofuran-3-carboxylic acid (**5**) $^{13}C$ NMR (126 MHz, $CDCl_3$)

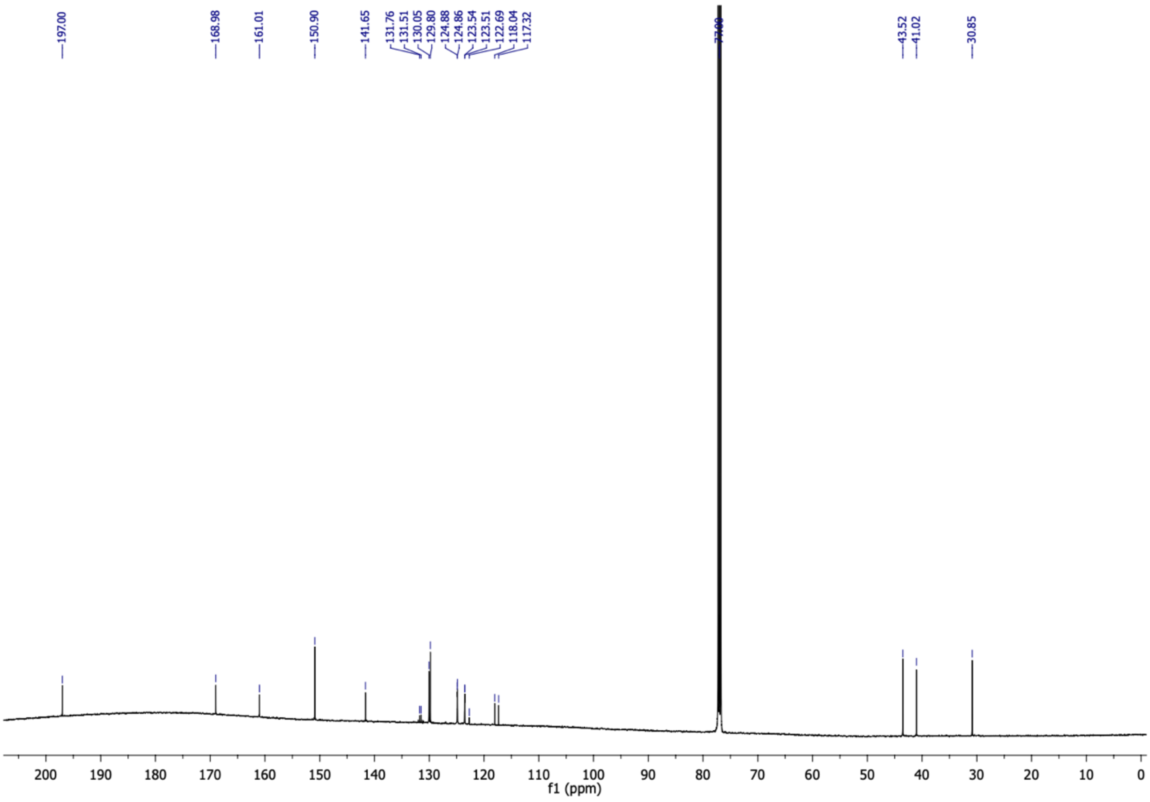

4-oxo-6-(3-(trifluoromethyl)phenyl)-4,5,6,7-tetrahydrobenzofuran-3-carboxylic acid (**5**) $^{19}F$ NMR (470 MHz, $CDCl_3$)

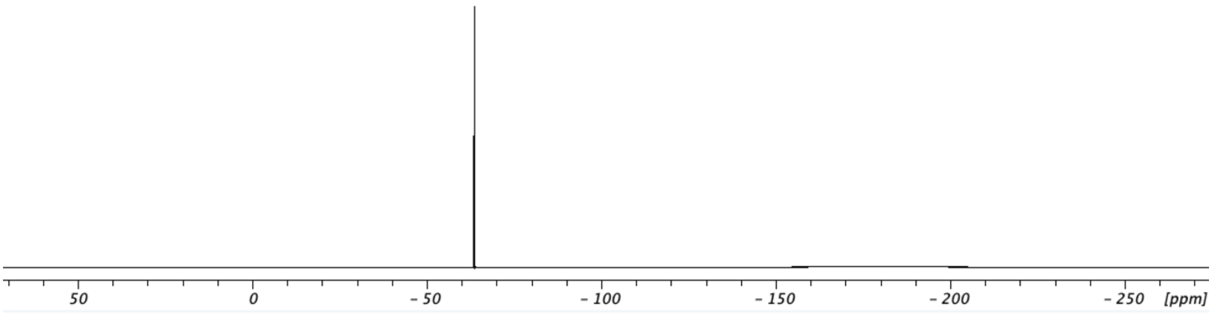

(E)-4-(benzofuran-2-yl)but-3-en-2-one (**6b**) $^{1}$H NMR (400 MHz, $CDCl_3$)

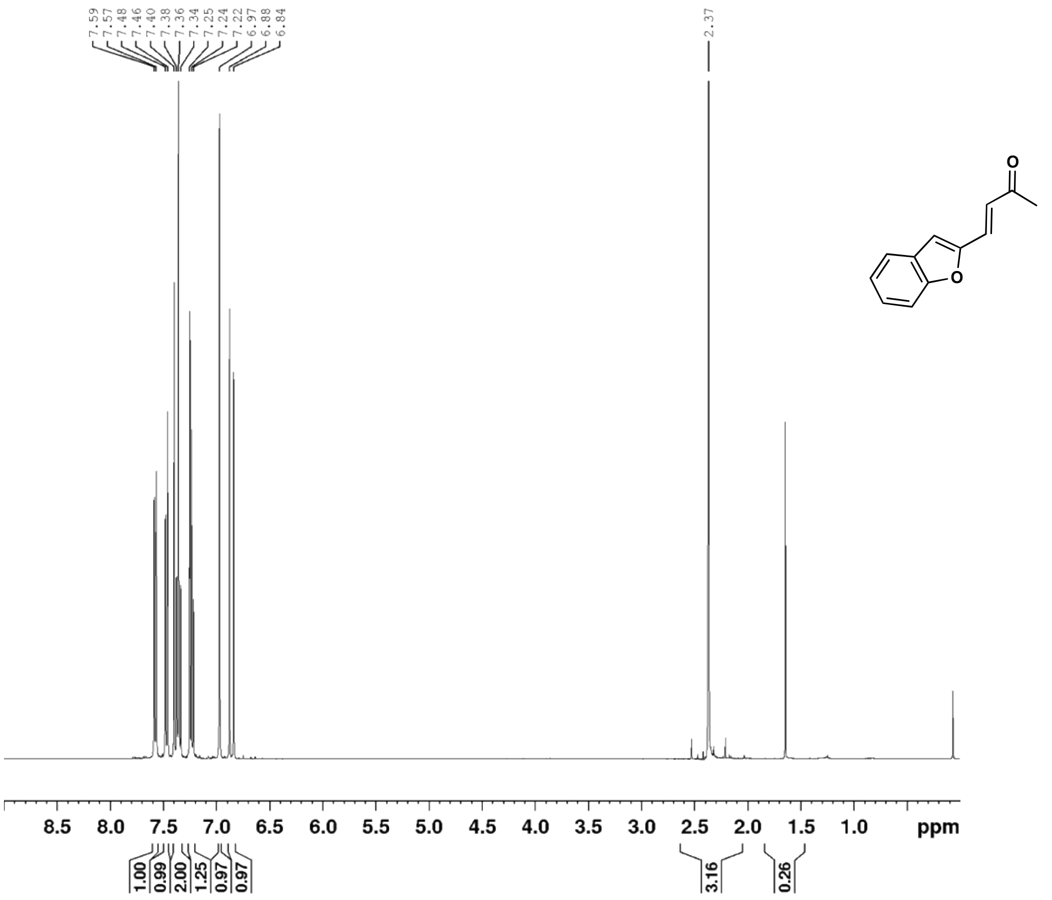

(E)-4-(benzofuran-2-yl)but-3-en-2-one (**6b**) $^{13}$C NMR (101 MHz, $CDCl_3$)

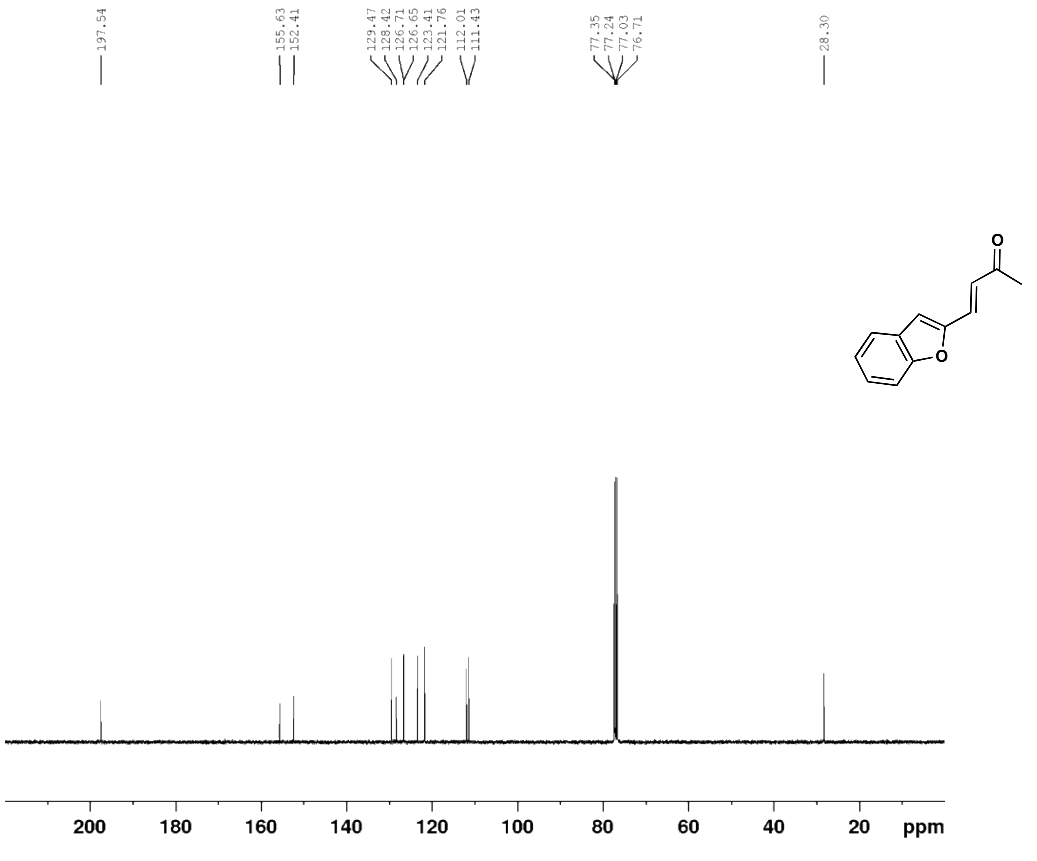

5-(benzofuran-2-yl)cyclohexane-1,3-dione (**6c**) $^{1}$H NMR (400 MHz, $CD_3OD$)

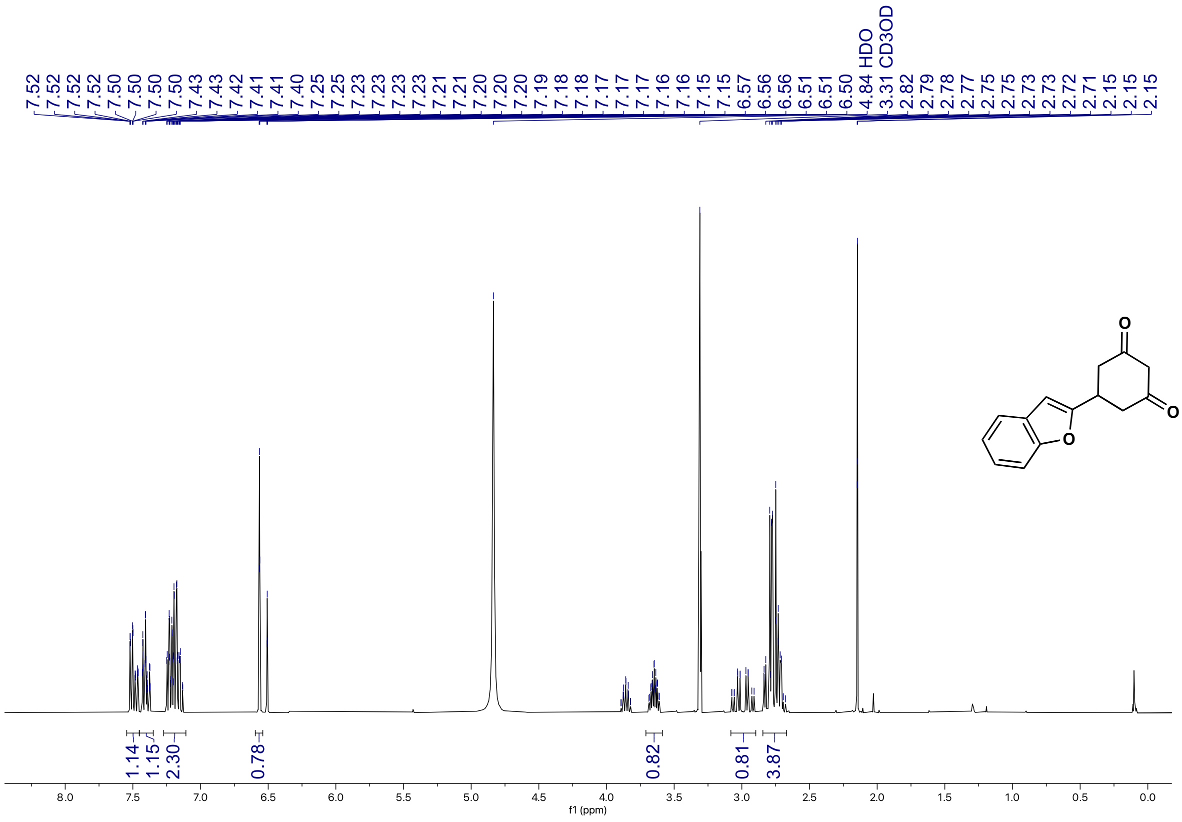

5-(benzofuran-2-yl)cyclohexane-1,3-dione (**6c**) $^{13}$C NMR (101 MHz, $CD_3OD$)

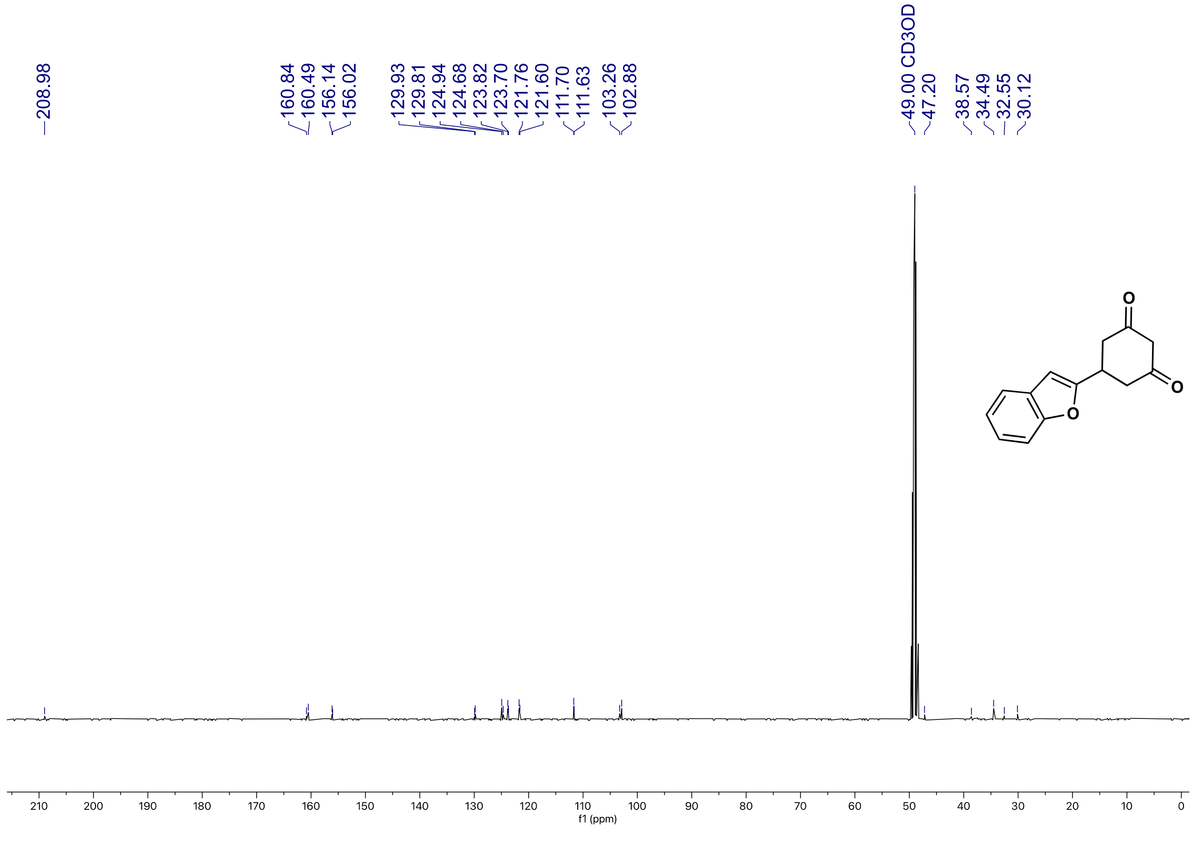

4'-oxo-4',5',6',7'-tetrahydro-[2,6'-bibenzofuran]-3'-carboxylic acid (**6**) $^{1}H$ NMR (500 MHz, $CD_3OD$)

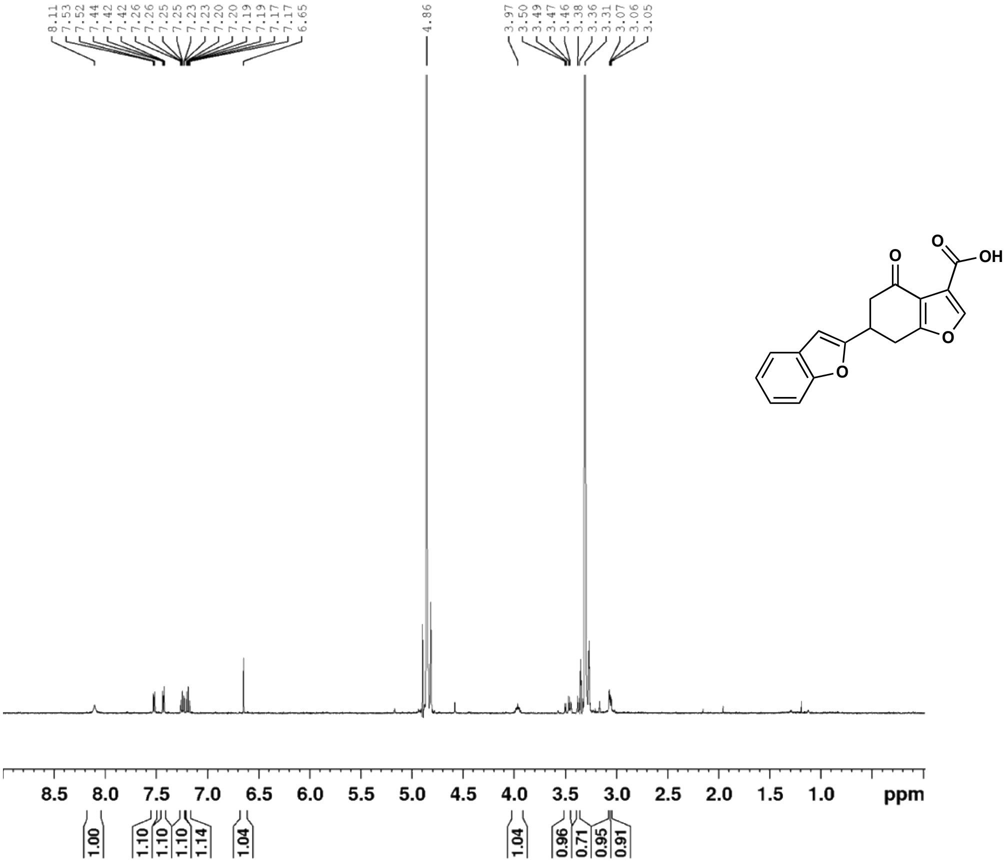

4'-oxo-4',5',6',7'-tetrahydro-[2,6'-bibenzofuran]-3'-carboxylic acid (**6**) $^{13}C$ DEPT NMR (125 MHz, $CD_3OD$)

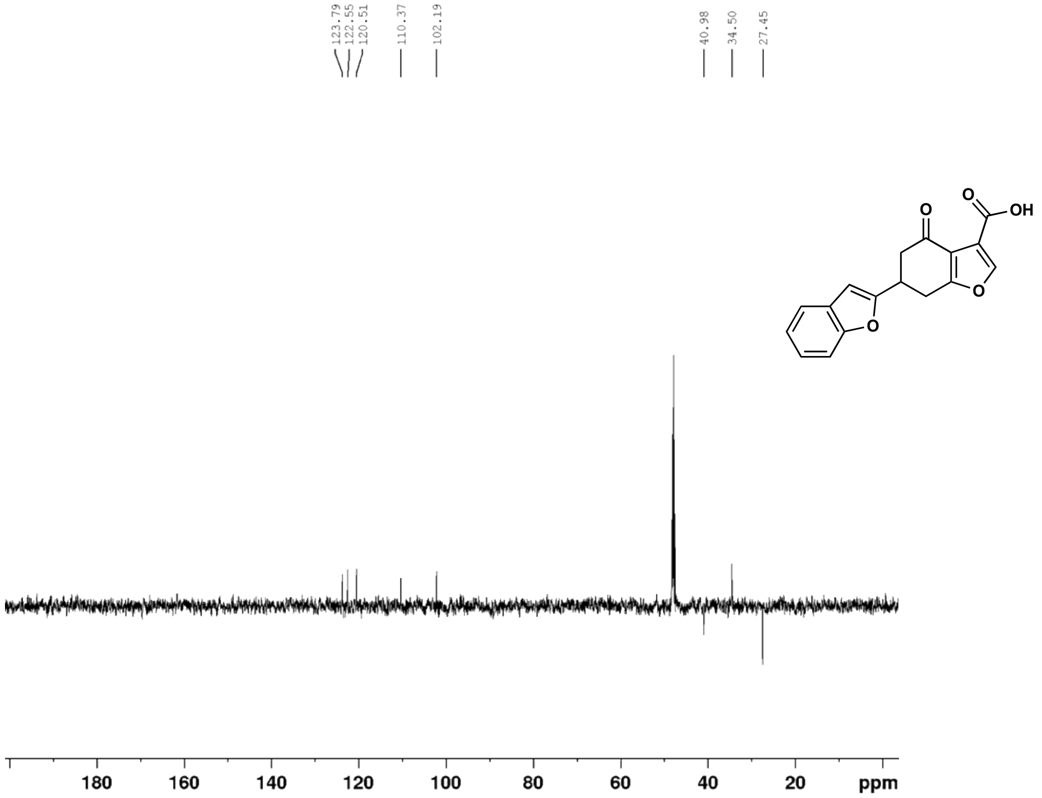

(*E*)-4-(2-bromophenyl)but-3-en-2-one (**7b**) $^{1}$H NMR (500 MHz, $CDCl_3$)

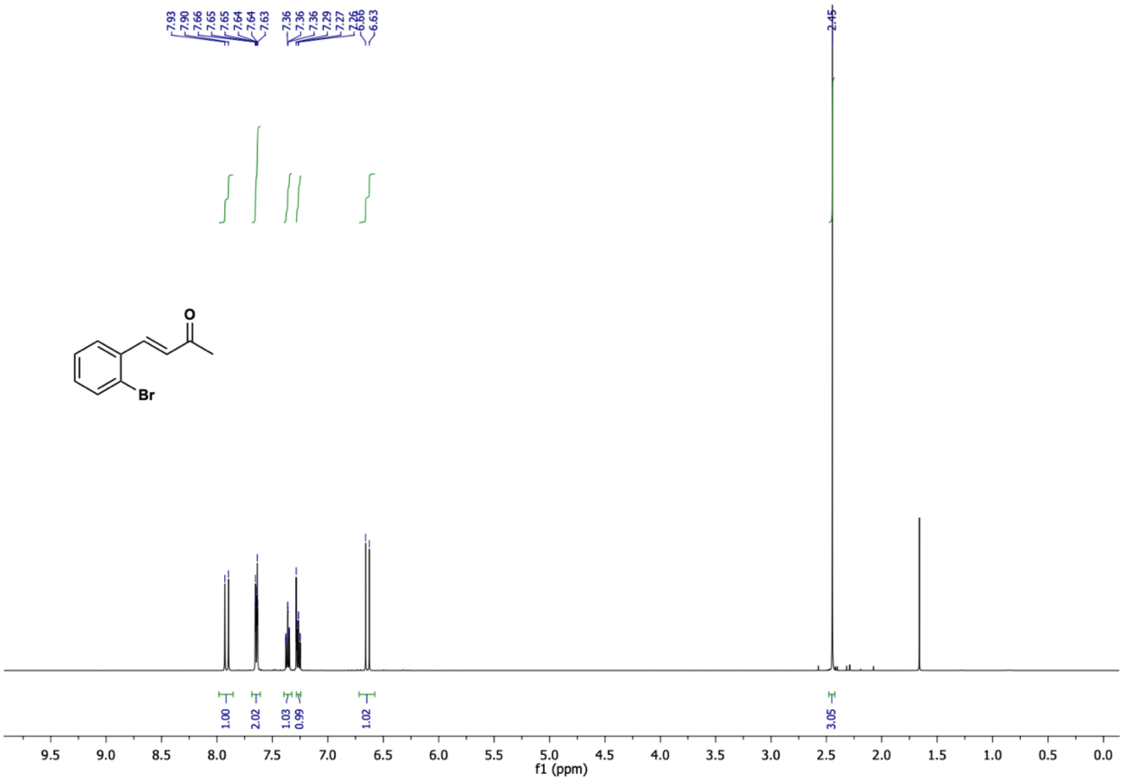

(*E*)-4-(2-bromophenyl)but-3-en-2-one (**7b**) $^{13}$C NMR (126 MHz, $CDCl_3$)

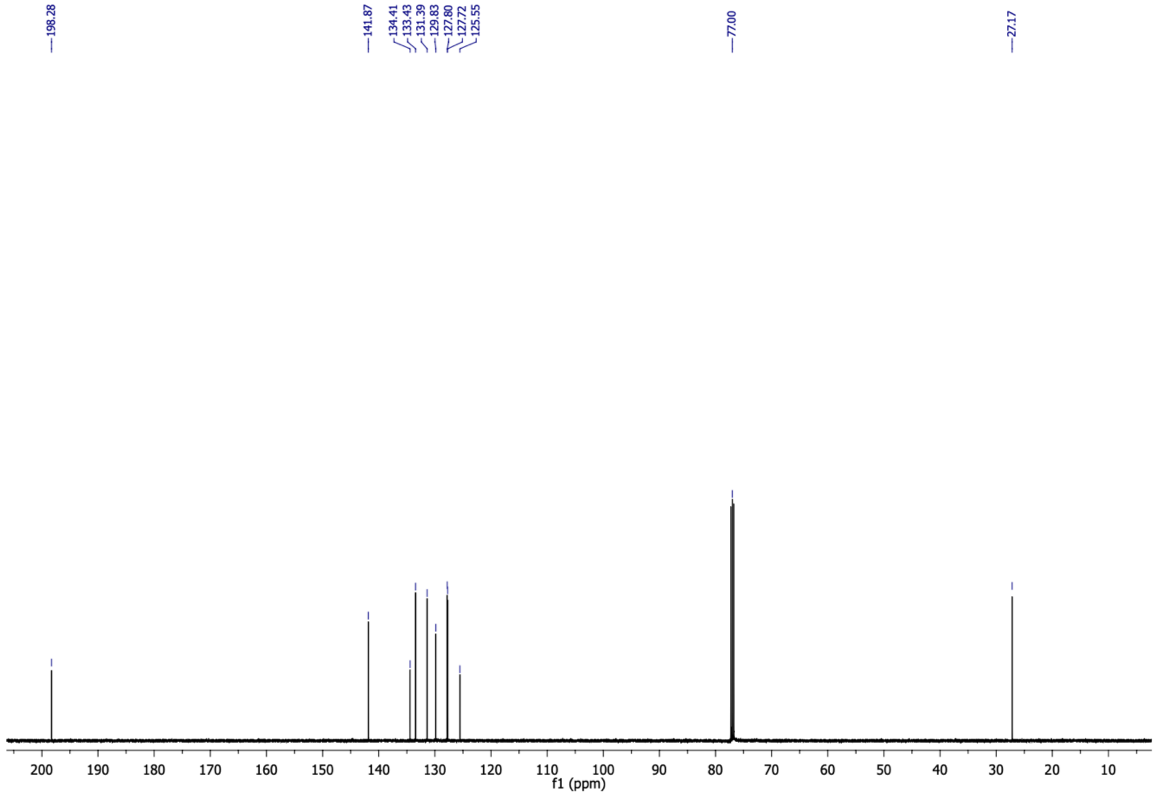

5-(2-bromophenyl)cyclohexane-1,3-dione (**7c**) $^{1}H$ NMR (500 MHz, $CD_3OD$)

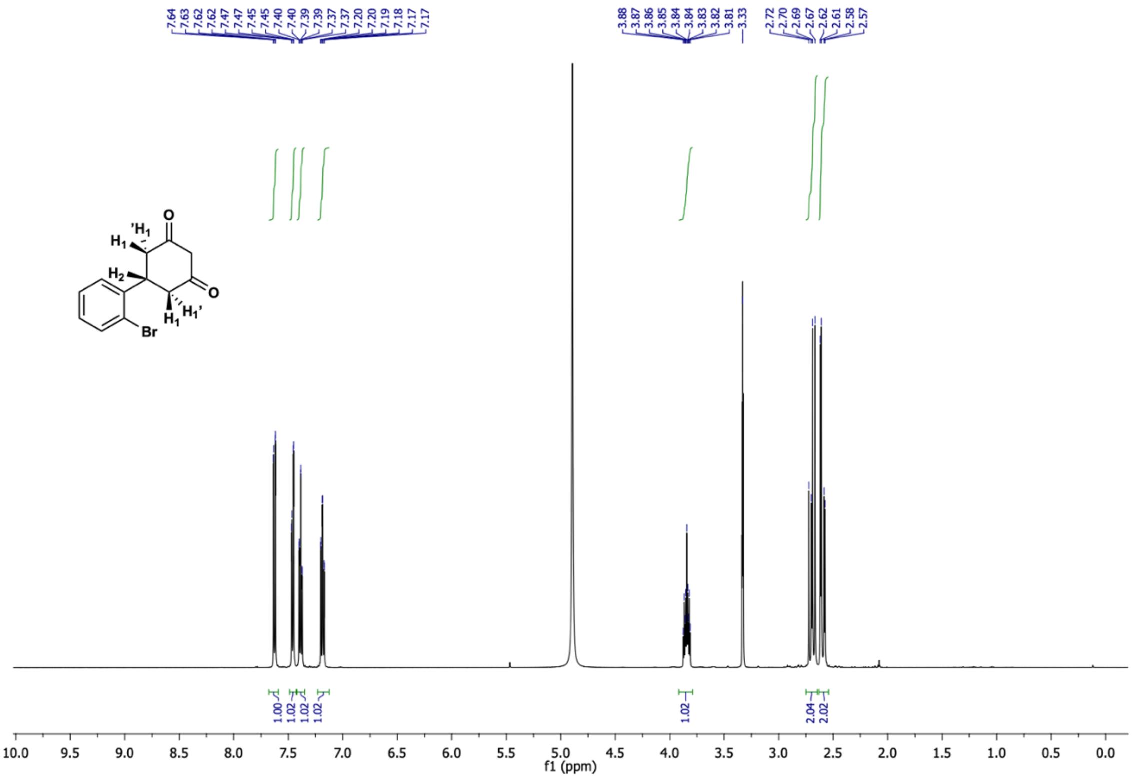

5-(2-bromophenyl)cyclohexane-1,3-dione (**7c**) $^{13}C$ NMR (126 MHz, $CD_3OD$)

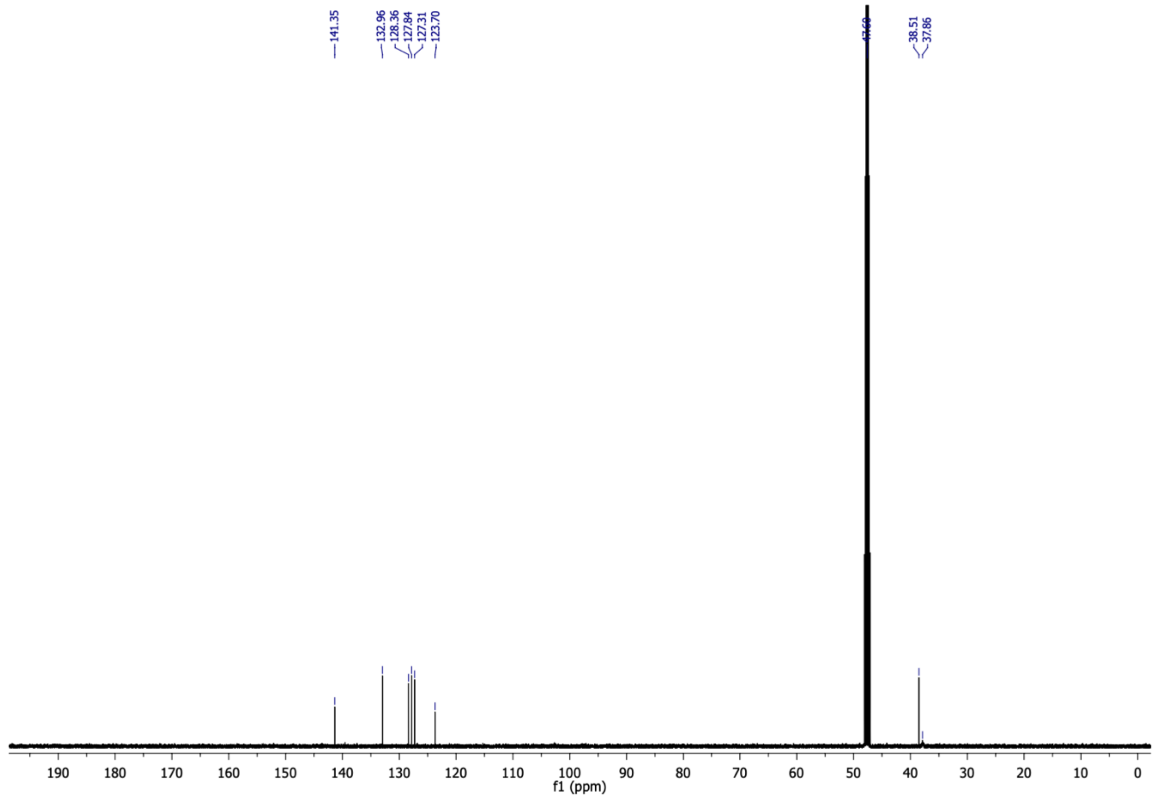

6-(2-bromophenyl)-4-oxo-4,5,6,7-tetrahydrobenzofuran-3-carboxylic acid (**7**) $^{1}$H NMR (500 MHz, $CDCl_3$)

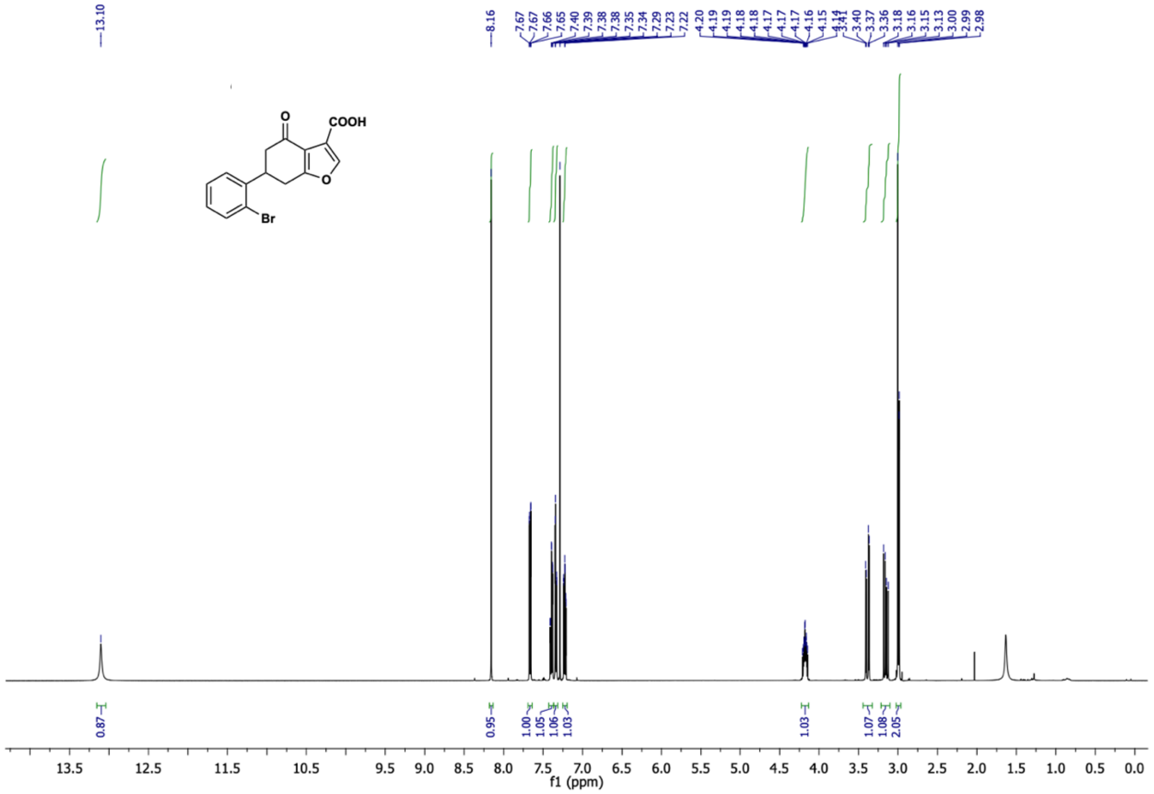

6-(2-bromophenyl)-4-oxo-4,5,6,7-tetrahydrobenzofuran-3-carboxylic acid (**7**) $^{13}$C NMR (126 MHz, $CDCl_3$)

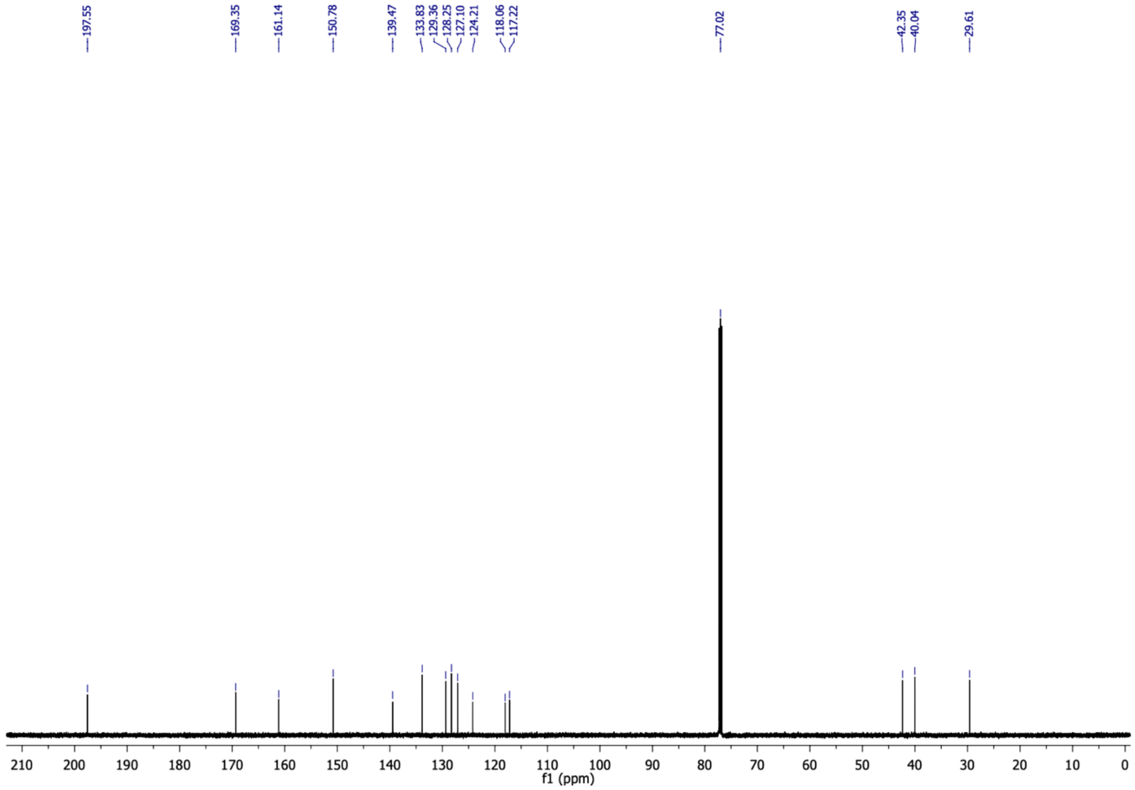

(*E*)-4-(2-fluorophenyl)but-3-en-2-one (**8b**) $^{1}$H NMR (500 MHz, $CDCl_3$)

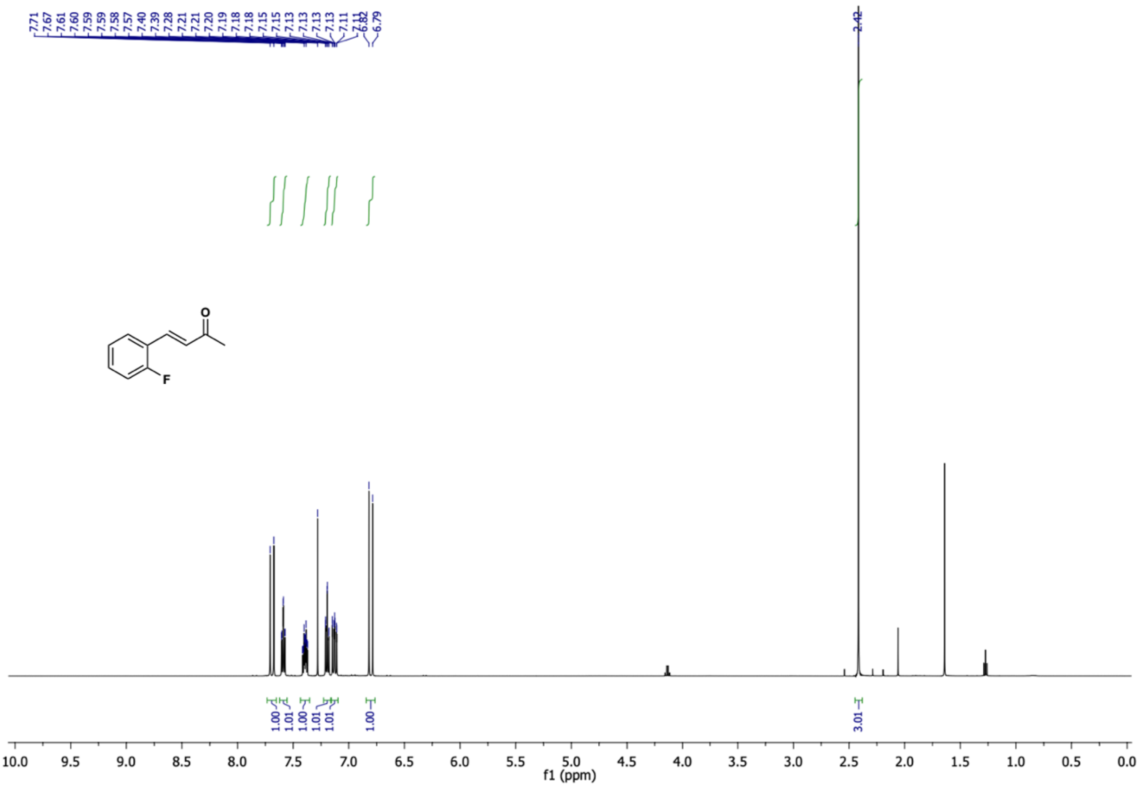

(*E*)-4-(2-fluorophenyl)but-3-en-2-one (**8b**) $^{13}$C NMR (126 MHz, $CDCl_3$)

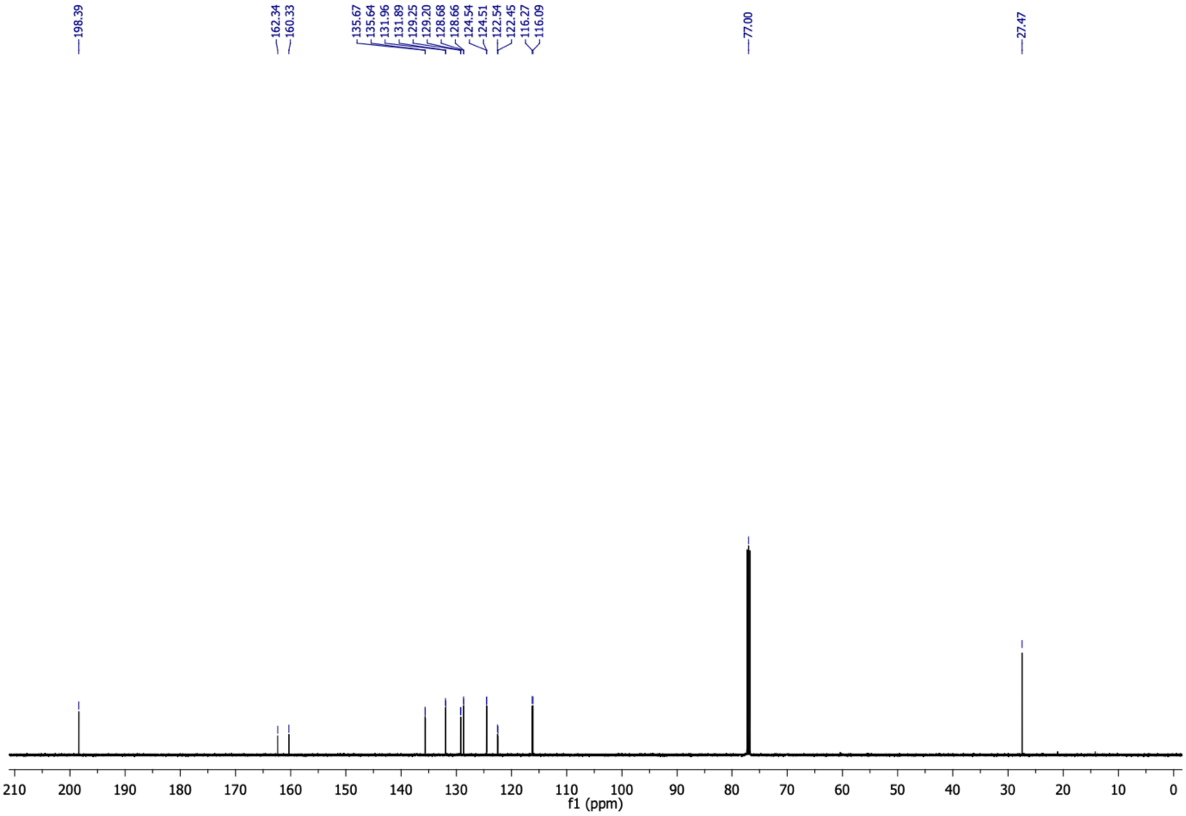

(*E*)-4-(2-fluorophenyl)but-3-en-2-one (**8b**) $^{19}$F NMR (376 MHz, $CDCl_3$)

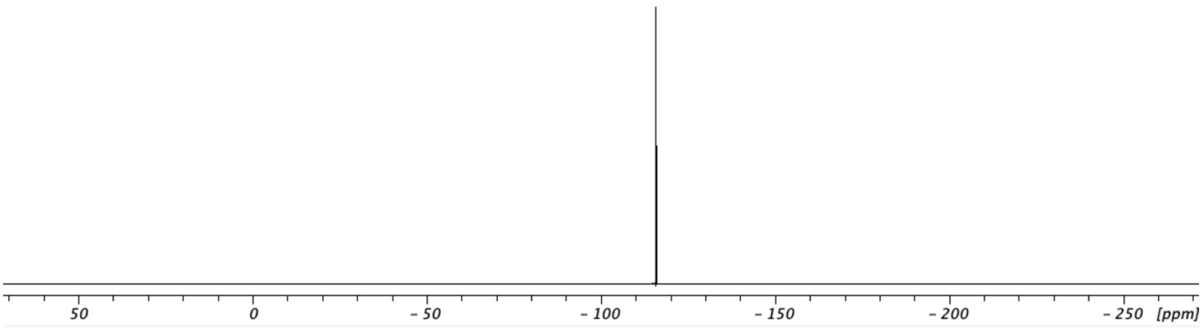

5-(2-fluorophenyl)cyclohexane-1,3-dione (**8c**) $^{1}$H NMR (500 MHz, $CD_3OD$)

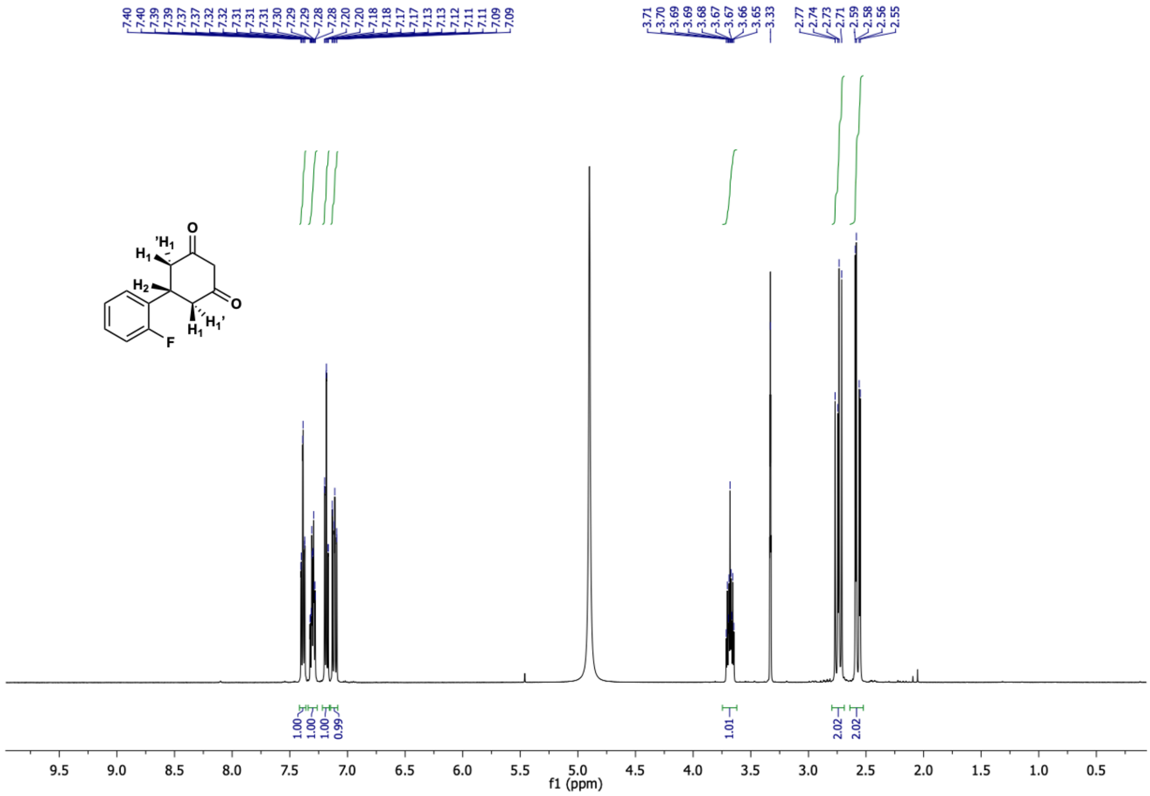

5-(2-fluorophenyl)cyclohexane-1,3-dione (**8c**) $^{13}C$ NMR (126 MHz, $CD_3OD$)

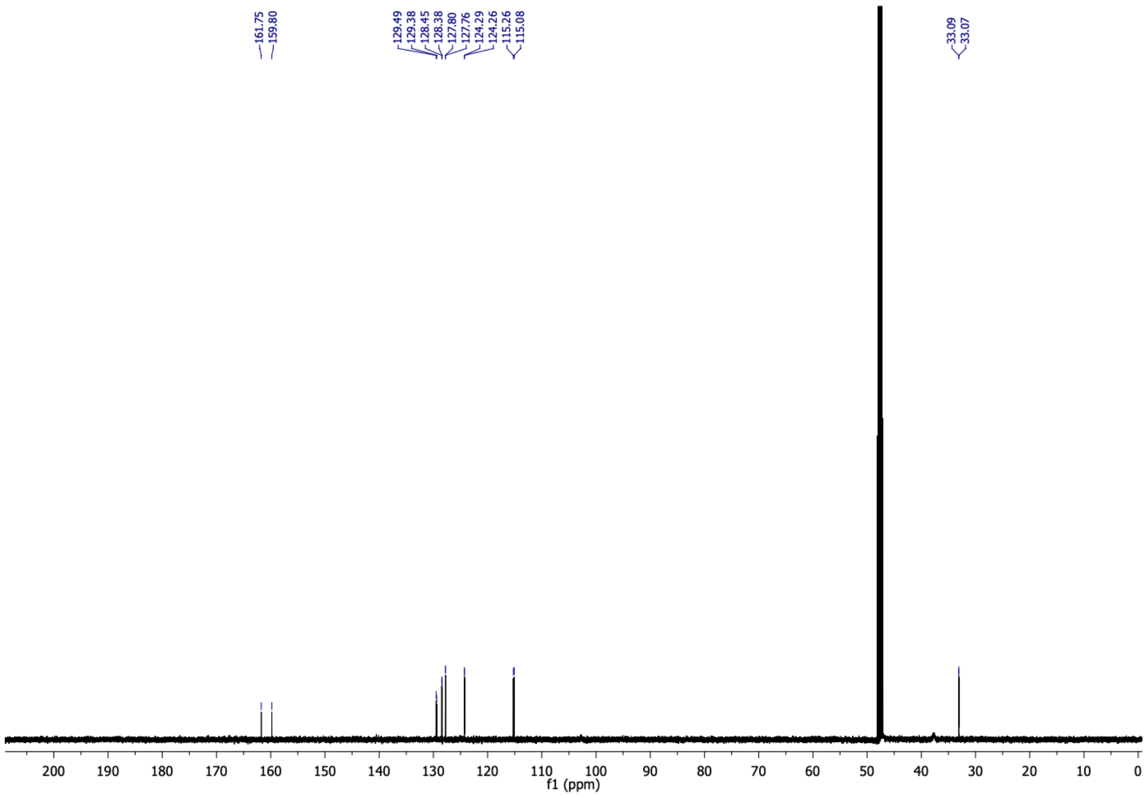

5-(2-fluorophenyl)cyclohexane-1,3-dione (**8c**) $^{19}F$ NMR (376 MHz, $CDCl_3$)

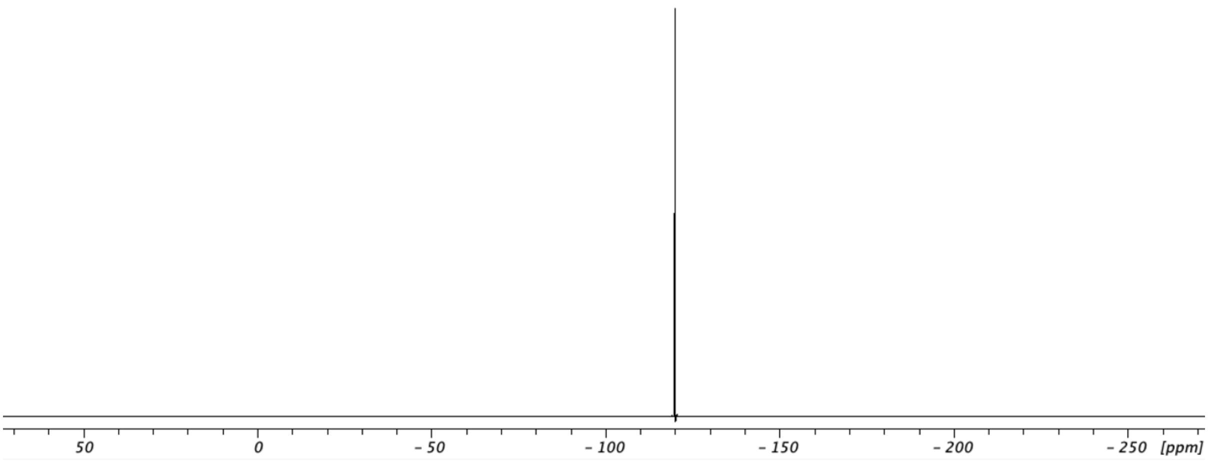

6-(2-fluorophenyl)-4-oxo-4,5,6,7-tetrahydrobenzofuran-3-carboxylic acid (**8**) $^{1}H$ NMR (500 MHz, $CDCl_3$)

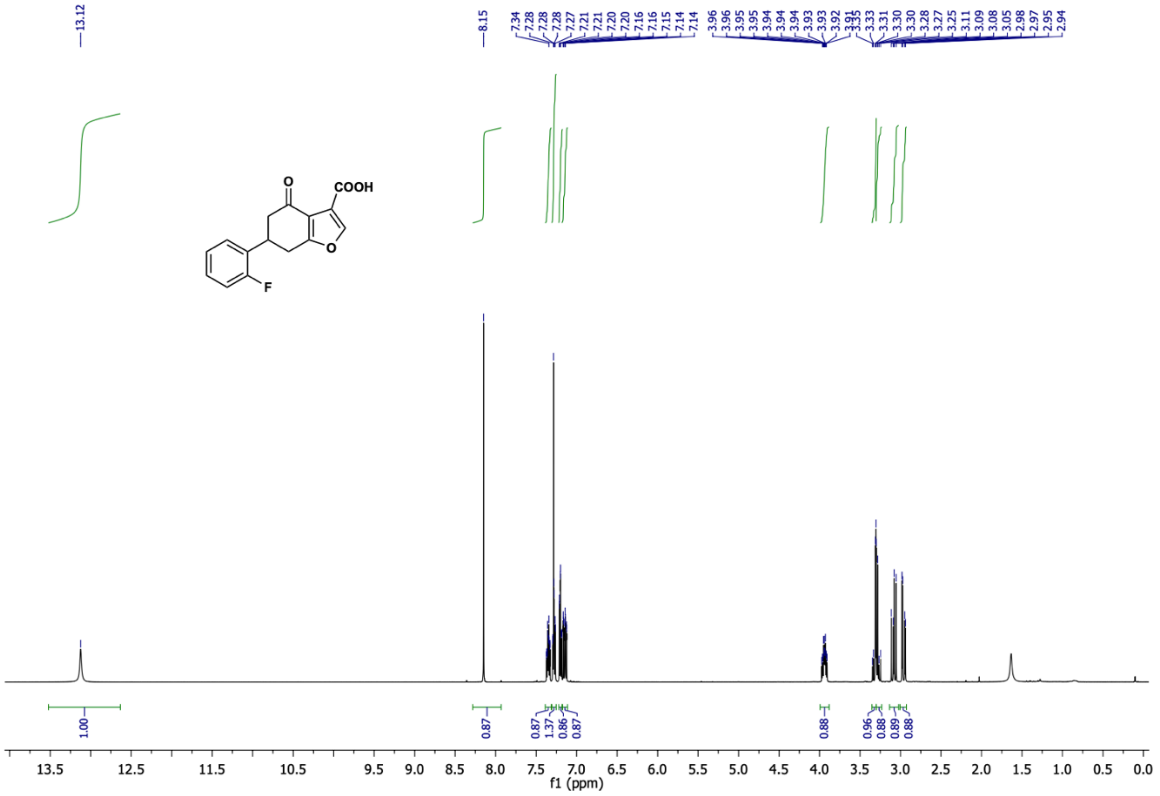

6-(2-fluorophenyl)-4-oxo-4,5,6,7-tetrahydrobenzofuran-3-carboxylic acid (**8**) $^{13}C$ NMR (126 MHz, $CDCl_3$)

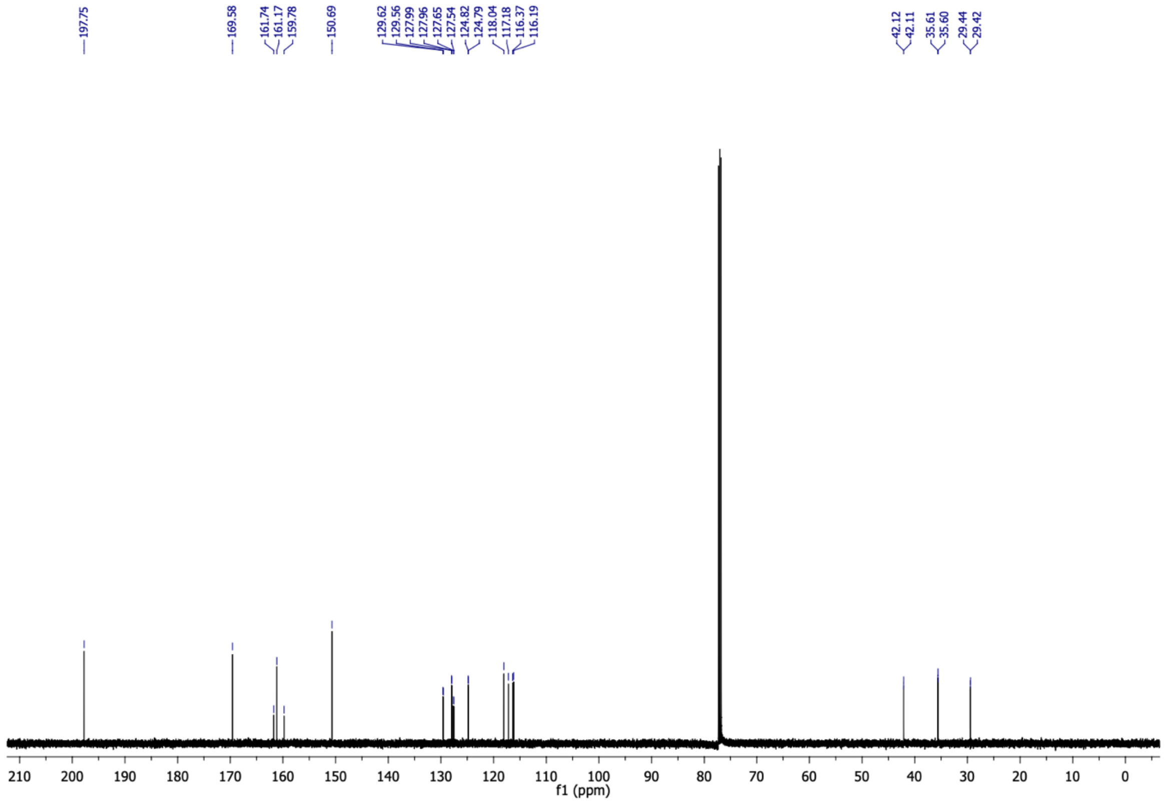

6-(2-fluorophenyl)-4-oxo-4,5,6,7-tetrahydrobenzofuran-3-carboxylic acid (**8**) $^{19}F$ NMR (376 MHz, $CDCl_3$)

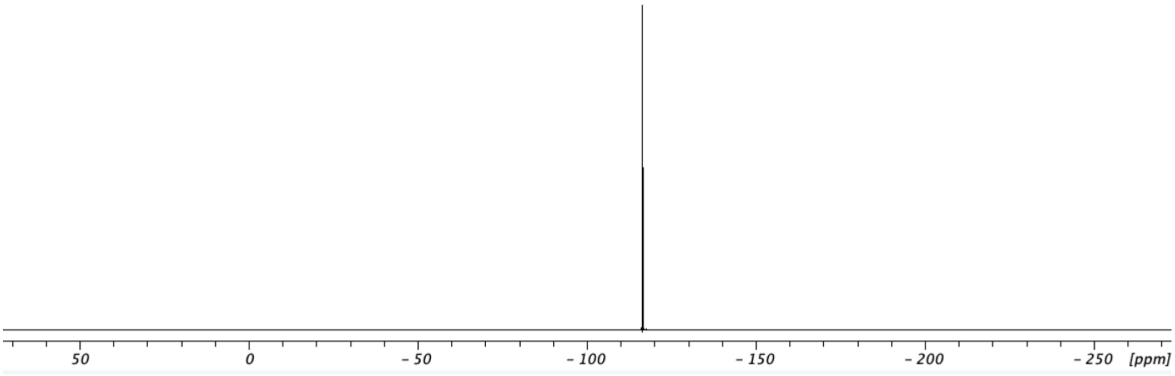

(*E*)-4-(2-phenoxyphenyl)but-3-en-2-one (**9b**) $^{1}H$ NMR (500 MHz, $CDCl_3$)

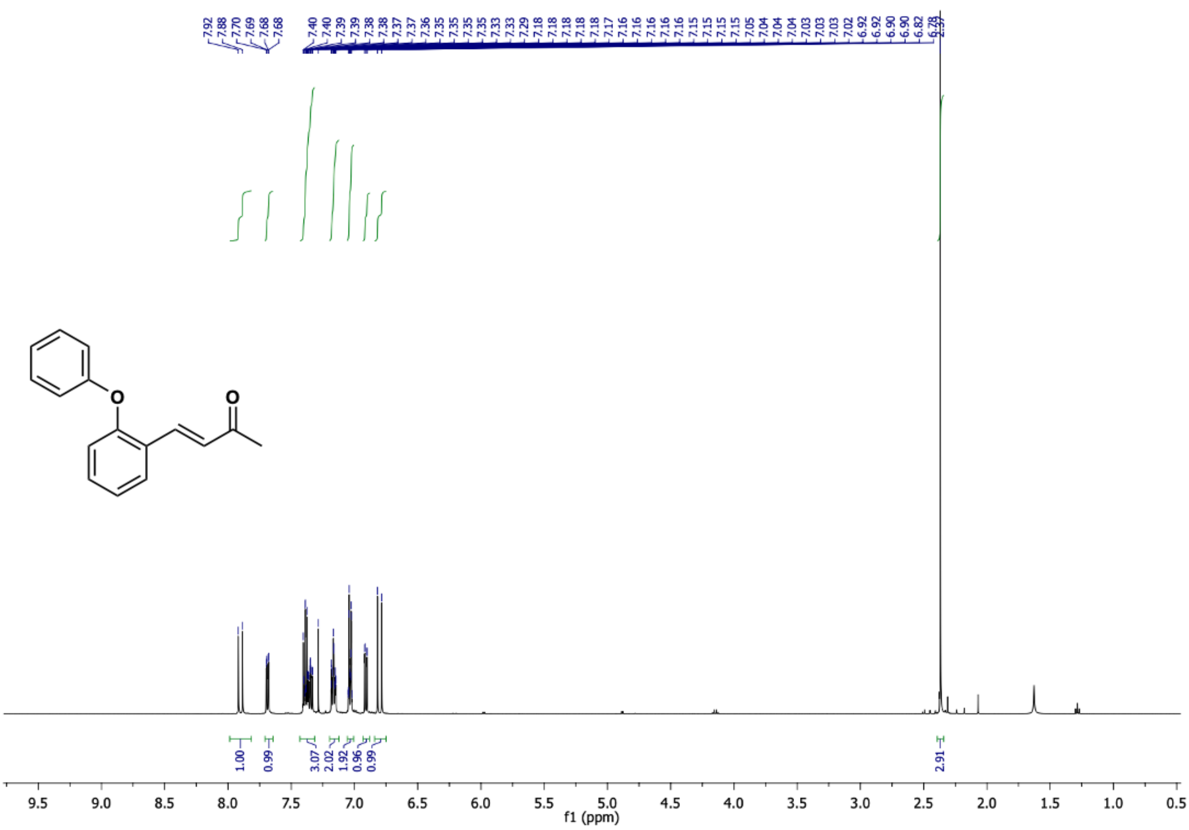

(*E*)-4-(2-phenoxyphenyl)but-3-en-2-one (**9b**) $^{13}C$ NMR (126 MHz, $CDCl_3$)

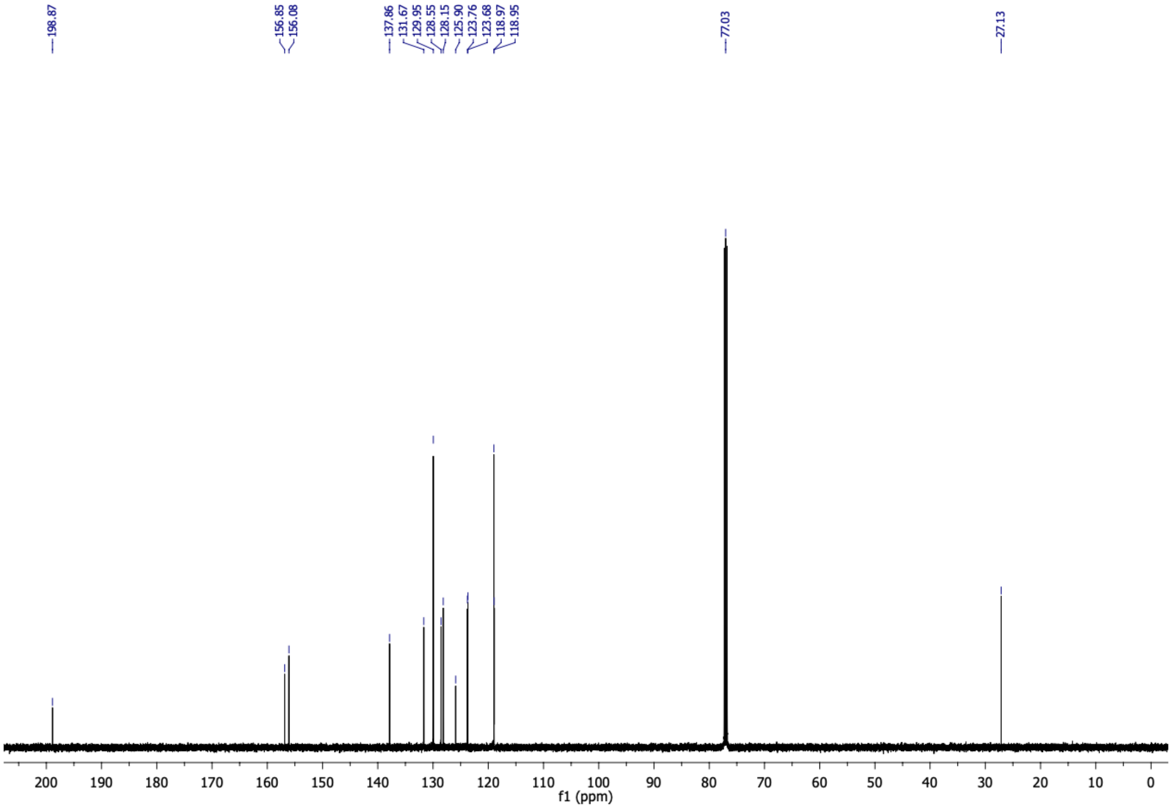

5-(2-phenoxyphenyl)cyclohexane-1,3-dione (**9c**) $^{1}H$ NMR (500 MHz, $CD_3OD$)

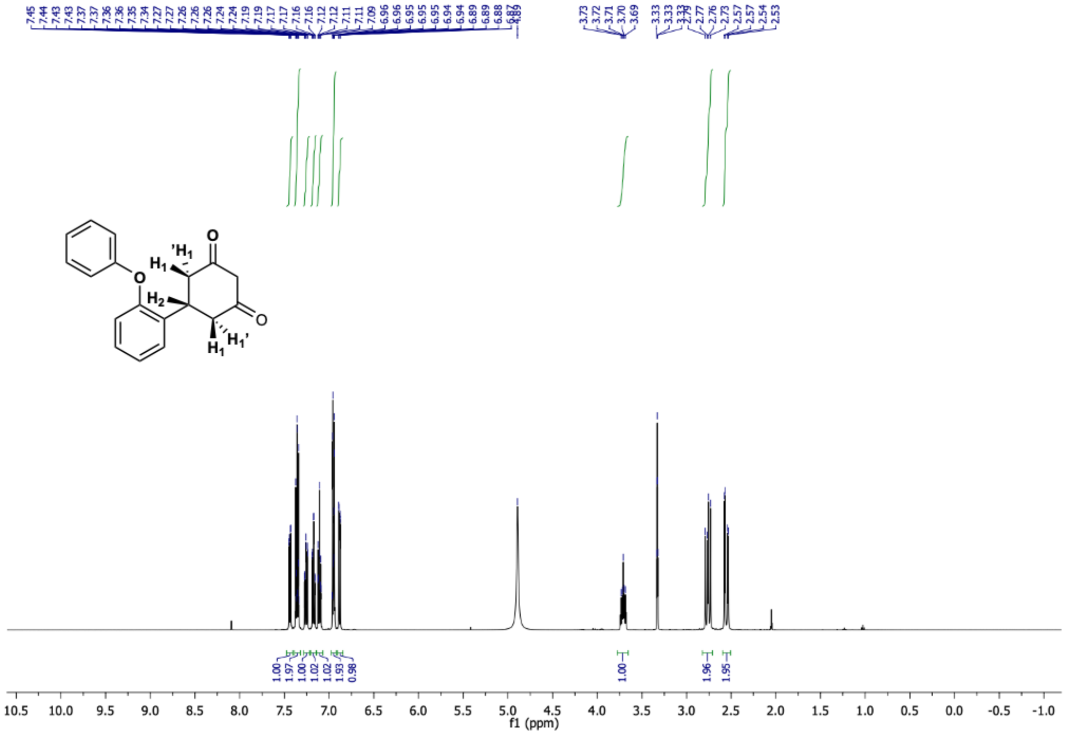

5-(2-phenoxyphenyl)cyclohexane-1,3-dione (**9c**) $^{13}C$ NMR (126 MHz, $CD_3OD$)

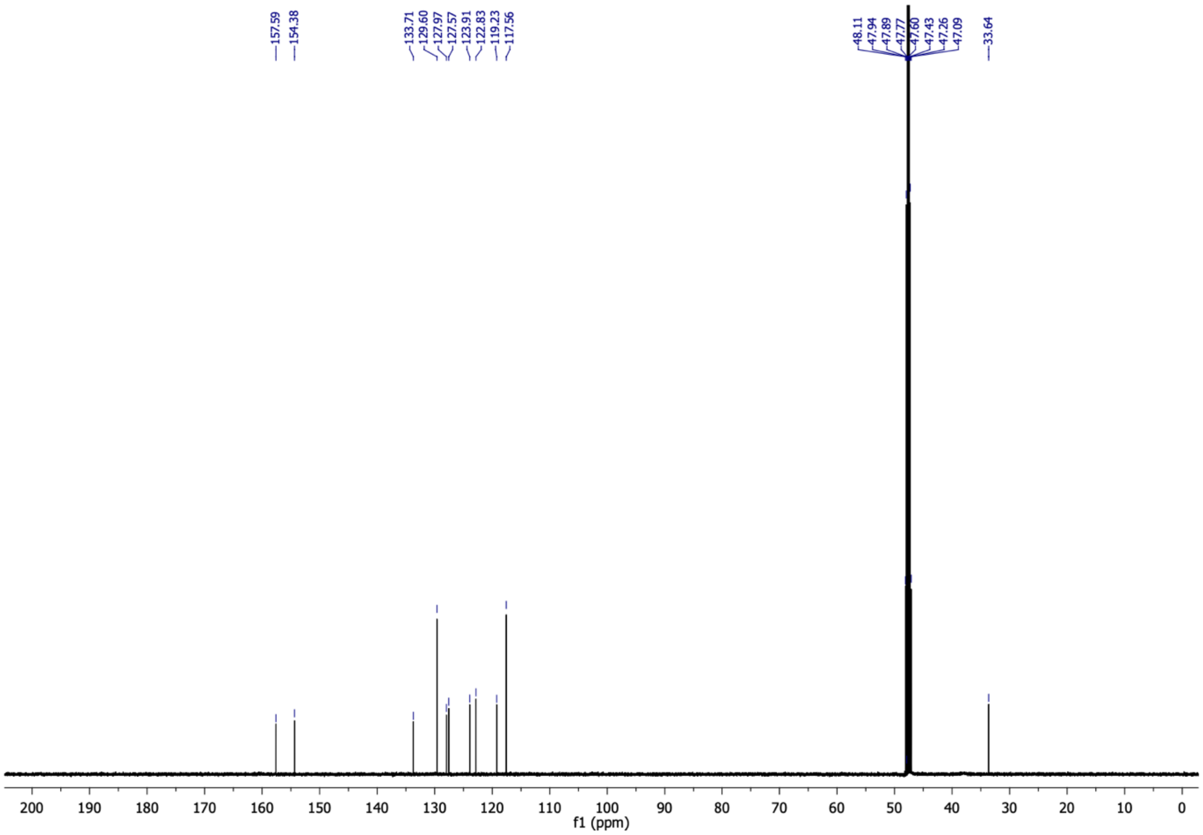

4-oxo-6-(2-phenoxyphenyl)-4,5,6,7-tetrahydrobenzofuran-3-carboxylic acid (**9**) $^{1}$H NMR (400 MHz, $CDCl_3$)

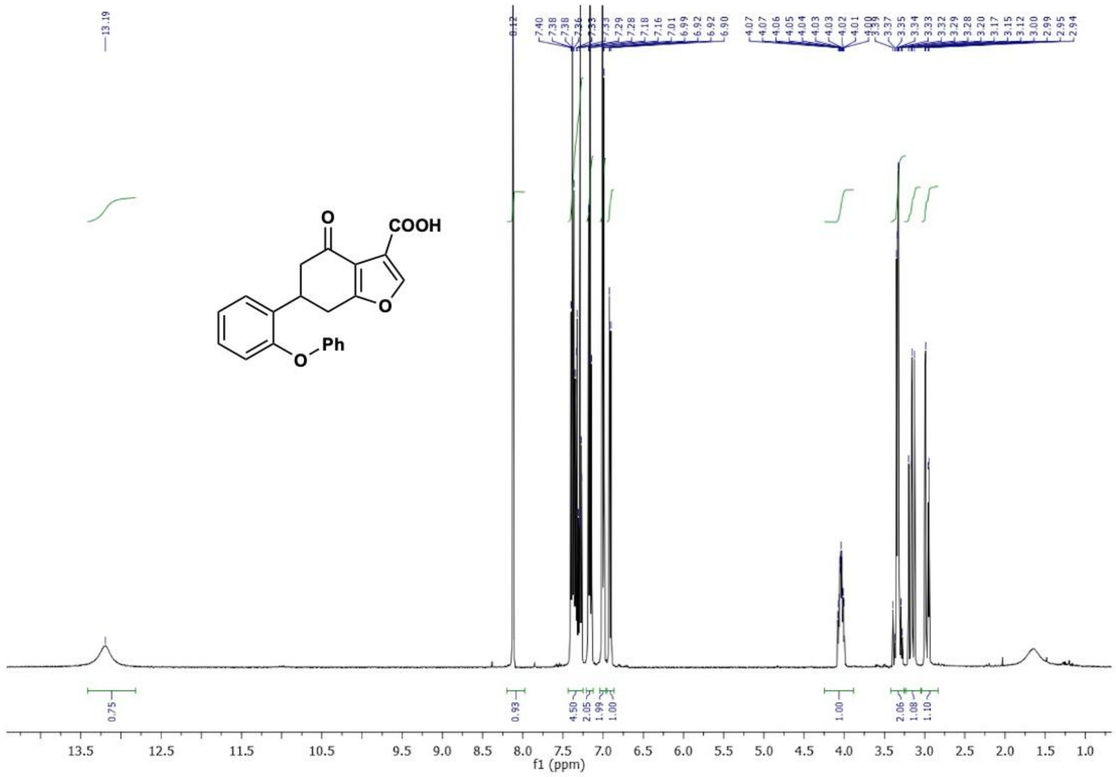

4-oxo-6-(2-phenoxyphenyl)-4,5,6,7-tetrahydrobenzofuran-3-carboxylic acid (**9**) $^{13}$C NMR (101 MHz, $CDCl_3$)

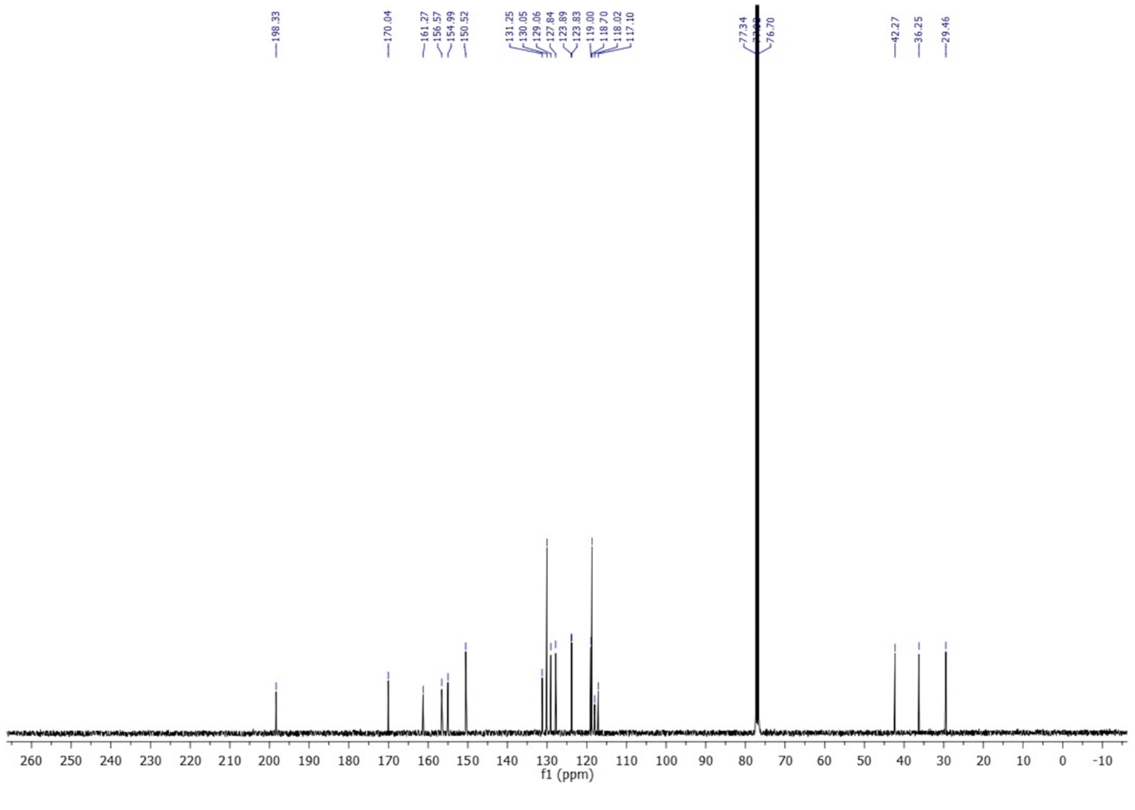

2-isopropoxybenzaldehyde (**10a**) $^{1}H$ NMR (500 MHz, $CDCl_3$)

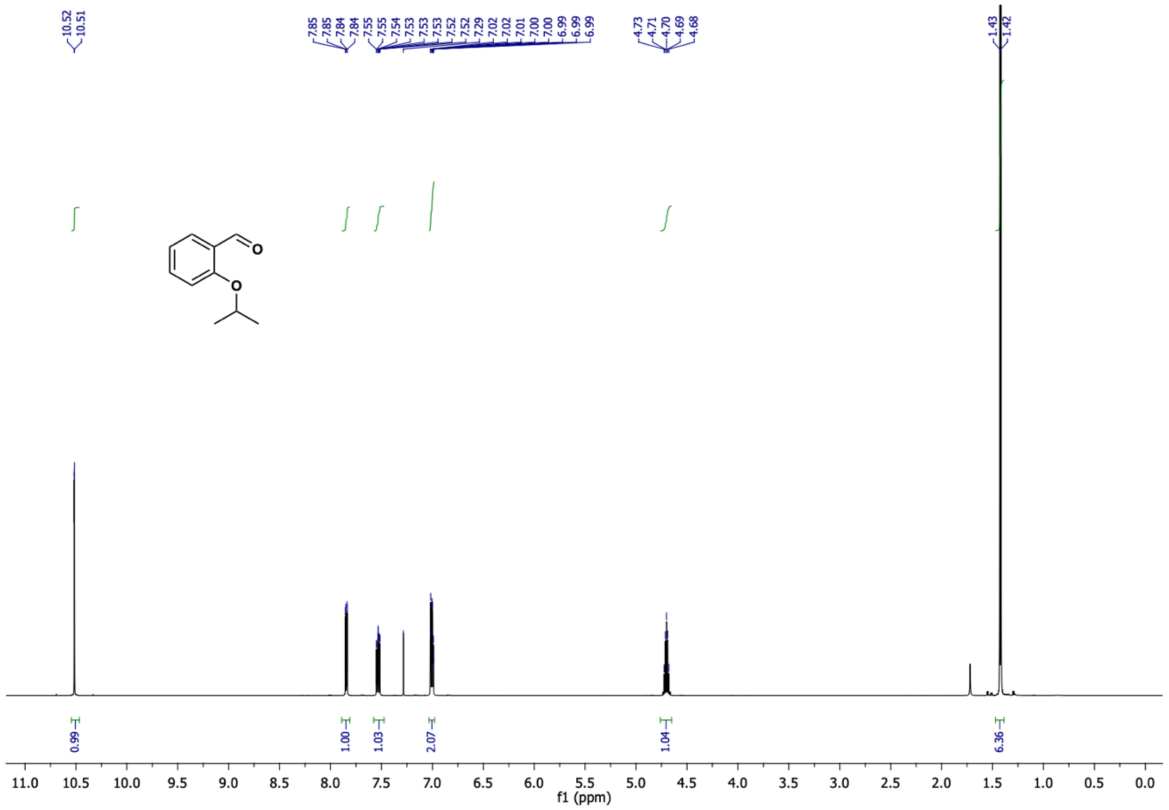

2-isopropoxybenzaldehyde (**10a**) $^{13}C$ NMR (126 MHz, $CDCl_3$)

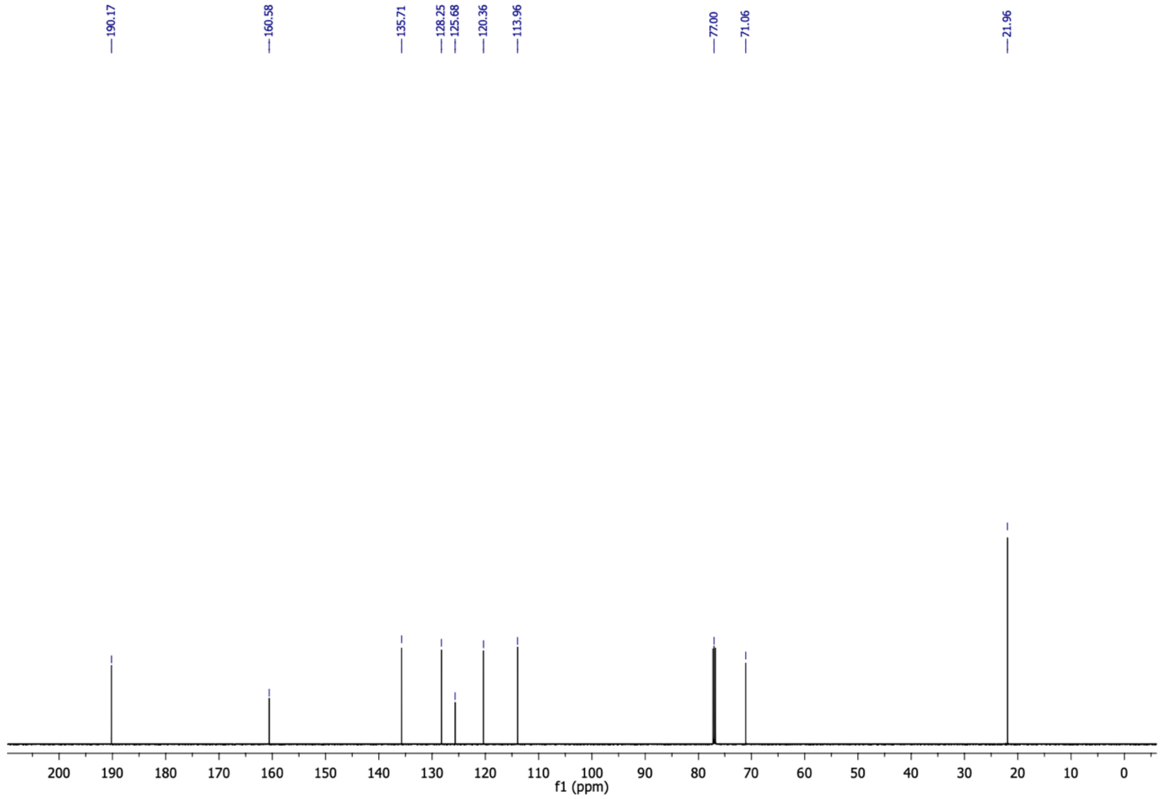

(E)-4-(2-isopropoxyphenyl)but-3-en-2-one (**10b**) $^{1}$H NMR (500 MHz, $CDCl_3$)

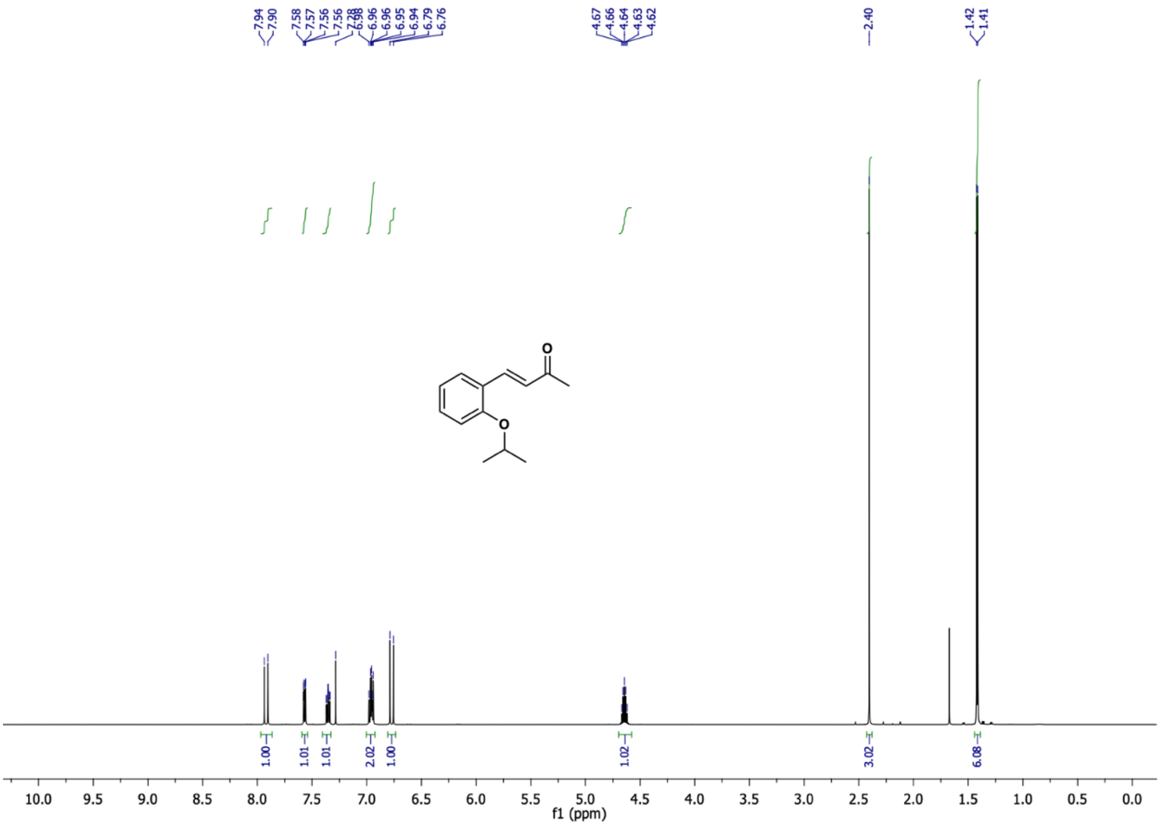

(E)-4-(2-isopropoxyphenyl)but-3-en-2-one (**10b**) $^{13}$C NMR (126 MHz, $CDCl_3$)

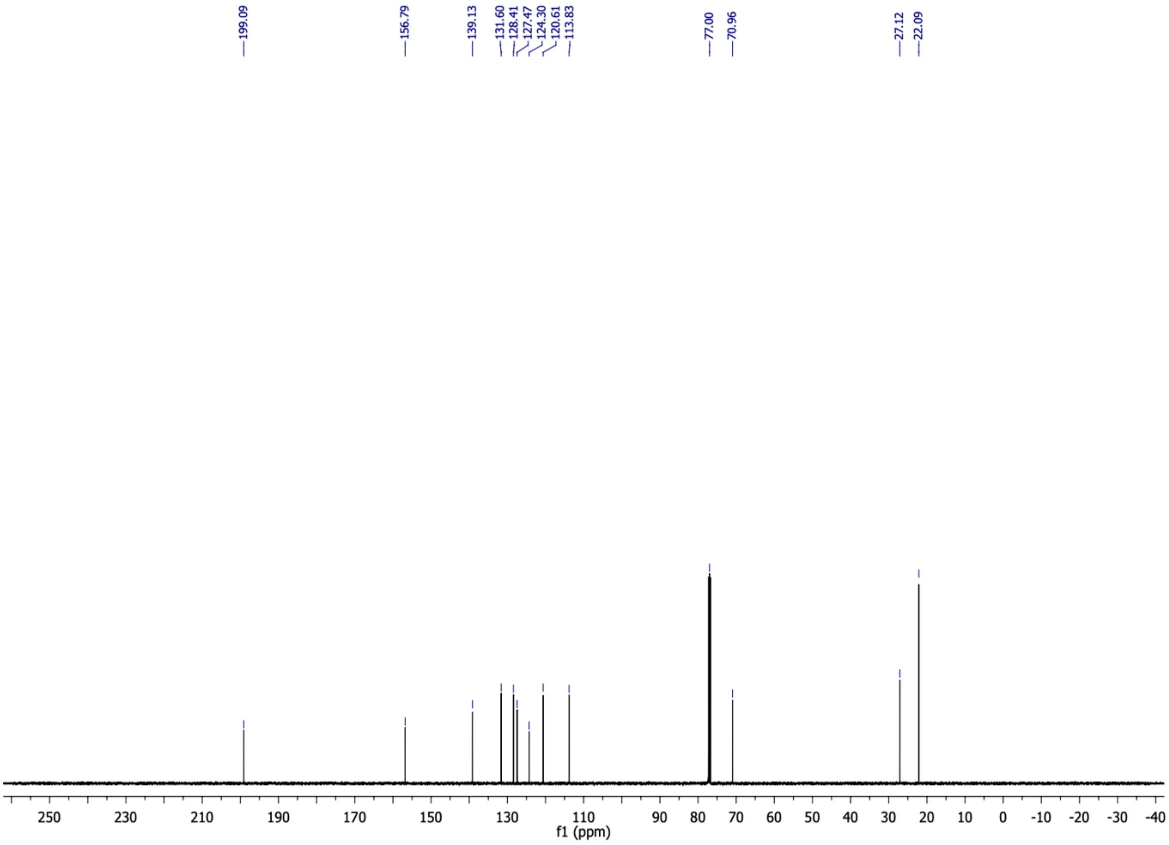

5-(2-isopropoxyphenyl)cyclohexane-1,3-dione (**10c**) $^{1}H$ NMR (500 MHz, $CD_3OD$)

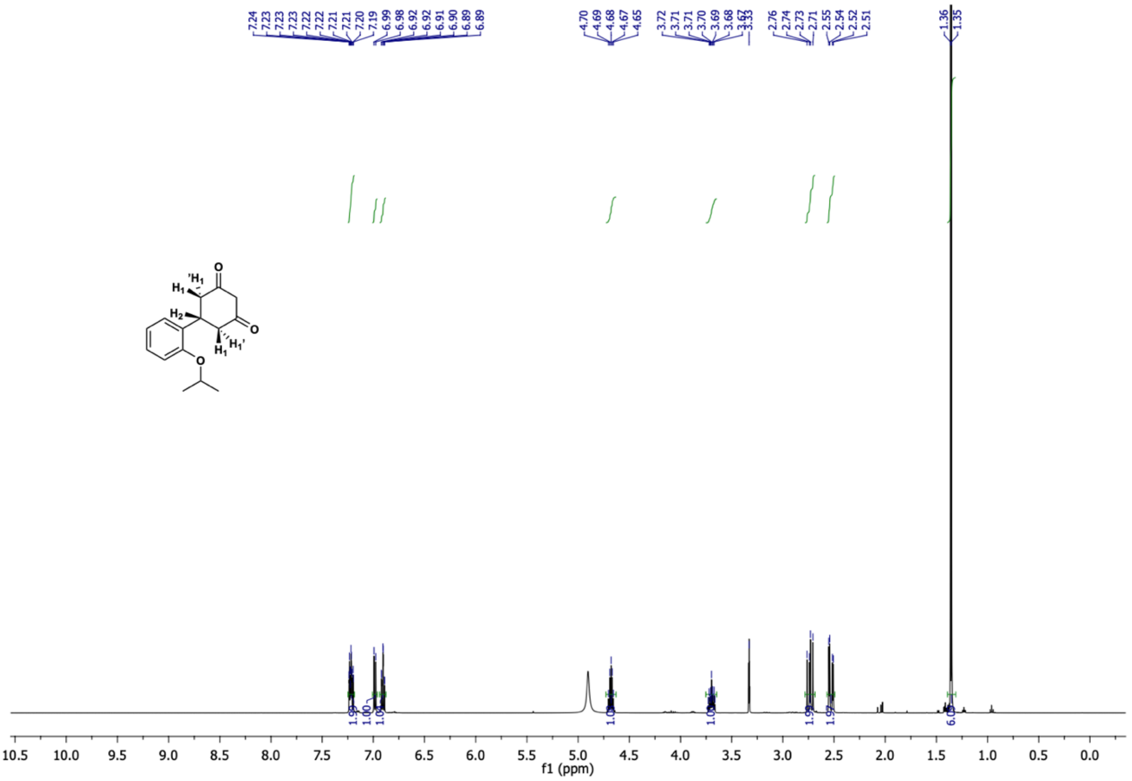

5-(2-isopropoxyphenyl)cyclohexane-1,3-dione (**10c**) $^{13}C$ NMR (126 MHz, $CD_3OD$)

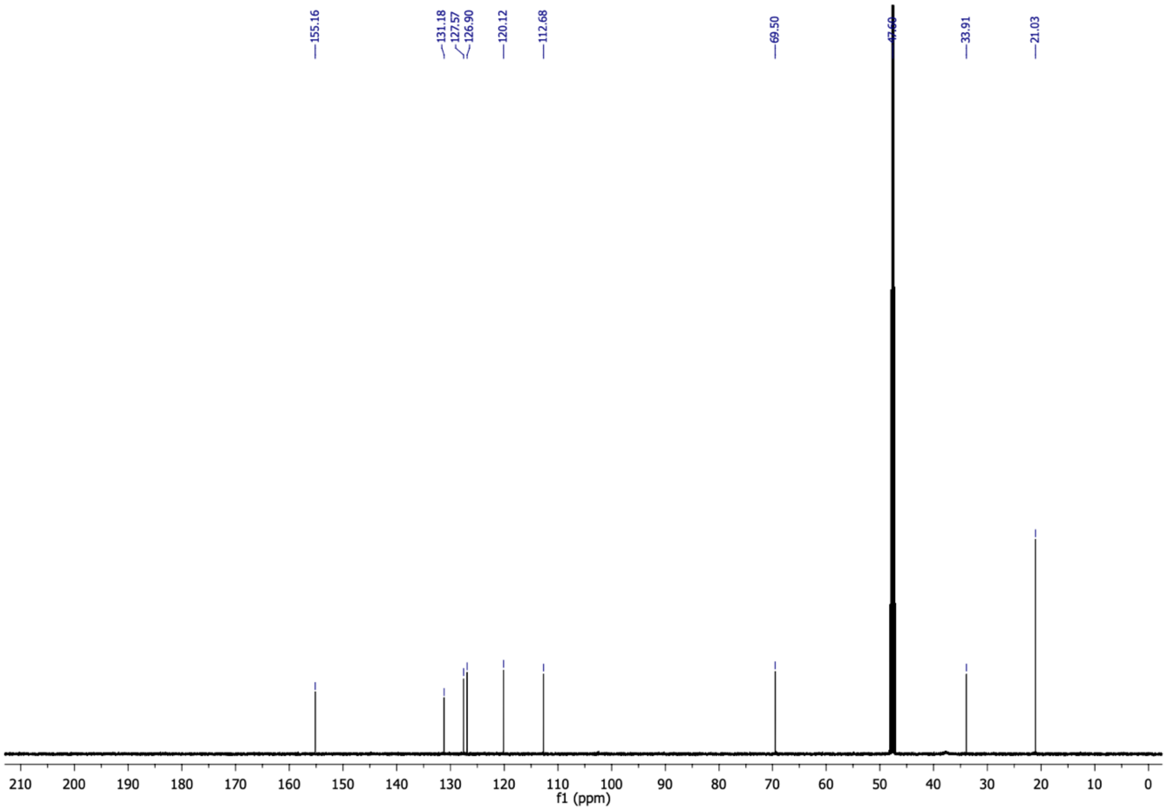

6-(2-isopropoxyphenyl)-4-oxo-4,5,6,7-tetrahydrobenzofuran-3-carboxylic acid (**10**) $^{1}H$ NMR (500 MHz, $CDCl_3$)

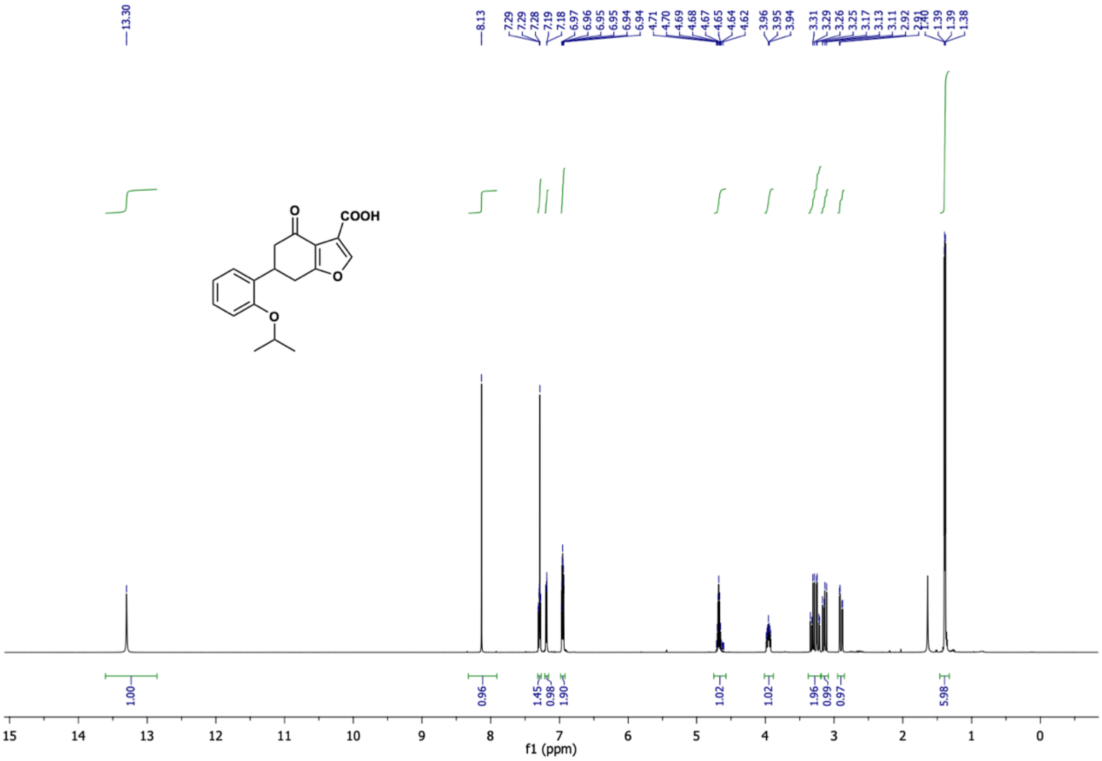

6-(2-isopropoxyphenyl)-4-oxo-4,5,6,7-tetrahydrobenzofuran-3-carboxylic acid (**10**) $^{13}C$ NMR (126 MHz, $CDCl_3$)

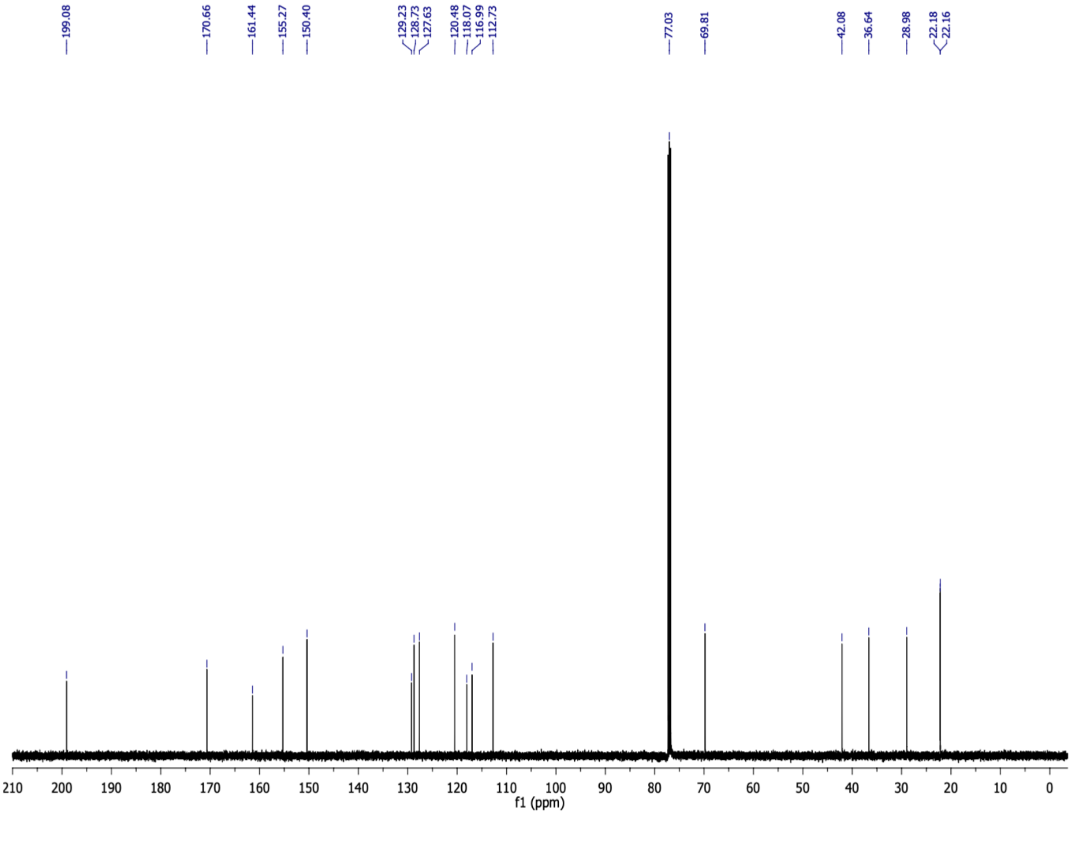

(*E*)-4-(2-methoxyphenyl)but-3-en-2-one (**11b**) $^{1}$H NMR (500 MHz, $CDCl_3$)

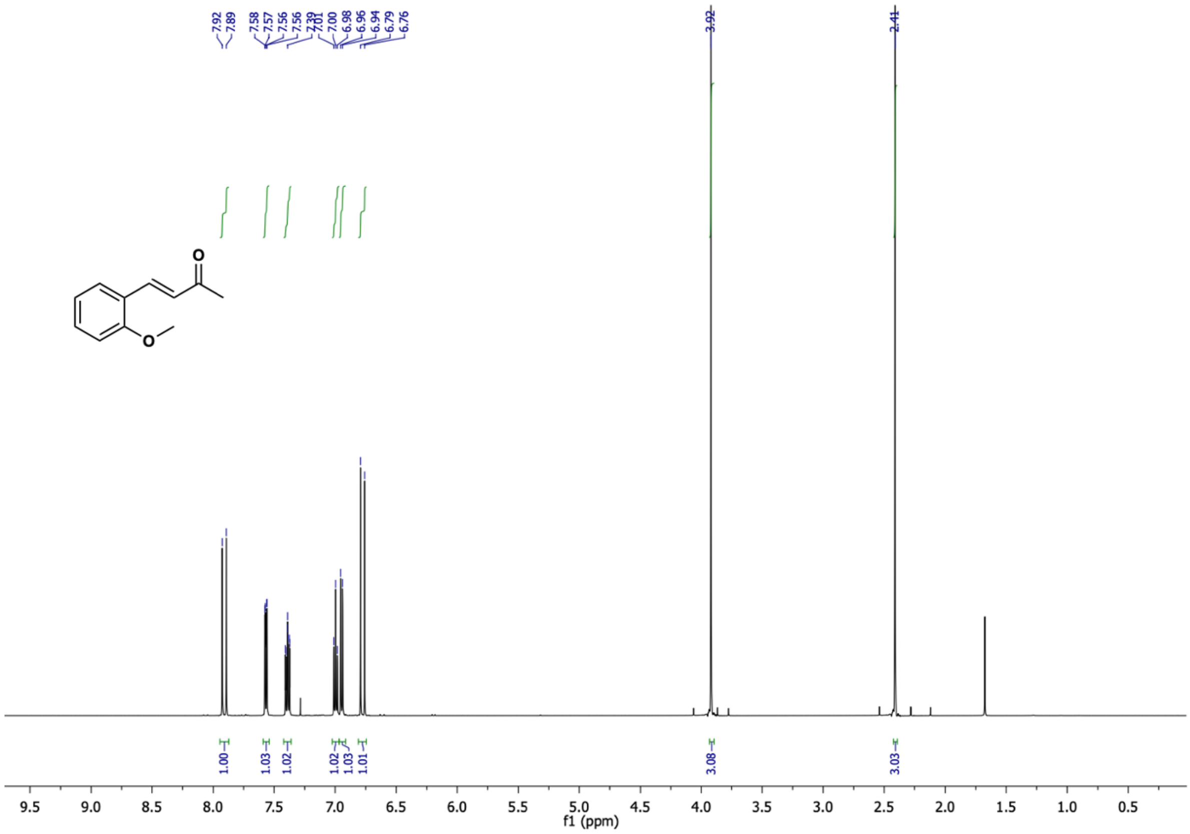

(*E*)-4-(2-methoxyphenyl)but-3-en-2-one (**11b**) $^{13}$C NMR (126 MHz, $CDCl_3$)

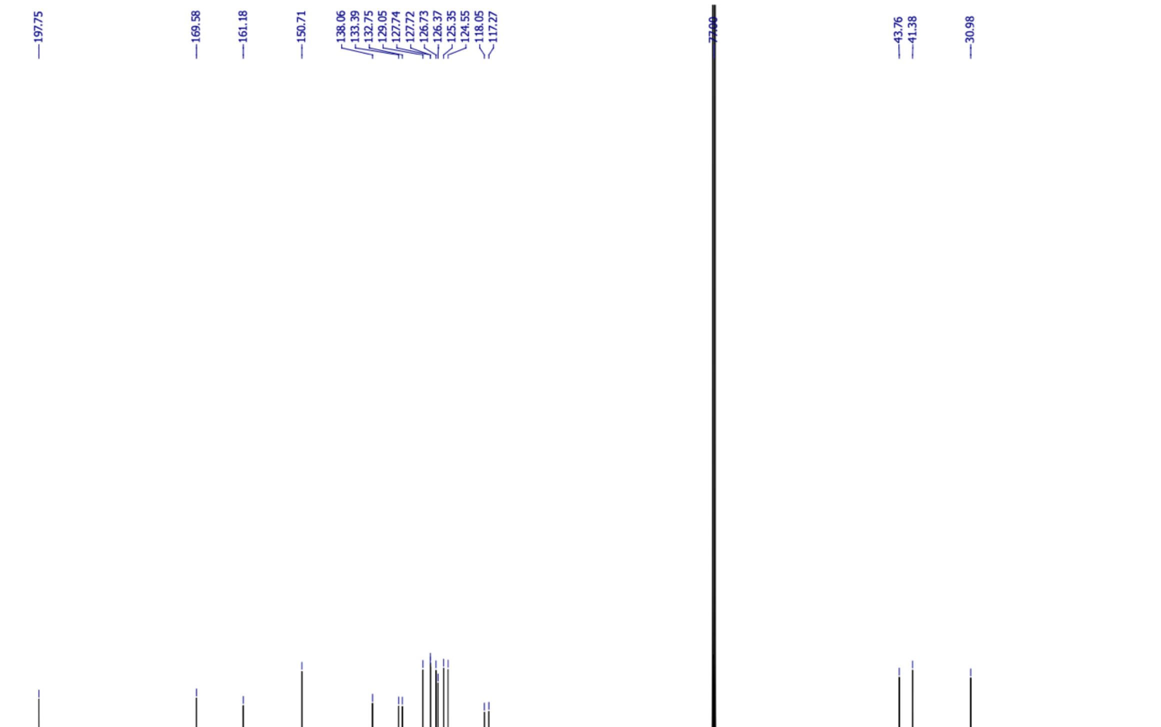

5-(2-methoxyphenyl)cyclohexane-1,3-dione (**11c**) $^{1}H$ NMR (500 MHz, $CD_3OD$)

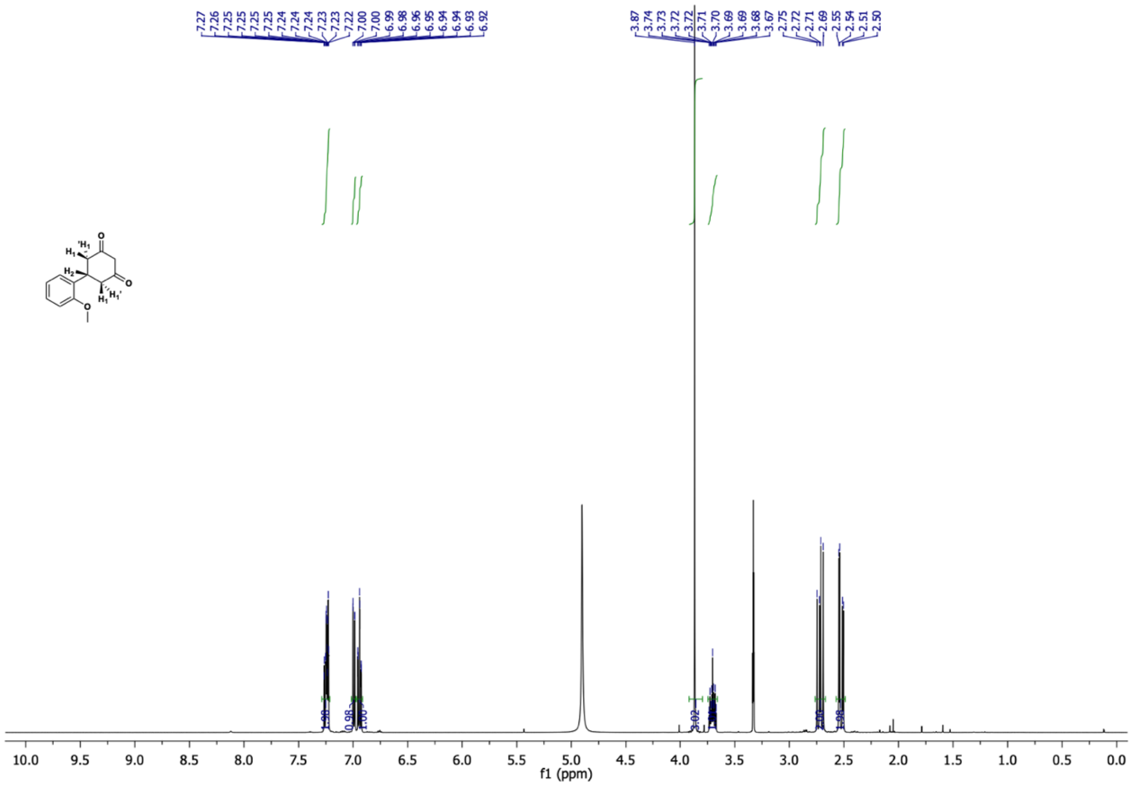

5-(2-methoxyphenyl)cyclohexane-1,3-dione (**11c**) $^{13}C$ NMR (126 MHz, $CD_3OD$)

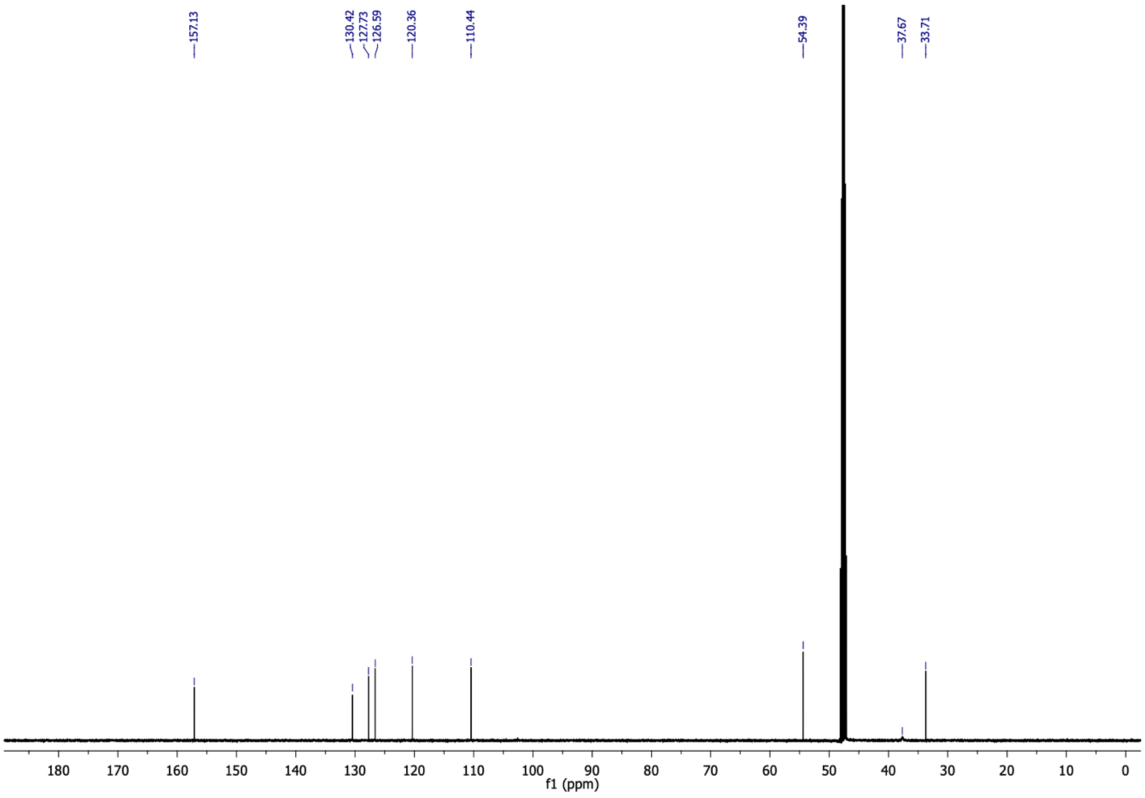

6-(2-methoxyphenyl)-4-oxo-4,5,6,7-tetrahydrobenzofuran-3-carboxylic acid (**11**) $^{1}$**H NMR** (500 MHz, $CDCl_3$)

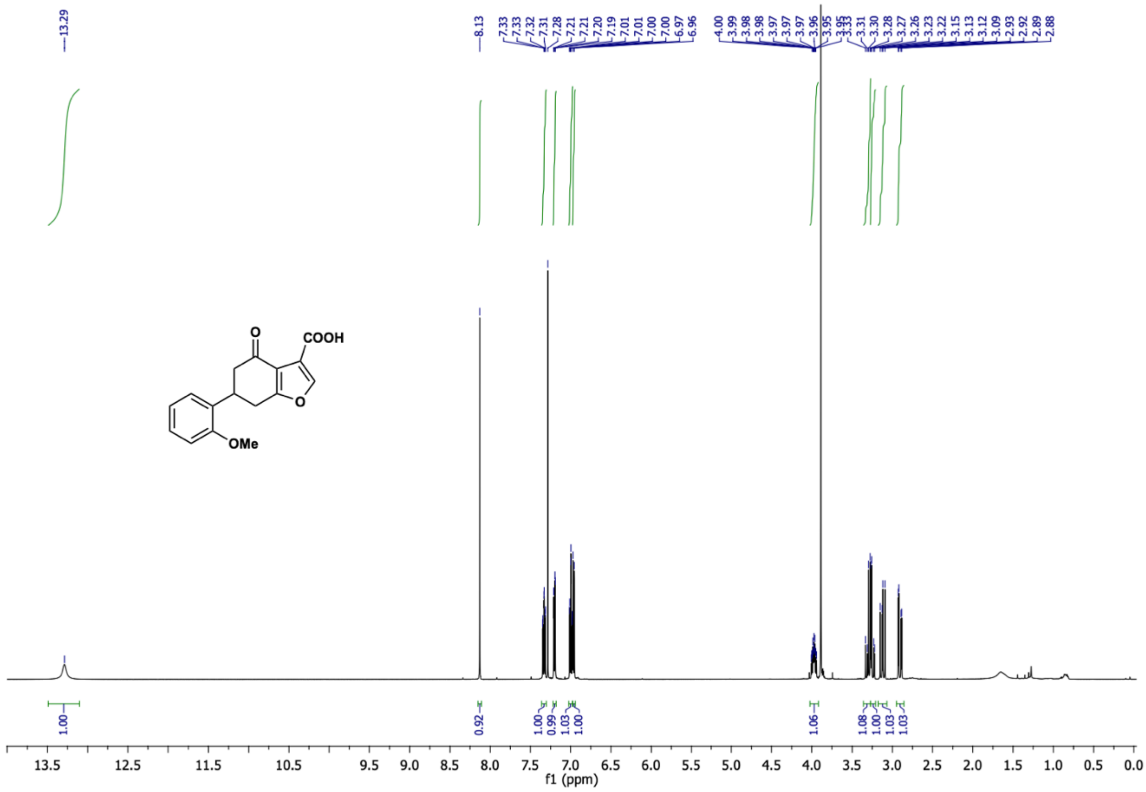

6-(2-methoxyphenyl)-4-oxo-4,5,6,7-tetrahydrobenzofuran-3-carboxylic acid (**11**) $^{13}$C NMR (126 MHz, $CDCl_3$)

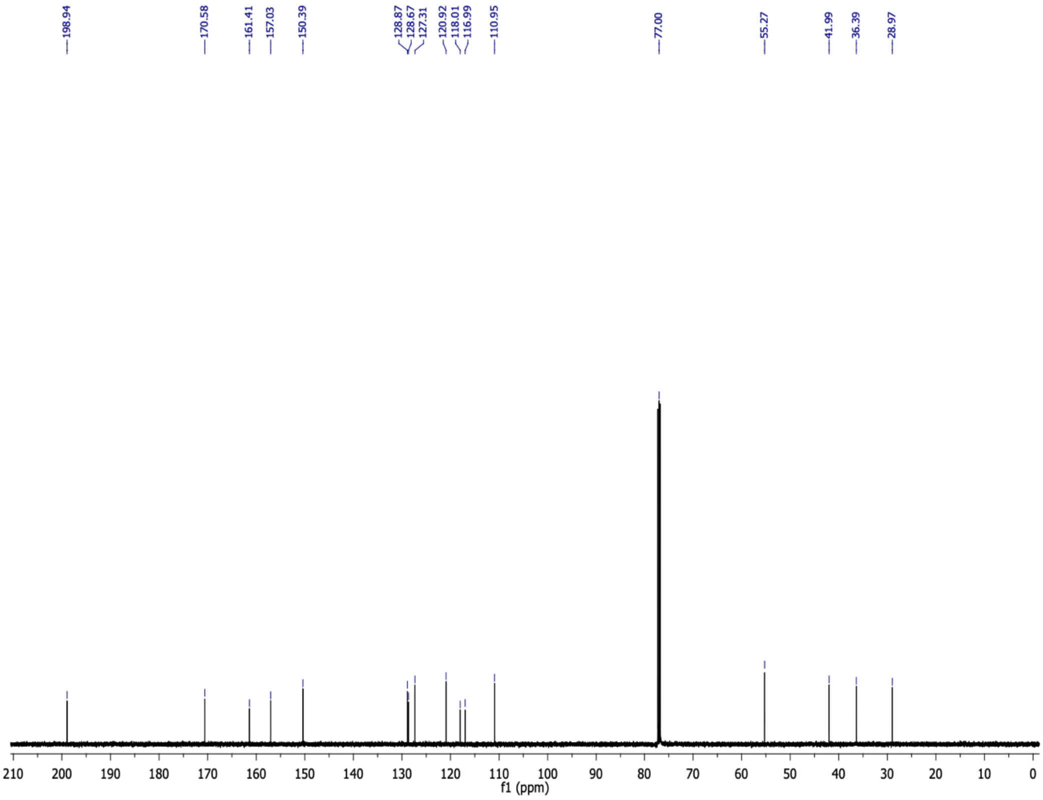

(*E*)-4-(pyridin-2-yl)but-3-en-2-one (**12b**) $^{1}H$ NMR (500 MHz, $CDCl_3$)

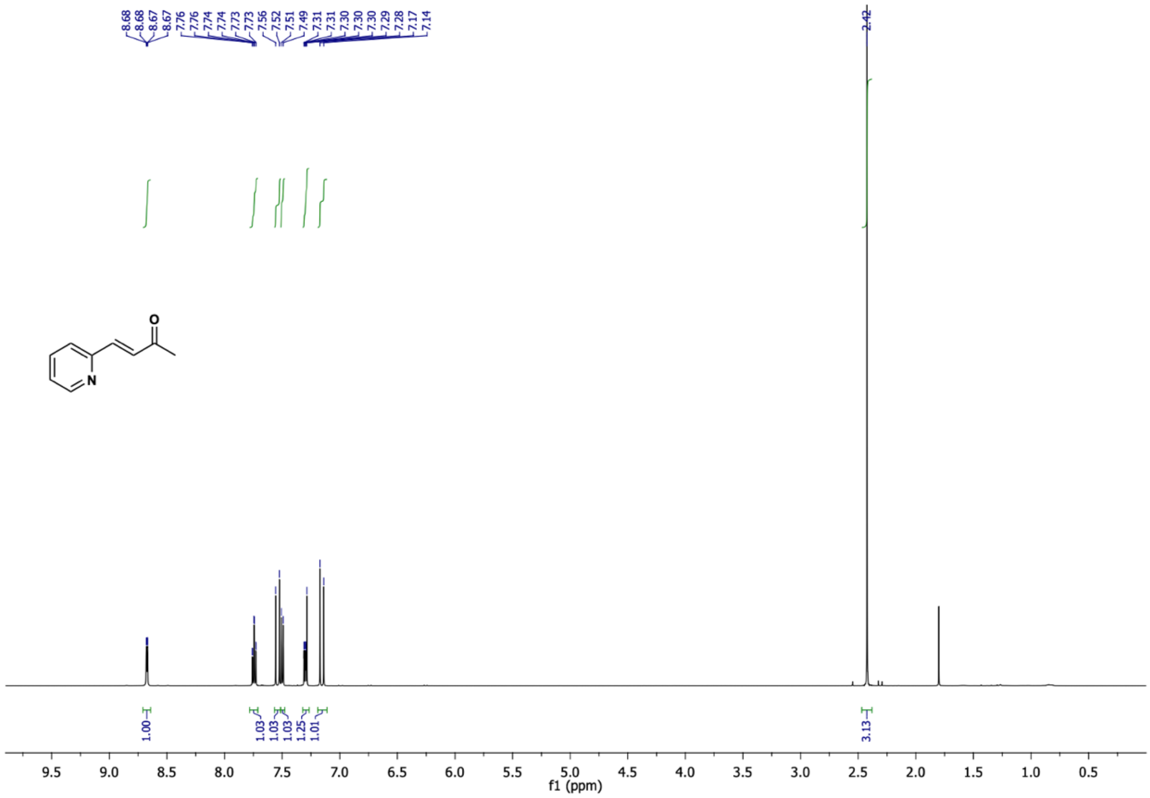

(*E*)-4-(pyridin-2-yl)but-3-en-2-one (**12b**) $^{13}C$ NMR (126 MHz, $CDCl_3$)

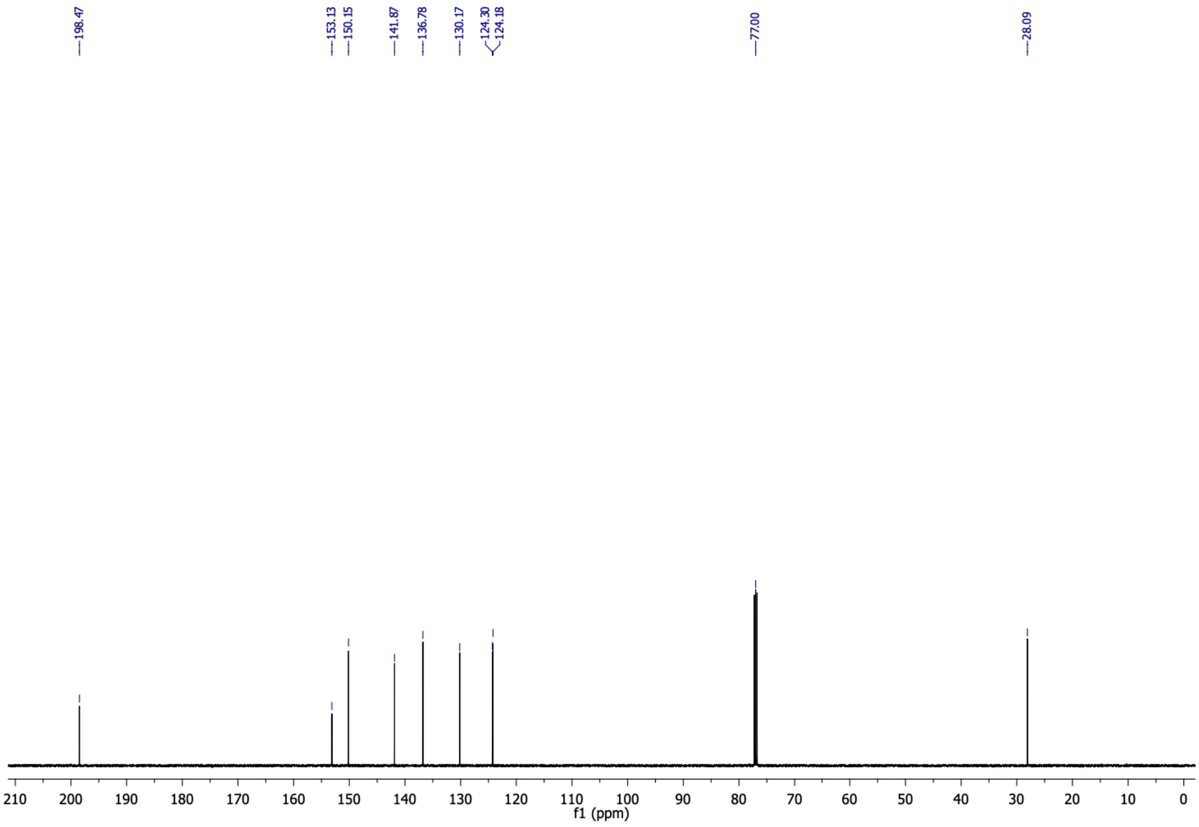

5-(pyridin-2-yl)cyclohexane-1,3-dione (**12c**) $^{1}$H NMR (500 MHz, $CD_3OD$)

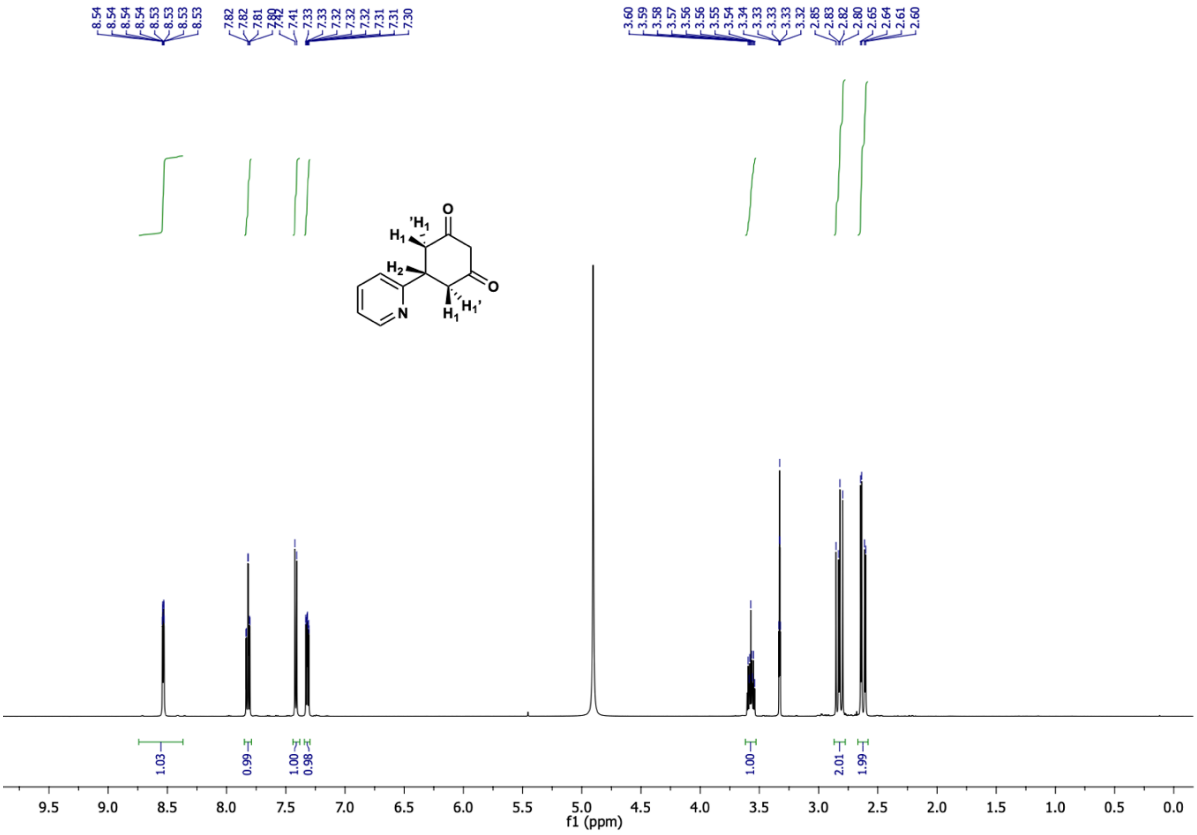

5-(pyridin-2-yl)cyclohexane-1,3-dione (**12c**) $^{13}$C NMR (126 MHz, $CD_3OD$)

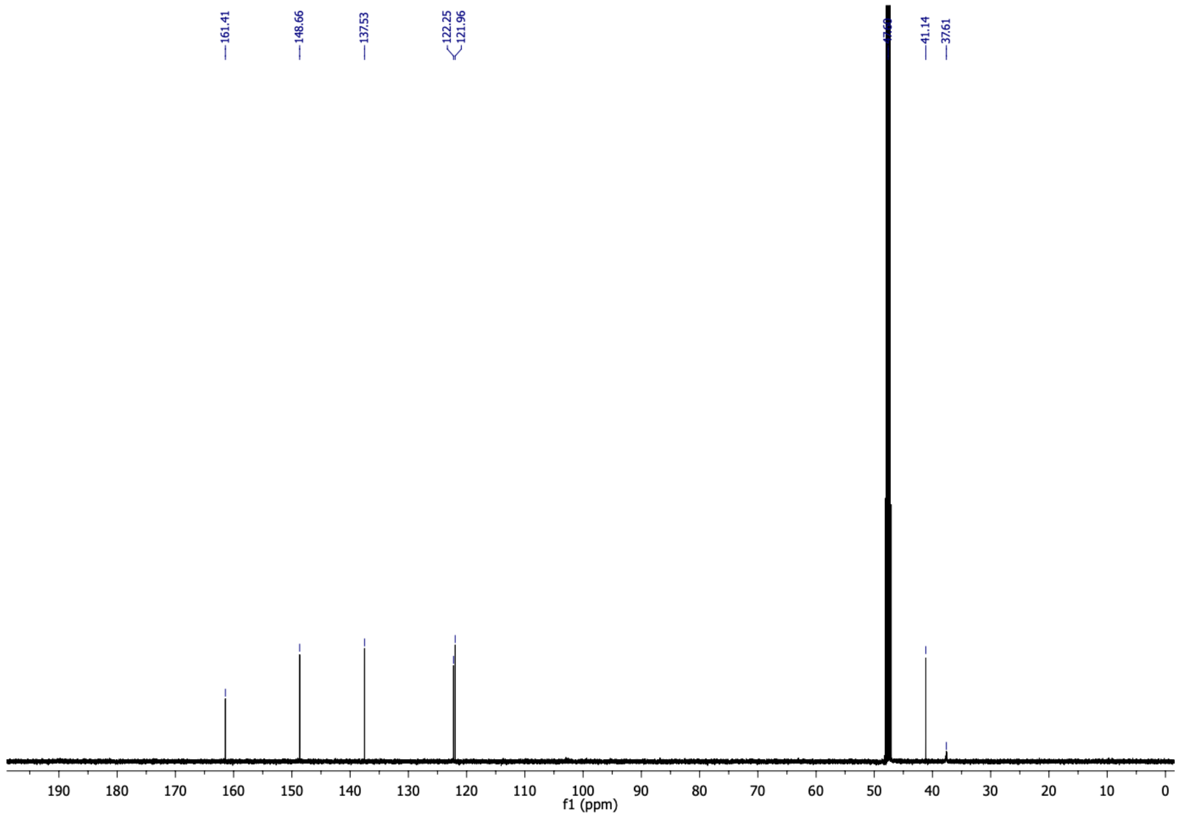

4-oxo-6-(pyridin-2-yl)-4,5,6,7-tetrahydrobenzofuran-3-carboxylic acid (**12**) $^{1}H$ NMR (500 MHz, $CDCl_3$)

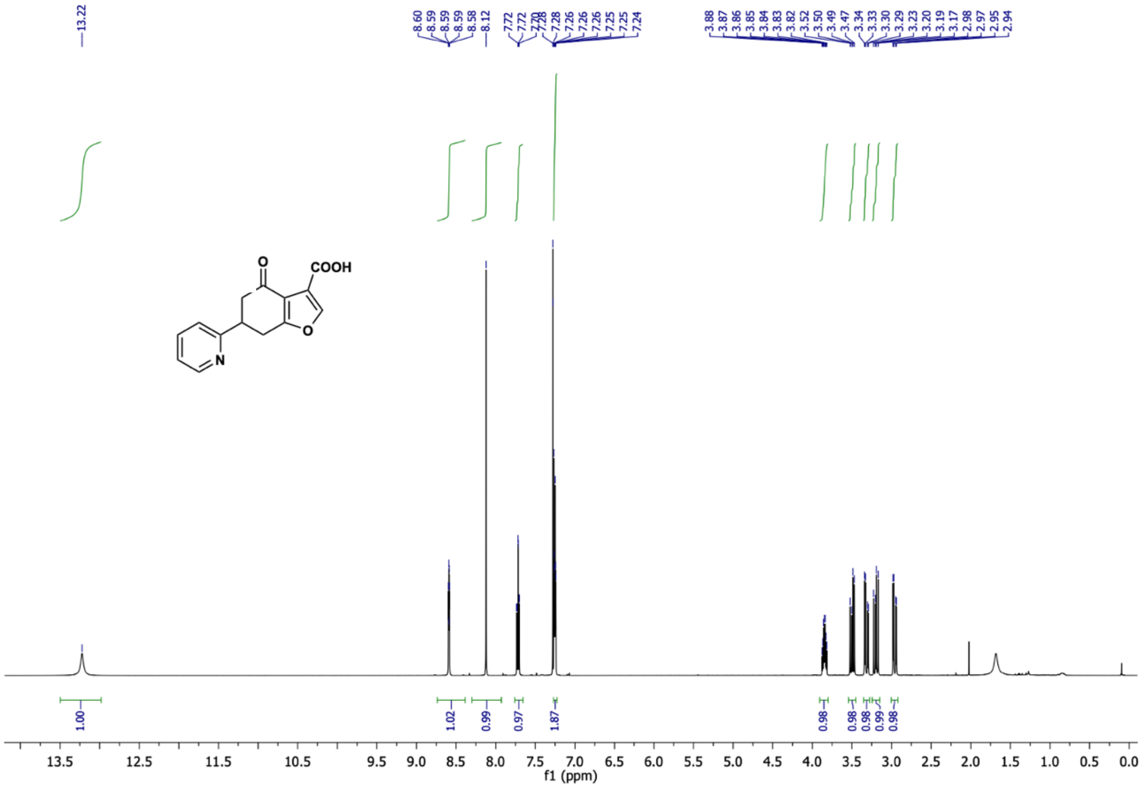

4-oxo-6-(pyridin-2-yl)-4,5,6,7-tetrahydrobenzofuran-3-carboxylic acid (**12**) $^{13}C$ NMR (126 MHz, $CDCl_3$)

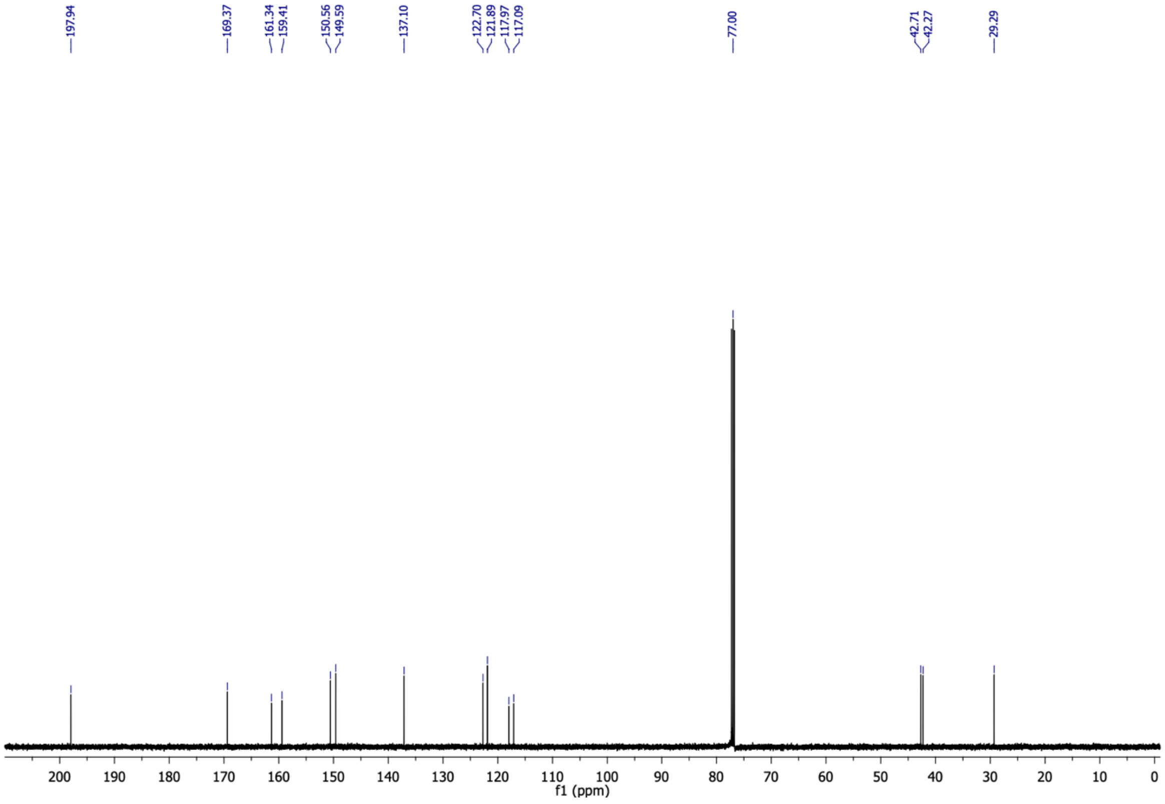

2-(3-carboxy-4-oxo-4,5,6,7-tetrahydrobenzofuran-6-yl)pyridine 1-oxide (**13**) $^{1}$H NMR (500 MHz, $CDCl_3$)

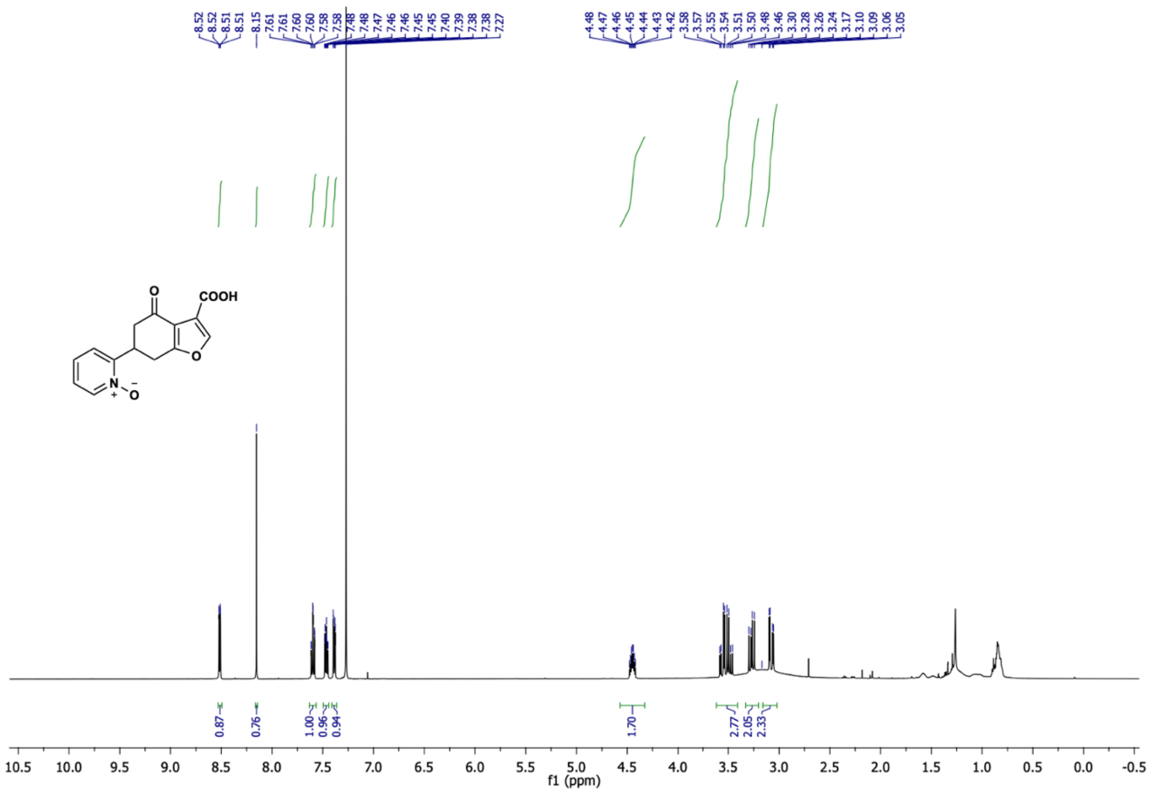

2-(3-carboxy-4-oxo-4,5,6,7-tetrahydrobenzofuran-6-yl)pyridine 1-oxide (**13**) $^{13}$C NMR (126 MHz, $CDCl_3$)

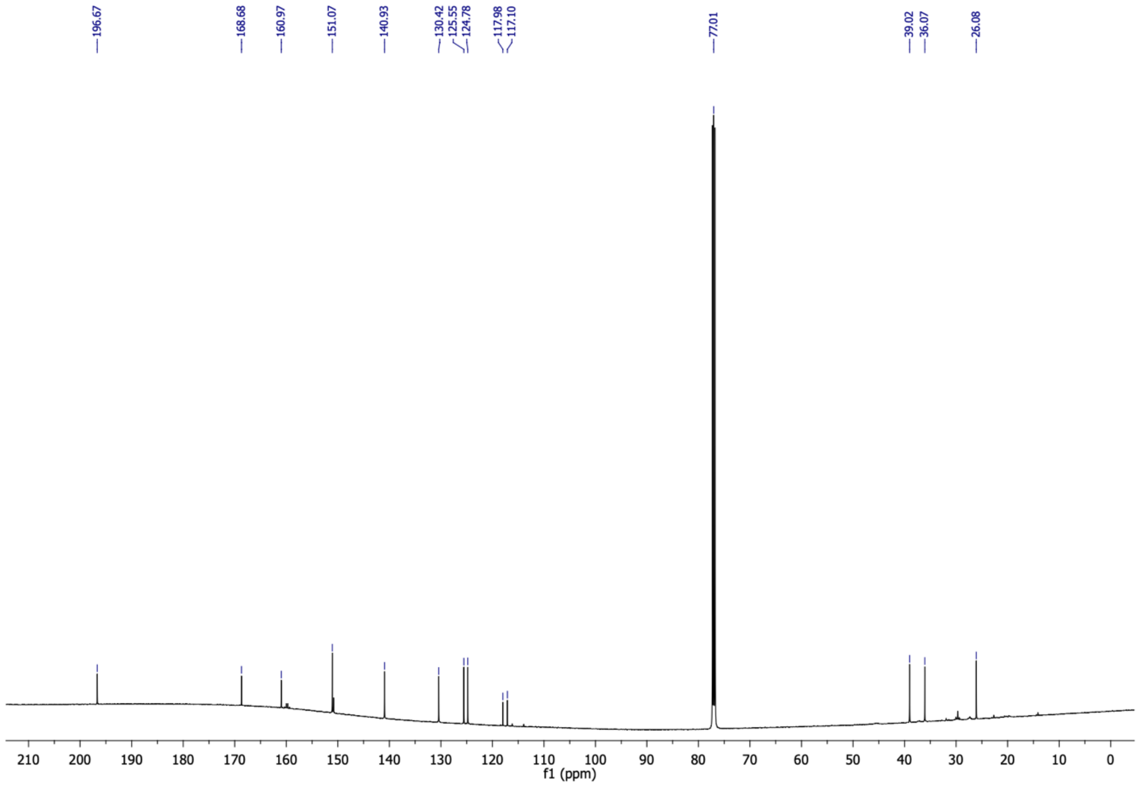

6-(2-hydroxyphenyl)-4-oxo-4,5,6,7-tetrahydrobenzofuran-3-carboxylic acid (**14**) $^{1}$H NMR (500 MHz, $CDCl_3$)

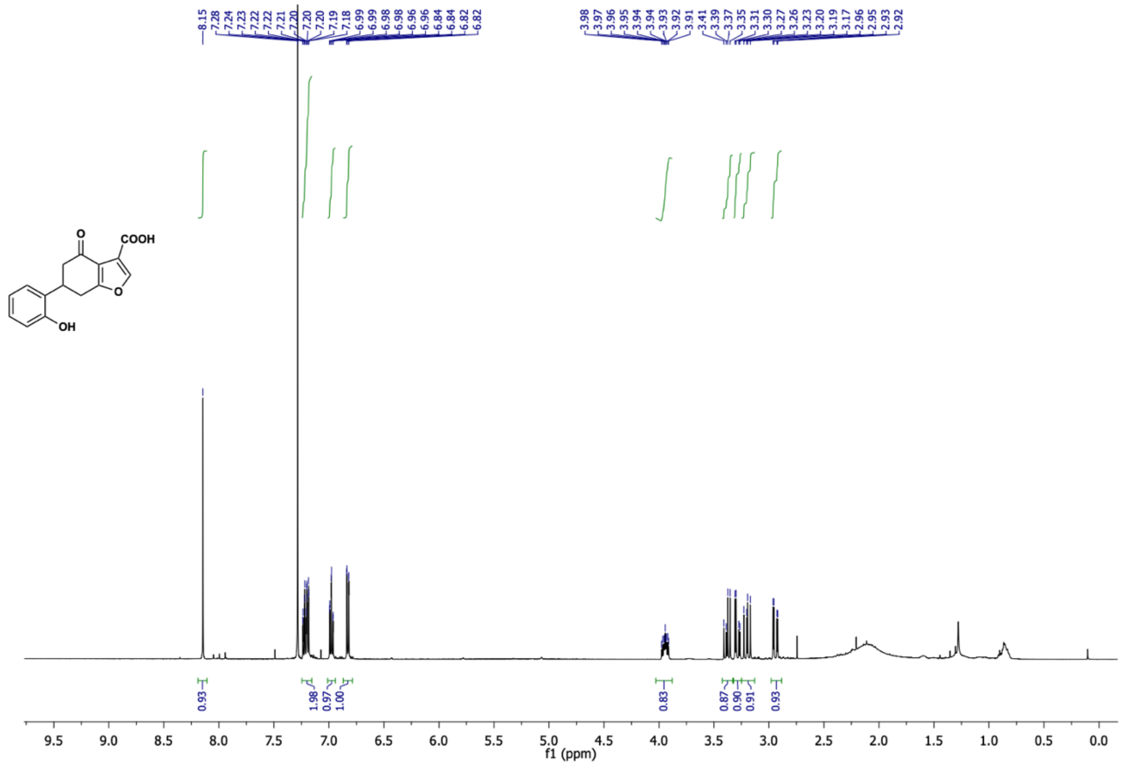

6-(2-hydroxyphenyl)-4-oxo-4,5,6,7-tetrahydrobenzofuran-3-carboxylic acid (**14**) $^{13}$C NMR (126 MHz, $CDCl_3$)

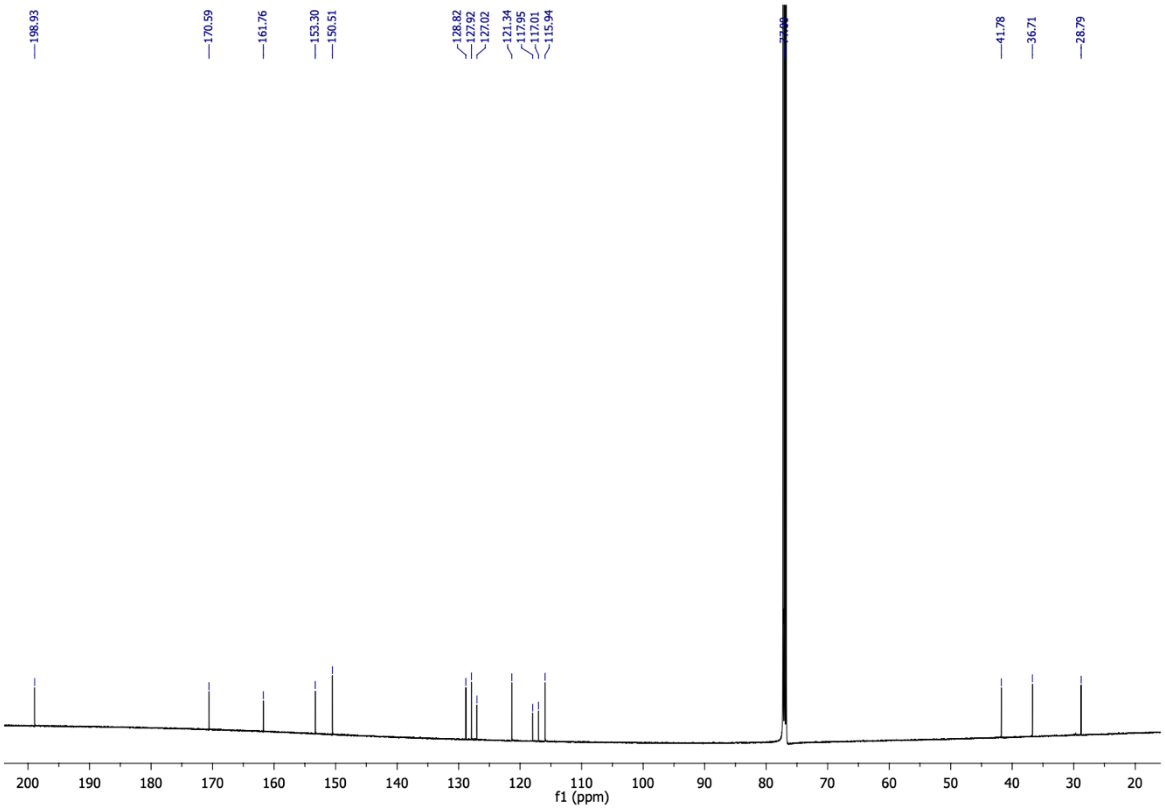

6-([1,1'-biphenyl]-2-yl)-4-oxo-4,5,6,7-tetrahydrobenzofuran-3-carboxylic acid (**15**) $^{1}$H NMR (500 MHz, $CDCl_3$)

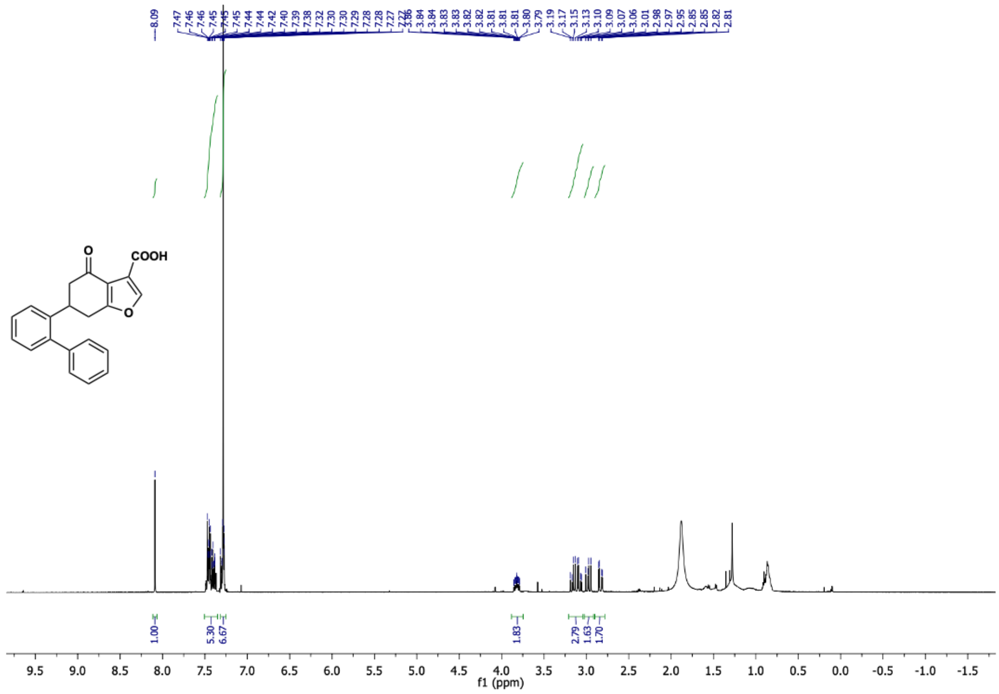

6-([1,1'-biphenyl]-2-yl)-4-oxo-4,5,6,7-tetrahydrobenzofuran-3-carboxylic acid (**15**) $^{13}$C NMR (126 MHz, $CDCl_3$)

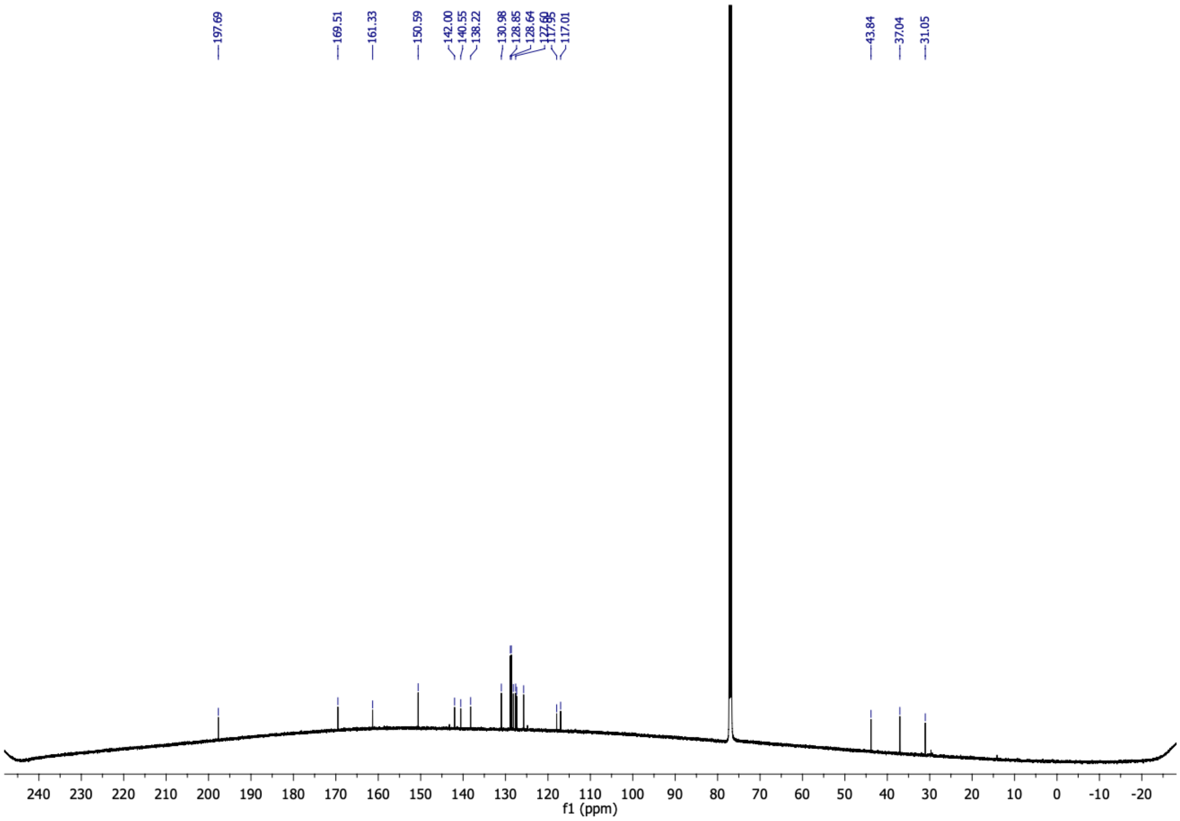

2-bromo-6-methoxybenzaldehyde (**16a**) $^{1}$H NMR (500 MHz, $CDCl_3$)

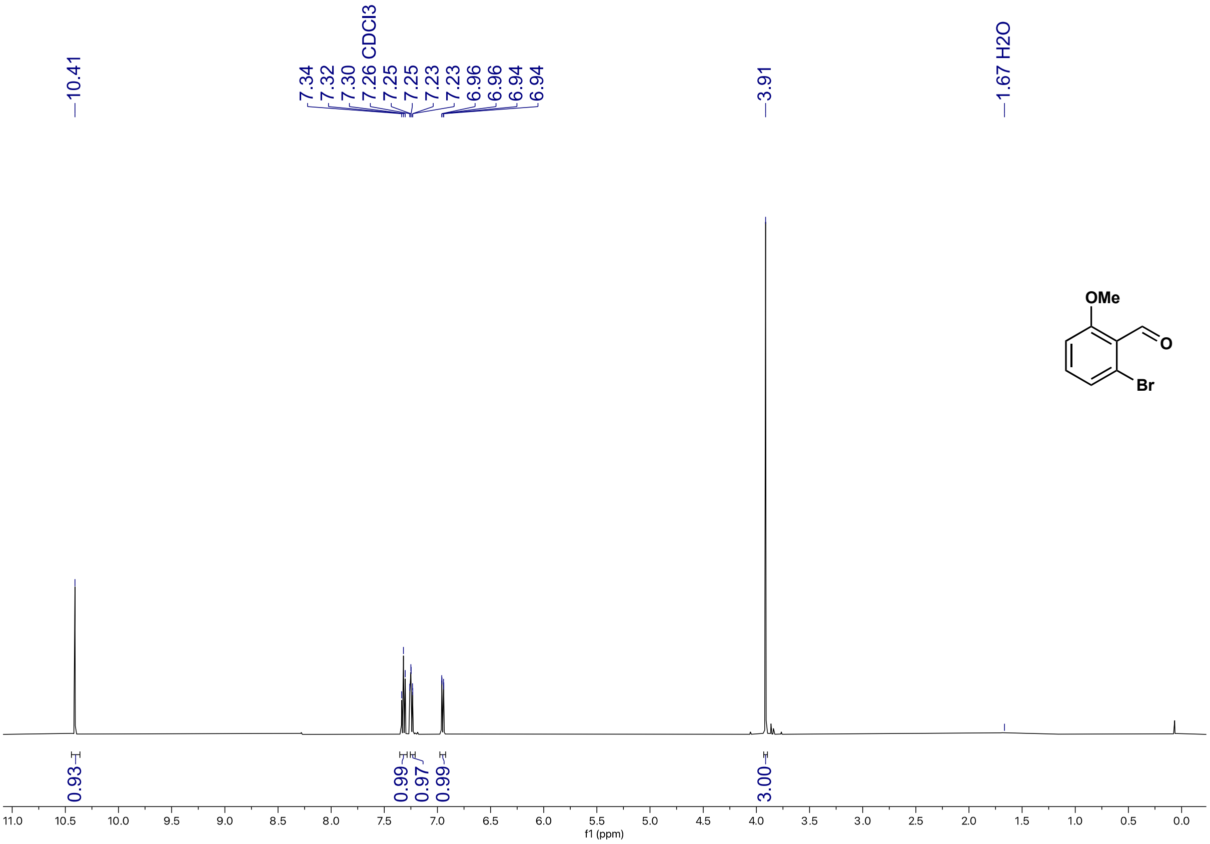

2-bromo-6-methoxybenzaldehyde (**16a**) $^{13}$C NMR (126 MHz, $CDCl_3$)

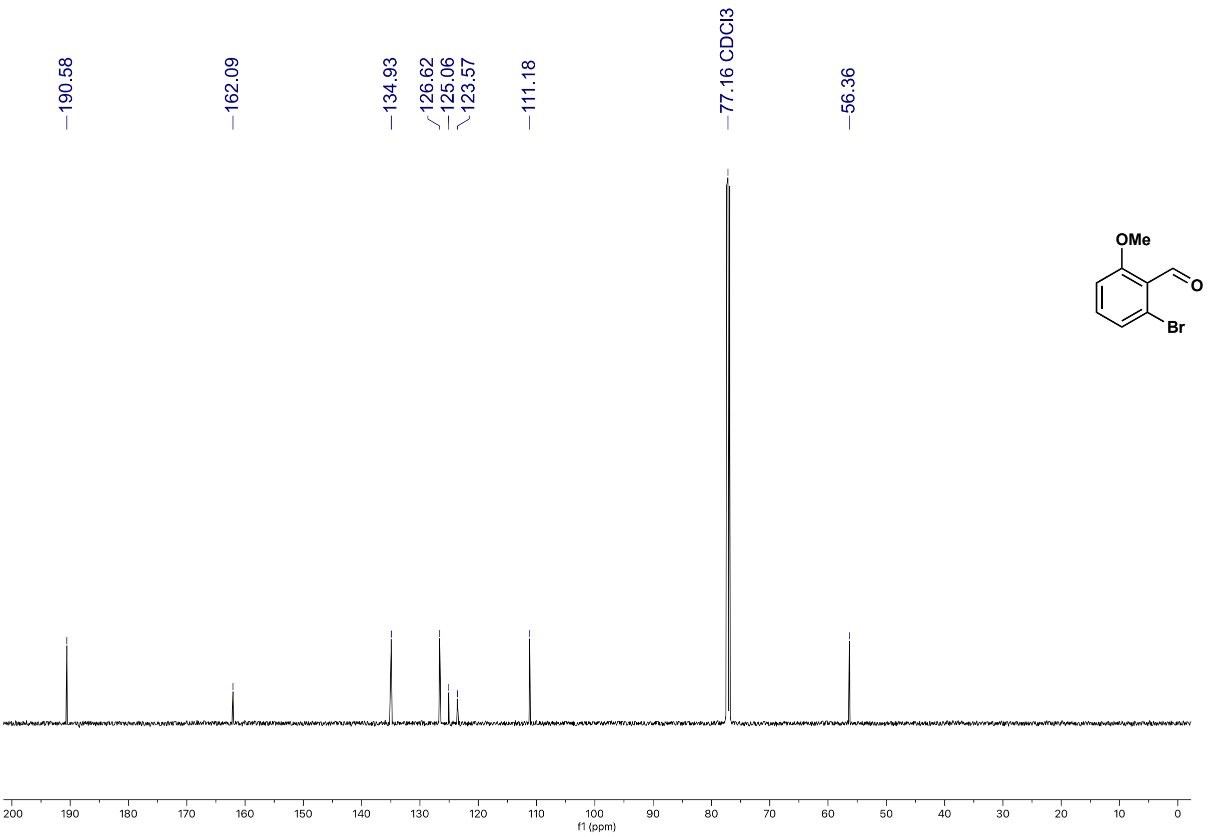

(E)-4-(2-bromo-6-methoxyphenyl)but-3-en-2-one (**16b**) $^{1}H$ NMR (500 MHz, $CDCl_3$)

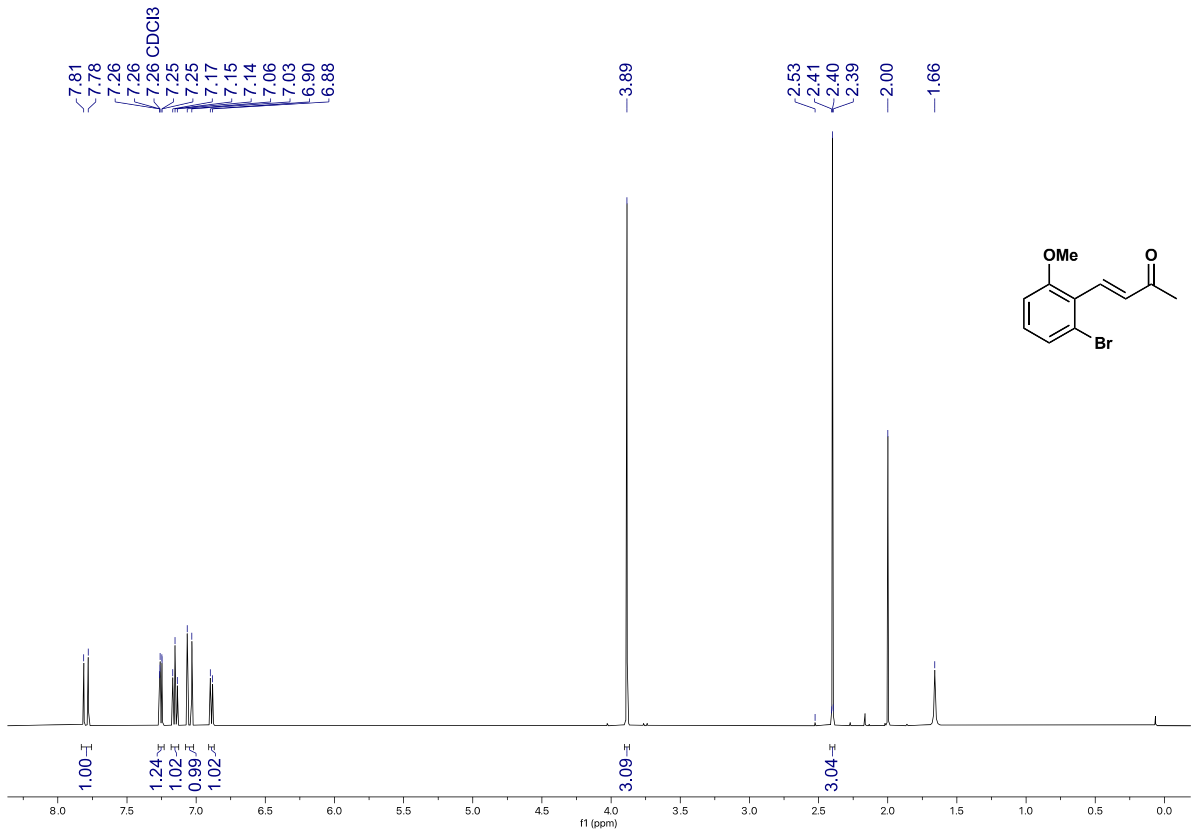

(E)-4-(2-bromo-6-methoxyphenyl)but-3-en-2-one (**16b**) $^{13}C$ NMR (126 MHz, $CDCl_3$)

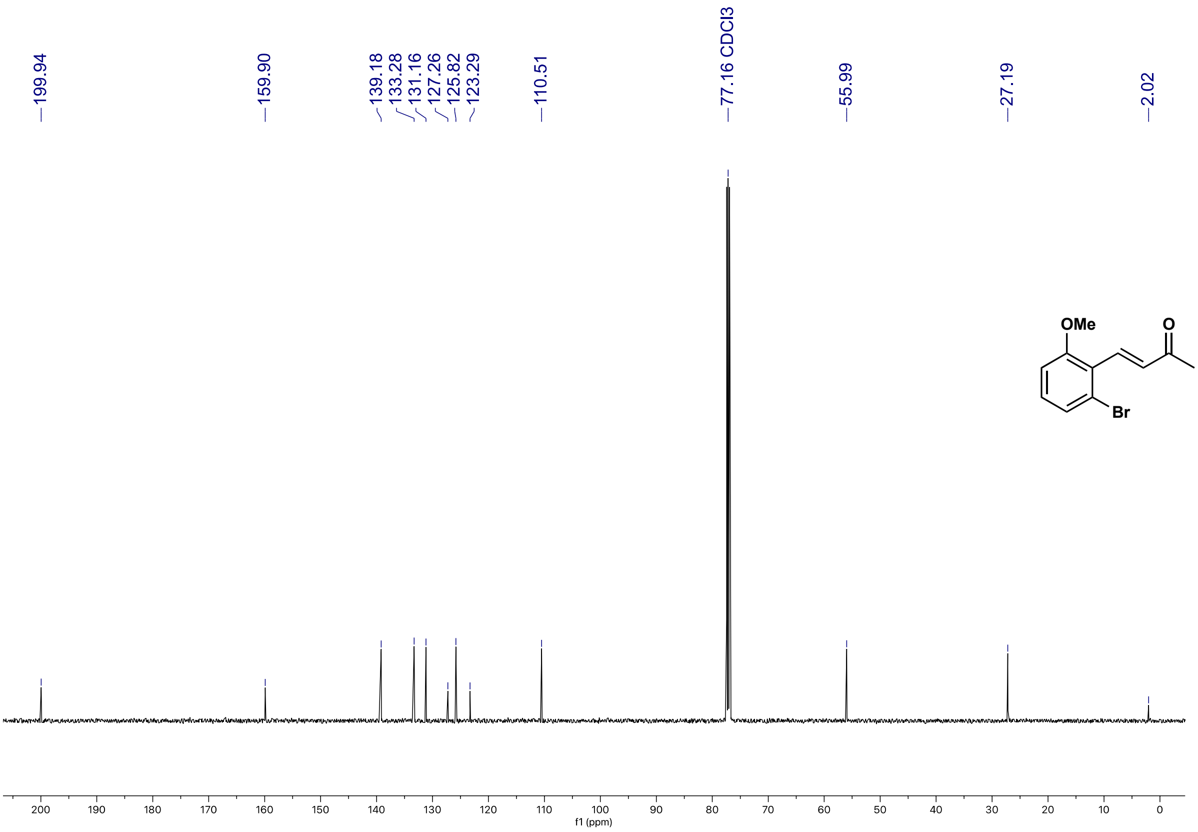

5-(2-bromo-6-methoxyphenyl)cyclohexane-1,3-dione (**16c**) $^{1}$H NMR (500 MHz, $d_6$-DMSO)

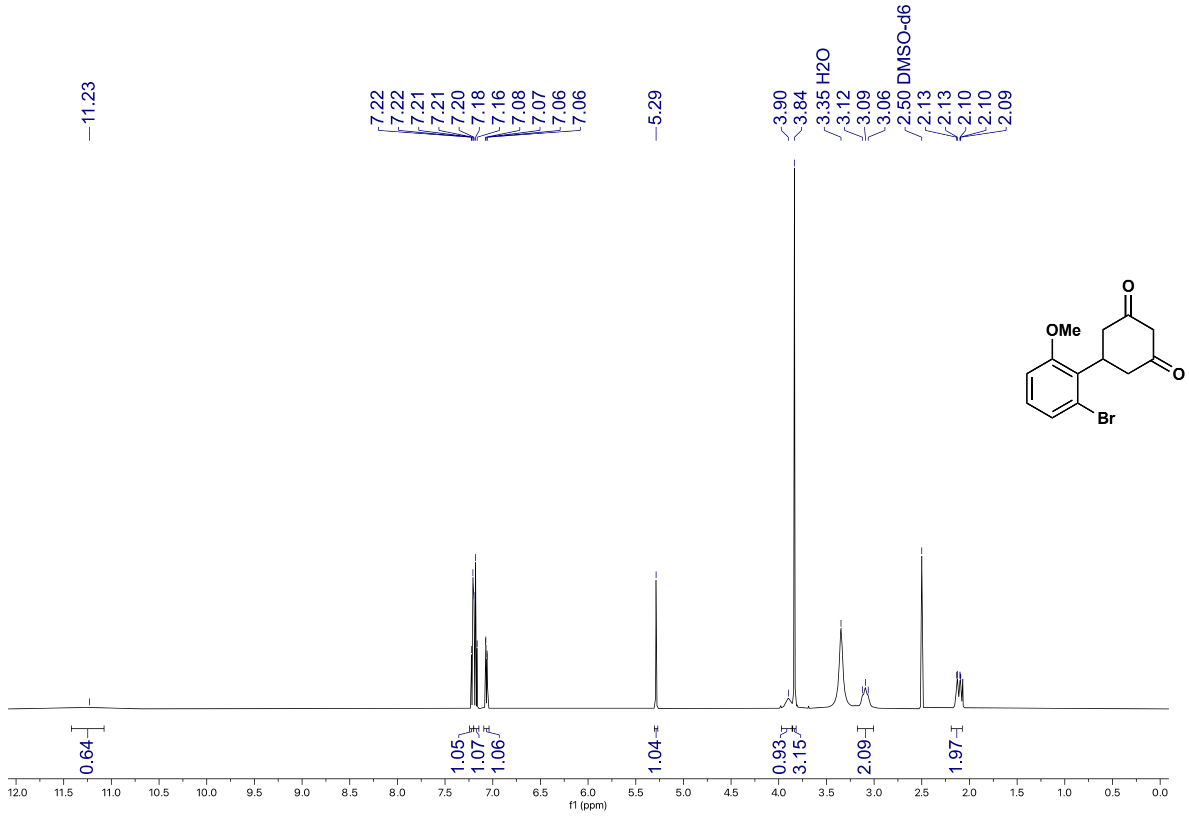

5-(2-bromo-6-methoxyphenyl)cyclohexane-1,3-dione (**16c**) $^{13}$C NMR (126 MHz, $d_6$-DMSO)

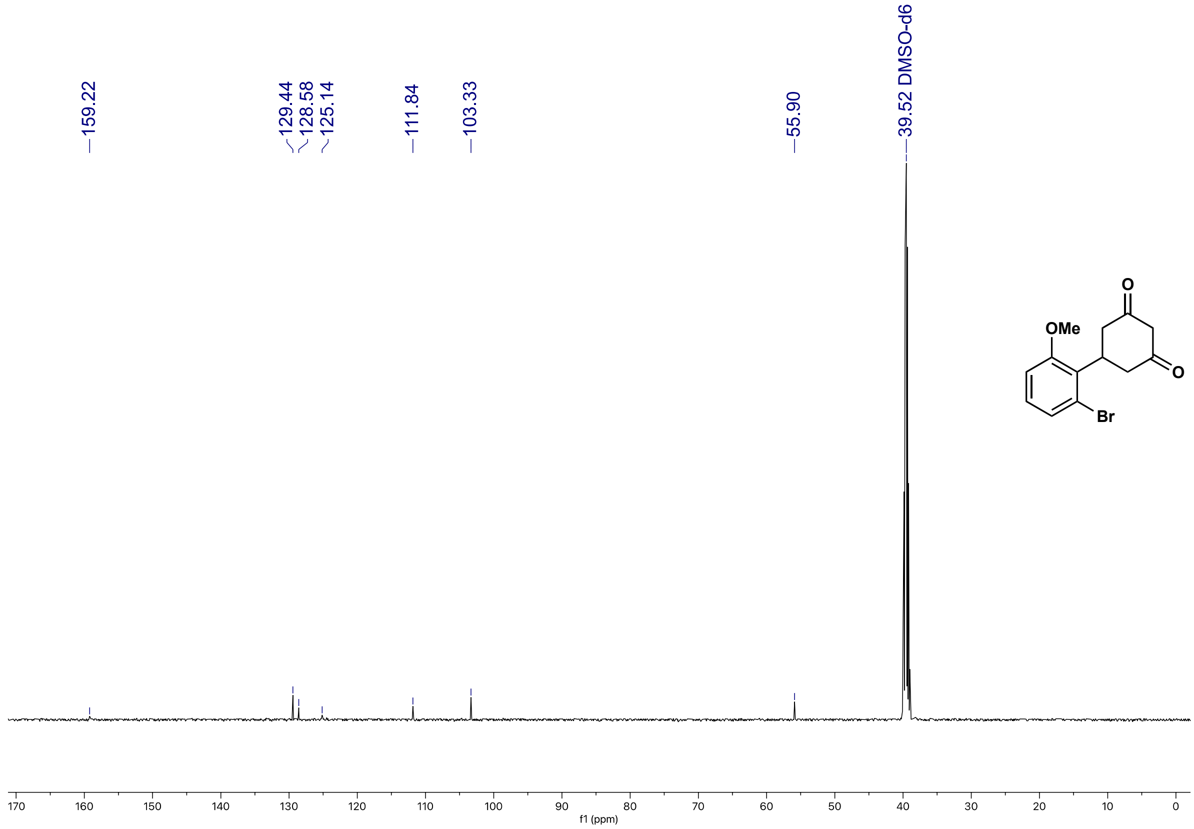

6-(2-bromo-6-methoxyphenyl)-4-oxo-4,5,6,7-tetrahydrobenzofuran-3-carboxylic acid (**16**) $^{1}H$ NMR (500 MHz, $CDCl_3$)

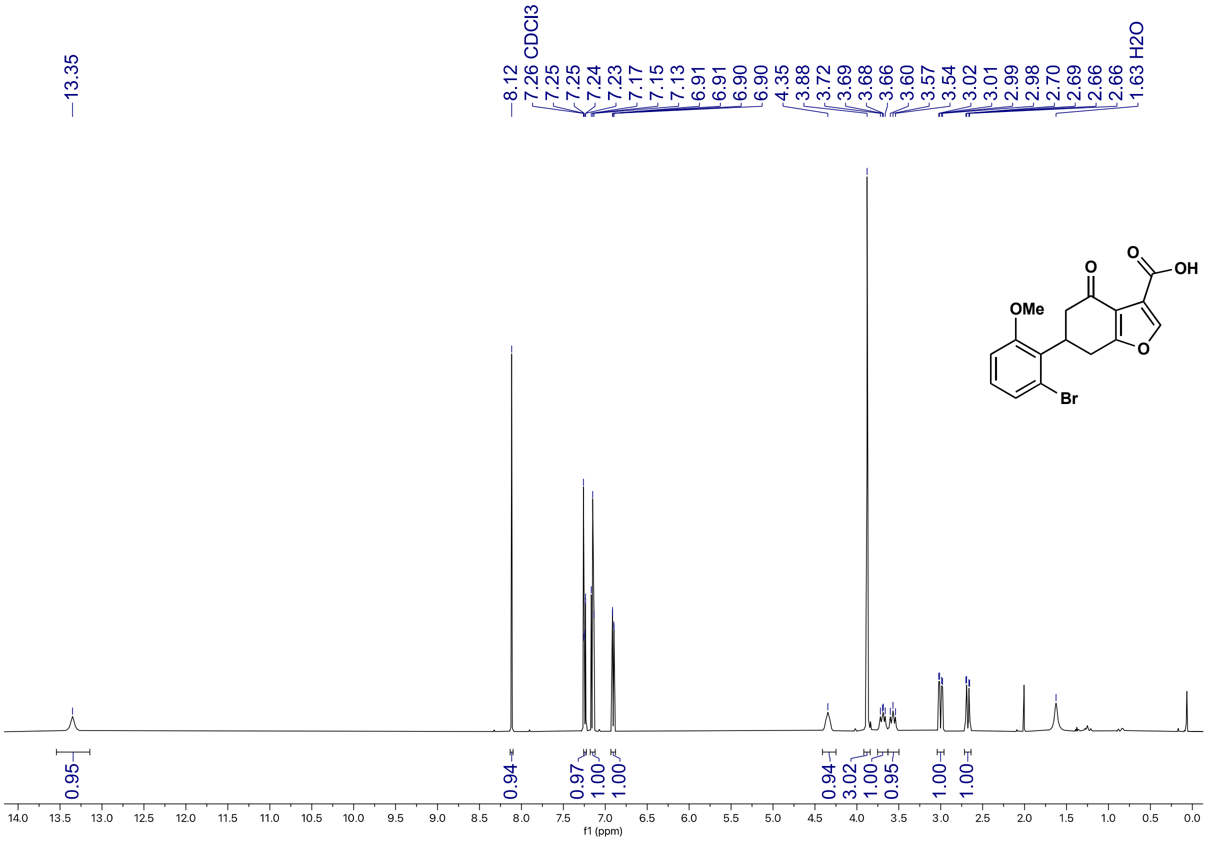

6-(2-bromo-6-methoxyphenyl)-4-oxo-4,5,6,7-tetrahydrobenzofuran-3-carboxylic acid (**16**) $^{13}C$ NMR (126 MHz, $CDCl_3$)

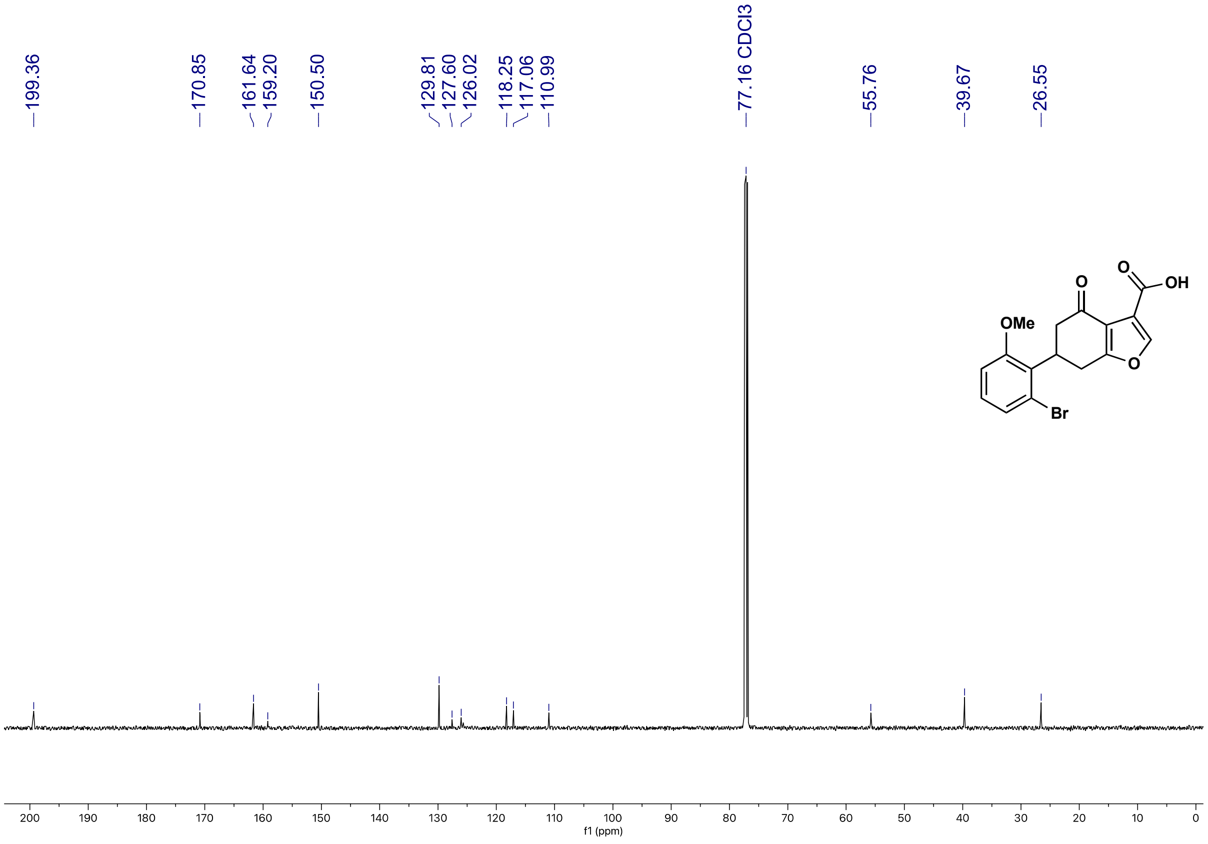

ethyl 6-(2-bromo-6-methoxyphenyl)-4-oxo-4,5,6,7-tetrahydrobenzofuran-3-carboxylate (**16d**)
$^{1}$H NMR (500 MHz, $CDCl_3$)

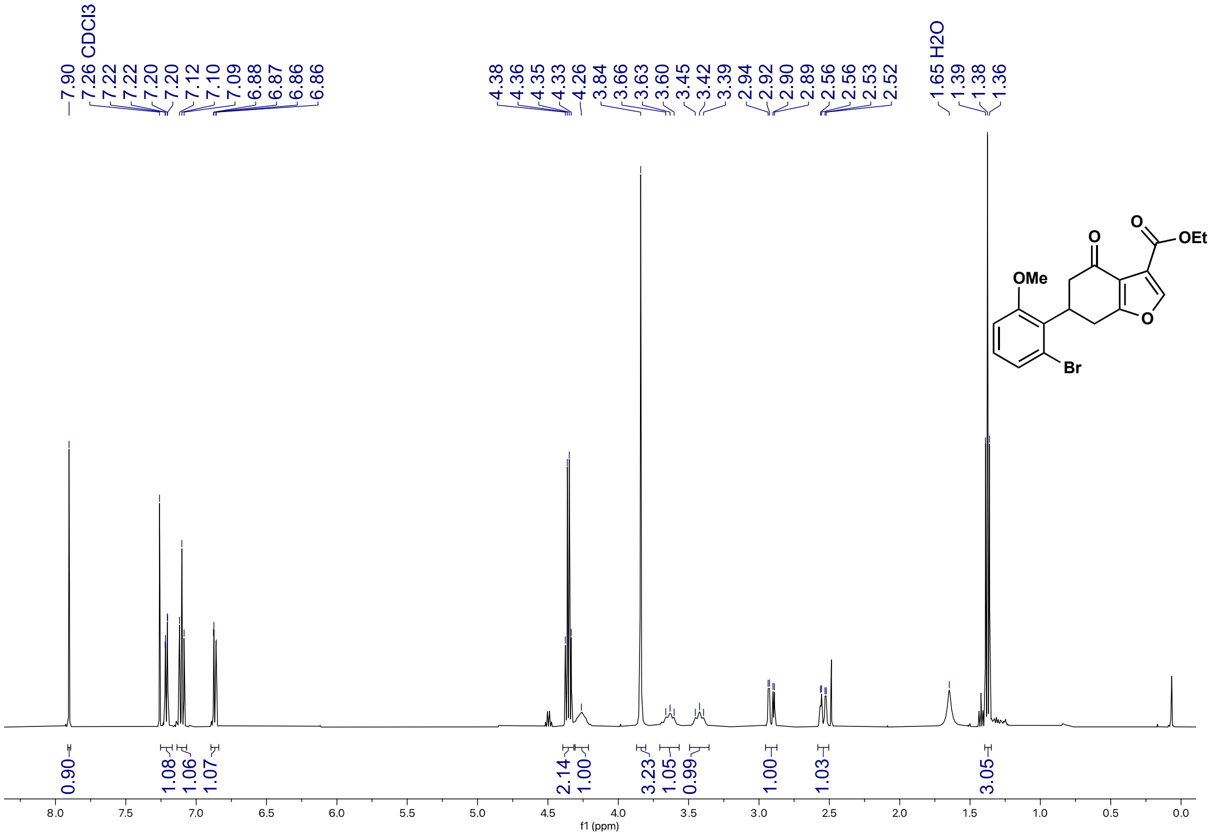

ethyl 6-(2-bromo-6-methoxyphenyl)-4-oxo-4,5,6,7-tetrahydrobenzofuran-3-carboxylate (**16d**)
$^{13}$C NMR (126 MHz, $CDCl_3$)

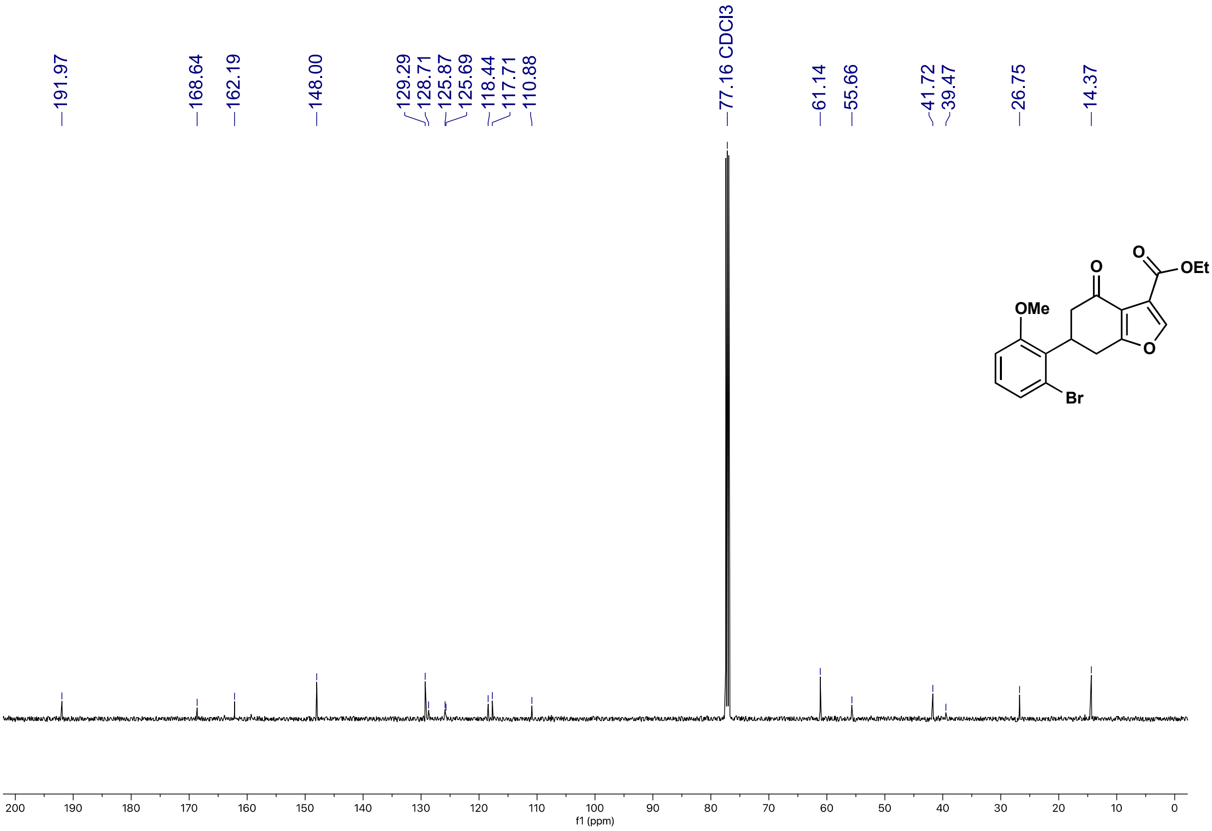

(*E*)-4-(2-chloro-6-fluorophenyl)but-3-en-2-one (**17b**) $^{1}$H NMR (500 MHz, $CDCl_3$)

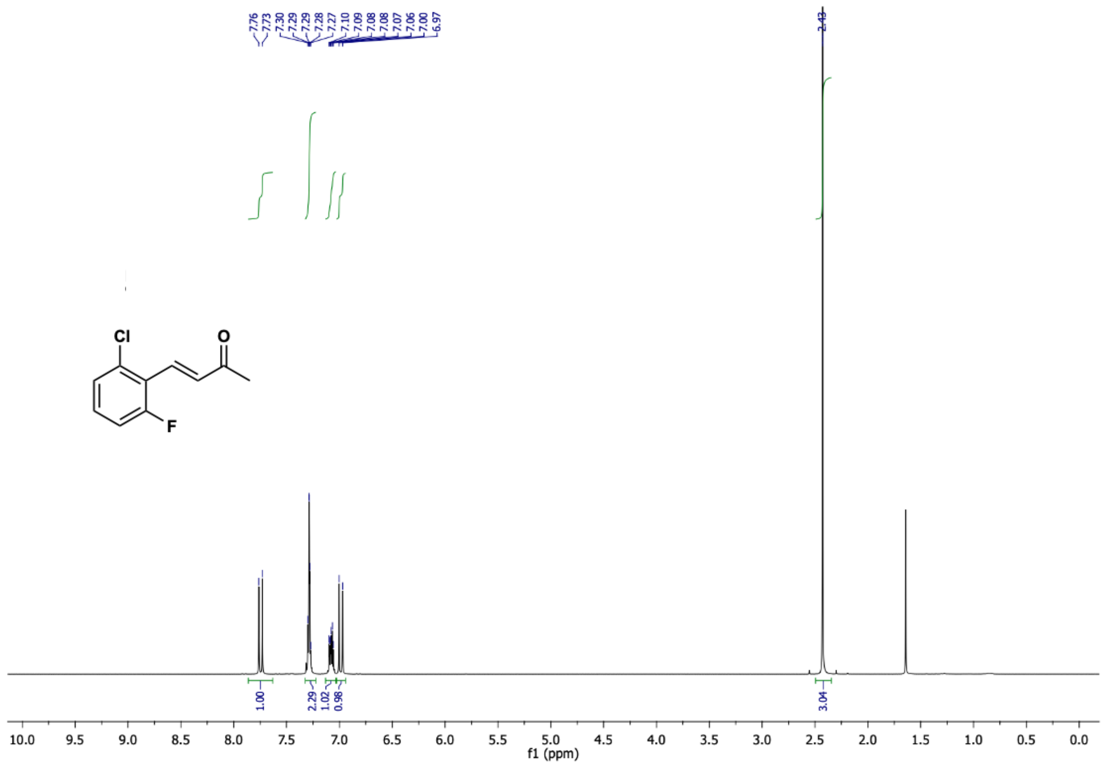

(*E*)-4-(2-chloro-6-fluorophenyl)but-3-en-2-one (**17b**) $^{13}$C NMR (126 MHz, $CDCl_3$)

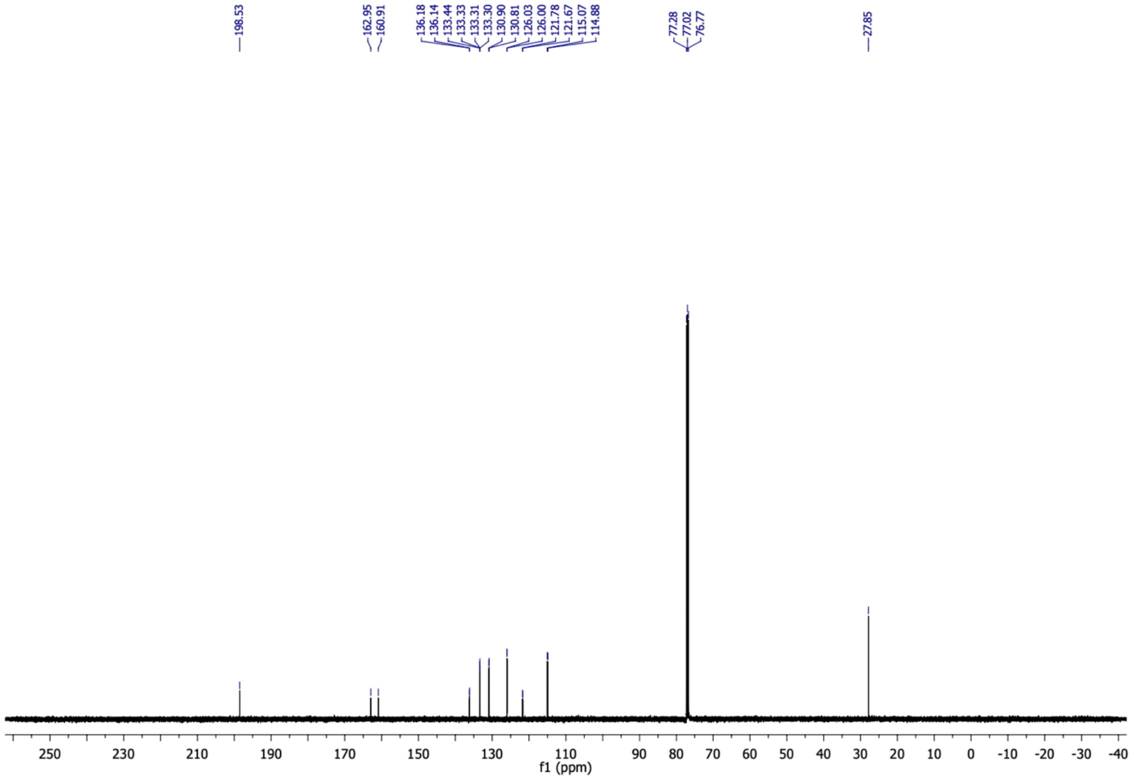

(*E*)-4-(2-chloro-6-fluorophenyl)but-3-en-2-one (**17b**) $^{19}F$ NMR (470 MHz, $CDCl_3$)

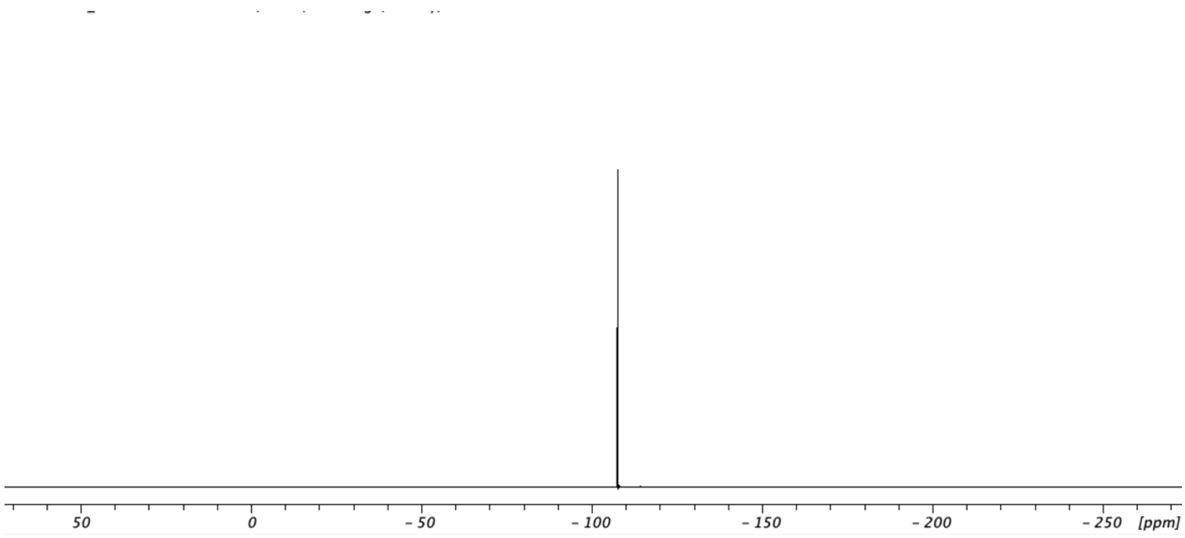

5-(2-chloro-6-fluorophenyl)cyclohexane-1,3-dione (**17c**) $^1H$ NMR (500 MHz, $CD_3OD$)

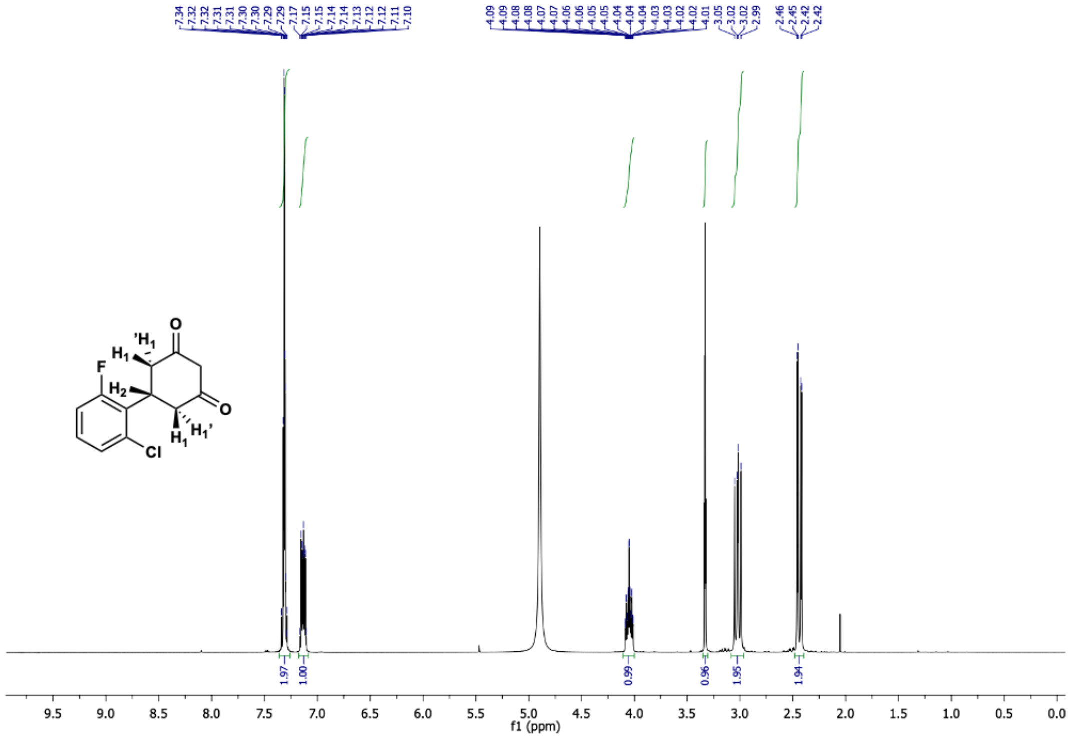

5-(2-chloro-6-fluorophenyl)cyclohexane-1,3-dione (**17c**) $^{13}C$ NMR (126 MHz, $CD_3OD$)

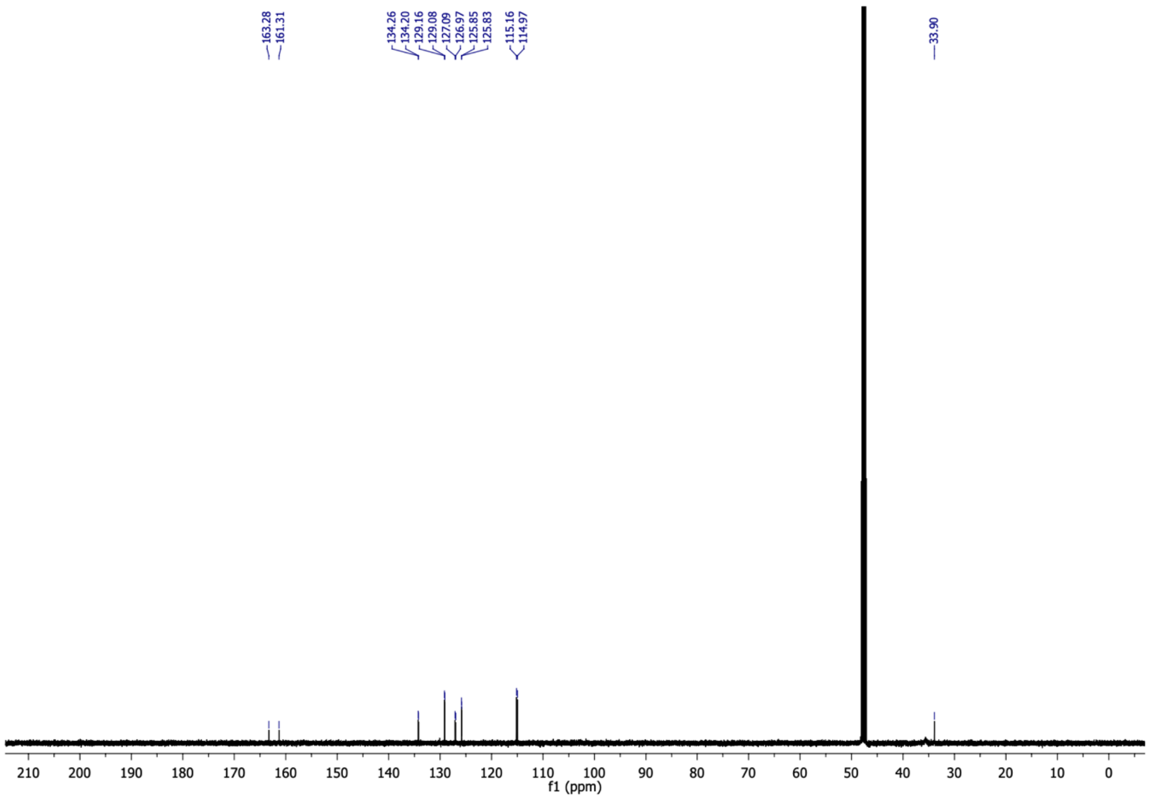

5-(2-chloro-6-fluorophenyl)cyclohexane-1,3-dione (**17c**) $^{19}F$ NMR (470 MHz, $CD_3OD$)

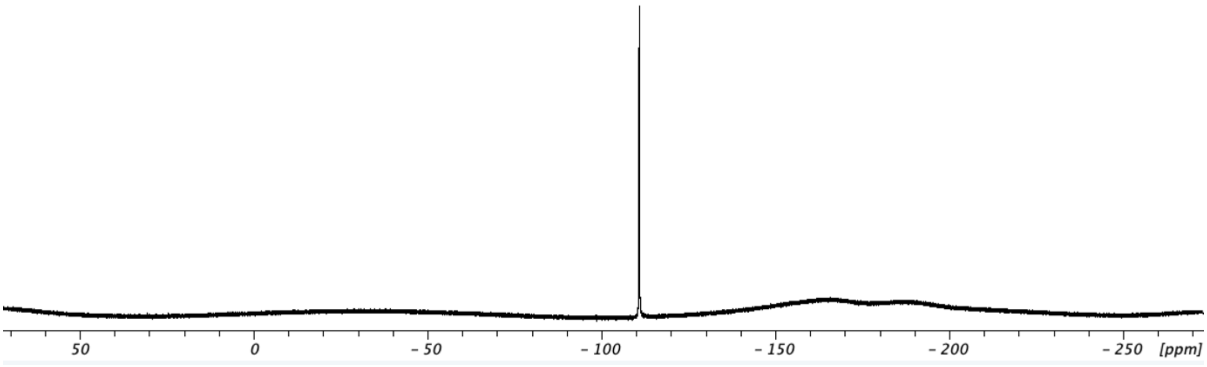

6-(2-chloro-6-fluorophenyl)-4-oxo-4,5,6,7-tetrahydrobenzofuran-3-carboxylic acid (**17**) $^{1}$H NMR (400 MHz, $CDCl_3$)

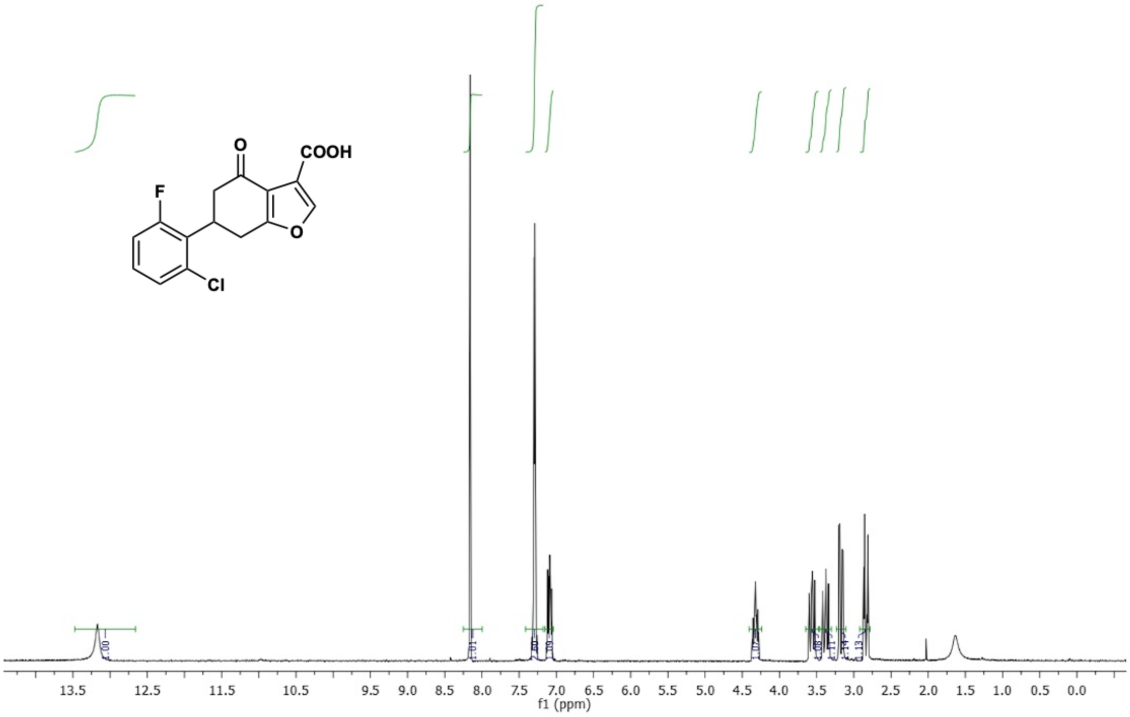

6-(2-chloro-6-fluorophenyl)-4-oxo-4,5,6,7-tetrahydrobenzofuran-3-carboxylic acid (**17**) $^{13}$C NMR (101 MHz, $CDCl_3$)

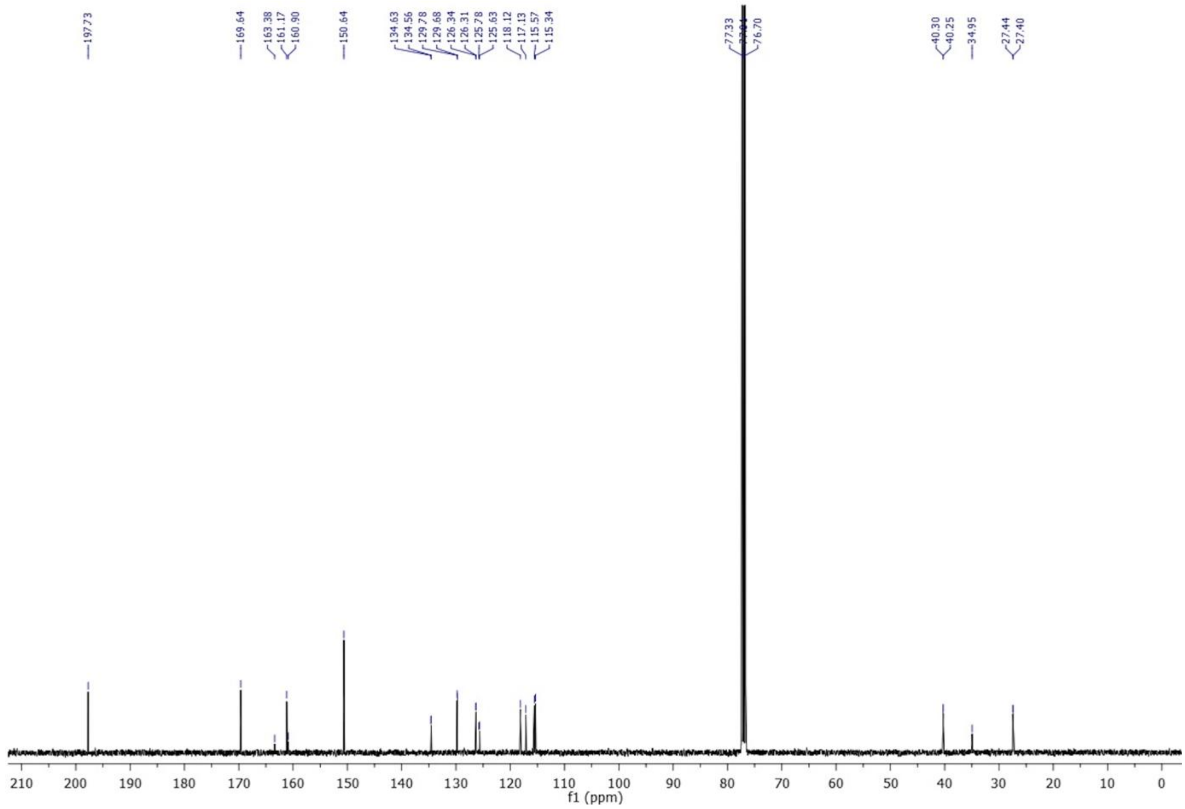

6-(2-chloro-6-fluorophenyl)-4-oxo-4,5,6,7-tetrahydrobenzofuran-3-carboxylic acid (**17**) $^{19}$F NMR (376 MHz, $CDCl_3$)

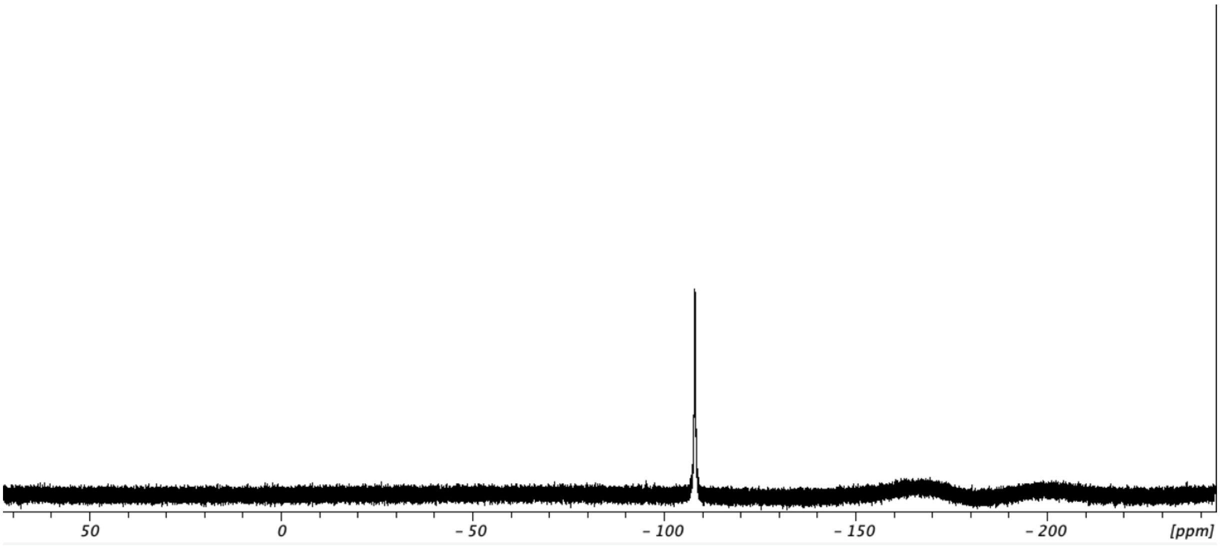

6-(2-chloro-6-fluorophenyl)-4-hydroxy-4,5,6,7-tetrahydrobenzofuran-3-carboxylic acid (**17e**) $^{1}$H NMR (500 MHz, $CDCl_3$)

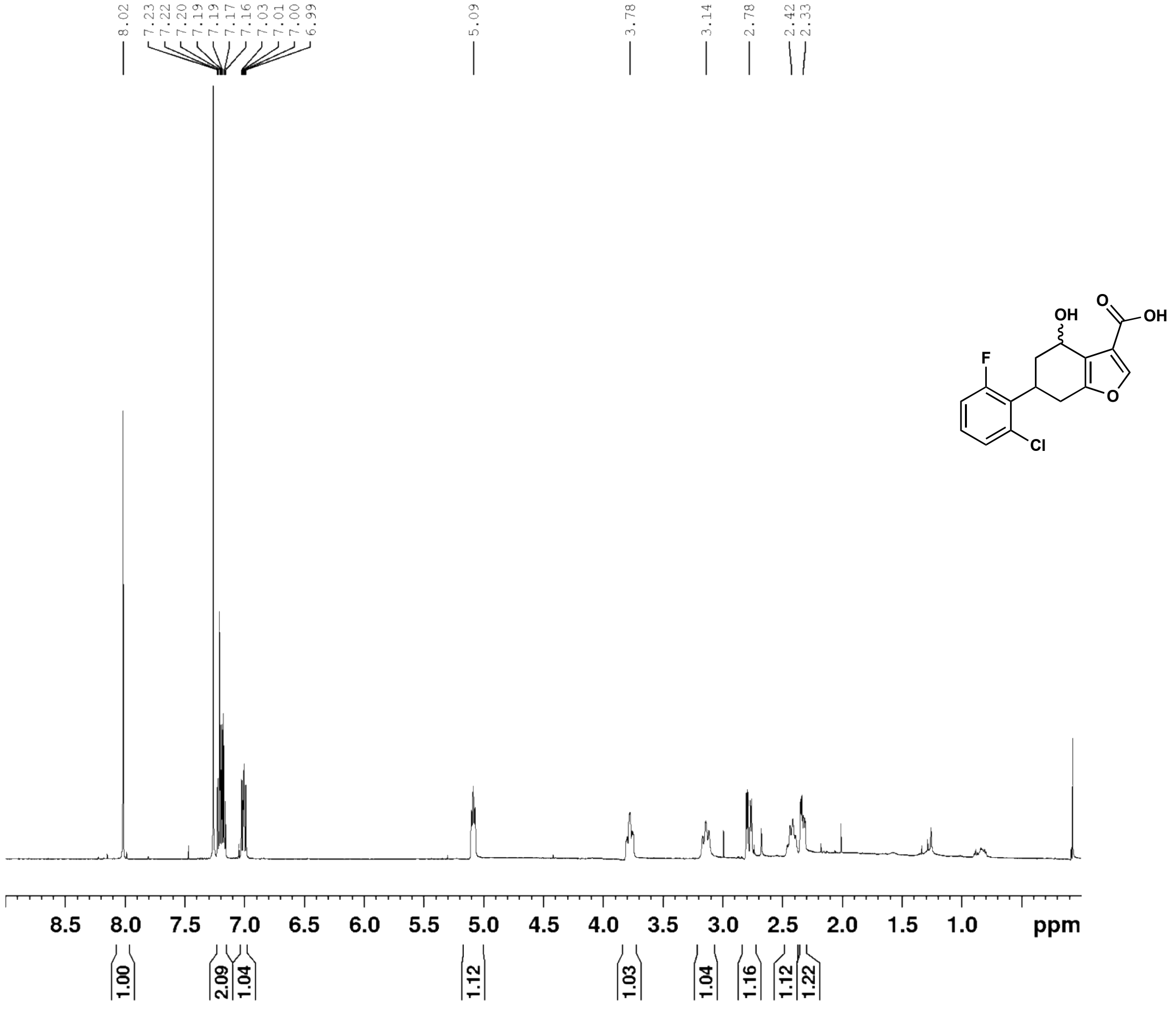

6-(2-chloro-6-fluorophenyl)-4-hydroxy-4,5,6,7-tetrahydrobenzofuran-3-carboxylic acid (**17e**)
$^{13}$C NMR (126 MHz, $CDCl_3$)

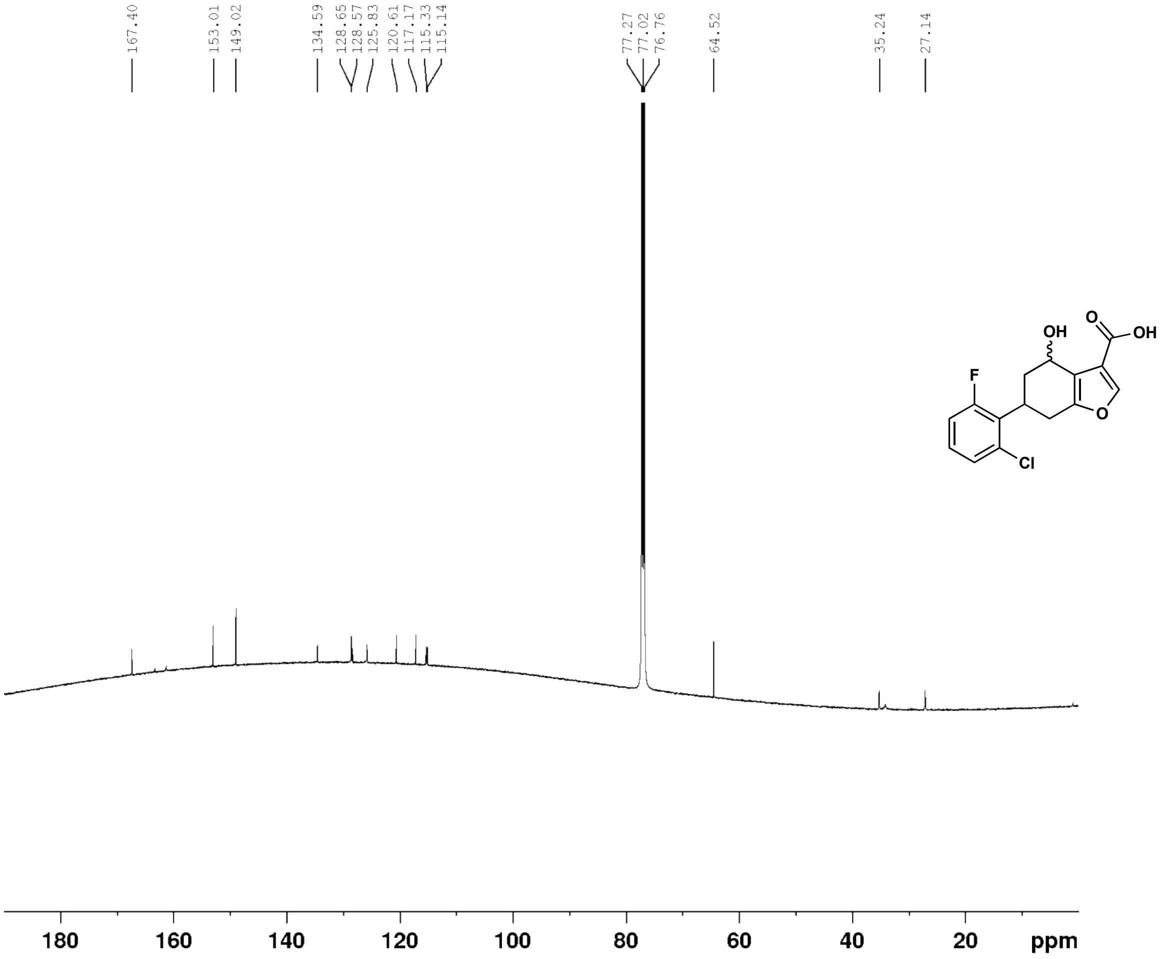

(*E*)-4-(2,6-dimethoxyphenyl)but-3-en-2-one (**18b**) $^{1}H$ NMR (500 MHz, $CDCl_3$)

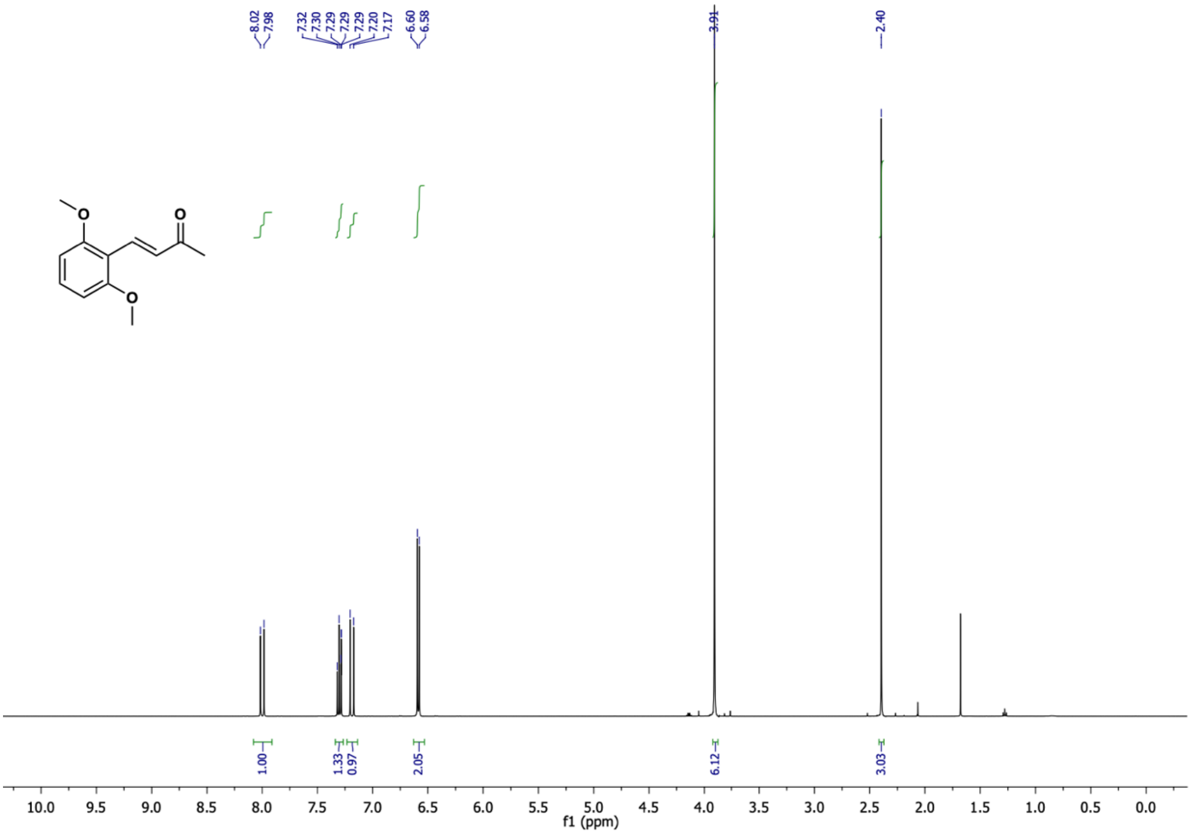

(*E*)-4-(2,6-dimethoxyphenyl)but-3-en-2-one (**18b**) $^{13}C$ NMR (126 MHz, $CDCl_3$)

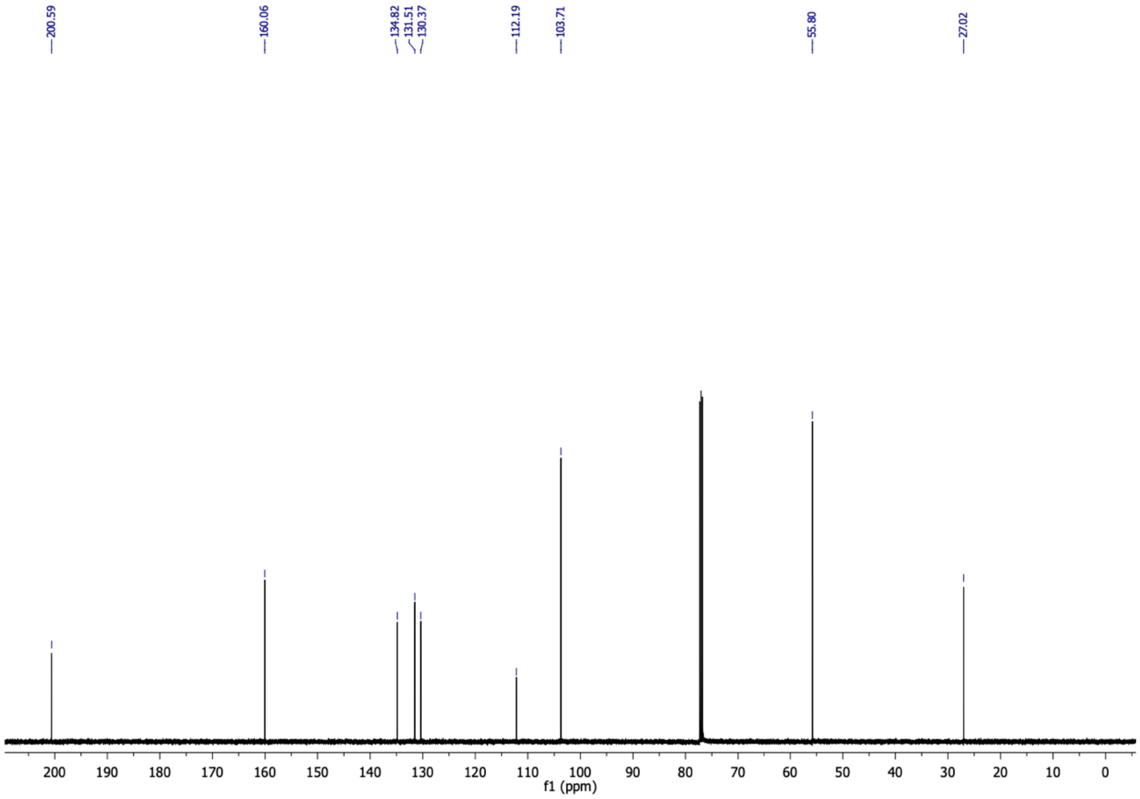

5-(2,6-dimethoxyphenyl)cyclohexane-1,3-dione (**18c**) $^{1}H$ NMR (500 MHz, $CD_3OD$)

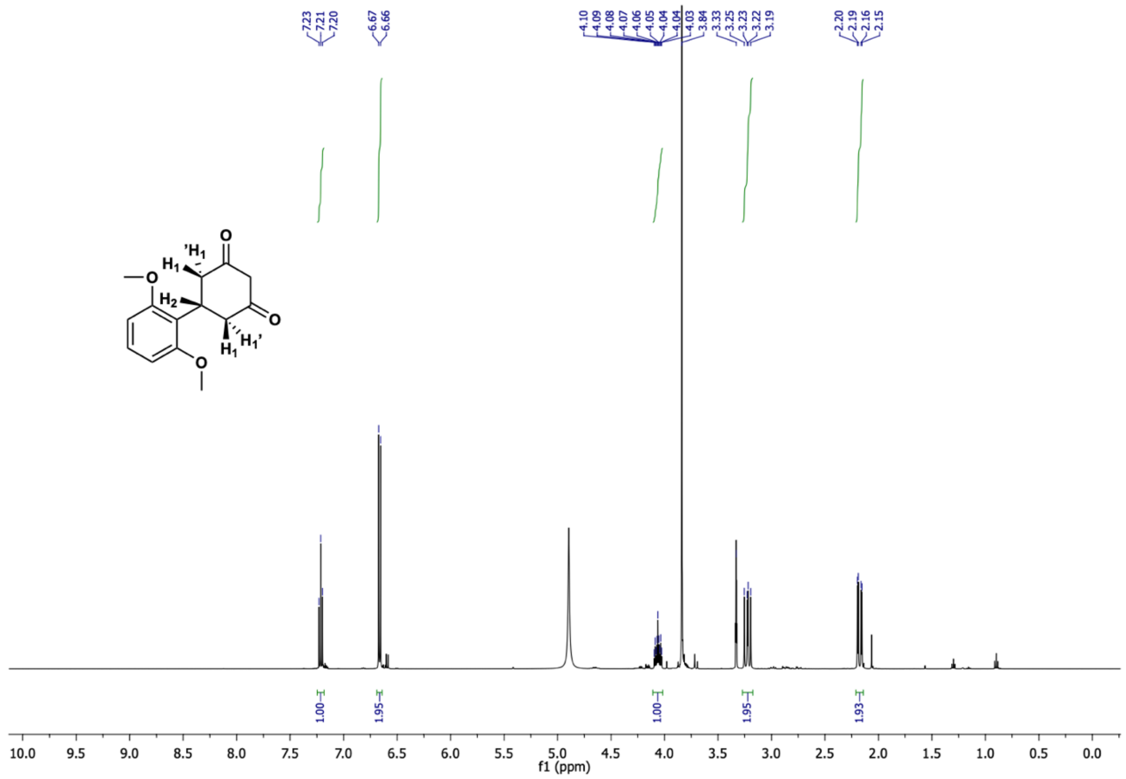

5-(2,6-dimethoxyphenyl)cyclohexane-1,3-dione (**18c**) $^{13}C$ NMR (126 MHz, $CD_3OD$)

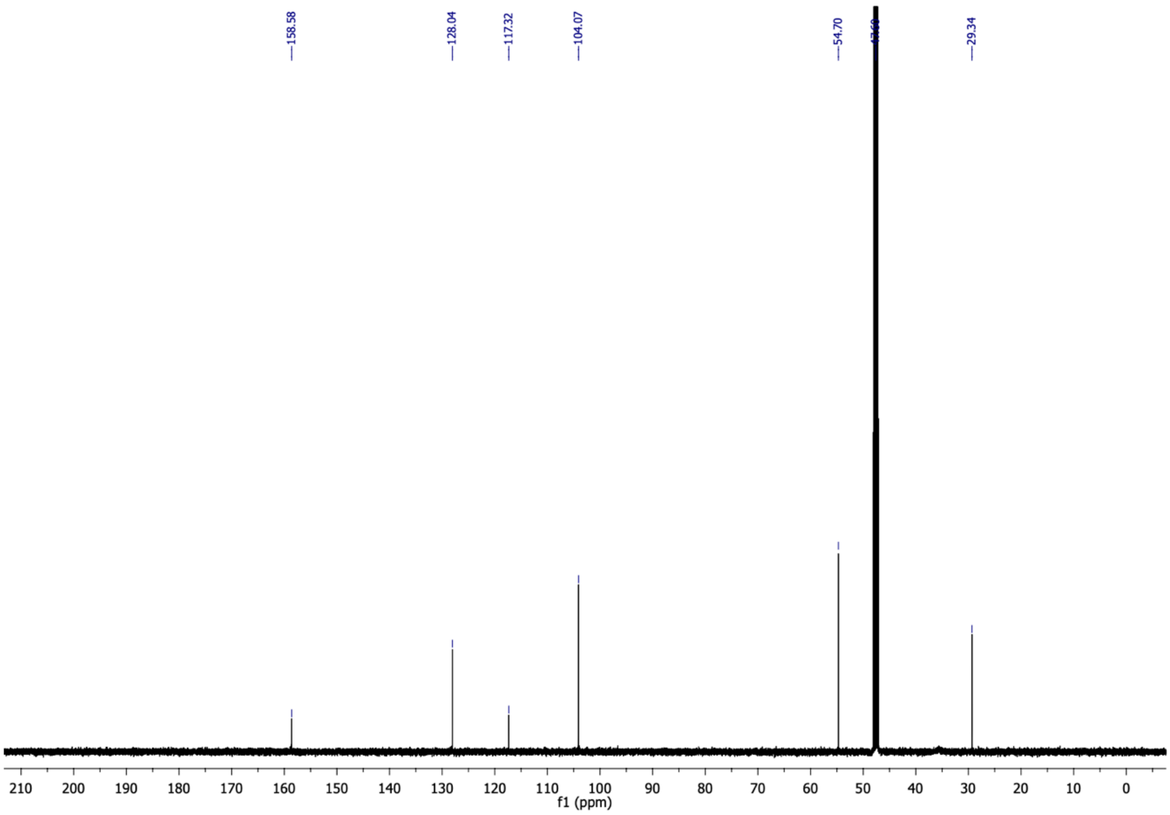

6-(2,6-dimethoxyphenyl)-4-oxo-4,5,6,7-tetrahydrobenzofuran-3-carboxylic acid (**18**) $^{1}$H NMR (500 MHz, $CDCl_3$)

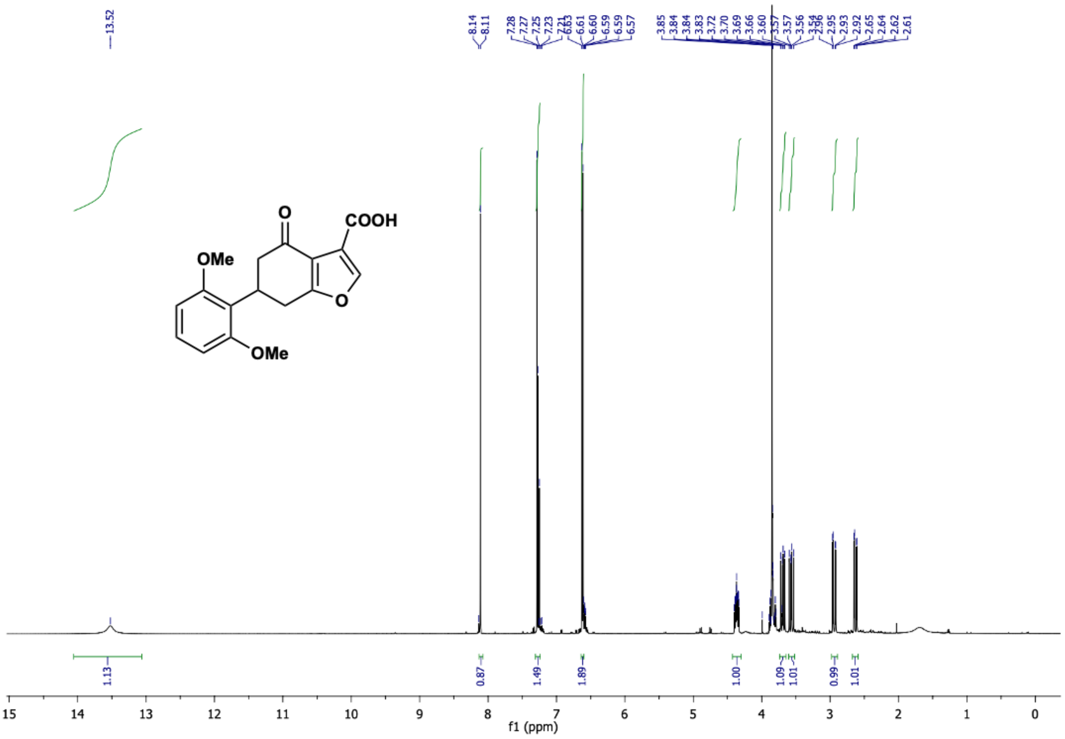

6-(2,6-dimethoxyphenyl)-4-oxo-4,5,6,7-tetrahydrobenzofuran-3-carboxylic acid (**18**) $^{13}$C NMR (126 MHz, $CDCl_3$)

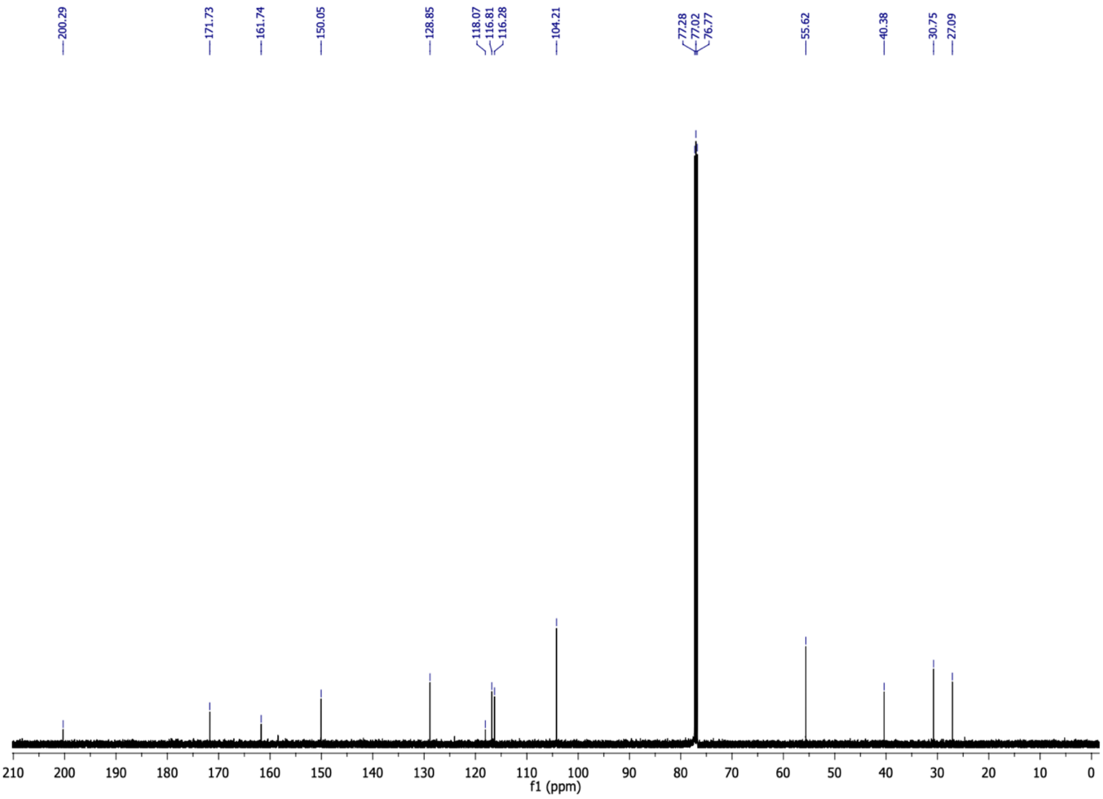

(E)-4-(2-bromo-6-chlorophenyl)but-3-en-2-one (**19b**) $^1$H NMR (500 MHz, $CDCl_3$)

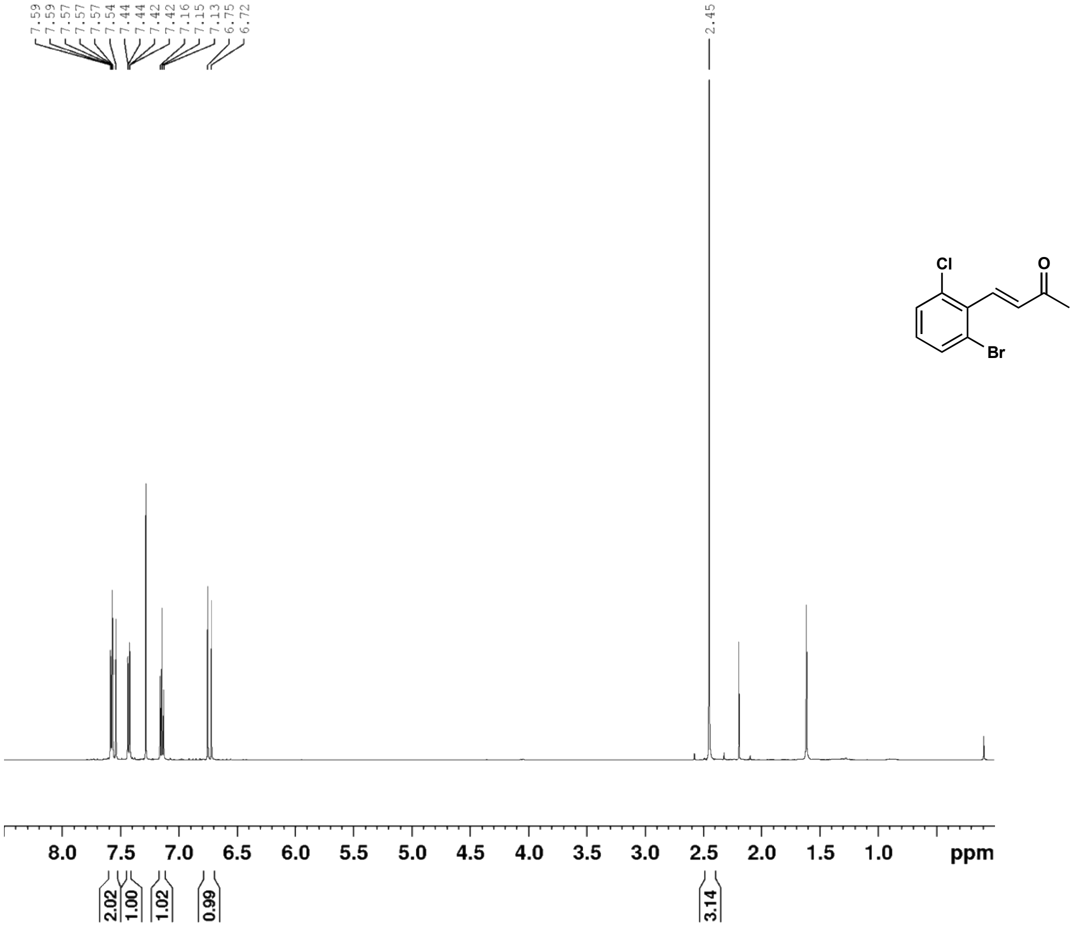

(E)-4-(2-bromo-6-chlorophenyl)but-3-en-2-one (**19b**) $^{13}$C NMR (126 MHz, $CDCl_3$)

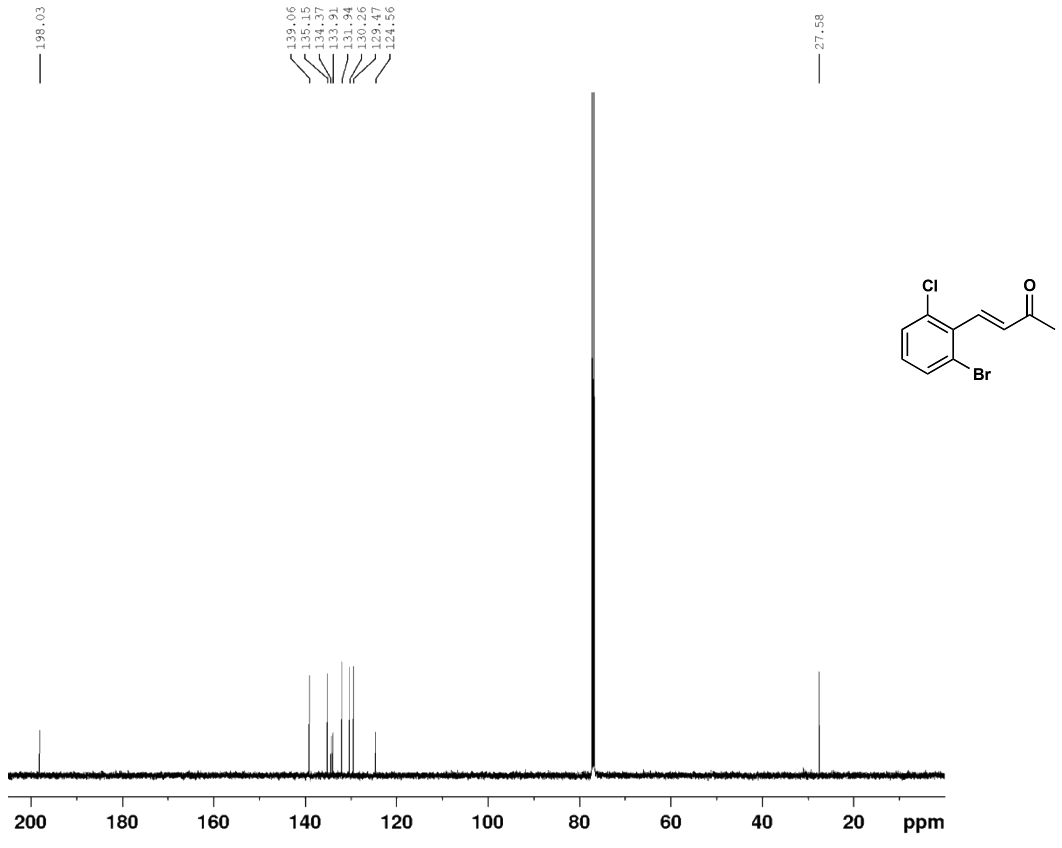

5-(2-bromo-6-chlorophenyl)cyclohexane-1,3-dione (**19c**) $^{1}$H NMR (500 MHz, $CD_3OD$)

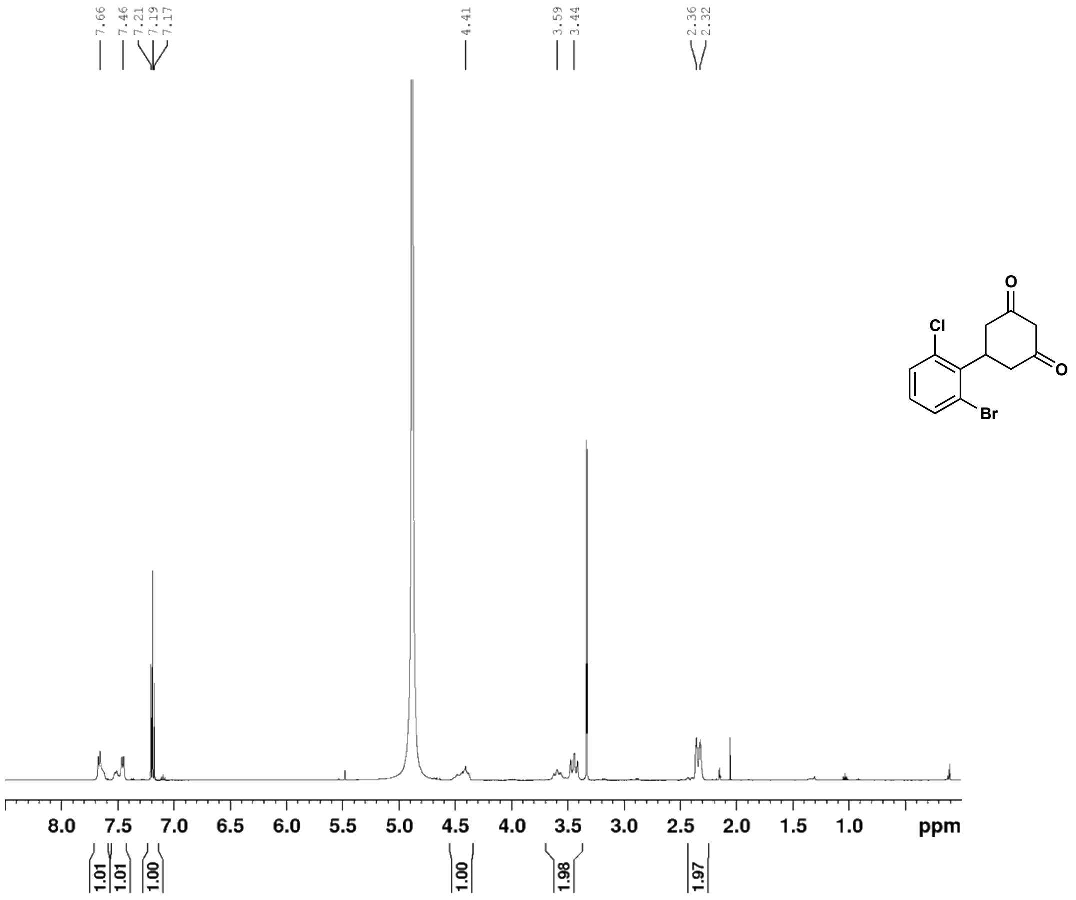

5-(2-bromo-6-chlorophenyl)cyclohexane-1,3-dione (**19c**) $^{13}$C NMR (126 MHz, $CD_3OD$)

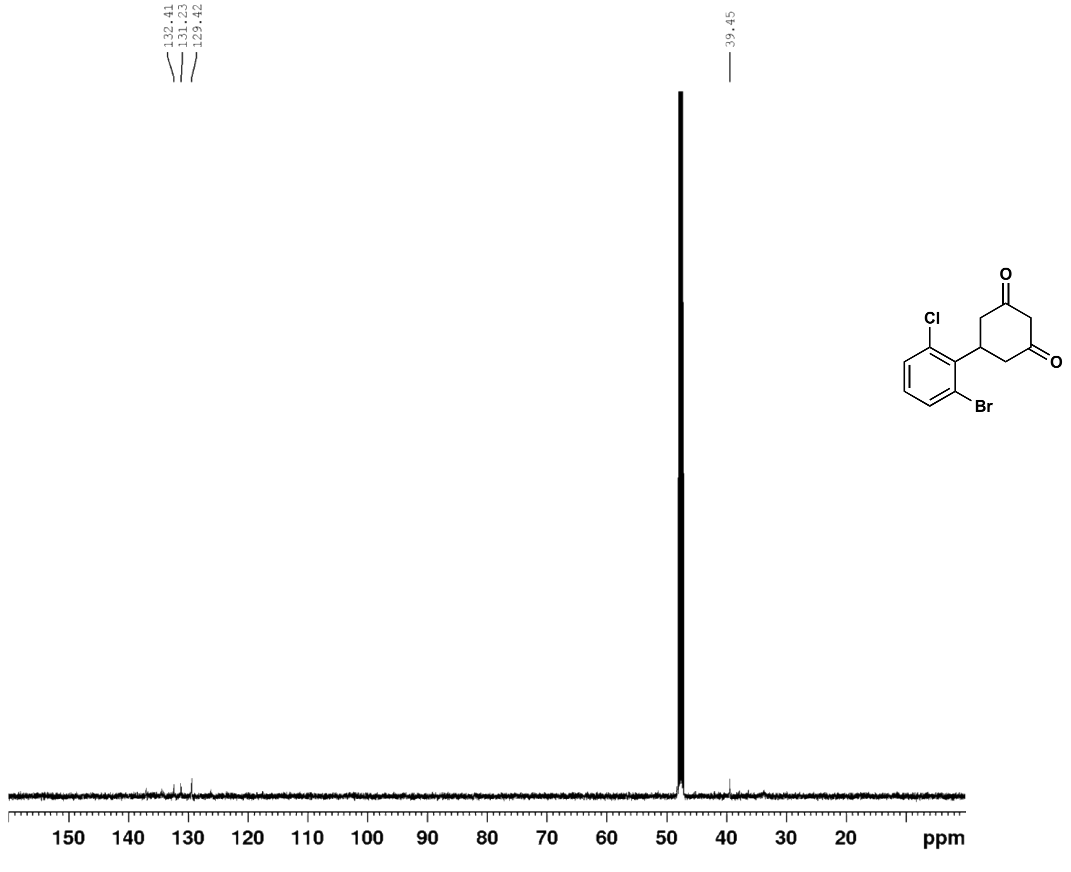

6-(2-bromo-6-chlorophenyl)-4-oxo-4,5,6,7-tetrahydrobenzofuran-3-carboxylic acid (**19**) $^{1}$H NMR (500 MHz, $CDCl_3$)

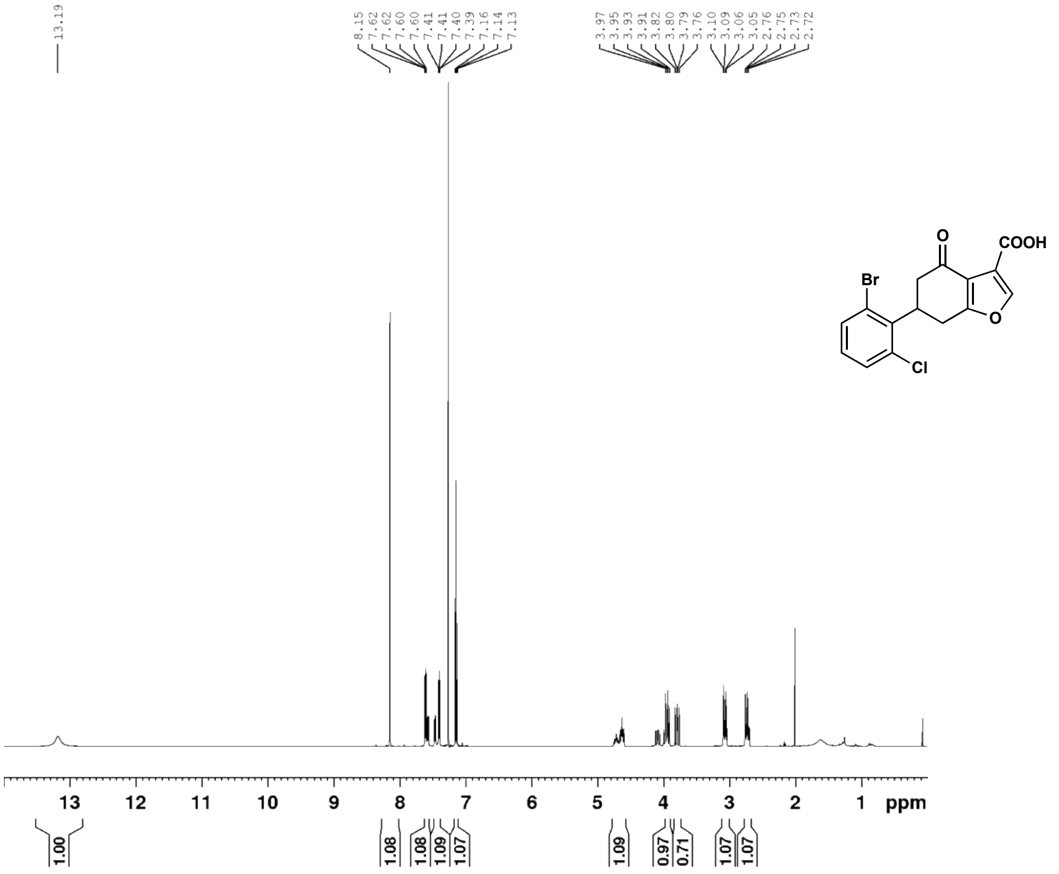

6-(2-bromo-6-chlorophenyl)-4-oxo-4,5,6,7-tetrahydrobenzofuran-3-carboxylic acid (**19**) $^{13}$C NMR (126 MHz, $CDCl_3$)

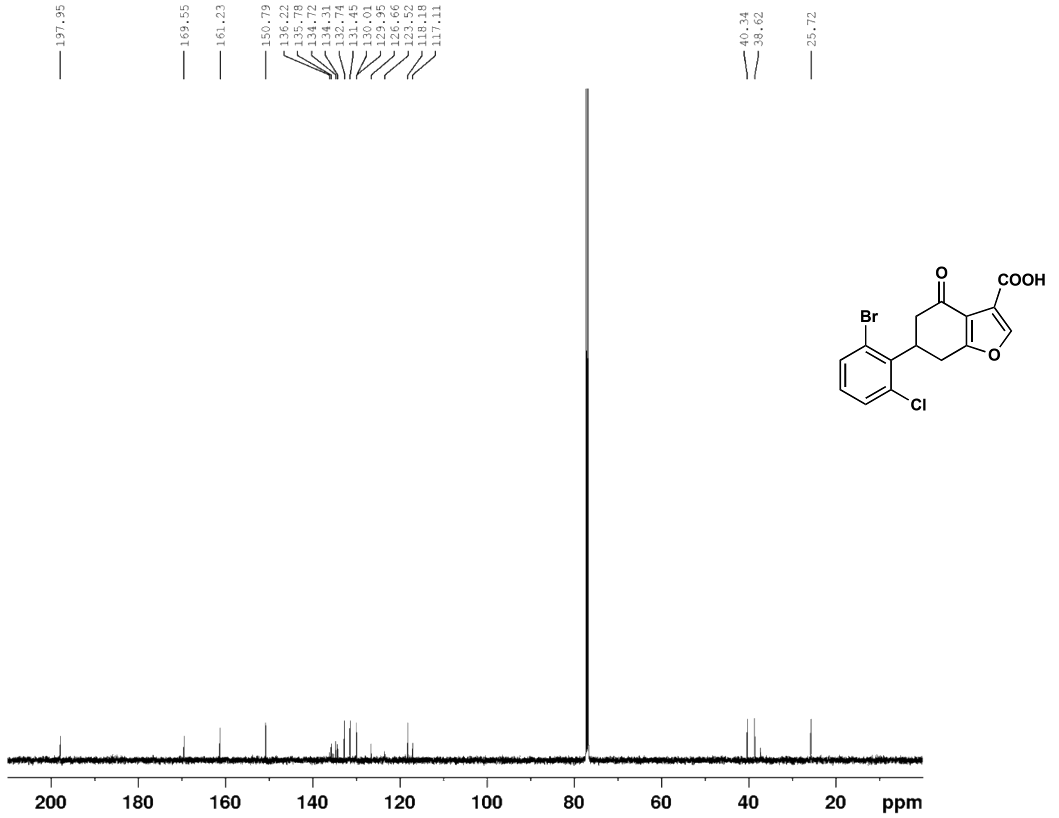

2-bromo-6-butoxybenzaldehyde (**20a**) $^{1}H$ NMR (500 MHz, $CDCl_3$)

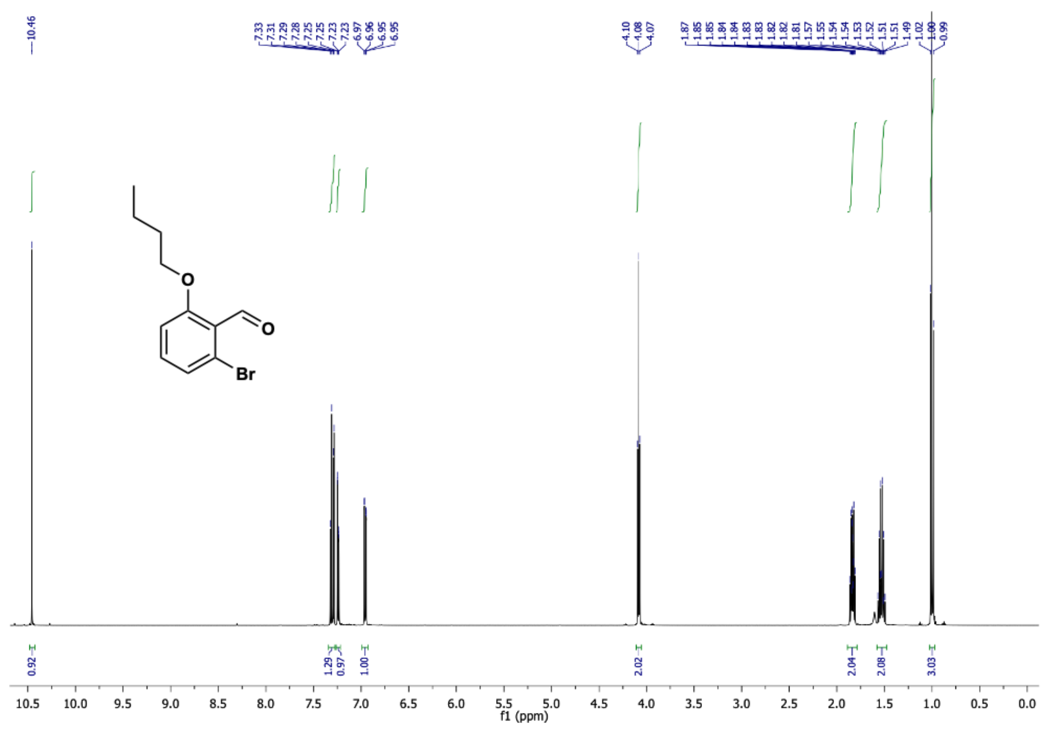

2-bromo-6-butoxybenzaldehyde (**20a**) $^{13}C$ NMR (126 MHz, $CDCl_3$)

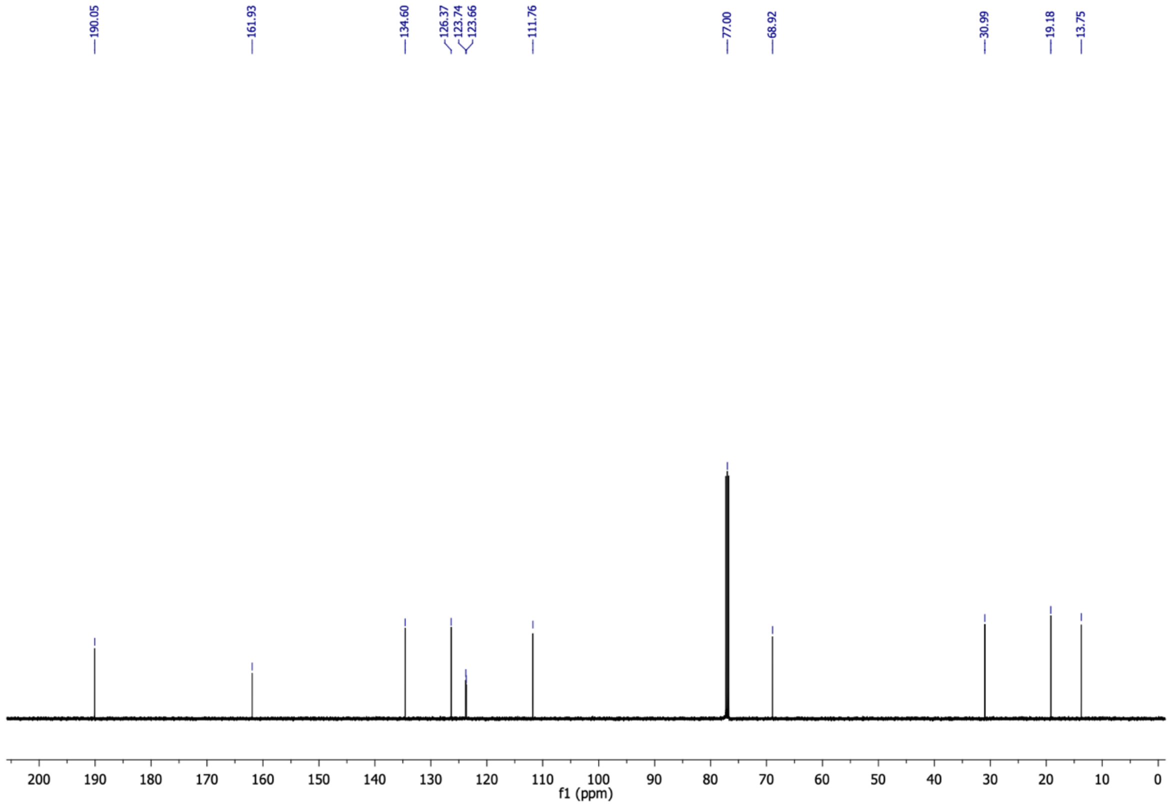

(*E*)-4-(2-bromo-6-butoxyphenyl)but-3-en-2-one (**20b**) $^{1}H$ NMR (500 MHz, $CDCl_3$)

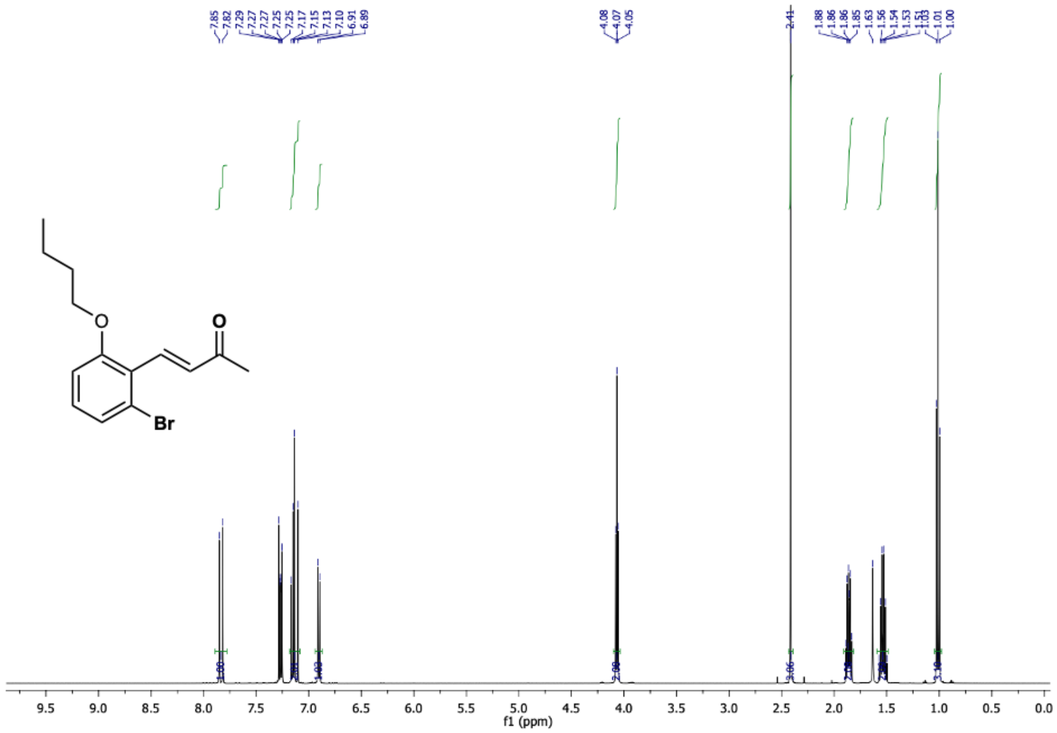

(*E*)-4-(2-bromo-6-butoxyphenyl)but-3-en-2-one (**20b**) $^{13}C$ NMR (126 MHz, $CDCl_3$)

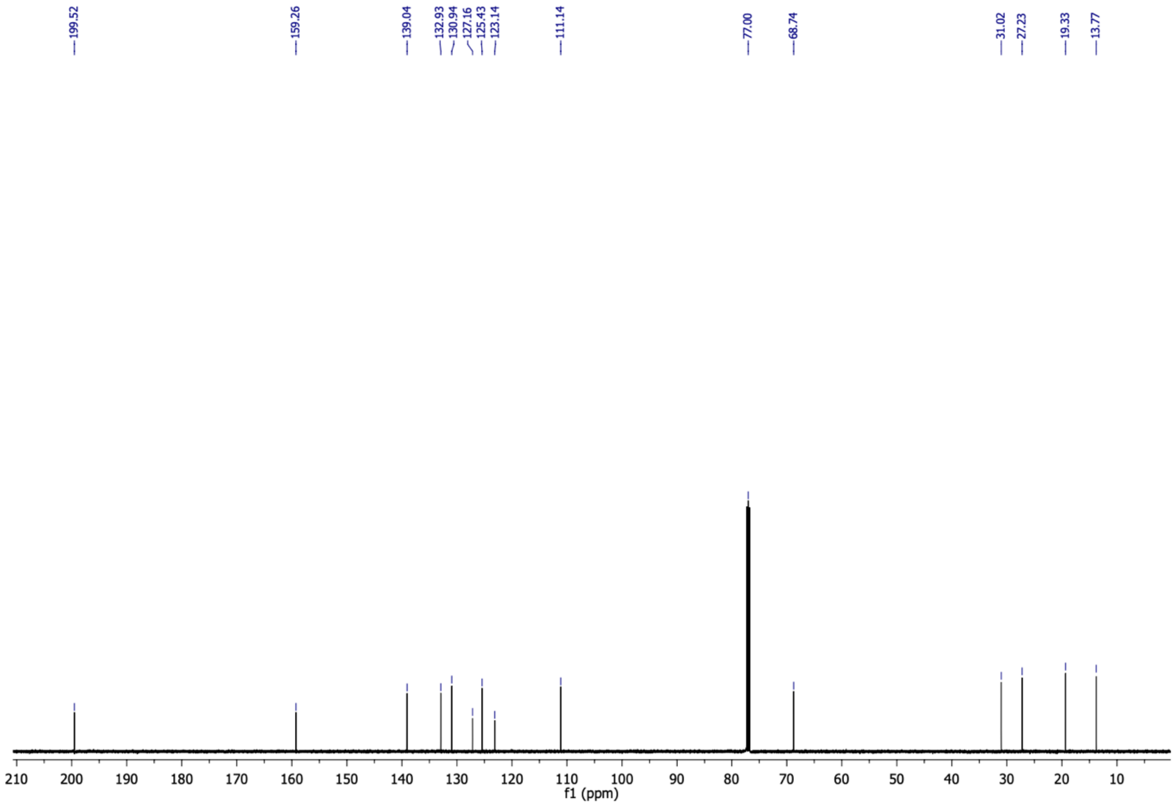

5-(2-bromo-6-butoxyphenyl)cyclohexane-1,3-dione (**20c**) $^{1}H$ NMR (500 MHz, $CD_3OD$)

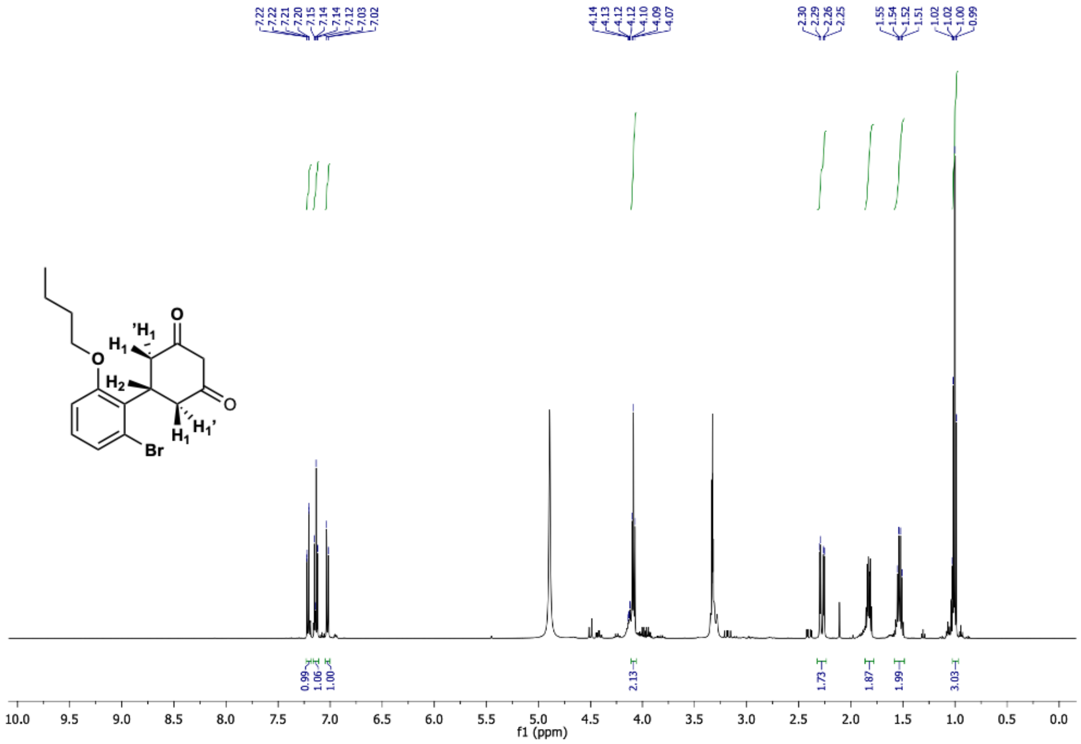

5-(2-bromo-6-butoxyphenyl)cyclohexane-1,3-dione (**20c**) $^{13}C$ NMR (126 MHz, $CD_3OD$)

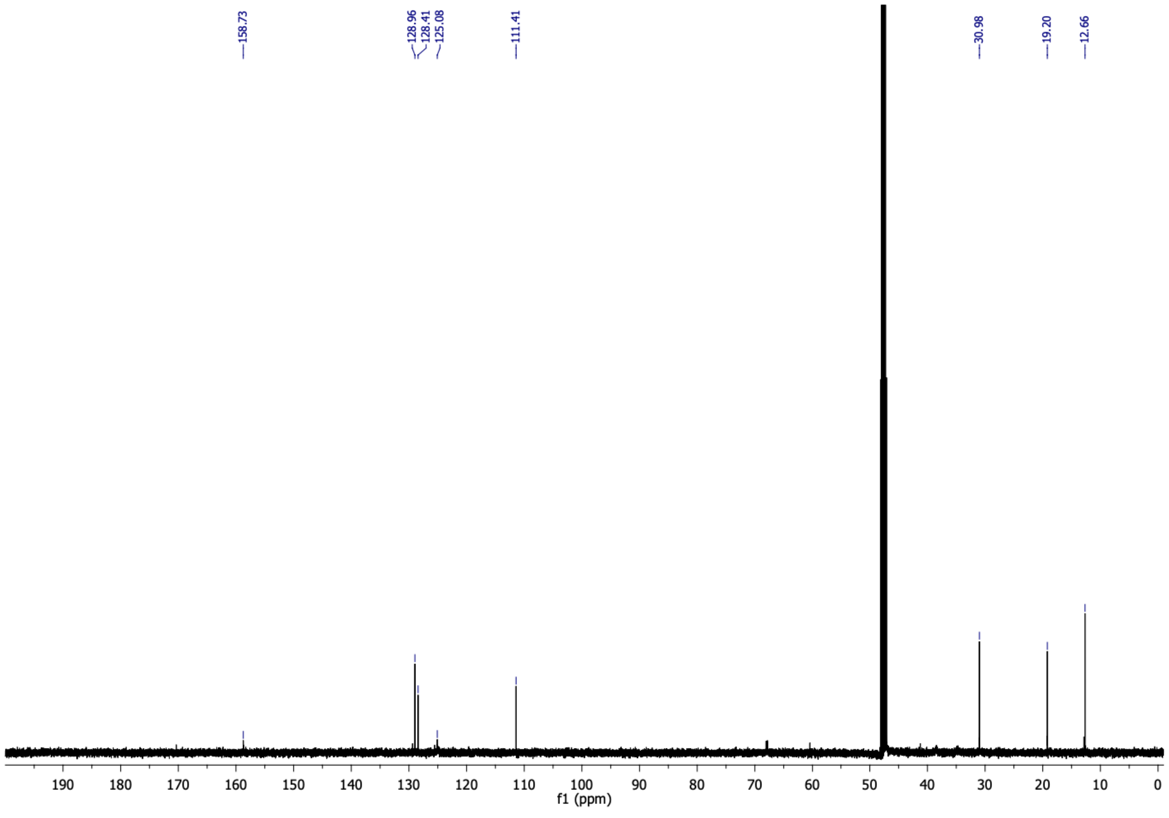

6-(2-bromo-6-butoxyphenyl)-4-oxo-4,5,6,7-tetrahydrobenzofuran-3-carboxylic acid (**20**) $^{1}$H NMR (500 MHz, $CDCl_3$)

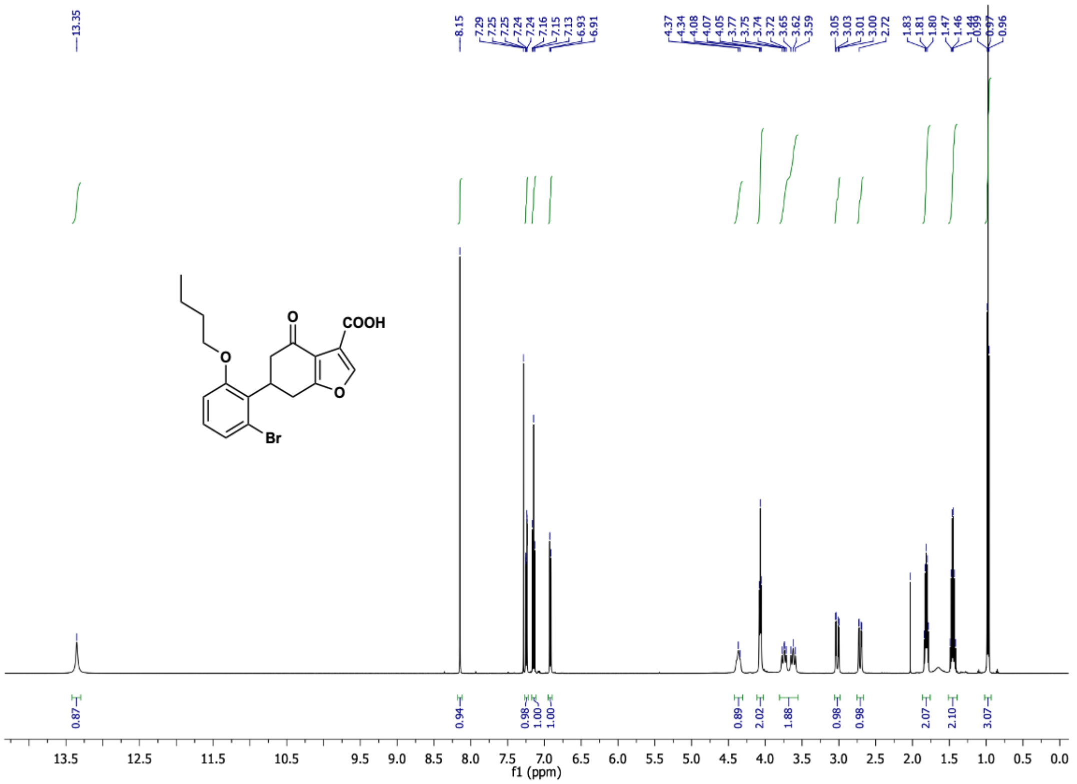

6-(2-bromo-6-butoxyphenyl)-4-oxo-4,5,6,7-tetrahydrobenzofuran-3-carboxylic acid (**20**) $^{13}$C NMR (126 MHz, $CDCl_3$)

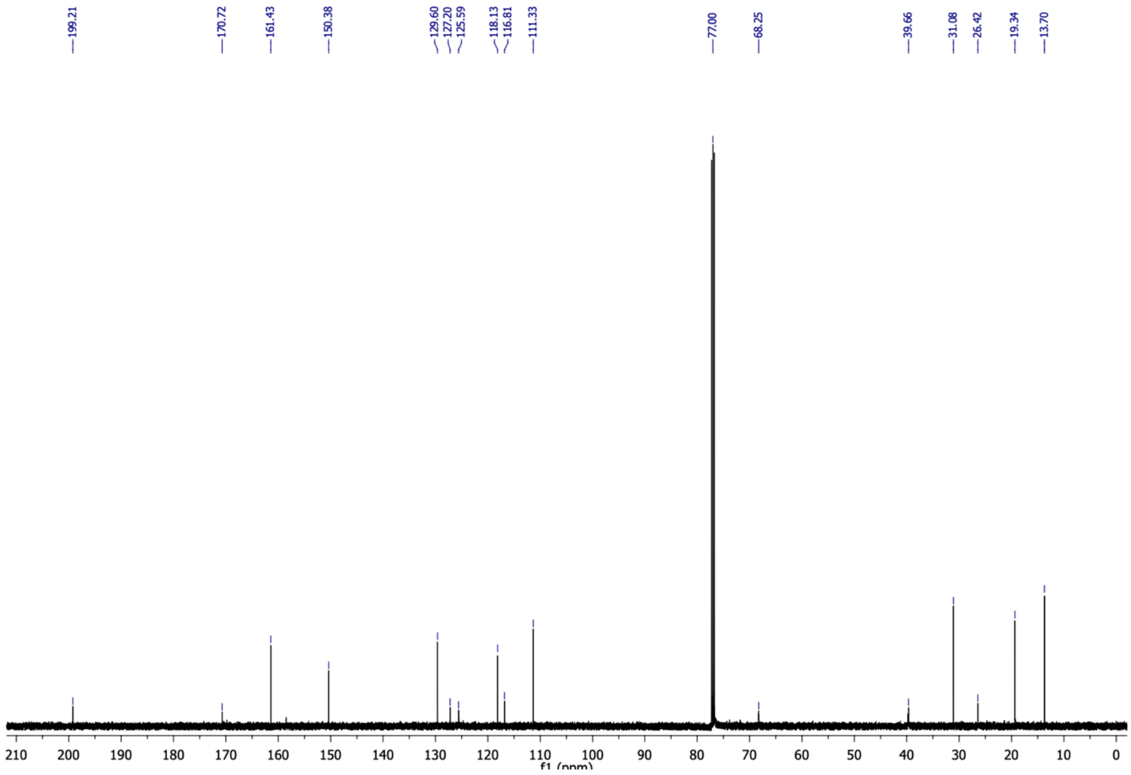

2-methoxy-6-(2-methoxyethoxy)benzaldehyde (**21a**) $^{1}$H NMR (500 MHz, $CDCl_3$)

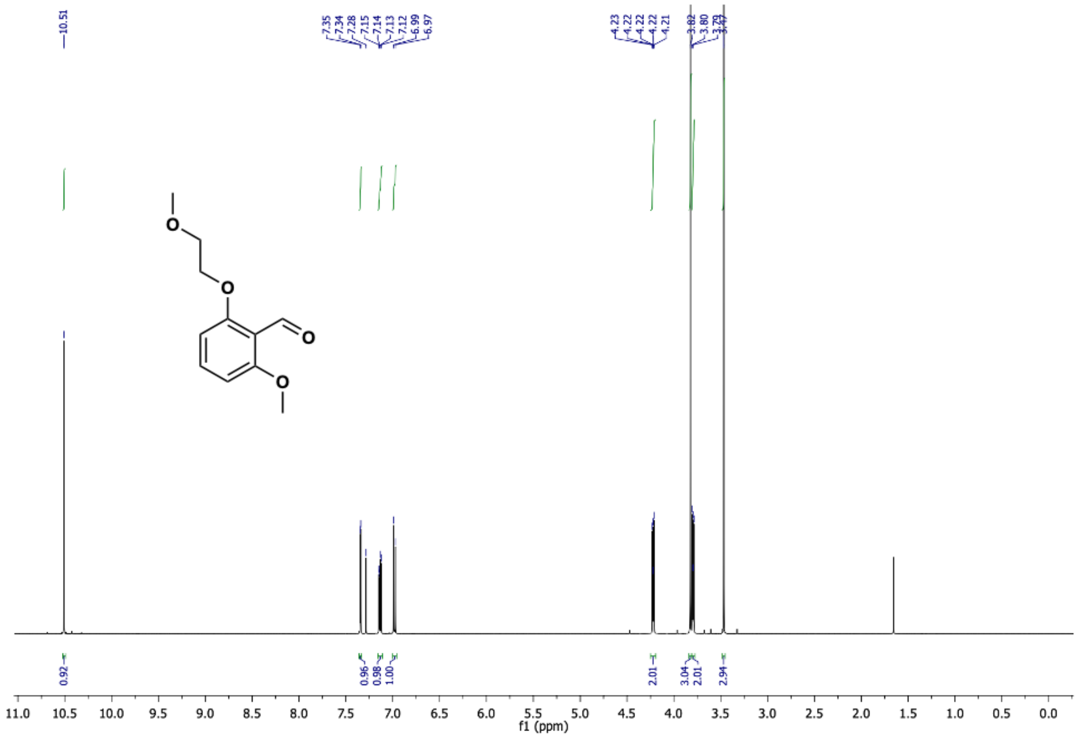

2-methoxy-6-(2-methoxyethoxy)benzaldehyde (**21a**) $^{13}$C NMR (126 MHz, $CDCl_3$)

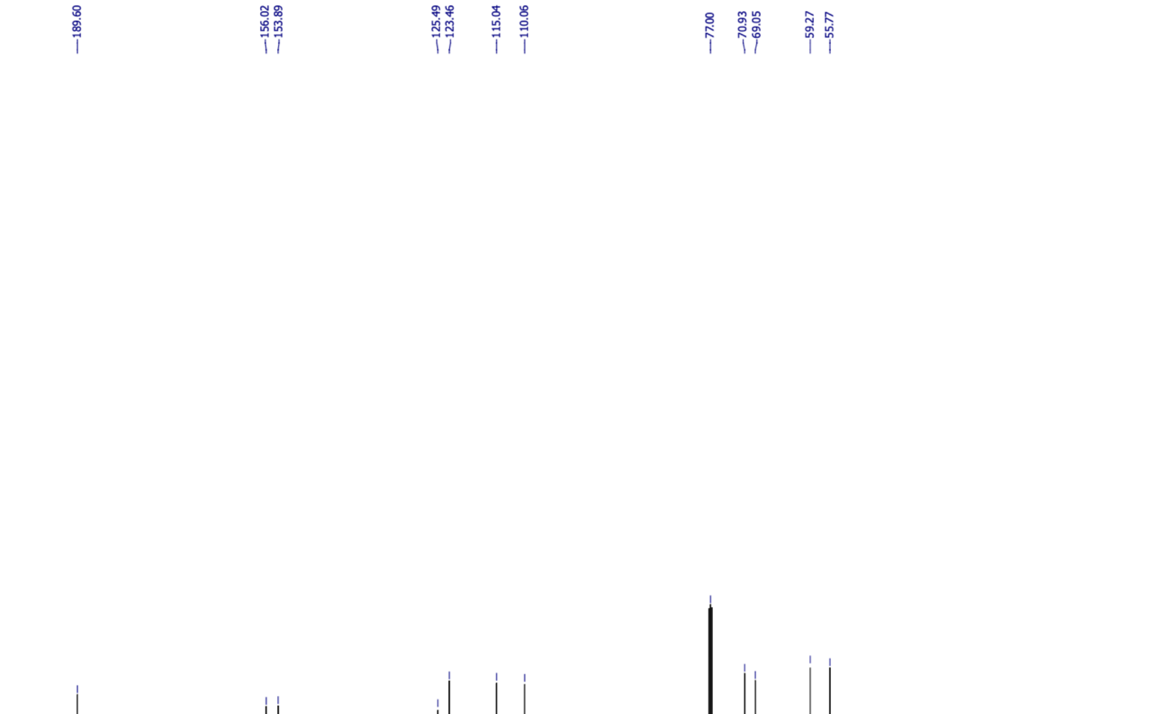

(*E*)-4-(2-methoxy-6-(2-methoxyethoxy)phenyl)but-3-en-2-one (**21b**) $^{1}H$ NMR (500 MHz, $CDCl_3$)

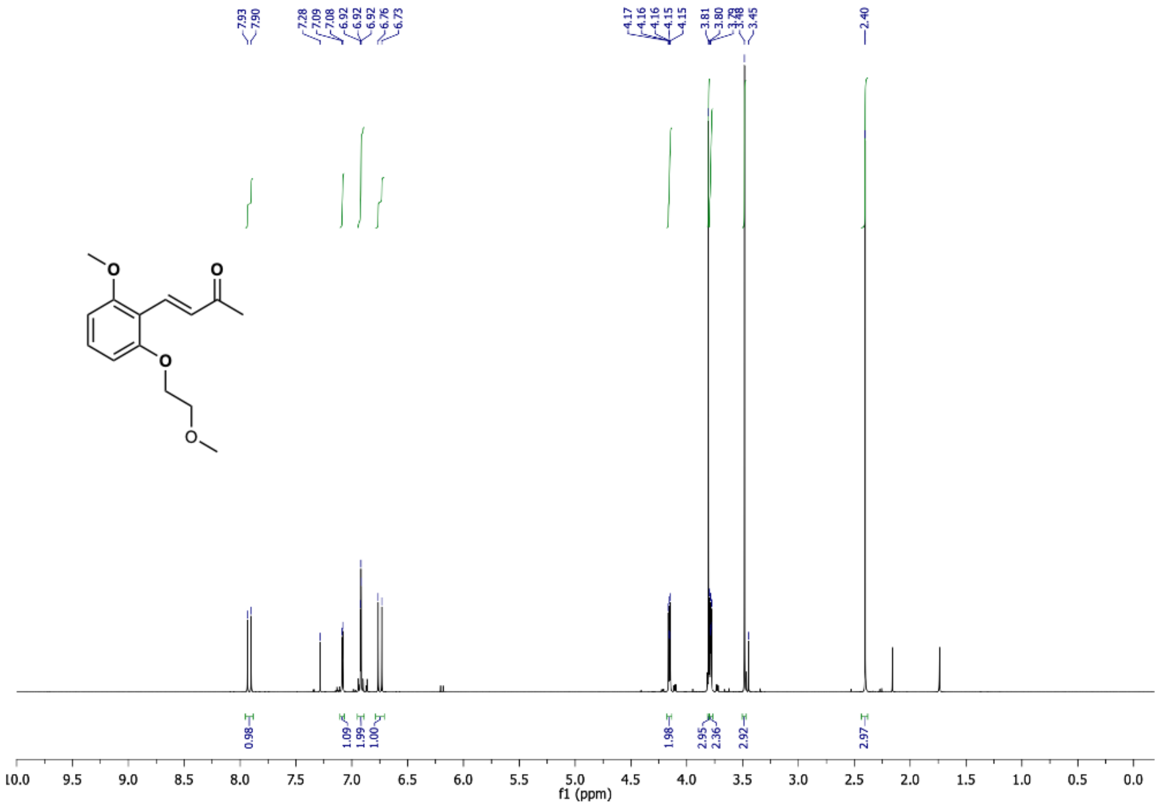

(*E*)-4-(2-methoxy-6-(2-methoxyethoxy)phenyl)but-3-en-2-one (**21b**) $^{13}C$ NMR (126 MHz, $CDCl_3$)

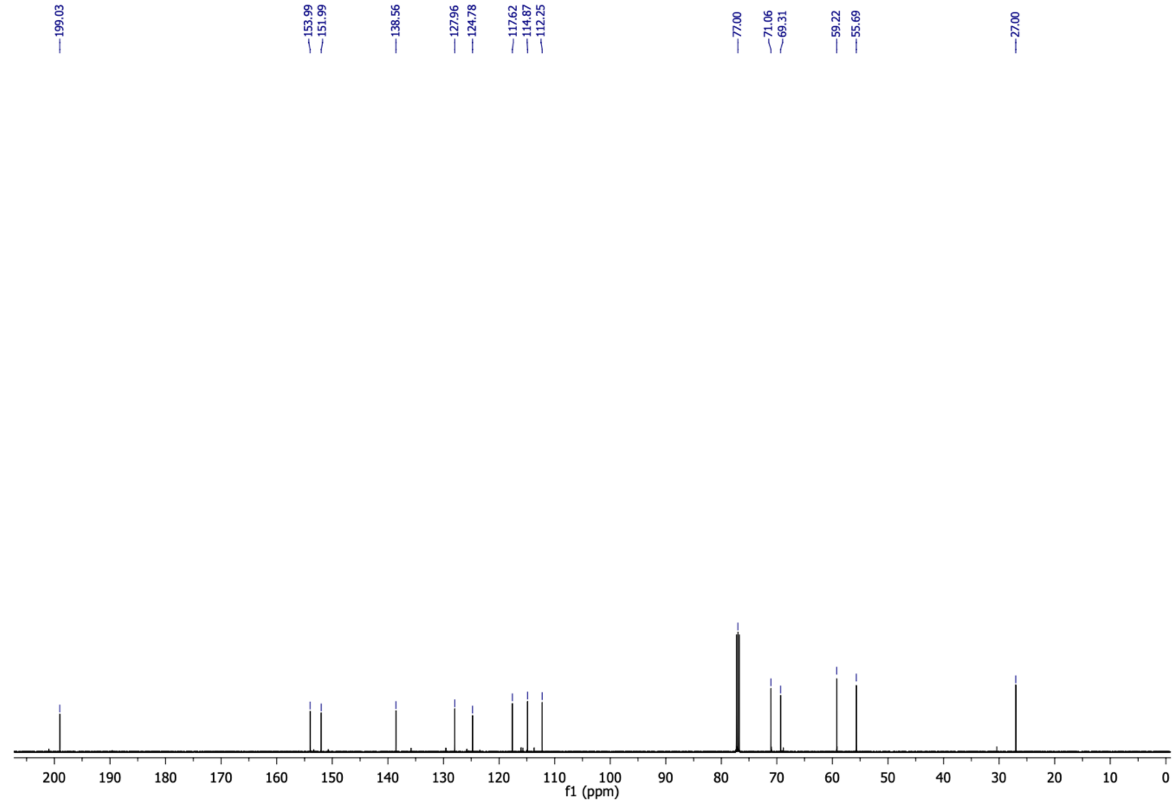

5-(2-methoxy-6-(2-methoxyethoxy)phenyl)cyclohexane-1,3-dione (**21c**) $^{1}$H NMR (500 MHz, $CD_3OD$)

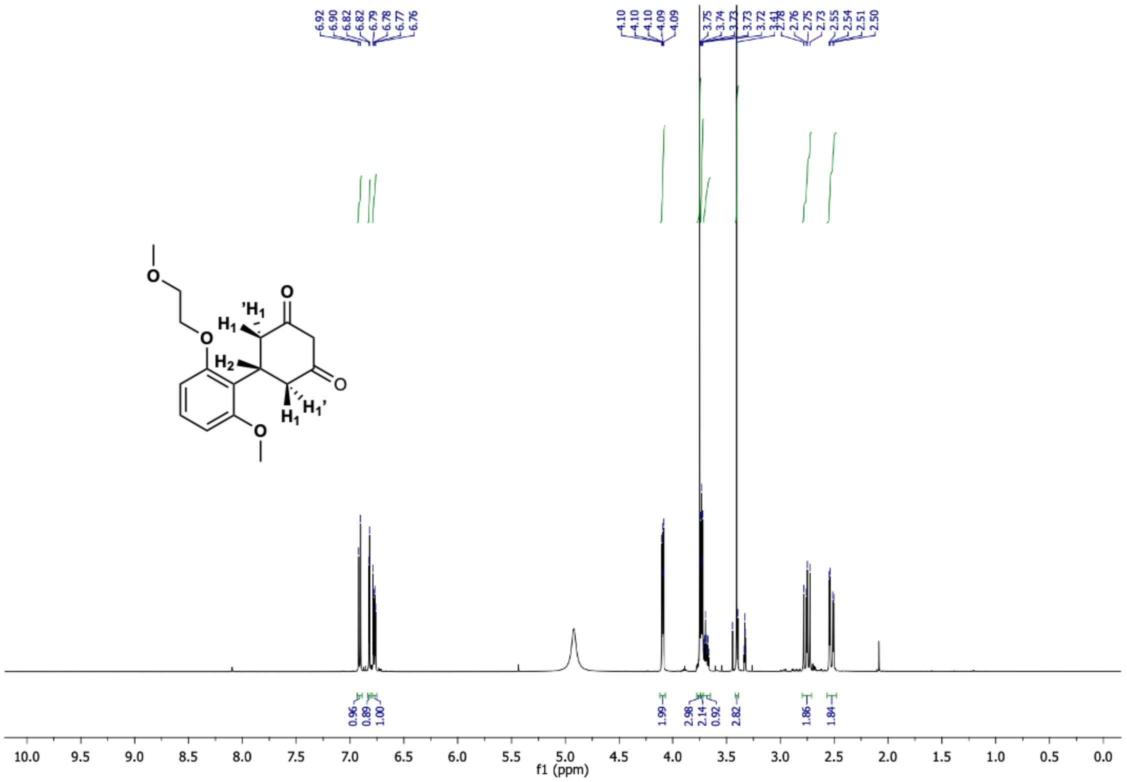

5-(2-methoxy-6-(2-methoxyethoxy)phenyl)cyclohexane-1,3-dione (**21c**) $^{13}$C NMR (126 MHz, $CD_3OD$)

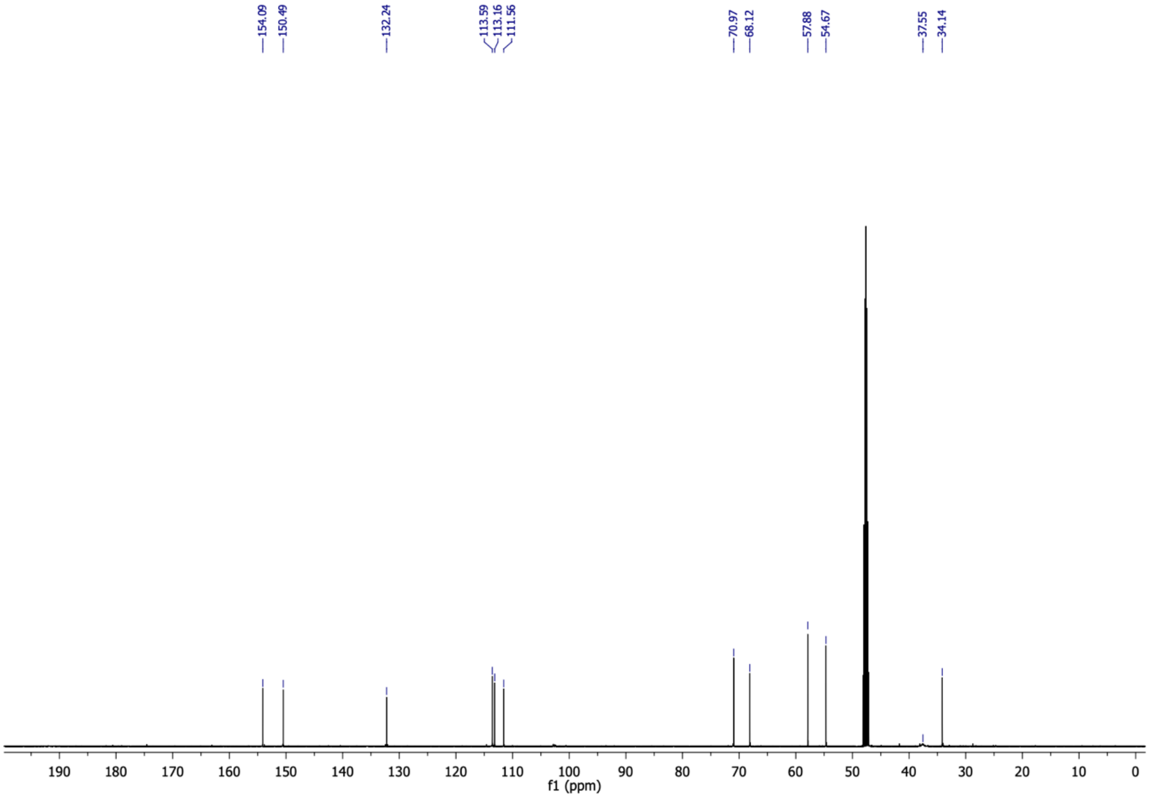

6-(2-methoxy-6-(2-methoxyethoxy)phenyl)-4-oxo-4,5,6,7-tetrahydrobenzofuran-3-carboxylic acid (**21**) $^{1}H$ NMR (500 MHz, $CDCl_3$)

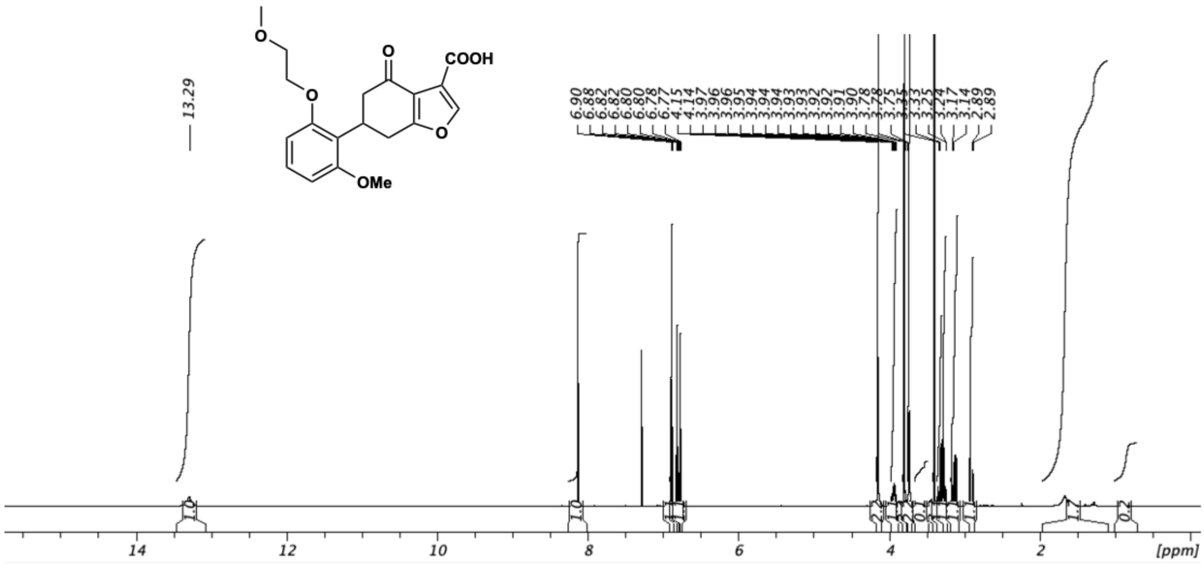

6-(2-methoxy-6-(2-methoxyethoxy)phenyl)-4-oxo-4,5,6,7-tetrahydrobenzofuran-3-carboxylic acid (**21**) $^{13}C$ NMR (126 MHz, $CDCl_3$)

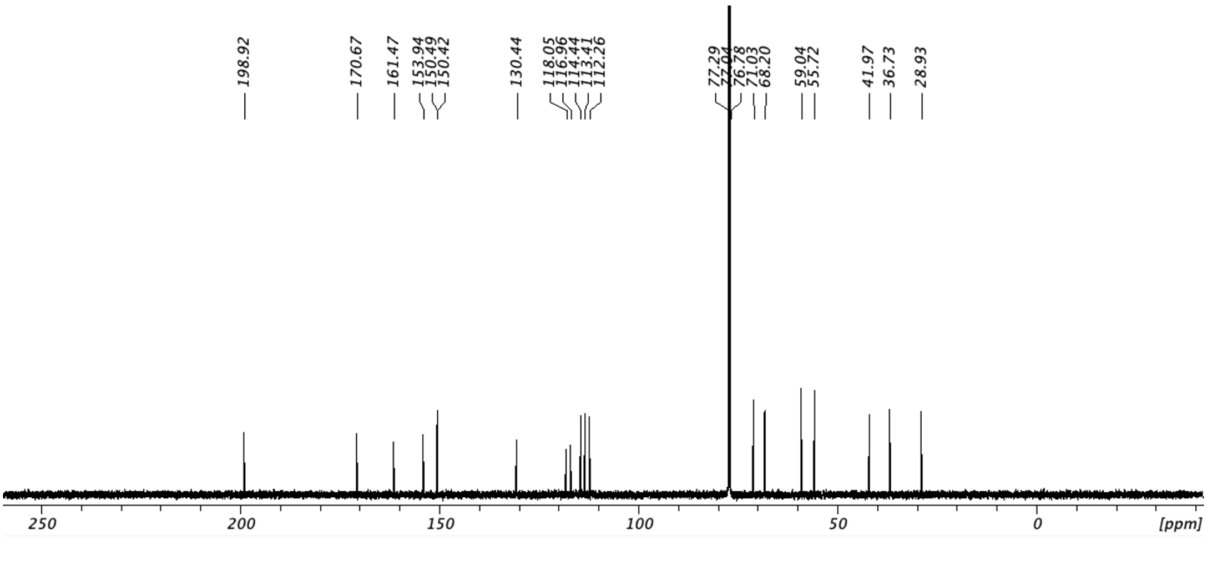

(*E*)-4-(2,3-dichlorophenyl)but-3-en-2-one (**22b**) $^{1}$H NMR (500 MHz, $CDCl_3$)

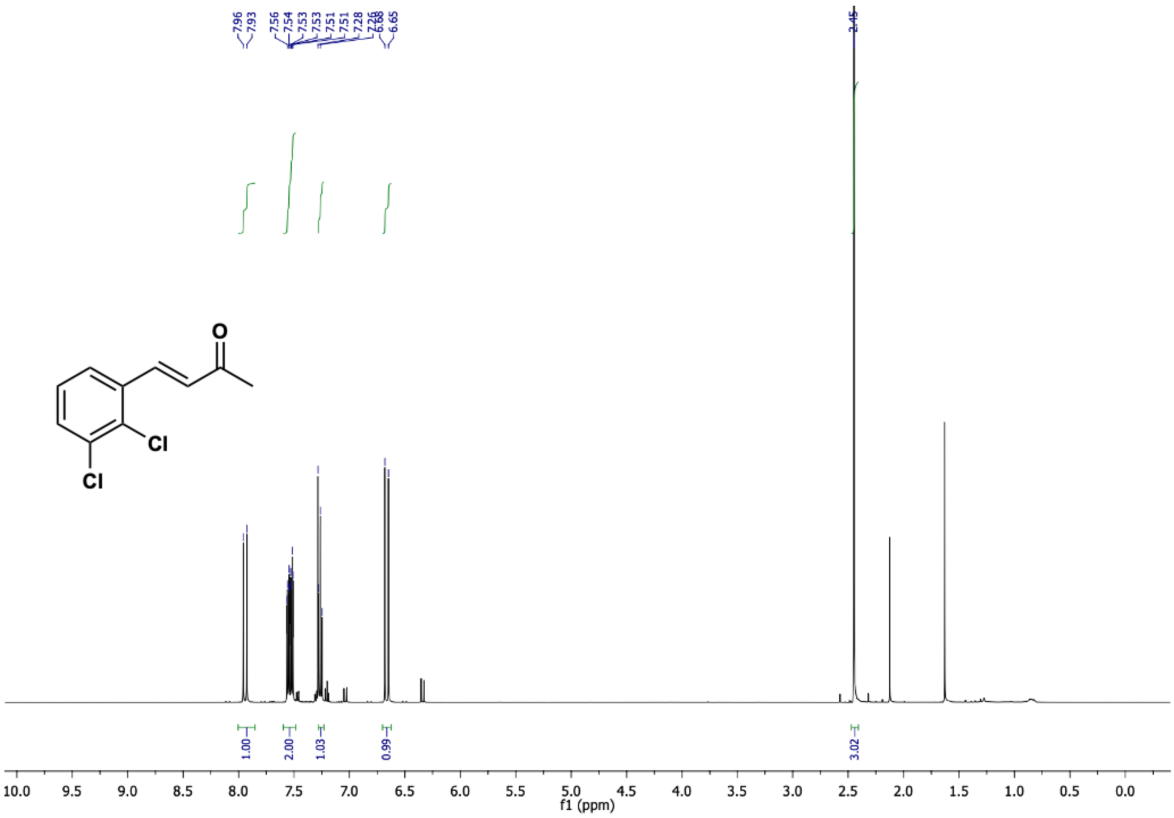

(*E*)-4-(2,3-dichlorophenyl)but-3-en-2-one (**22b**) $^{13}$C NMR (126 MHz, $CDCl_3$)

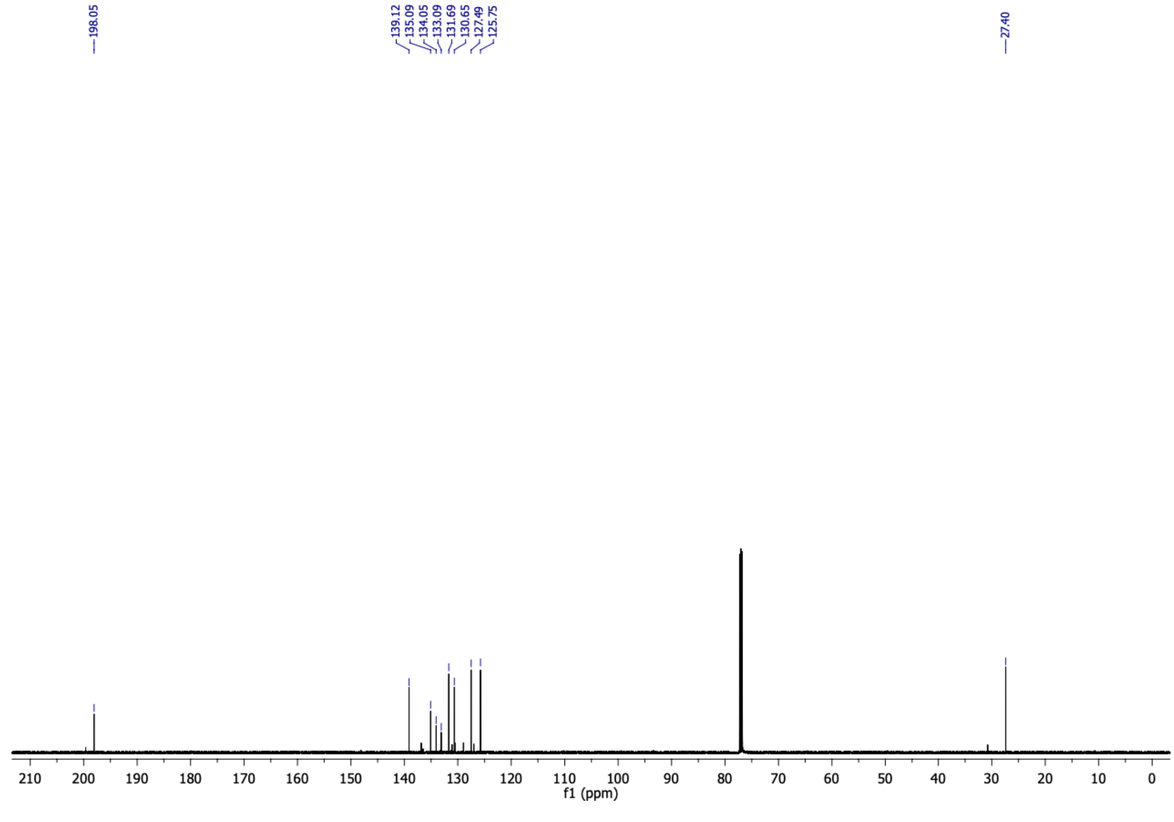

5-(2,3-dichlorophenyl)cyclohexane-1,3-dione (**22c**) $^1$H NMR (500 MHz, $d_6$-DMSO)

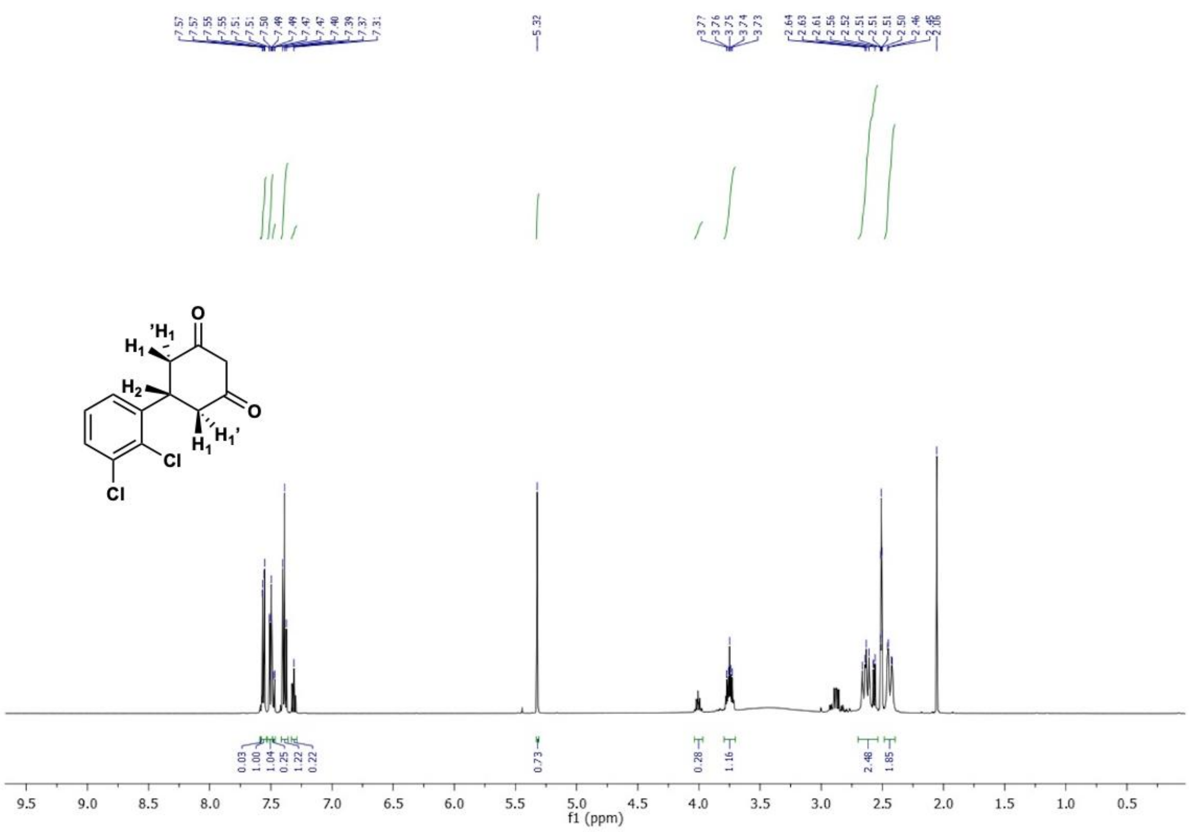

5-(2,3-dichlorophenyl)cyclohexane-1,3-dione (**22c**) $^{13}$C NMR (126 MHz, $d_6$-DMSO)

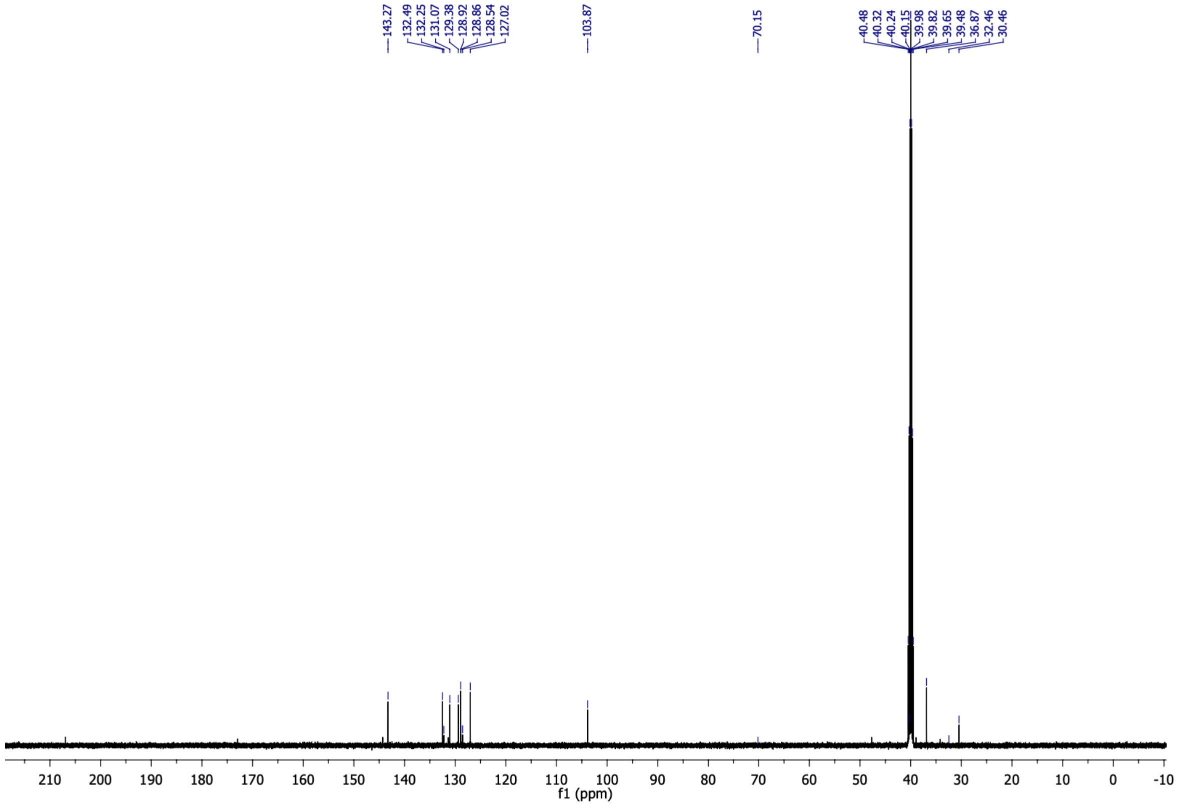

6-(2,3-dichlorophenyl)-4-oxo-4,5,6,7-tetrahydrobenzofuran-3-carboxylic acid (**22**) $^{1}$H NMR (500 MHz, $CDCl_3$)

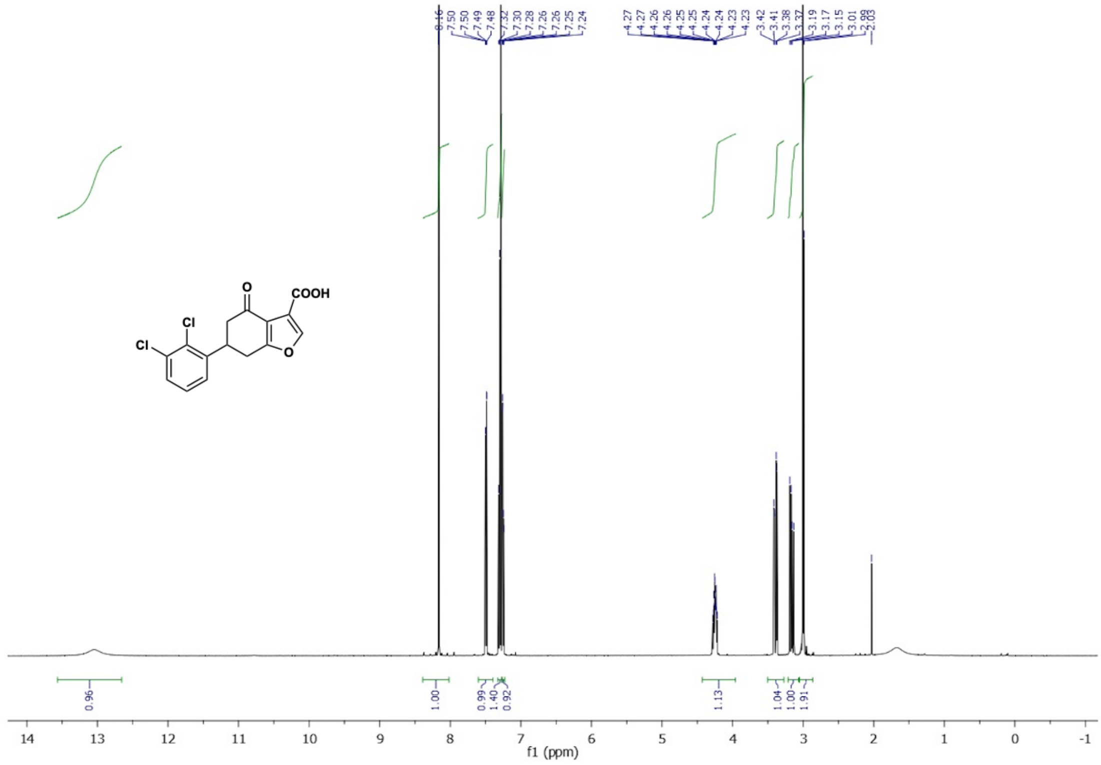

6-(2,3-dichlorophenyl)-4-oxo-4,5,6,7-tetrahydrobenzofuran-3-carboxylic acid (**22**) $^{13}$C NMR (126 MHz, $CDCl_3$)

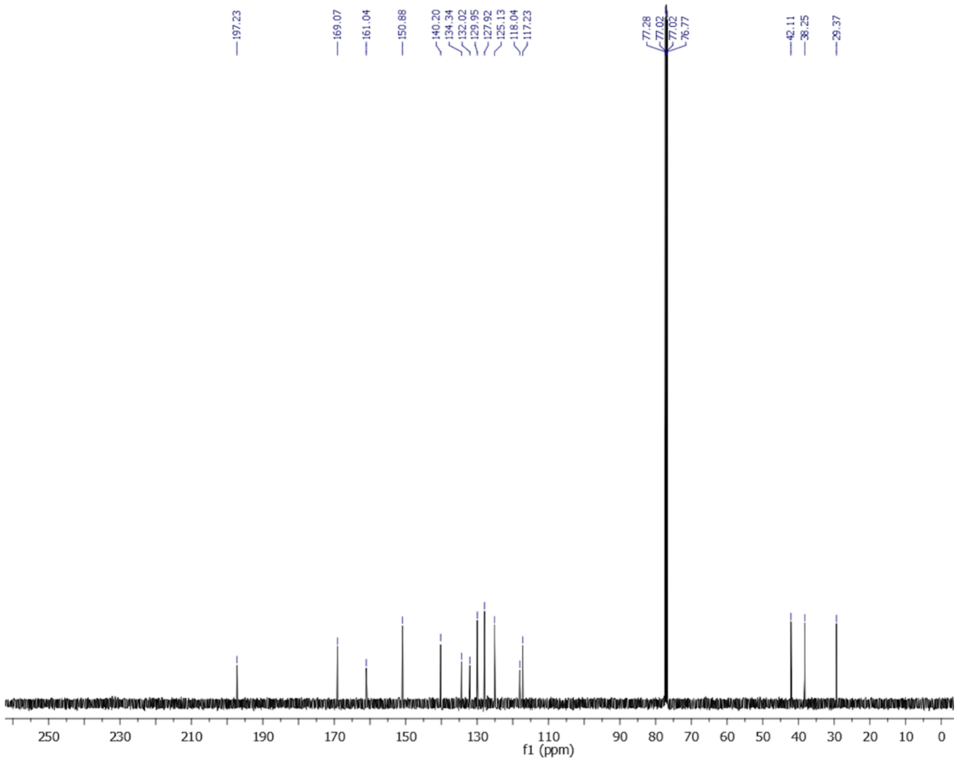

(E)-4-(2,3,6-trichlorophenyl)but-3-en-2-one (**23b**) $^{1}H$ NMR (500 MHz, $CDCl_3$)

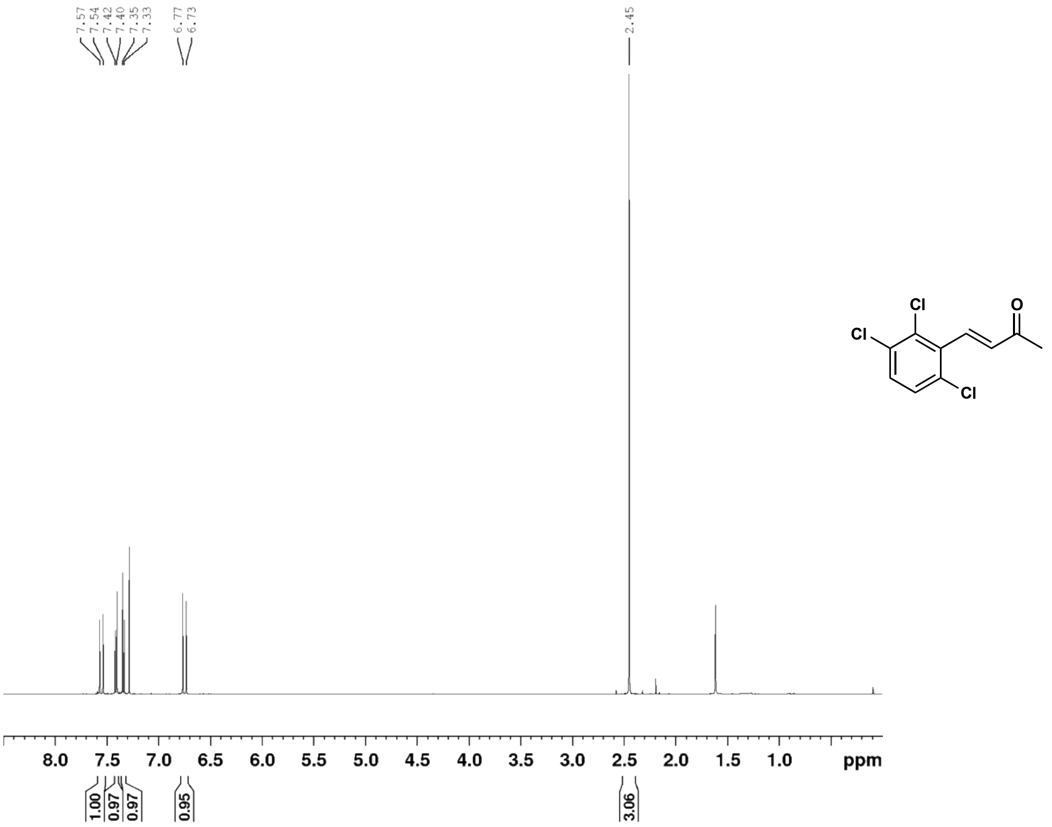

(E)-4-(2,3,6-trichlorophenyl)but-3-en-2-one (**23b**) $^{13}C$ NMR (126 MHz, $CDCl_3$)

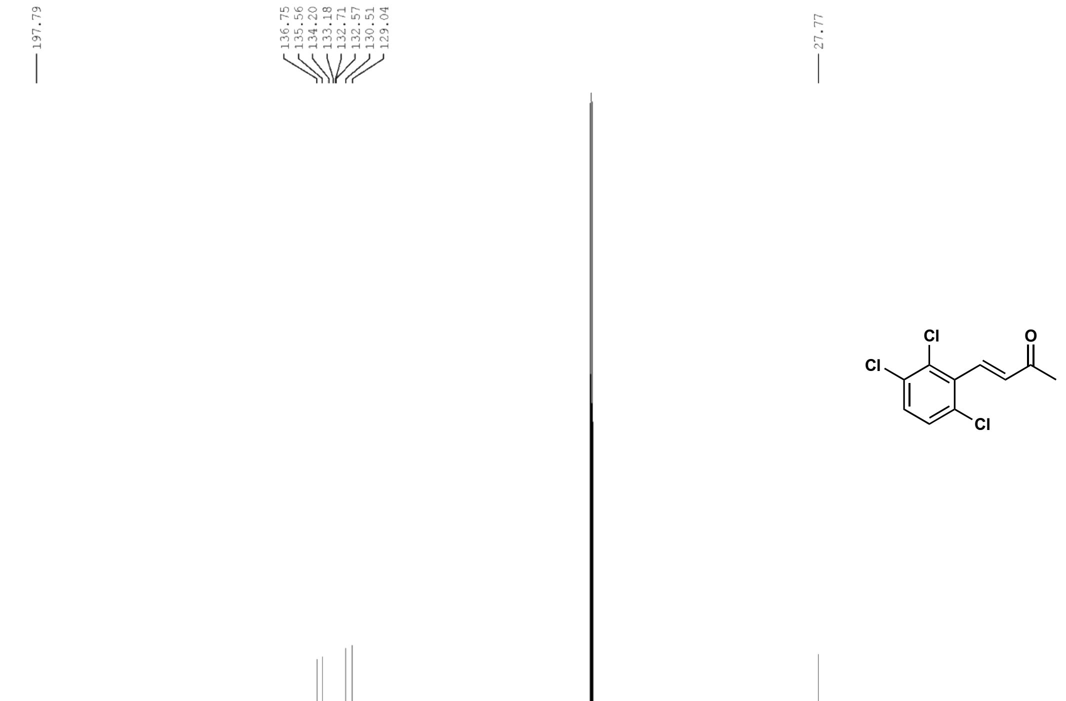

5-(2,3,6-trichlorophenyl)cyclohexane-1,3-dione (**23c**) $^{1}$H NMR (500 MHz, $CD_3OD$)

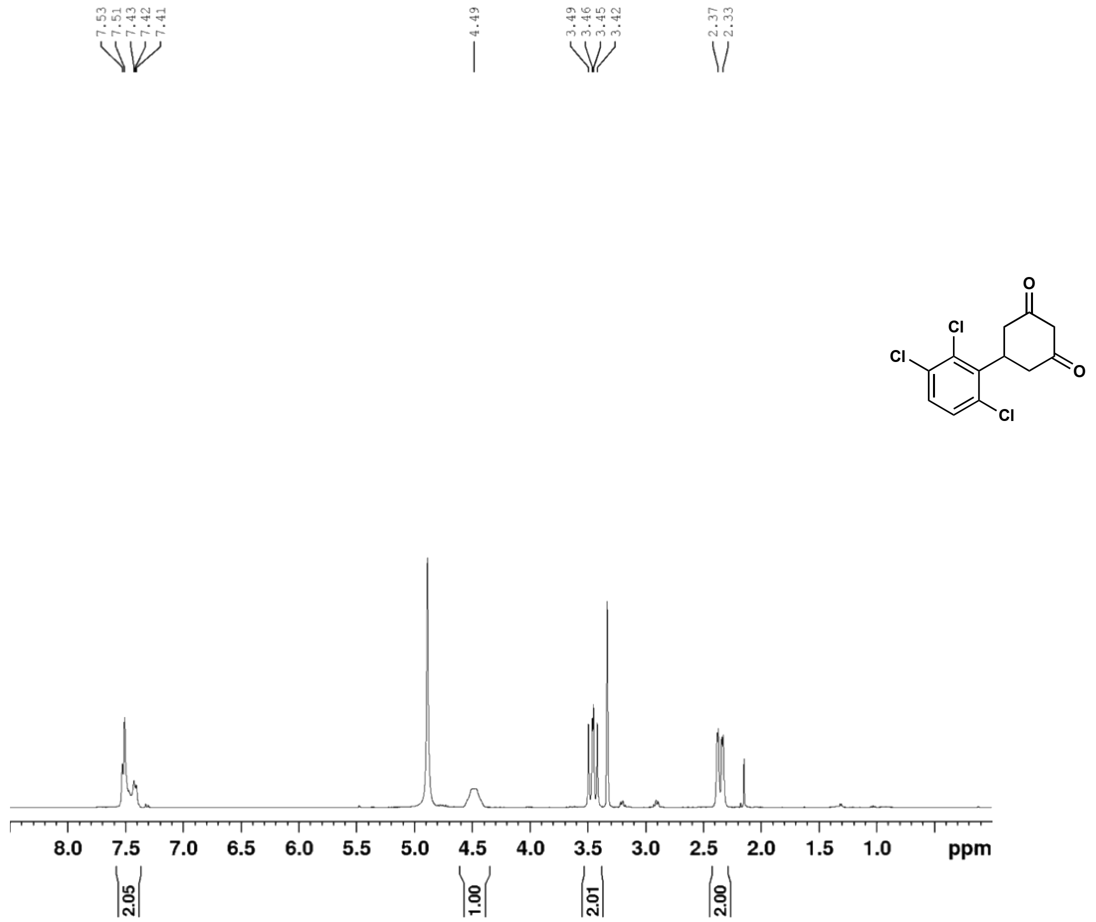

5-(2,3,6-trichlorophenyl)cyclohexane-1,3-dione (**23c**) $^{13}$C NMR (126 MHz, $CD_3OD$)

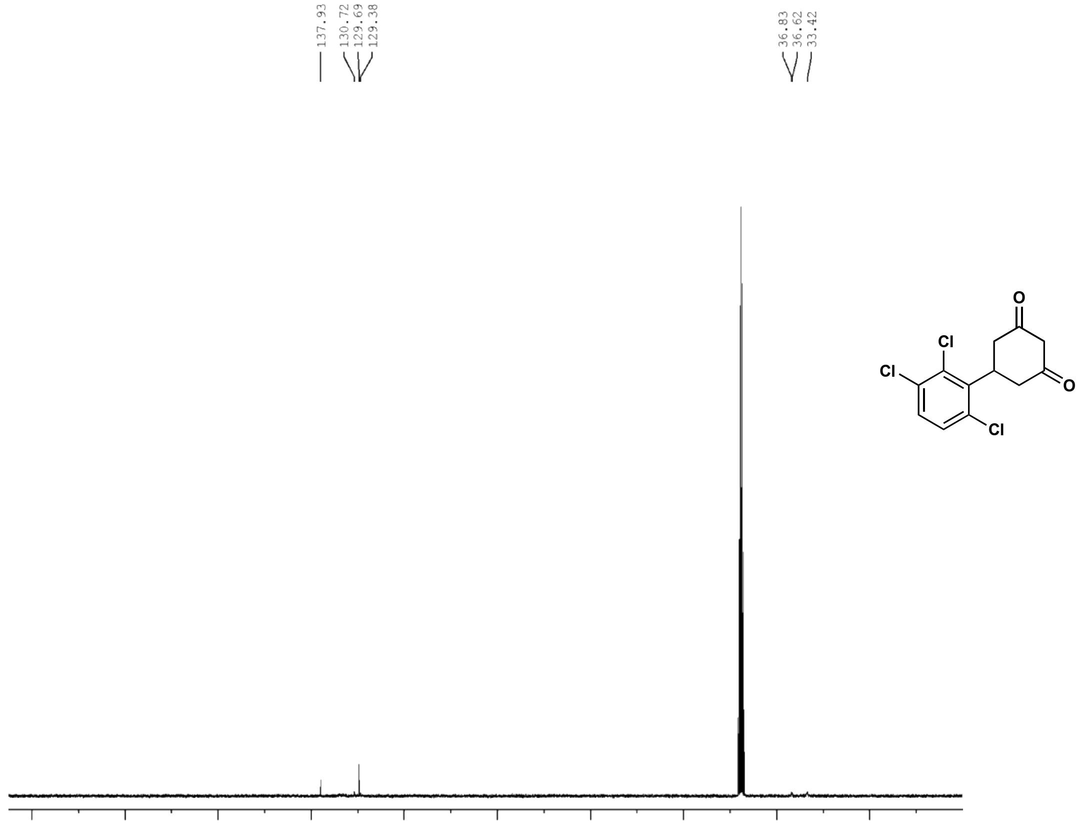

4-oxo-6-(2,3,6-trichlorophenyl)-4,5,6,7-tetrahydrobenzofuran-3-carboxylic acid (**23**) $^{1}$H NMR (500 MHz, $CDCl_3$)

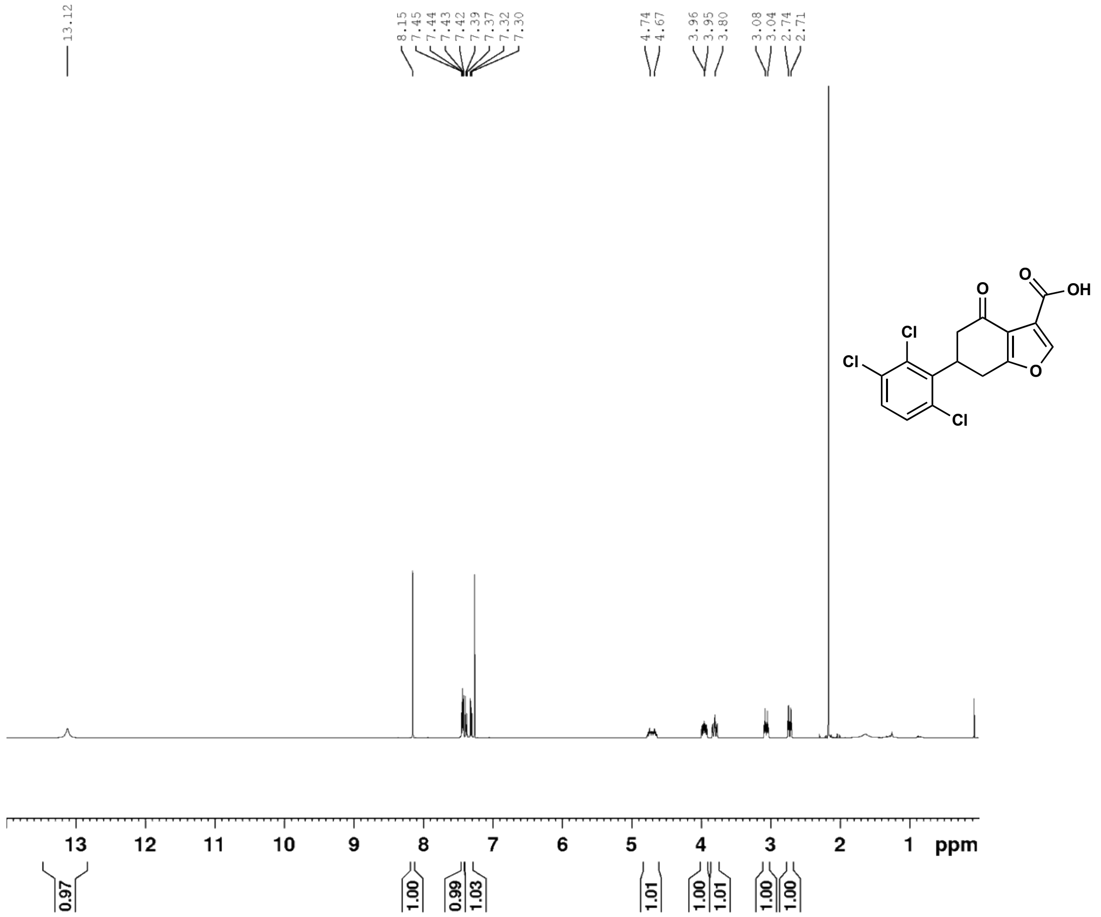

4-oxo-6-(2,3,6-trichlorophenyl)-4,5,6,7-tetrahydrobenzofuran-3-carboxylic acid (**23**) $^{13}$C NMR (126 MHz, $CDCl_3$)

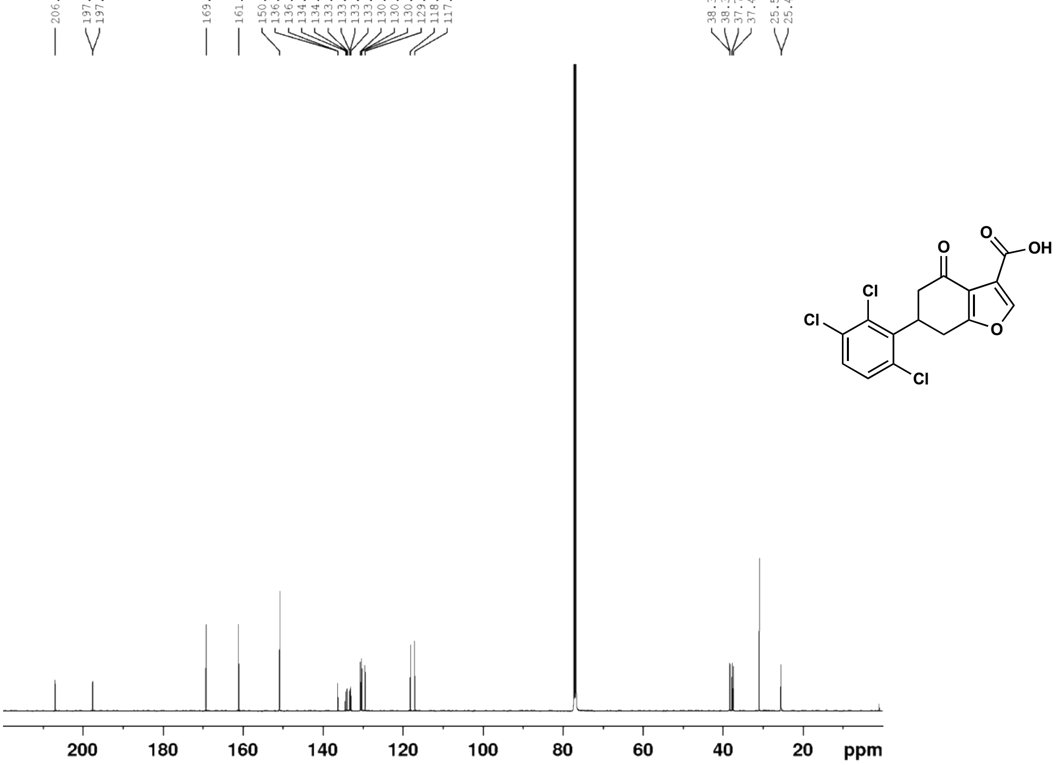

(E)-4-(4-chlorophenyl)but-3-en-2-one (**24b**) $^{1}$H NMR (400 MHz, $CDCl_3$)

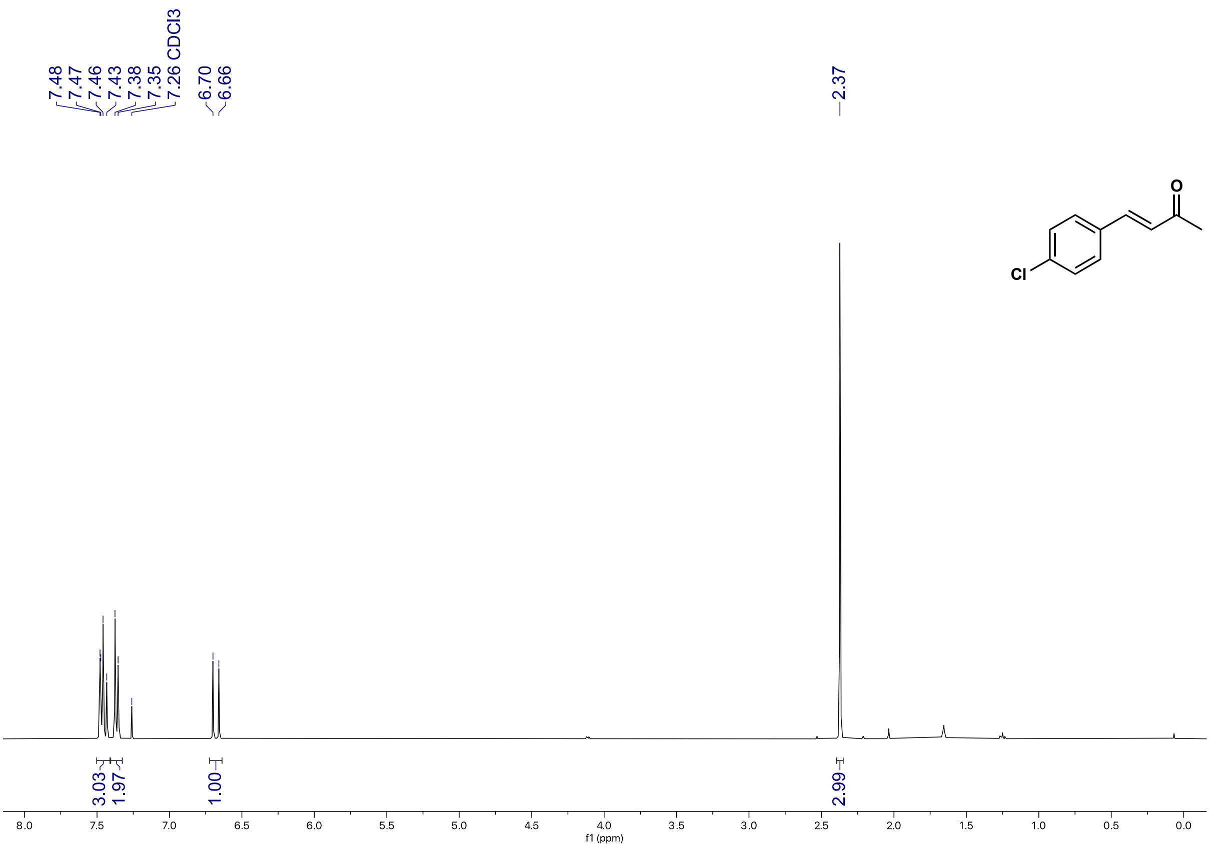

(E)-4-(4-chlorophenyl)but-3-en-2-one (**24b**) $^{13}$C NMR (101 MHz, $CDCl_3$)

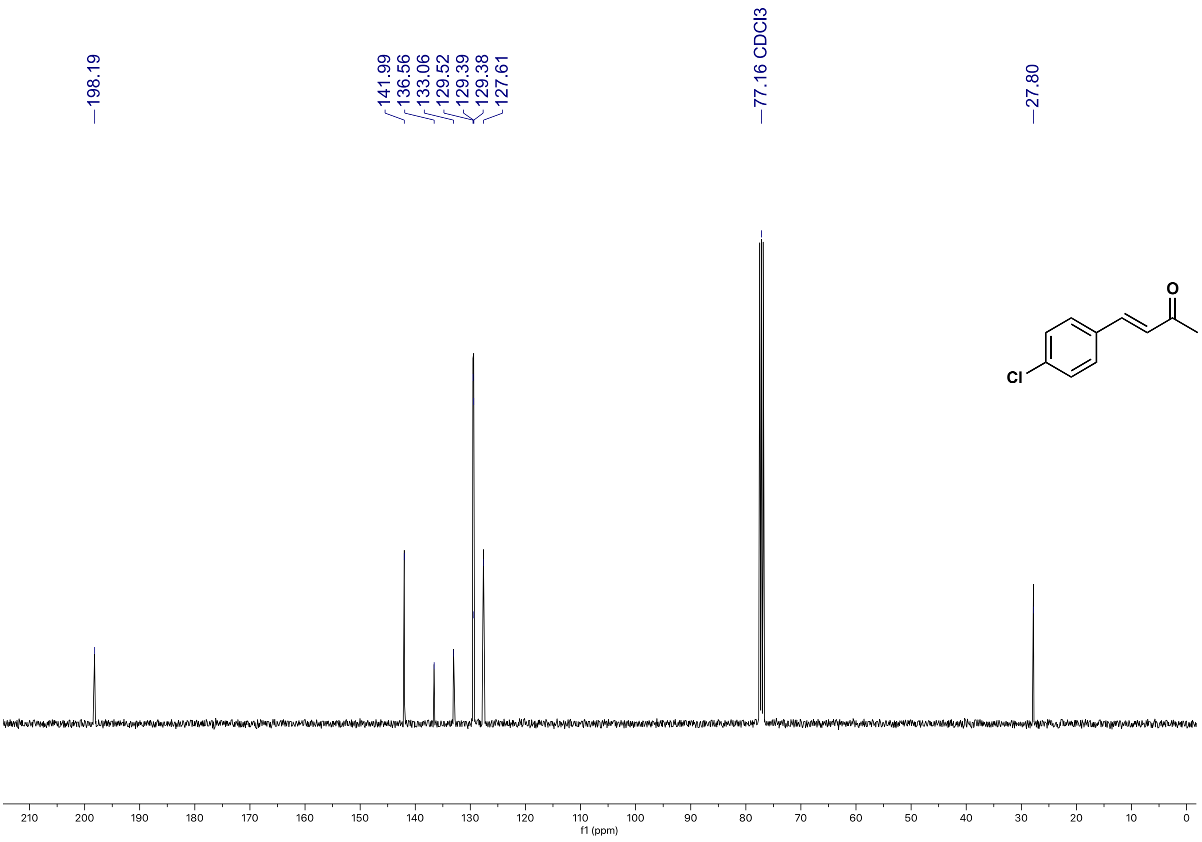

5-(4-chlorophenyl)cyclohexane-1,3-dione (**24c**) $^{1}$H NMR (500 MHz, $d_6$-DMSO)

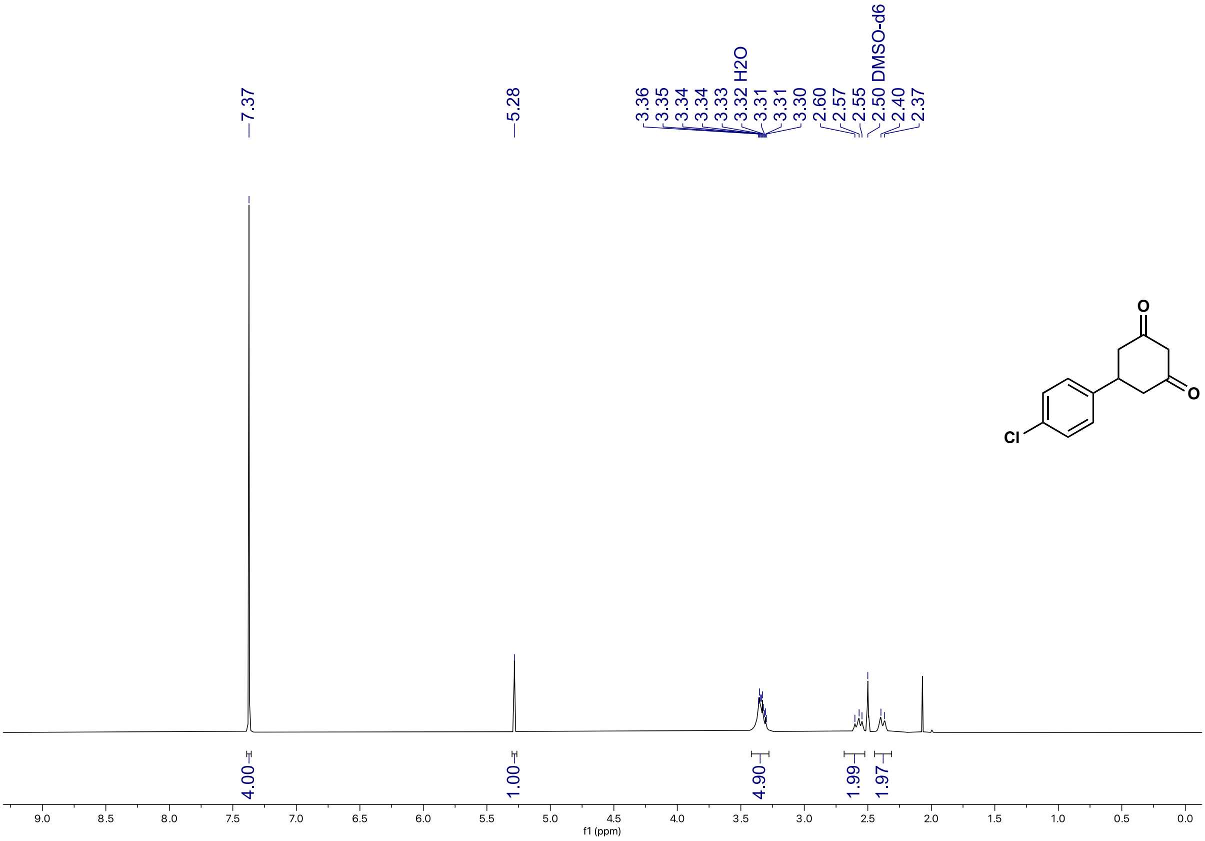

5-(4-chlorophenyl)cyclohexane-1,3-dione (**24c**) $^{13}$C NMR (126 MHz, $d_6$-DMSO)

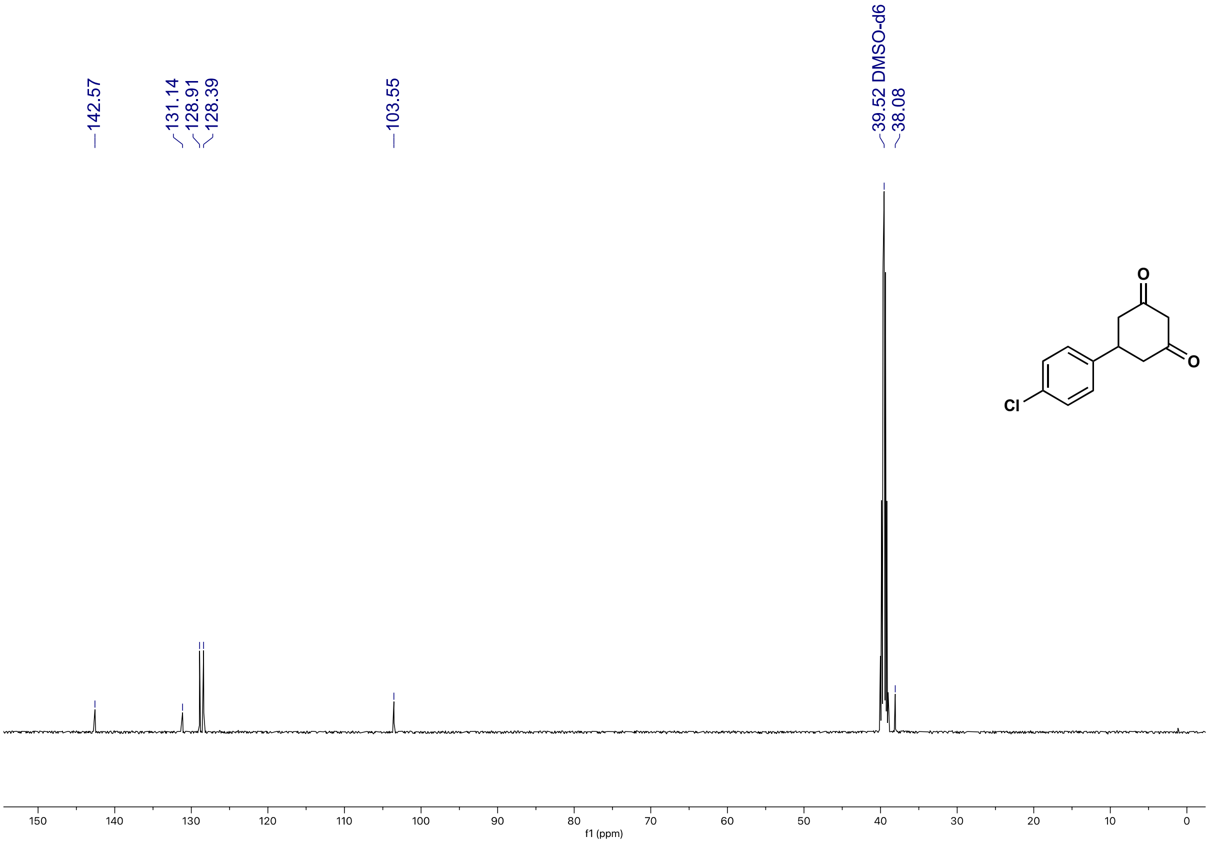

6-(4-chlorophenyl)-4-oxo-4,5,6,7-tetrahydrobenzofuran-3-carboxylic acid (**24**) $^{1}H$ NMR (500 MHz, $CDCl_3$)

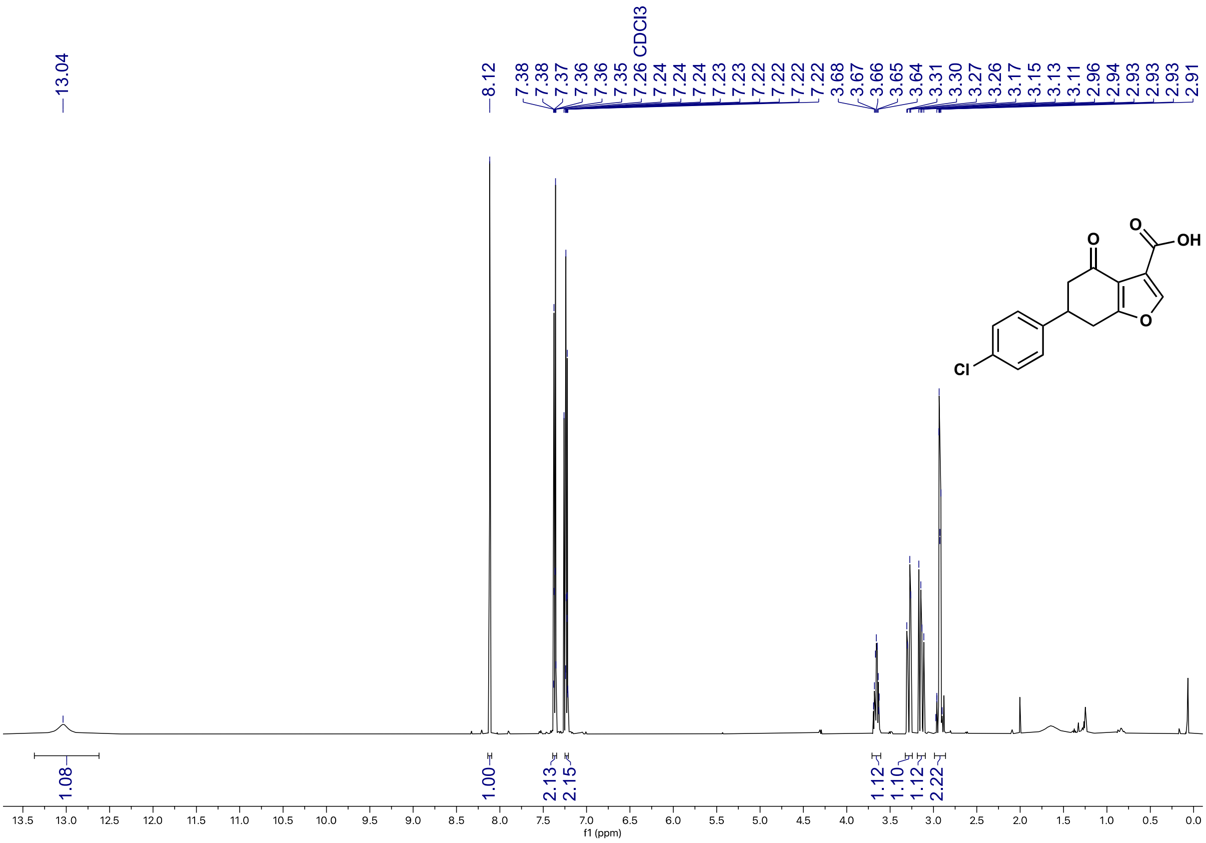

6-(4-chlorophenyl)-4-oxo-4,5,6,7-tetrahydrobenzofuran-3-carboxylic acid (**24**) $^{13}C$ NMR (126 MHz, $CDCl_3$)

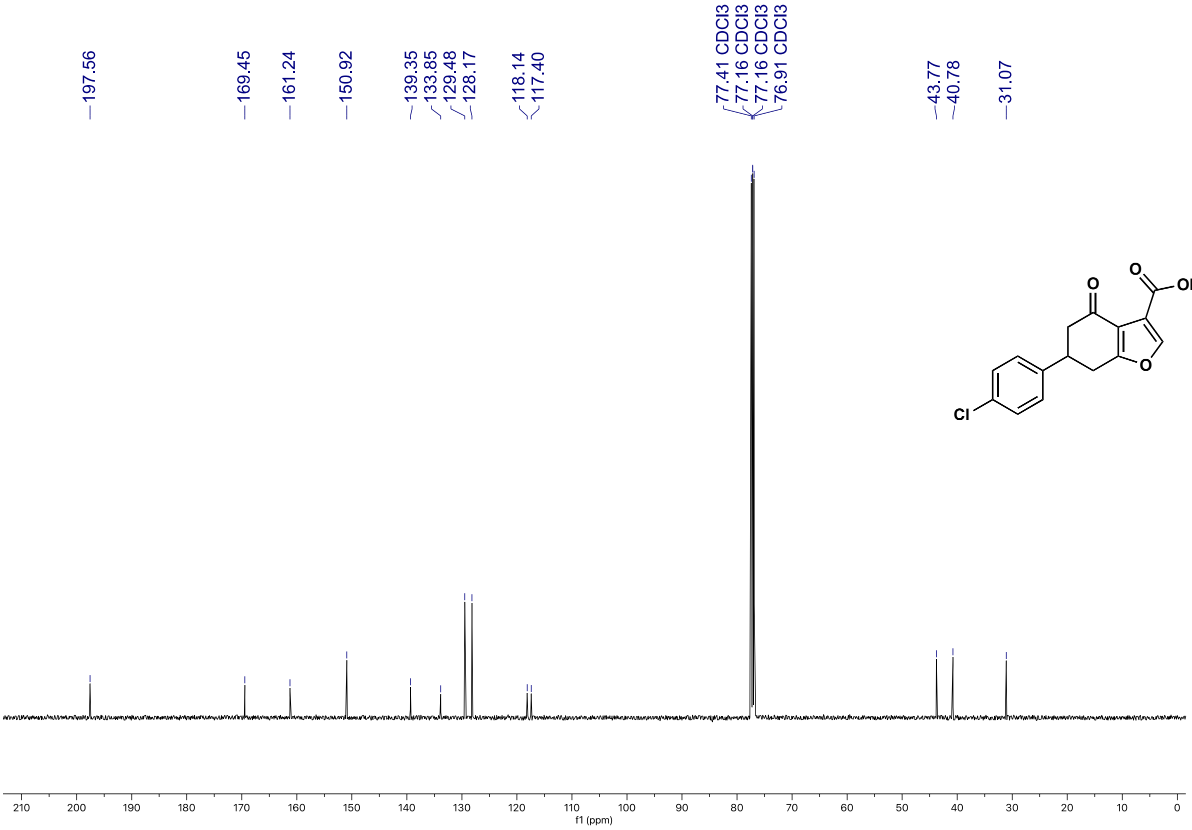

ethyl 6-(4-chlorophenyl)-4-oxo-4,5,6,7-tetrahydrobenzofuran-3-carboxylate (**24d**) $^{1}$H NMR (700 MHz, $CDCl_3$)

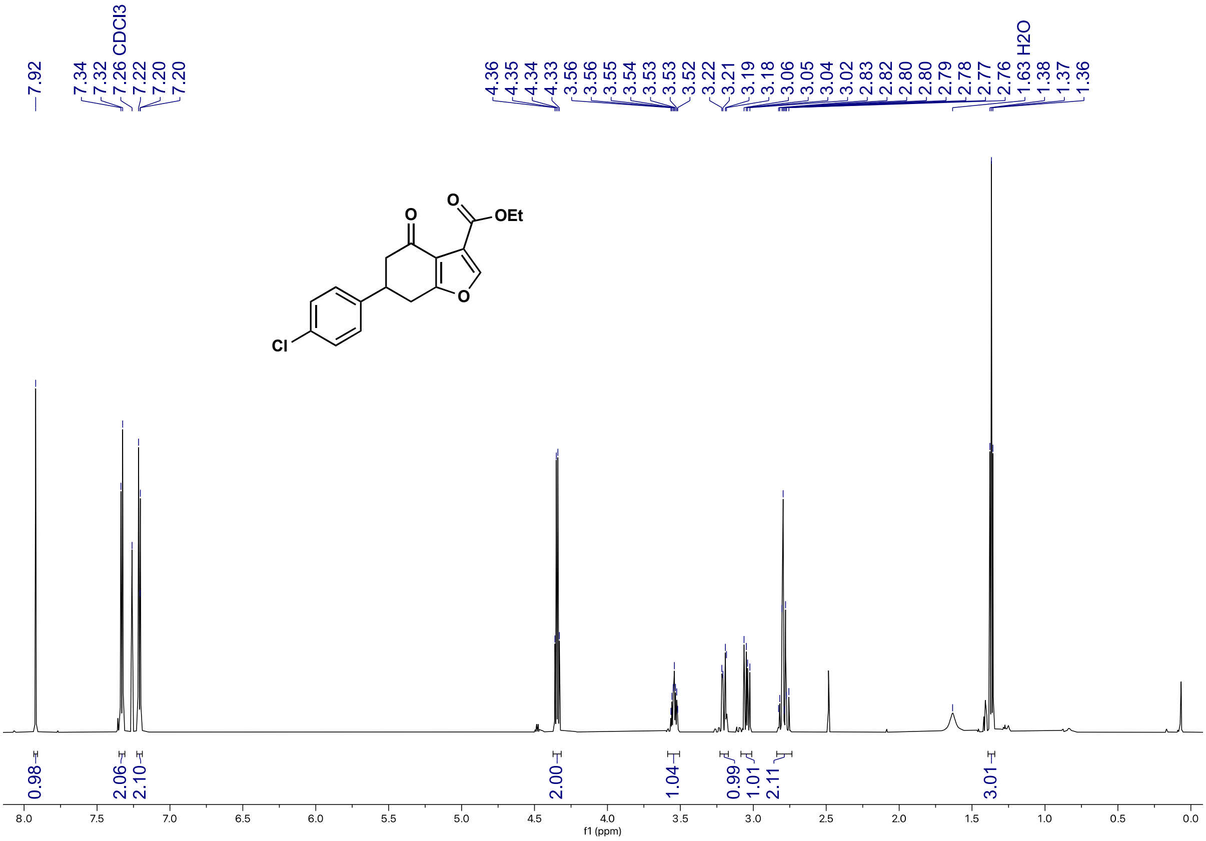

ethyl 6-(4-chlorophenyl)-4-oxo-4,5,6,7-tetrahydrobenzofuran-3-carboxylate (**24d**) $^{13}$C NMR (176 MHz, $CDCl_3$)

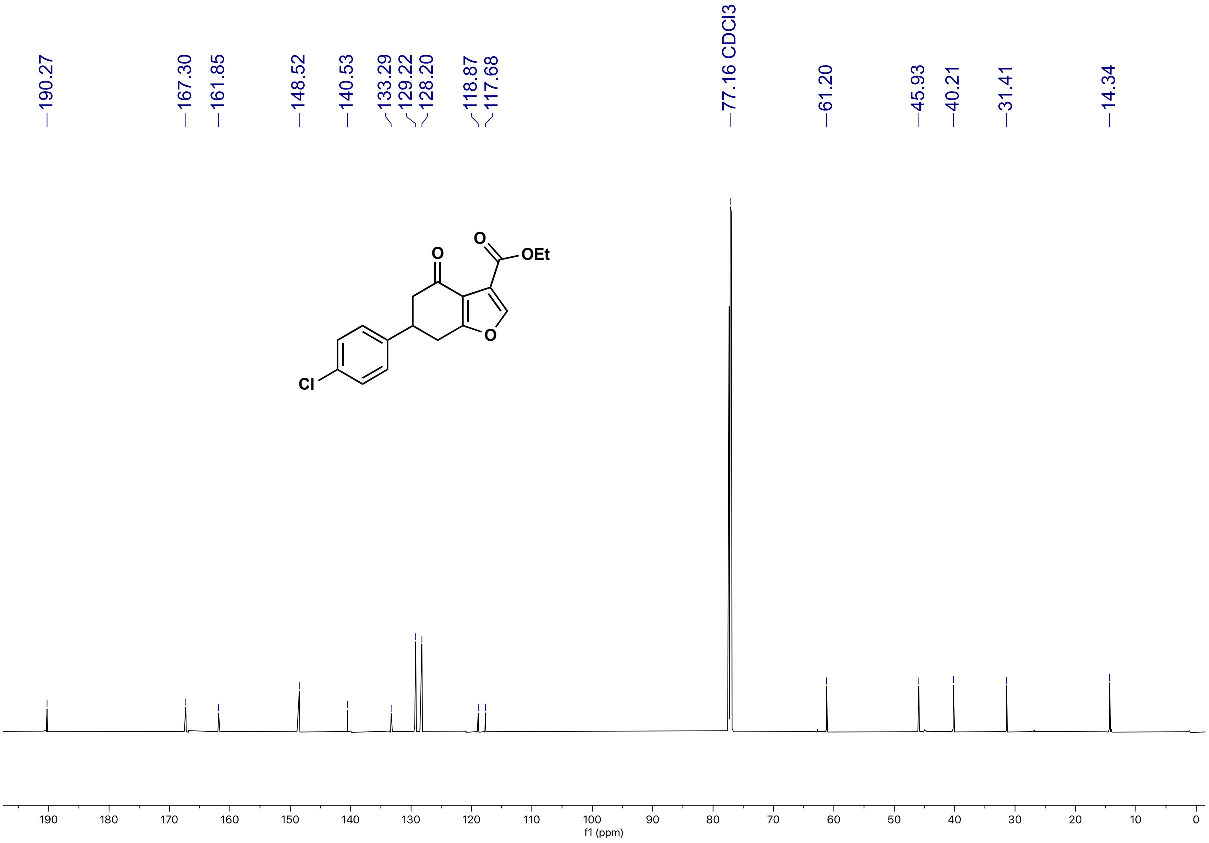

(E)-4-(2-chloro-3-methoxyphenyl)but-3-en-2-one (**25b**) $^{1}H$ NMR (500 MHz, $CDCl_3$)

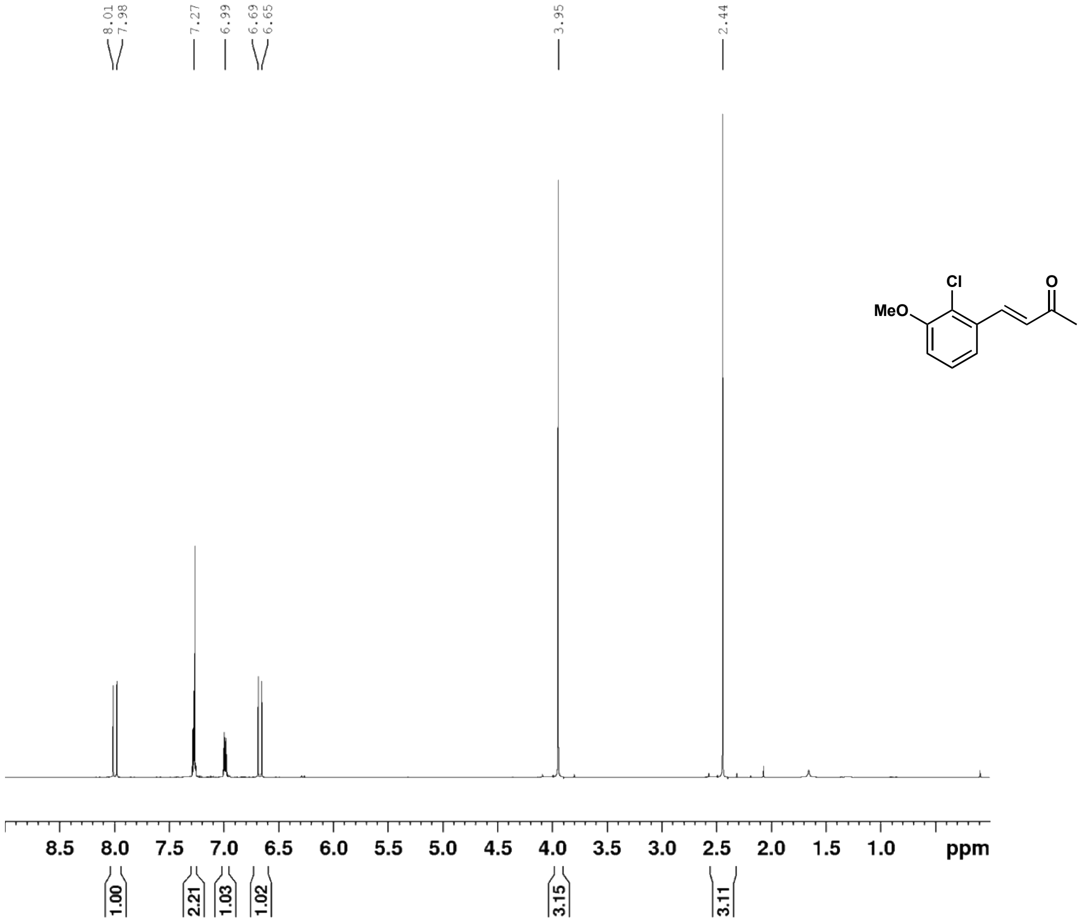

(E)-4-(2-chloro-3-methoxyphenyl)but-3-en-2-one (**25b**) $^{13}C$ NMR (126 MHz, $CDCl_3$)

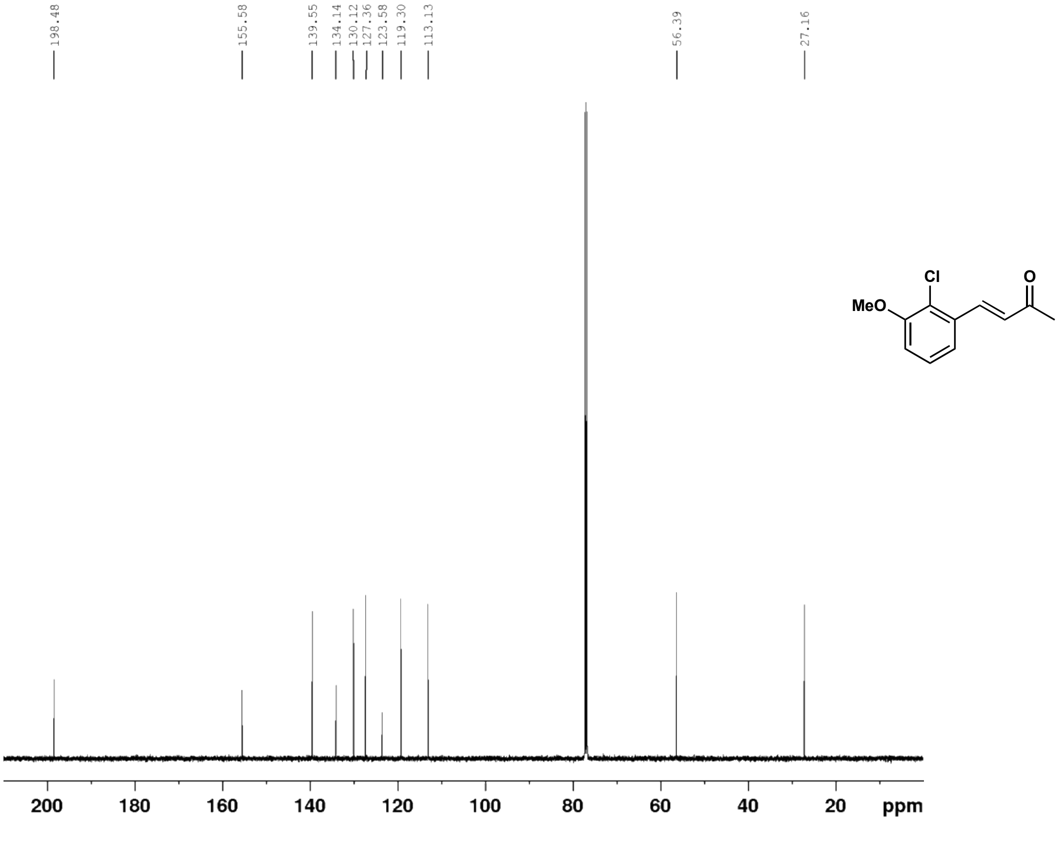

5-(2-chloro-3-methoxyphenyl)cyclohexane-1,3-dione (**25c**) $^{1}$H NMR (500 MHz, $CD_3OD$)

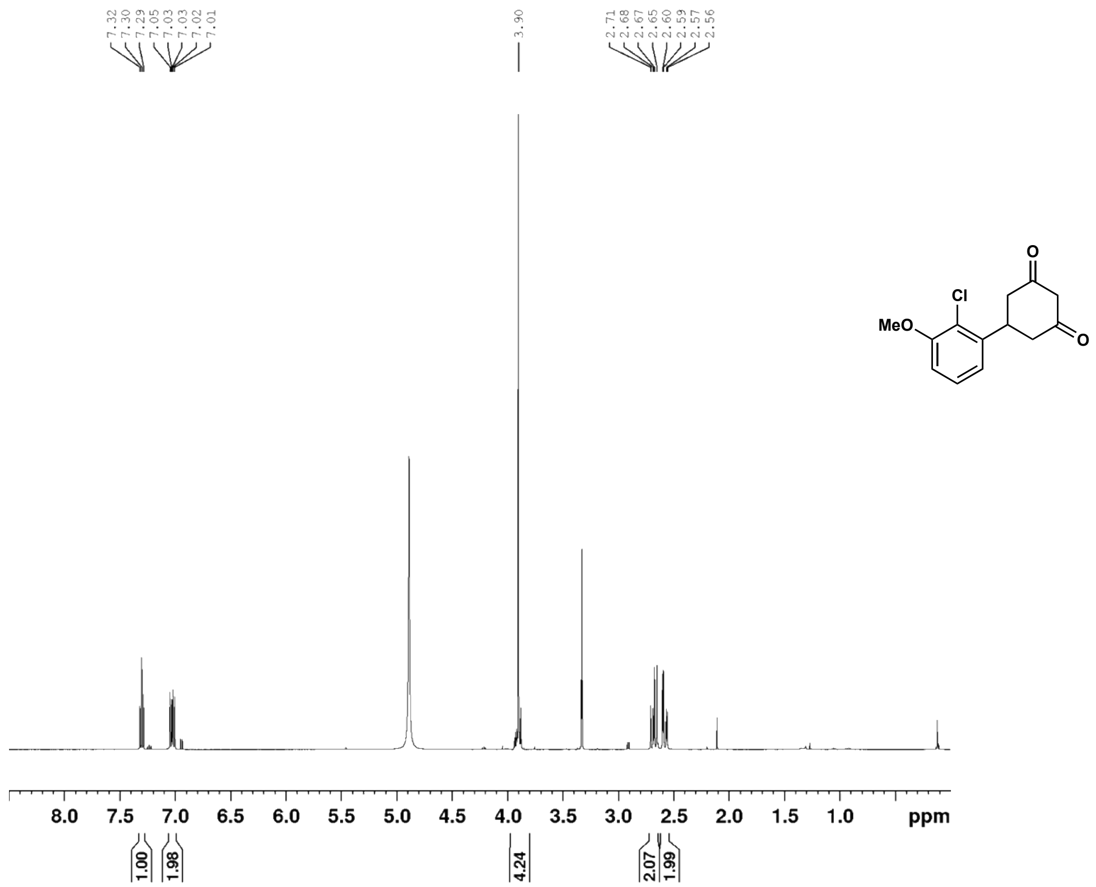

5-(2-chloro-3-methoxyphenyl)cyclohexane-1,3-dione (**25c**) $^{13}$C NMR (126 MHz, $CD_3OD$)

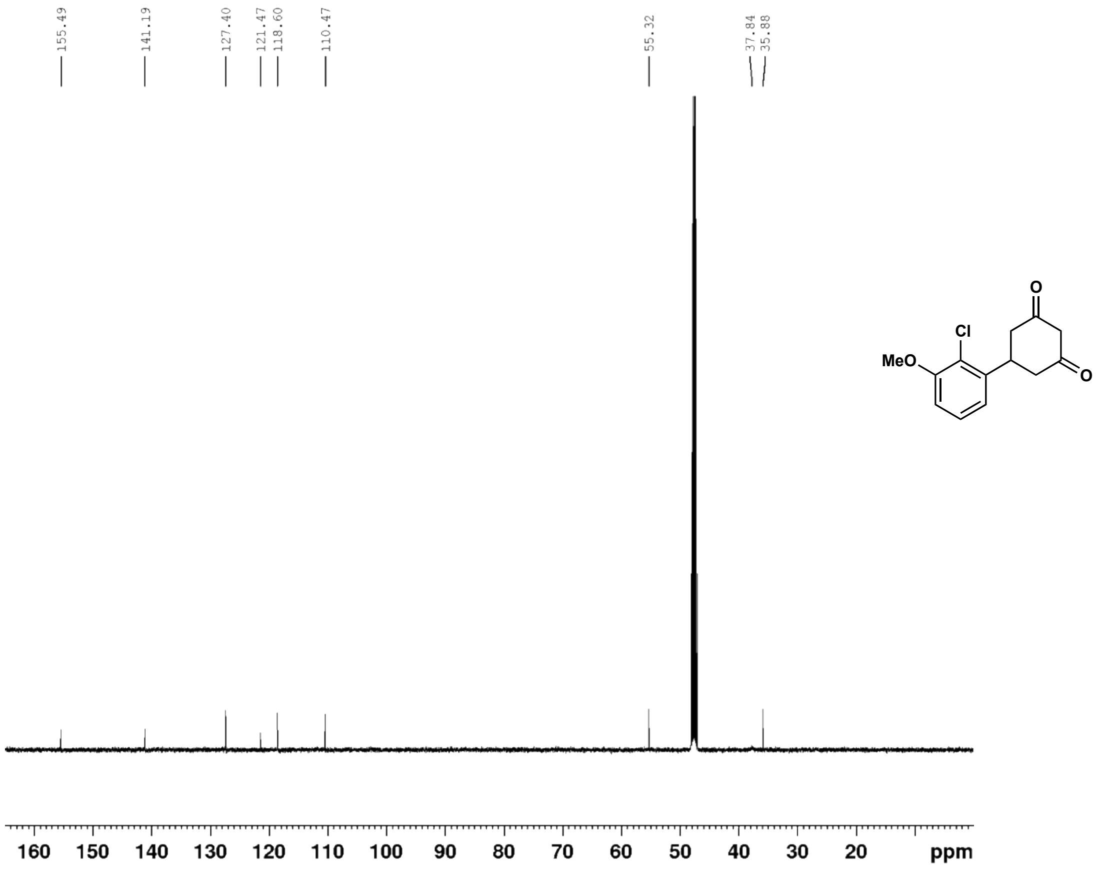

6-(2-chloro-3-methoxyphenyl)-4-oxo-4,5,6,7-tetrahydrobenzofuran-3-carboxylic acid (**25**) $^{1}$H NMR (500 MHz, $CDCl_3$)

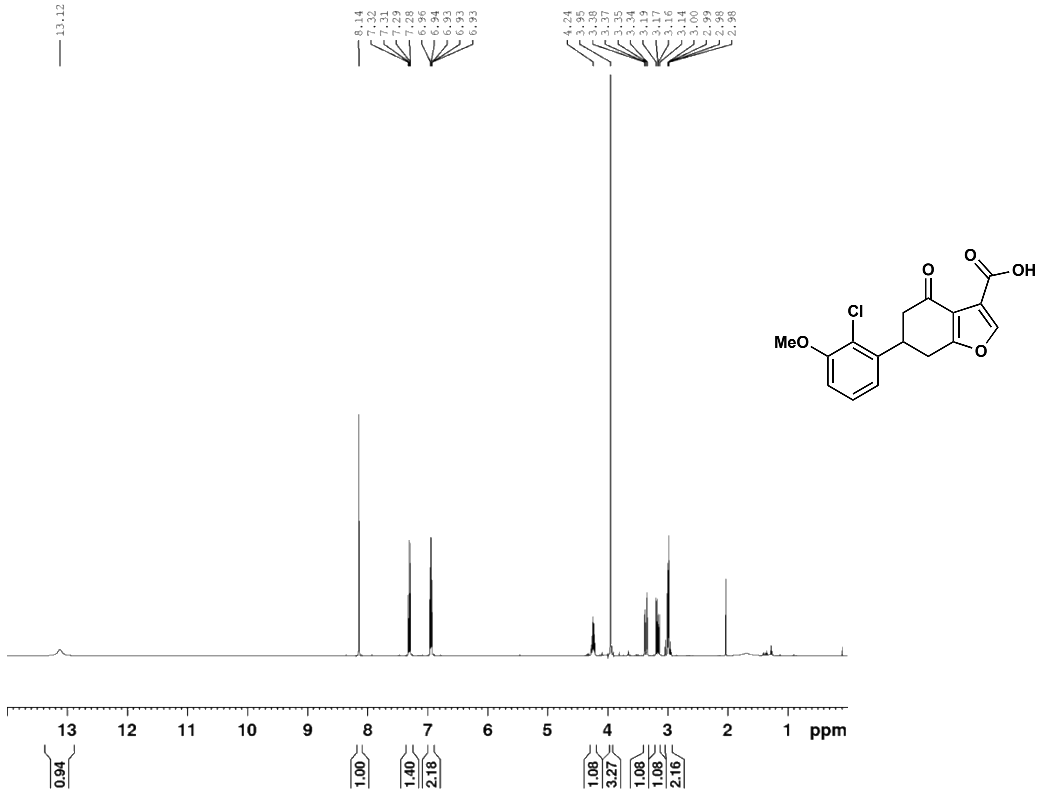

6-(2-chloro-3-methoxyphenyl)-4-oxo-4,5,6,7-tetrahydrobenzofuran-3-carboxylic acid (**25**) $^{13}$C NMR (126 MHz, $CDCl_3$)

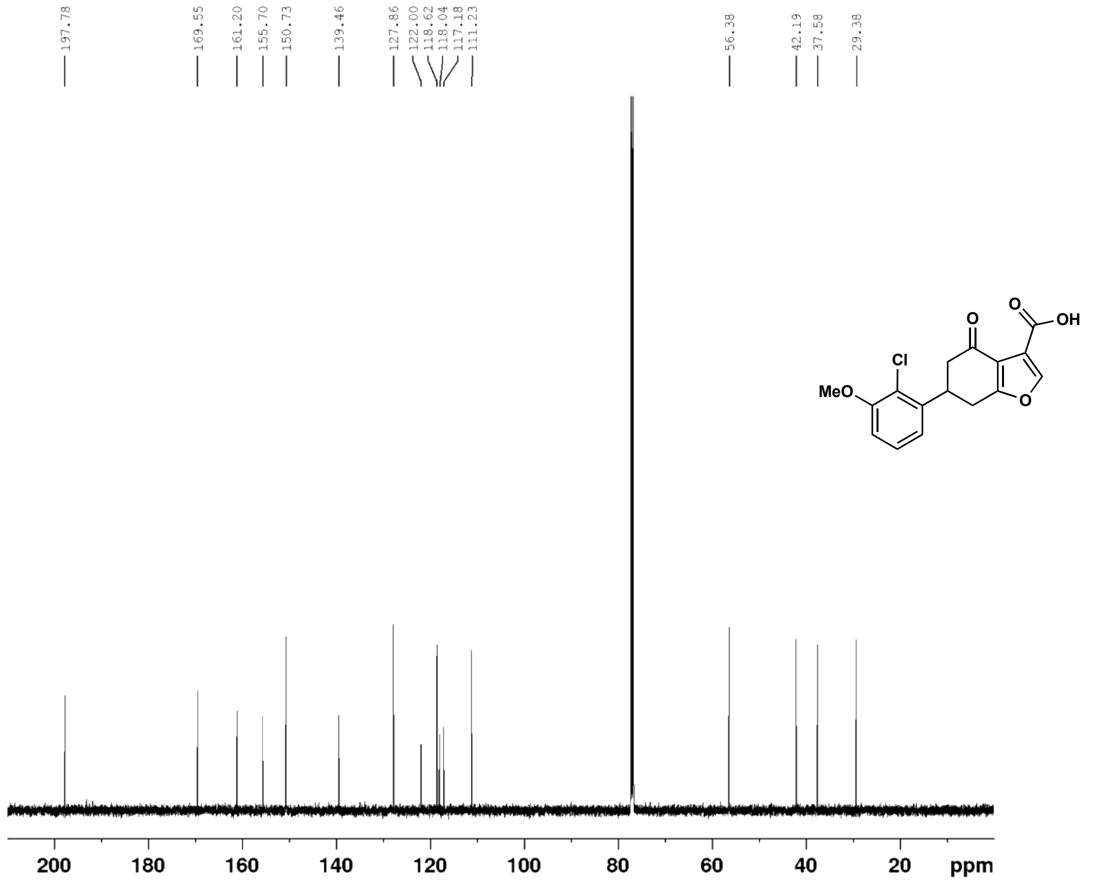

(E)-4-(2,5-dimethoxyphenyl)but-3-en-2-one (**26b**) $^{1}$H NMR (500 MHz, $CDCl_3$)

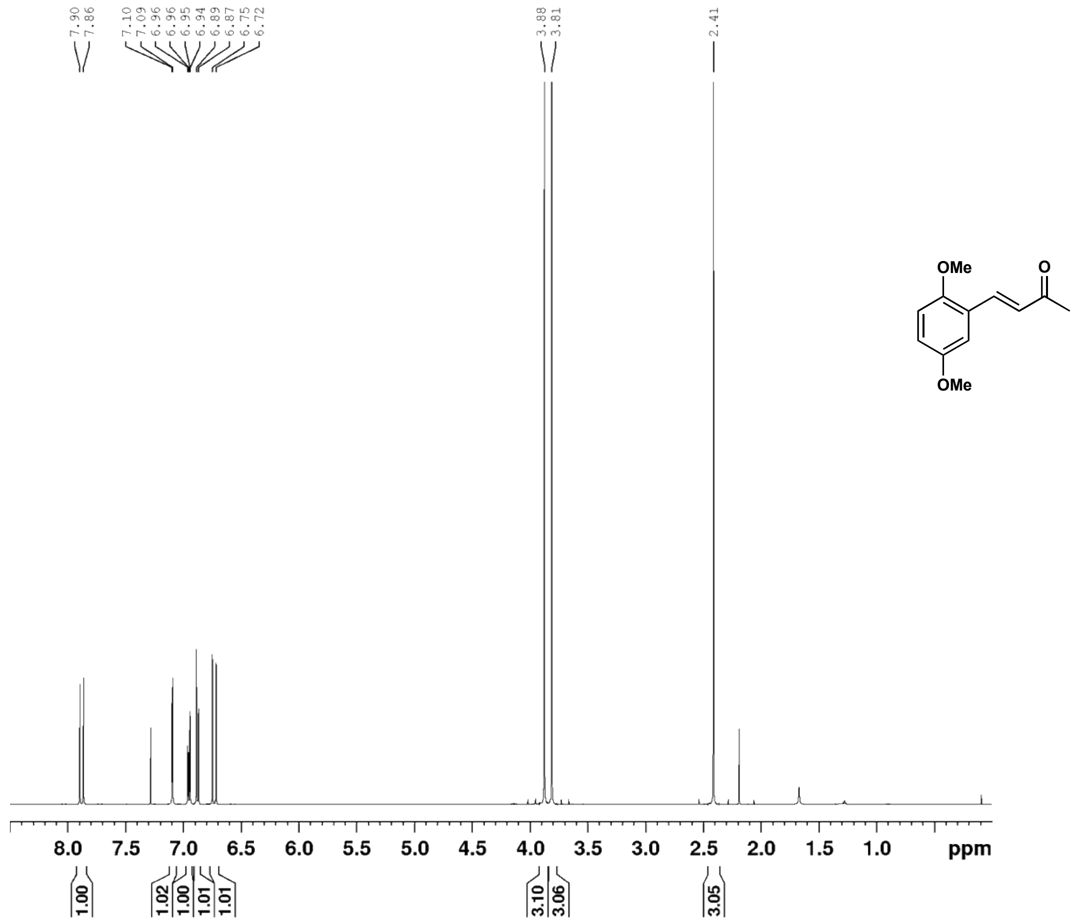

(E)-4-(2,5-dimethoxyphenyl)but-3-en-2-one (**26b**) $^{13}$C NMR (126 MHz, $CDCl_3$)

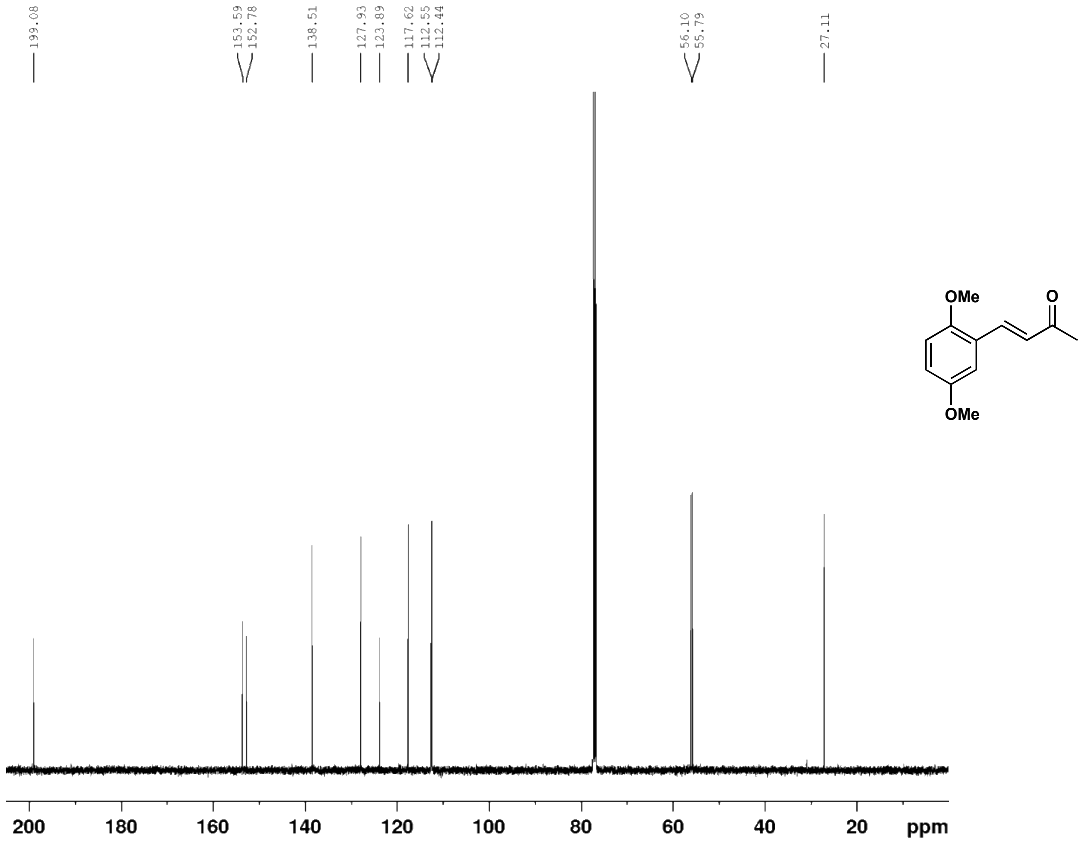

5-(2,5-dimethoxyphenyl)cyclohexane-1,3-dione (**26c**) $^{1}H$ NMR (500 MHz, $CD_3OD$)

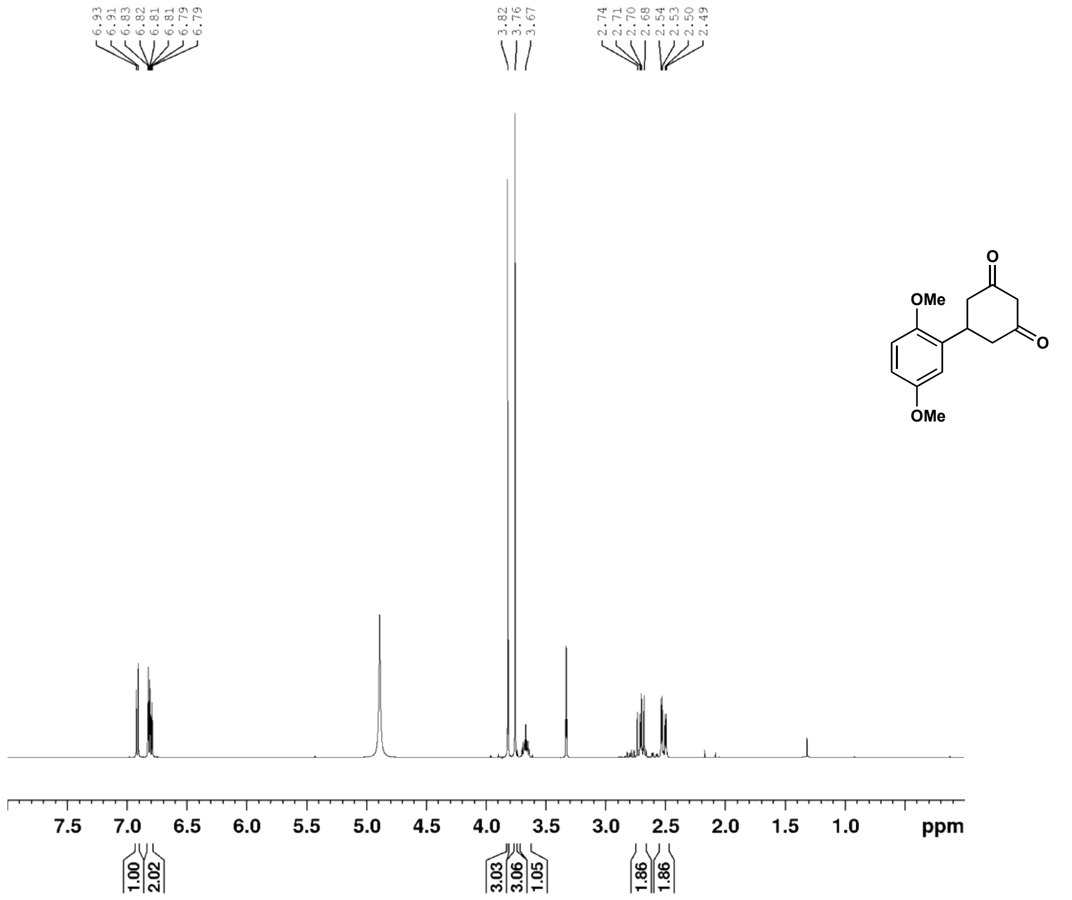

5-(2,5-dimethoxyphenyl)cyclohexane-1,3-dione (**26c**) $^{13}C$ NMR (126 MHz, $CD_3OD$)

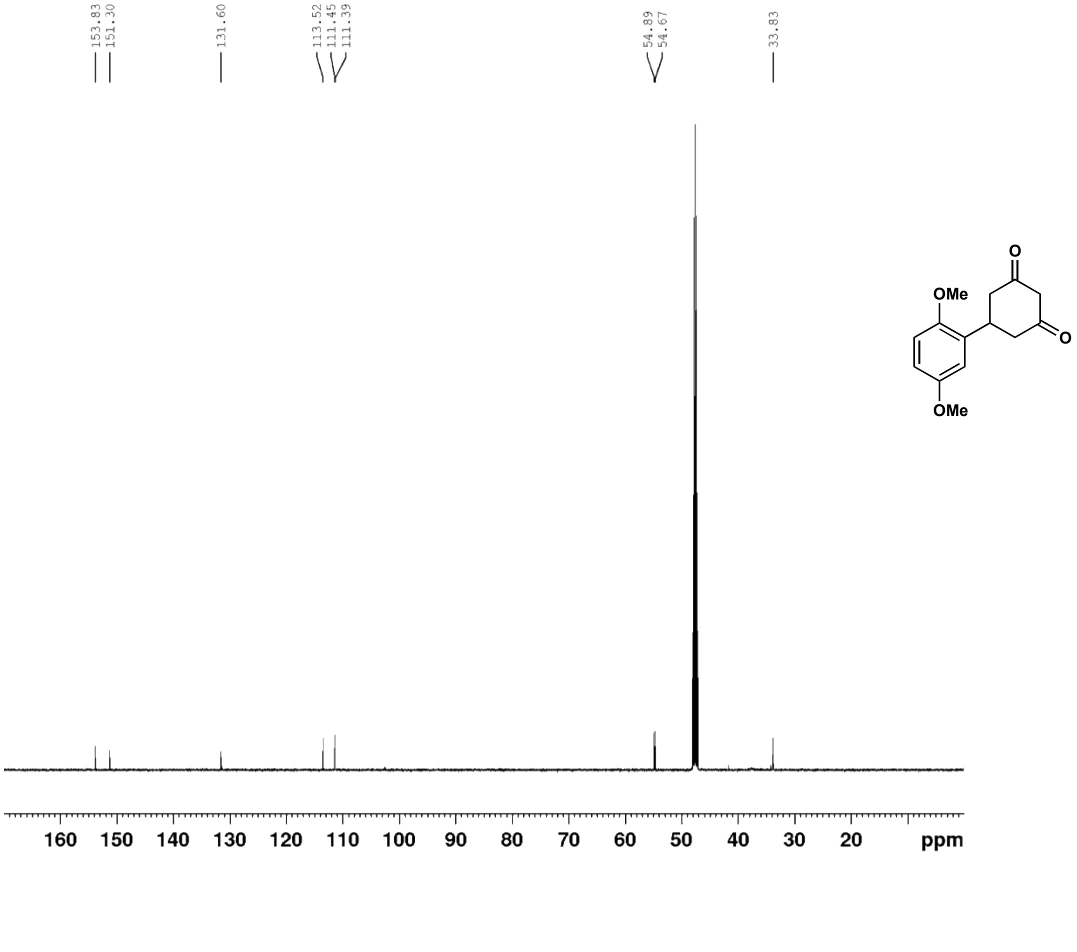

6-(2,5-dimethoxyphenyl)-4-oxo-4,5,6,7-tetrahydrobenzofuran-3-carboxylic acid (**26**) $^{1}$H NMR (500 MHz, $d_6$-DMSO)

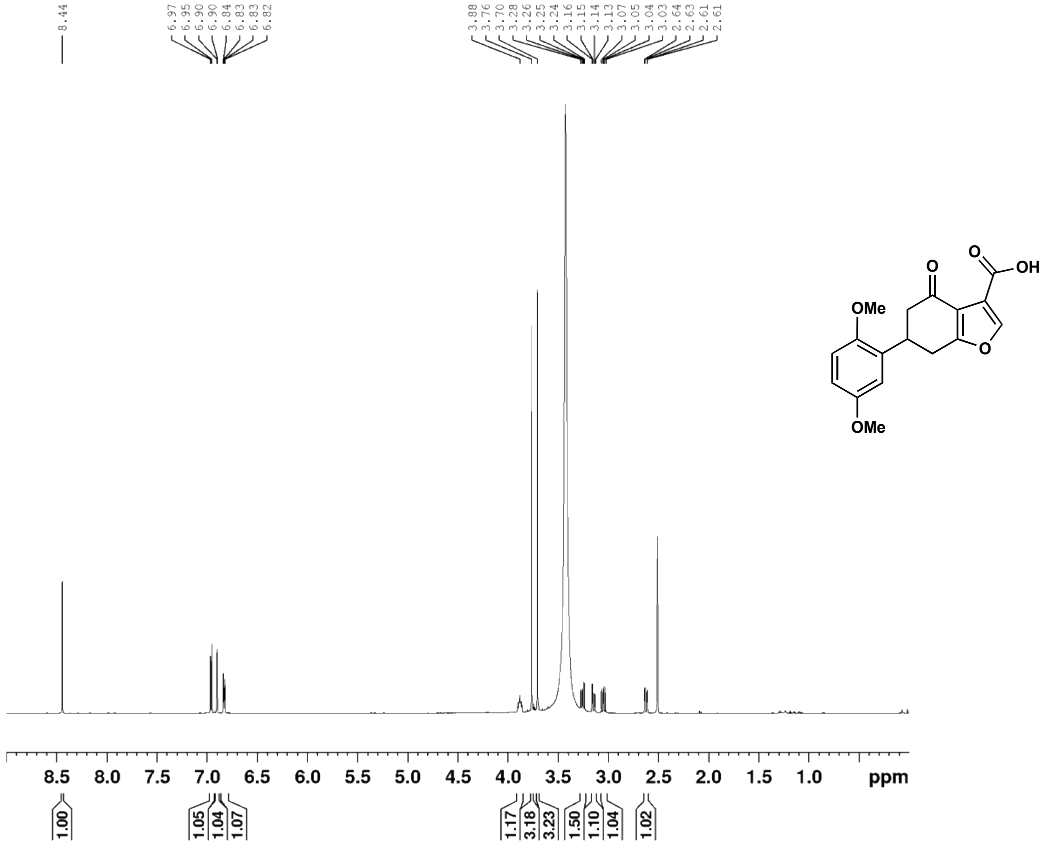

6-(2,5-dimethoxyphenyl)-4-oxo-4,5,6,7-tetrahydrobenzofuran-3-carboxylic acid (**26**) $^{13}$C NMR (126 MHz, $d_6$-DMSO)

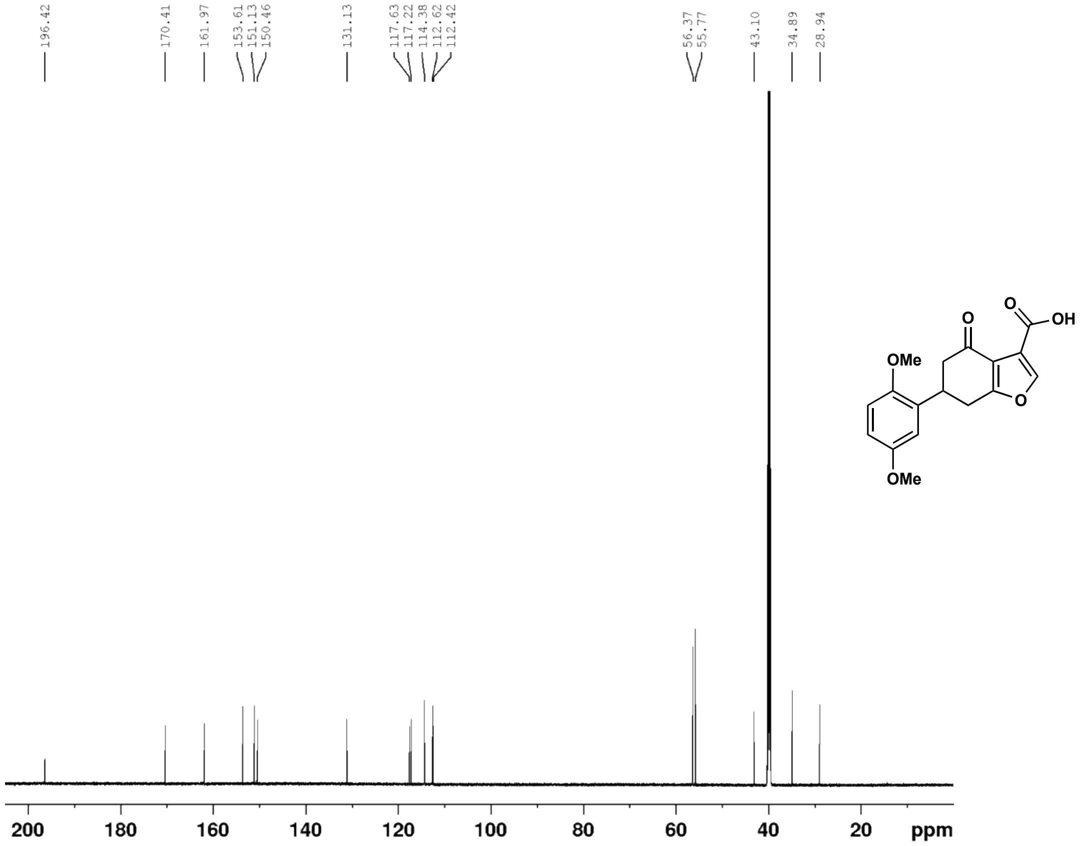

(E)-4-(4-chloro-2-methoxyphenyl)but-3-en-2-one (**27b**) $^{1}$H NMR (500 MHz, $CDCl_3$)

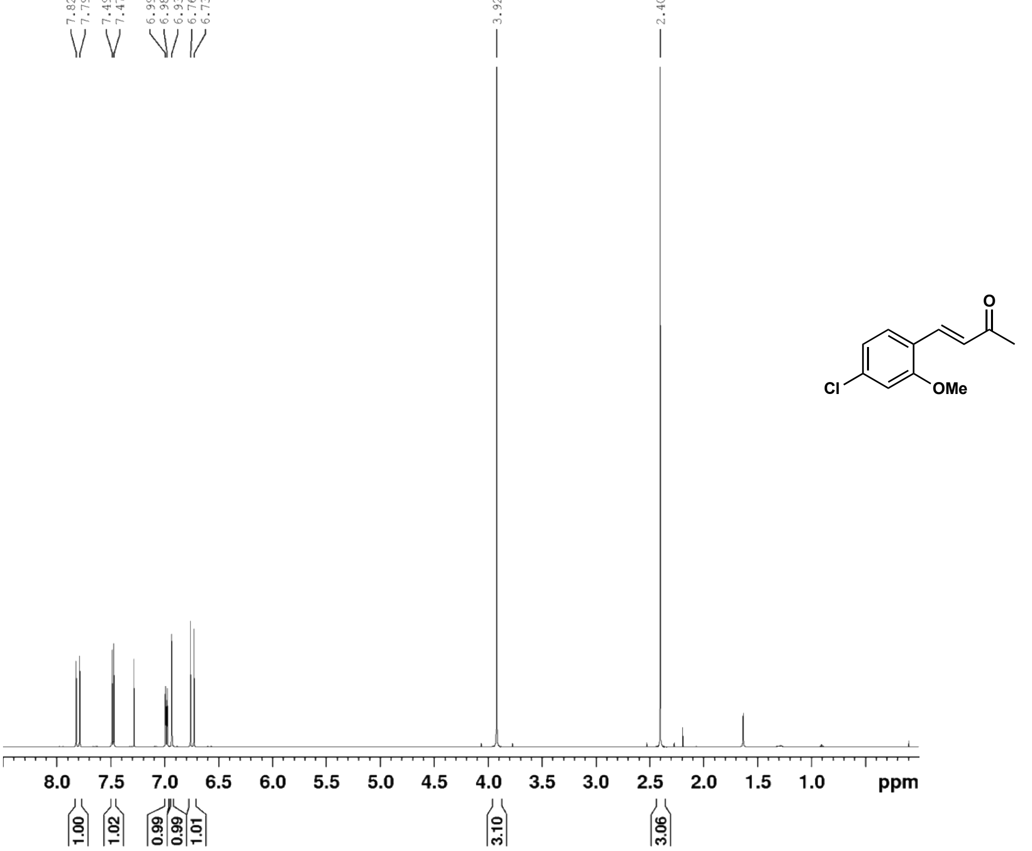

(E)-4-(4-chloro-2-methoxyphenyl)but-3-en-2-one (**27b**) $^{13}$C NMR (126 MHz, $CDCl_3$)

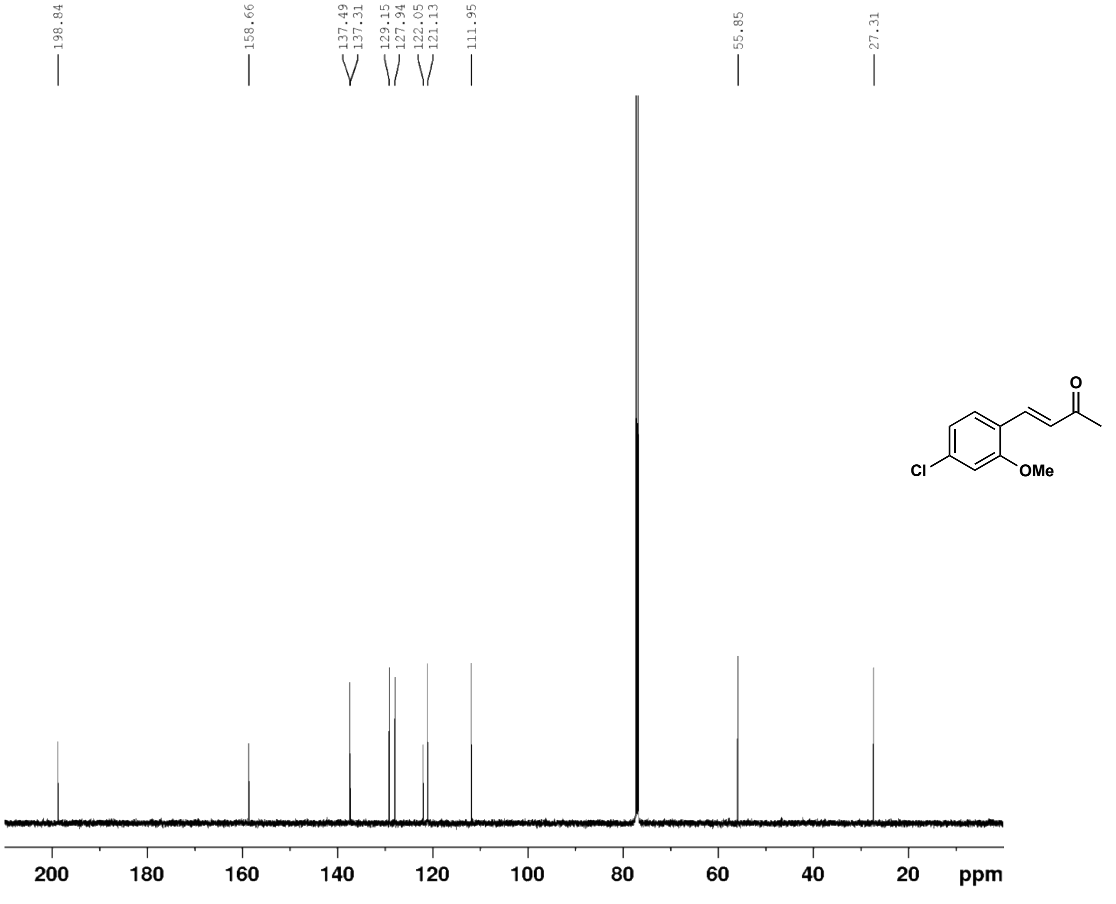

5-(4-chloro-2-methoxyphenyl)cyclohexane-1,3-dione (**27c**) $^{1}$H NMR (500 MHz, $d_6$-DMSO)

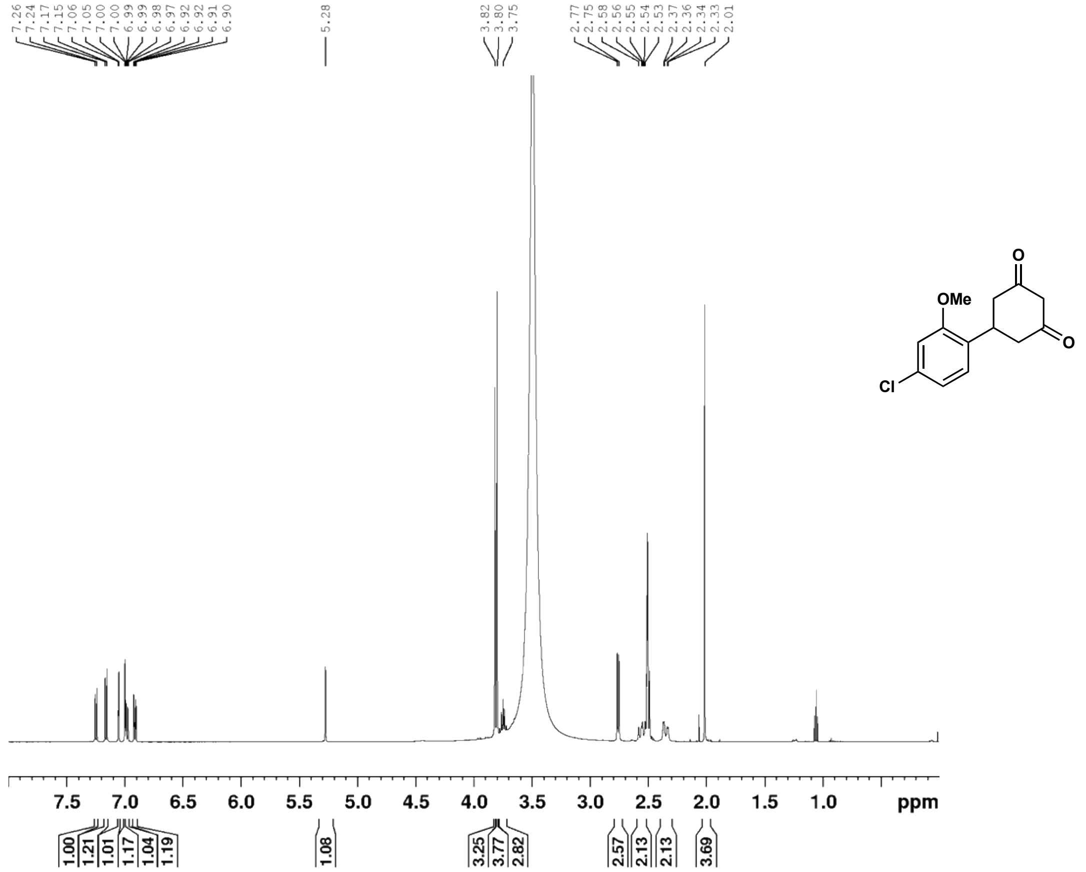

5-(4-chloro-2-methoxyphenyl)cyclohexane-1,3-dione (**27c**) $^{13}$C NMR (126 MHz, $d_6$-DMSO)

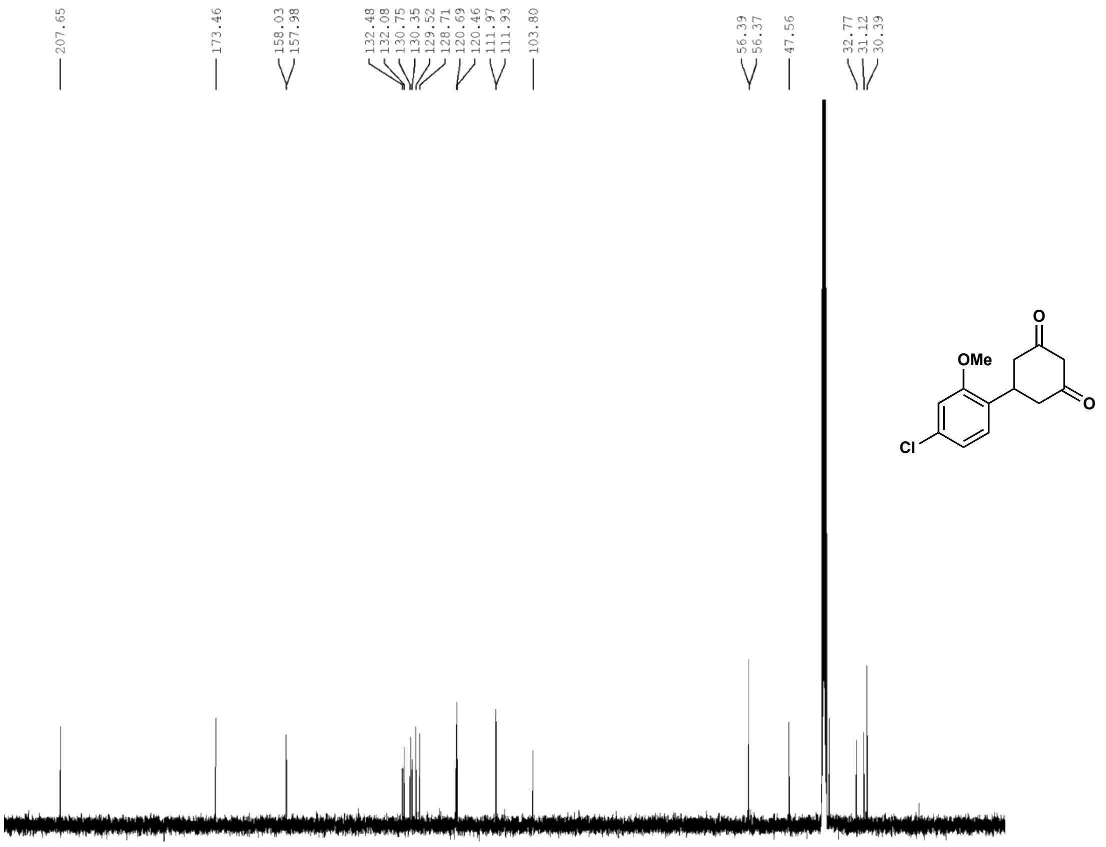

6-(4-chloro-2-methoxyphenyl)-4-oxo-4,5,6,7-tetrahydrobenzofuran-3-carboxylic acid (**27**) $^{1}$H NMR (500 MHz, $CDCl_3$)

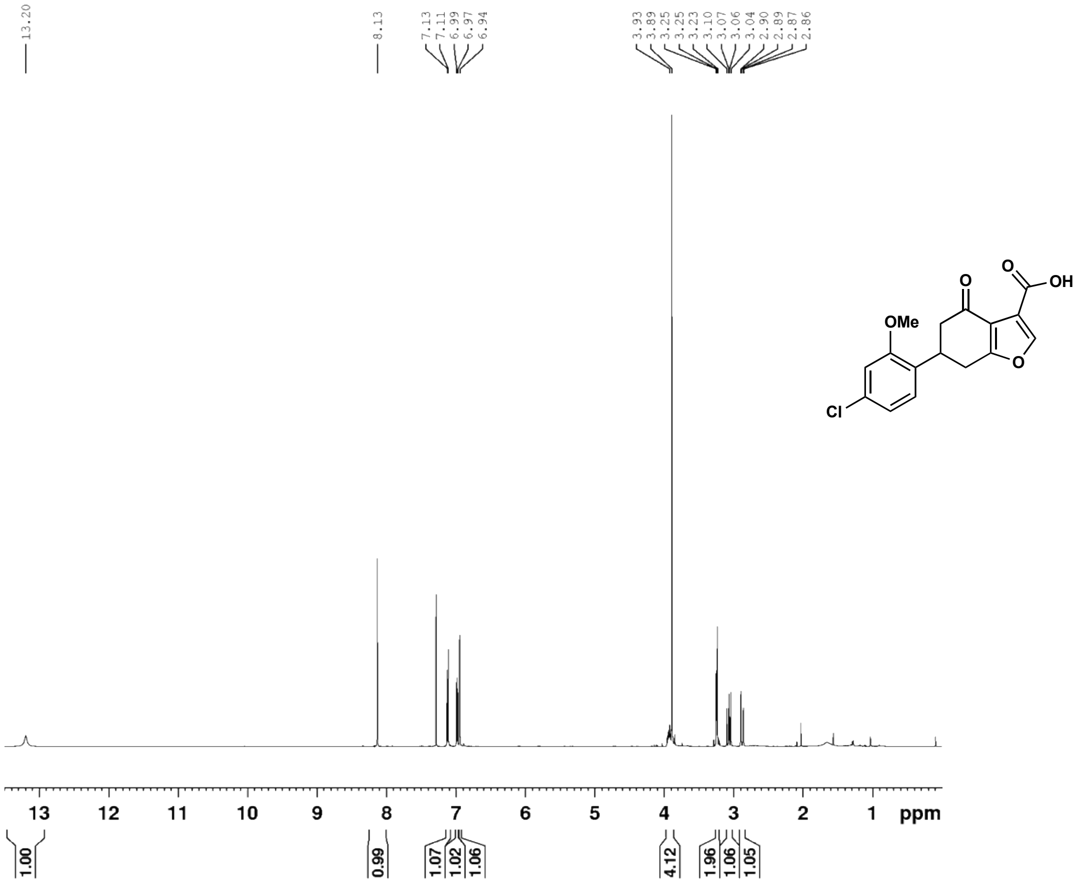

6-(4-chloro-2-methoxyphenyl)-4-oxo-4,5,6,7-tetrahydrobenzofuran-3-carboxylic acid (**27**) $^{13}$C NMR (126 MHz, $CDCl_3$)

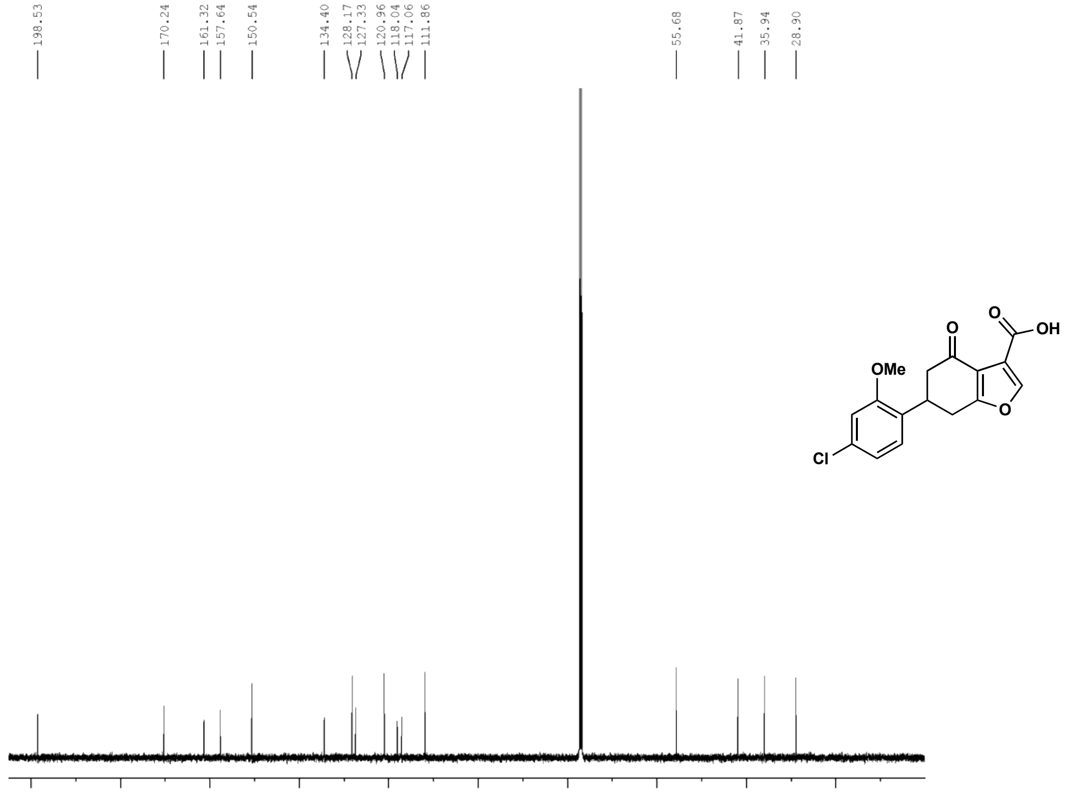

(E)-4-(4-chloro-3-methoxyphenyl)but-3-en-2-one (**28b**) $^{1}H$ NMR (500 MHz, $CDCl_3$)

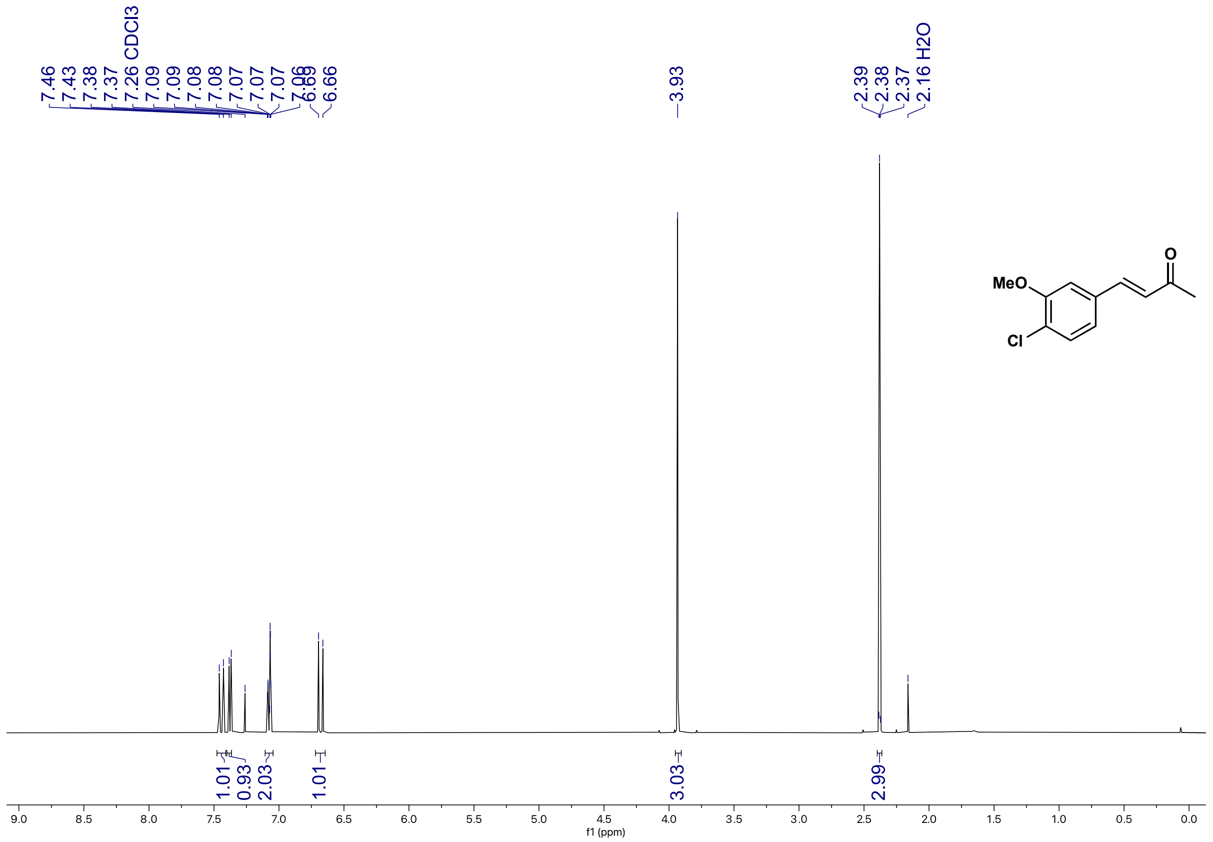

(E)-4-(4-chloro-3-methoxyphenyl)but-3-en-2-one (**28b**) $^{13}C$ NMR (126 MHz, $CDCl_3$)

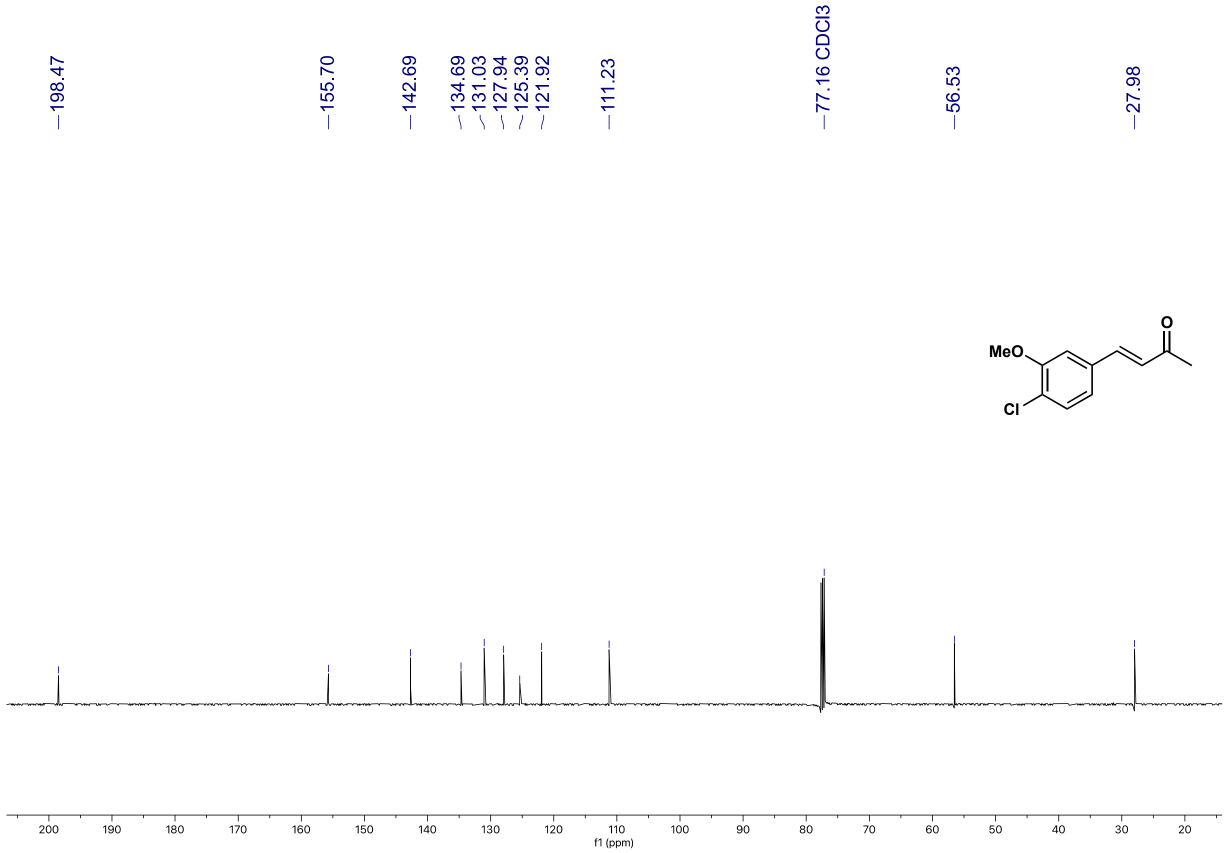

5-(4-chloro-3-methoxyphenyl)cyclohexane-1,3-dione (**28c**) $^{1}$H NMR (500 MHz, $CD_3OD$)

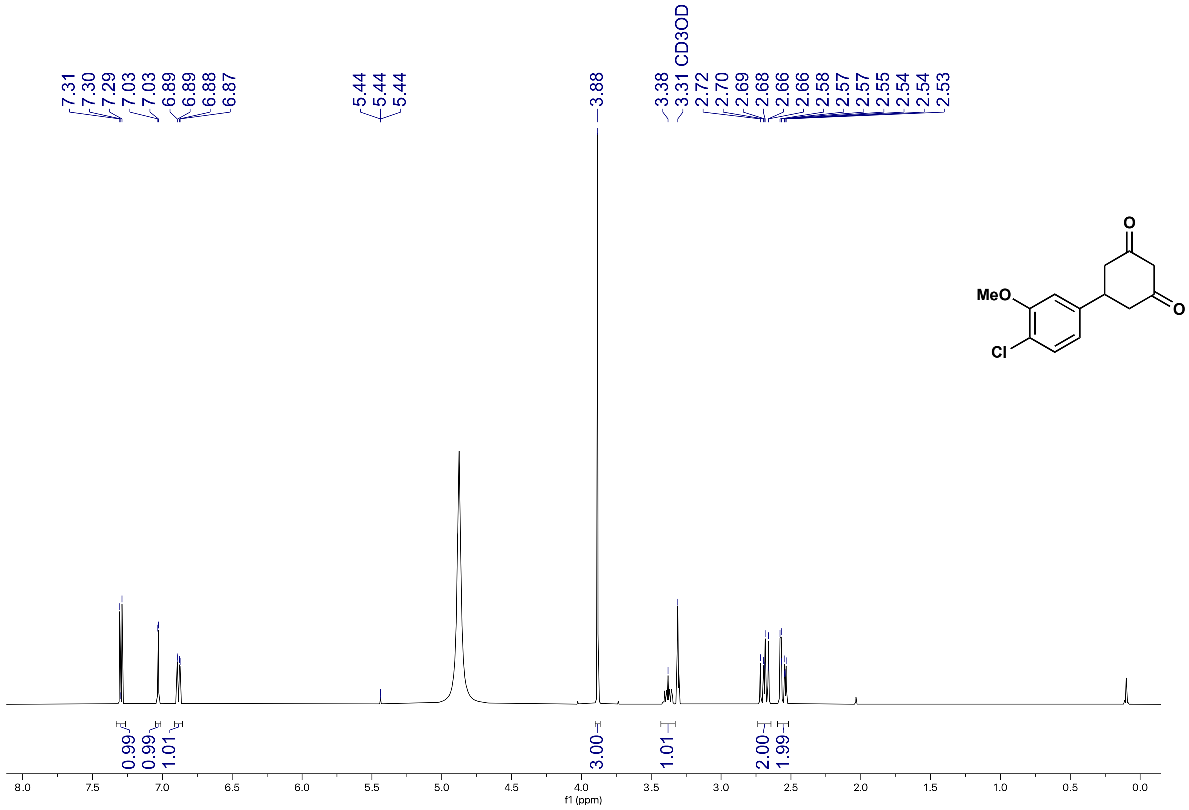

5-(4-chloro-3-methoxyphenyl)cyclohexane-1,3-dione (**28c**) $^{13}$C NMR (126 MHz, $CD_3OD$)

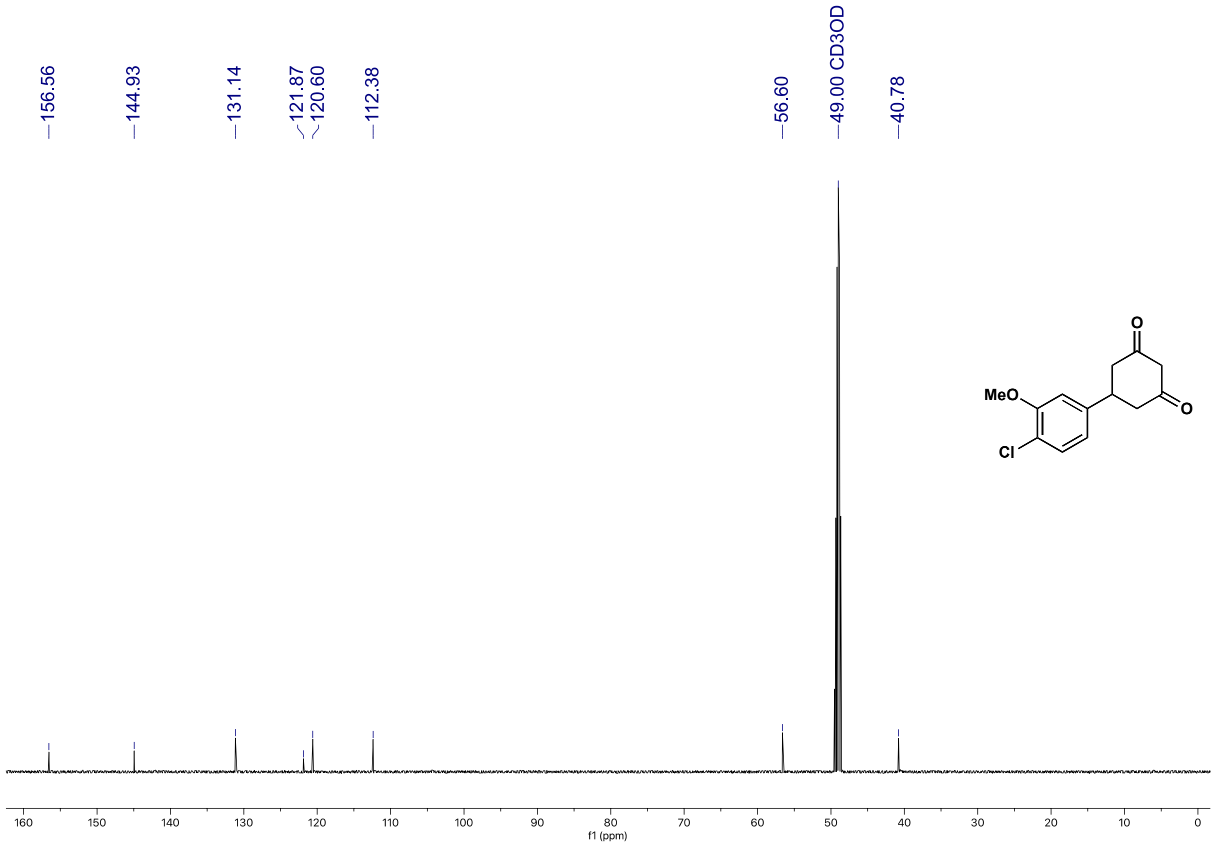

6-(4-chloro-3-methoxyphenyl)-4-oxo-4,5,6,7-tetrahydrobenzofuran-3-carboxylic acid (**28**) $^{1}$H NMR (500 MHz, $d_6$-DMSO)

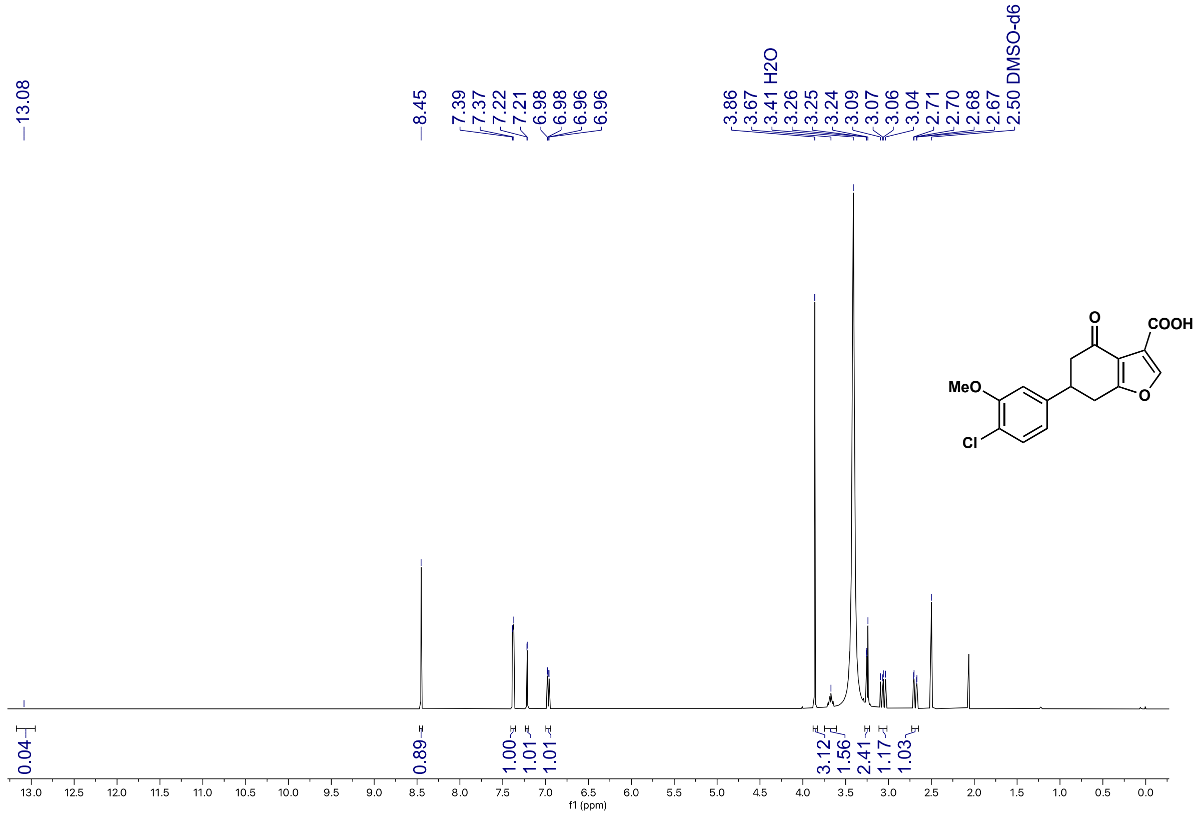

6-(4-chloro-3-methoxyphenyl)-4-oxo-4,5,6,7-tetrahydrobenzofuran-3-carboxylic acid (**28**) $^{13}$C NMR (126 MHz, $d_6$-DMSO)

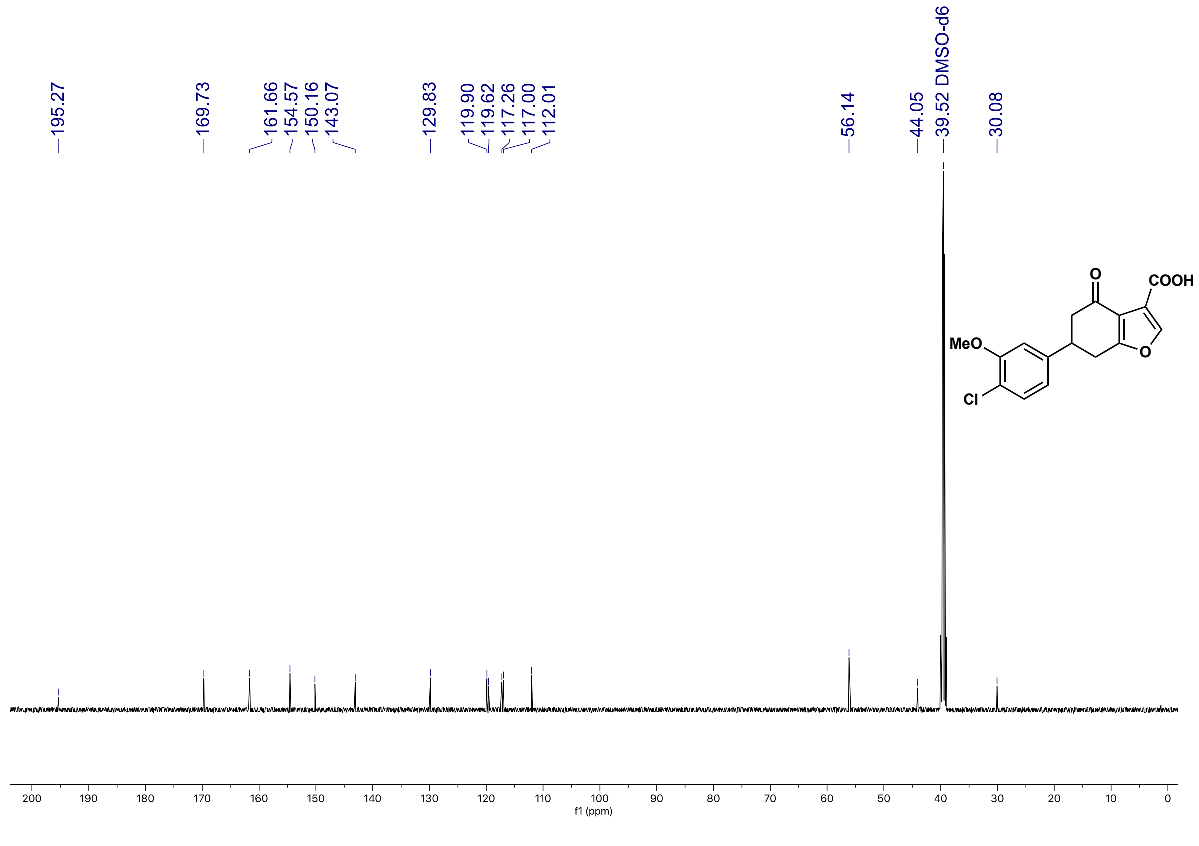

5-(thiophen-2-yl)cyclohexane-1,3-dione (**29c**) $^{1}$H NMR (500 MHz, $CD_3OD$)

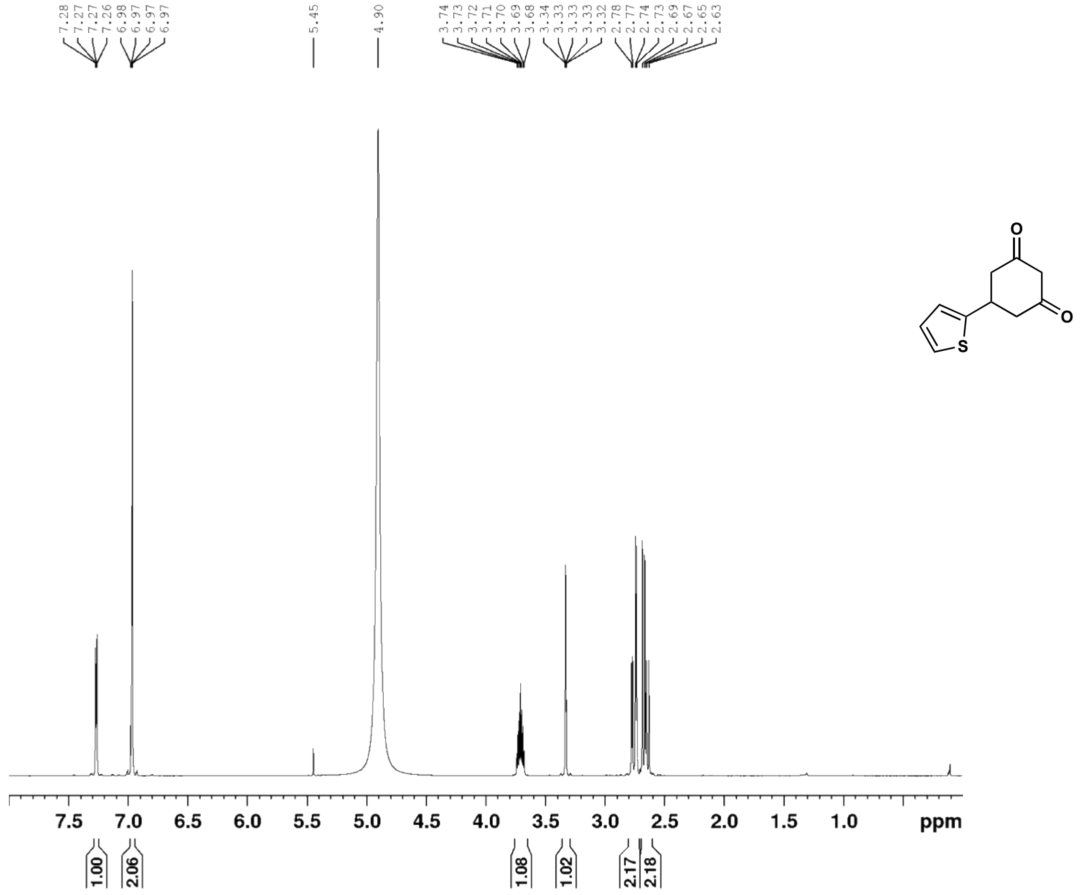

5-(thiophen-2-yl)cyclohexane-1,3-dione (**29c**) $^{13}$C NMR (126 MHz, $CD_3OD$)

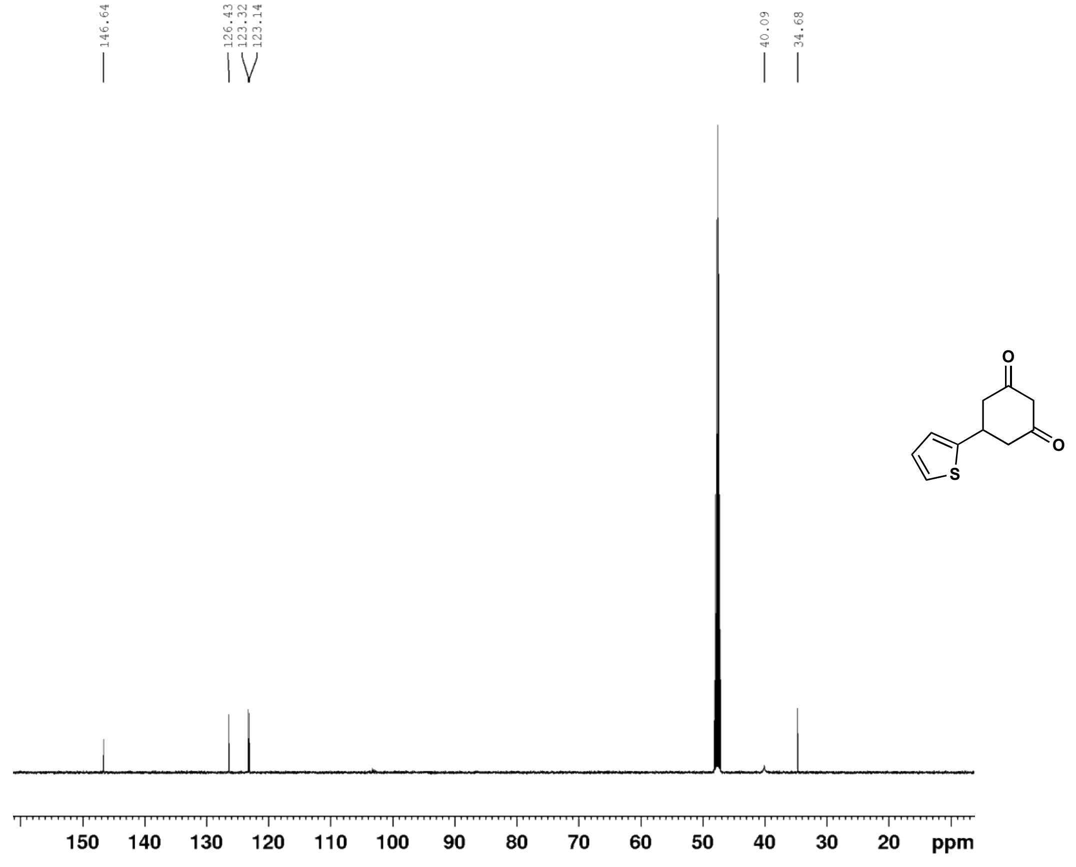

4-oxo-6-(thiophen-2-yl)-4,5,6,7-tetrahydrobenzofuran-3-carboxylic acid (**29**) $^{1}$H NMR (500 MHz, $CDCl_3$)

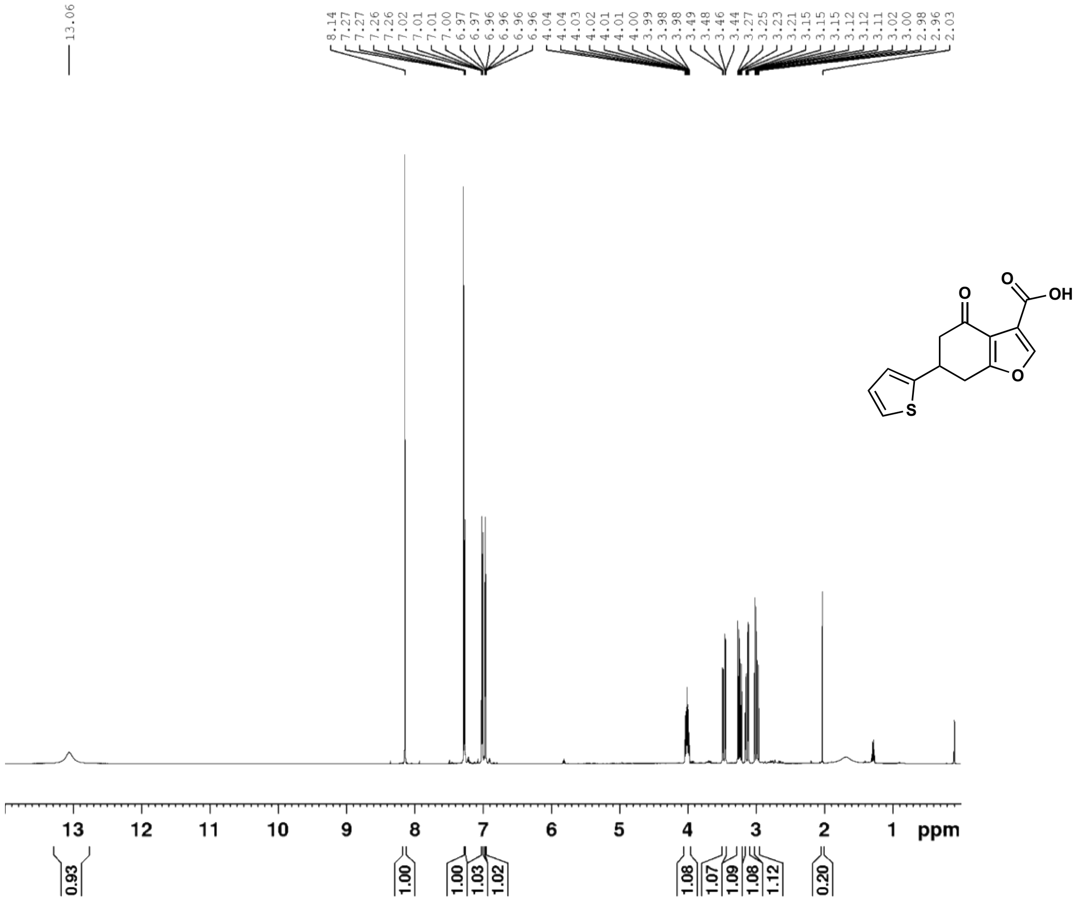

4-oxo-6-(thiophen-2-yl)-4,5,6,7-tetrahydrobenzofuran-3-carboxylic acid (**29**) $^{13}$C NMR (126 MHz, $CDCl_3$)

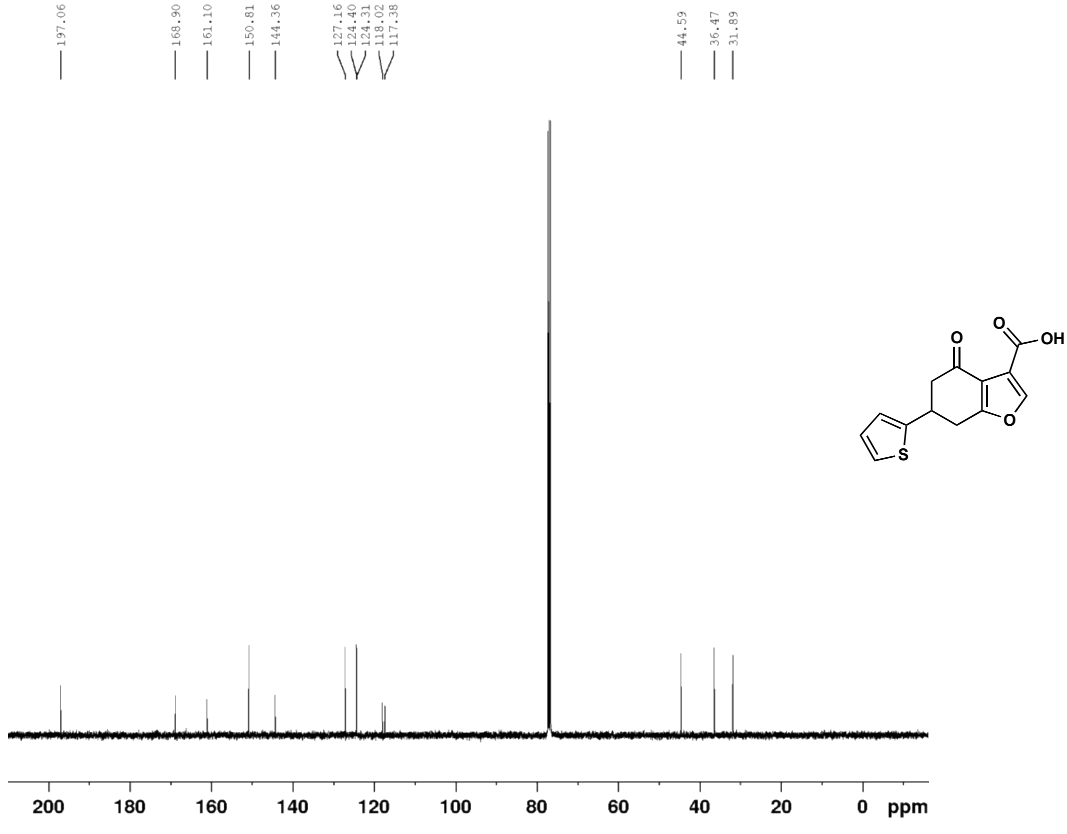

(E)-4-(3-bromothiophen-2-yl)but-3-en-2-one (**30b**) $^{1}$H NMR (500 MHz, $CDCl_3$)

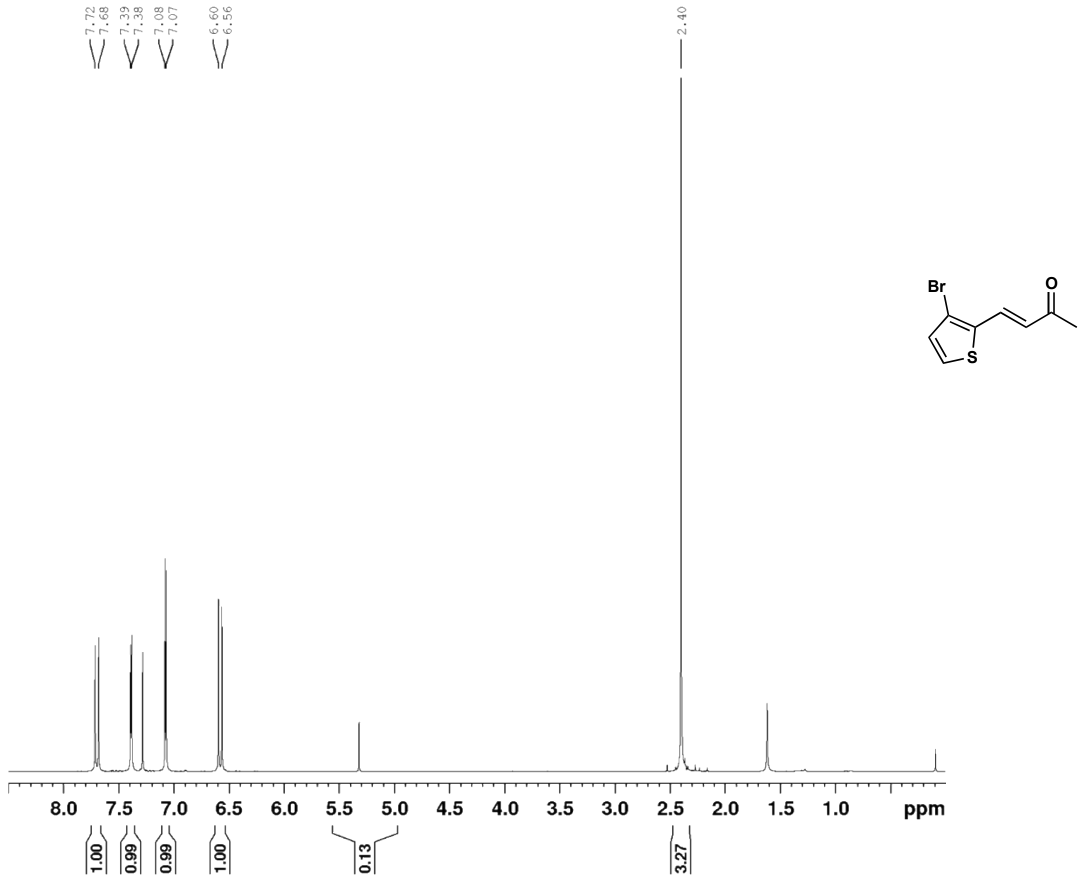

(E)-4-(3-bromothiophen-2-yl)but-3-en-2-one (**30b**) $^{13}$C NMR (126 MHz, $CDCl_3$)

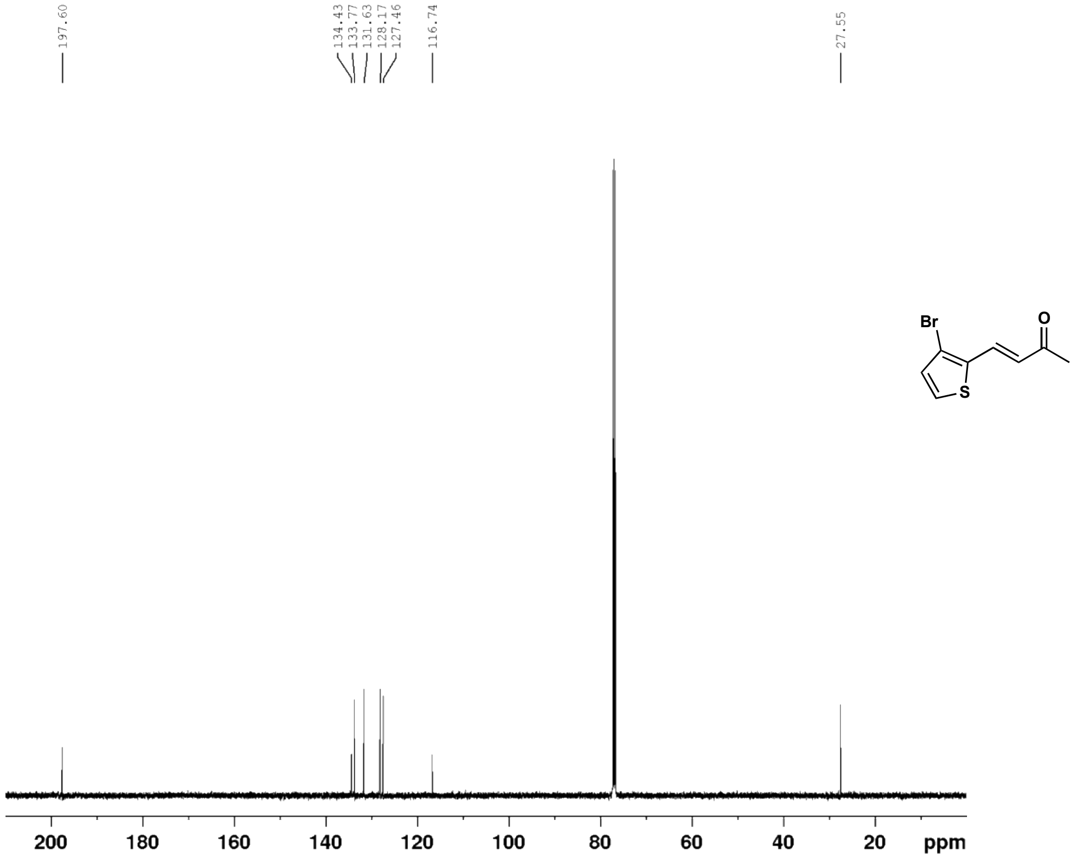

5-(3-bromothiophen-2-yl)cyclohexane-1,3-dione (**30c**) $^{1}H$ NMR (500 MHz, $CD_3OD$)

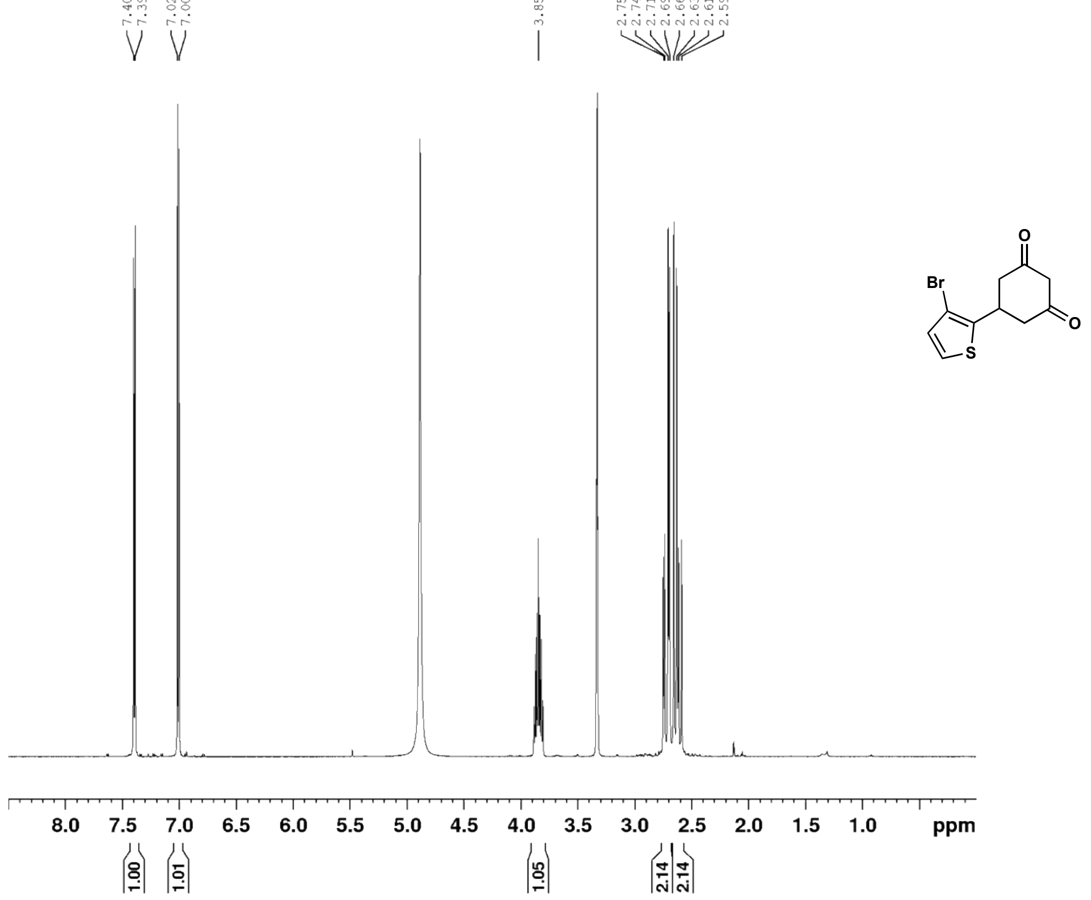

5-(3-bromothiophen-2-yl)cyclohexane-1,3-dione (**30c**) $^{13}C$ NMR (126 MHz, $CD_3OD$)

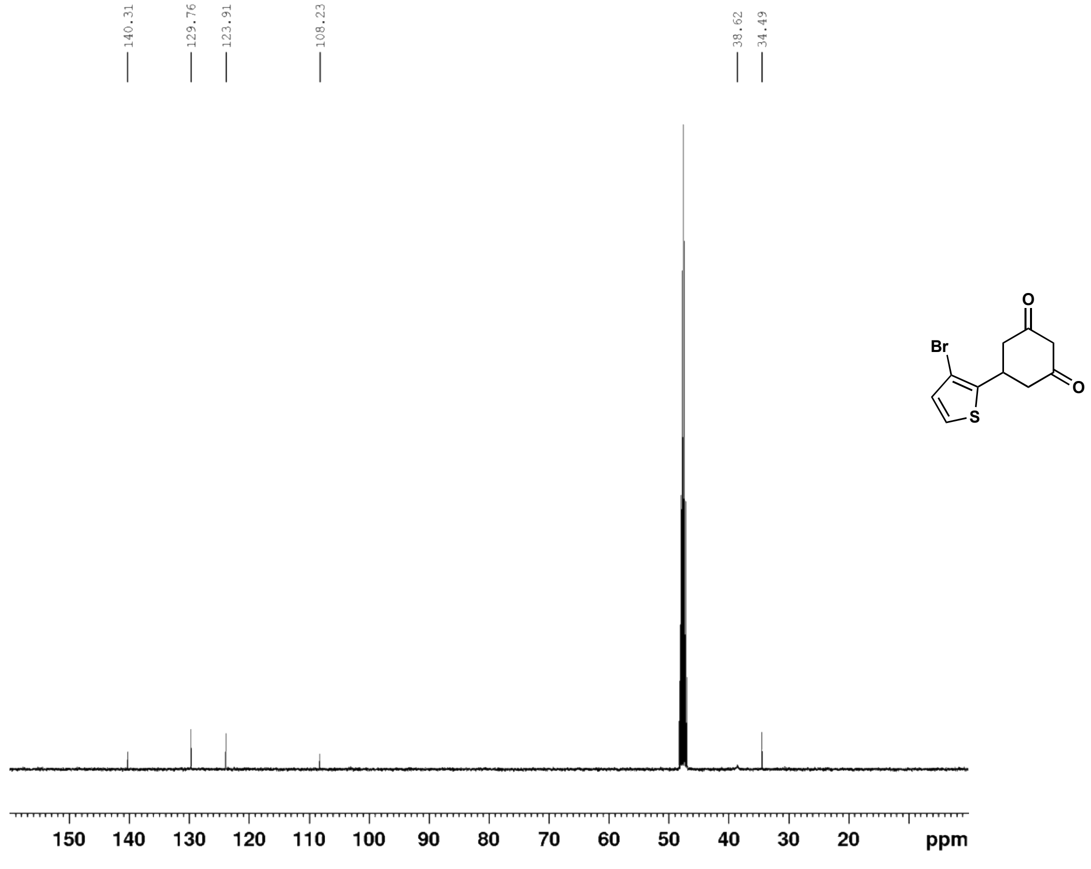

6-(3-bromothiophen-2-yl)-4-oxo-4,5,6,7-tetrahydrobenzofuran-3-carboxylic acid (**30**) $^{1}$H NMR (500 MHz, $CDCl_3$)

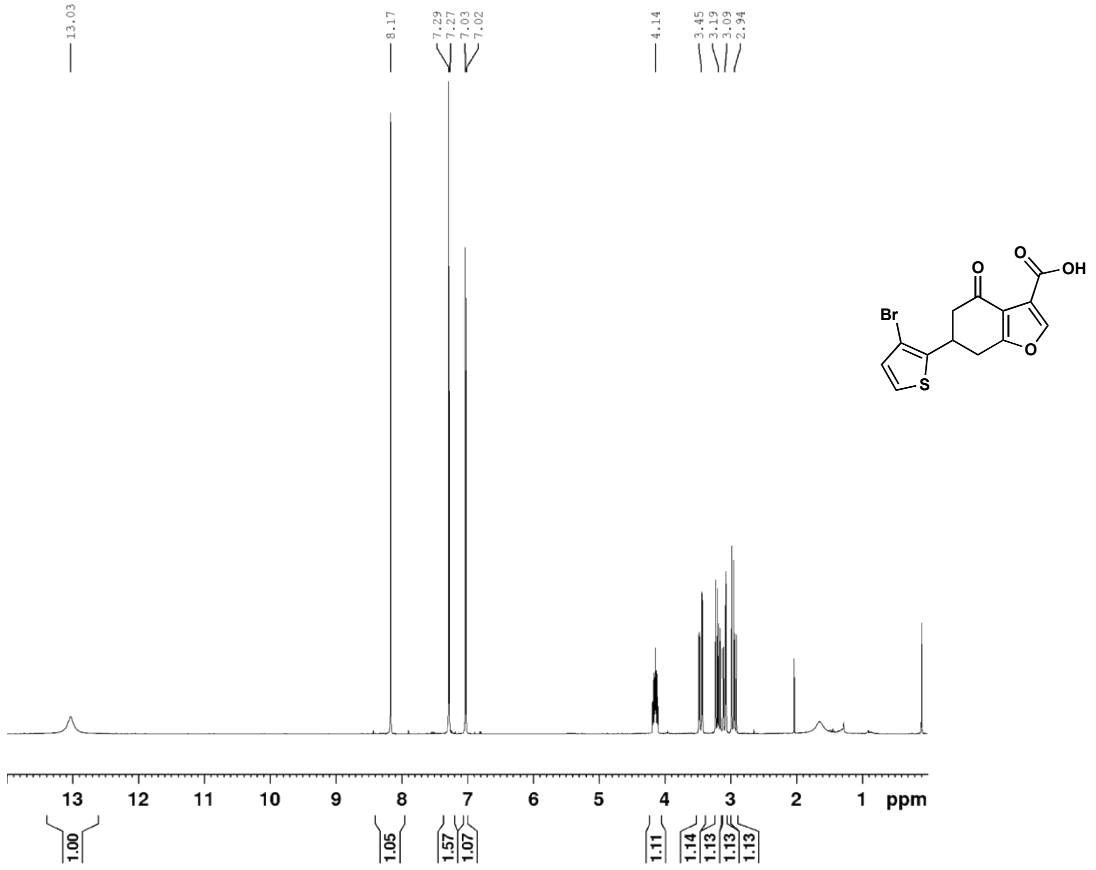

6-(3-bromothiophen-2-yl)-4-oxo-4,5,6,7-tetrahydrobenzofuran-3-carboxylic acid (**30**) $^{13}$C NMR (126 MHz, $CDCl_3$)

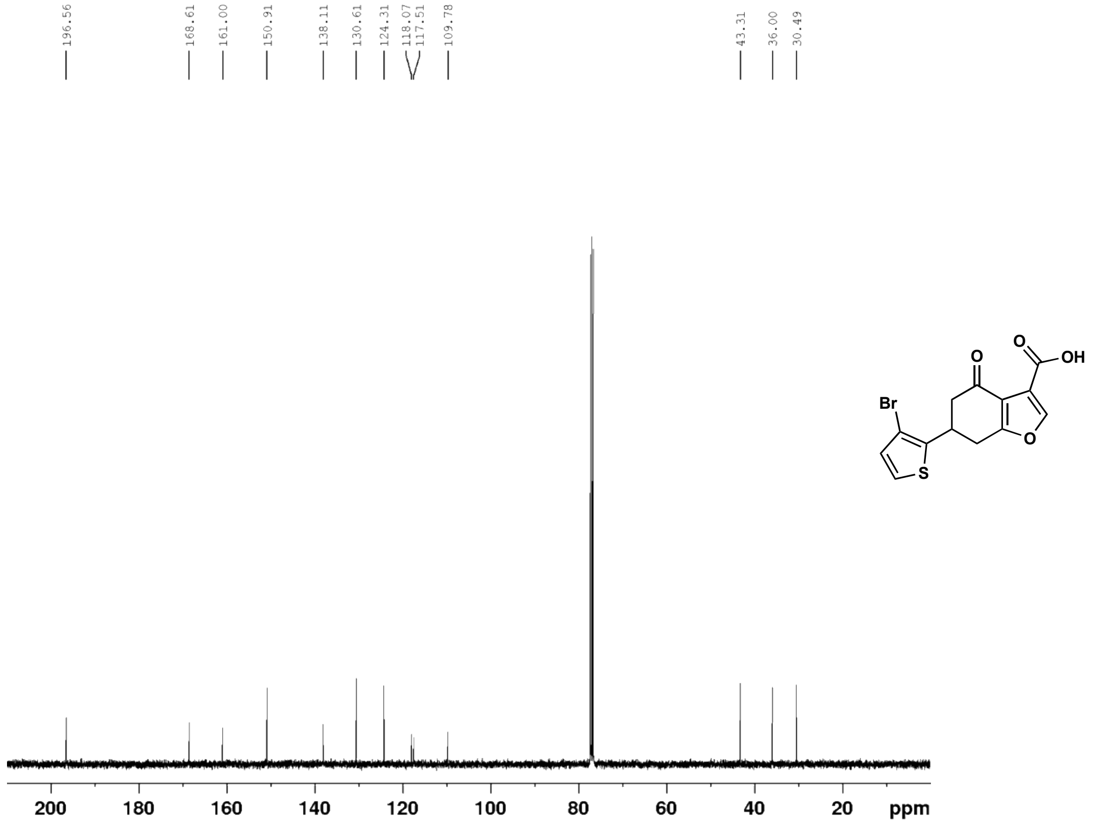

(E)-4-(5-bromothiophen-2-yl)but-3-en-2-one (**31b**) $^1$H NMR (500 MHz, $CDCl_3$)

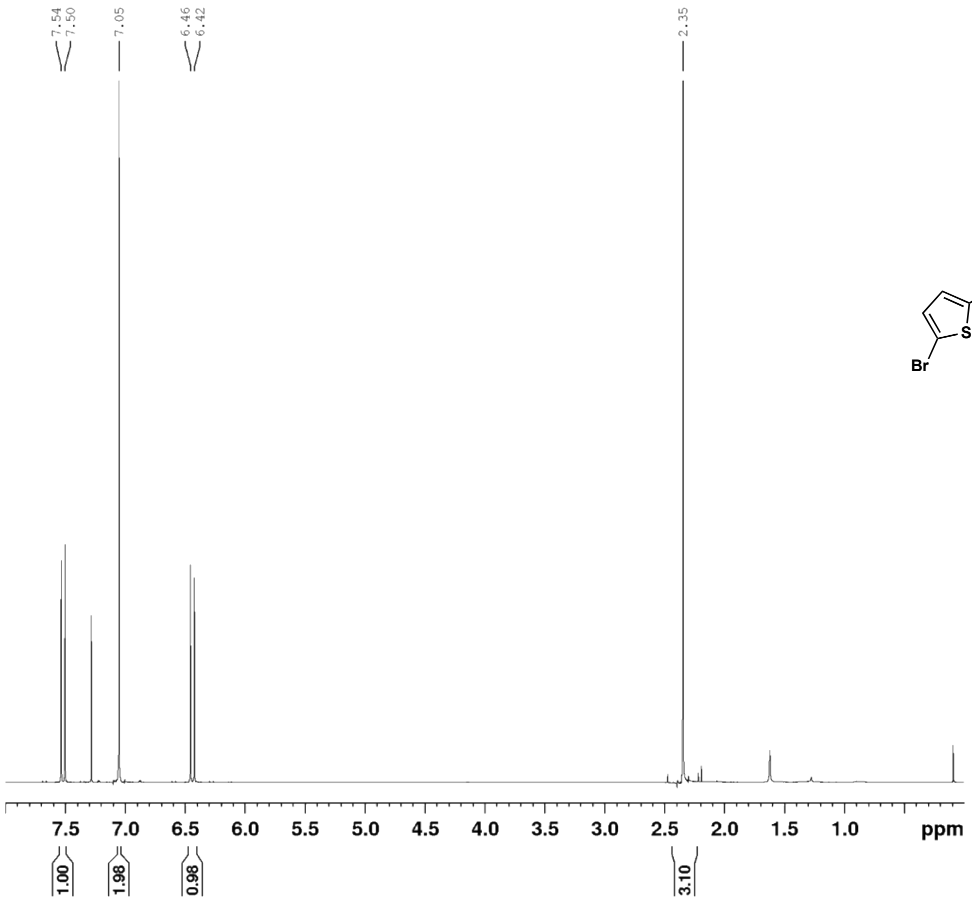

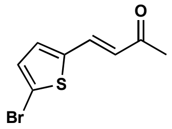

(E)-4-(5-bromothiophen-2-yl)but-3-en-2-one (**31b**) $^{13}$C NMR (126 MHz, $CDCl_3$)

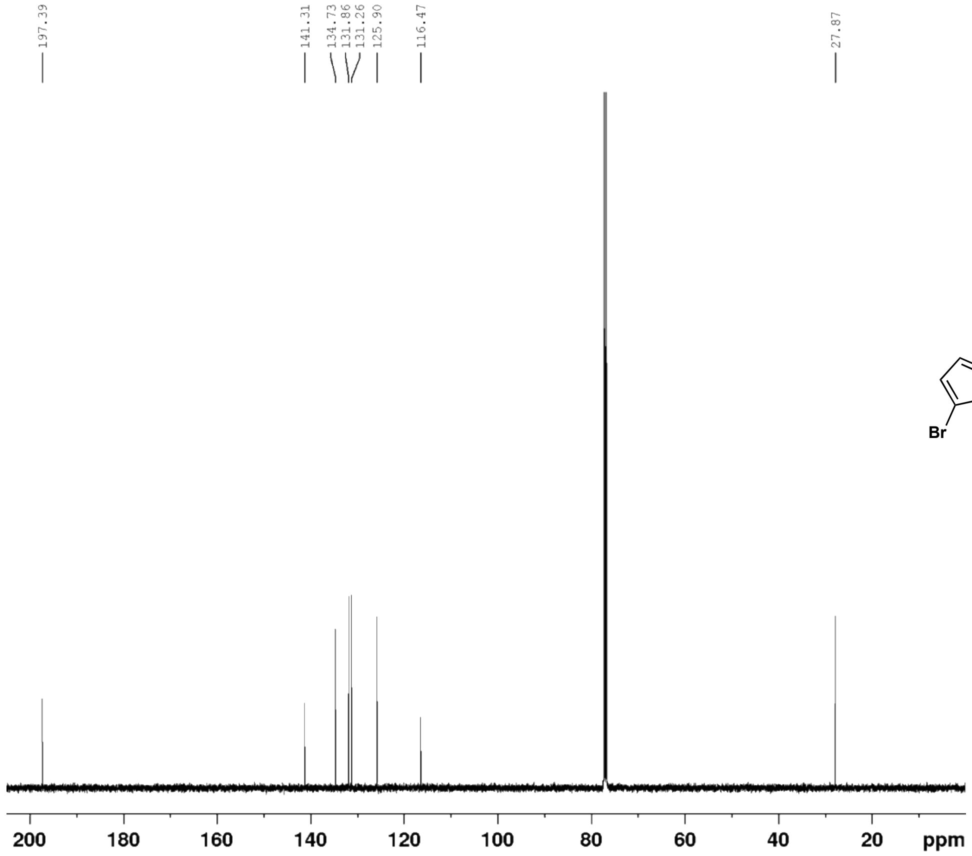

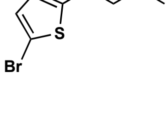

5-(5-bromothiophen-2-yl)cyclohexane-1,3-dione (**31c**) $^{1}$H NMR (500 MHz, $CD_3OD$)

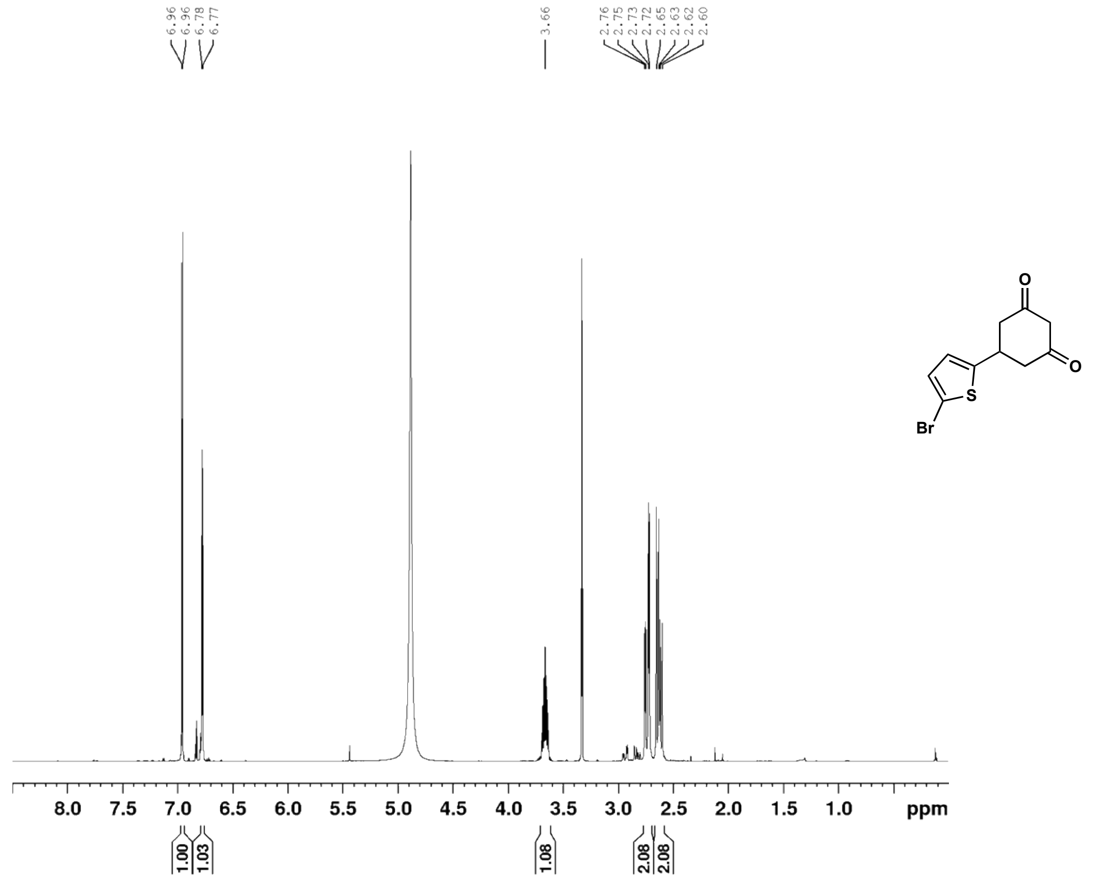

5-(5-bromothiophen-2-yl)cyclohexane-1,3-dione (**31c**) $^{13}$C NMR (126 MHz, $CD_3OD$)

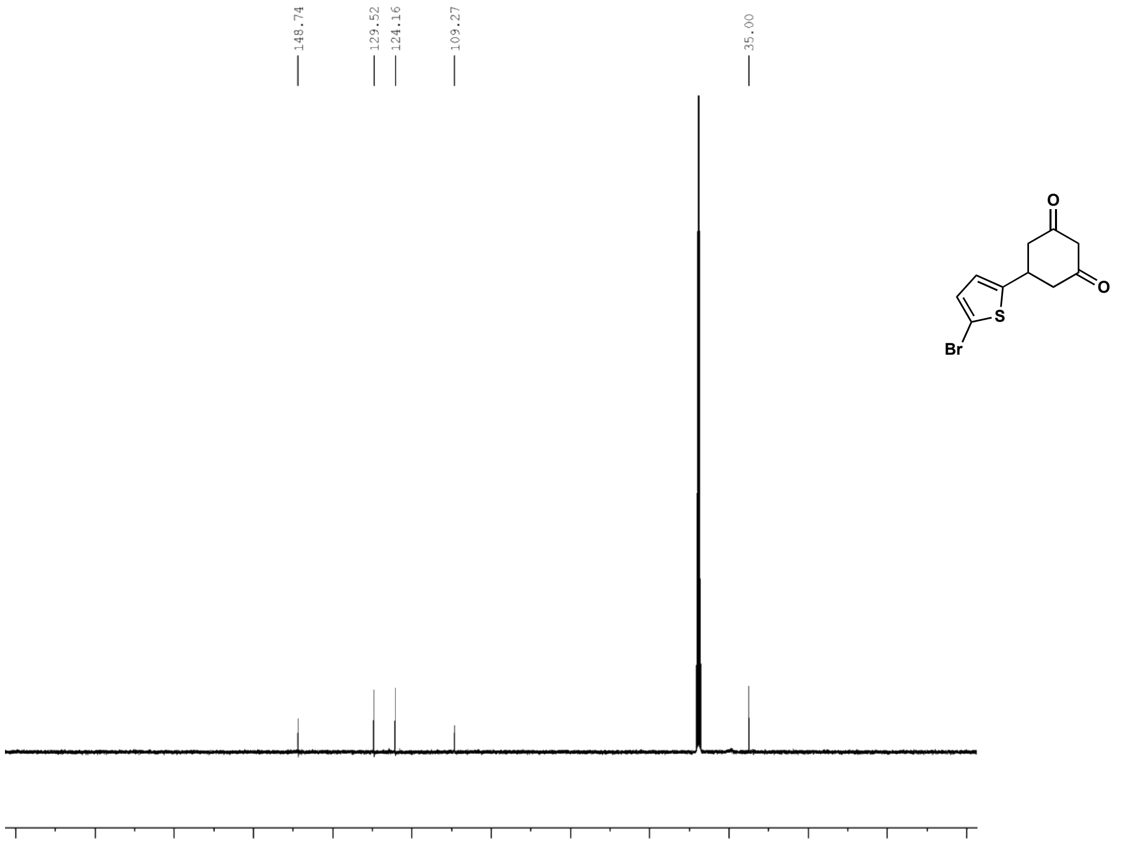

6-(5-bromothiophen-2-yl)-4-oxo-4,5,6,7-tetrahydrobenzofuran-3-carboxylic acid (**31**) $^{1}$H NMR (500 MHz, $d_6$-DMSO)

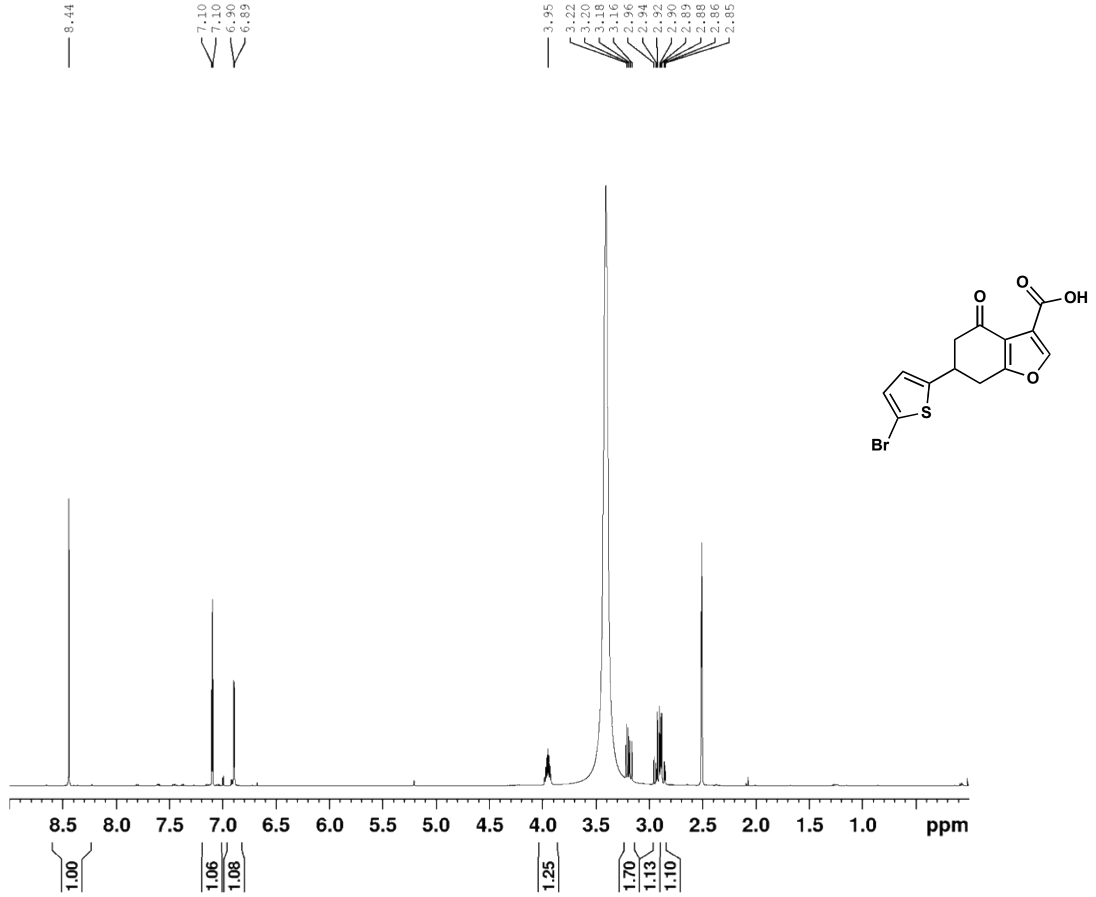

6-(5-bromothiophen-2-yl)-4-oxo-4,5,6,7-tetrahydrobenzofuran-3-carboxylic acid (**31**) $^{13}$C NMR (126 MHz, $d_6$-DMSO)

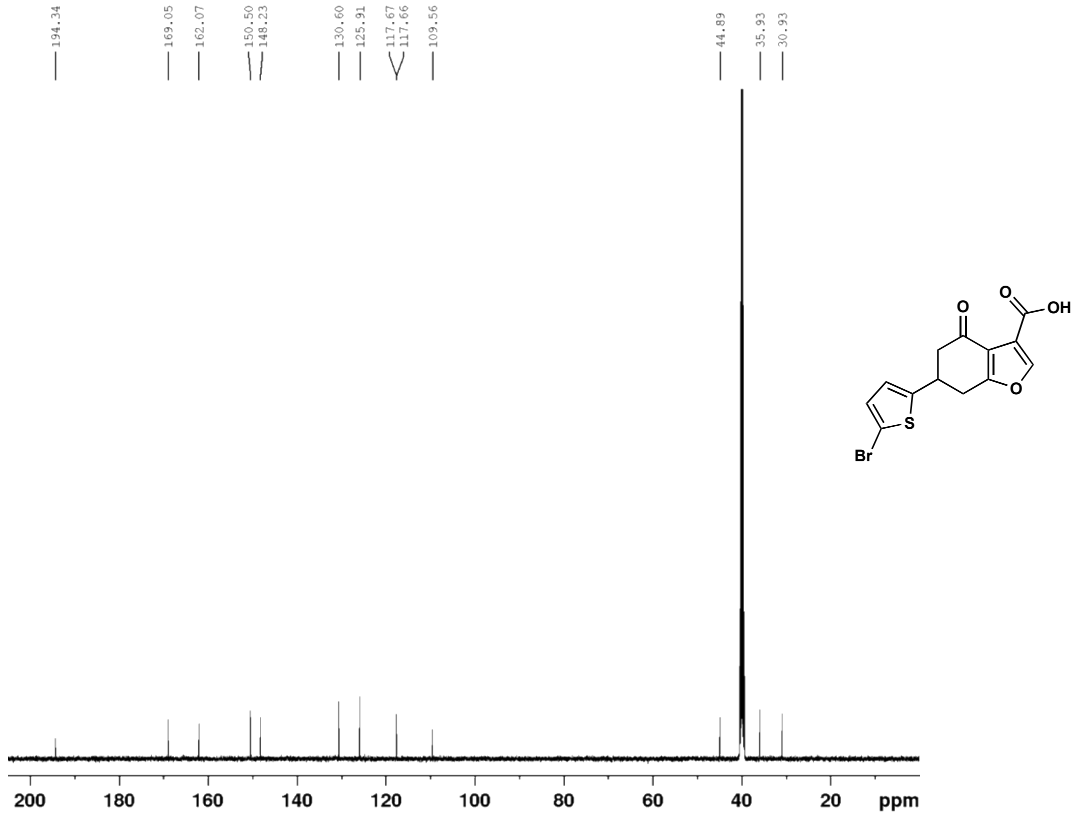

(E)-4-(5-chlorothiophen-2-yl)but-3-en-2-one (**32b**) $^{1}$H NMR (500 MHz, $CDCl_3$)

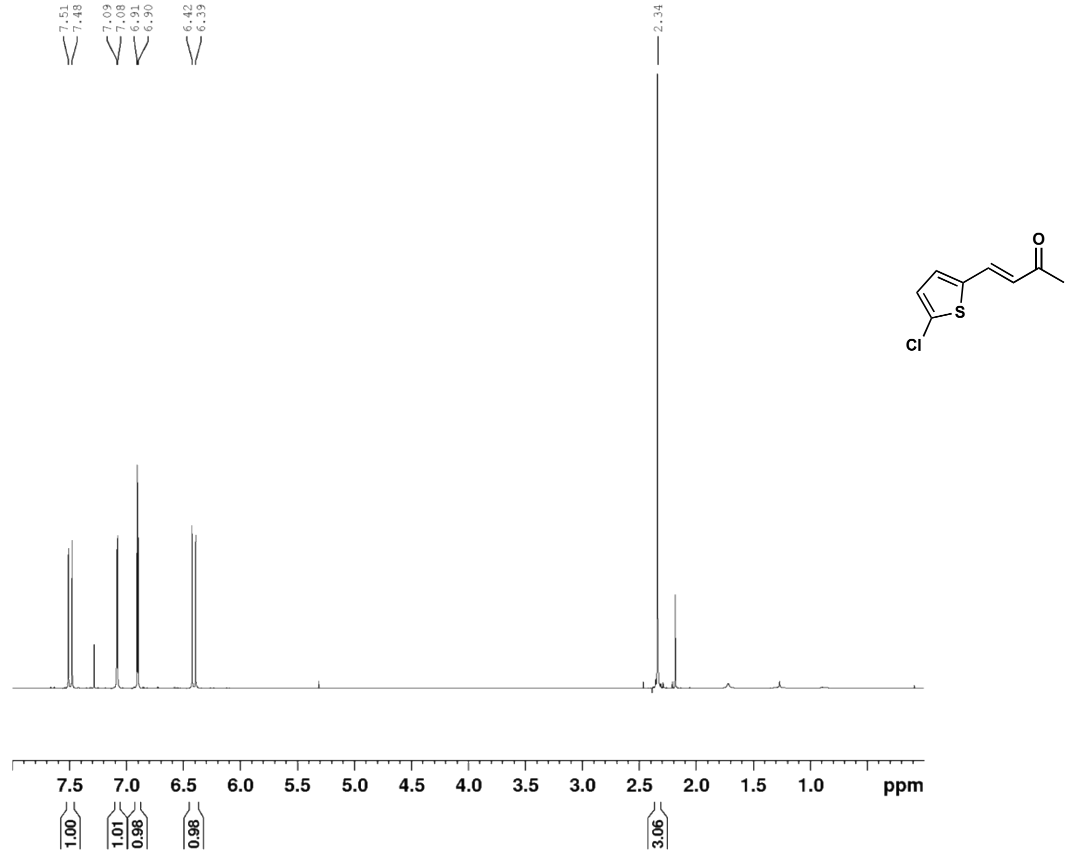

(E)-4-(5-chlorothiophen-2-yl)but-3-en-2-one (**32b**) $^{13}$C NMR (126 MHz, $CDCl_3$)

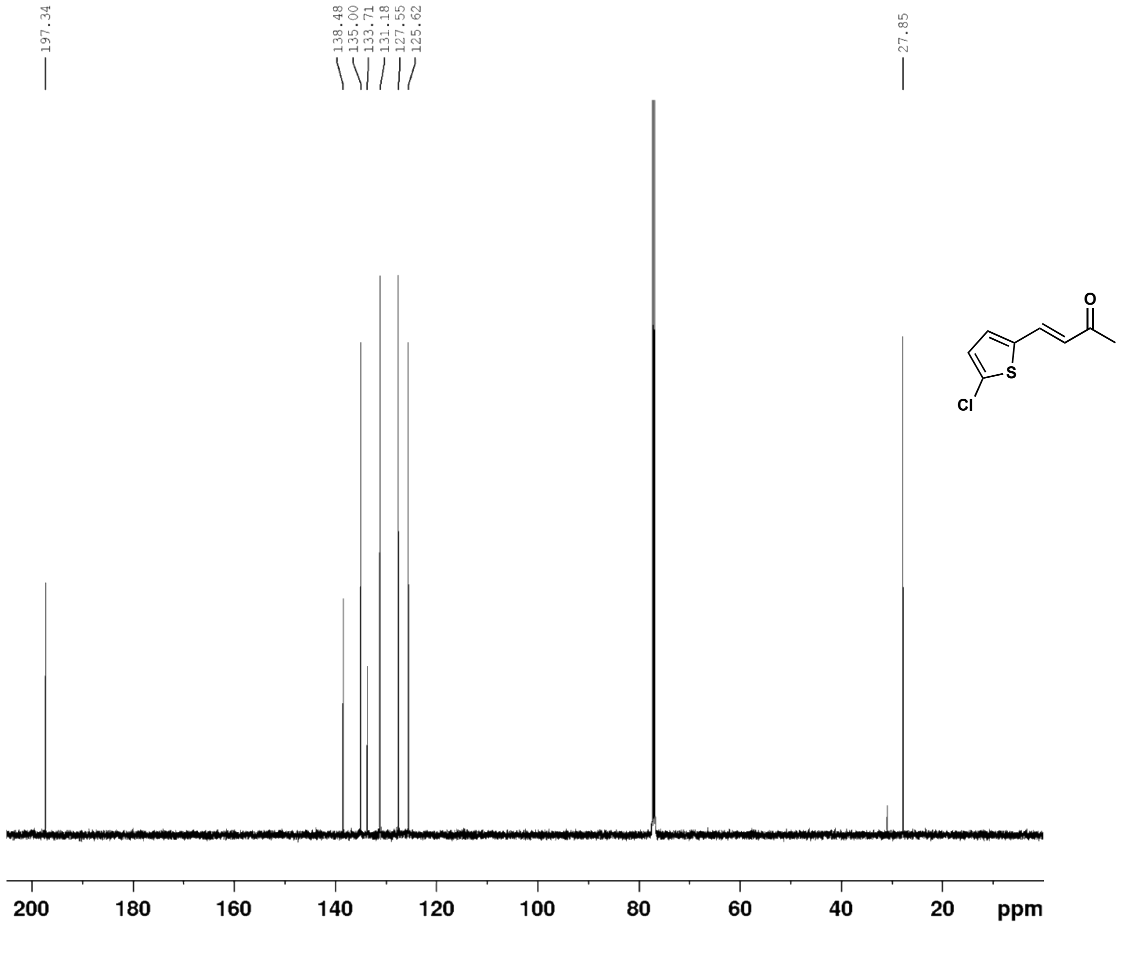

5-(5-chlorothiophen-2-yl)cyclohexane-1,3-dione (**32c**) $^{1}$H NMR (500 MHz, $CD_3OD$)

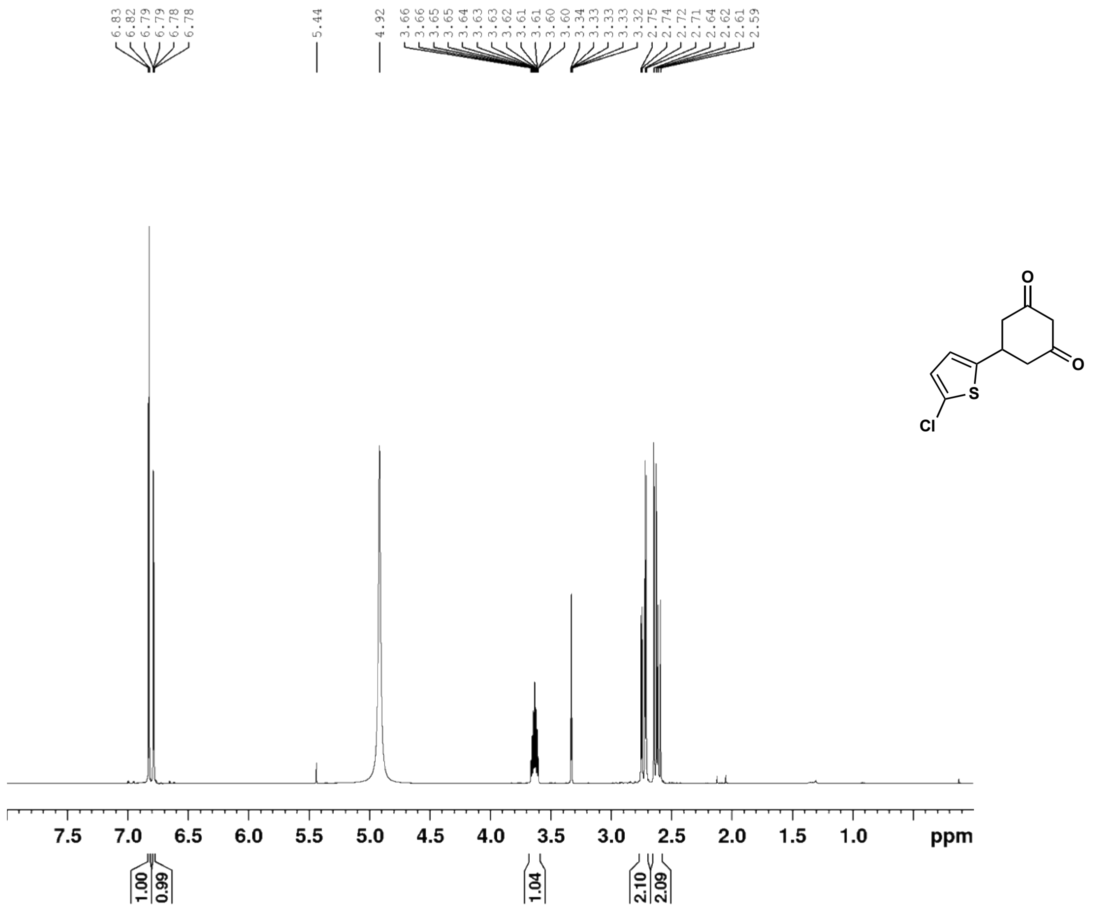

5-(5-chlorothiophen-2-yl)cyclohexane-1,3-dione (**32c**) $^{13}$C NMR (126 MHz, $CD_3OD$)

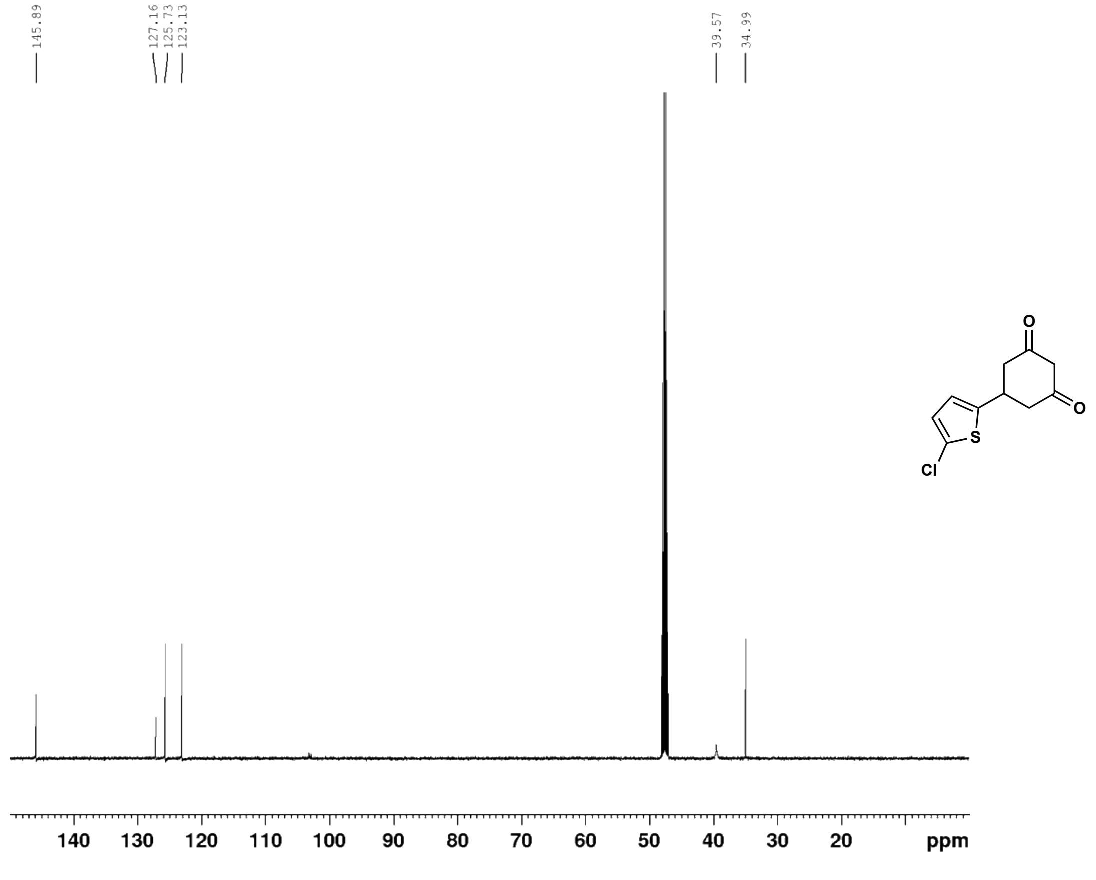

6-(5-chlorothiophen-2-yl)-4-oxo-4,5,6,7-tetrahydrobenzofuran-3-carboxylic acid (**32**) $^{1}$H NMR (500 MHz, $d_6$-DMSO)

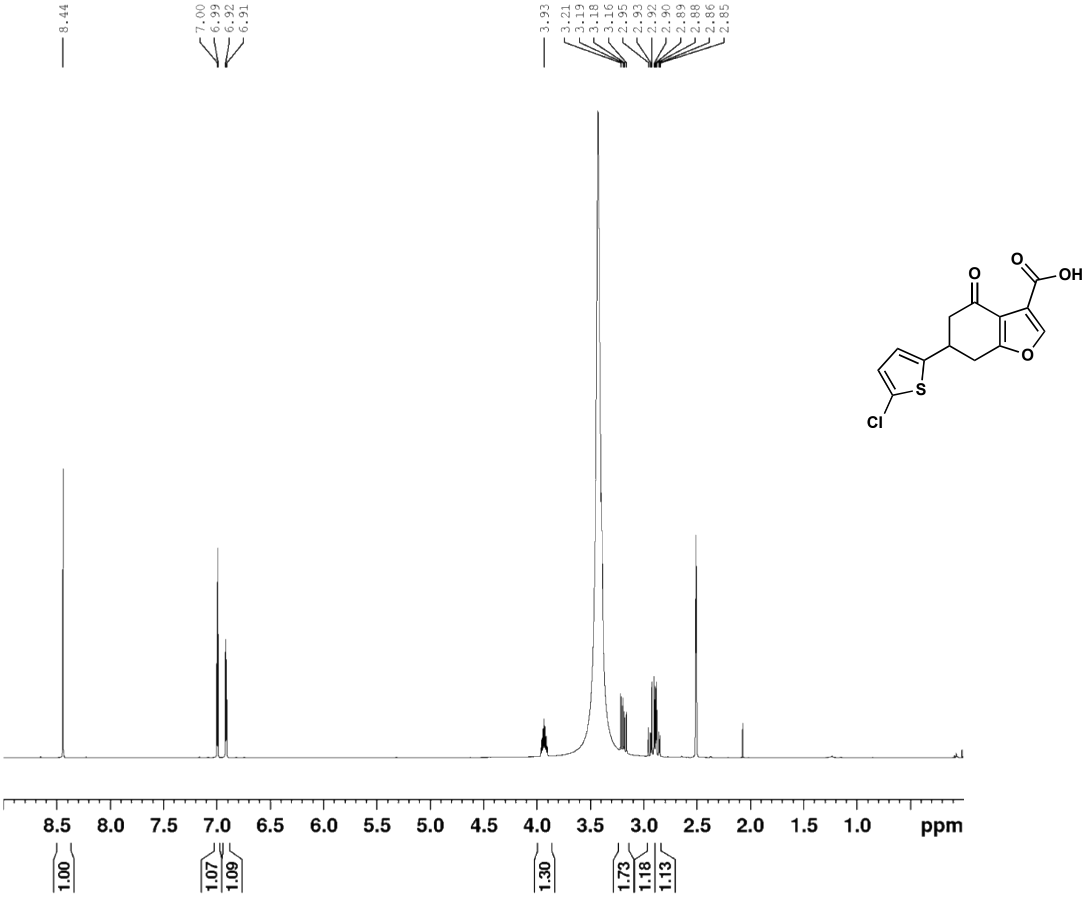

6-(5-chlorothiophen-2-yl)-4-oxo-4,5,6,7-tetrahydrobenzofuran-3-carboxylic acid (**32**) $^{13}$C NMR (126 MHz, $d_6$-DMSO)

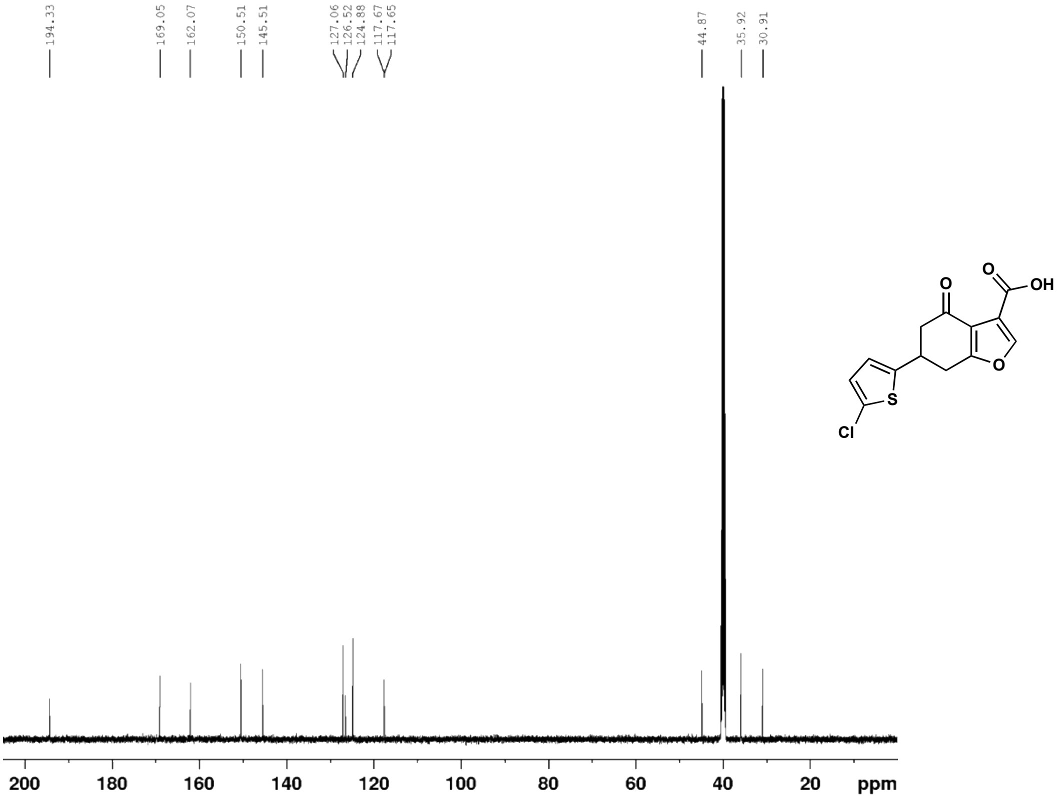

(E)-4-(4-bromothiophen-2-yl)but-3-en-2-one (**33b**) $^{1}H$ NMR (500 MHz, $CDCl_3$)

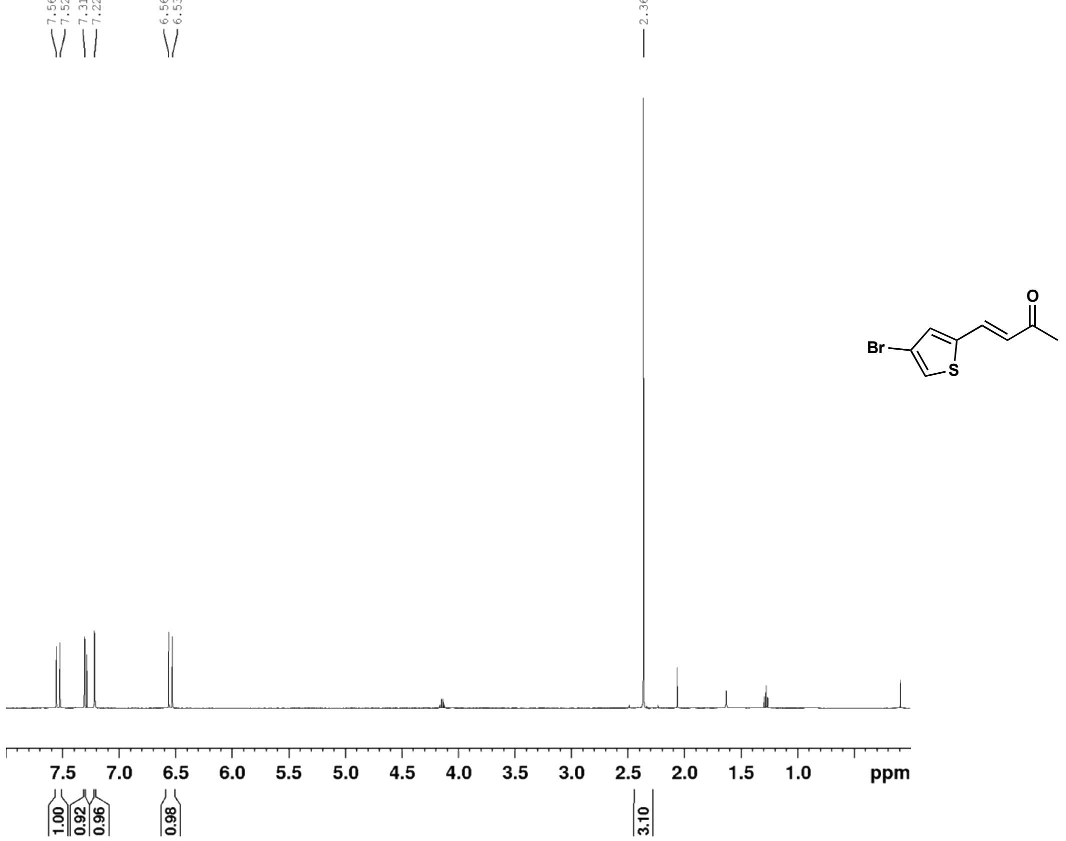

(E)-4-(4-bromothiophen-2-yl)but-3-en-2-one (**33b**) $^{13}C$ NMR (126 MHz, $CDCl_3$)

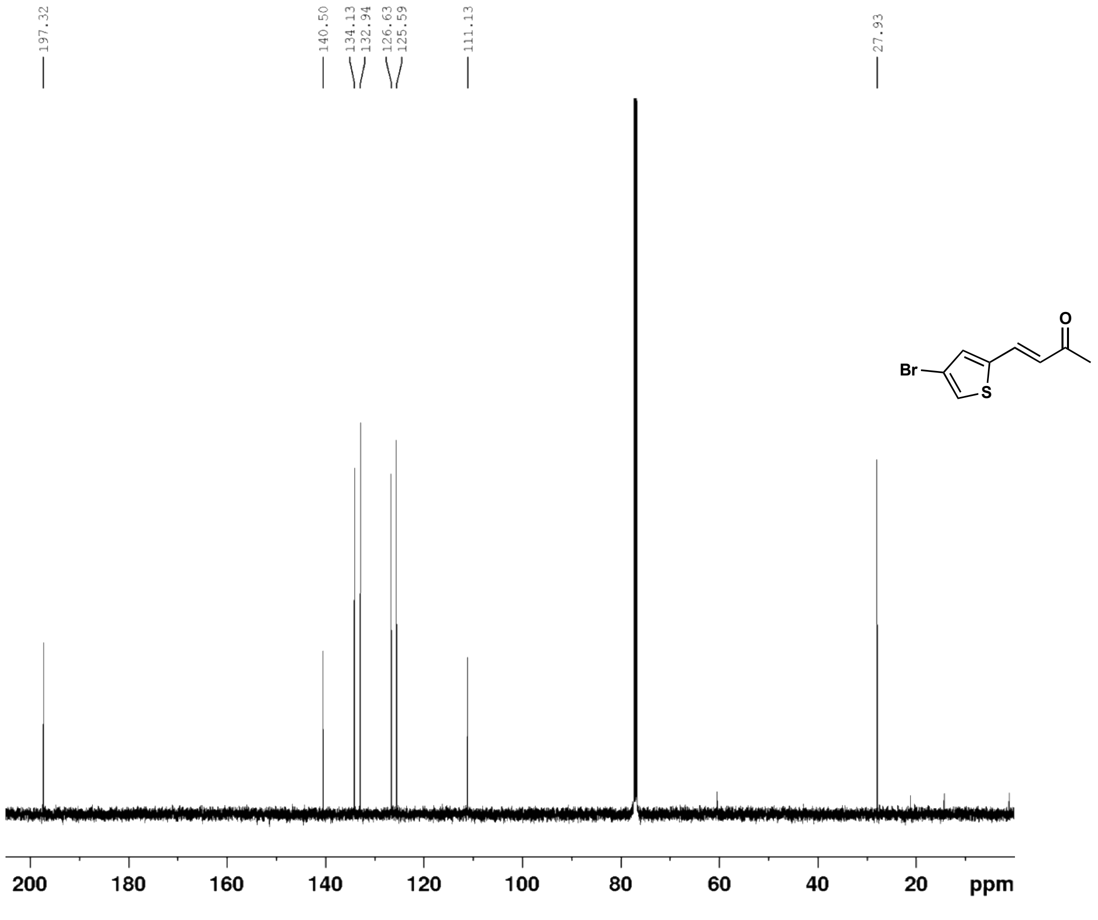

5-(4-bromothiophen-2-yl)cyclohexane-1,3-dione (**33c**) $^{1}H$ NMR (500 MHz, $CD_3OD$)

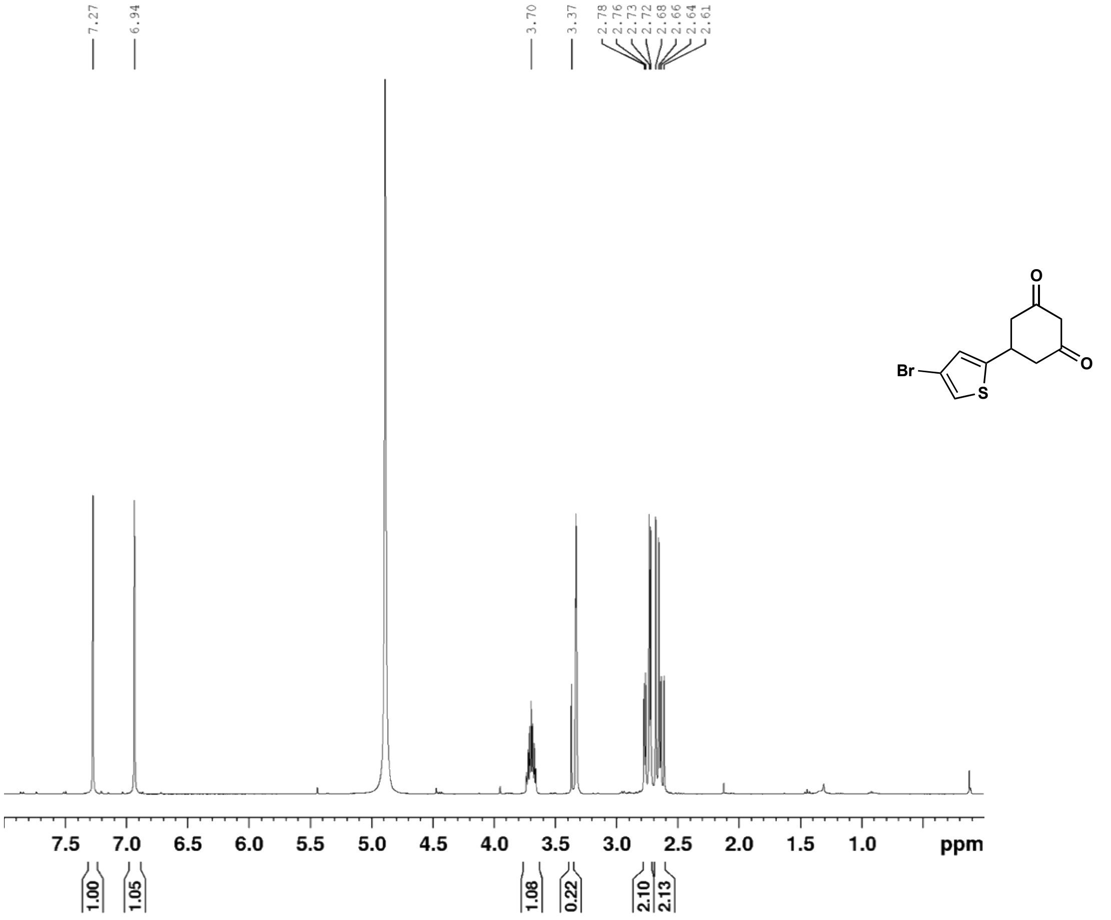

5-(4-bromothiophen-2-yl)cyclohexane-1,3-dione (**33c**) $^{13}C$ NMR (126 MHz, $CD_3OD$)

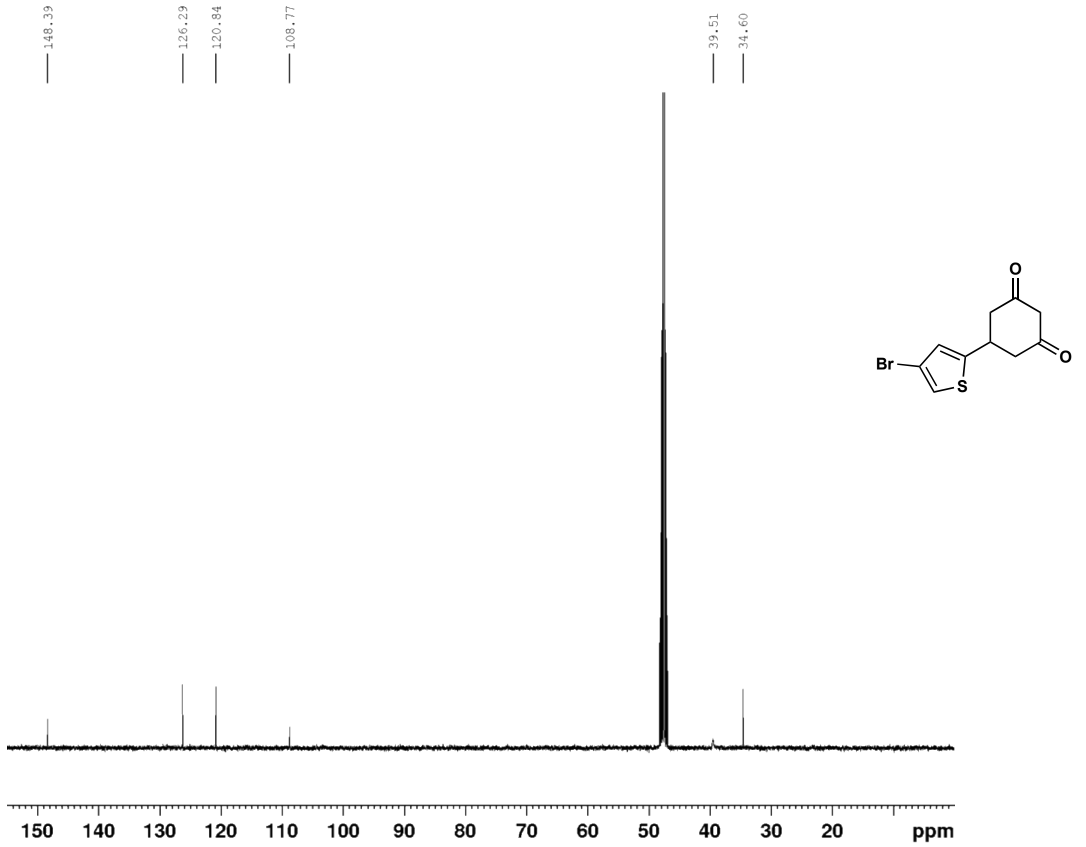

6-(4-bromothiophen-2-yl)-4-oxo-4,5,6,7-tetrahydrobenzofuran-3-carboxylic acid (**33**) $^{1}$H NMR (500 MHz, $d_6$-DMSO)

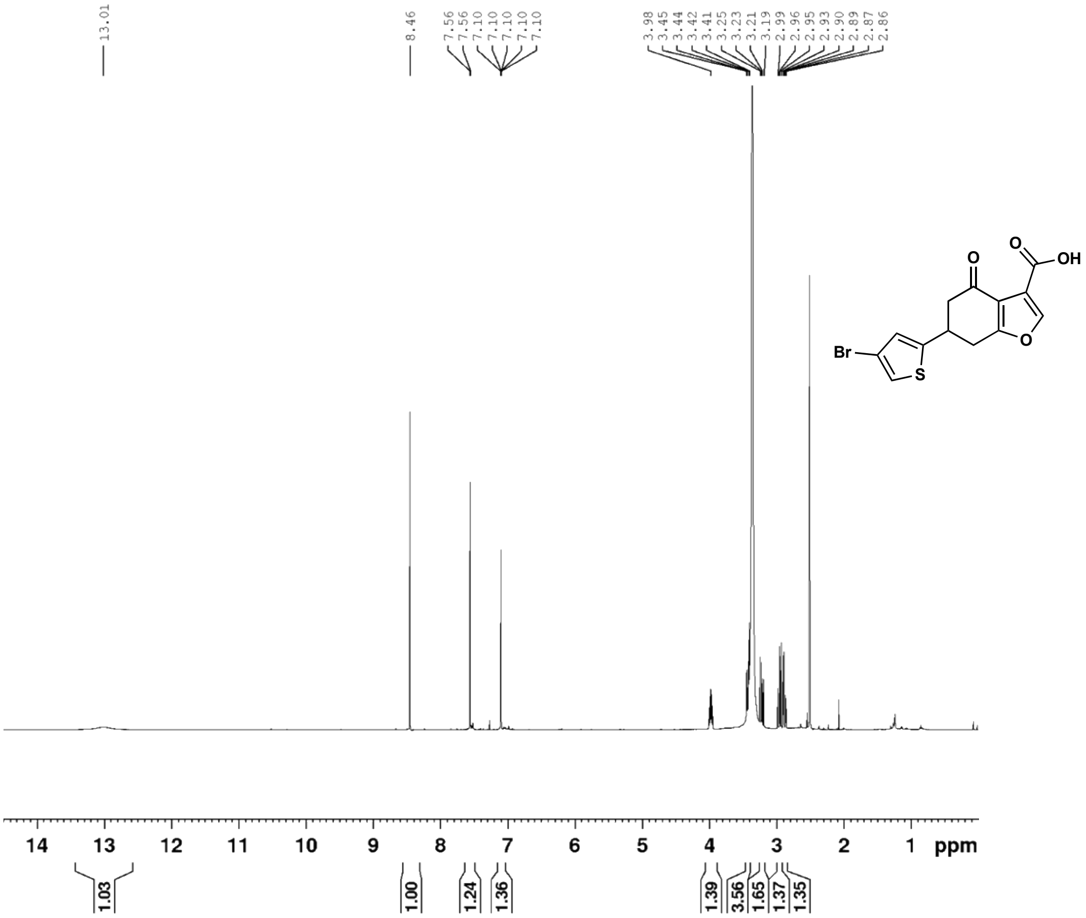

6-(4-bromothiophen-2-yl)-4-oxo-4,5,6,7-tetrahydrobenzofuran-3-carboxylic acid (**33**) $^{13}$C NMR (126 MHz, $d_6$-DMSO)

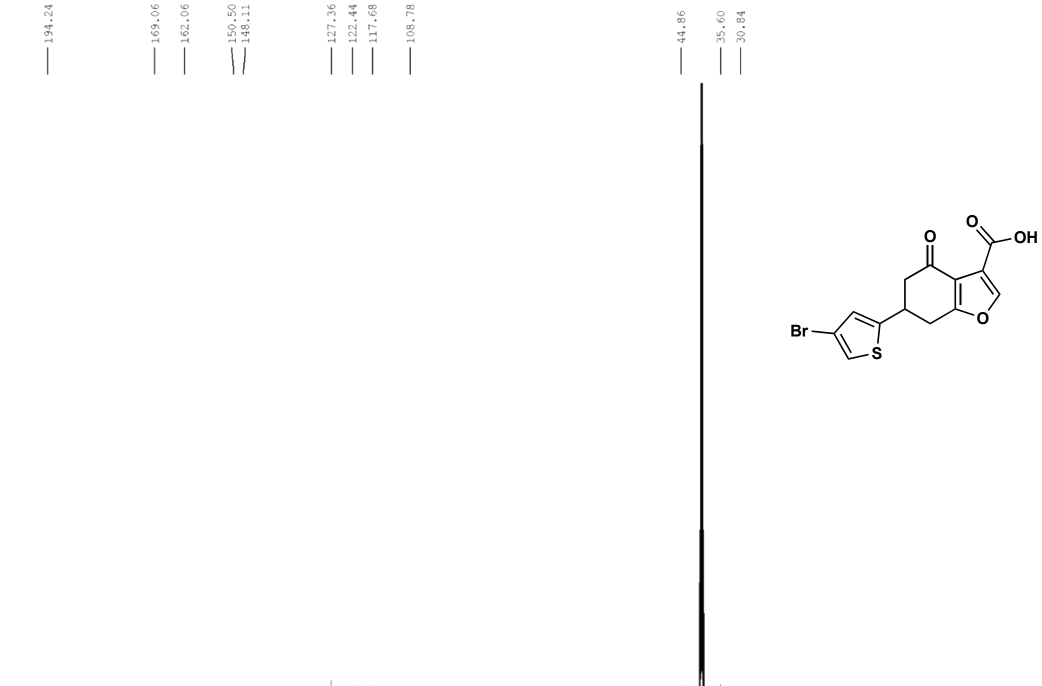

ethyl 6-(4-bromothiophen-2-yl)-4-oxo-4,5,6,7-tetrahydrobenzofuran-3-carboxylate (**33d**) $^{1}$H NMR (500 MHz, $CDCl_3$)

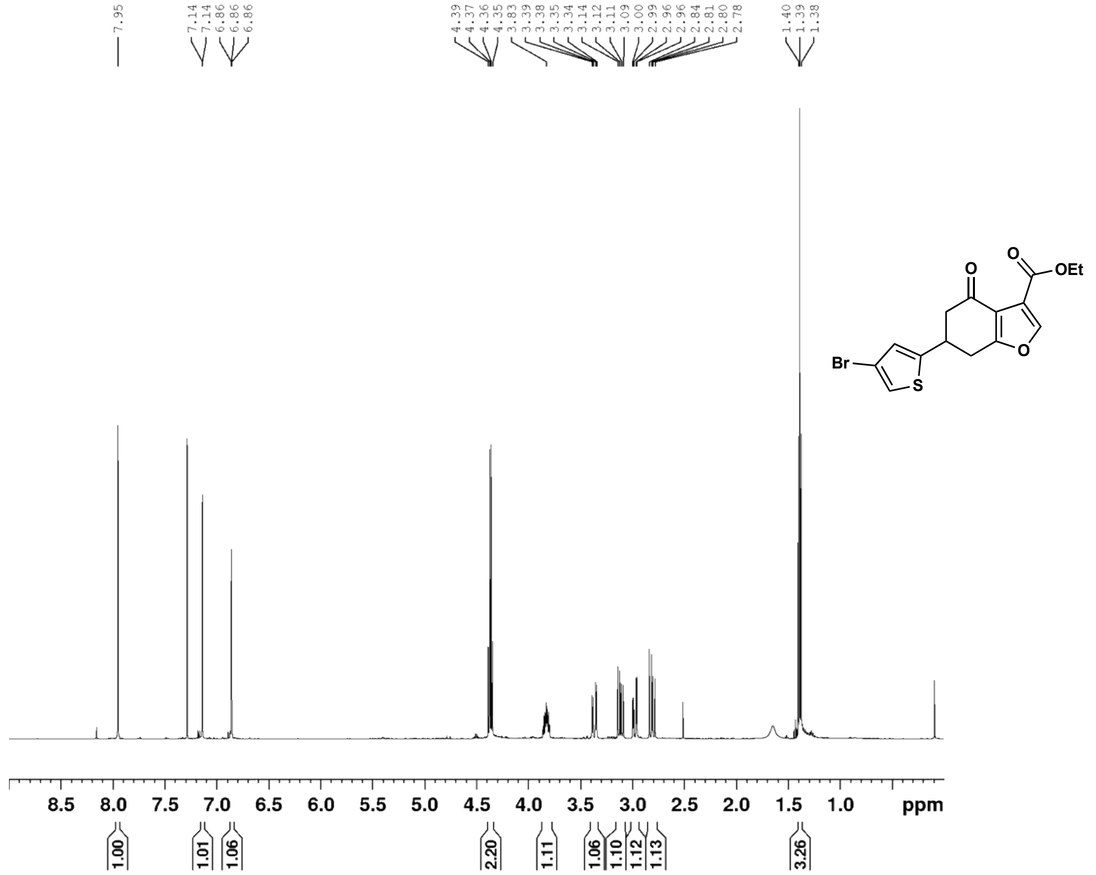

ethyl 6-(4-bromothiophen-2-yl)-4-oxo-4,5,6,7-tetrahydrobenzofuran-3-carboxylate (**33d**) $^{13}$C NMR (126 MHz, $CDCl_3$)

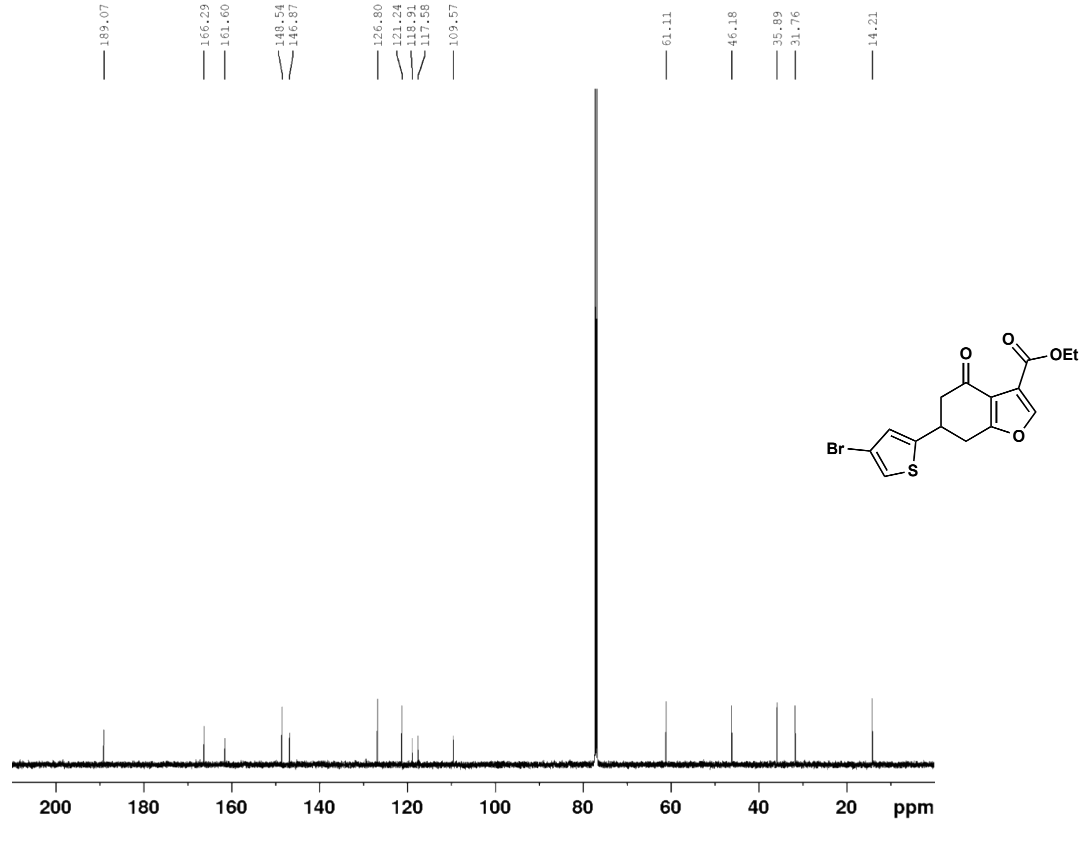

(E)-4-(4,5-dibromothiophen-2-yl)but-3-en-2-one (**34b**) $^{1}H$ NMR (500 MHz, $CDCl_3$)

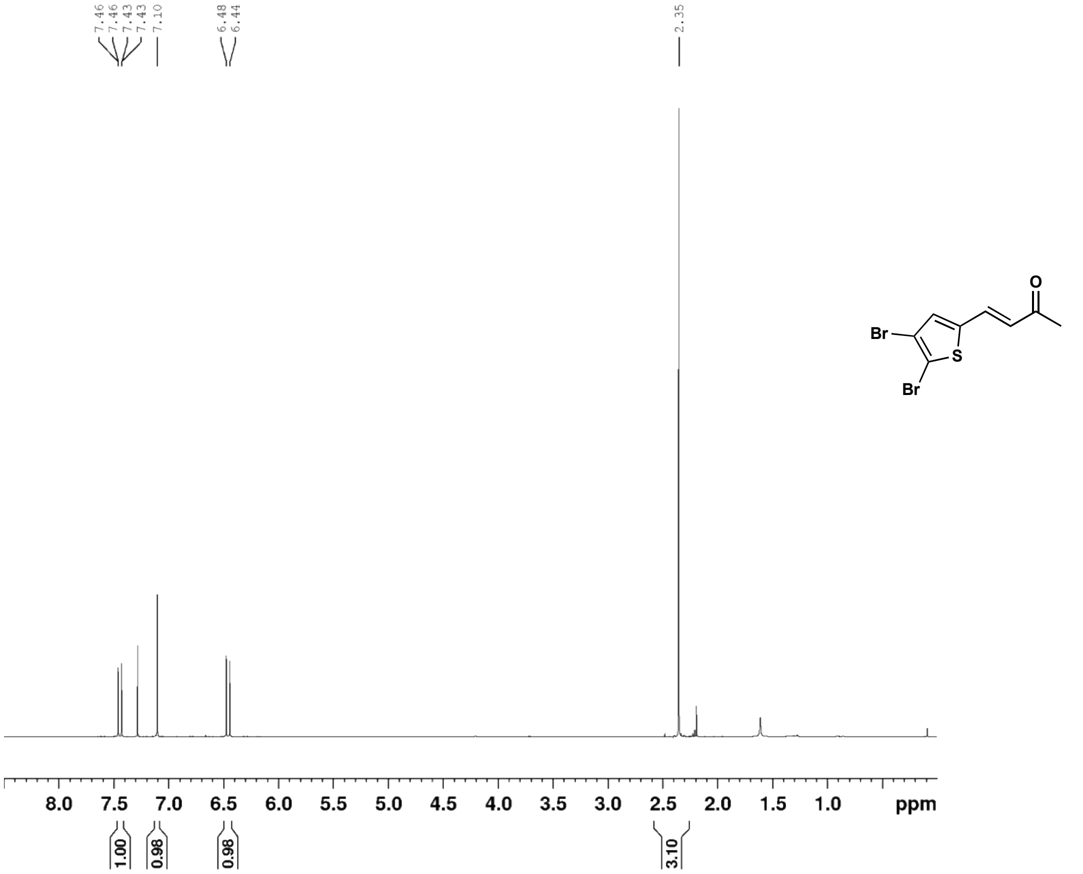

(E)-4-(4,5-dibromothiophen-2-yl)but-3-en-2-one (**34b**) $^{13}C$ NMR (126 MHz, $CDCl_3$)

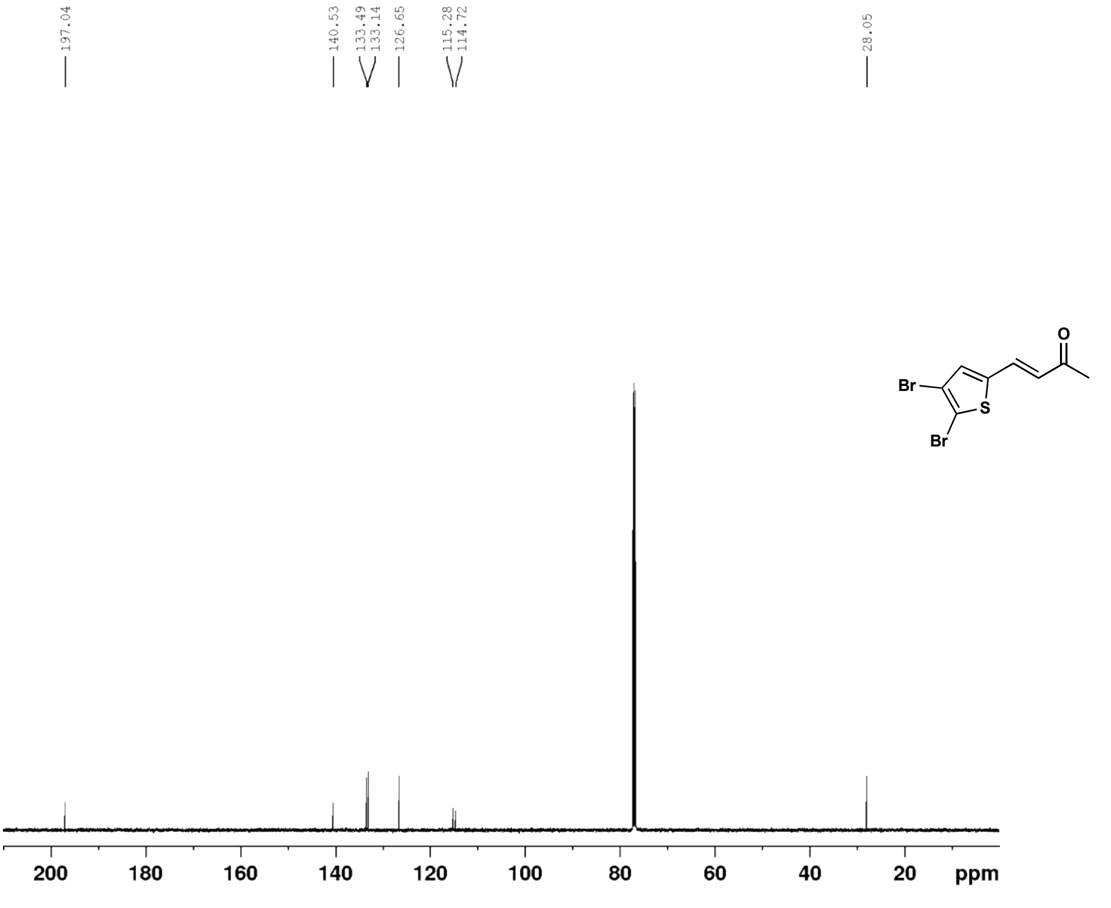

5-(4,5-dibromothiophen-2-yl)cyclohexane-1,3-dione (**34c**) $^{1}$H NMR (500 MHz, $d_6$-DMSO)

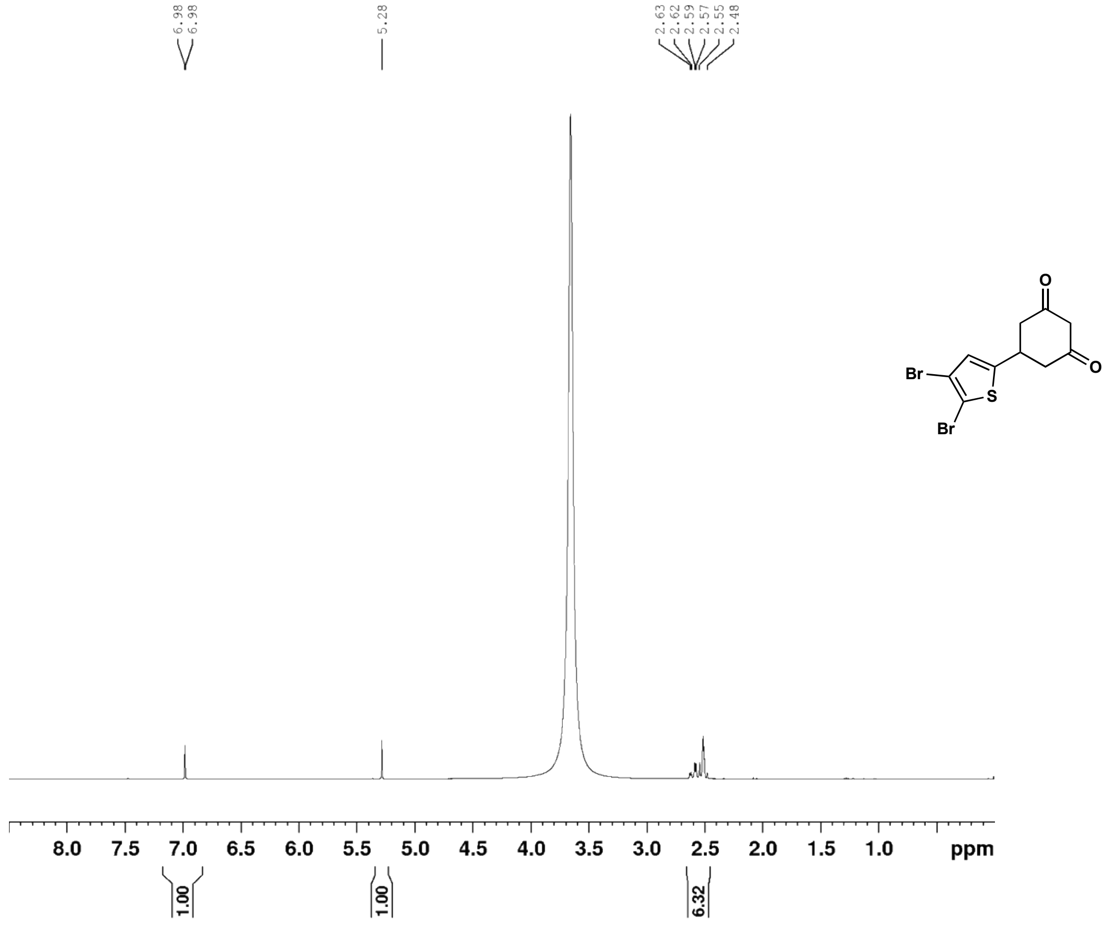

5-(4,5-dibromothiophen-2-yl)cyclohexane-1,3-dione (**34c**) $^{13}$C NMR (126 MHz, $d_6$-DMSO)

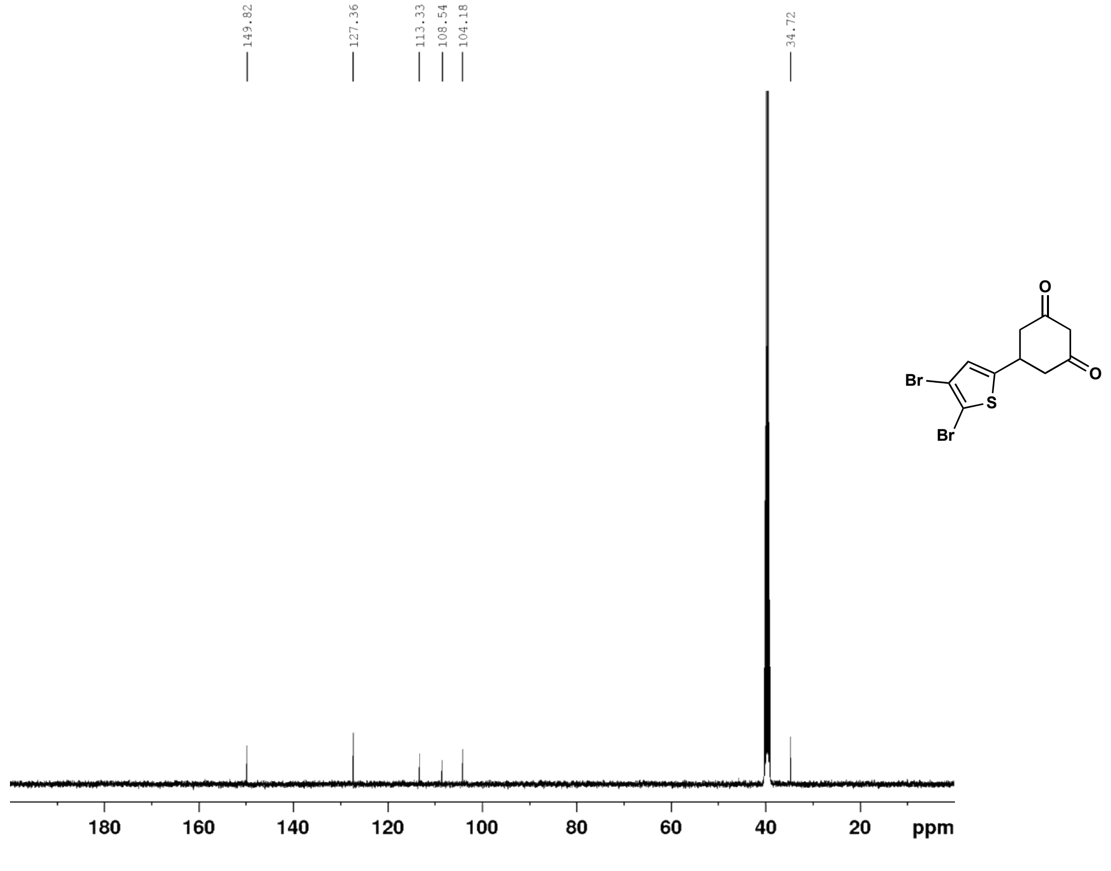

6-(4,5-dibromothiophen-2-yl)-4-oxo-4,5,6,7-tetrahydrobenzofuran-3-carboxylic acid (**34**) $^{1}$H NMR (500 MHz, $d_6$-DMSO)

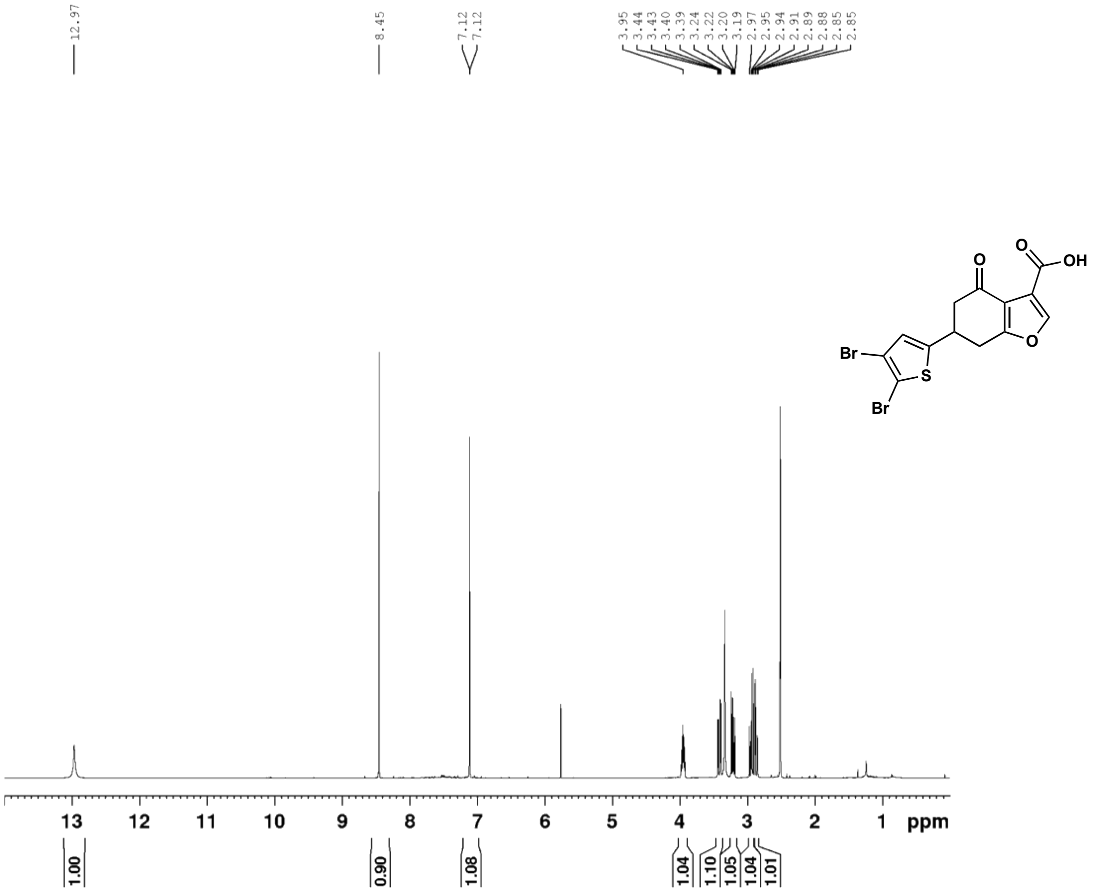

6-(4,5-dibromothiophen-2-yl)-4-oxo-4,5,6,7-tetrahydrobenzofuran-3-carboxylic acid (**34**) $^{13}$C NMR (126 MHz, $d_6$-DMSO)

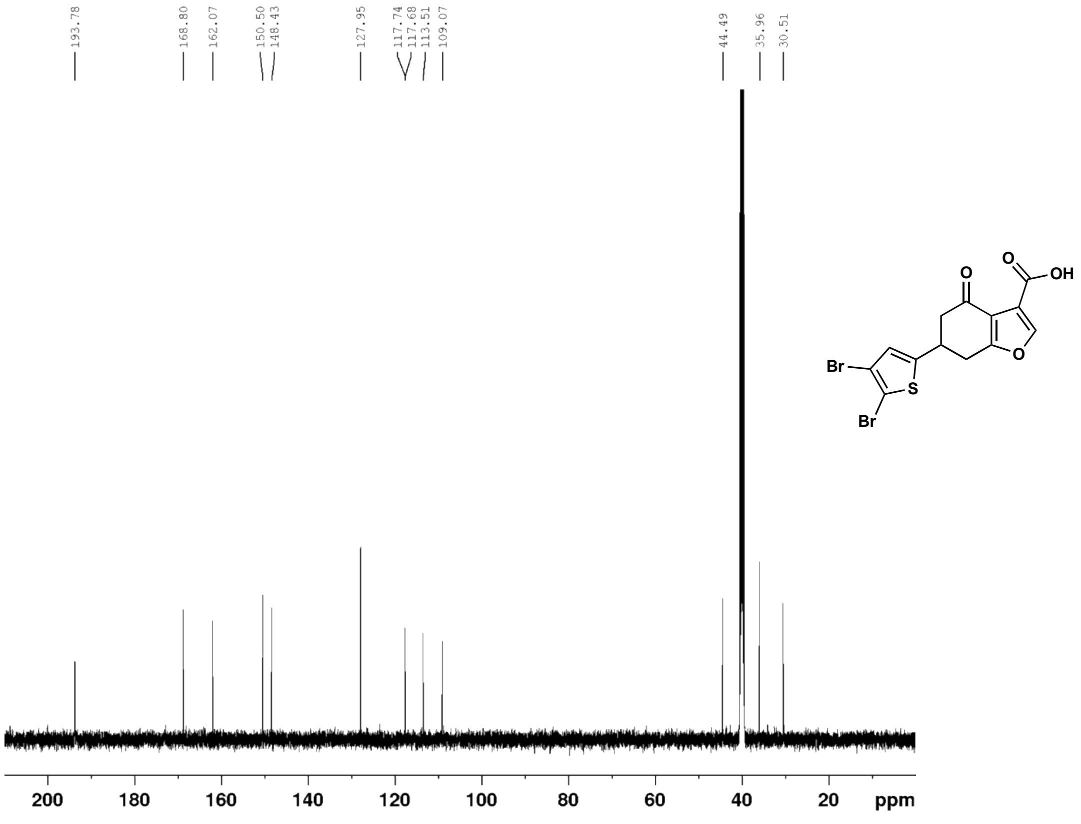

ethyl 2-(1-oxo-1,2,3,4-tetrahydronaphthalen-2-yl)acetate (**36**) $^{1}$H NMR (500 MHz, $CDCl_3$)

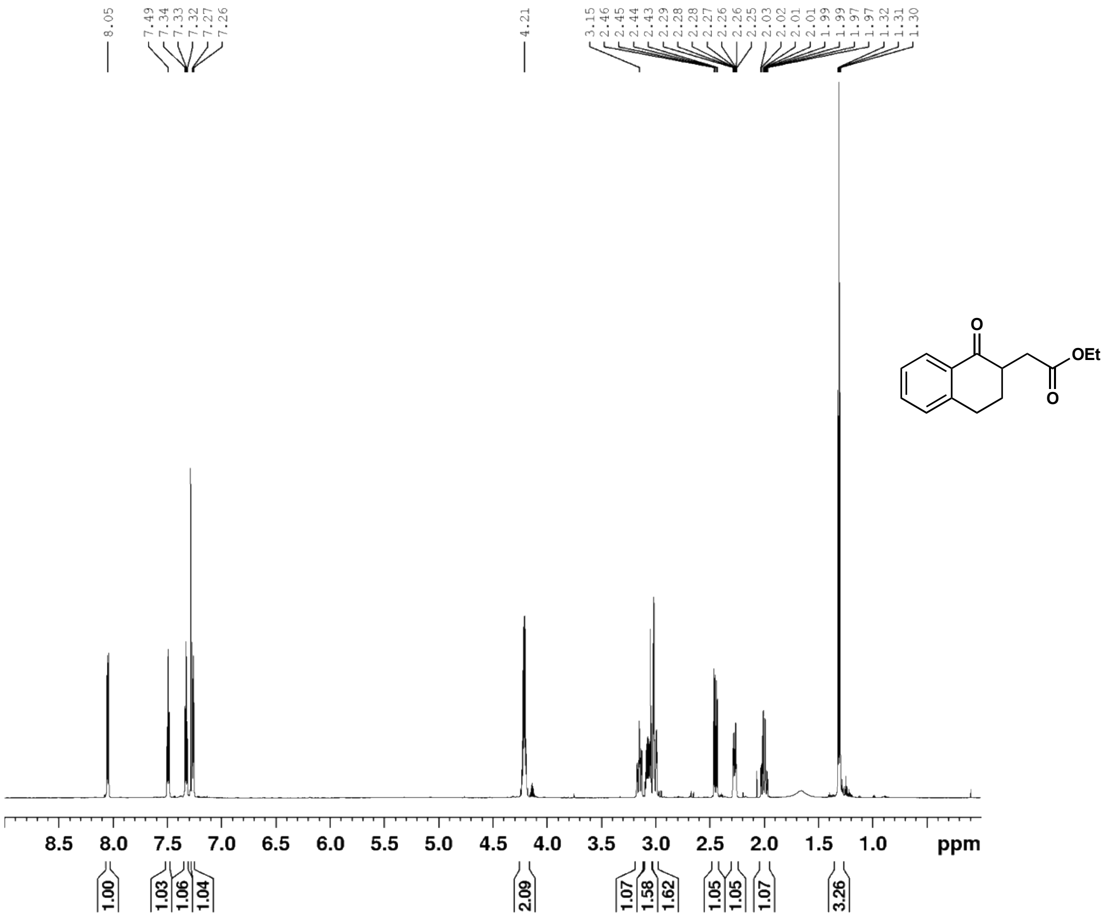

ethyl 2-(1-oxo-1,2,3,4-tetrahydronaphthalen-2-yl)acetate (**36**) $^{13}$C NMR (126 MHz, $CDCl_3$)

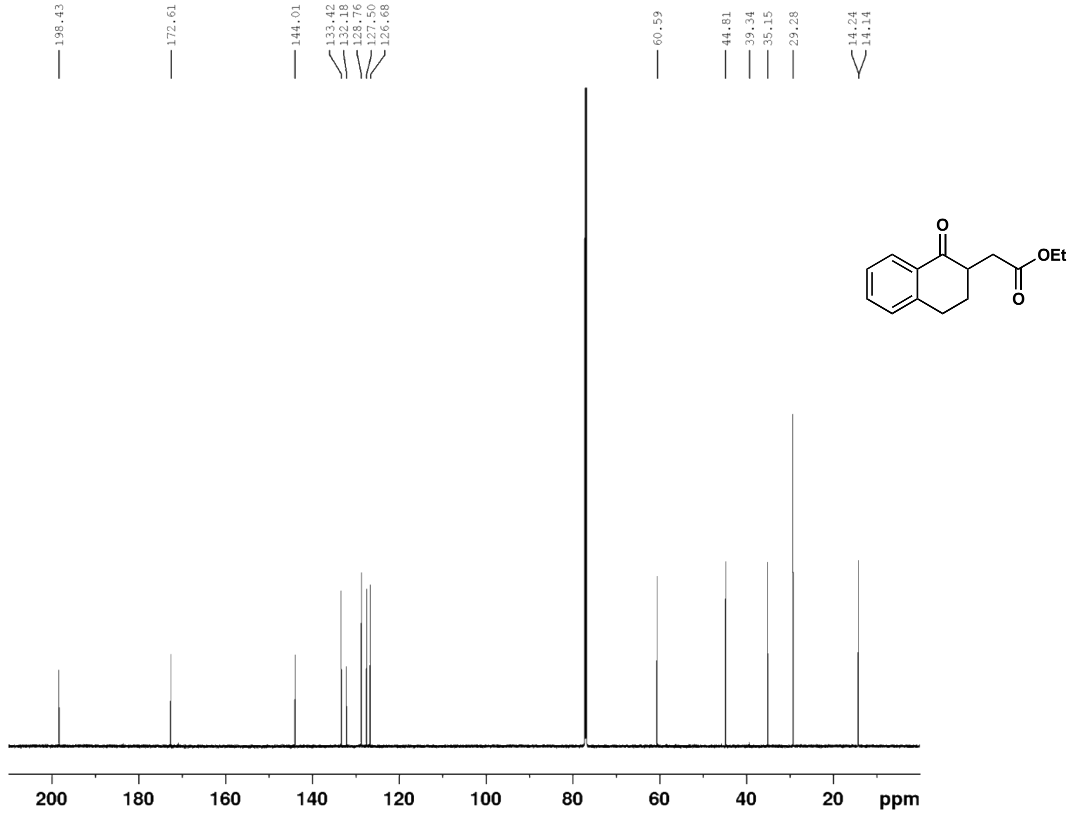

2-(1-oxo-1,2,3,4-tetrahydronaphthalen-2-yl)acetic acid (**37**) $^{1}$H NMR (500 MHz, $CDCl_3$)

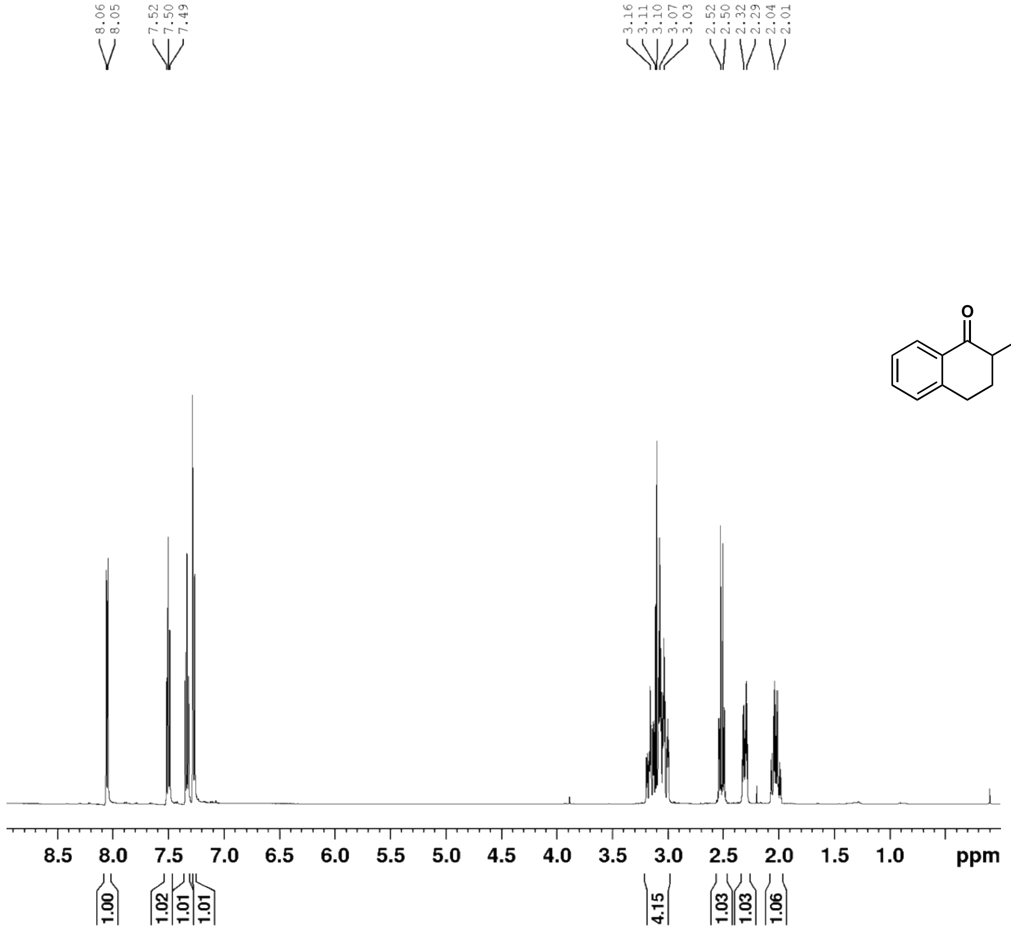

2-(1-oxo-1,2,3,4-tetrahydronaphthalen-2-yl)acetic acid (**37**) $^{13}$C NMR (126 MHz, $CDCl_3$)

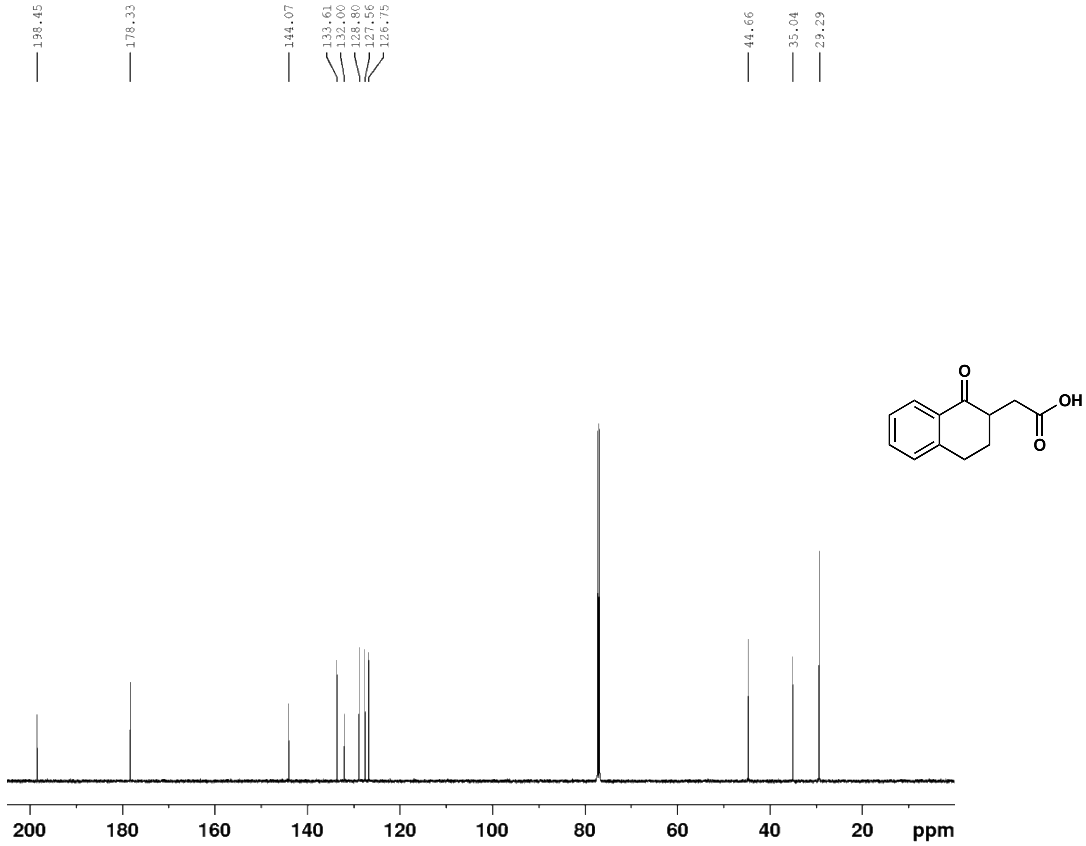

### S3 Compounds HPLC Traces

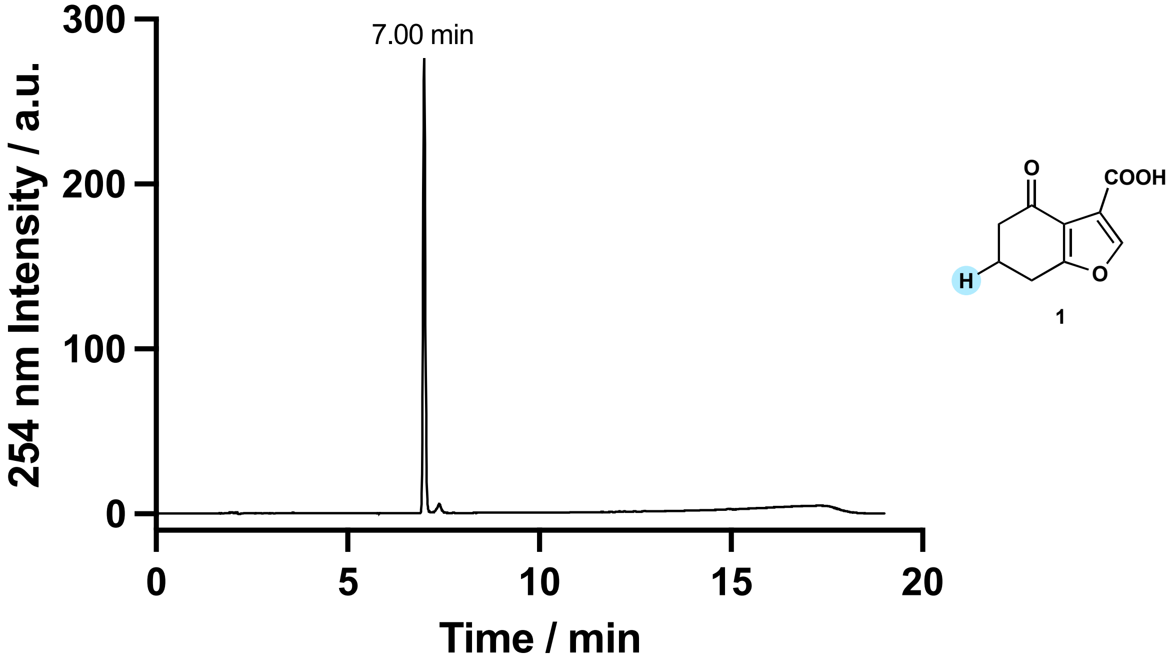

Figure S3 -Compound **1**, HPLC 5–95 % MeCN in $H_2O$. Purity based on integration of 100 %.

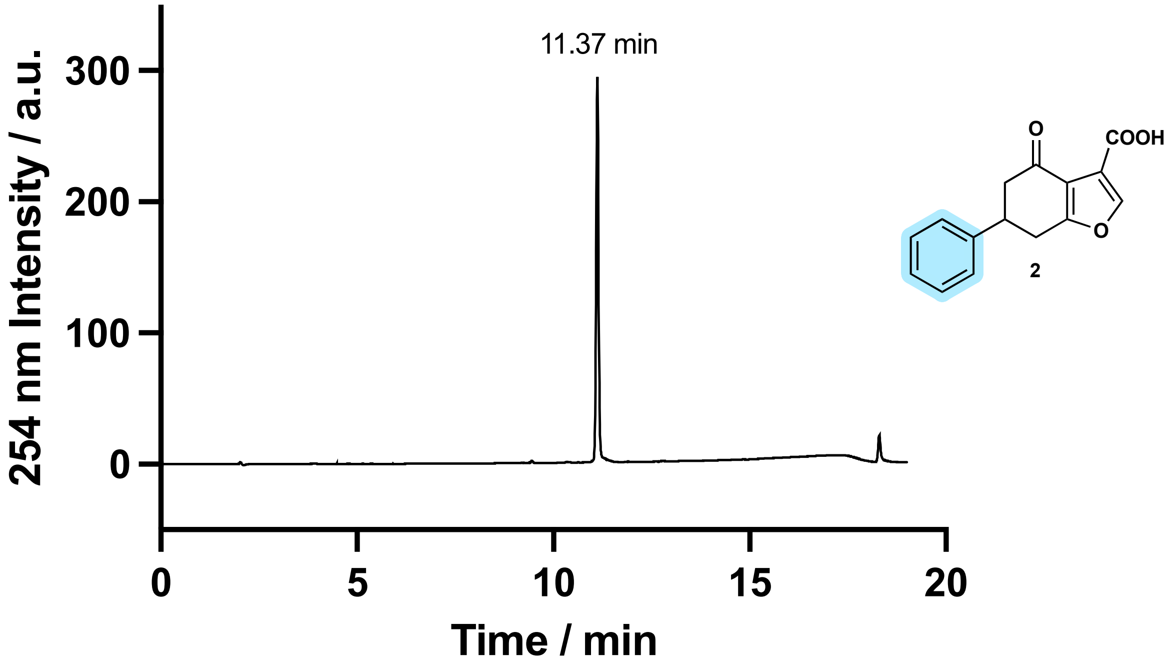

Figure S4 - Compound **2**, HPLC 5–95 % MeCN in $H_2O$. Purity based on integration of 96 %.

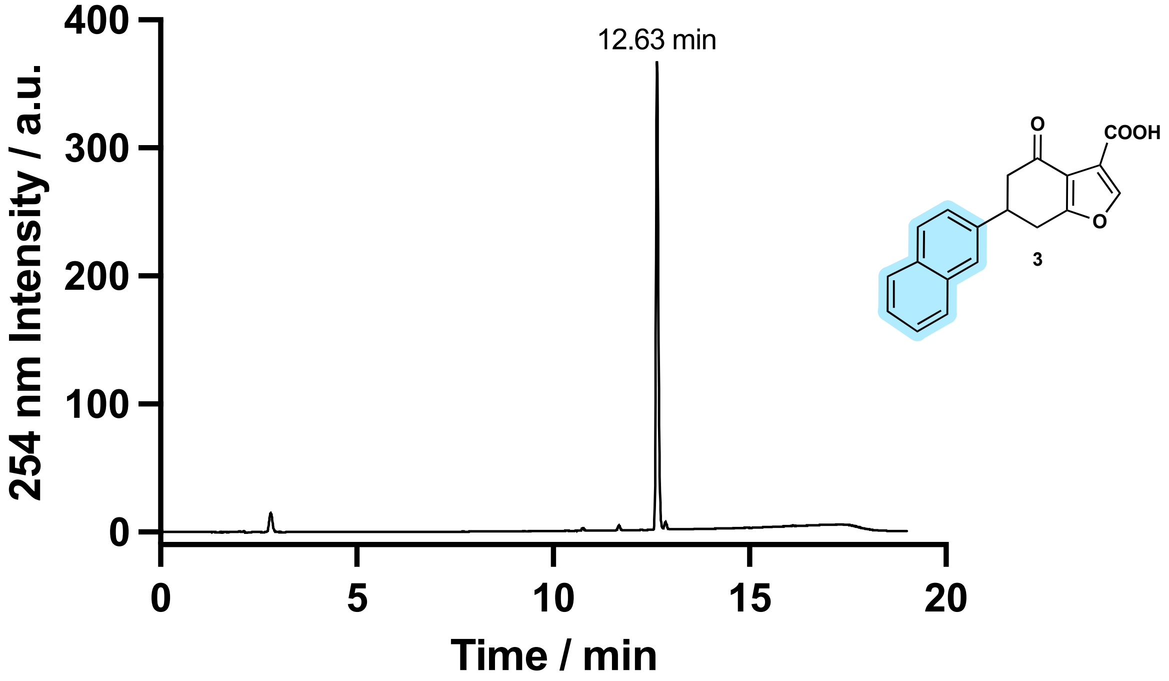

Figure S5 - Compound **3**, HPLC 5–95 % MeCN in $H_2O$. Purity based on integration of 98 %.

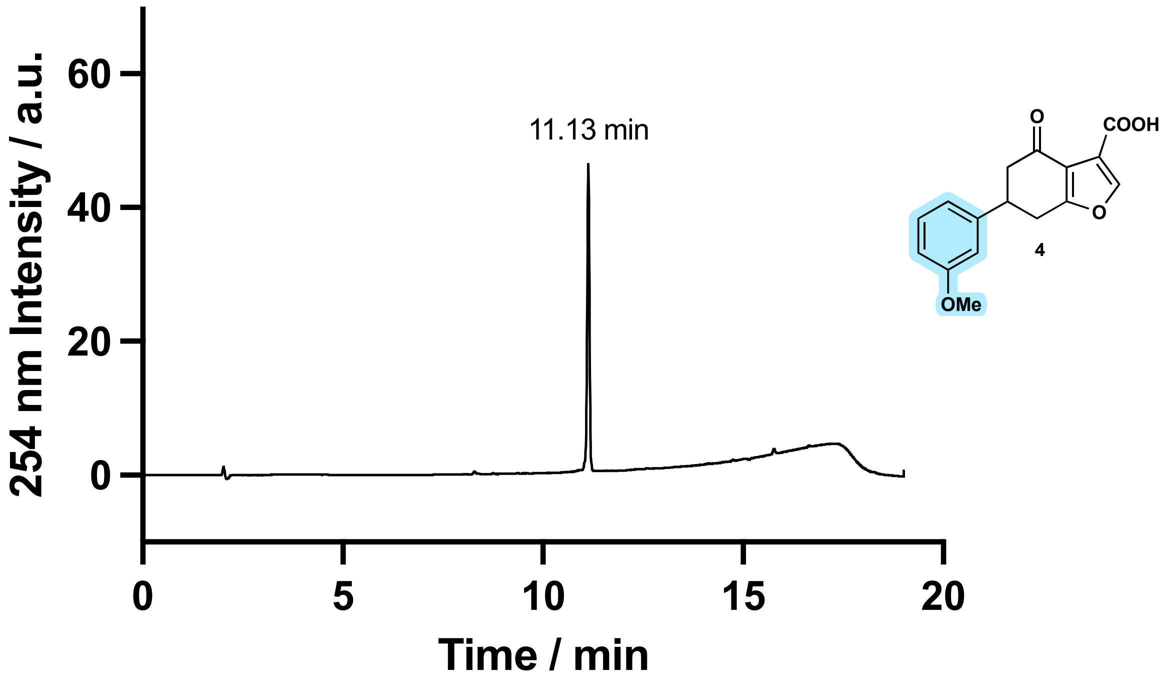

Figure S6 - Compound **4**, HPLC 5–95 % MeCN in $H_2O$. Purity based on integration of 98 %.

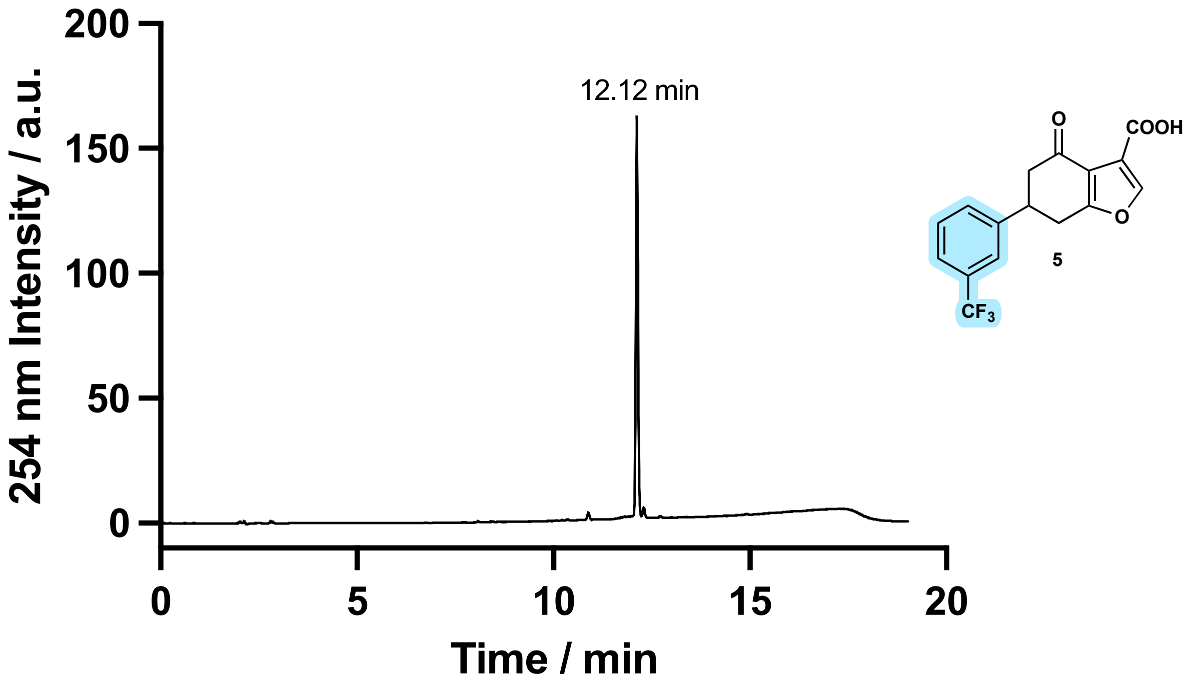

Figure S7 - Compound **5**, HPLC 5–95 % MeCN in $H_2O$. Purity based on integration of 96 %.

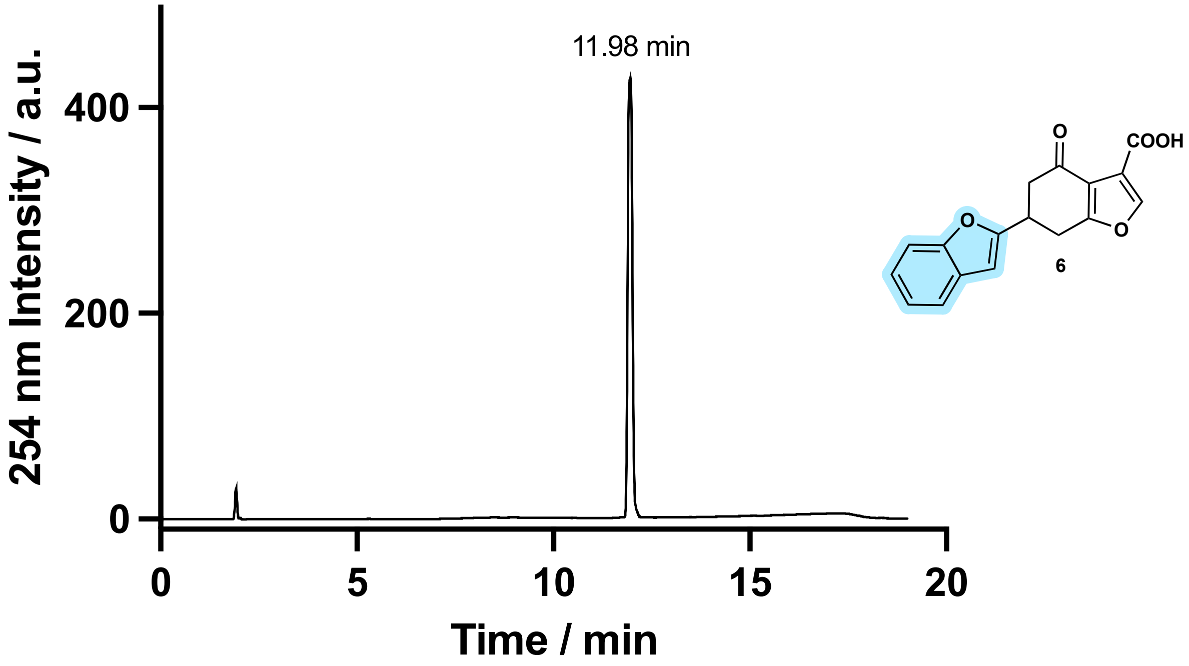

Figure S8 - Compound **6**, HPLC 5–95 % MeCN in $H_2O$. Purity based on integration of 97 %.

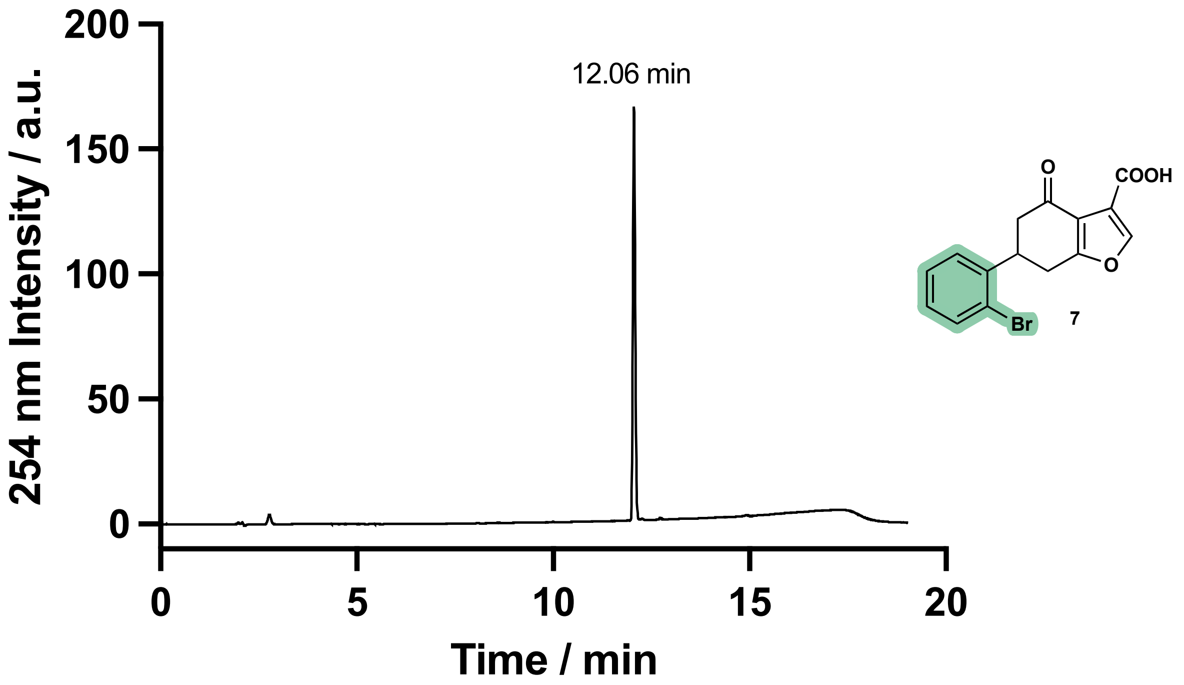

Figure S9 - Compound **7**, HPLC 5–95 % MeCN in $H_2O$. Purity based on integration of 97 %.

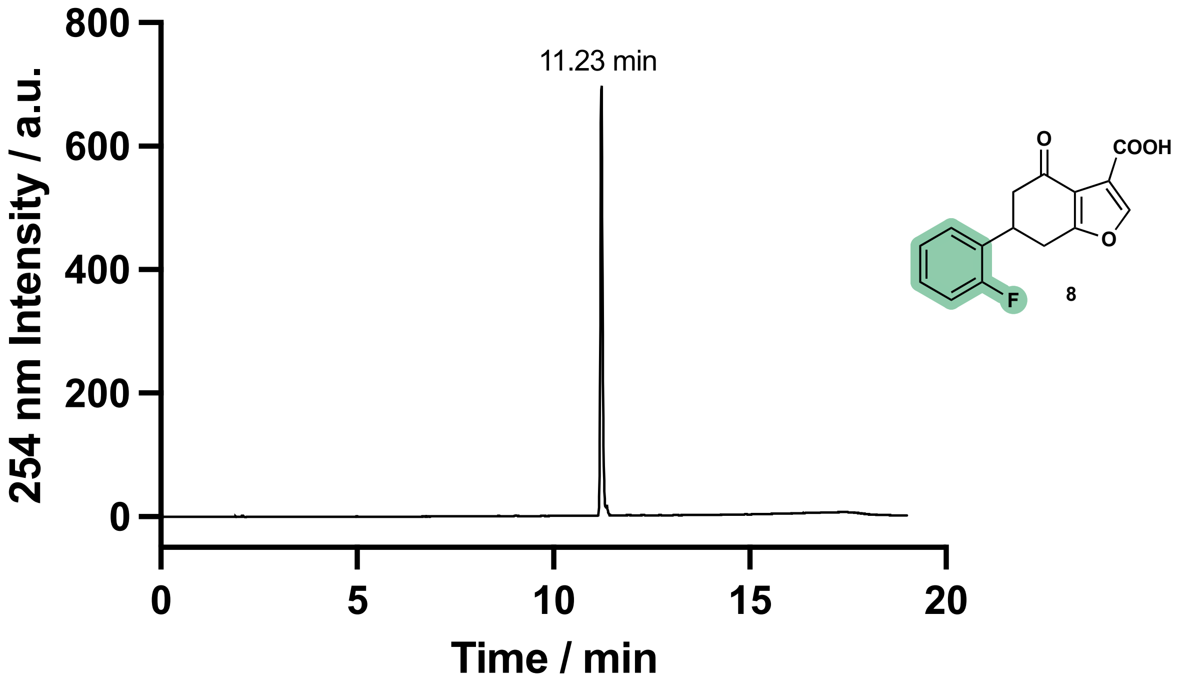

Figure S10 - Compound **8**, HPLC 5–95 % MeCN in $H_2O$. Purity based on integration of 100 %.

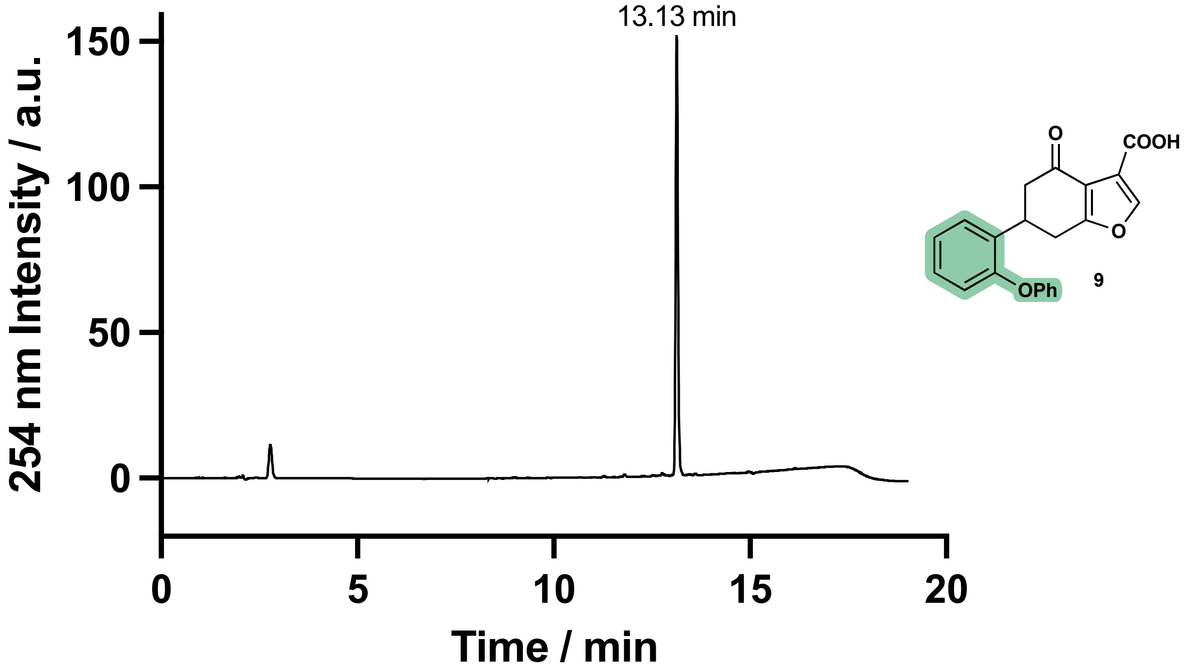

Figure S11 - Compound **9**, HPLC 5–95 % MeCN in $H_2O$. Purity based on integration of 91 %.

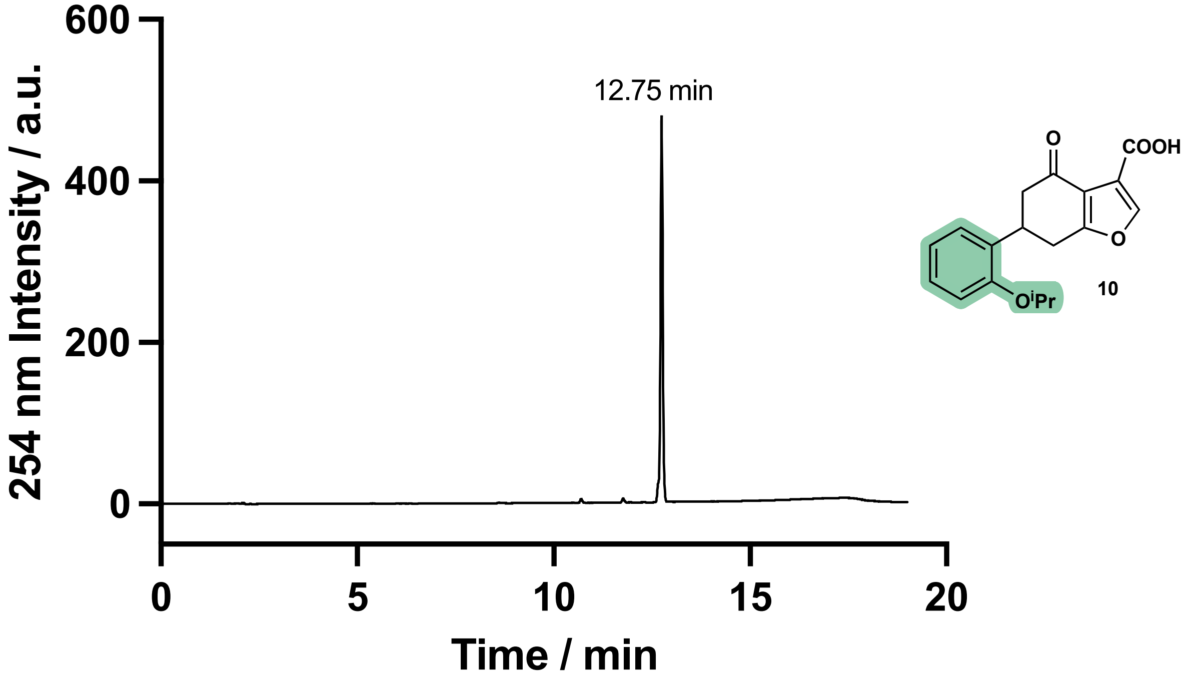

Figure S12 - Compound **10**, HPLC 5–95 % MeCN in $H_2O$. Purity based on integration of 97 %.

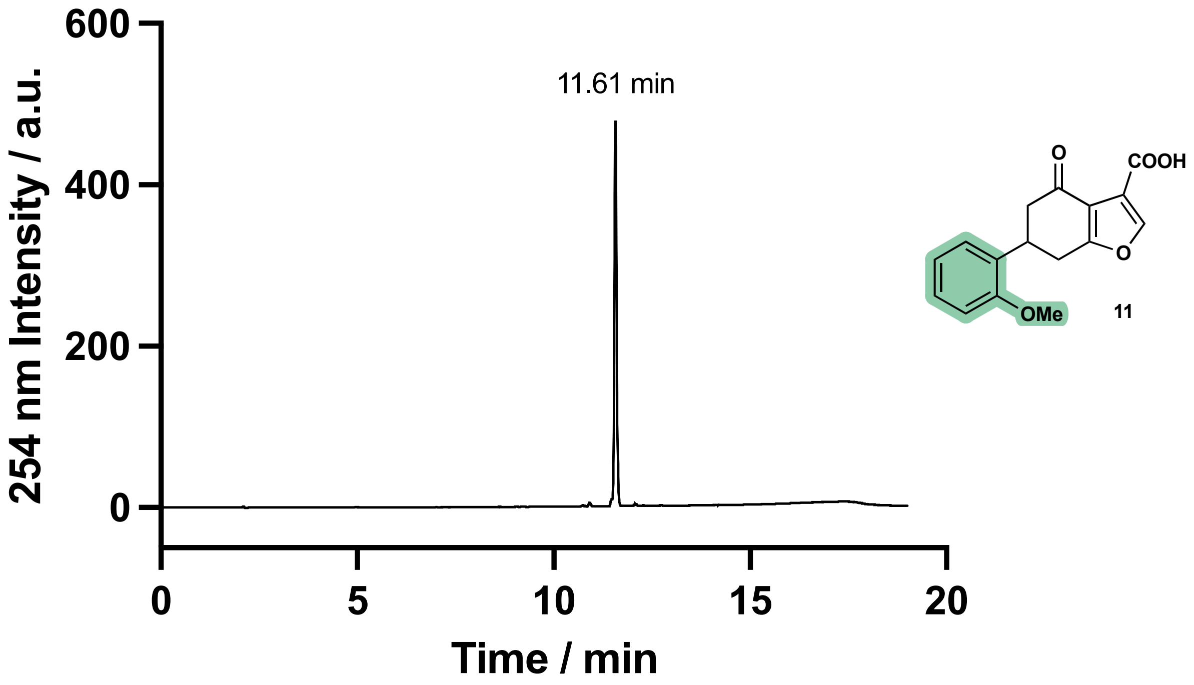

Figure S13 - Compound **11**, HPLC 5–95 % MeCN in $H_2O$. Purity based on integration of 97 %.

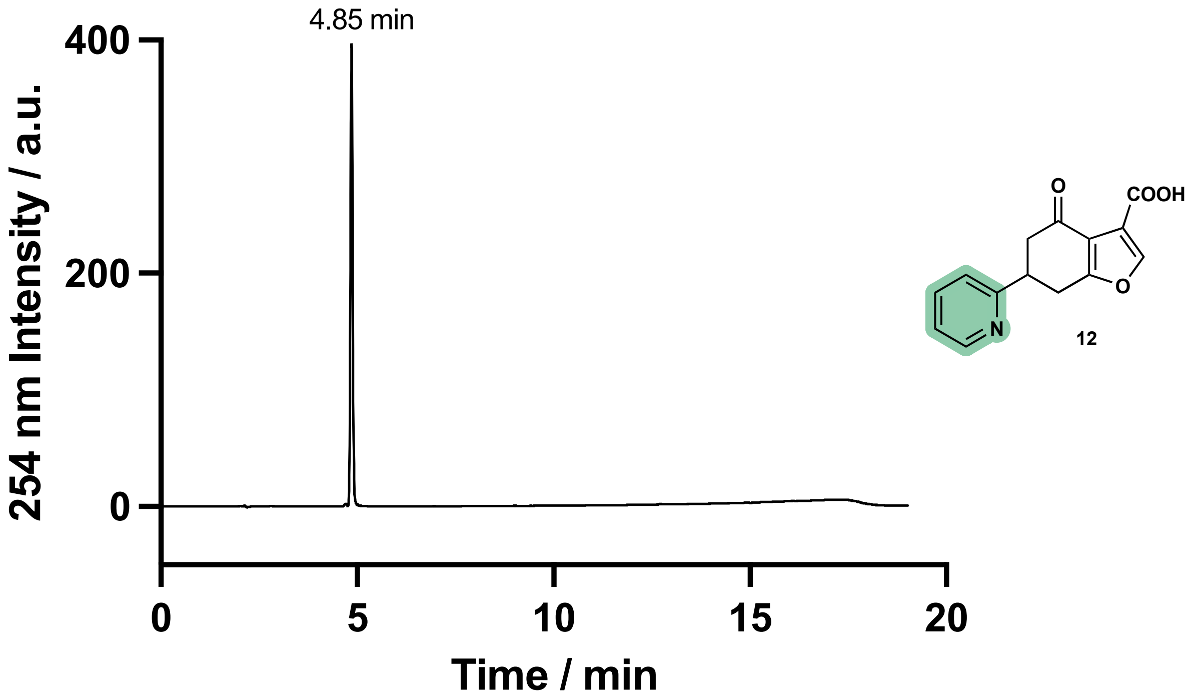

Figure S14 - Compound **12**, HPLC 5–95 % MeCN in $H_2O$. Purity based on integration of 100 %.

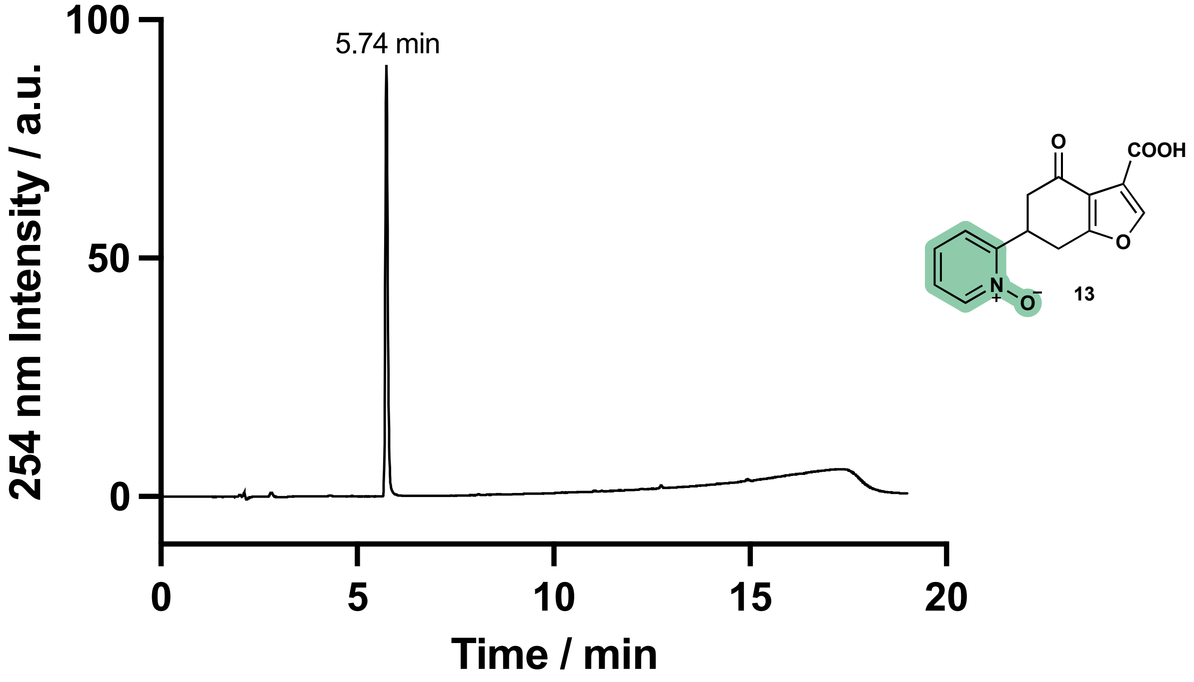

Figure S15 - Compound **13**, HPLC 5–95 % MeCN in $H_2O$. Purity based on integration of 100 %.

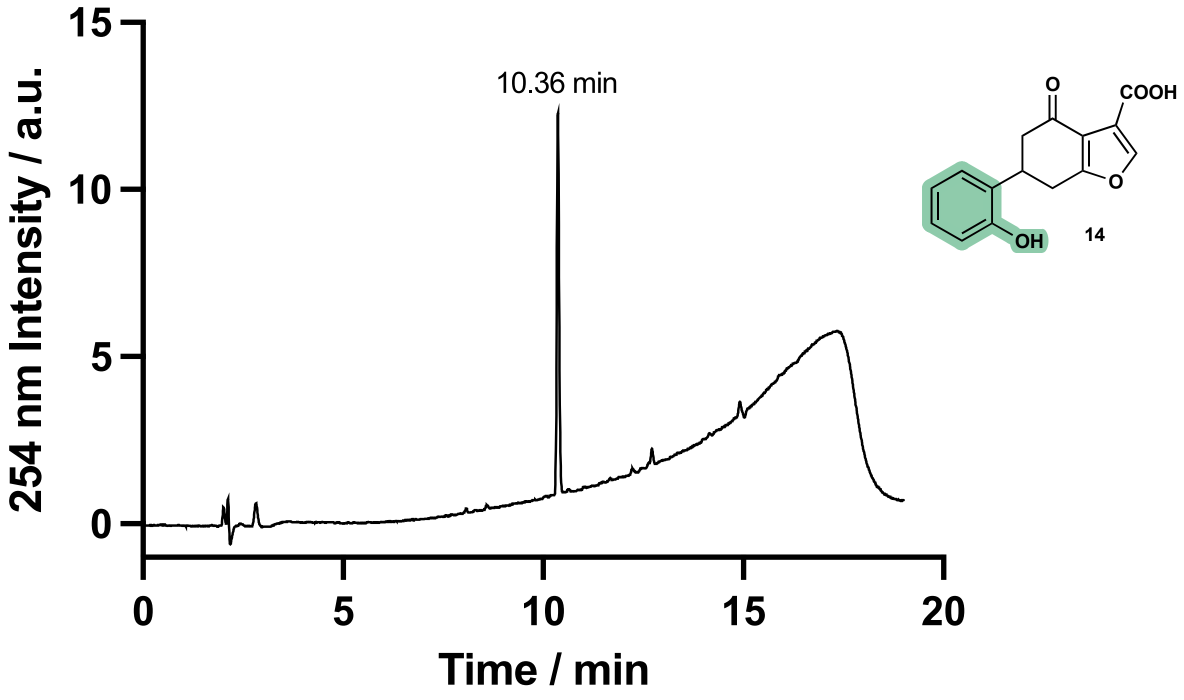

Figure S16 - Compound **14**, HPLC 5–95 % MeCN in $H_2O$. Purity based on integration of 100 %.

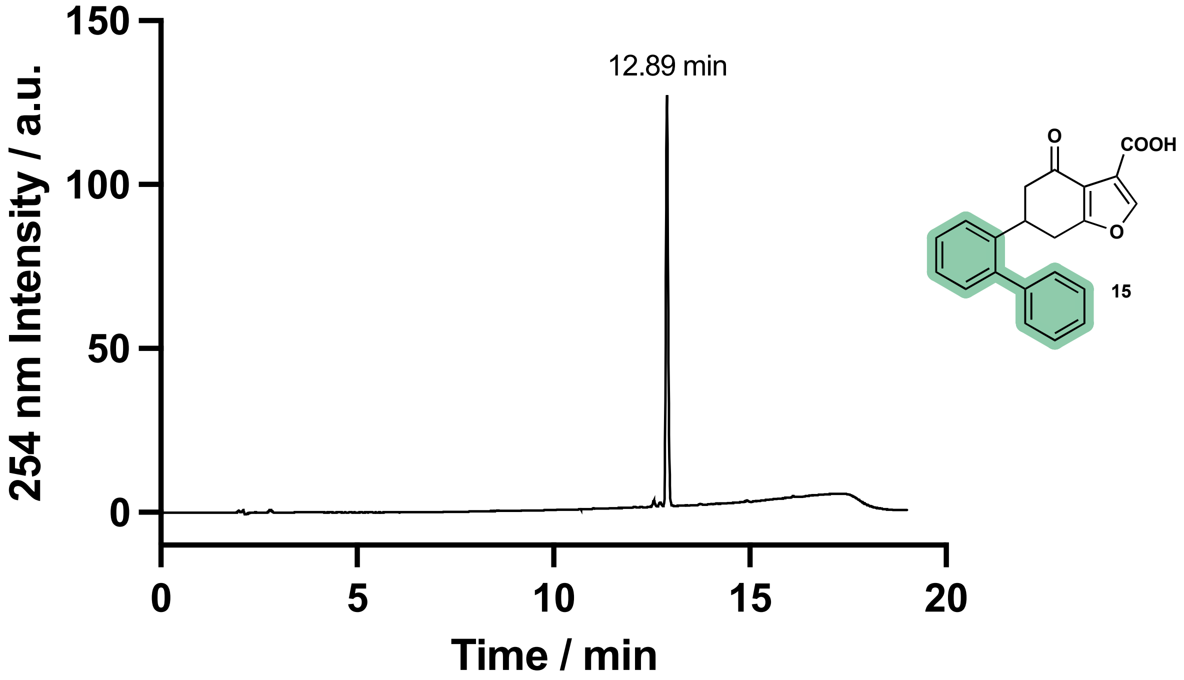

Figure S17 - Compound **15**, HPLC 5–95 % MeCN in $H_2O$. Purity based on integration of 97 %.

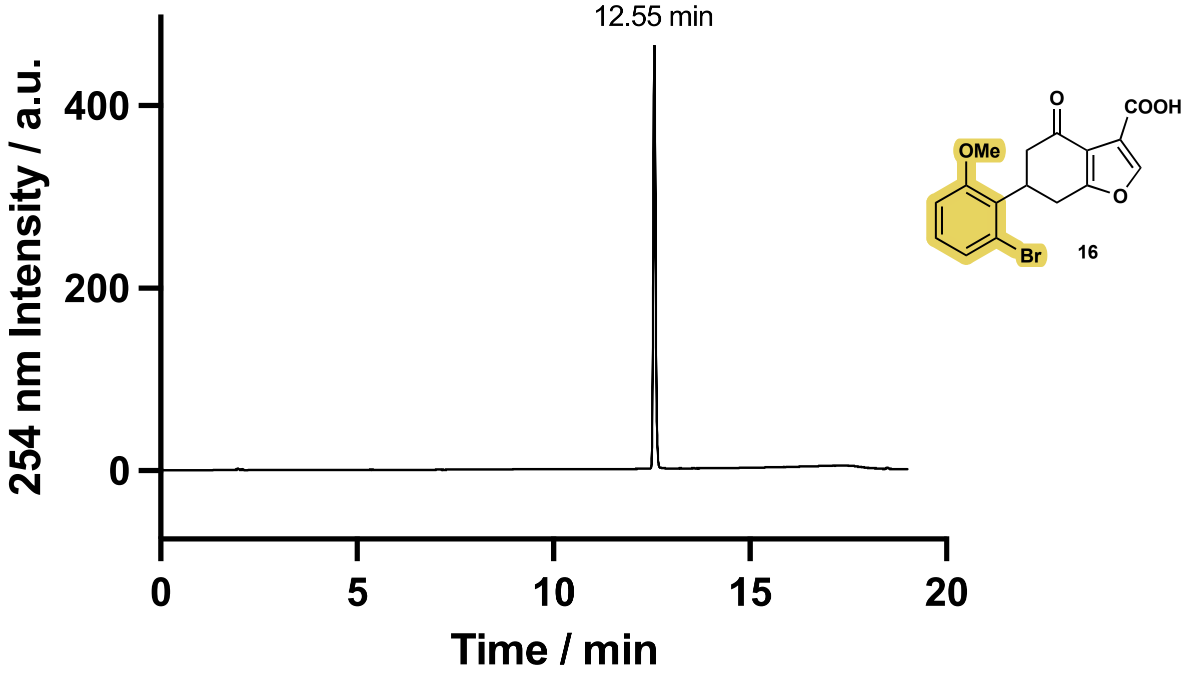

Figure S18 - Compound **16**, HPLC 5–95 % MeCN in $H_2O$. Purity based on integration of 100 %.

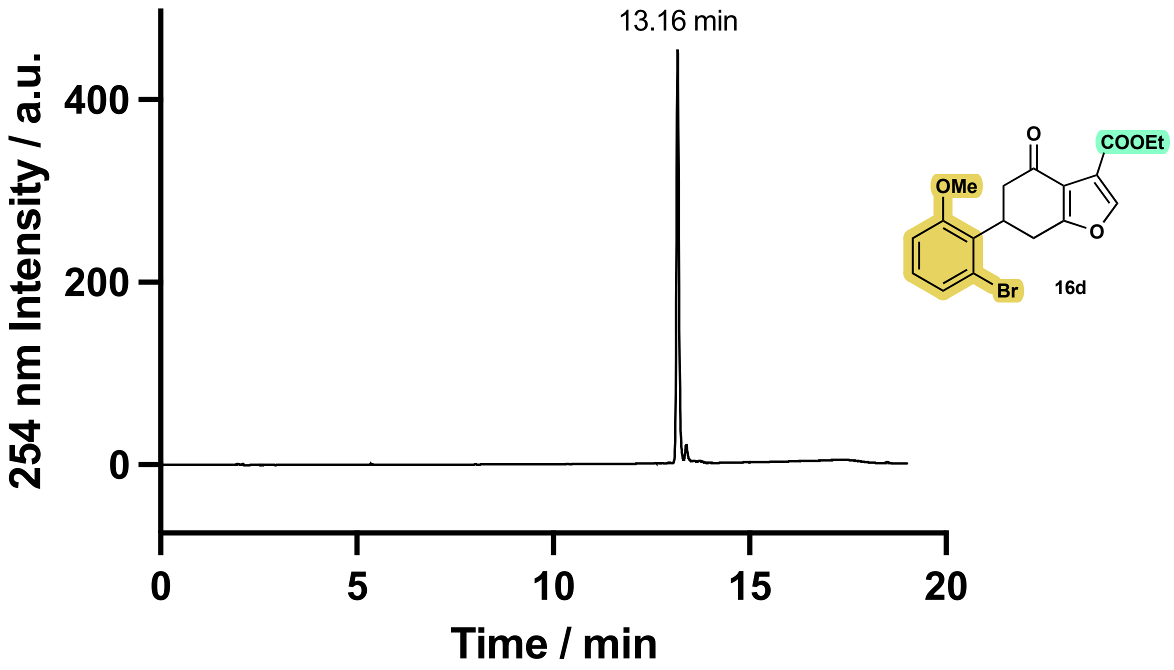

Figure S19 - Compound **16d**, HPLC 5–95 % MeCN in $H_2O$. Purity based on integration of 95 %.

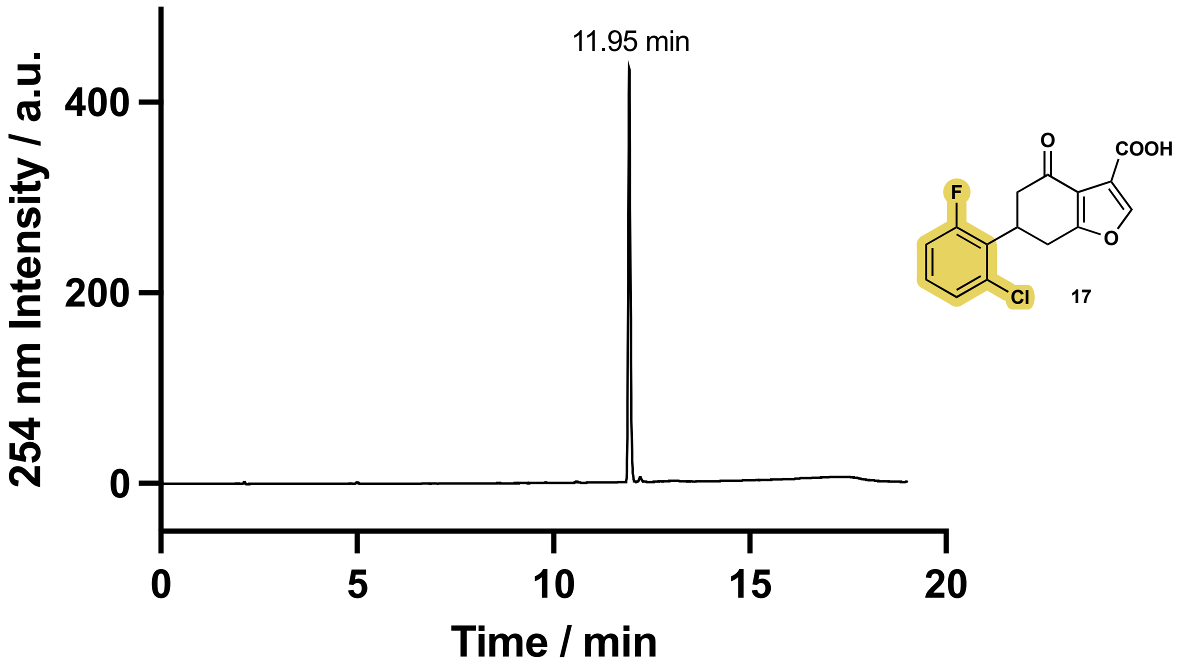

Figure S20 - Compound **17**, HPLC 5–95 % MeCN in $H_2O$. Purity based on integration of 99 %.

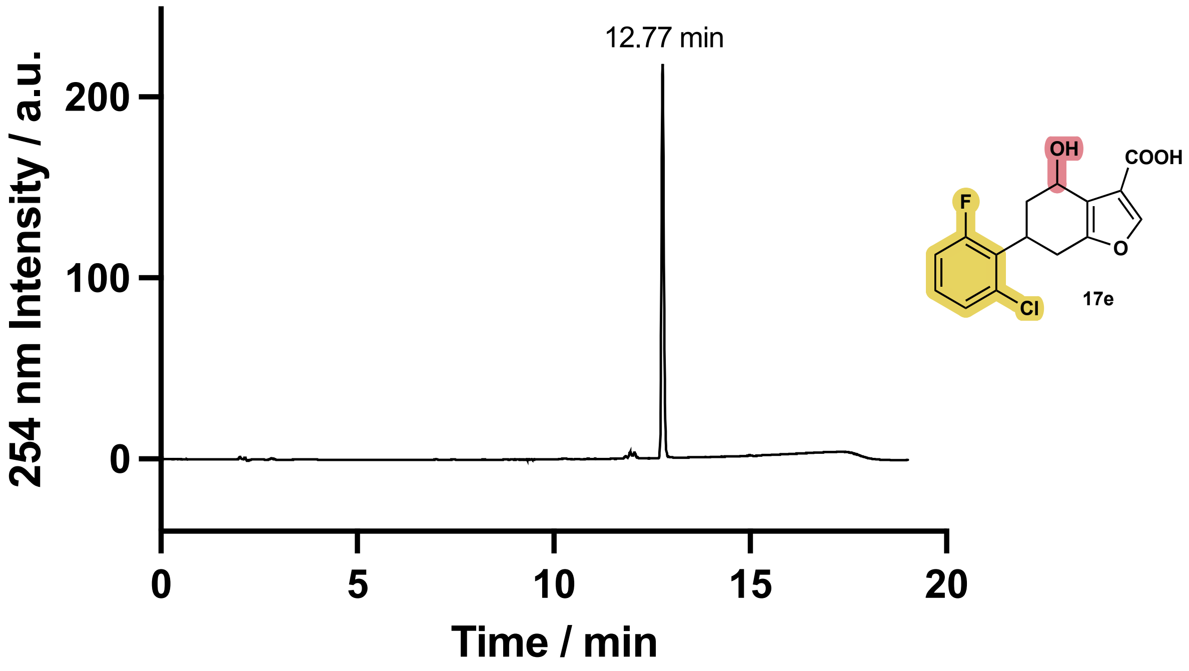

Figure S21 - Compound **17e**, HPLC 5–95 % MeCN in $H_2O$. Purity based on integration of 96 %.

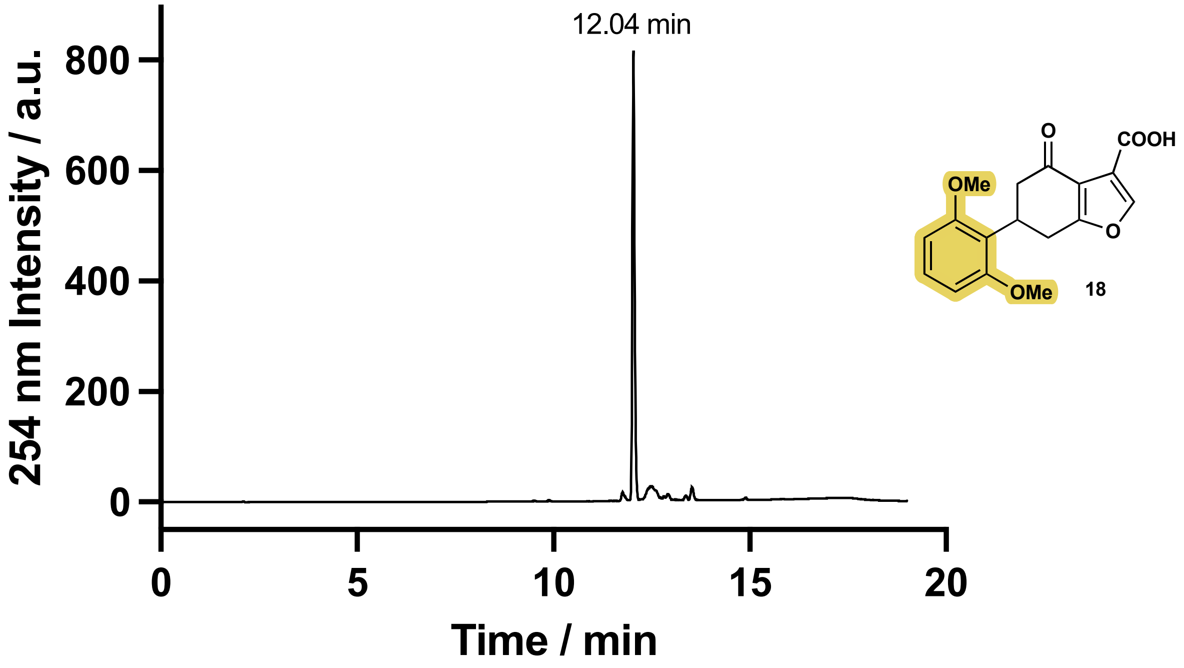

Figure S22 - Compound **18**, HPLC 5–95 % MeCN in $H_2O$. Purity based on integration of 95 %.

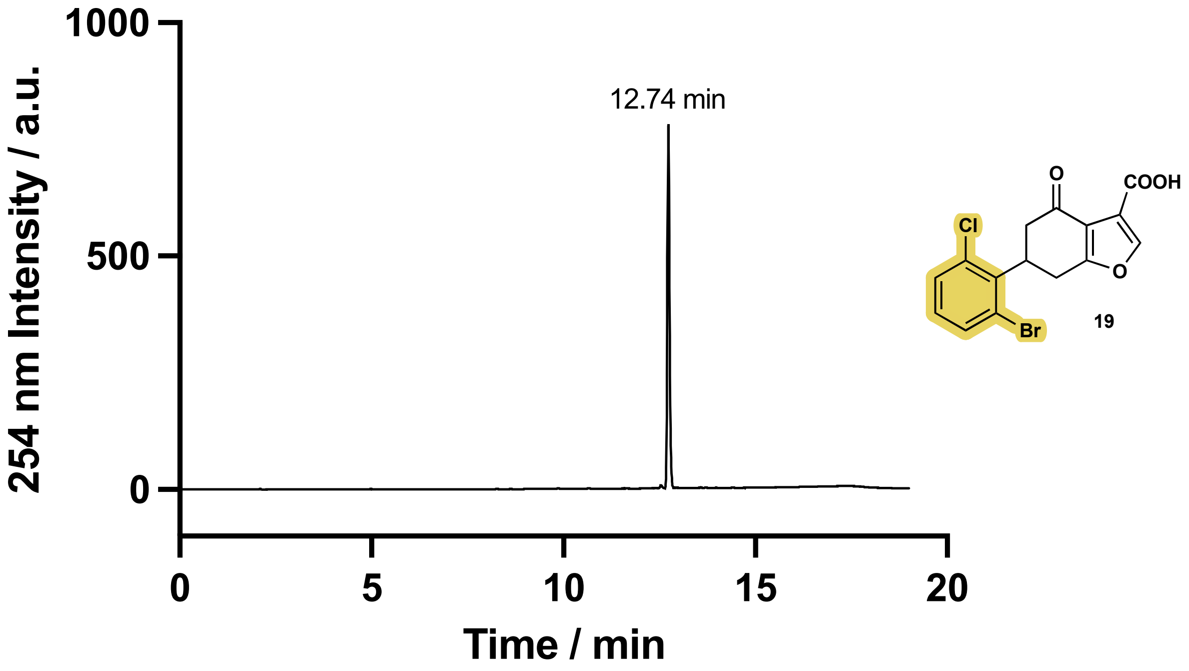

Figure S23 - Compound **19**, HPLC 5–95 % MeCN in $H_2O$. Purity based on integration of 98 %.

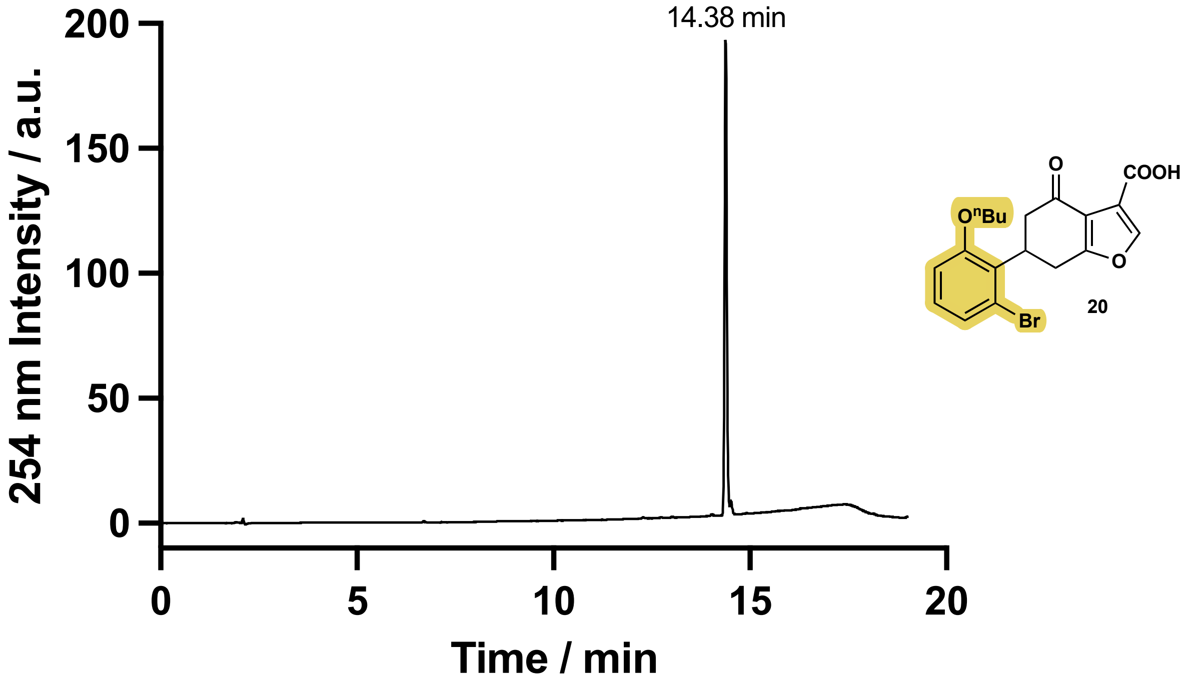

Figure S24 - Compound **20**, HPLC 5–95 % MeCN in $H_2O$. Purity based on integration of 96 %.

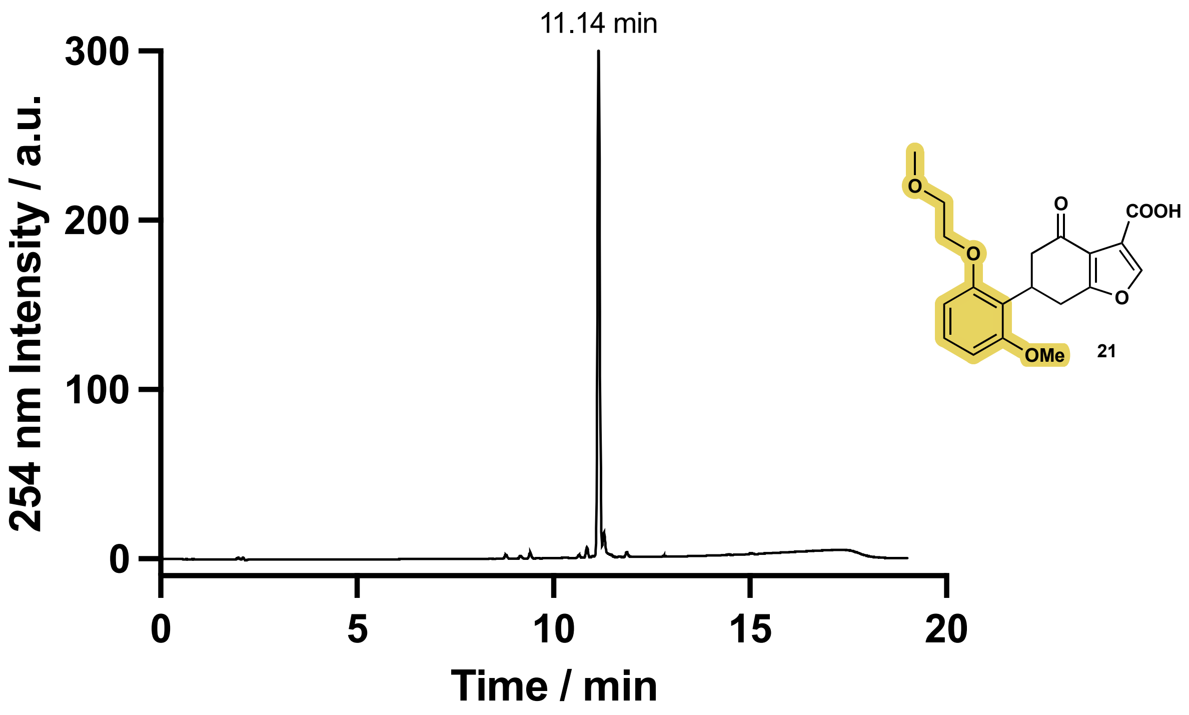

Figure S25 - Compound **21**, HPLC 5–95 % MeCN in $H_2O$. Purity based on integration of 94 %.

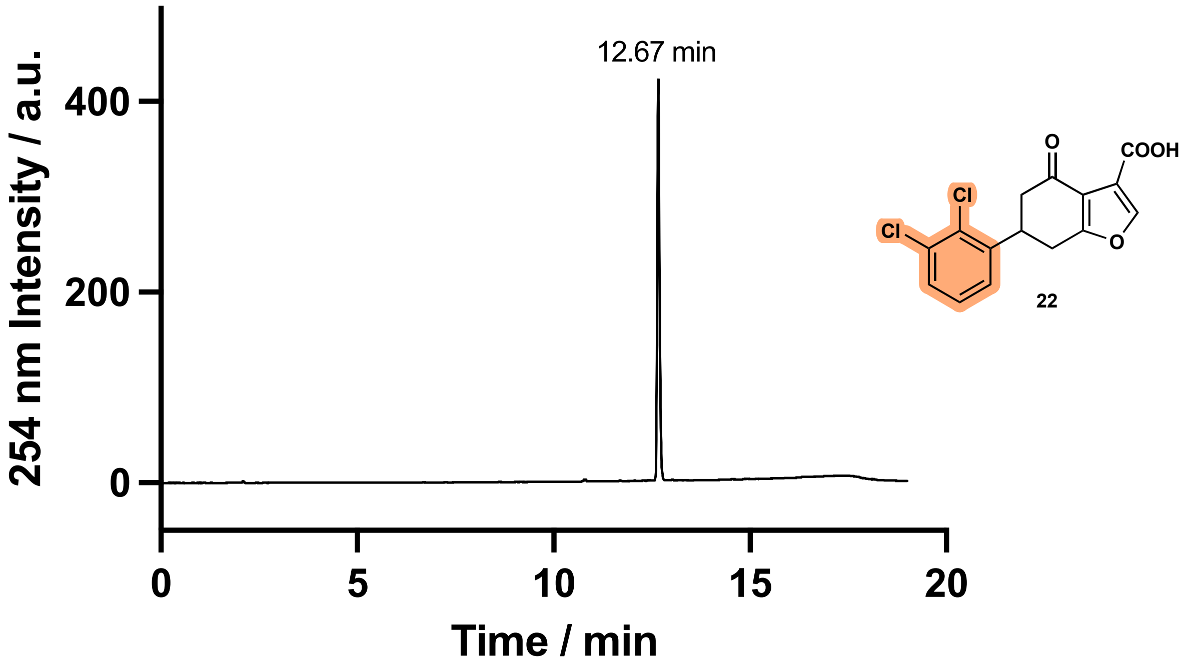

Figure S26 - Compound **22**, HPLC 5–95 % MeCN in $H_2O$. Purity based on integration of 100 %.

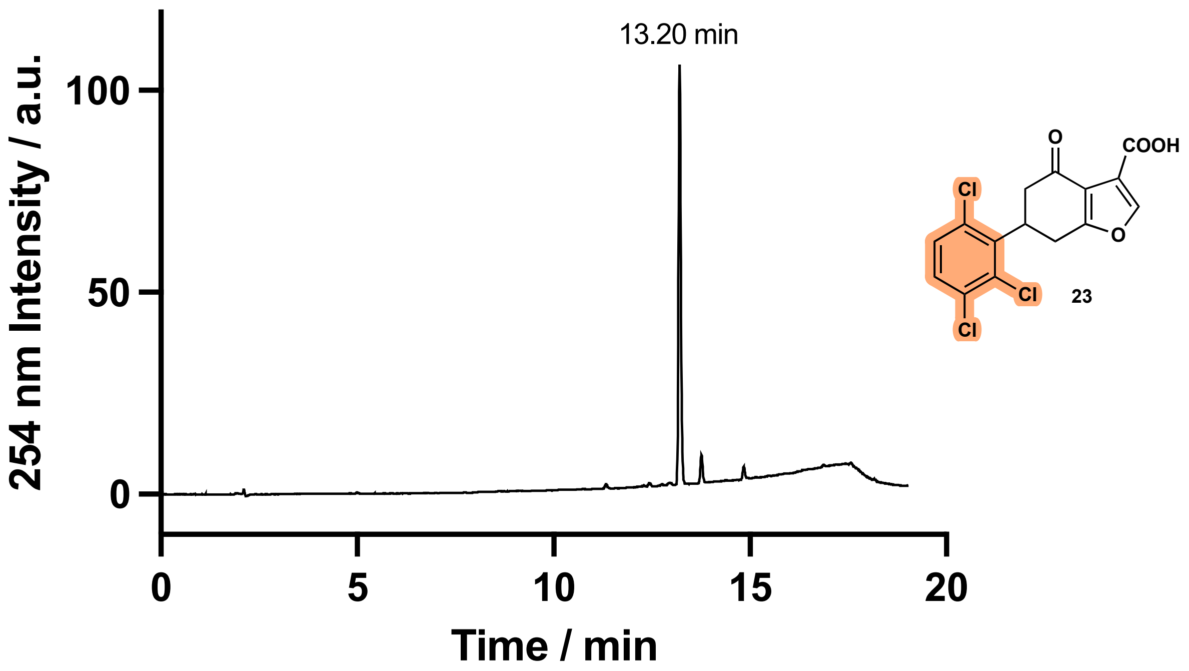

Figure S27 - Compound **23**, HPLC 5–95 % MeCN in $H_2O$. Purity based on integration of 99 %.

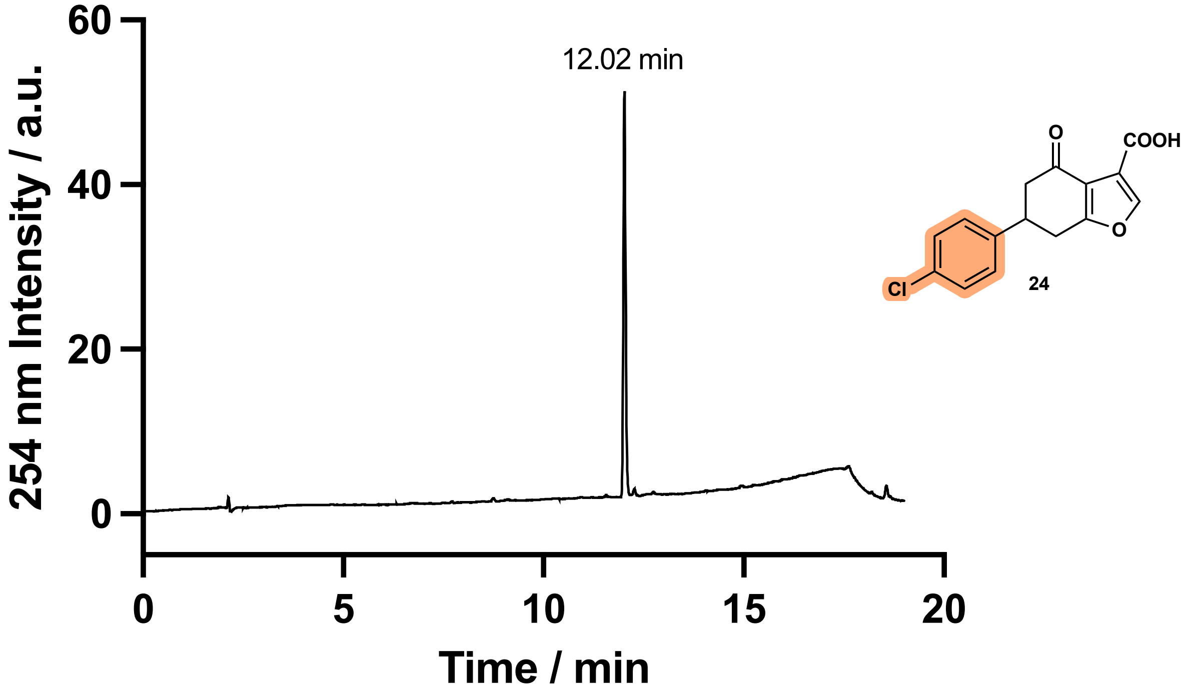

Figure S28 - Compound **24**, HPLC 5–95 % MeCN in $H_2O$. Purity based on integration of 100 %.

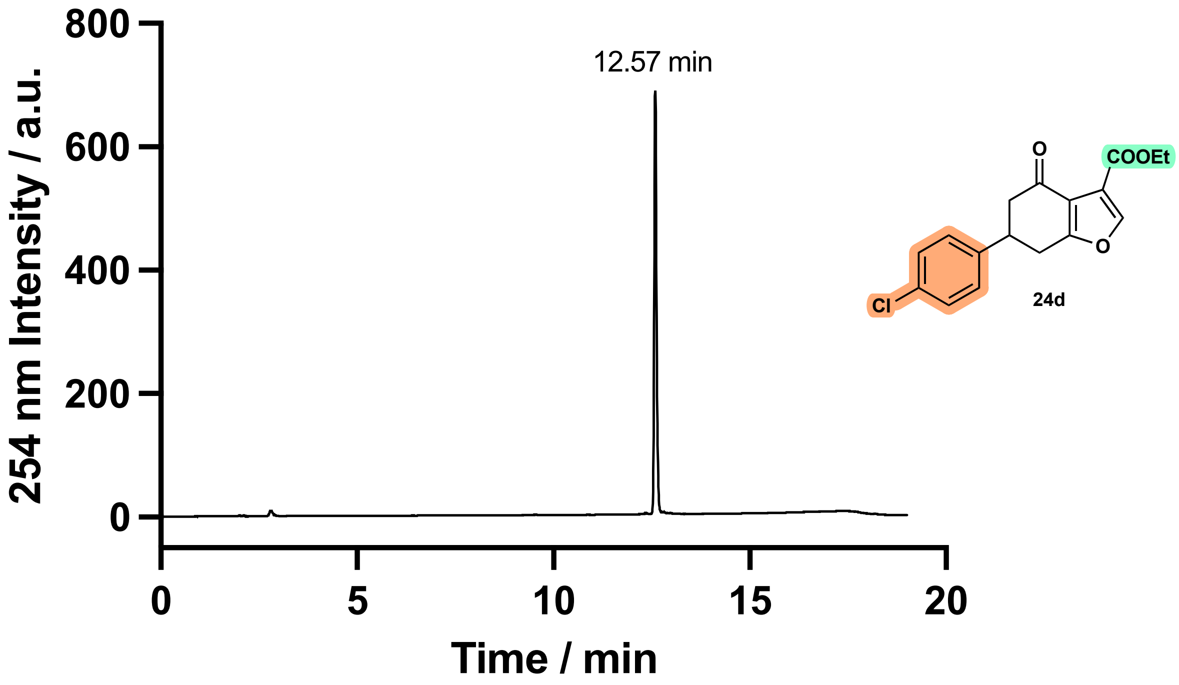

Figure S29 - Compound **24d**, HPLC 5–95 % MeCN in $H_2O$. Purity based on integration of 99 %.

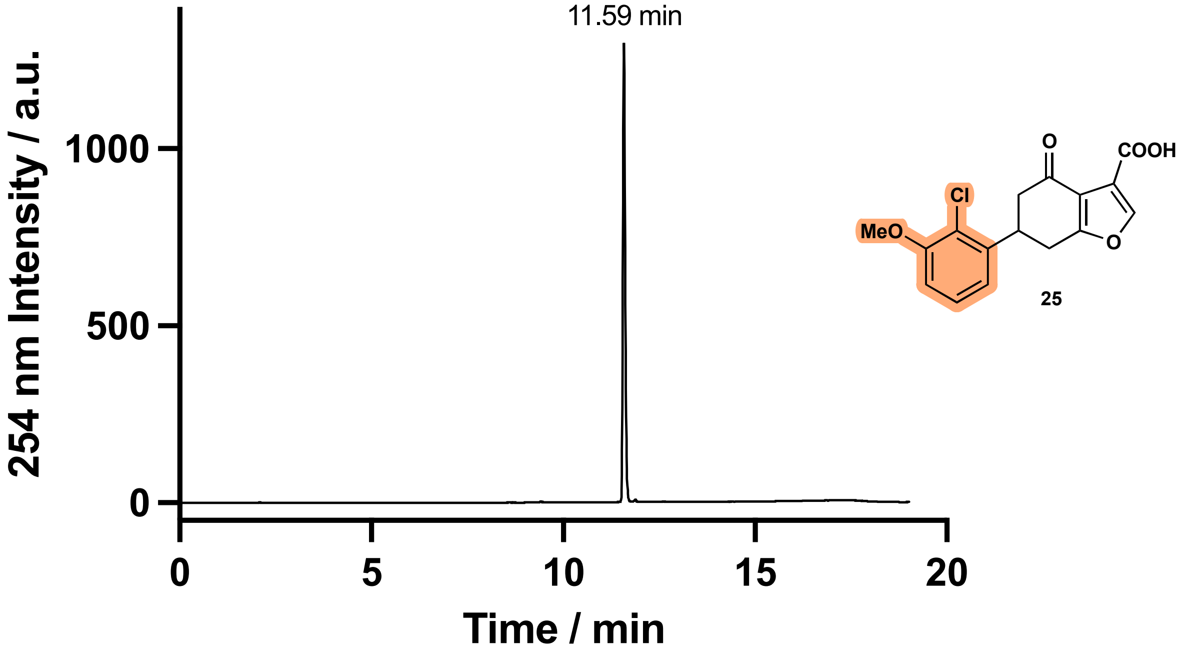

Figure S30 - Compound **25**, HPLC 5–95 % MeCN in $H_2O$. Purity based on integration of 98 %.

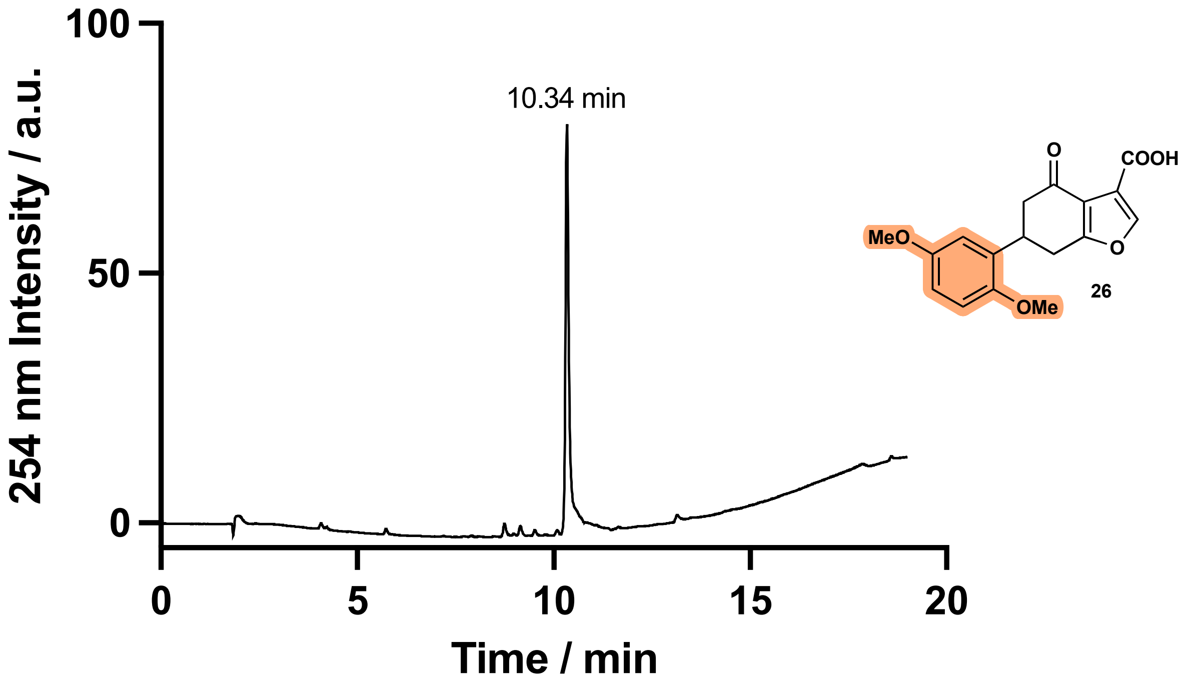

Figure S31 - Compound **26**, HPLC 5–95 % MeCN in $H_2O$. Purity based on integration of 94 %.

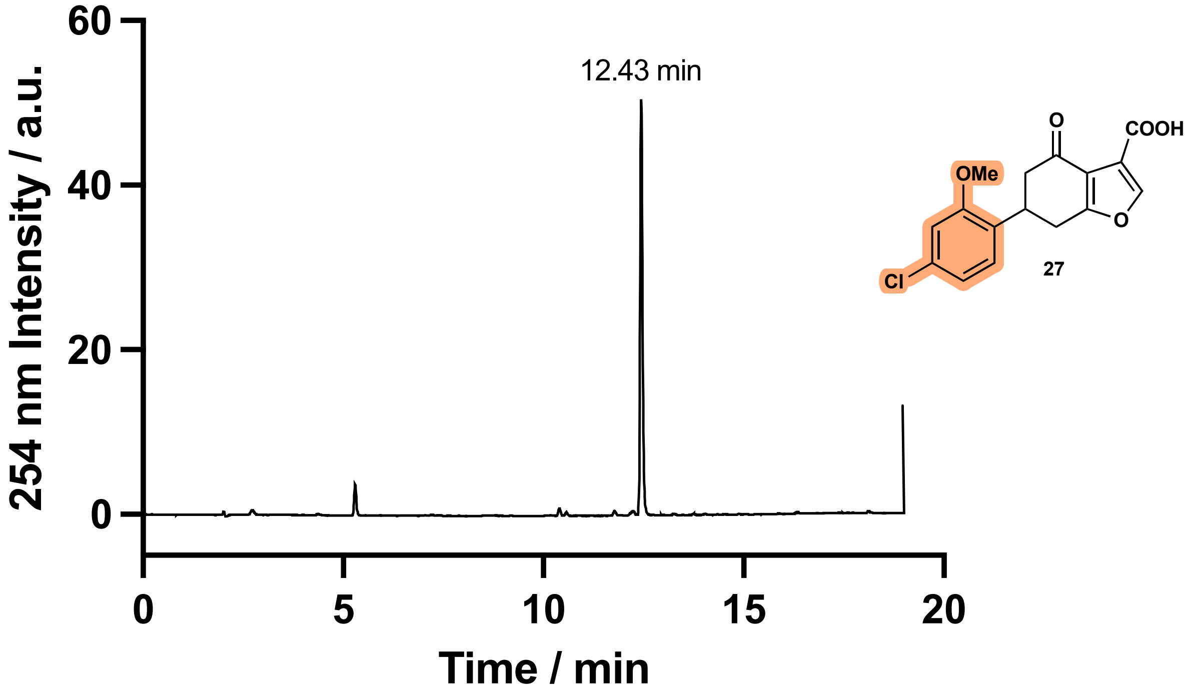

Figure S32 - Compound **27**, HPLC 5–95 % MeCN in $H_2O$. Purity based on integration of 93 %.

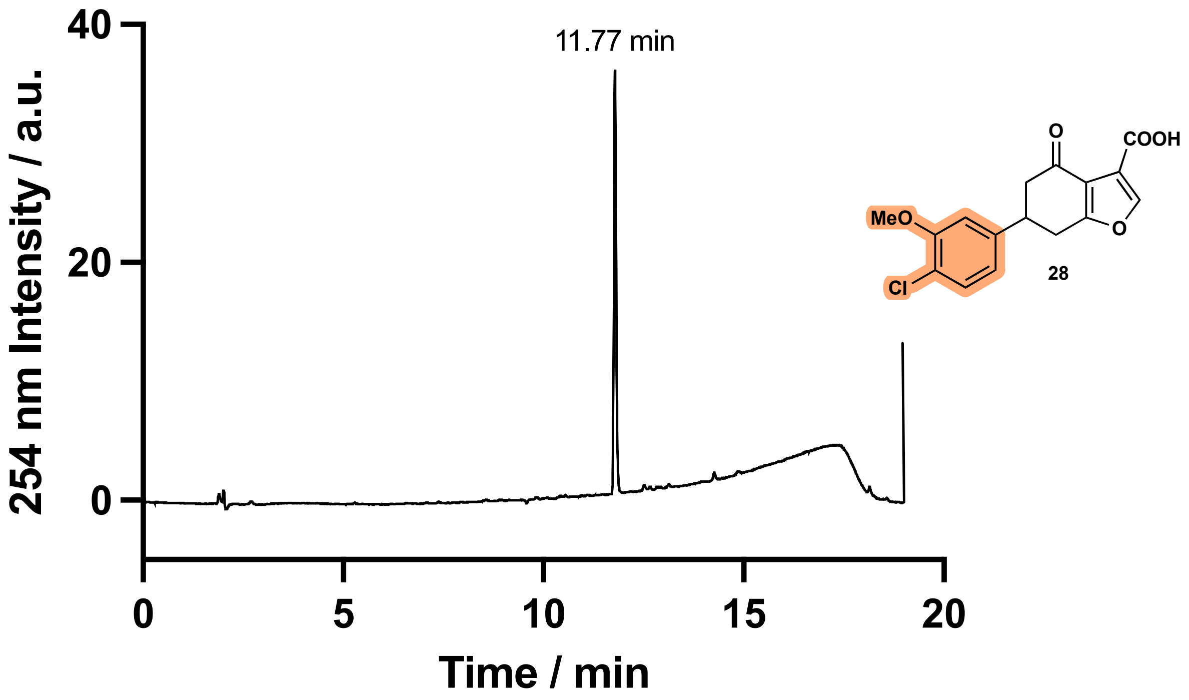

Figure S33 - Compound **28**, HPLC 5–95 % MeCN in $H_2O$. Purity based on integration of 100 %.

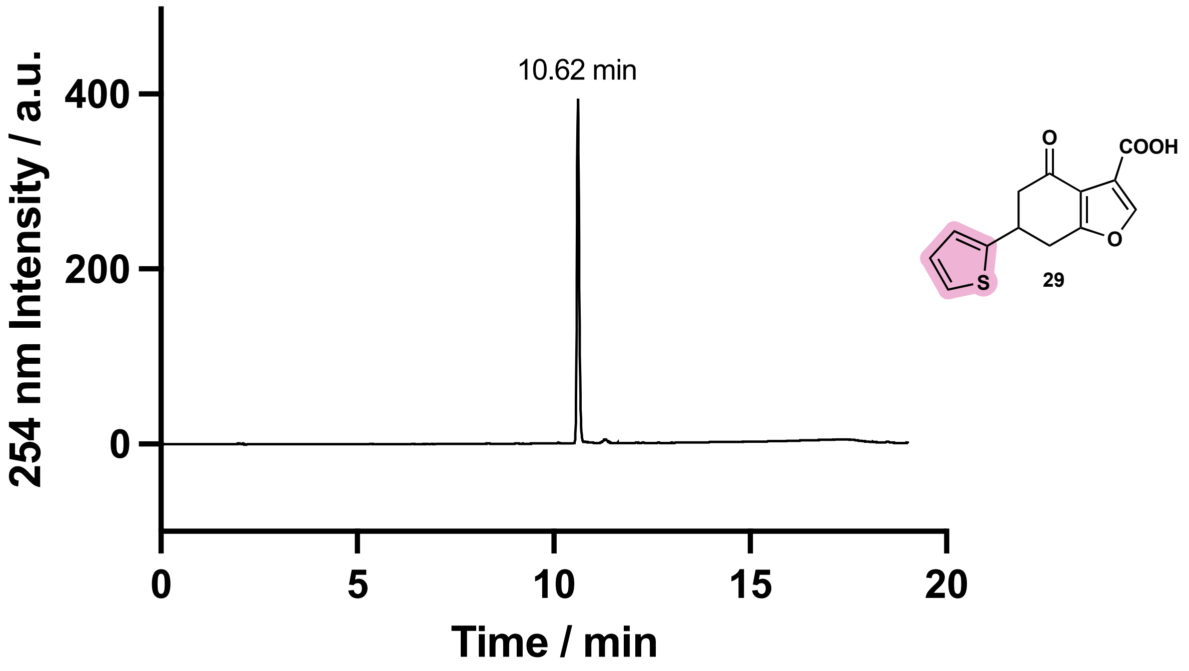

Figure S34 - Compound **29**, HPLC 5–95 % MeCN in $H_2O$. Purity based on integration of 96 %.

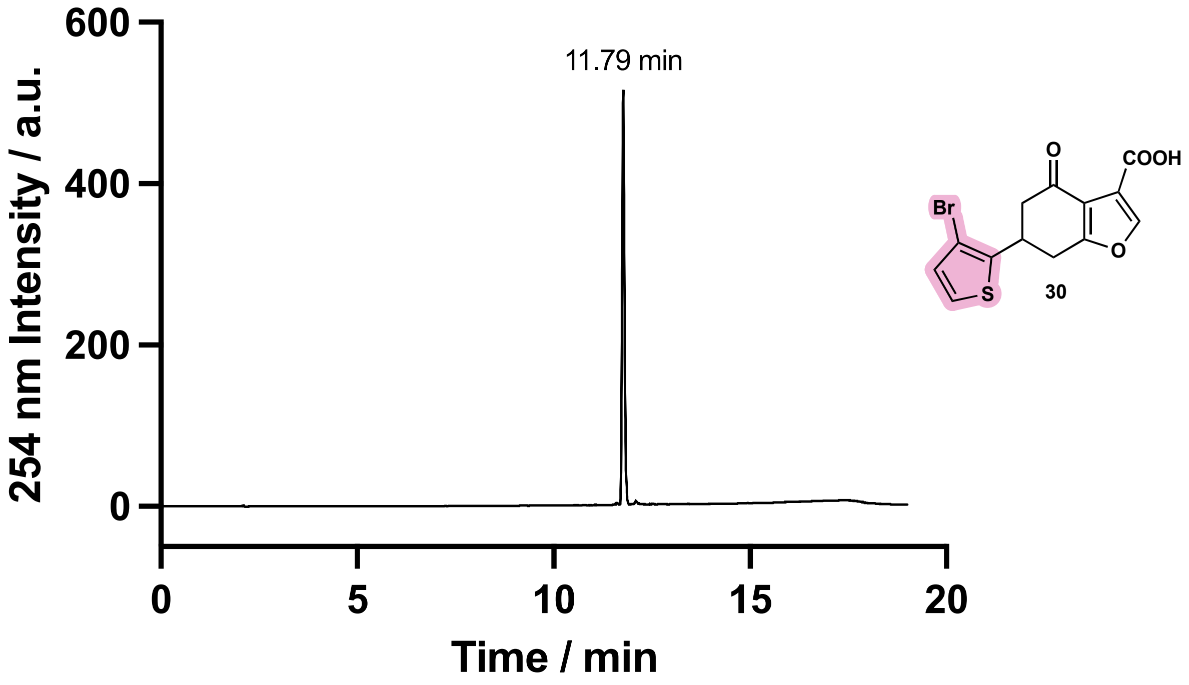

Figure S35 - Compound **30**, HPLC 5–95 % MeCN in $H_2O$. Purity based on integration of 100 %.

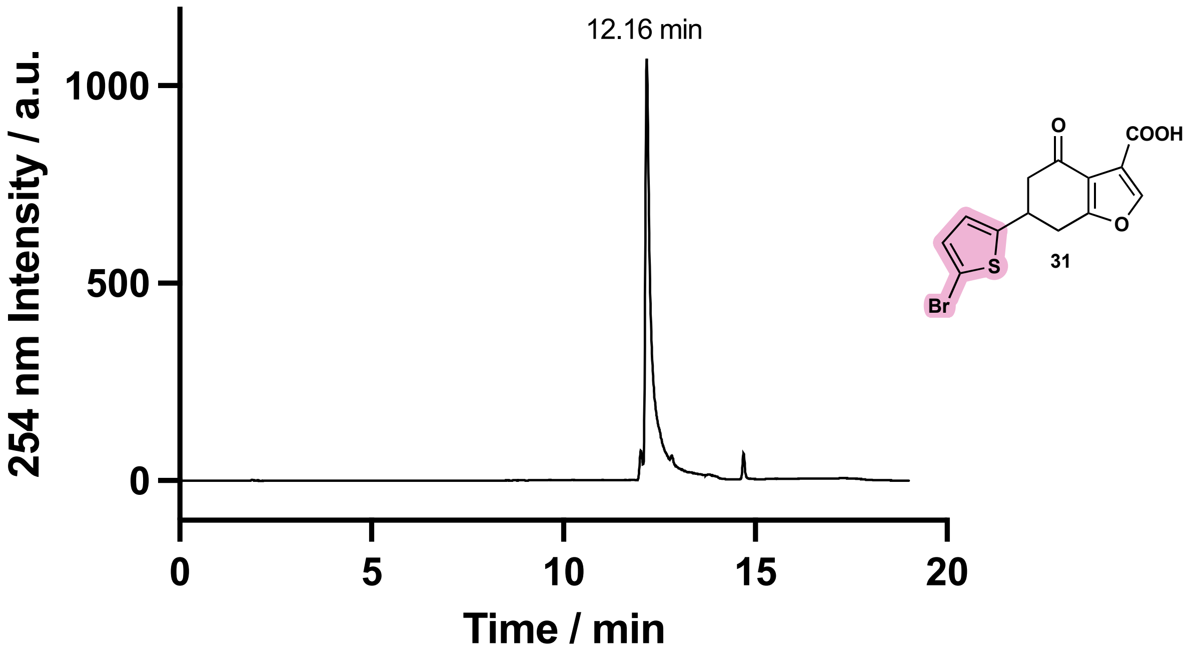

Figure S36 - Compound **31**, HPLC 5–95 % MeCN in $H_2O$. Purity based on integration of 90 %.

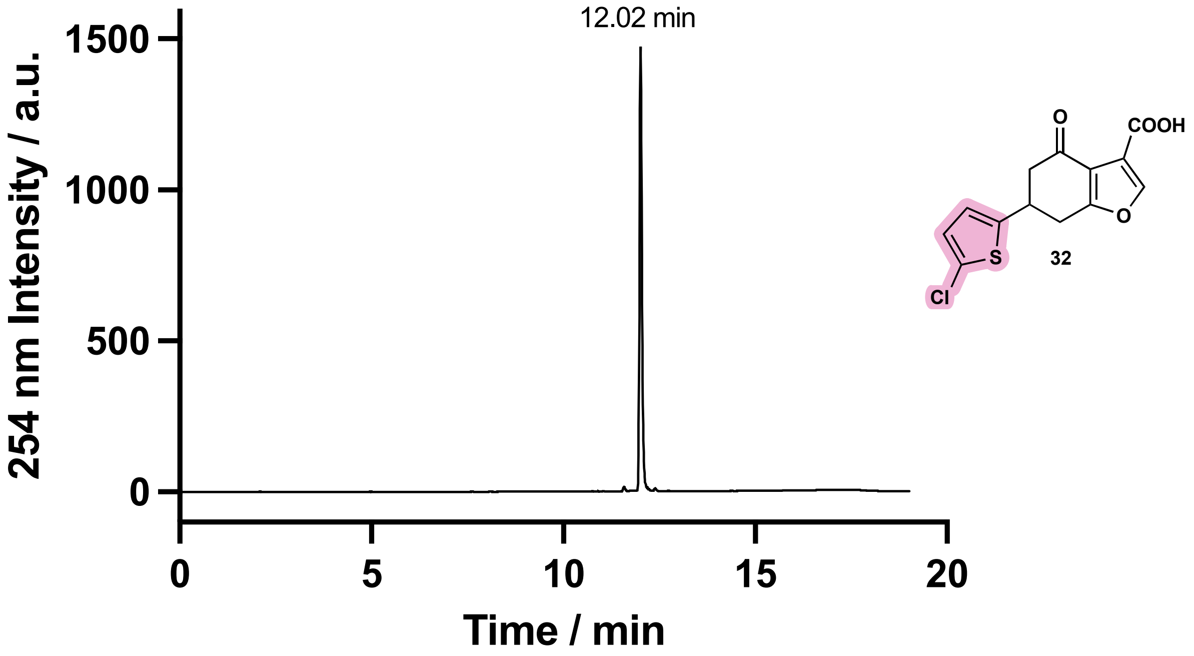

Figure S37 - Compound **32**, HPLC 5–95 % MeCN in $H_2O$. Purity based on integration of 95 %.

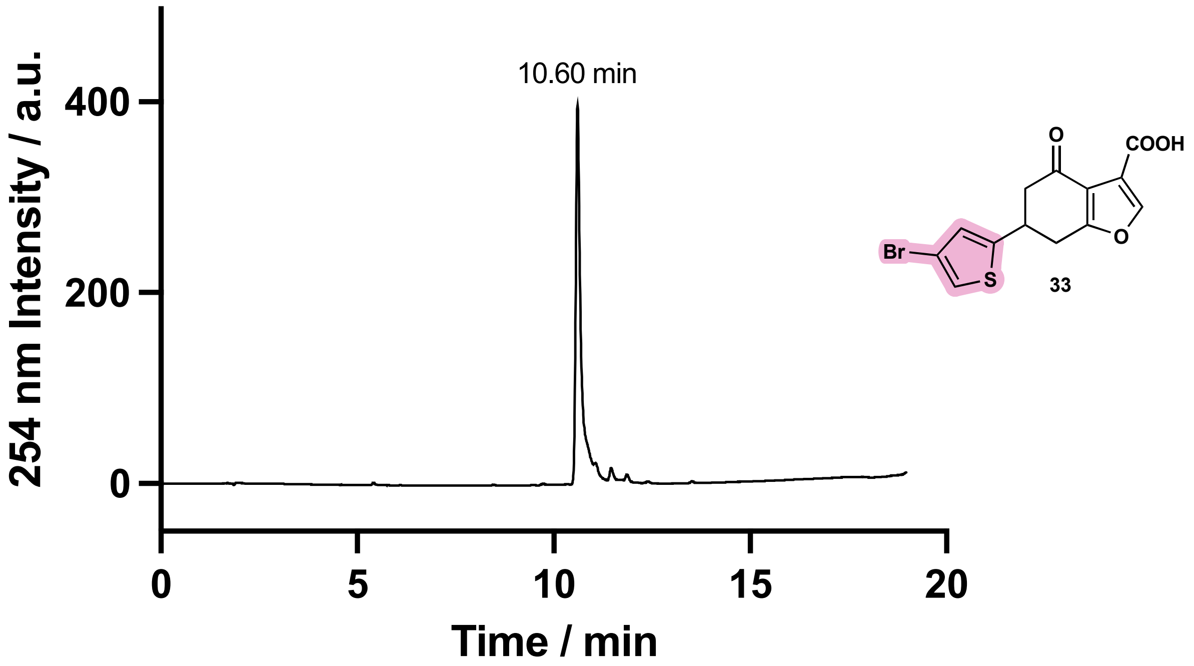

Figure S38 - Compound **33**, HPLC 5–95 % MeCN in $H_2O$. Purity based on integration of 100 %.

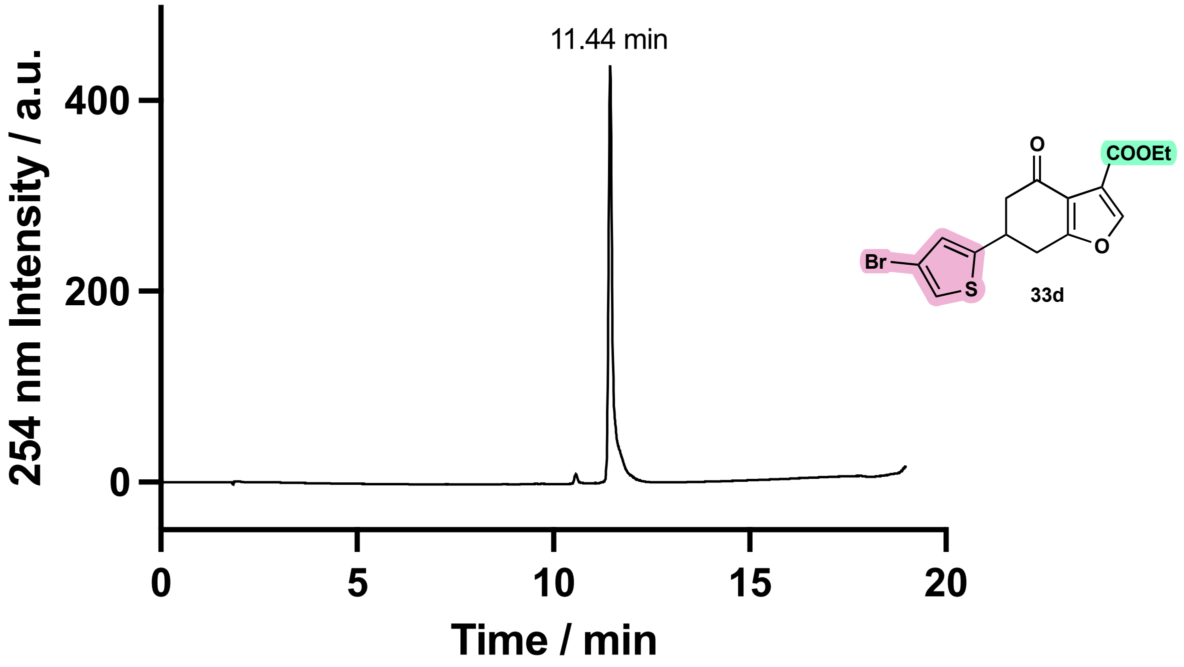

Figure S39 -Compound **33d**, HPLC 5–95 % MeCN in $H_2O$. Purity based on integration of 95 %.

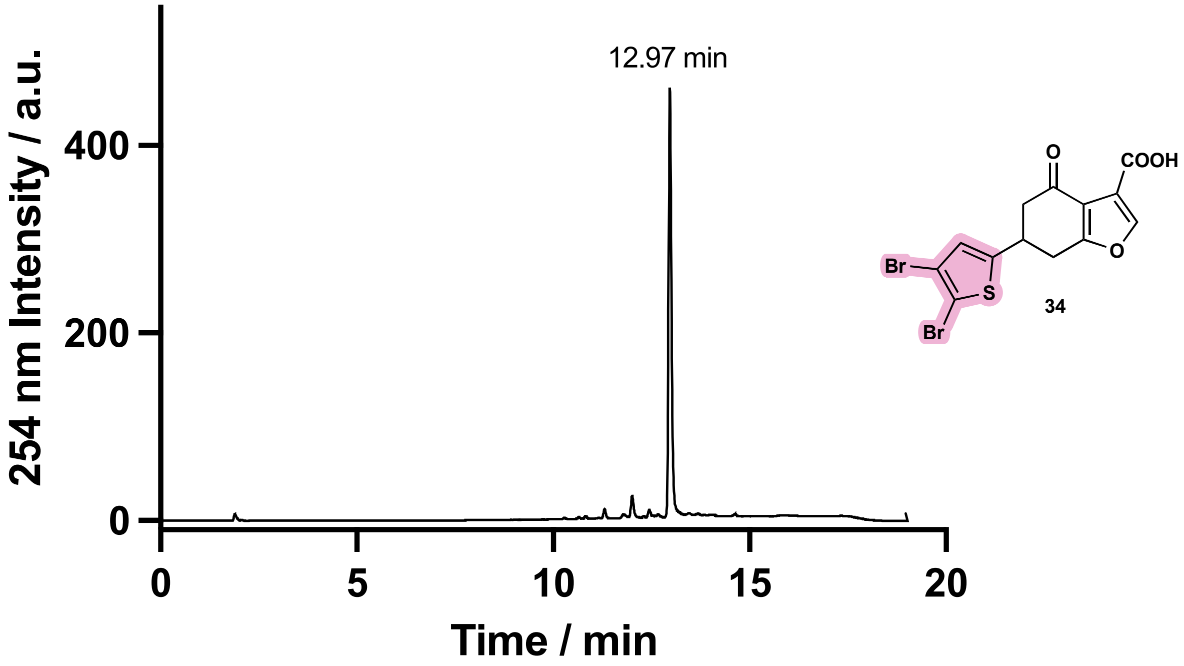

Figure S40 - Compound **34**, HPLC 5–95 % MeCN in $H_2O$. Purity based on integration of 90 %.

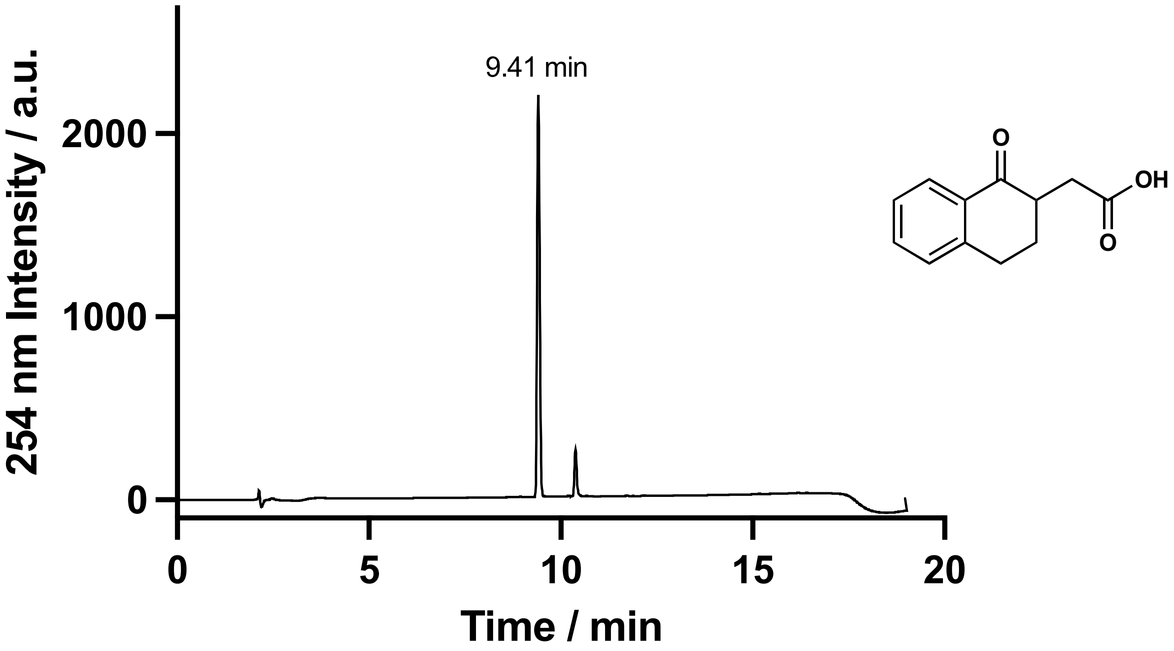

Figure S41 - Compound **37**, HPLC 5–95 % MeCN in $H_2O$. Purity based on integration of 100 %.

### S4 Mass Spectrometry Studies

#### S4.1 Deconvoluted Spectra for Figure 8A and 8B

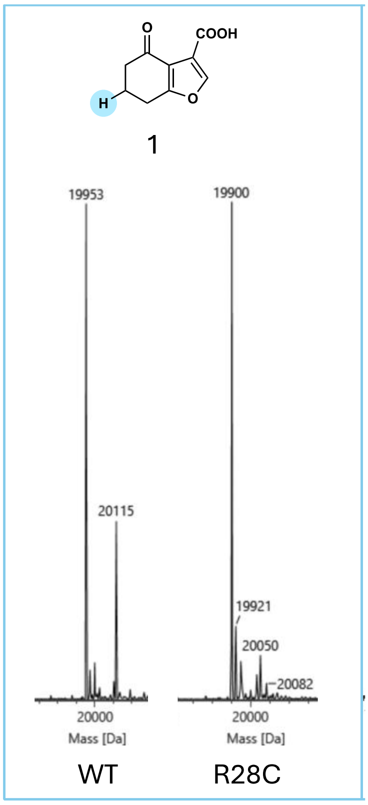

Figure S42 – Mass spectrum of compound **1** (180 Da, 150 μM) incubated at 37 ºC for 5 minutes with WT protein (19953 Da, 2.5 μM) and R28C mutant (19900 Da, 2.5 μM) in pH 8 buffer (20 mM HEPES, TCEP 0.5 mM, NaCl 100 mM). Covalent adduct with WT = 19953 + 180 – 18 = 20115 Da.

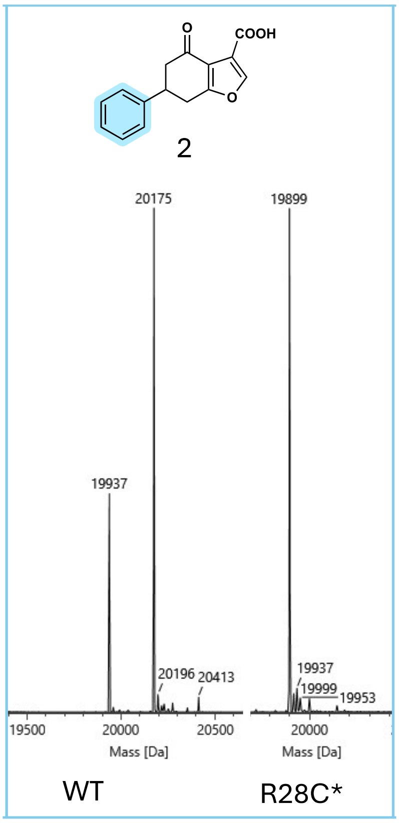

Figure S43 – Mass spectrum of compound **2** (256 Da, 150 µM) incubated at 37 $^{o}$C for 5 minutes with WT protein (19937 Da, 2.5 µM) and R28C mutant (19900 Da, 2.5 µM) in pH 8 buffer (20 mM HEPES, TCEP 0.5 mM, NaCl 100 mM). Covalent adduct with WT = 19937 + 256 – 18 = 20175 Da.

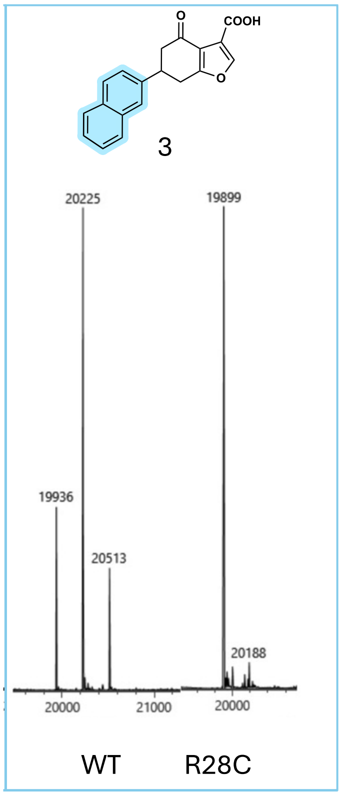

Figure S44 – Mass spectrum of compound **3** (306 Da, 150 µM) incubated at 37 $^{o}$C for 5 minutes with WT protein (19936 Da, 2.5 µM) and R28C mutant (19900 Da, 2.5 µM) in pH 8 buffer (20 mM HEPES, TCEP 0.5 mM, NaCl 100 mM). Covalent adduct with WT = 19936 + 306 – 18 = 20225 Da.

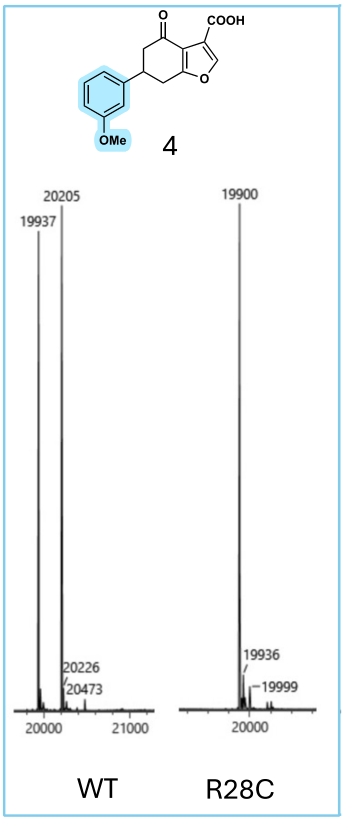

Figure S45 – Mass spectrum of compound **4** (286 Da, 150 µM) incubated at 37 °C for 5 minutes with WT protein (19937 Da, 2.5 µM) and R28C mutant (19900 Da, 2.5 µM) in pH 8 buffer (20 mM HEPES, TCEP 0.5 mM, NaCl 100 mM). Covalent adduct with WT = 19937 + 286 – 18 = 20205 Da.

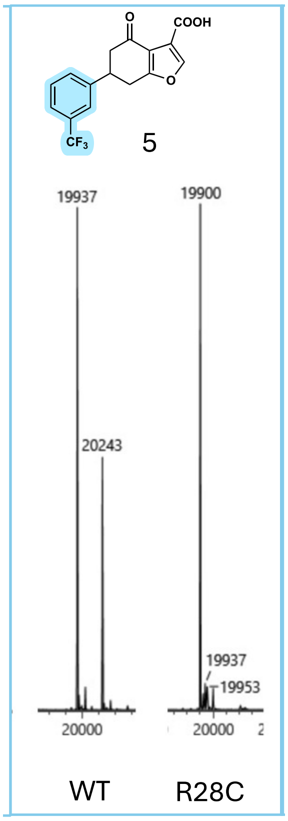

Figure S46 – Mass spectrum of compound **5** (324 Da, 150 µM) incubated at 37 °C for 5 minutes with WT protein (19937 Da, 2.5 µM) and R28C mutant (19900 Da, 2.5 µM) in pH 8 buffer (20 mM HEPES, TCEP 0.5 mM, NaCl 100 mM). Covalent adduct with WT = 19937 + 324 – 18 = 20243 Da.

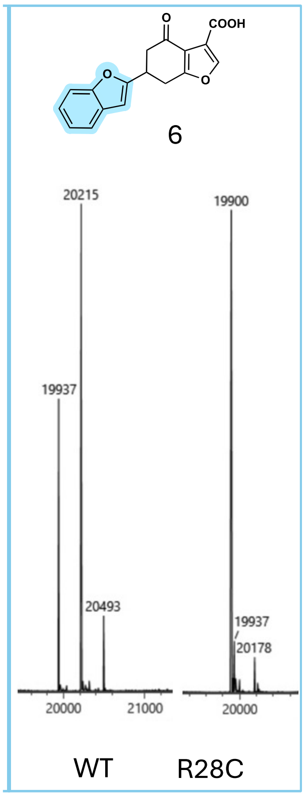

Figure S47 – Mass spectrum of compound **6** (296 Da, 150 µM) incubated at 37 ºC for 5 minutes with WT protein (19937 Da, 2.5 µM) and R28C mutant (19900 Da, 2.5 µM) in pH 8 buffer (20 mM HEPES, TCEP 0.5 mM, NaCl 100 mM). Covalent adduct with WT = 19937 + 296 – 18 = 20215 Da.

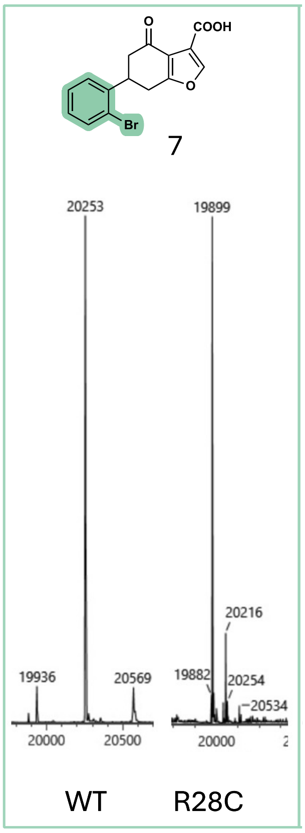

Figure S48 – Mass spectrum of compound **7** (335 Da, 150 µM) incubated at 37 ºC for 5 minutes with WT protein (19936 Da, 2.5 µM) and R28C mutant (19900 Da, 2.5 µM) in pH 8 buffer (20 mM HEPES, TCEP 0.5 mM, NaCl 100 mM). Covalent adduct with WT = 19936 + 335 – 18 = 20253 Da.

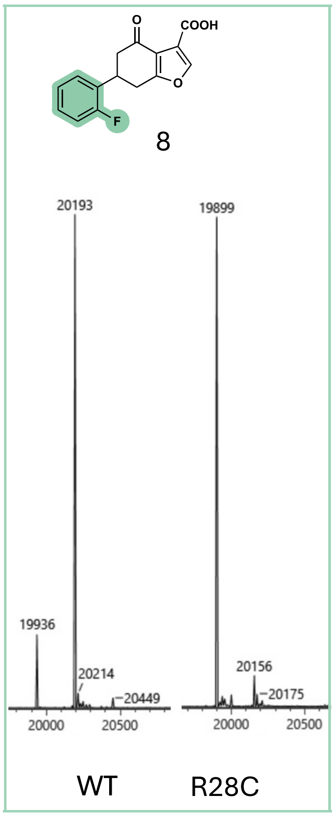

Figure S49 – Mass spectrum of compound **8** (274 Da, 150 µM) incubated at 37 ºC for 5 minutes with WT protein (19936 Da, 2.5 µM) and R28C mutant (19900 Da, 2.5 µM) in pH 8 buffer (20 mM HEPES, TCEP 0.5 mM, NaCl 100 mM). Covalent adduct with WT = 19936 + 274 – 18 = 20192 Da.

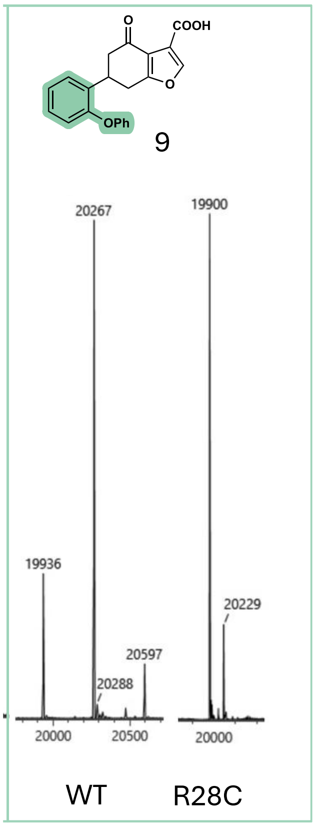

Figure S50 – Mass spectrum of compound **9** (348 Da, 150 µM) incubated at 37 ºC for 5 minutes with WT protein (19936 Da, 2.5 µM) and R28C mutant (19900 Da, 2.5 µM) in pH 8 buffer (20 mM HEPES, TCEP 0.5 mM, NaCl 100 mM). Covalent adduct with WT = 19936 + 348 – 18 = 20266 Da.

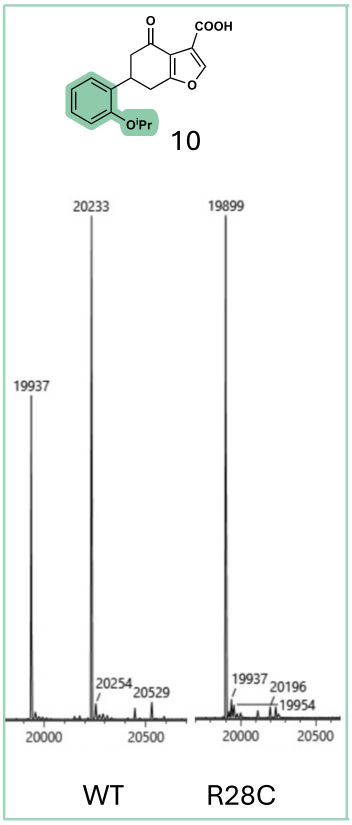

Figure S51 – Mass spectrum of compound **10** (314 Da, 150 μM) incubated at 37 ºC for 5 minutes with WT protein (19937 Da, 2.5 μM) and R28C mutant (19899 Da, 2.5 μM) in pH 8 buffer (20 mM HEPES, TCEP 0.5 mM, NaCl 100 mM). Covalent adduct with WT = 19937 + 314 – 18 = 20233 Da.

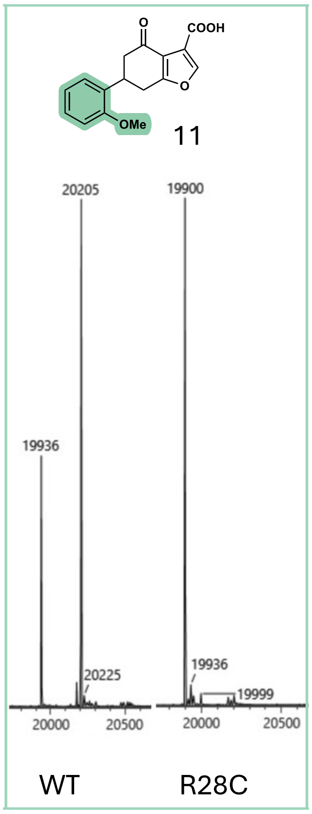

Figure S52 – Mass spectrum of compound **11** (286 Da, 150 μM) incubated at 37 ºC for 5 minutes with WT protein (19936 Da, 2.5 μM) and R28C mutant (19900 Da, 2.5 μM) in pH 8 buffer (20 mM HEPES, TCEP 0.5 mM, NaCl 100 mM). Covalent adduct with WT = 19936 + 286 – 18 = 20204 Da.

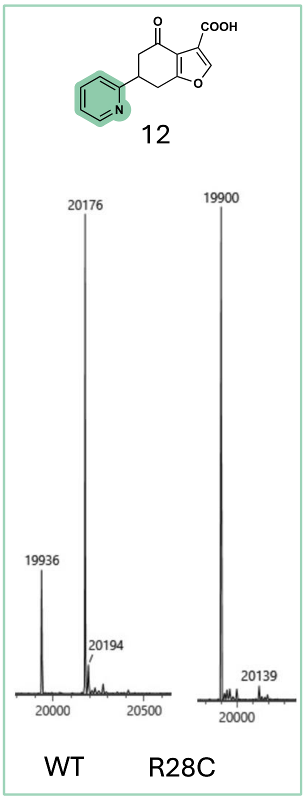

Figure S53 – Mass spectrum of compound **12** (257 Da, 150 μM) incubated at 37 °C for 5 minutes with WT protein (19936 Da, 2.5 μM) and R28C mutant (19900 Da, 2.5 μM) in pH 8 buffer (20 mM HEPES, TCEP 0.5 mM, NaCl 100 mM). Covalent adduct with WT = 19936 + 257 – 18 = 20175 Da.

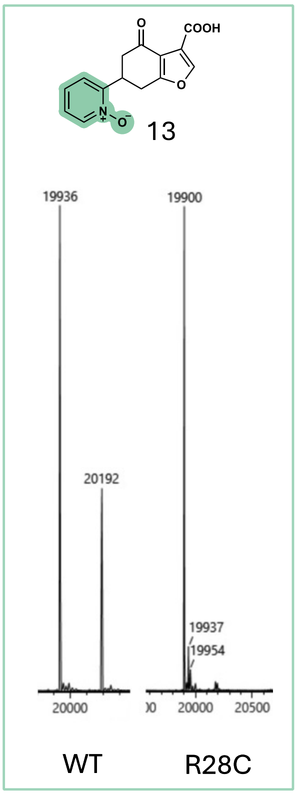

Figure S54 – Mass spectrum of compound **13** (273 Da, 150 μM) incubated at 37 °C for 5 minutes with WT protein (19936 Da, 2.5 μM) and R28C mutant (19900 Da, 2.5 μM) in pH 8 buffer (20 mM HEPES, TCEP 0.5 mM, NaCl 100 mM). Covalent adduct with WT = 19936 + 273 – 18 = 20191 Da.

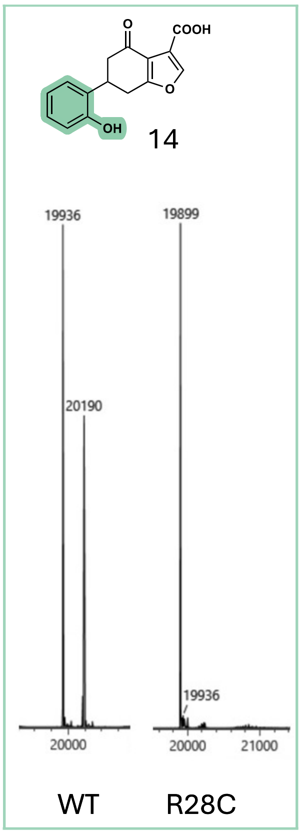

Figure S55 – Mass spectrum of compound **14** (272 Da, 150 μM) incubated at 37 ºC for 5 minutes with WT protein (19936 Da, 2.5 μM) and R28C mutant (19900 Da, 2.5 μM) in pH 8 buffer (20 mM HEPES, TCEP 0.5 mM, NaCl 100 mM). Covalent adduct with WT = 19936 + 272 – 18 = 20190 Da.

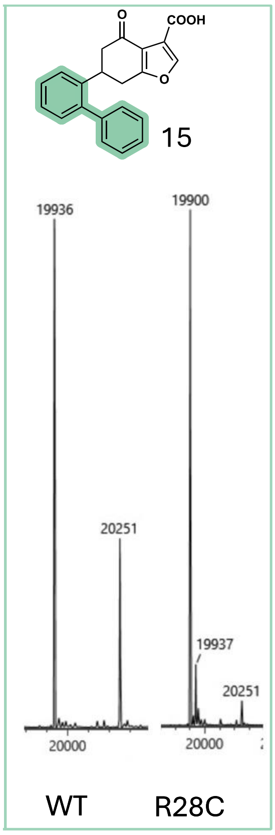

Figure S56 – Mass spectrum of compound **15** (332 Da, 150 μM) incubated at 37 ºC for 5 minutes with WT protein (19936 Da, 2.5 μM) and R28C mutant (19900 Da, 2.5 μM) in pH 8 buffer (20 mM HEPES, TCEP 0.5 mM, NaCl 100 mM). Covalent adduct with WT = 19936 + 332 – 18 = 20250 Da.

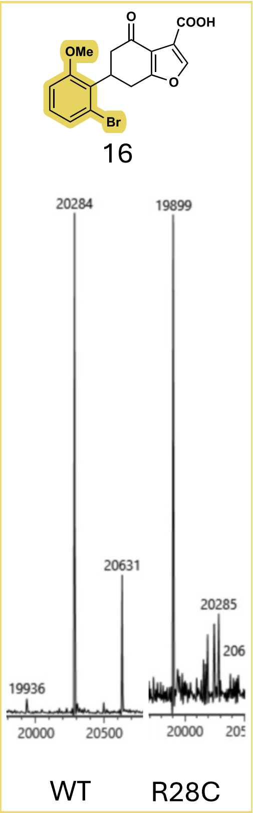

Figure S57 – Mass spectrum of compound **16** (365 Da, 150 µM) incubated at 37 ºC for 5 minutes with WT protein (19936 Da, 2.5 µM) and R28C mutant (19900 Da, 2.5 µM) in pH 8 buffer (20 mM HEPES, TCEP 0.5 mM, NaCl 100 mM). Covalent adduct with WT = 19936 + 365 – 18 = 20283 Da.

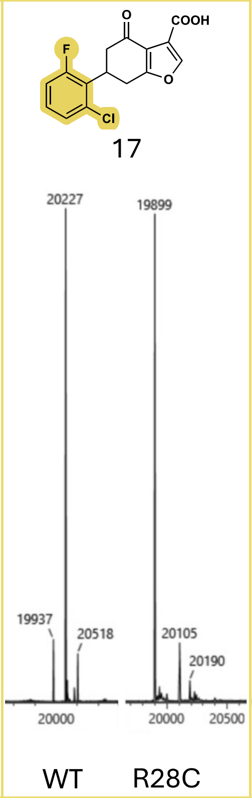

Figure S58 – Mass spectrum of compound **17** (309 Da, 150 µM) incubated at 37 ºC for 5 minutes with WT protein (19937 Da, 2.5 µM) and R28C mutant (19900 Da, 2.5 µM) in pH 8 buffer (20 mM HEPES, TCEP 0.5 mM, NaCl 100 mM). Covalent adduct with WT = 19937 + 309 – 18 = 20228 Da.

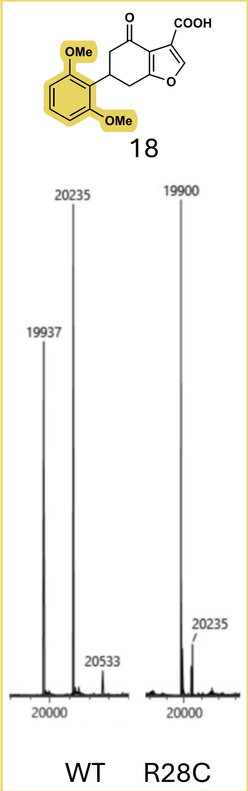

Figure S59 – Mass spectrum of compound **18** (316 Da, 150 µM) incubated at 37 °C for 5 minutes with WT protein (19937 Da, 2.5 µM) and R28C mutant (19900 Da, 2.5 µM) in pH 8 buffer (20 mM HEPES, TCEP 0.5 mM, NaCl 100 mM). Covalent adduct with WT = 19937 + 316 – 18 = 20235 Da.

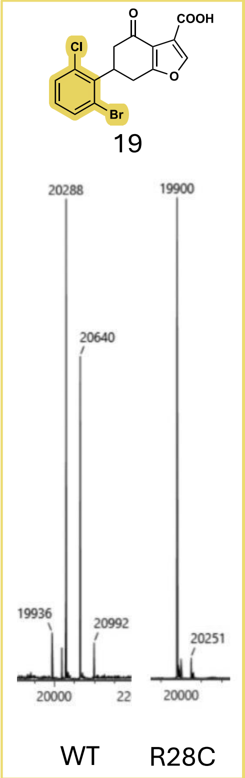

Figure S60 – Mass spectrum of compound **19** (370 Da, 150 µM) incubated at 37 °C for 5 minutes with WT protein (19936 Da, 2.5 µM) and R28C mutant (19900 Da, 2.5 µM) in pH 8 buffer (20 mM HEPES, TCEP 0.5 mM, NaCl 100 mM). Covalent adduct with WT = 19936 + 370 – 18 = 20288 Da.

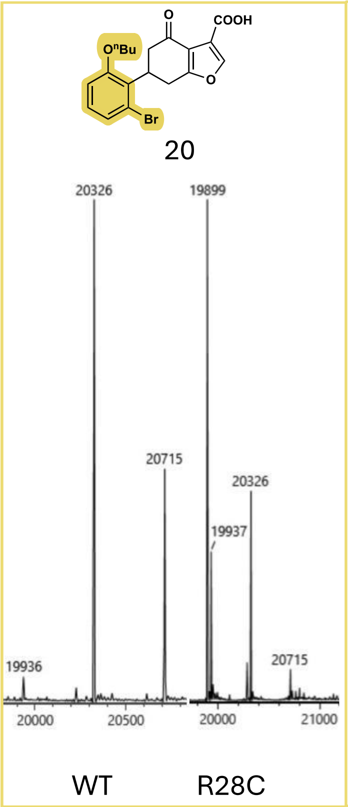

Figure S61 – Mass spectrum of compound **20** (407 Da, 150 μM) incubated at 37 ºC for 5 minutes with WT protein (19936 Da, 2.5 μM) and R28C mutant (19900 Da, 2.5 μM) in pH 8 buffer (20 mM HEPES, TCEP 0.5 mM, NaCl 100 mM). Covalent adduct with WT = 19936 + 407 – 18 = 20325 Da.

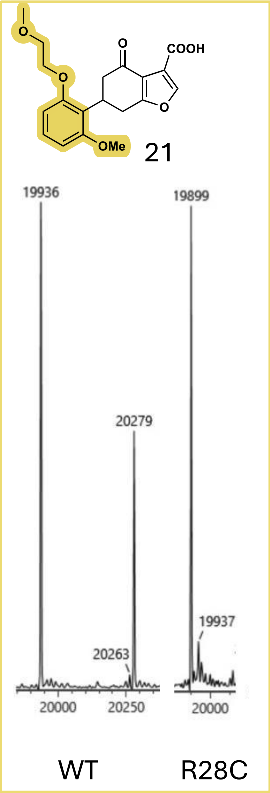

Figure S62 – Mass spectrum of compound **21** (360 Da, 150 μM) incubated at 37 ºC for 5 minutes with WT protein (19936 Da, 2.5 μM) and R28C mutant (19900 Da, 2.5 μM) in pH 8 buffer (20 mM HEPES, TCEP 0.5 mM, NaCl 100 mM). Covalent adduct with WT = 19936 + 360 – 18 = 20278 Da.

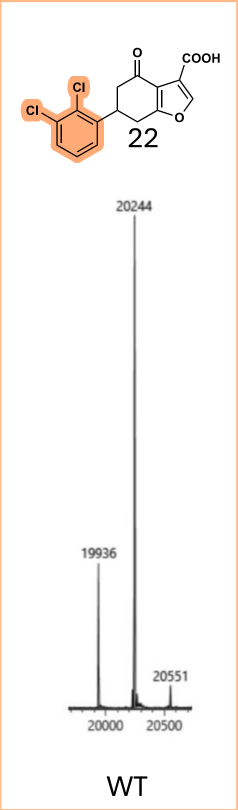

Figure S63 – Mass spectrum of compound **22** (325 Da, 150 µM) incubated at 37 ºC for 5 minutes with WT protein (19936 Da, 2.5 µM) in pH 8 buffer (20 mM HEPES, TCEP 0.5 mM, NaCl 100 mM). Covalent adduct with WT = 19936 + 325 – 18 = 20243 Da.

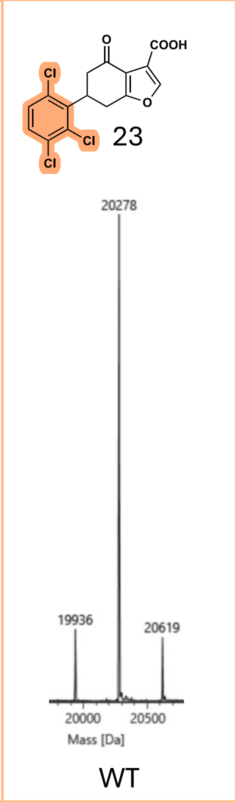

Figure S64 – Mass spectrum of compound **23** (360 Da, 150 µM) incubated at 37 ºC for 5 minutes with WT protein (19936 Da, 2.5 µM) in pH 8 buffer (20 mM HEPES, TCEP 0.5 mM, NaCl 100 mM). Covalent adduct with WT = 19936 + 360 – 18 = 20278 Da.

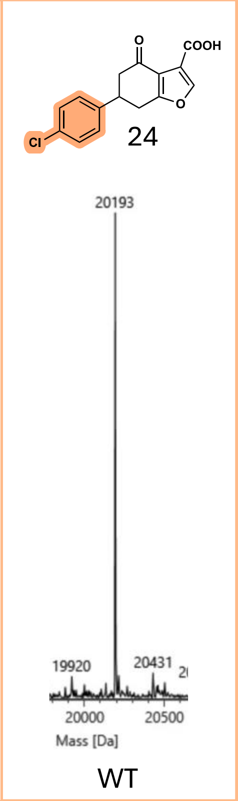

Figure S65 – Mass spectrum of compound **24** (291 Da, 150 μM) incubated at 37 ºC for 5 minutes with Ala mutant, binding unaffected, of the WT protein (19920 Da, 2.5 μM) in pH 8 buffer (20 mM HEPES, TCEP 0.5 mM, NaCl 100 mM). Covalent adduct with WT = 19920 + 291 – 18 = 20193 Da.

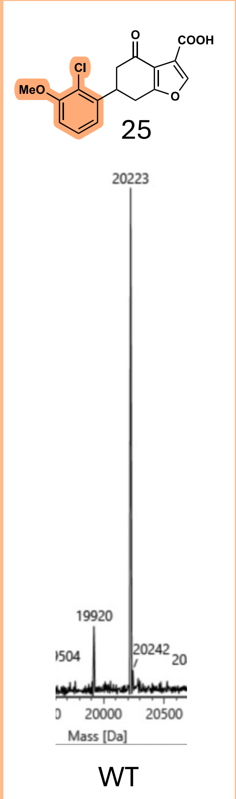

Figure S66 – Mass spectrum of compound **25** (321 Da, 150 μM) incubated at 37 ºC for 5 minutes with Ala mutant, binding unaffected, of the WT protein (19920 Da, 2.5 μM) in pH 8 buffer (20 mM HEPES, TCEP 0.5 mM, NaCl 100 mM). Covalent adduct with WT = 19920 + 321 – 18 = 20223 Da.

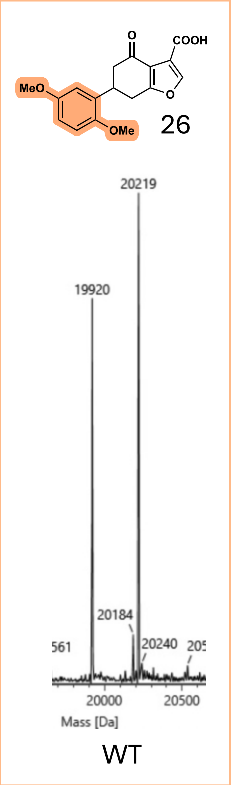

Figure S67 – Mass spectrum of compound **26** (316 Da, 150 μM) incubated at 37 ºC for 5 minutes with Ala mutant, binding unaffected, of the WT protein (19920 Da, 2.5 μM) in pH 8 buffer (20 mM HEPES, TCEP 0.5 mM, NaCl 100 mM). Covalent adduct with WT = 19920 + 316 – 18 = 20218 Da.

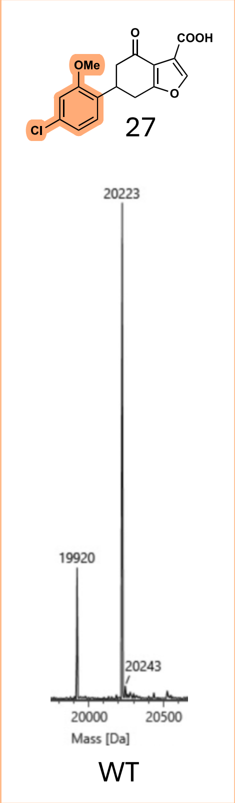

Figure S68 – Mass spectrum of compound **27** (321 Da, 150 μM) incubated at 37 ºC for 5 minutes with Ala mutant, binding unaffected, of the WT protein (19920 Da, 2.5 μM) in pH 8 buffer (20 mM HEPES, TCEP 0.5 mM, NaCl 100 mM). Covalent adduct with WT = 19920 + 321 – 18 = 20223 Da.

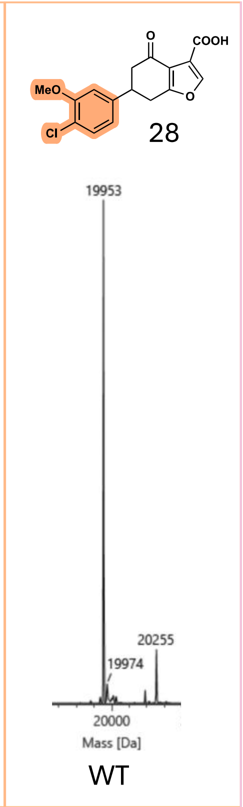

Figure S69 – Mass spectrum of compound **28** (321 Da, 150 µM) incubated at 37 ºC for 5 minutes with WT protein (19953 Da, 2.5 µM) in pH 8 buffer (20 mM HEPES, TCEP 0.5 mM, NaCl 100 mM). Covalent adduct with WT = 19953 + 321 – 18 = 20256 Da.

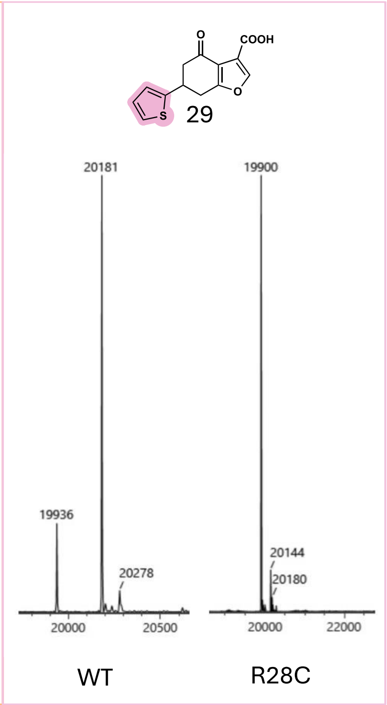

Figure S70 – Mass spectrum of compound **29** (262 Da, 150 µM) incubated at 37 ºC for 5 minutes with WT protein (19936 Da, 2.5 µM) and R28C mutant (19900 Da, 2.5 µM) in pH 8 buffer (20 mM HEPES, TCEP 0.5 mM, NaCl 100 mM). Covalent adduct with WT = 19936 + 262 – 18 = 20180 Da.

Table S2 – Mass spectrum of thiophenes **30** - **34** (150 µM) in competition with the unsubstituted thiophene **29** (150 µM), incubated at 37 ºC for 1 hr with WT protein (19936 Da, 2.5 µM) or Ala mutant, binding unaffected, of the WT protein (19920 Da, 2.5 µM) in pH 8 buffer (20 mM HEPES, TCEP 0.5 mM, NaCl 100 mM).

| 29 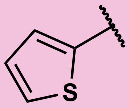 20181/20165 vs. | 30 | 31 | 32 | 33 | 34 |
|---|---|---|---|---|---|
| | 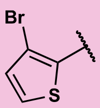 | 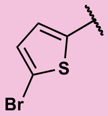 | 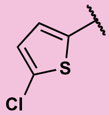 | 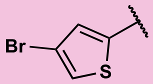 | 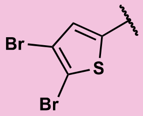 |
| WT/Ala* Mass / Da | 19936 | 19920 | 19920 | 19920 | 19936 |
| WT + lig – 18 / Da | 20259 | 20243 | 20199 | 20243 | 20338 |
| Deconvoluted Spectra | 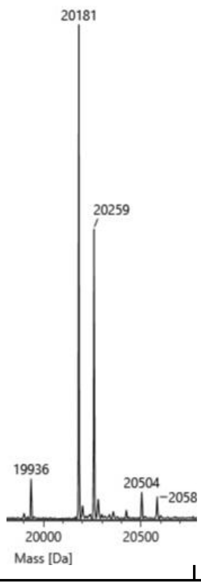 | 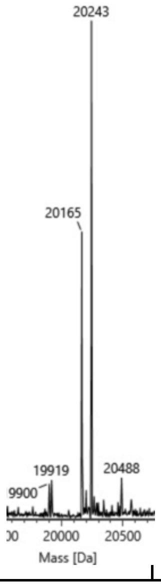 | 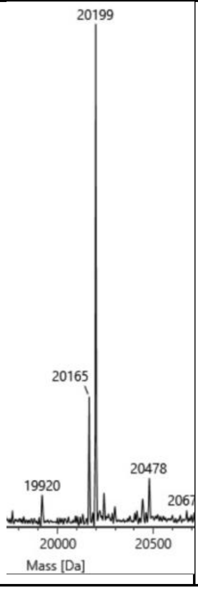 | 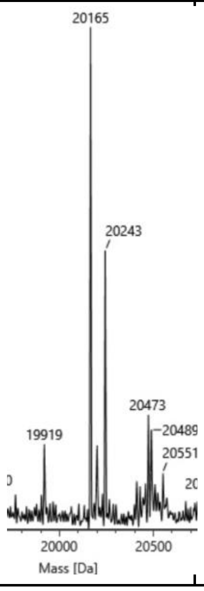 | 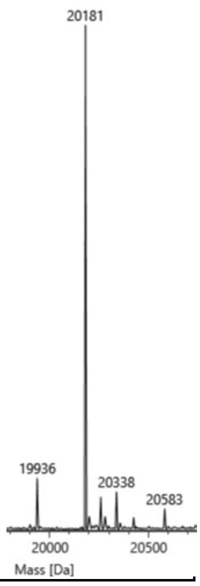 |

#### S4.2 Deconvoluted Spectra for Figure 8C, Reversibility Study

Table S3 – A summary of conditions for the reversibility experiment, displaying the structure of compounds **2** and **24** used in the study.

| Time / hr | Item Name | | |
|---|---|---|---|
| | rm204a – **2** then **24** | 204b – **24** then **2** | 204e – **2** and **24** |
| 1 | 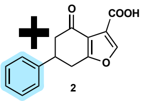 | 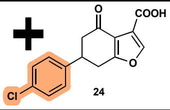 | 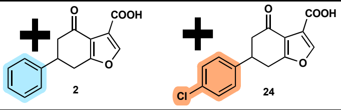 |
| 2 | 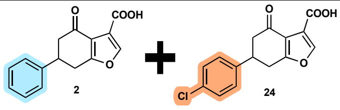 | 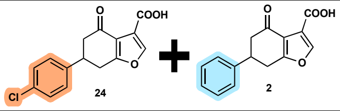 | 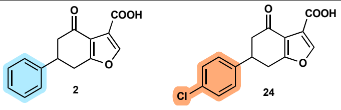 |
| 24 | 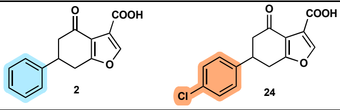 | 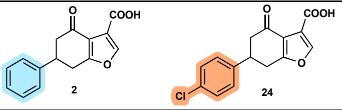 | 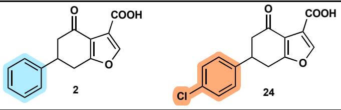 |

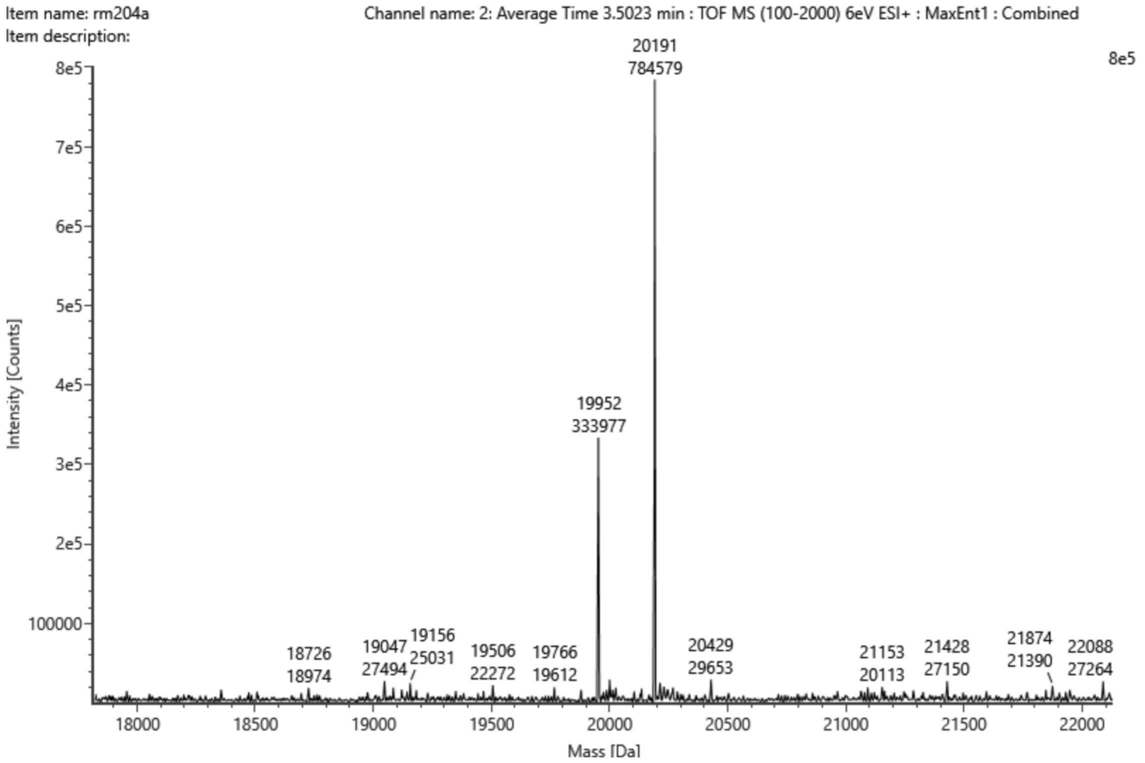

Figure S71 – Mass spectrum with intensities of compound **2** (256 Da, 150 µM) incubated at 37 ºC for 1 hr with WT protein (19952 Da, 2.5 µM) in pH 8 buffer (20 mM HEPES, TCEP 0.5 mM, NaCl 100 mM). Covalent adduct of WT protein + **2** = 19952 + 256 – 18 = 20191 Da.

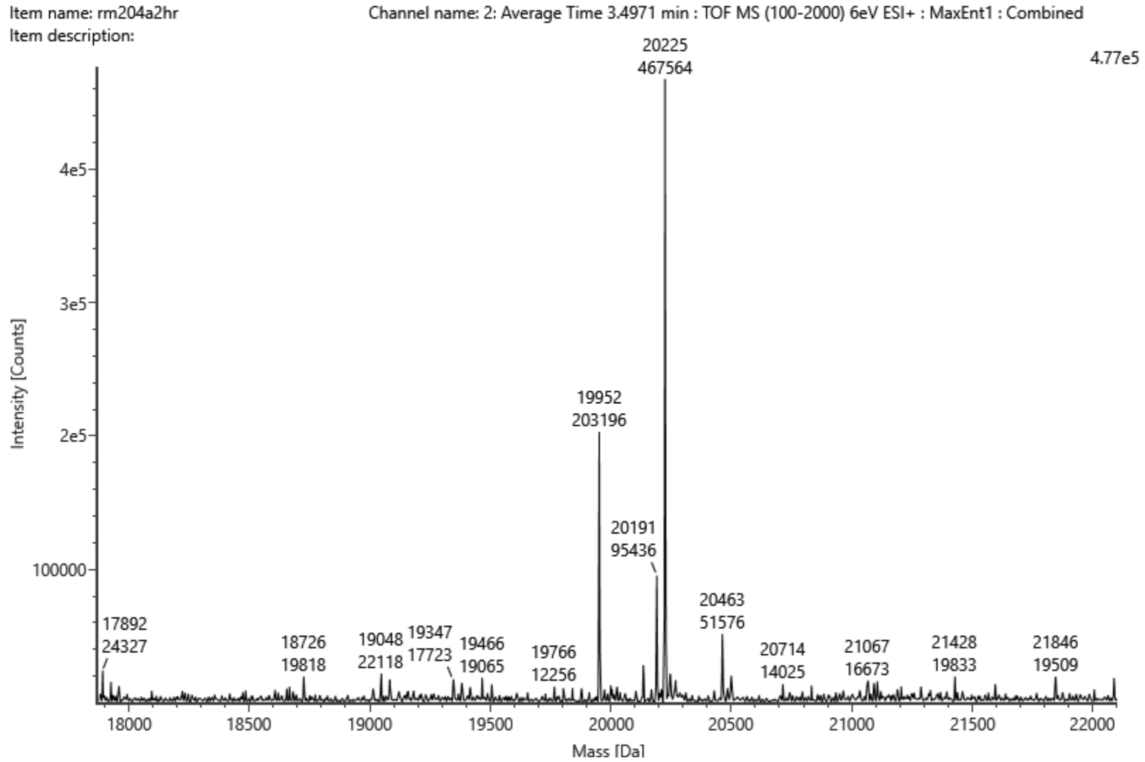

Figure S72 – Mass spectrum with intensities of compound **2** (256 Da, 150 µM) incubated at 37 ºC for 2 hrs, adding compound **24** (291 Da, 150 µM) after 1 hr, with WT protein (19952 Da, 2.5 µM) in pH 8 buffer (20 mM HEPES, TCEP 0.5 mM, NaCl 100 mM). Covalent adduct of WT protein + **2** = 19952 + 256 – 18 = 20191 Da. Covalent adduct of WT protein + **24** = 19952 + 291 – 18 = 20225 Da.

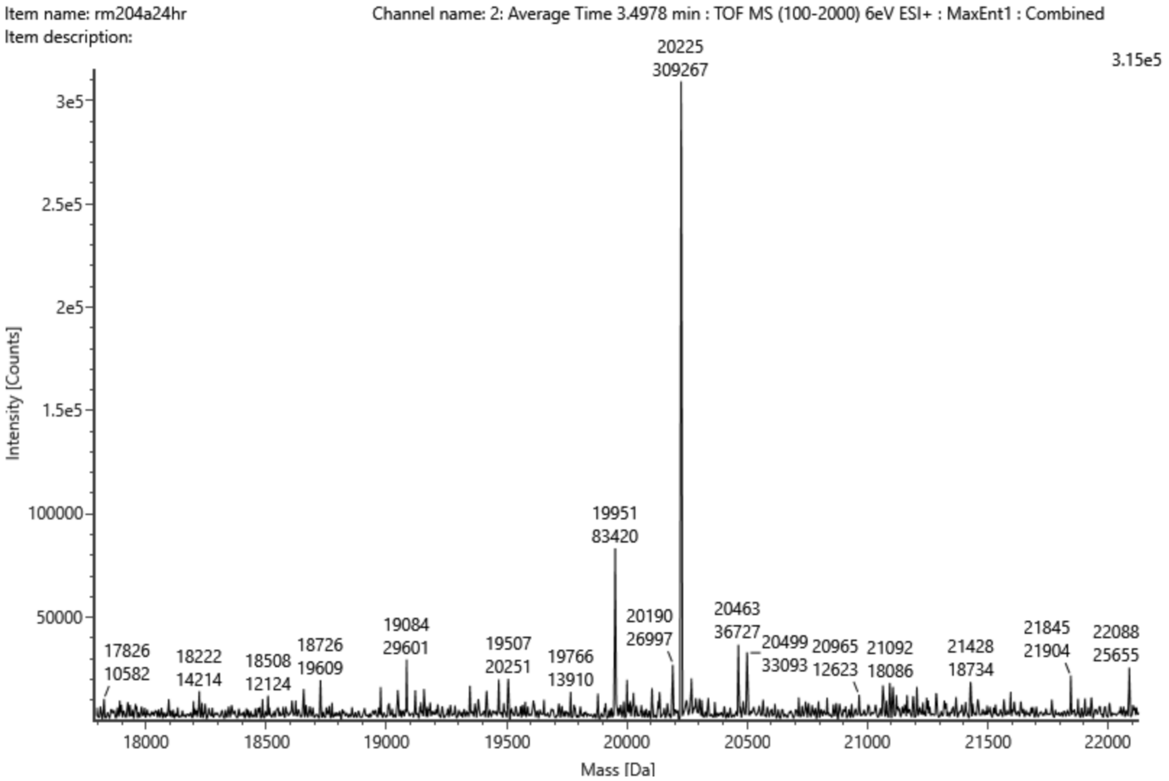

Figure S73 – Mass spectrum with intensities of compound **2** (256 Da, 150 µM) incubated at 37 ºC for 24 hrs, adding compound **24** (291 Da, 150 µM) after 1 hr, with WT protein (19952 Da, 2.5 µM) in pH 8 buffer (20 mM HEPES, TCEP 0.5 mM, NaCl 100 mM). Covalent adduct of WT protein + **2** = 19952 + 256 – 18 = 20191 Da. Covalent adduct of WT protein + **24** = 19952 + 291 – 18 = 20225 Da.

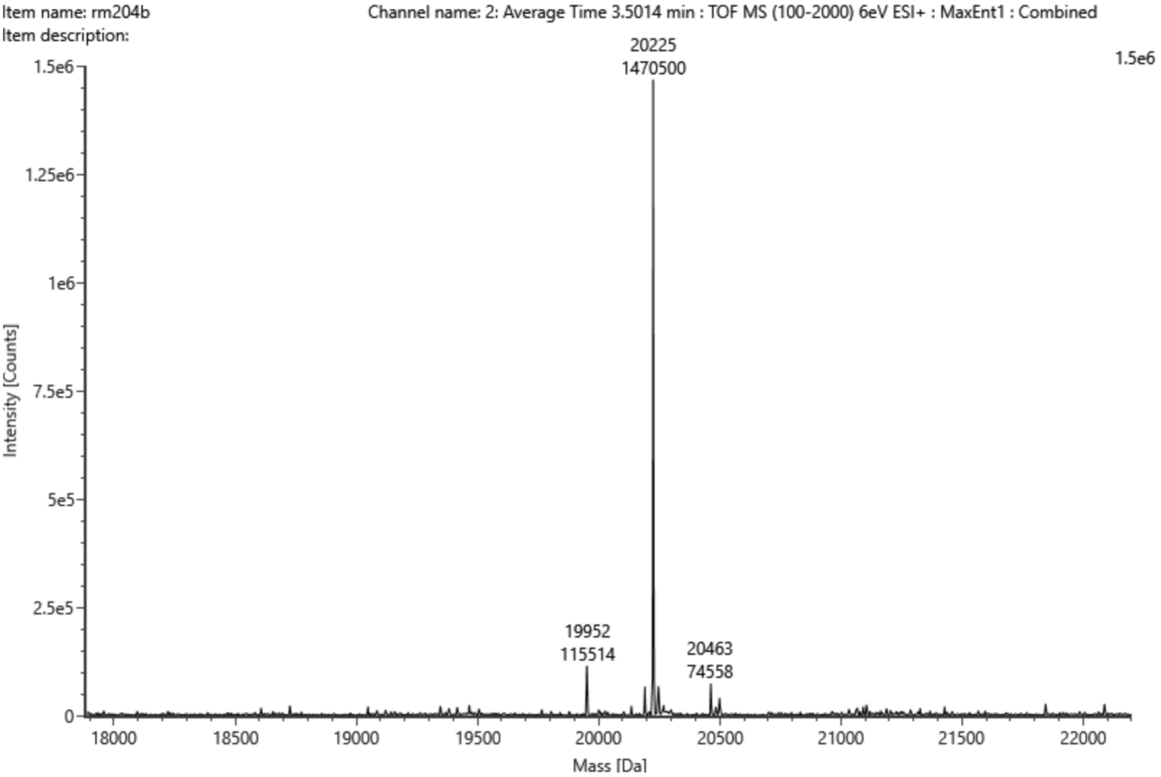

Figure S74 – Mass spectrum with intensities of compound **24** (291 Da, 150 µM) incubated at 37 ºC for 1 hr with WT protein (19952 Da, 2.5 µM) in pH 8 buffer (20 mM HEPES, TCEP 0.5 mM, NaCl 100 mM). Covalent adduct of WT protein + **24** = 19952 + 291 – 18 = 20225 Da.

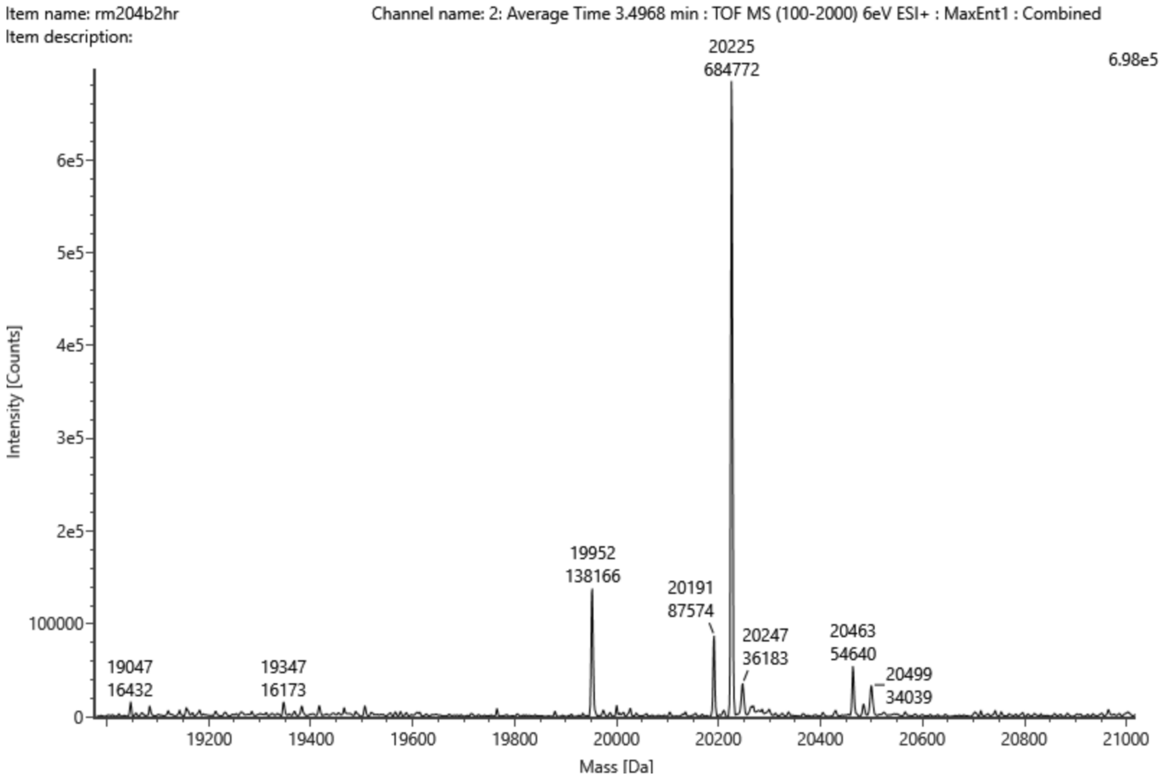

Figure S75 – Mass spectrum with intensities of compound **24** (291 Da, 150 µM) incubated at 37 ºC for 2 hrs, adding compound **2** (256 Da, 150 µM) after 1 hr, with WT protein (19952 Da, 2.5 µM) in pH 8 buffer (20 mM HEPES, TCEP 0.5 mM, NaCl 100 mM). Covalent adduct of WT protein + **24** = 19952 + 291 – 18 = 20225 Da. Covalent adduct of WT protein + **2** = 19952 + 256 – 18 = 20191 Da.

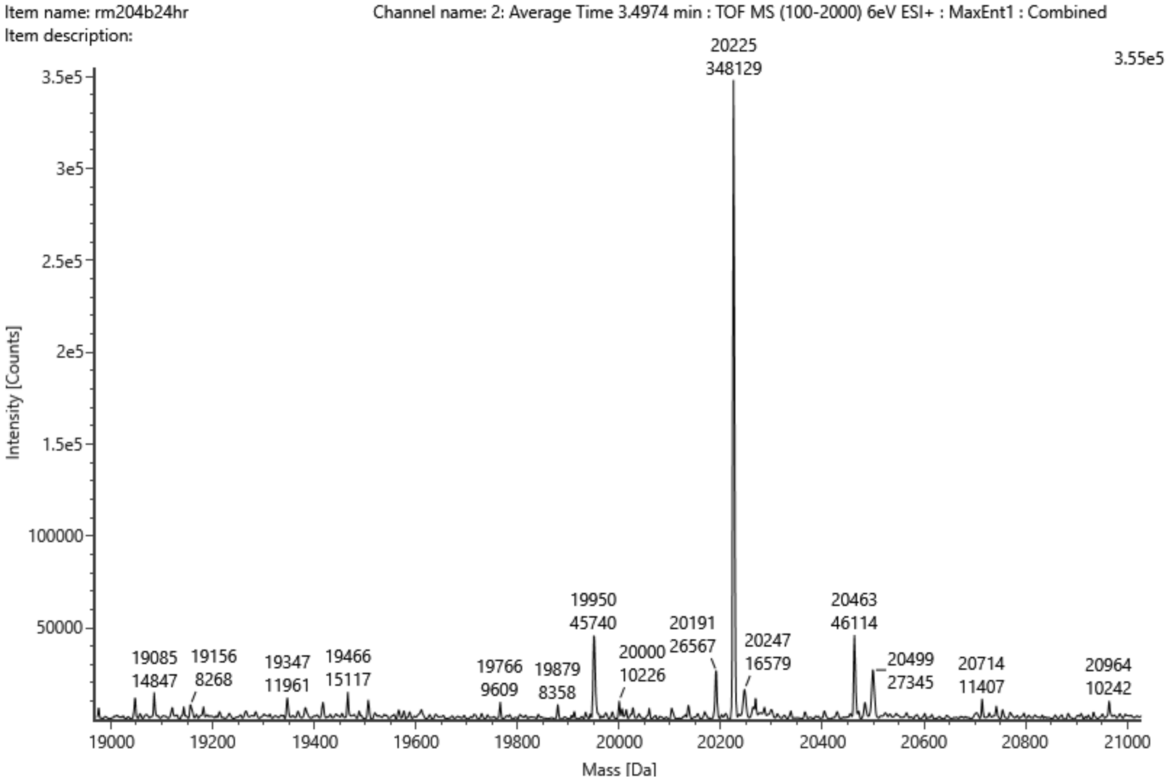

Figure S76 – Mass spectrum with intensities of compound **24** (291 Da, 150 µM) incubated at 37 ºC for 24 hrs, adding compound **2** (256 Da, 150 µM) after 1 hr, with WT protein (19952 Da, 2.5 µM) in pH 8 buffer (20 mM HEPES, TCEP 0.5 mM, NaCl 100 mM). Covalent adduct of WT protein + **24** = 19952 + 291 – 18 = 20225 Da. Covalent adduct of WT protein + **2** = 19952 + 256 – 18 = 20191 Da.

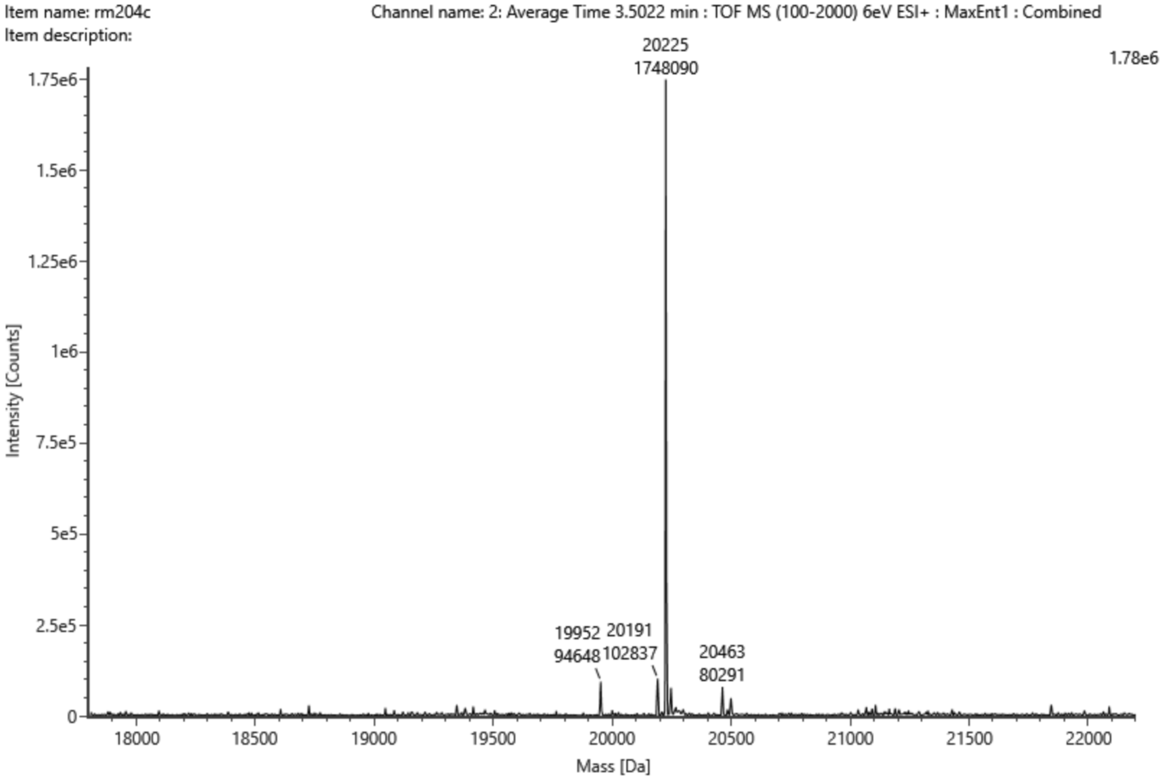

Figure S77 – Mass spectrum with intensities of compound **2** (256 Da, 150 μM) and compound **24** (291 Da, 150 μM) incubated at 37 ºC for 1 hr with WT protein (19952 Da, 2.5 μM) in pH 8 buffer (20 mM HEPES, TCEP 0.5 mM, NaCl 100 mM). Covalent adduct of WT protein + **2** = 19952 + 256 – 18 = 20191 Da. Covalent adduct of WT protein + **24** = 19952 + 291 – 18 = 20225 Da.

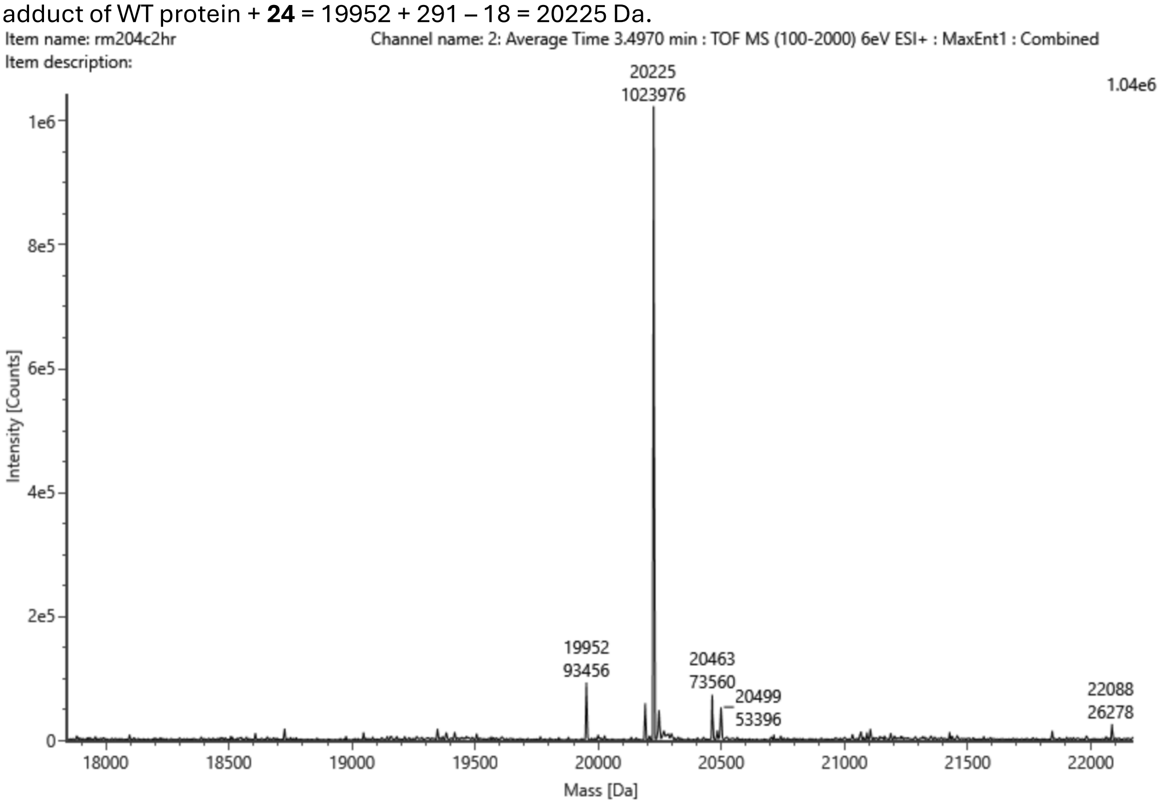

Figure S78 – Mass spectrum with intensities of compound **2** (256 Da, 150 μM) and compound **24** (291 Da, 150 μM) incubated at 37 ºC for 2 hrs with WT protein (19952 Da, 2.5 μM) in pH 8 buffer (20 mM HEPES, TCEP 0.5 mM, NaCl 100 mM). Covalent adduct of WT protein + **2** = 19952 + 256 – 18 = 20191 Da. Covalent adduct of WT protein + **24** = 19952 + 291 – 18 = 20225 Da.

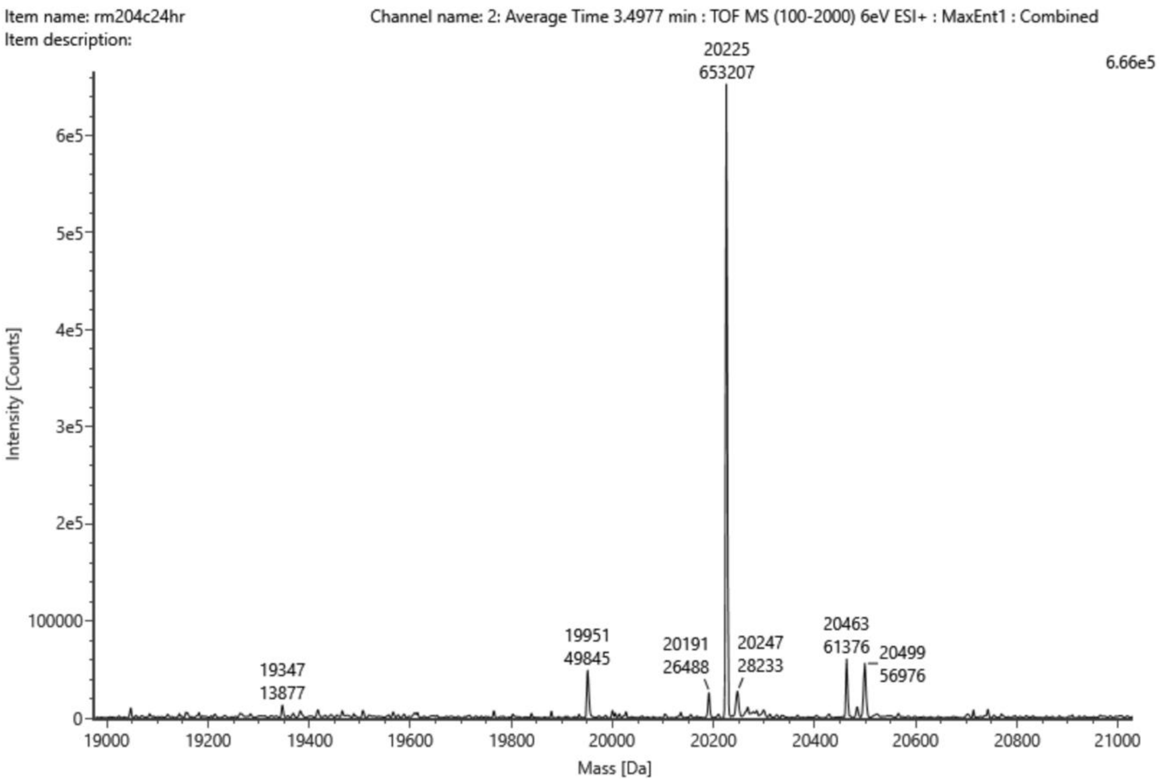

Figure S79 – Mass spectrum with intensities of compound **2** (256 Da, 150 µM) and compound **24** (291 Da, 150 µM) incubated at 37 ºC for 1 hr with WT protein (19952 Da, 2.5 µM) in pH 8 buffer (20 mM HEPES, TCEP 0.5 mM, NaCl 100 mM). Covalent adduct of WT protein + **2** = 19952 + 256 – 18 = 20191 Da. Covalent adduct of WT protein + **24** = 19952 + 291 – 18 = 20225 Da.

#### S4.3 pH and time dependence of reaction, Figure S80 and Deconvoluted Spectra

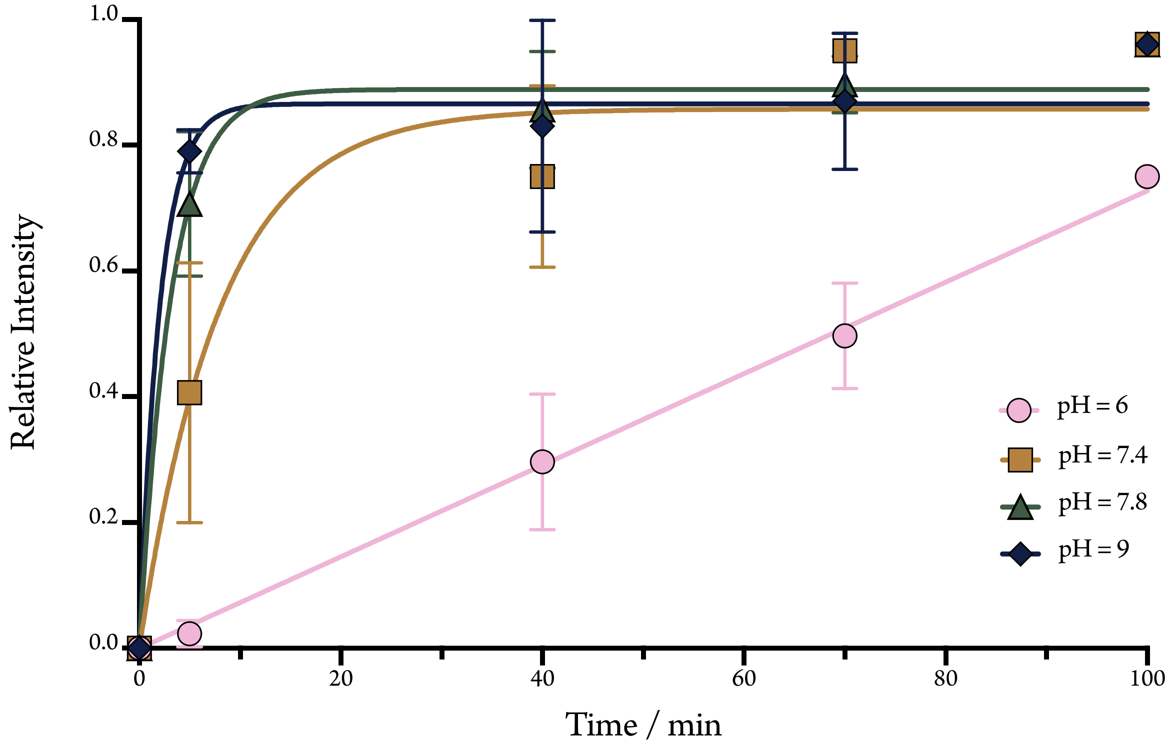

Figure S80 – Investigating pH and time dependence of labelling by comparing the intensity of peaks after incubation of **16** (150 µM) with the WT BTK PH domain (2.5 µM) for 5, 40, 70 and 100 minutes in 20 mM HEPES buffers (100 mM NaCl, 0.5 mM TCEP) of various pH values (pH 6, 7.4, 7.8 and 9). The data does not distinguish single and non-specific labelling as it shows the intensity of total labelling (singly and doubly labelled protein) relative to total protein labelling (singly, doubly and unlabelled protein).

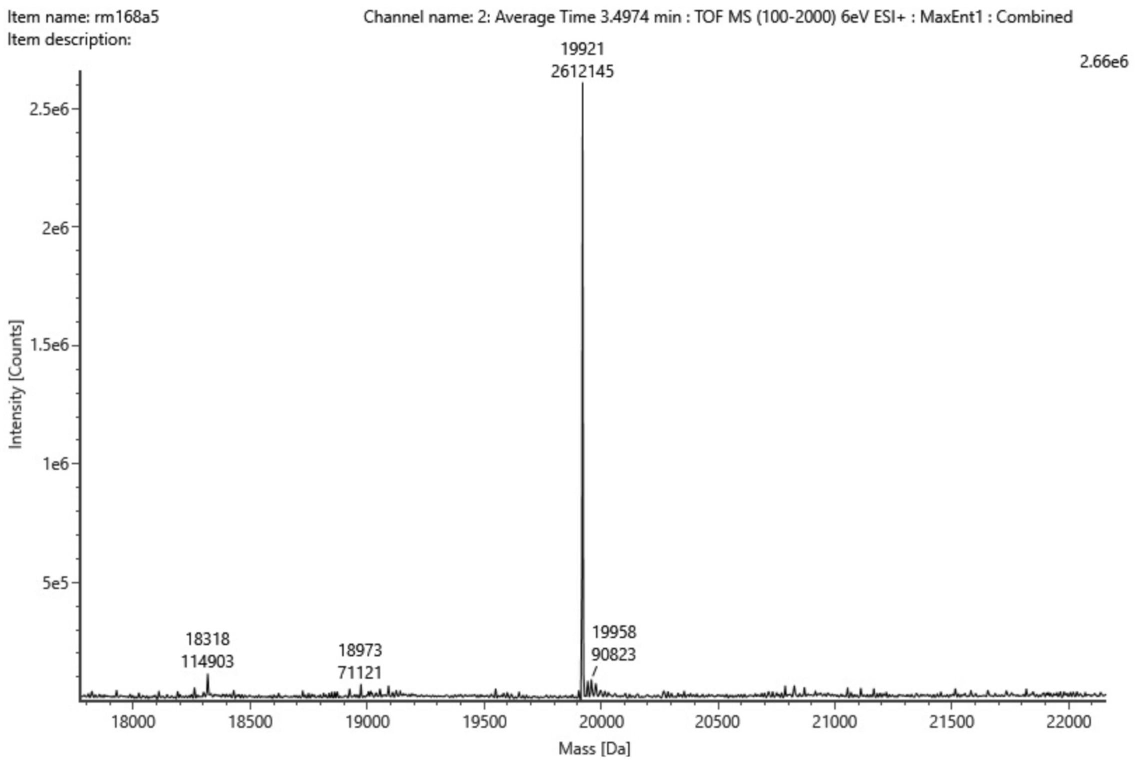

Figure S81 – Mass spectrum with intensities of compound **16** (365 Da, 150 µM) incubated at 37 ºC for 5 minutes with WT protein (19921 Da, 2.5 µM) with pH 6 buffer (20 mM HEPES, TCEP 0.5 mM, NaCl 100 mM).

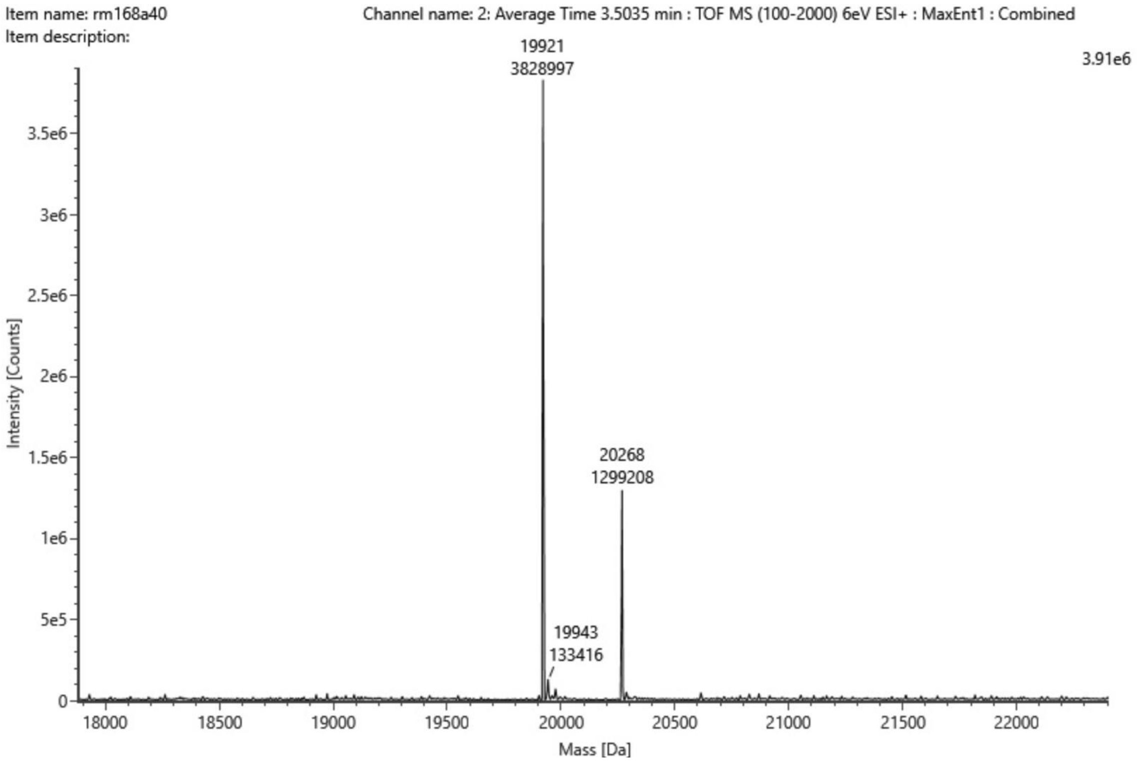

Figure S82 – Mass spectrum with intensities of compound **16** (365 Da, 150 µM) incubated at 37 ºC for 40 minutes with WT protein (19921 Da, 2.5 µM) with pH 6 buffer (20 mM HEPES, TCEP 0.5 mM, NaCl 100 mM). Covalent adduct of WT protein + **16** = 19921 + 365 – 18 = 20267 Da.

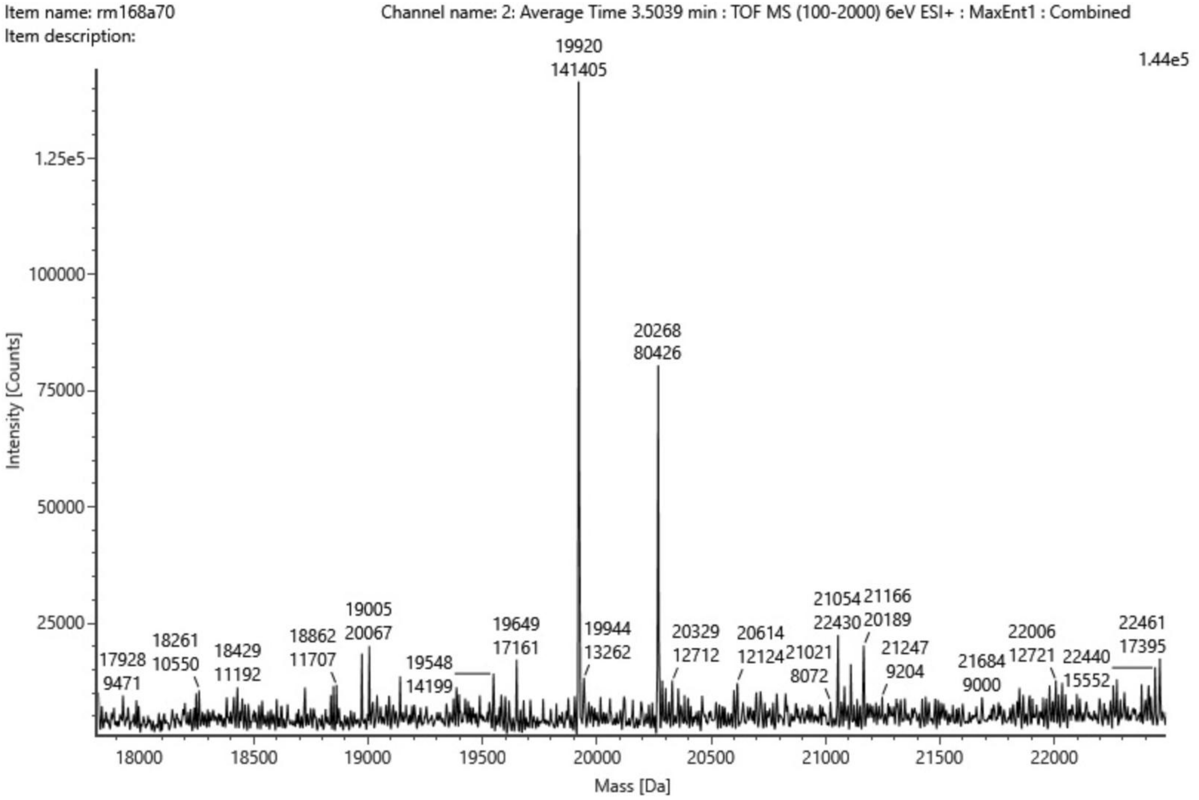

Figure S83 – Mass spectrum with intensities of compound **16** (365 Da, 150 µM) incubated at 37 ºC for 70 minutes with WT protein (19920 Da, 2.5 µM) with pH 6 buffer (20 mM HEPES, TCEP 0.5 mM, NaCl 100 mM). Covalent adduct of WT protein + **16** = 19920 + 365 – 18 = 20267 Da.

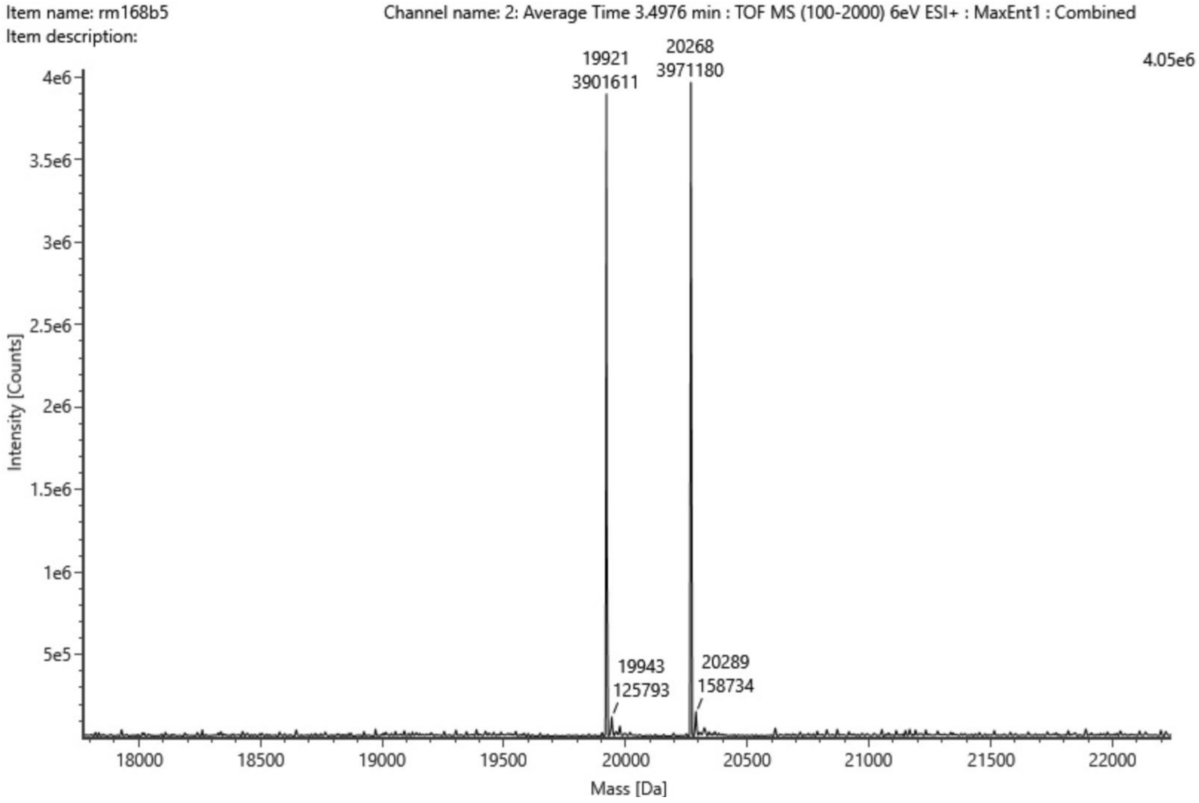

Figure S84 – Mass spectrum with intensities of compound **16** (365 Da, 150 µM) incubated at 37 ºC for 5 minutes with WT protein (19921 Da, 2.5 µM) with pH 7.4 buffer (20 mM HEPES, TCEP 0.5 mM, NaCl 100 mM). Covalent adduct of WT protein + **16** = 19921 + 365 – 18 = 20268 Da.

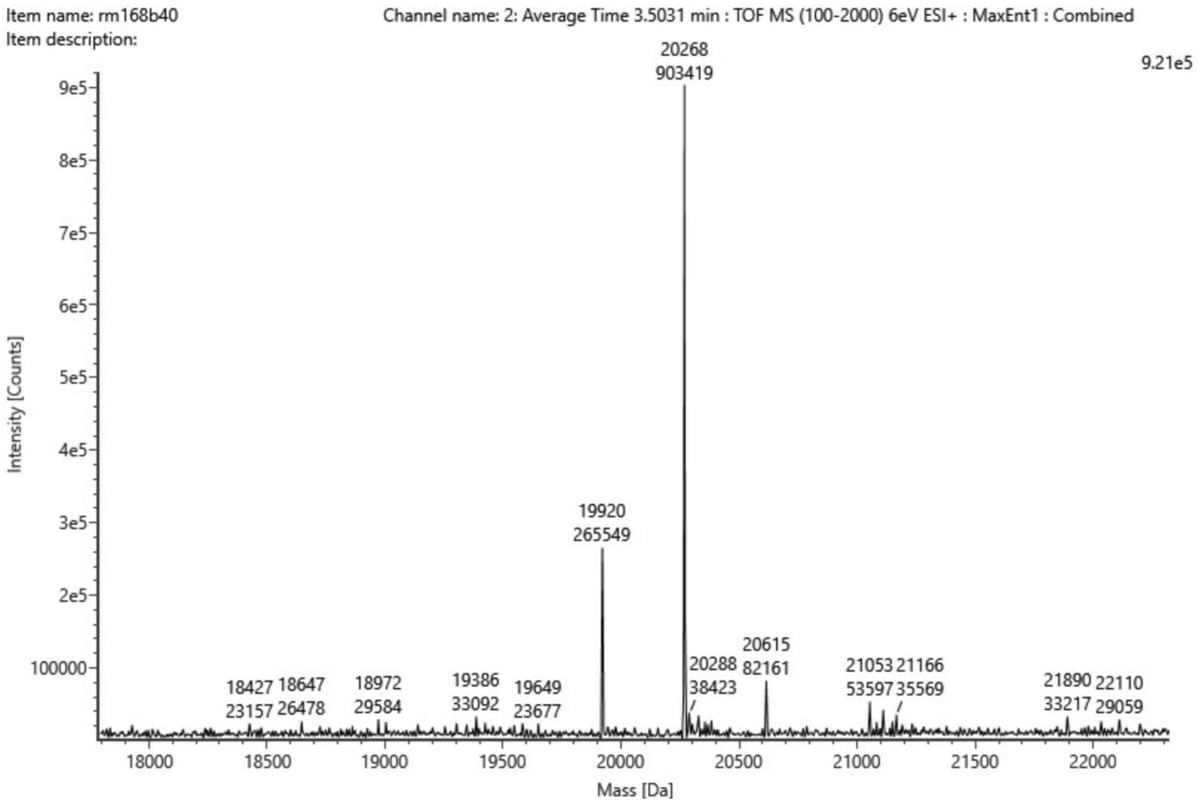

Figure S85 – Mass spectrum with intensities of compound **16** (365 Da, 150 µM) incubated at 37 ºC for 40 minutes with WT protein (19920 Da, 2.5 µM) with pH 7.4 buffer (20 mM HEPES, TCEP 0.5 mM, NaCl 100 mM). Covalent adduct of WT protein + **16** = 19920 + 365 – 18 = 20267 Da.

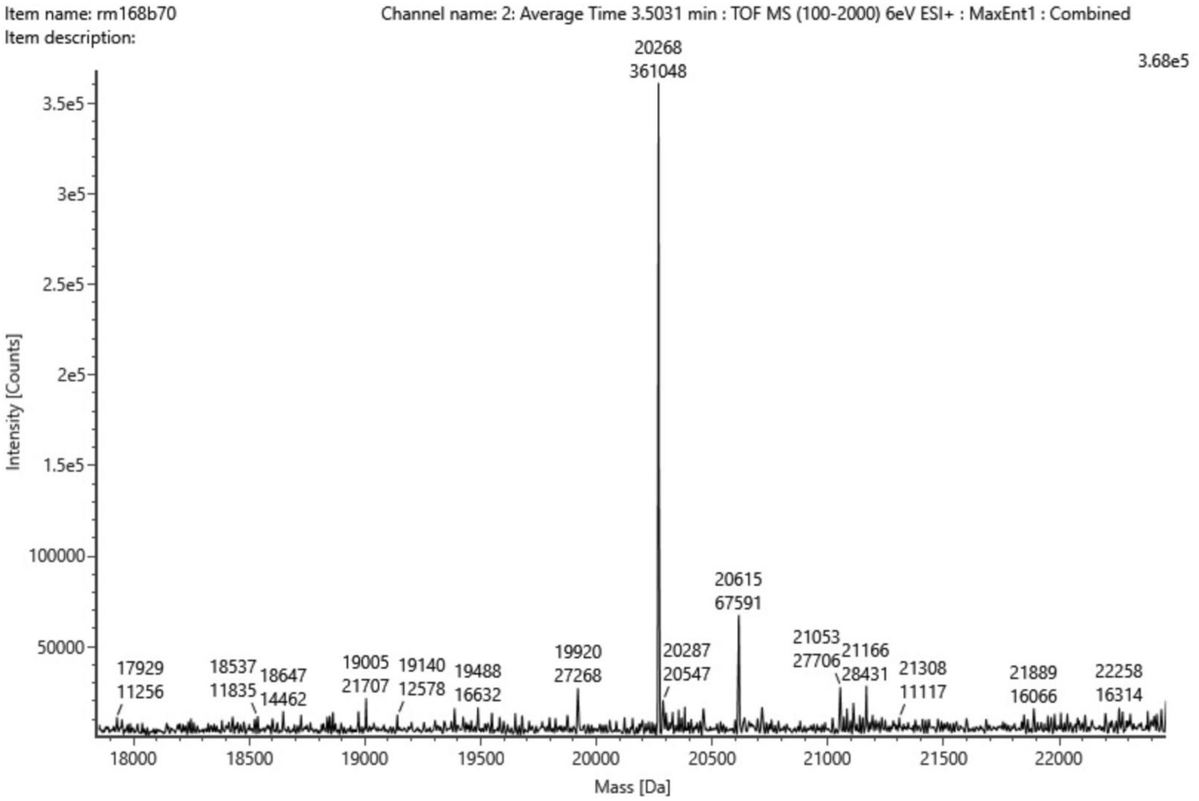

Figure S86 – Mass spectrum with intensities of compound **16** (365 Da, 150 µM) incubated at 37 ºC for 70 minutes with WT protein (19920 Da, 2.5 µM) with pH 7.4 buffer (20 mM HEPES, TCEP 0.5 mM, NaCl 100 mM). Covalent adduct of WT protein + **16** = 19920 + 365 – 18 = 20267 Da.

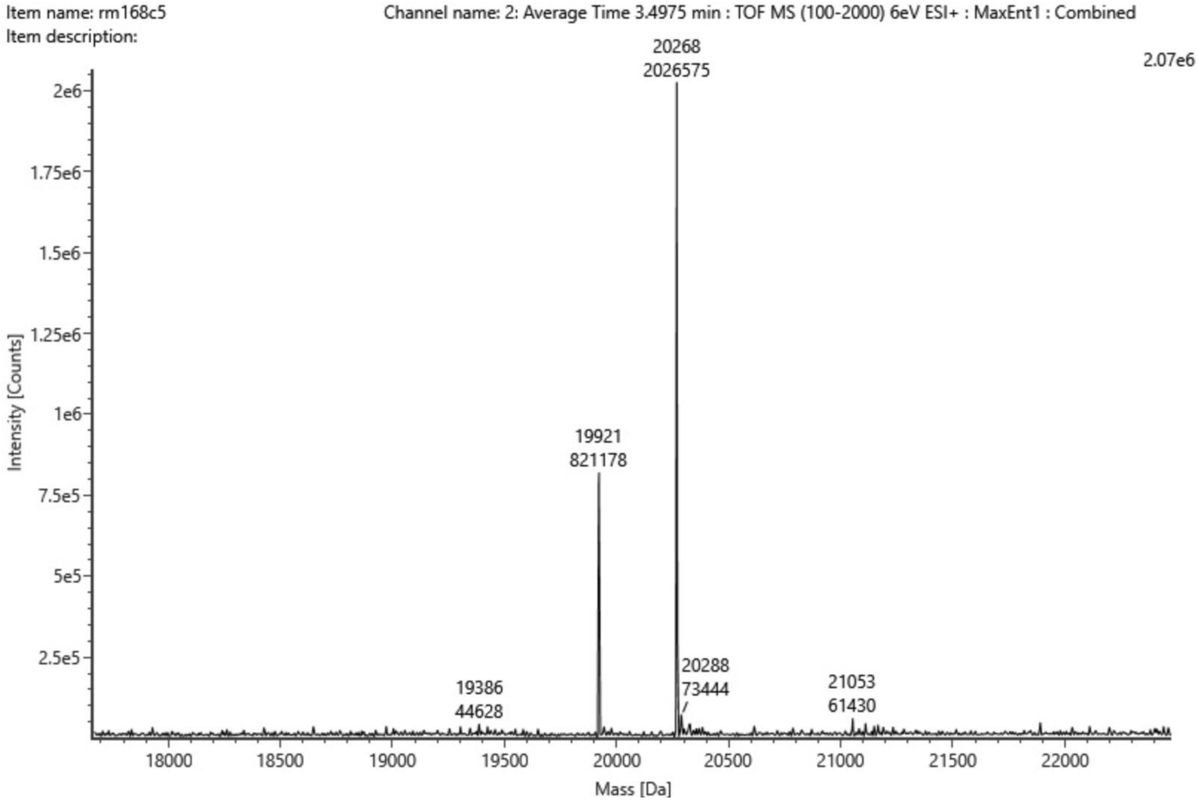

Figure S87 – Mass spectrum with intensities of compound **16** (365 Da, 150 µM) incubated at 37 ºC for 5 minutes with WT protein (19921 Da, 2.5 µM) with pH 7.8 buffer (20 mM HEPES, TCEP 0.5 mM, NaCl 100 mM). Covalent adduct of WT protein + **16** = 19921 + 365 – 18 = 20268 Da.

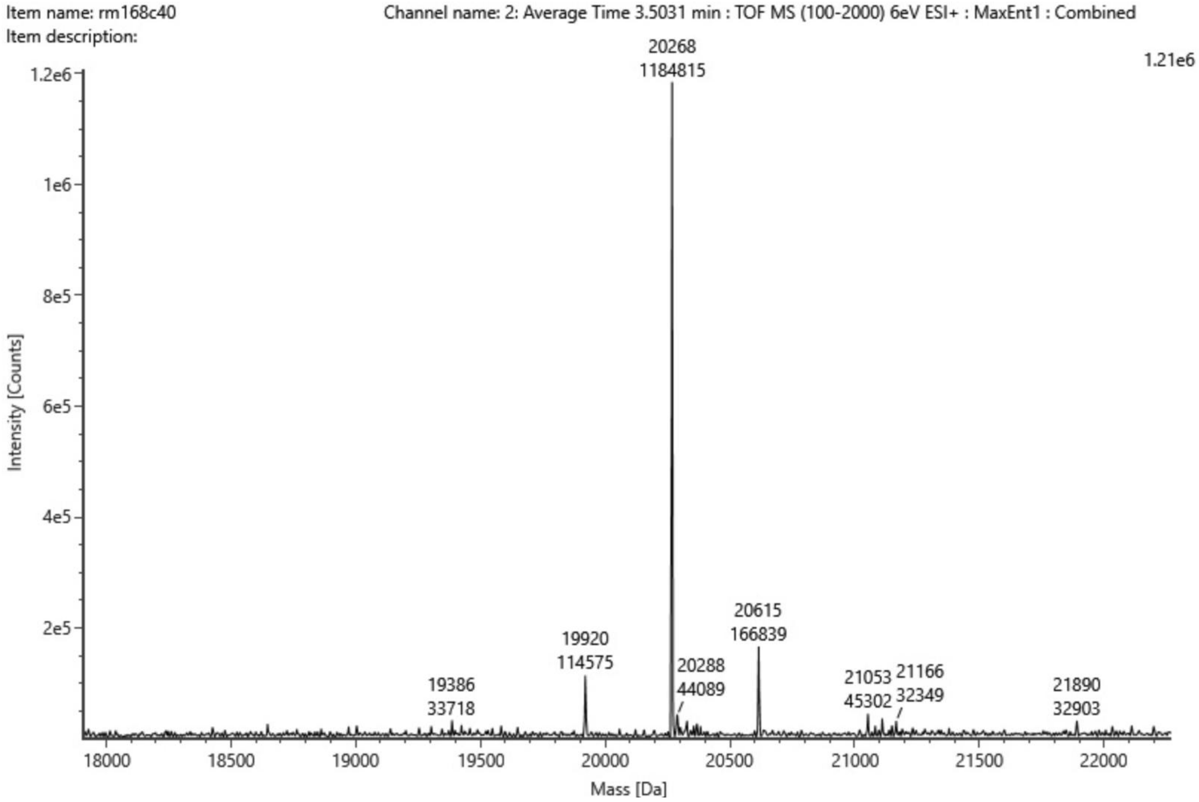

Figure S88 – Mass spectrum with intensities of compound **16** (365 Da, 150 µM) incubated at 37 ºC for 40 minutes with WT protein (19920 Da, 2.5 µM) with pH 7.8 buffer (20 mM HEPES, TCEP 0.5 mM, NaCl 100 mM). Covalent adduct of WT protein + **16** = 19920 + 365 – 18 = 20268 Da.

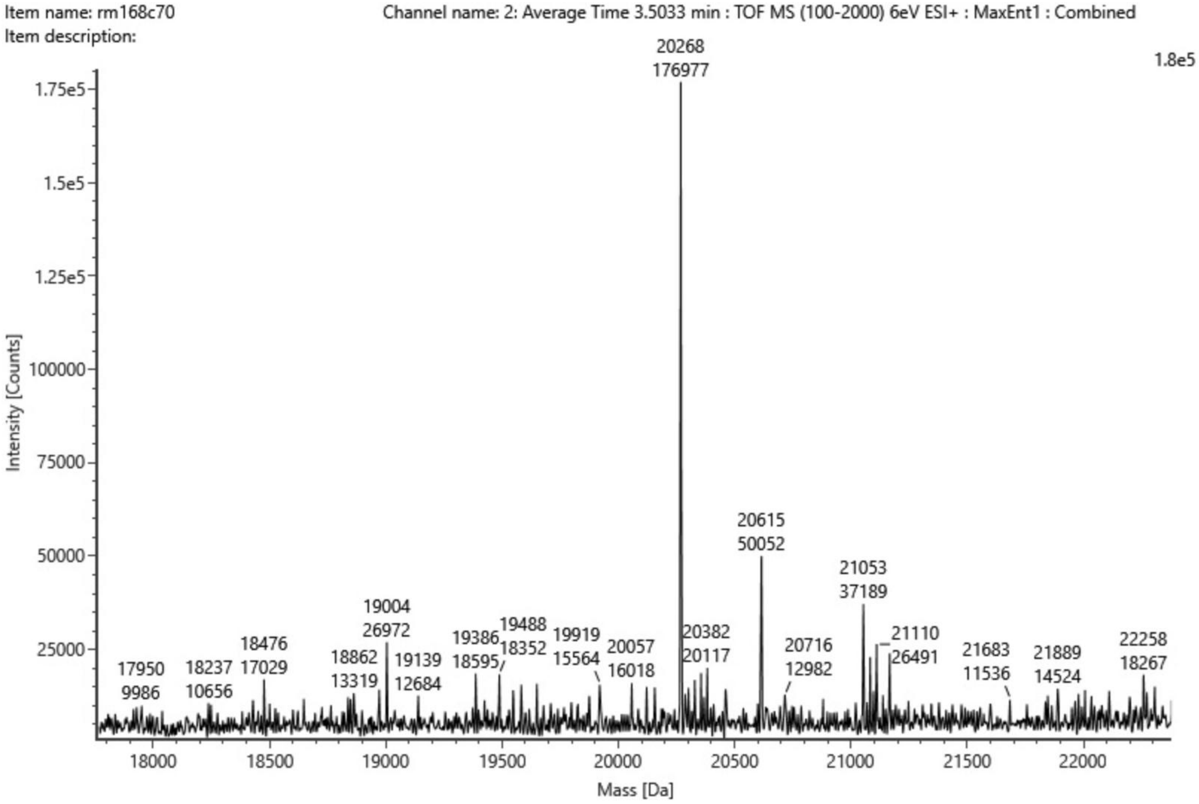

Figure S89 – Mass spectrum with intensities of compound **16** (365 Da, 150 µM) incubated at 37 ºC for 70 minutes with WT protein (19921 Da, 2.5 µM) with pH 7.8 buffer (20 mM HEPES, TCEP 0.5 mM, NaCl 100 mM). Covalent adduct of WT protein + **16** = 19921+ 365 – 18 = 20268 Da.

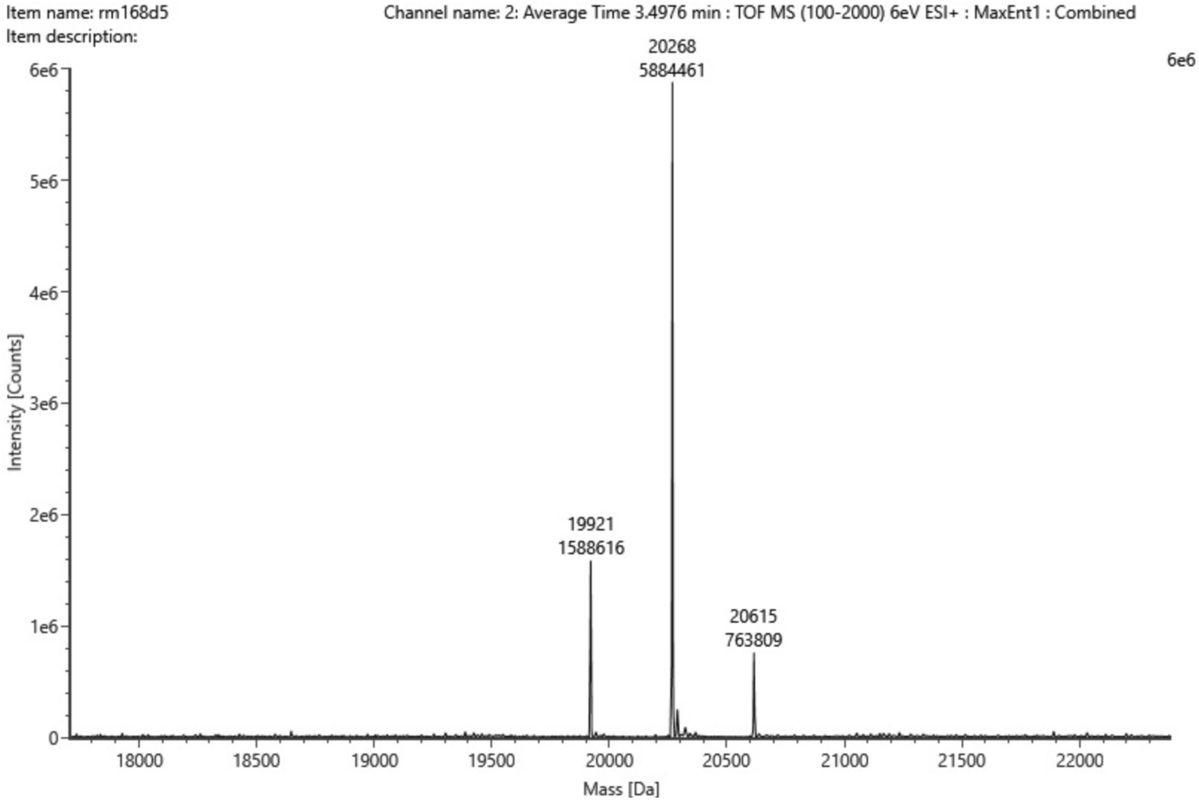

Figure S90 – Mass spectrum with intensities of compound **16** (365 Da, 150 µM) incubated at 37 ºC for 5 minutes with WT protein (19921 Da, 2.5 µM) with pH 9 buffer (20 mM HEPES, TCEP 0.5 mM, NaCl 100 mM). Covalent adduct of WT protein + **16** = 19921+ 365 – 18 = 20268 Da.

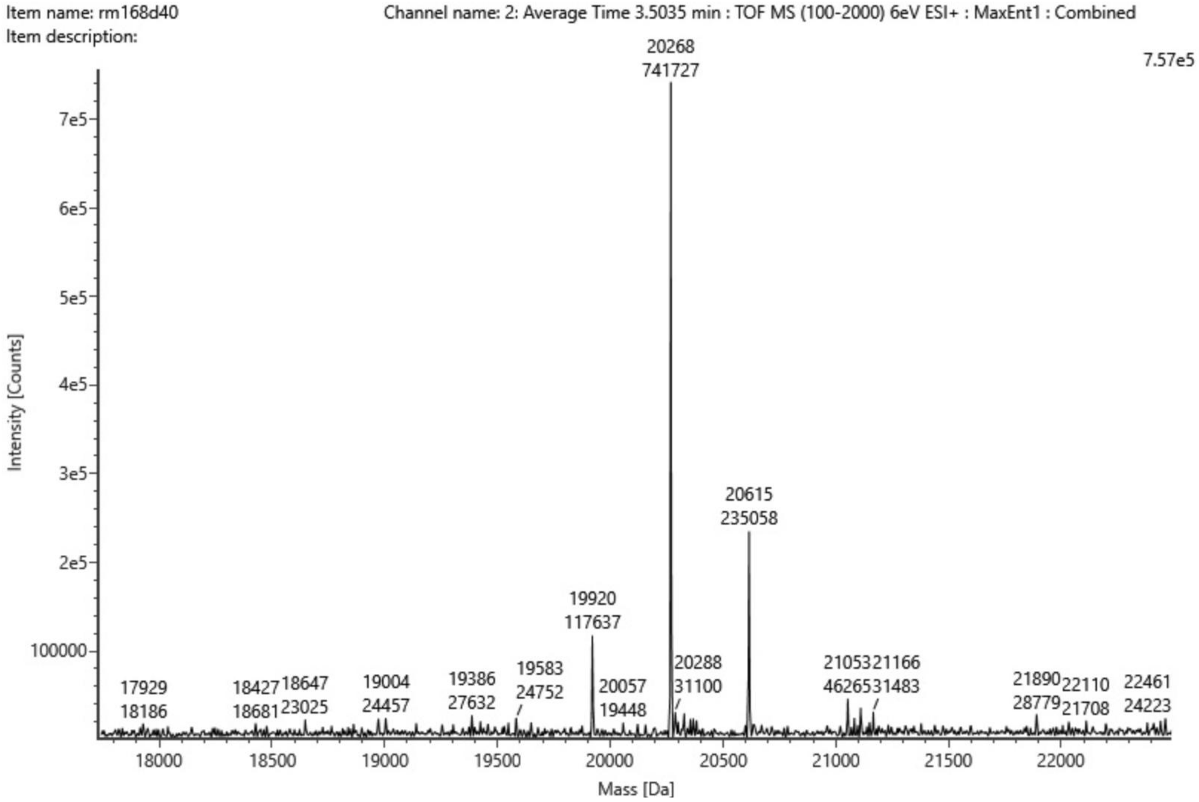

Figure S91 – Mass spectrum with intensities of compound **16** (365 Da, 150 µM) incubated at 37 ºC for 40 minutes with WT protein (19920 Da, 2.5 µM) with pH 9 buffer (20 mM HEPES, TCEP 0.5 mM, NaCl 100 mM). Covalent adduct of WT protein + **16** = 19920 + 365 – 18 = 20267 Da.

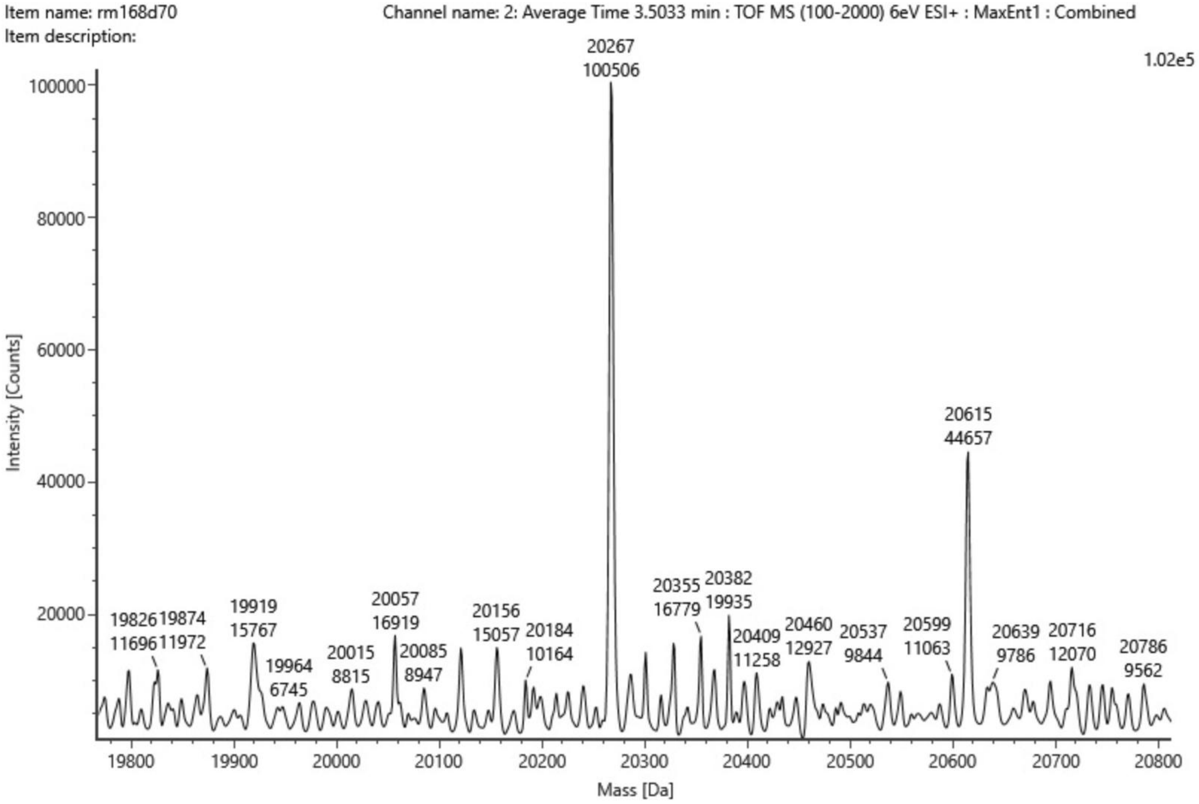

Figure S92 – Mass spectrum with intensities of compound **16** (365 Da, 150 µM) incubated at 37 ºC for 70 minutes with WT protein (19920 Da, 2.5 µM) with pH 9 buffer (20 mM HEPES, TCEP 0.5 mM, NaCl 100 mM). Covalent adduct of WT protein + **16** = 19920 + 365 – 18 = 20267 Da.

#### S4.4 Negative Controls - Deconvoluted Spectra

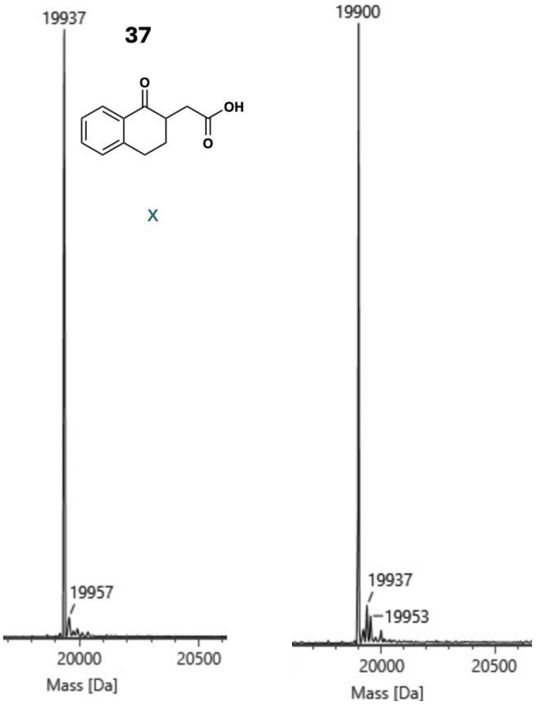

**WT** **R28C**

Figure S93 – Mass spectrum of compound **37** (204 Da, 150 µM) incubated at 37 ºC for 5 minutes with WT protein (19936 Da, 2.5 µM) and R28C mutant (19900 Da, 2.5 µM) in pH 8 buffer (20 mM HEPES, TCEP 0.5 mM, NaCl 100 mM).

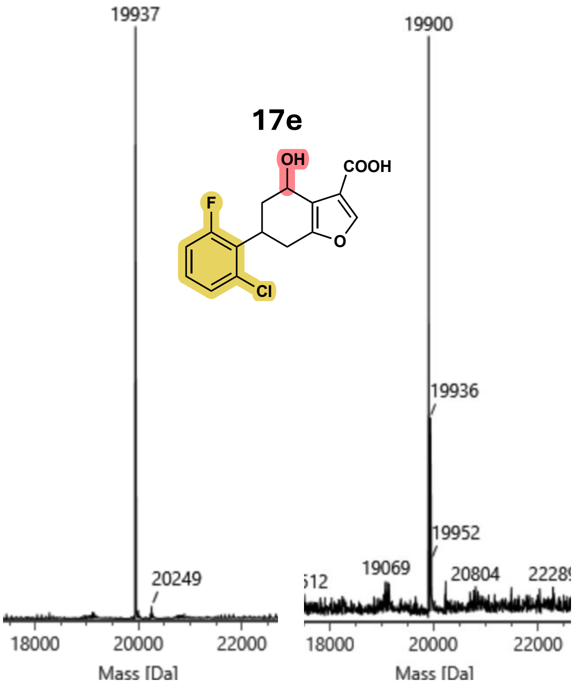

**WT** **R28C**

Figure S94 – Mass spectrum of compound **17e** (311 Da, 150 µM) incubated at 37 ºC for 5 minutes with WT protein (19937 Da, 2.5 µM) and R28C mutant (19900 Da, 2.5 µM) in pH 8 buffer (20 mM HEPES, TCEP 0.5 mM, NaCl 100 mM). Non-covalent adduct with WT = 19937 + 311 = 20249 Da.

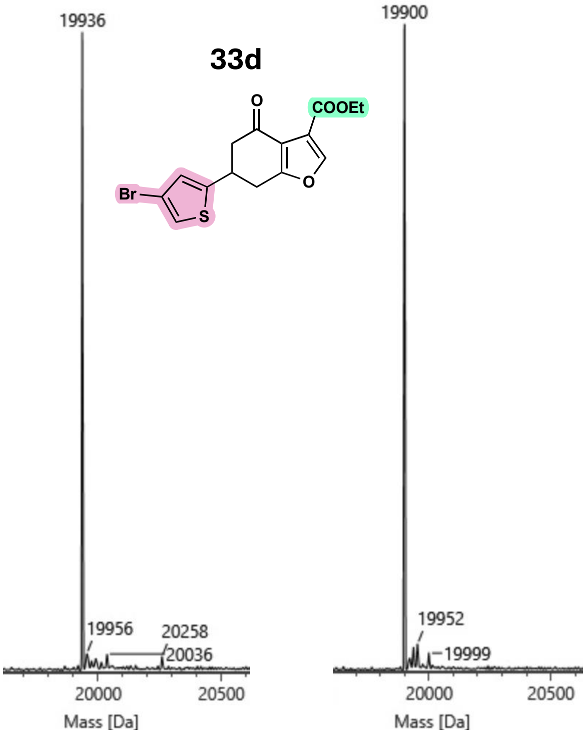

**WT** **R28C**

Figure S95 – Mass spectrum of compound **29** (369 Da, 150 µM) incubated at 37 °C for 5 minutes with WT protein (19936 Da, 2.5 µM) and R28C mutant (19900 Da, 2.5 µM) in pH 8 buffer (20 mM HEPES, TCEP 0.5 mM, NaCl 100 mM). Not observed: covalent adduct with WT = 19936 + 369 – 18 = 20287 Da, non-covalent adduct with WT = 19936 + 369 = 20305 Da.

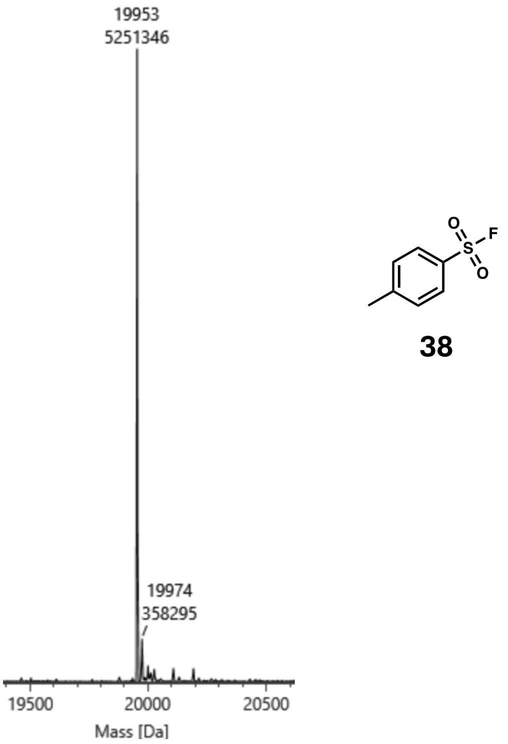

**WT**

Figure S96 – Mass spectrum with intensities of compound **38** (174 Da, 150 µM) incubated at 37 °C for 5 minutes with WT protein (19953 Da, 2.5 µM) in pH 8 buffer (20 mM HEPES, TCEP 0.5 mM, NaCl 100 mM). Not observed: covalent adduct with WT = 19953 + 174 – 19 = 20108 Da.

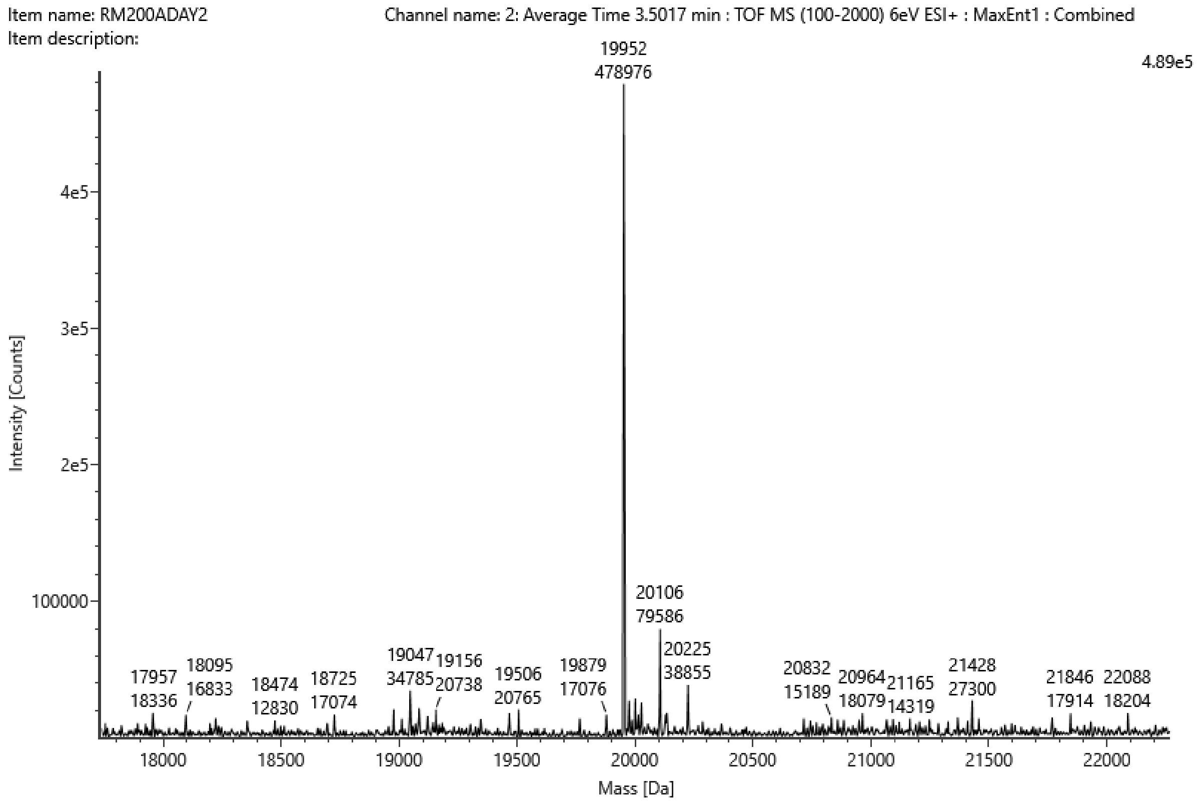

Figure S97 – Mass spectrum with intensities of compound **38** (174 Da, 1 mM) incubated at 37 ºC for 5 minutes and then at RT overnight with WT protein (19952 Da, 2.5 µM) in pH 8 buffer (20 mM HEPES, TCEP 0.5 mM, NaCl 100 mM). Observed: covalent adduct with WT = 19952 + 174 – 19 = 20107 Da.

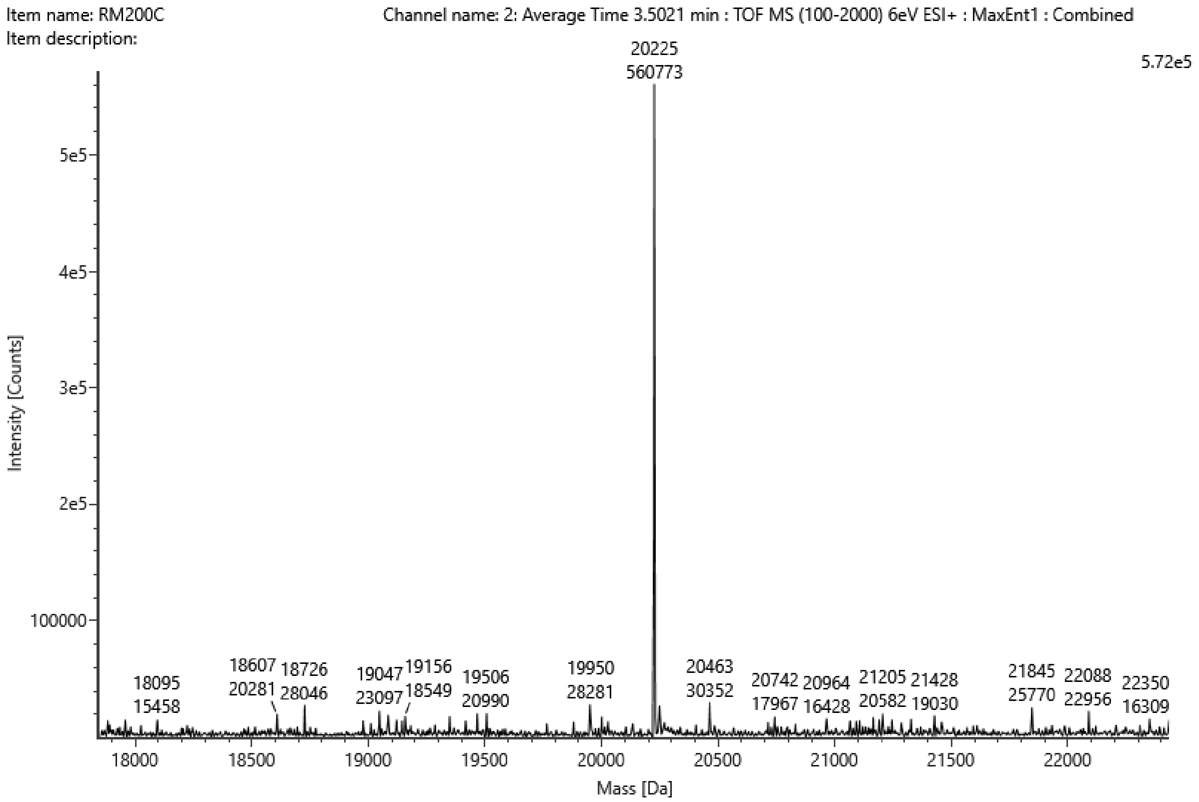

Figure S98 – Mass spectrum with intensities of compound **24** (291 Da, 150 µM) incubated at 37 ºC for 5 minutes and then at RT overnight with WT protein (19952 Da, 2.5 µM) in pH 8 buffer (20 mM HEPES, TCEP 0.5 mM, NaCl 100 mM). Covalent adduct with WT = 19952 + 291 – 18 = 20225 Da..

#### S4.5 Enantiomerically Pure Ligands

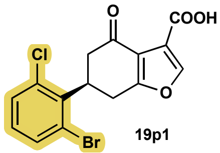

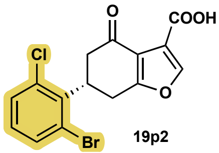

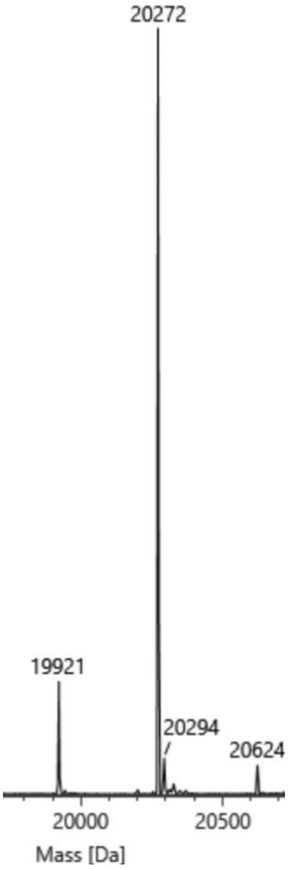

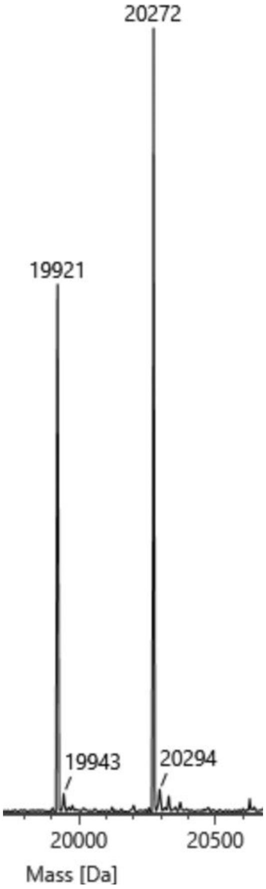

Figure S99 – Mass spectra of enantiomerically pure compounds **19p1** and **19p2** (369 Da, 150 μM) incubated at 37 ºC for 5 minutes with WT protein (19921 Da, 2.5 μM) in pH 8 buffer (20 mM HEPES, TCEP 0.5 mM, NaCl 100 mM). Covalent adduct with WT = 19921 + 369 – 18 = 20272 Da.

### S5 MALDI-TOF

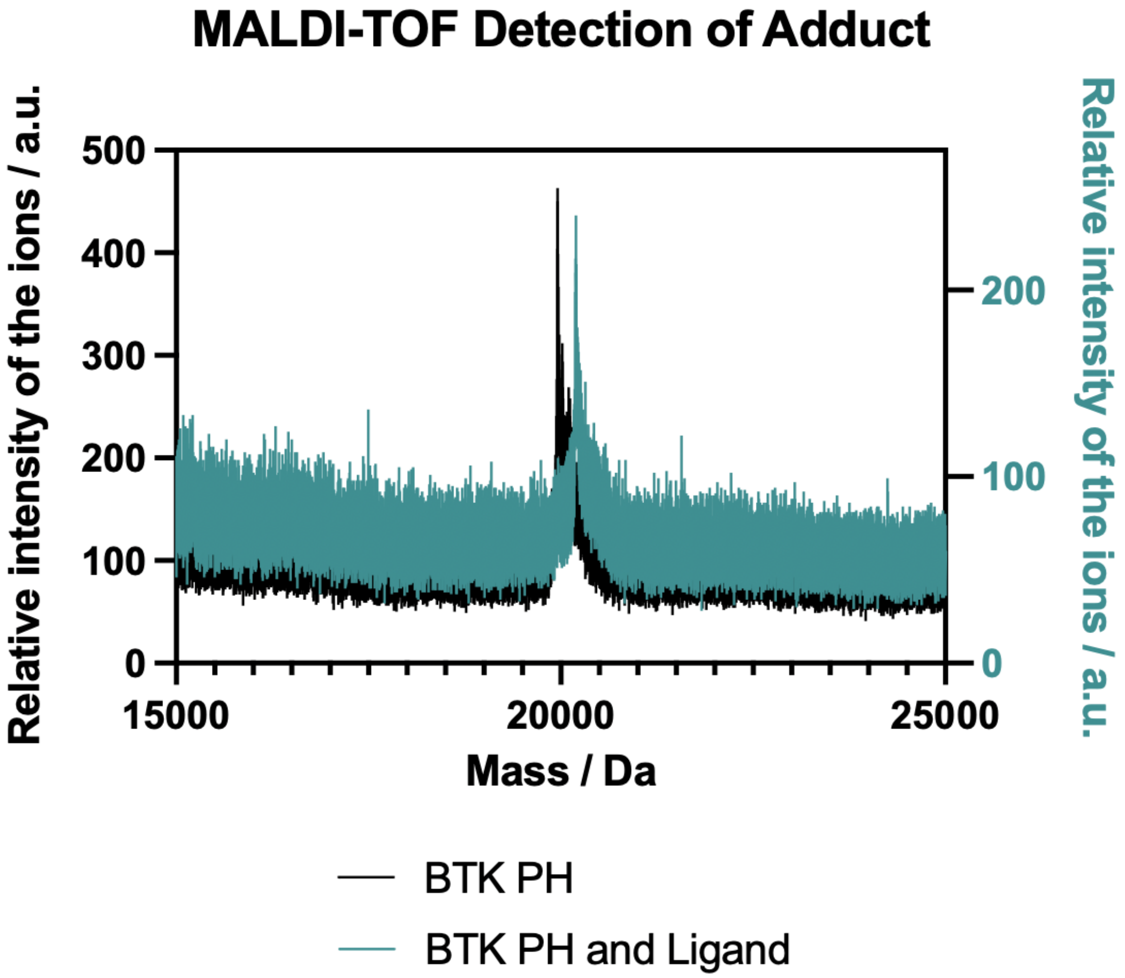

Figure S100 – Matrix Assisted Laser Desorption/Ionization – Time of Flight results with BTK PH domain alone (black) and BTK PH domain with compound **2** (teal). Covalent adduct with WT = 19957 + 256 – 18 = 20195 Da.

### S6 Differential Scanning Fluorimetry (DSF) Studies

#### S6.1 Raw Data for Figure 9

Table S4 – A table to show WT protein (5 µM) $T_m$ values obtained in DSF with parent phenyl, **2**, the ortho fluoro chloro analogue, **17**, and its reduced counterpart **17e** at varying ligand concentration.

| **Conc/ µM** | **$T_m$ values (°C) for Analogue** | | |
|---|---|---|---|
| | **2** | **17e** | **17** |
| **3000** | 51.5 | | 59.0 |
| **1500** | 51.5 | 37.0 | 58.5 |
| **750** | 51.0 | 37.5 | 57.0 |
| **375** | 50.0 | 38.0 | 55.5 |
| **187.5** | 48.0 | | 53.0 |
| **93.75** | 46.0 | 40.0 | 51.0 |
| **46.88** | 45.0 | 41.5 | 49.5 |
| **23.44** | 44.5 | 42.0 | 47.5 |
| **11.72** | 44.0 | 42.5 | 45.5 |
| **5.86** | 43.5 | 42.5 | 44.0 |
| **2.93** | 43.5 | 42.5 | 44.0 |
| **1.46** | 43.5 | 42.0 | 43.5 |
| **0.73** | 43.0 | 42.5 | 43.0 |
| **0.10** | 44.0 | 42.5 | 43.0 |

S6.2 Supplementary Figures S101A and S101B

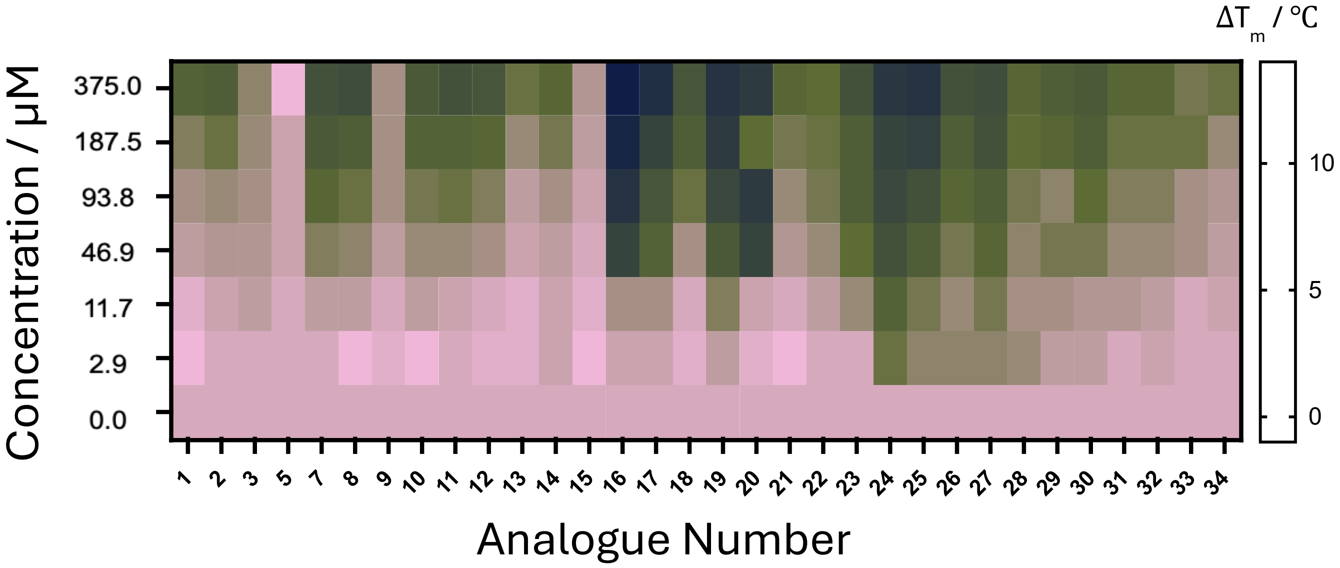

Figure S101A - Differential Scanning Fluorimetry with WT (5 μM).

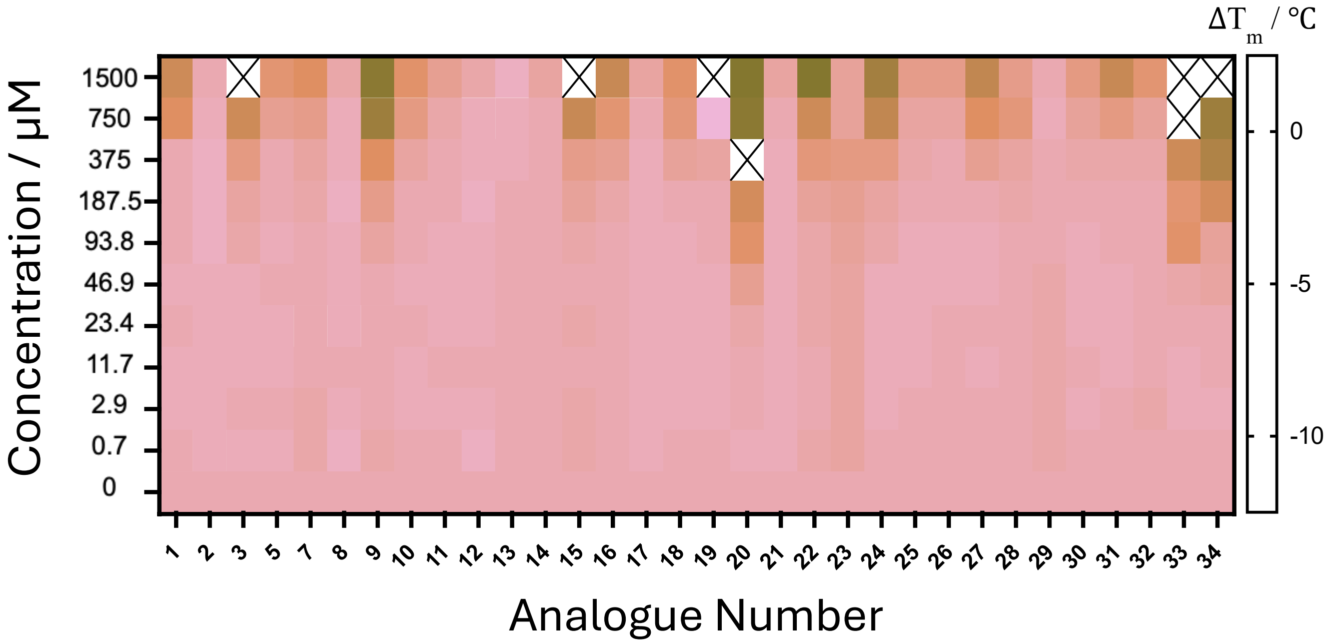

Figure S101B - Differential Scanning Fluorimetry with R28C Mutant (5 μM). Conditions marked as “X” showed now clear melting curve.

#### S6.3 Raw data for Supplementary Figures S101A and S101B

##### Table S5 – Raw Data used to calculate Δ $T_m$ for Supp. Figure S101A- $T_m$ values (°C) from DSF with WT (5 μM)

| Conc/ μM | $T_m$ values (°C) for Analogue | | | | | | | | | | | | | | | | | | | | | | | | | | | | | | |
|---|---|---|---|---|---|---|---|---|---|---|---|---|---|---|---|---|---|---|---|---|---|---|---|---|---|---|---|---|---|---|---|
| | 1 | 2 | 3 | 5 | 7 | 8 | 9 | 10 | 11 | 12 | 13 | 14 | 15 | 16 | 17 | 18 | 19 | 20 | 21 | 22 | 23 | 24 | 25 | 26 | 27 | 28 | 29 | 30 | 31 | 32 | 33 | 34 |
| 3000 | | 51.5 | 45.5 | 39.5 | 52.5 | 55.5 | | 51.0 | 53.0 | 56.5 | 55.0 | 53.5 | | 59.0 | 59.0 | | 55.5 | | 52.0 | 43.0 | 54.0 | | | | | | | | | | | |
| 1500 | | 51.5 | 45.5 | 42.0 | 52.5 | 55.0 | | 51.5 | 52.5 | 55.5 | 52.0 | 52.5 | | 59.5 | 58.5 | 53.0 | 57.0 | | 51.5 | 46.0 | 52.5 | 55.5 | 56.0 | 53.5 | | 49.0 | 55.0 | | 51.5 | 52.5 | | |
| 750 | | 51.0 | 46.0 | 44.0 | 53.0 | 53.5 | 44.0 | 51.5 | 52.0 | 53.0 | 50.0 | 51.0 | 45.5 | 58.0 | 57.0 | 52.5 | 57.0 | 53.0 | 50.0 | 48.0 | 52.0 | 55.5 | 55.0 | 52.5 | | 49.5 | 54.0 | 54.5 | 52.5 | 53.0 | | 50.0 |
| 375 | 49.5 | 50.0 | 46.5 | 42.5 | 51.5 | 52.0 | 45.5 | 50.5 | 51.5 | 51.0 | 48.0 | 49.0 | 45.0 | 57.5 | 55.5 | 51.0 | 55.0 | 54.0 | 49.0 | 48.5 | 51.5 | 54.5 | 55.0 | 51.5 | 52.0 | 49.0 | 53.5 | 54.0 | 52.5 | 52.5 | 51.0 | 51.5 |
| 187.5 | 47.0 | 48.0 | 46.0 | 44.0 | 50.5 | 50.0 | 45.5 | 49.5 | 49.5 | 49.0 | 46.0 | 47.5 | 44.5 | 56.5 | 53.0 | 50.0 | 54.0 | 48.5 | 47.5 | 48.0 | 50.0 | 53.0 | 53.5 | 50.0 | 51.5 | 48.5 | 52.5 | 53.5 | 51.5 | 51.5 | 51.5 | 49.5 |
| 93.75 | 45.5 | 46.0 | 45.5 | 44.0 | 49.0 | 48.0 | 45.5 | 47.5 | 48.0 | 47.0 | 44.5 | 45.5 | 44.0 | 55.0 | 51.0 | 48.0 | 52.5 | 54.0 | 46.0 | 47.5 | 50.0 | 52.5 | 51.5 | 49.0 | 50.0 | 47.5 | 50.0 | 52.0 | 50.5 | 50.5 | 49.0 | 48.5 |
| 46.88 | 44.5 | 45.0 | 45.0 | 44.0 | 47.0 | 46.5 | 44.5 | 46.0 | 46.0 | 45.5 | 44.0 | 44.5 | 43.5 | 53.0 | 49.5 | 45.5 | 50.5 | 53.0 | 45.0 | 46.0 | 48.5 | 51.5 | 50.0 | 47.5 | 49.0 | 46.5 | 51.0 | 51.0 | 49.5 | 49.5 | 49.0 | 48.0 |
| 23.44 | 44.0 | 44.5 | 44.5 | 44.0 | 45.5 | 45.0 | 44.5 | 45.0 | 45.0 | 45.5 | 43.5 | 44.5 | 43.5 | 50.0 | 47.5 | 44.5 | 49.5 | 48.0 | 44.0 | 45.0 | 47.0 | 50.5 | 49.0 | 47.0 | 48.0 | 46.0 | 50.5 | | | | | |
| 11.72 | 43.0 | 44.0 | 44.5 | 43.5 | 44.5 | 44.5 | 43.5 | 44.5 | 44.0 | 43.5 | 43.0 | 44.0 | 43.0 | 45.5 | 45.5 | 43.5 | 47.0 | 44.0 | 43.5 | 44.5 | 46.0 | 49.5 | 47.5 | 46.0 | 47.5 | 45.5 | 49.0 | 48.5 | 48.5 | 48.0 | 47.0 | 47.5 |
| 5.86 | 43.0 | 43.5 | 43.5 | 43.5 | 43.5 | 43.5 | 43.5 | 43.5 | 44.0 | 43.0 | 43.0 | 44.0 | 43.0 | 44.0 | 44.0 | 43.0 | 46.0 | 43.5 | 43.0 | 43.5 | 44.0 | 48.5 | | | | | | | | | | |
| 2.93 | 42.5 | 43.5 | 43.5 | 43.5 | 43.5 | 42.5 | 43.0 | 42.5 | 43.5 | 43.0 | 43.0 | 44.0 | 42.5 | 44.0 | 44.0 | 43.0 | 44.5 | 43.0 | 42.5 | 43.5 | 43.5 | 48.0 | 46.5 | 46.5 | 46.5 | 46.0 | 48.0 | 48.0 | 47.0 | 47.5 | 47.0 | 47.0 |
| 1.46 | 42.5 | 43.5 | 43.0 | 43.5 | 43.0 | 43.0 | 43.0 | 42.5 | 44.0 | 42.5 | 43.0 | 43.5 | 43.0 | 43.5 | 43.5 | 43.0 | 43.5 | 43.0 | 42.5 | 43.0 | 43.5 | 47.5 | | | | | | | | | | |
| 0.73 | 42.5 | 43.0 | 43.5 | 43.0 | 43.0 | 43.0 | 43.0 | 42.5 | 43.0 | 42.5 | 42.5 | 44.0 | 42.5 | 43.0 | 43.0 | 43.0 | 44.5 | 43.0 | 42.5 | 43.0 | 43.0 | 46.5 | 46.5 | 46.0 | 46.0 | 46.0 | 47.5 | | | 47.5 | 47.0 | |
| 0.00 | | 44.0 | | | | | | | 43.5 | | | 43.5 | | 43.5 | | | | 43.0 | | | | 46.0 | | 43.0 | 44.5 | | 47.0 | | | | | |

Table S6 – Raw Data used to calculate Δ $T_m$ for Supp. Figure S101B - $T_m$ values (°C) from DSF with Mutant (5 μM)

| Conc/ μM | $T_m$ values (°C) for Analogue | | | | | | | | | | | | | | | | | | | | | | | | | | | | | | |
|---|---|---|---|---|---|---|---|---|---|---|---|---|---|---|---|---|---|---|---|---|---|---|---|---|---|---|---|---|---|---|---|
| | 1 | 2 | 3 | 5 | 7 | 8 | 9 | 10 | 11 | 12 | 13 | 14 | 15 | 16 | 17 | 18 | 19 | 20 | 21 | 22 | 23 | 24 | 25 | 26 | 27 | 28 | 29 | 30 | 31 | 32 | 33 | 34 |
| 3000 | 41.5 | 49.0 | 41.5 | 42.0 | 40.5 | 47.0 | 36.5 | 41.5 | 45.0 | 47.0 | 49.0 | 45.0 | | 38.0 | 44.0 | | | 32.5 | 45.5 | | 50.5 | 41.5 | 43.0 | 44.5 | | | 49.5 | | 40.5 | | | |
| 1500 | 42.5 | 49.5 | | 45.5 | 44.5 | 48.5 | 37.0 | 44.5 | 47.0 | 48.5 | 50.5 | 48.5 | | 42.5 | 47.5 | 44.5 | | 36.5 | 47.5 | 37.0 | 50.5 | 40.5 | 48.0 | 48.0 | 43.0 | 48.0 | 50.0 | 47.5 | 43.5 | 46.5 | | |
| 750 | 44.0 | 50.0 | 42.5 | 47.5 | 47.0 | 49.5 | 38.5 | 46.0 | 48.5 | 49.5 | 50.0 | 49.5 | 42.5 | 45.5 | 48.5 | 45.5 | 53.0 | 37.0 | 48.5 | 43.0 | 50.5 | 43.0 | 49.5 | 49.5 | 45.5 | 47.0 | 50.5 | 49.0 | 47.5 | 49.0 | | 40.0 |
| 375 | 49.0 | 50.5 | 46.0 | 49.5 | 48.5 | 49.5 | 44.0 | 48.0 | 49.0 | 49.5 | 50.0 | 49.5 | 47.0 | 47.5 | 49.0 | 47.5 | 49.5 | 27.0 | 49.0 | 46.0 | 49.0 | 47.5 | 50.0 | 50.5 | 48.5 | 49.5 | 50.0 | 50.0 | 50.0 | 50.0 | 44.0 | 41.5 |
| 187.5 | 49.0 | 50.5 | 48.0 | 49.5 | 49.0 | 50.0 | 46.5 | 49.0 | 49.0 | 50.0 | 49.5 | 49.5 | 48.0 | 49.0 | 49.0 | 49.0 | 50.5 | 43.0 | 49.0 | 48.0 | 50.0 | 49.5 | 50.5 | 50.5 | 50.5 | 50.0 | 50.0 | 50.5 | 50.5 | 50.5 | 46.5 | 44.5 |
| 93.75 | 49.0 | 50.5 | 48.5 | 50.0 | 49.5 | 49.5 | 48.0 | 49.0 | 49.5 | 49.5 | 49.5 | 49.5 | 49.0 | 49.5 | 49.0 | 49.5 | 50.5 | 44.5 | 49.0 | 49.0 | 50.5 | 50.0 | 51.0 | 51.0 | 51.0 | 50.5 | 50.0 | 51.0 | 50.5 | 50.5 | 46.0 | 49.0 |
| 46.88 | 49.5 | 50.0 | 49.5 | 49.5 | 49.5 | 49.5 | 49.0 | 49.5 | 49.5 | 49.5 | 49.5 | 49.5 | 49.5 | 49.5 | 49.0 | 49.5 | 51.0 | 47.0 | 49.0 | 49.0 | 51.0 | 51.0 | 51.0 | 51.0 | 51.0 | 50.5 | 49.5 | 51.0 | 51.0 | 50.5 | 50.0 | 49.5 |
| 23.44 | 49.0 | 50.0 | 49.5 | 50.0 | 49.5 | 49.5 | 49.0 | 49.0 | 49.5 | 49.5 | 49.5 | 49.5 | 49.0 | 49.5 | 49.0 | 49.5 | 51.0 | 48.5 | 49.0 | 49.0 | 51.0 | 51.0 | 51.0 | 50.5 | 50.5 | 50.5 | 49.5 | 51.0 | 51.0 | 50.5 | 50.5 | 50.5 |
| 11.72 | 49.5 | 50.0 | 49.5 | 50.0 | 49.5 | 49.0 | 49.0 | 49.5 | 49.0 | 49.0 | 49.5 | 49.5 | 49.5 | 49.5 | 49.0 | 49.5 | 51.0 | 49.0 | 49.0 | 49.5 | 51.0 | 51.0 | 51.0 | 50.5 | 51.0 | 50.5 | 49.5 | 50.5 | 51.0 | 50.5 | 51.0 | 50.5 |
| 5.86 | 49.5 | 50.0 | 49.5 | 50.0 | 49.0 | 49.5 | 49.0 | 49.5 | 49.5 | 49.5 | 49.5 | 49.5 | 49.5 | 49.5 | 49.0 | 49.0 | | 49.0 | 49.0 | 49.0 | | | | | | | 49.5 | 50.5 | 50.5 | 50.5 | 50.5 | 50.5 |
| 2.93 | 49.5 | 50.0 | 49.0 | 49.5 | 49.0 | 49.5 | 49.0 | 49.5 | 49.5 | 49.5 | 49.5 | 49.5 | 49.0 | 49.5 | 49.0 | 49.5 | 51.0 | 49.0 | 49.0 | 49.5 | 51.0 | 51.0 | 50.5 | 50.5 | 50.5 | 50.5 | 49.5 | 51.0 | 50.5 | 50.0 | 51.0 | 51.0 |
| 1.46 | 49.0 | 50.0 | 49.0 | 49.5 | 49.0 | 49.5 | 49.0 | 49.5 | 49.0 | 49.5 | 49.5 | 49.5 | 49.0 | 49.5 | 49.0 | 49.5 | | 49.0 | 49.0 | 49.0 | | | | | | | 49.5 | | | 50.5 | | |
| 0.73 | 49.0 | 50.0 | 49.5 | 50.0 | 49.0 | 50.0 | 48.5 | 49.0 | 49.0 | 50.0 | 49.5 | 49.5 | 49.0 | 49.5 | 49.0 | 49.0 | 50.5 | 49.5 | 49.0 | 49.0 | 51.0 | 50.5 | 50.5 | 50.5 | 50.5 | 50.5 | 49.5 | 50.5 | 50.5 | 50.5 | 50.5 | 50.5 |
| 0.00 | 49.0 | 49.5 | 49.0 | 49.5 | 49.5 | 49.0 | 49.0 | 49.0 | | 49.0 | 49.5 | 49.5 | 49.5 | 49.5 | 48.5 | 49.0 | 50.5 | 49.0 | 48.5 | 49.5 | 52.0 | 50.5 | 50.5 | 50.5 | 50.5 | 50.5 | 50.0 | 50.5 | 50.5 | 50.5 | 50.5 | 50.5 |

Table S7 – Fitting DSF data to $Y = B_{max}*X/(K_d + X)$ using Prism 10.

| **Analogue** | **$B_{max}$** | **$K_d$** | **$R^2$** | **$\log(K_d)$ from DSF** |
|---|---|---|---|---|
| 1 | 18.4 | 795.8 | 0.98 | 2.90 |
| 2 | 9.0 | 190.7 | 0.99 | 2.28 |
| 3 | 2.5 | 29.2 | 0.90 | 1.47 |
| 4 | No data | | | |
| 5 | Undetermined | | | |
| 6 | No data | | | |
| 7 | 9.8 | 81.3 | 0.99 | 1.91 |
| 8 | 12.6 | 175.3 | 0.99 | 2.24 |
| 9 | 1.7 | 30.8 | 0.66 | 1.49 |
| 10 | 8.5 | 99.6 | 0.97 | 2.00 |
| 11 | 9.9 | 121.6 | 0.99 | 2.08 |
| 12 | 14.2 | 312.9 | 0.98 | 2.50 |
| 13 | 15.1 | 1015.0 | 0.98 | 3.01 |
| 14 | 11.3 | 375.9 | 0.99 | 2.58 |
| 15 | 4.1 | 731.9 | 0.70 | 2.86 |
| 16 | 15.9 | 41.5 | 0.99 | 1.62 |
| 17 | 15.5 | 92.9 | 0.99 | 1.97 |
| 18 | 10.8 | 156.8 | 0.98 | 2.20 |
| 19 | 13.0 | 33.9 | 0.99 | 1.53 |
| 20 | 10.4 | 30.8 | 0.82 | 1.49 |
| 21 | 9.3 | 270.1 | 0.97 | 2.43 |
| 22 | 5.4 | 51.9 | 0.96 | 1.71 |
| 23 | 9.4 | 47.3 | 0.98 | 1.67 |
| 24 | 10.5 | 5.8 | 0.88 | 0.76 |
| 25 | 11.5 | 25.8 | 0.90 | 1.41 |
| 26 | 8.6 | 38.3 | 0.81 | 1.58 |
| 27 | 7.8 | 10.6 | 0.85 | 1.02 |
| 28 | 5.6 | 24.0 | 0.70 | 1.38 |
| 29 | 6.7 | 34.5 | 0.91 | 1.54 |
| 30 | 7.9 | 47.7 | 0.99 | 1.68 |
| 31 | 5.4 | 44.6 | 0.96 | 1.65 |
| 32 | 6.2 | 65.3 | 0.99 | 1.82 |
| 33 | 5.7 | 112.6 | 0.93 | 2.05 |
| 34 | 4.4 | 130.8 | 0.87 | 2.12 |

### S7 Statistics

#### S7.1 Normality Tests

Table S8 - Results of the Shapiro-Wilk test performed on the log($K_d$) values approximated from DSF.

| **Approximate log($K_d$) from DSF, Shapiro-Wilk test** | |
|---|---|
| W | 0.9702 |
| P value | 0.5245 |
| Passed normality test (alpha=0.05)? | Yes |
| P value summary | ns |

Table S9 - Results of the Shapiro-Wilk test performed on the total labelling relative intensity values from mass spectrometry.

| **Total Labelling Relative Intensity from MS, Shapiro-Wilk test** | |
|---|---|
| W | 0.9406 |
| P value | 0.0855 |
| Passed normality test (alpha=0.05)? | Yes |
| P value summary | ns |

#### S7.2 Correlation Tests

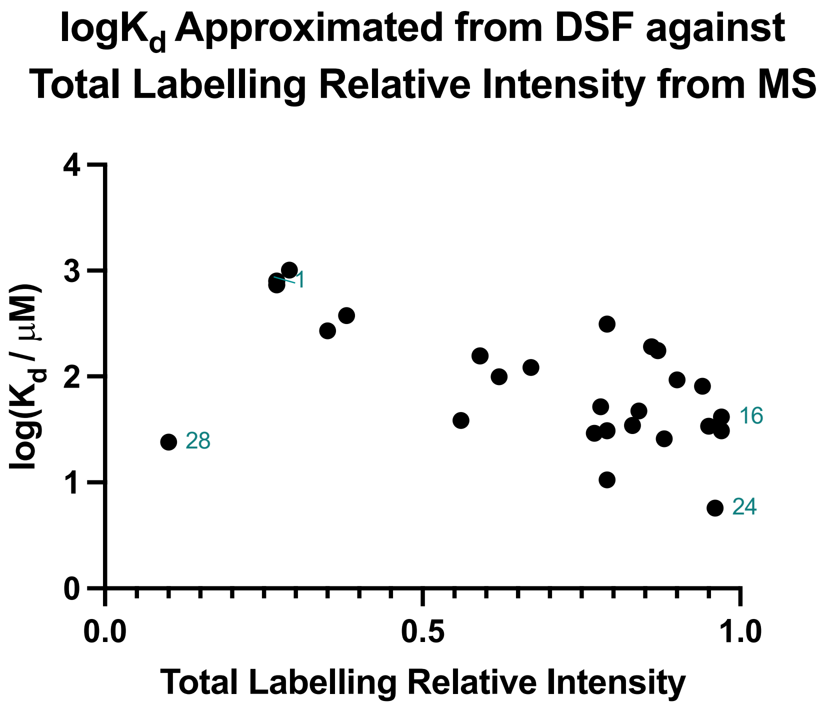

Figure S102 – A graph to show log($K_d$) values approximated from DSF against Total Labelling Relative Intensity values from mass spectrometry. Pearson r = -0.56, 95 % confidence interval = -0.78 to -0.22, R squared = 0.32, P value (two-tailed) = 0.003. Pearson correlation analysis, appropriate because the data points are independent and the expected relationship is linearly covariant, with x and y values recorded independently and sampled from populations that follow Gaussian distributions, gives a P value < 0.05 suggesting a correlation. The R squared value is low (R > 0.5 implies the model fits the data well). Analogue **28** appears to be anomalous.

### S8 Separation of Enantiomers

#### S8.1 Supercritical fluid chromatography

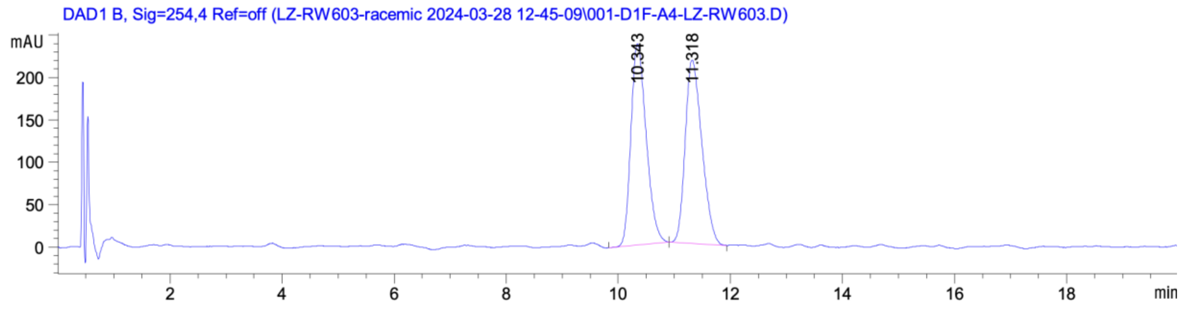

Figure S103 – Chiral LCMS spectrum for a racemic mixture of **19**. Peak 1 and 2 in buffer (20 mM HEPES pH 8, TCEP 0.5 mM, NaCl 100 mM).

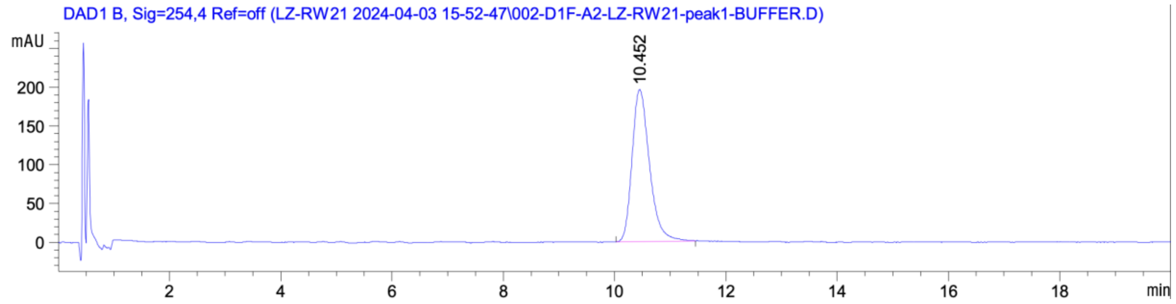

Figure S104 – Chiral LCMS spectrum for t = 0 hr, peak 1 in buffer (20 mM HEPES pH 8, TCEP 0.5 mM, NaCl 100 mM).

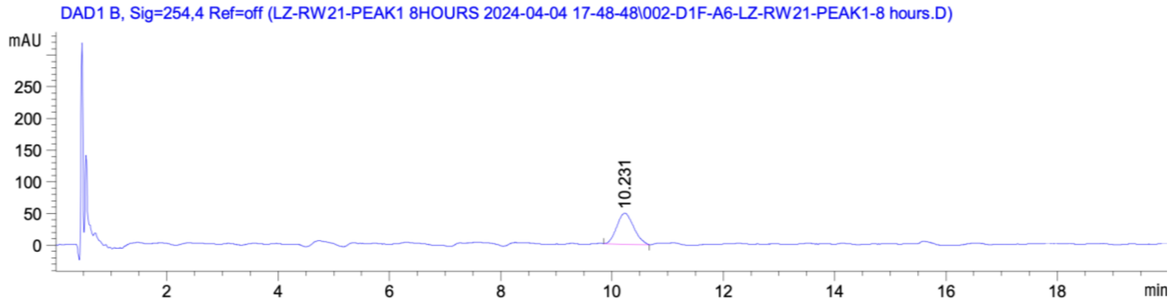

Figure S105 – Chiral LCMS spectrum for t = 8 hr, peak 1 in buffer (20 mM HEPES pH 8, TCEP 0.5 mM, NaCl 100 mM).

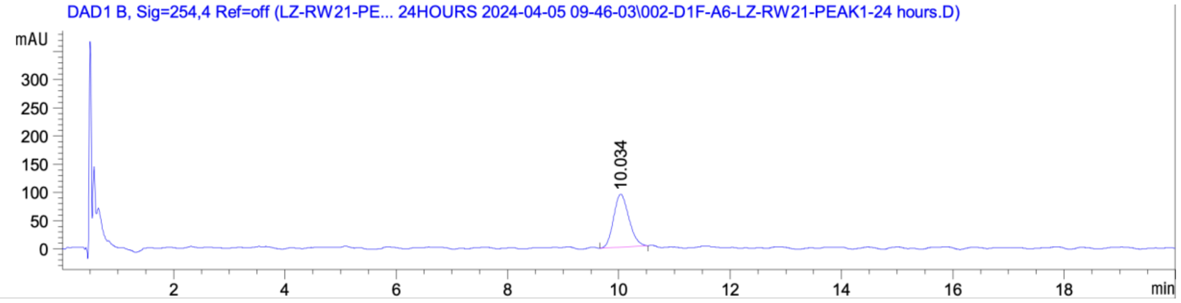

Figure S106 – Chiral LCMS spectrum for t = 24 hr, peak 1 in buffer (20 mM HEPES pH 8, TCEP 0.5 mM, NaCl 100 mM).

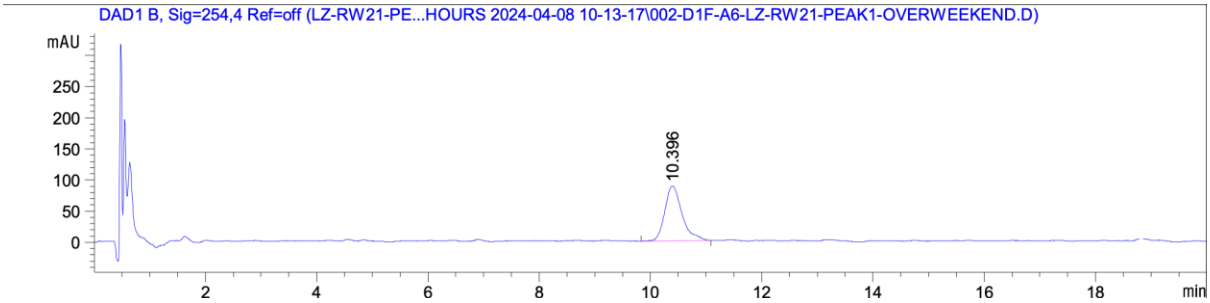

Figure S107 – Chiral LCMS spectrum for t = 96 hr, peak 1 in buffer (20 mM HEPES pH 8, TCEP 0.5 mM, NaCl 100 mM).

### S9 Computational Work

#### S9.1 Mixed Solvent Molecular Dynamics

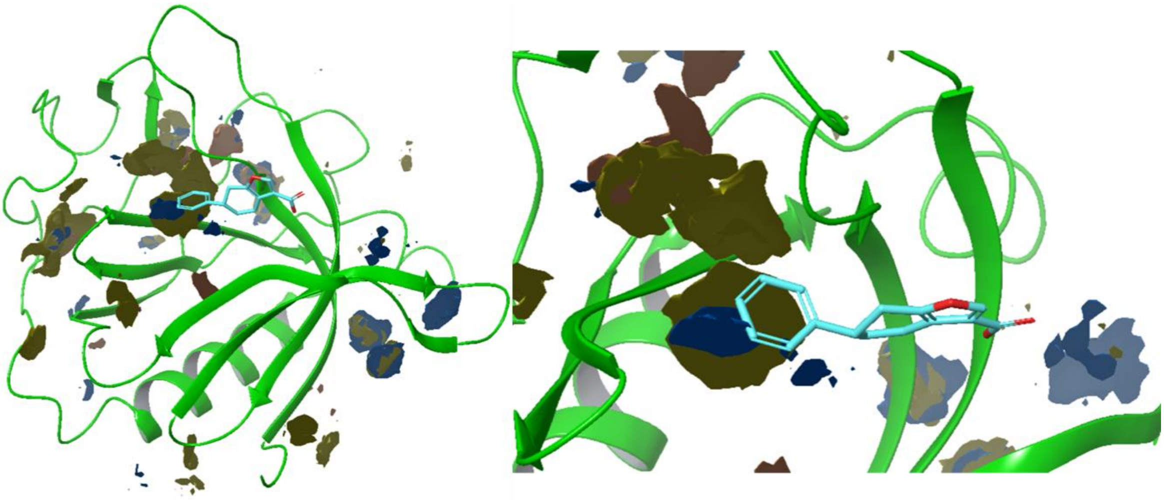

Figure S108 – Hot spot probing of the BTK PH domain with mixed solvent MD simulations. The colour of the pockets represents the type of co-solvent clustering: maroon – acetonitrile; yellow-brown – pyrimidine; blue – isopropanol.

#### S9.2 Virtual Screening

Virtual screening has been performed (using Schrödinger) in order to probe the pocket with various electronics, H-bonding potentials and degrees of hydrophobicity. R-group enumeration was used to enumerate a library of different substituents (halogens, alkyl and aromatic) on the three positions of the phenyl ring. The enumerated library of mono and di-substituted analogues (n = 192) was then used in virtual screening, with a cut-off for pose retainment of 2.5 kcal/mol. Given the rigorous cut-off, virtual screening yielded valid poses for 95 analogues with various mono and di-substitution patterns. Docking scores were then analysed in boxplots (Figure S109), in order to determine which substituents induce low and high docking scores. Covalent docking score analysis shows halogens in the *ortho* and *para* position are privileged scaffolds, inducing the most negative docking scores. Other emerging privileged groups are isopropoxy and methoxy. Phenoxy, phenyl and trifluoromethyl substituents emerge as destabilising across all three positions. The emerging trends were confirmed by MS and DSF, which highlighted the privileged nature of the halogens in *ortho* and *para* positions, as well as the detrimental effects of extra phenyl rings or trifluoromethyl.

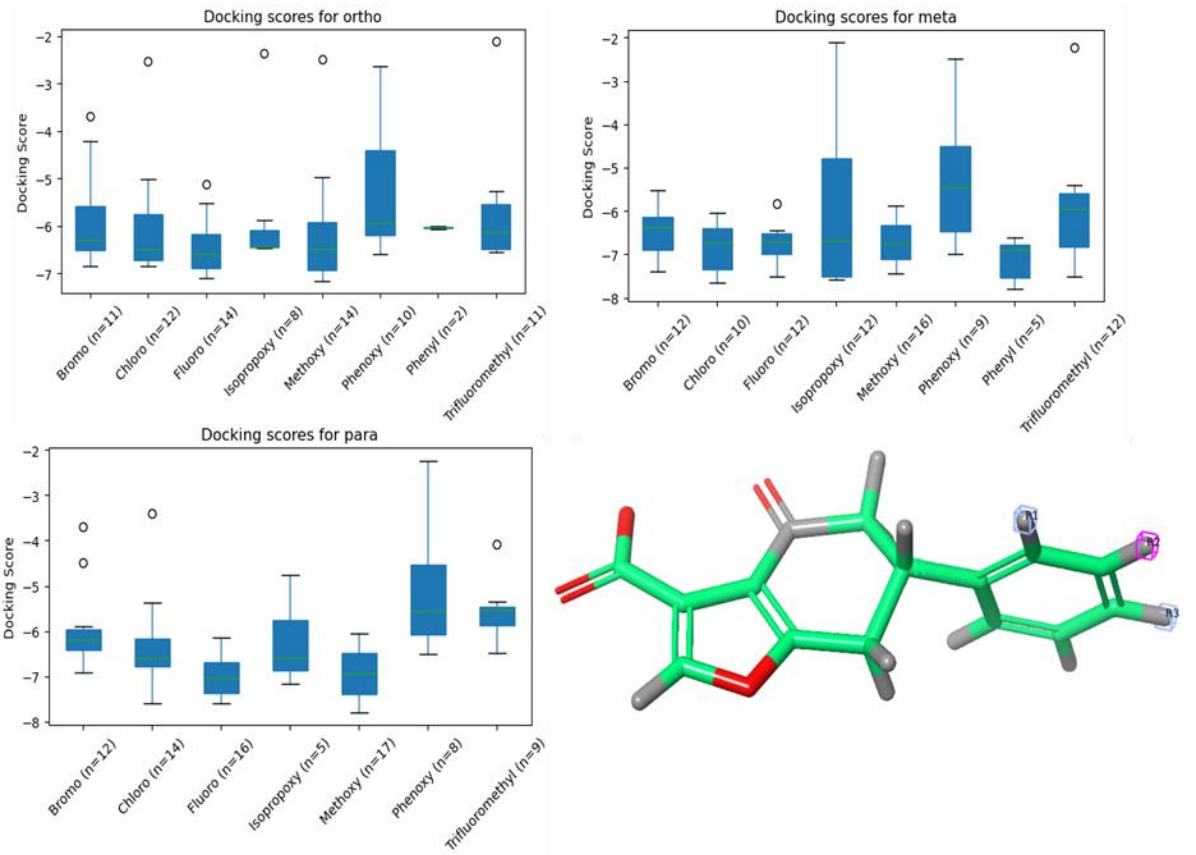

Figure S109 – Virtual screening results of the outcome of R group enumeration across the three positions of the phenyl ring.

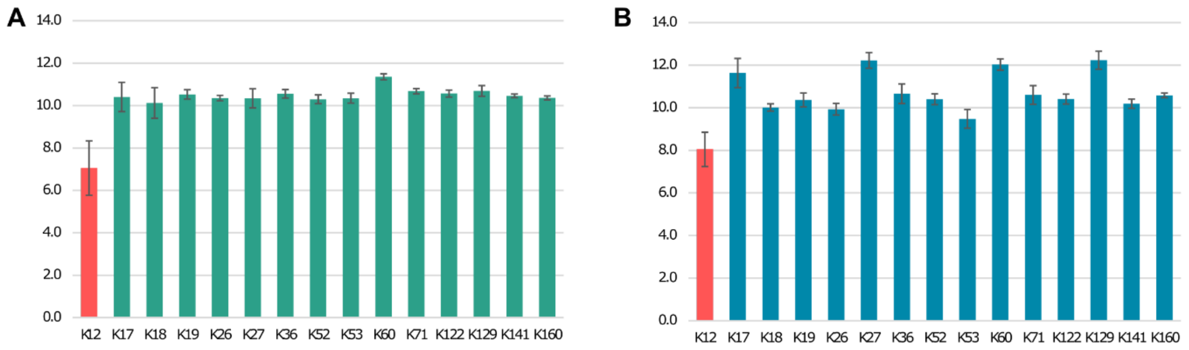

Figure S110. Calculated pKa values for all the lysines in Btk PH domain, calculated with DeepKa (A) and pypKa (B) servers using structures 1B55, 1BWN, 2Z0B and 4Y94 from PDB.

**Table S10**. Calculated pKa values for Btk K12 and equivalent lysine residues in other PH domains. The PDB coordinates used were 1B55, 1BWN, 2Z0B and 4Y94 for Btk, 1H10, 1UNQ and 2UZR for Akt, 1FGZ, 1FHW, 1FHX and 1FGY for GRP1, 1FAO and 1FB8 for DAPP1, 5D27, 5D3X, 5D3V and 5D3Y for p-Rex1 and 1mai for PLCδ. Number of domains in the table refers to total number of PH domains was used in calculations from all the structures. The pKa difference is calculated using the reference value for lysine amino group used in the two programmes, 10.40 and 10.46 for DeepKa and pypKa, respectively.

| Method | Protein | Number of domains | Lys12 equivalent | Average pKa for Lys12 | Average pKa difference | Number of other lysines per domain | Average pKa for other lysines |
|---|---|---|---|---|---|---|---|
| pypKa | Btk | 12 | K12 | 8.83 ± 0.80 | -2.26 | 14 | 10.84 ± 0.79 |
| DeepKa | Btk | 12 | K12 | 7.05 ± 1.29 | -3.35 | 14 | 10.51 ± 0.97 |
| DeepKa | Akt | 3 | K14 | 8.24 ± 0.87 | -2.16 | 7 | 10.71 ± 0.33 |
| DeepKa | GRP1 | 6 | K273 | 7.32 ± 1.02 | -3.19 | 10 | 10.46 ± 0.35 |
| DeepKa | Dapp1 | 2 | K173 | 7.82 ± 0.18 | -2.59 | 8 | 10.54 ± 0.26 |
| DeepKa | P-Rex1 | 7 | K280 | 8.08 ± 0.62 | -2.32 | 14 | 10.42 ±0.32 |
| DeepKa | PLCδ | 1 | K30 | 9.42 | -0.98 | 8 | 10.78 ±0.45 |
